# In silico reconstruction of primary and metastatic tumor architecture using GIS-augmented spatial transcriptomics

**DOI:** 10.1101/2025.10.16.682879

**Authors:** Jin Young Yoo, Sabrina Akter, Qianying Zuo, Amir Kazemi, Wenjun Wu, Mahima Goel, Audrey Lam, Debapriya Dutta, Betsy Barnick, Melike Pekmezci, Maria Grosse Perdekamp, Aiman Soliman, Zeynep Madak-Erdogan

**Author notes:** Corresponding Author: Zeynep Madak Erdogan, 1201 W Gregory Dr., Urbana, IL 61801, USA.

## Abstract

The tumor microenvironment (TME) comprises different cell populations that interact, contributing to tumor heterogeneity and therapy response. Spatial transcriptomics offers valuable insights into transcriptional complexity and heterogeneity of the TME. We established Geographic Information System (GIS)-augmented In-Silico Reconstruction of Tumor Architecture (GIS-ROTA), a biologically informed analytic framework that integrates pathway or cell type-based enrichment analysis with local Moran’s I to uncover functional spatial domains. In our Visium dataset of primary and metastatic estrogen receptor-positive breast tumor samples, GIS-ROTA revealed extensive co-localization of estrogen response with metabolic pathway gene sets and mutual exclusivity with metastasis-related and specific immune-related pathway gene sets. The novelty of our approach lies in considering biological functions prior to identifying any spatial domains, providing direct interpretability and minimizing the subjectivity of interpreting clusters observed from conventional analytic methods. Overall, our GIS-ROTA framework integrates biological knowledge first, yielding spatial patterns with functional relevance and enabling identification of novel targets for development of therapeutic strategies.

## Introduction

Breast tumors are heterogeneous tissues, with diverse cells displaying distinct molecular signatures and differing therapeutic sensitivity(1). Such heterogeneity poses a significant challenge in managing cancer treatment and prognosis. Breast cancer is classified into four subtypes, of which luminal A [estrogen receptor alpha positive (ER+)/progesterone receptor positive (PR+)/human epidermal growth factor negative (HER2-)] is the most common. Therapies targeting estrogen receptor signaling have been the standard-of-care primary approach for metastatic hormone receptor-positive breast cancer patients(2), yet prolonged application results in development of resistance. Metastasis is the primary cause of death in >90% of cancer patients(3,4), highlighting the importance of novel strategies to delay or overcome therapy resistance in cancer cells.

The tumor microenvironment (TME) is a complex ecosystem of cellular and stromal constituents, whose interactions and paracrine signaling contribute to therapy resistance and cancer aggressiveness(5,6). With tumor heterogeneity contributing to therapy resistance and tumor progression, high-throughput methods such as bulk or single-cell RNA sequencing (RNAseq) have been widely used for molecular profiling of cells in the TME(7–9). Such high-throughput data may reveal key molecular signatures or potential biomarkers but do not reflect the TME’s spatial context(10). Spatial transcriptomics (ST) generally refers to methodologies that simultaneously obtain tissue or cell transcriptomic information and spatial information(11). ST methods include sequencing-based, probe-based, imaging-based, and image-guided spatially resolved single-cell RNAseq(12). Although ST has existed for some time(13), the 10X Genomics Visium platform has emerged as a powerful high-throughput tool to visualize the spatial distribution of gene expression across tissue samples and is widely used to study various cancers(14–17). Such rise of spatial transcriptomics enabling incorporation of spatial information in gene expression analysis emphasizes the need to fully utilize the spatial context to better understand cellular networks within the tissue microenvironment.

Geographic Information System (GIS) is a robust geospatial technology that organizes, analyzes, and stores spatial datasets and has been widely applied in agriculture, environmental monitoring, and urban planning(18). A primary advantage of spatial omics data is the ability to identify spatially variable genes (SVGs), genes that exhibit heterogeneous expression patterns across tissues. Several computational approaches have been developed to identify SVGs by applying Local spatial association indices utilized in GIS. For example, saSpatial utilizes Local Moran’s I to detect regions with spatially different enrichment for specific genes(19), while SpatialDM employs bivariate Global Moran’s I statistics to identify co-expression patterns between gene pairs or ligand-receptor pairs(20). Alternative approaches, such as SpatialDE(21) and SPARK(22), model SVG expression using covariance functions (e.g., Gaussian or exponential kernels) to capture spatial effects. SPARK further expands this framework by employing a generalized linear spatial model that directly models observed UMI counts. To reduce computational complexity from cubic to nearly linear scaling, methods such as SPARK-X test whether residuals from generalized linear models exhibit spatial independence using autocorrelation measures(23). More recent approaches for SVG identification include DESpace(24), a two-step framework connecting clustering and negative binomial model, and HEARTSVG(25), an autocorrelation-based nonparametric spatial analytic approach.

Unsupervised clustering based on the most significant SVGs often outperforms non-selective approaches that apply clustering after dimensionality reduction (e.g., principal component analysis) of whole transcriptome profiles. However, interpreting the biological relevance of the identified spatial domains remains challenging and often requires post hoc analysis such as differential biomarker detection or gene set enrichment analysis. To overcome such limitation, we developed GIS-augmented In-Silico Reconstruction of Tumor Architecture (GIS-ROTA), a biologically informed and hypothesis-driven analytic framework that considers specific biological functions prior to identifying spatial domains. GIS-ROTA begins with mapping tissues with biologically curated pathway and cell type signatures, followed by spatial localization analysis using local Moran’s I metrics, yielding biologically relevant spatial domains without any need for further annotations. We applied GIS-ROTA to Visium dataset of primary and metastatic ER+ breast tumors and identified biological networks at both molecular signaling and near-cellular levels. Our approach enables targeted biomarker discovery, paving the way for development of personalized treatments while avoiding the subjectivity of interpreting clusters as shown in conventional approaches.

## Results

### Overview of GIS-ROTA framework

We established GIS-ROTA, a hypothesis-driven computational framework that integrates spatial analytic approaches with geospatial metrics to identify spatial patterns with pre-defined biological relevance (**Figure 1A**). The analytic workflow begins with pre-processed spatial -omics data subjected to gene set coregulation analysis (GESECA) which enables spatial mapping of pathway or cell type gene sets. GESECA assigns a z-score to each barcode within a sample, reflecting the extent to which genes associated with the given pathway/cell type are collectively expressed at that location relative to other spots. Higher z-scores indicate greater pathway or cell type activities in that region. The outputs from GESECA are subsequently subjected to local spatial autocorrelation analysis, following reconstruction into an AnnData object for compatibility. Local spatial autocorrelation is assessed using Local Moran’s I, which classifies each spot into one of four spatial patterns based on its z-score: High-High, Low-Low, High-Low, and Low-High (**Figure 1B**). These patterns, respectively, outline regions of co-localization (concordant high expressions), co-absence (concordant low expressions), and mutual exclusivity (discordant expressions). Overall, the spatial domains identified in our framework represent targeted spatial maps that are both hypothesis-driven and biologically interpretable. In parallel, conventional methods identify broader, exploratory domains that may reflect potential cellular networks but require further analyses for biological validation.

**Figure 1.**
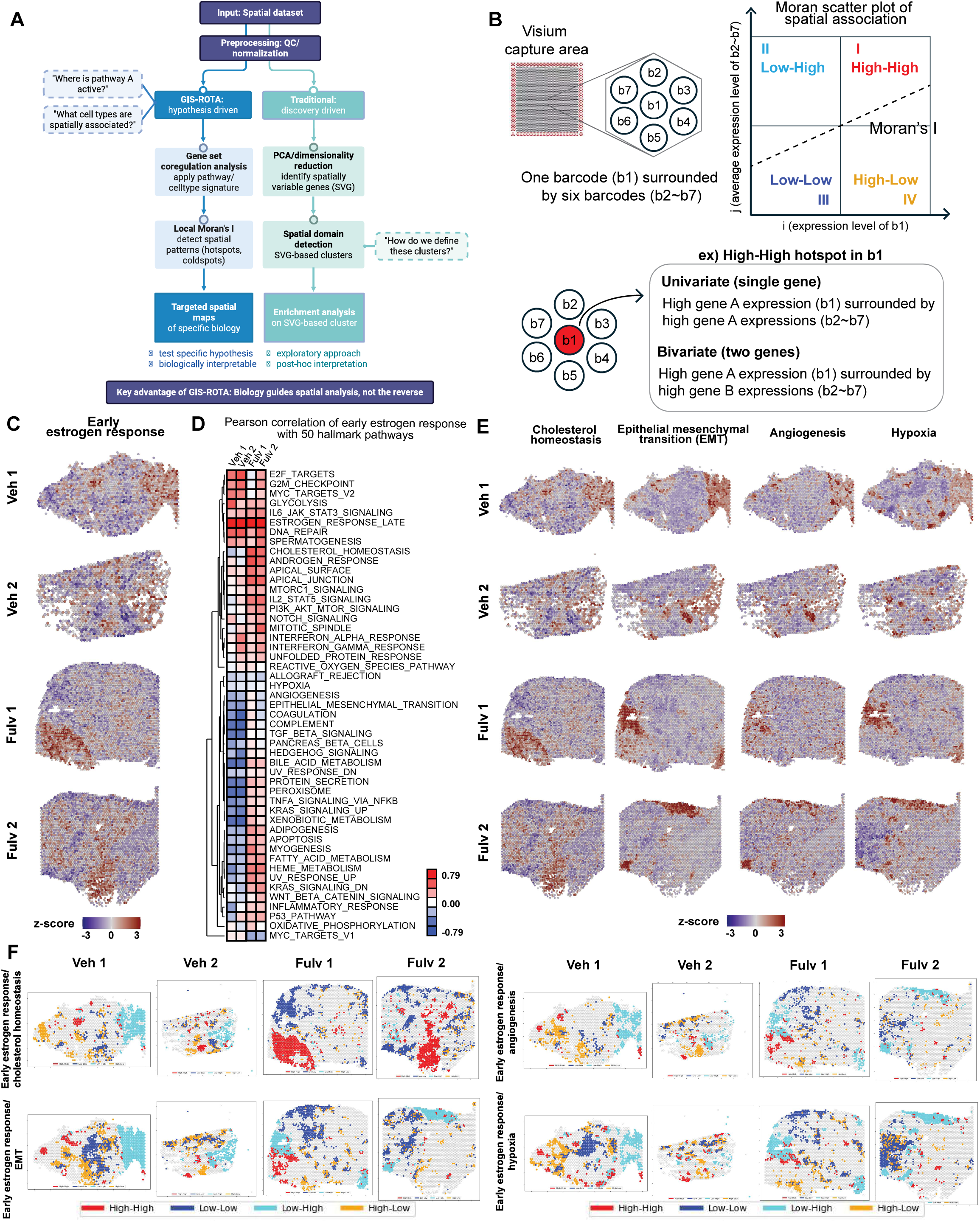
Geographic Information System-augmented In-Silico Reconstruction of Tumor Architecture (GIS-ROTA) framework and application to metastatic xenograft tumor analysis. (A) Visual representation of GIS-ROTA workflow and conventional method. (B) Schematic representation of Local Moran’s I analysis. Each barcode in the Visium capture area is surrounded by six barcodes. Four spatial clusters are defined based on deviation of expression level from global average expression for the central and its six neighboring barcodes. (C) Spatial mapping of GESECA results for early estrogen response. (D) Heatmap plot of the correlation matrix between GESECA z-scores of early estrogen response and hallmark pathways. (E) Spatial mapping of GESECA results for cholesterol homeostasis, EMT, angiogenesis, and hypoxia. (F) Visualization of local spatial cross-correlation based on bivariate local Moran’s I at p < 0.05.

### GIS-ROTA highlights biologically distinct spatial domains missed by unsupervised clustering

We applied GIS-ROTA and conventional unsupervised clustering methods (SNN, BIRCH, FlowSOM, and *k*-means) to 10X Visium spatial gene expression libraries constructed from four fresh-frozen metastatic tumor samples from xenograft mouse models injected with MCF7-*ESR1*^Y537S^ cells(26) and compared the results of our framework with those of traditional approaches. Two samples were from vehicle-treated mice, and two samples were from mice treated with fulvestrant, a selective estrogen degrader clinically used as standard-of-care for patients with ER+ metastatic disease. In this model, fulvestrant failed to reduce metastatic burden, enabling examination of endocrine resistance mechanisms in the metastatic setting(27). Among unsupervised clustering approaches run with default settings, shared nearest neighbor (SNN) clustering most efficiently captured the increased transcriptional diversity in fulvestrant-treated compared to vehicle-treated samples (**Supplementary Figure 1A**). The Uniform manifold approximation and projection (UMAP) plot and SpatialDimPlot of SNN clustering showed increased clusters as well as nonuniform spatial distribution of clusters compared to the vehicle-treated group (**Supplementary Figure 1B**). Gene set enrichment analysis (GSEA) on these clusters further revealed increased enrichment of estrogen response and cholesterol homeostasis in the fulvestrant-treated group except for cluster 8, capturing not only endocrine resistance but also altered metabolic phenotype as compared to other clusters within the same group (**Supplementary Table 1**). SNN clustering, however, is an unsupervised expression-based method, and clusters are often discontinuous as the method does not fully utilize spatial information(28). As such, we observed upregulated early estrogen response across several clusters, indicating a limited capacity of unsupervised clustering to delineate spatial domains with distinct biological functions.

Endocrine-resistant breast cancers characteristically fail to downregulate estrogen response genes following treatment. To investigate whether this dysregulation exhibits spatial organization within the tumor microenvironment, we applied gene set coregulation analysis (GESECA) to map the early estrogen response gene set onto our Visium dataset of the xenograft model (**Figure 1C**). GESECA is a computational function within the *fgsea* R package that estimates z-scores for each barcode, reflecting the variance of gene sets across a sample(29). The z-score per barcode indicates the degree to which gene set members are collectively highly expressed at a given spatial location (barcode) compared to all other locations within the same sample. Barcodes with high z-scores indicate spatial locations where pathway member genes are collectively highly expressed, suggesting increased pathway activity at those positions. To better capture the coordinated biological behavior of cellular networks, we incorporated GESECA into one of the initial steps of the GIS-ROTA to assess spatial autocorrelations at the gene set level. We used pathway gene sets from the Molecular Signatures Database (MSigDB) to compute and map z-scores, visualizing the relative activity of each pathway across the sample. Using the z-scores per barcode calculated from GESECA for all hallmark pathways, we performed Pearson’s correlation analysis to comprehensively assess the spatial relationships between early estrogen response and other pathway gene sets. We then visualized these correlation coefficients as heatmaps, depicting the spatial association of early estrogen response with 50 hallmark gene sets (**Figure 1D**) and 179 KEGG pathways (**Supplementary Figure 1C**). Distinct patterns of positive and negative spatial correlations were observed between fulvestrant and vehicle-treated groups for several metabolic and metastasis-related pathways, including angiogenesis, epithelial–mesenchymal transition (EMT), and hypoxia. However, some of these spatial correlations may result from spatial randomness, where dispersed barcodes show concordant or discordant expressions by chance. To determine whether the correlations observed in the heatmap reflected significant spatially co-regulations of these gene sets rather than mere randomness, we applied Local Moran’s I to the GESECA z-scores of these pathway gene sets. While visual comparison of GESECA z-score maps of these gene sets may provide an approximate degree of spatial co-regulations (**Figure 1E**), Local Moran’s I provides a quantitative metric that formally defines such spatial patterns.

### GIS-ROTA identifies spatial co-regulated patterns between biological pathways

We applied univariate Local Moran’s I to check local spatial autocorrelation for an individual gene set and bivariate Local Moran’s I for a pair of gene sets. Local Moran’s I assesses how strongly expression level at a given location (barcode) correlates with the average expression of its neighboring barcodes. This local autocorrelation is compared to a null distribution generated by randomly shuffling expression values across barcodes to simulate spatial randomness. For a given gene set of interest, each barcode is assigned a label based on its local autocorrelation index. “High” indicates expression above the global average, whereas “Low” indicates expression below the global average. Labels are assigned to each central barcode and its six neighboring barcodes based on combinations of “High” and “Low” expression levels, resulting in four possible categories. Positive autocorrelation indicates concordant expression patterns between the central and its neighboring barcodes. Specifically, High–High regions have higher expression in the central barcode surrounded by similarly higher expression in neighboring barcodes. Low–Low regions have lower expression in the central barcode surrounded by neighboring barcodes with similarly lower expression. On contrary, negative autocorrelation indicates discordant expression patterns between the central and its neighboring barcodes. High–Low regions have higher expression in the central barcode surrounded by neighboring barcodes with lower expression, while Low–High regions have lower expression in the central barcode surrounded by neighboring barcodes with higher expression.

Visualization of bivariate Moran’s I application on the z-scores of gene sets revealed robust co-localization between estrogen response and cholesterol homeostasis gene sets in fulvestrant-treated samples, as evidenced by extensive High-High spots (**Figure 1F**). This finding further supports the positive spatial correlation observed between these gene sets in the heatmap. In contrast, metastasis-related pathway gene sets that exhibited negative spatial correlations in the heatmap, including EMT, angiogenesis, and hypoxia, demonstrated mutual exclusivity with estrogen response gene set, as noted by prominent Low–High spots observed across all four samples. As such, GIS-ROTA assessed spatial co-regulation among pre-defined, biologically meaningful gene sets, eliminating the need for additional interpretation. These co-regulatory patterns were not distinctly captured by conventional unsupervised clustering approaches. We also applied GIS-ROTA to selected members of the early estrogen response gene set to validate our method at the single-gene level. **Supplementary Figure 1D** and 1E show mapping of univariate and bivariate Local Moran’s I results for *TFF1*, *GREB1*, and *FASN*—classical ER target genes(30,31) that displayed distinct spatial variation across samples—in vehicle and fulvestrant-treated samples. We found extensive High-Low spots for TFF1 and GREB1 pair in fulvestrant-treated sample, indicative of partial endocrine responsiveness, along with High-High spots for TFF1 and FASN pair in the same region. This region closely overlapped with High-High spots for early estrogen response and cholesterol homeostasis gene set pair, and this concordance highlights the robustness of the gene set-based approach within GIS-ROTA. This concordance between single-gene and gene set patterns highlights the robustness of the gene set-based approach within GIS-ROTA. Importantly, by aggregating signals from functionally related genes, gene sets reduce noise from individual gene variability and more effectively capture coordinated, pathway-level activity, which is a critical consideration given that cellular phenotypes typically emerge from networks of coregulated genes rather than individual molecular events.

### Generalizability of GIS-ROTA to ER+ breast cancer patient tumors

To evaluate generalizability of GIS-ROTA, we applied it to formalin-fixed paraffin-embedded primary or metastatic tumor samples from ER+ breast cancer patients (**Supplementary Table 2**). Visium gene expression libraries were prepared for five primary breast tumors and seven metastatic tumors. SNN clustering on pre-processed dataset revealed 30 clusters, and the UMAP plot showed minimal overlap of these clusters mapped between samples, hampering examination of biological pathways and comparison of cell populations (**Supplementary Figure 2**). Functional enrichment analyses on each cluster showed consistent mutually exclusive expression of estrogen response and EMT in clusters 0, 1, 5, 10, and 13 from primary breast tumors, whereas most clusters from metastatic tumors had downregulation of both pathways (**Supplementary Table 3**). Consistent with the results from xenograft tumors (**Supplementary Figure 1A**), the unsupervised clustering approach failed to reveal biologically distinct spatial patterns, as evidenced by the enrichment of biological pathways shared across multiple clusters.

We then transitioned to our approach, applying GESECA to hallmark pathway gene sets and estimating Pearson’s correlation coefficients of z-scores per barcode to identify potential spatial correlations across all hallmark gene sets (**Figure 2A** and **Supplementary Figure 3**). We found two distinct groups of pathways from the heatmap, where the pathways from each group displayed strong intra-group spatial correlations: 1) estrogen signaling, cell cycle, and metabolic signaling pathways; and 2) metastasis-related and immune response pathways. Interestingly, such spatial correlation pattern was more evident in primary breast tumors than in metastatic tumors. To assess how spatial correlation of endocrine responsiveness with hallmark pathways differed between primary and metastatic tumors, we focused on early estrogen response gene set and merged individual heatmaps from each sample into a single, consolidated heatmap (**Figure 2B**). Almost all primary breast tumors were closely clustered, with the exception of one ER+/HER2+ sample that clustered with its matching metastatic tumor. Overall, the heatmap showed positive correlation of early estrogen response with metabolic pathways (cholesterol homeostasis, bile acid metabolism, fatty acid metabolism, and oxidative phosphorylation) and cell cycle-related gene sets (MYC targets v1/2, E2F targets, and G2M checkpoint) in primary breast tumors. On the other hand, negative spatial correlations were observed with immune-related pathways (IL6 JAK STAT3 signaling, inflammatory response, and interferon alpha/gamma response), angiogenesis, and EMT gene sets. Metastatic tumors were not as closely clustered as primary tumors did on the heatmap, suggesting increased inter-tumor heterogeneity. ER+/HER2+ primary and metastatic tumors showed overall similar correlation patterns in the heatmap, except for EMT and cell cycle-related gene sets. While these positive and negative correlations in the heatmap suggest potential co-localization and mutual exclusivity between gene sets, further validation using Local Moran’s I is warranted to quantitatively assess and visualize statistically significant spatial co-regulation.

**Figure 2.**
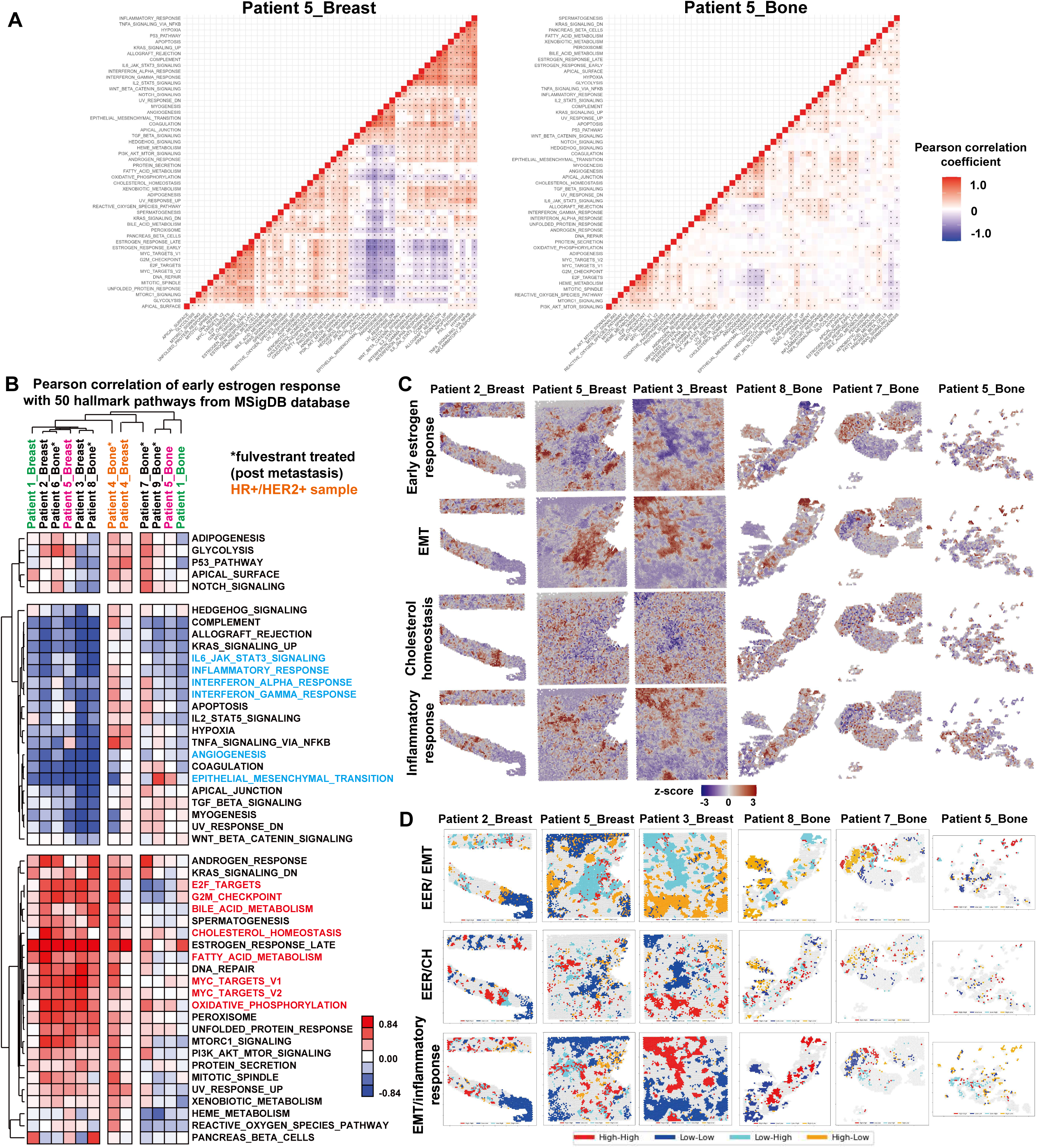
Generalizability of GIS-ROTA to ER+ breast cancer patient tumors. (A) Heatmap plot representing the correlation matrix between gene set coregulation analysis (GESECA) z-scores of hallmark pathways in primary and bone tumors from Patient 5. Grids with asterisks represent statistically significant correlation coefficient values at *p* < 0.05. (B) Heatmap plot representing the correlation matrix between GESECA z-scores of early estrogen response and 49 hallmark pathways for primary breast and metastatic bone tumor samples. Matching pairs are color-coded. (C) Spatial mapping of GESECA results for early estrogen response, cholesterol homeostasis, and EMT gene sets. Barcodes with higher z-scores indicate where pathway member genes are collectively highly expressed. (D) Visualization of local spatial autocorrelation based on bivariate Local Moran’s I of early estrogen response with EMT, early estrogen response with cholesterol homeostasis, and EMT with inflammatory response at *p* < 0.05. MSigDB: Molecular Signature Database; EMT: epithelial–mesenchymal transition; EER: early estrogen response; CH: cholesterol homeostasis

Guided by the prominent correlation patterns observed in the heatmap, we mapped GESECA z-scores for the estrogen response, EMT, and cholesterol homeostasis gene sets for visual comparison (**Figure 2C**) and subsequently applied bivariate Local Moran’s I to these gene sets (**Figure 2D**). Bivariate Local Moran’s I analysis showed extensive High–Low and Low–High regions for early estrogen response and EMT gene set pair in primary tumors, highlighting mutual exclusivity of cell populations expressing these transcriptional programs. Such mutual exclusivity was not as clearly defined in ER+/HER2+ or metastatic tumors (**Supplementary Figure 4**).In contrast, bivariate Local Moran’s I revealed robust co-localization between estrogen response and several metabolic pathway gene sets, including cholesterol homeostasis, fatty acid metabolism, and oxidative phosphorylation. Interestingly, although extensive co-localization of the estrogen response with these metabolic pathways was observed, the co-localized regions were not confined to a single tumor area, suggesting intra-tumor metabolic heterogeneity. As such, GIS-ROTA successfully identified distinct metabolic phenotypes of endocrine-responsive cell populations across different regions of the TME.

### Deciphering spatial architecture of TME through cell type mapping

Intra-tumor metabolic heterogeneity is often attributable to interactions of tumor cells with other components of the TME(32–34). To identify possible cellular interactions driving such heterogeneity, we compared different cell type deconvolution approaches in our Visium dataset, including annotation by a pathologist, two reference-free approaches (SNN clustering and STDeconvolve), and GIS-ROTA. A certified pathologist manually annotated tumor features, which further showed alignment of inflammatory cells along the border between adipocyte and tumor region in primary breast tumor from Patient 5 (**Figure 3A** and **Supplementary Figure 5A**). Adipocyte distribution was not only confirmed by the pathologist but also validated via mapping of *FABP4* mRNA expression, an adipocyte marker, as well as GESECA of the adipogenesis gene set (**Supplementary Figure 5B**). Similarly, tumor cell distribution was supported by mapping of *TFF1* and the estrogen response gene set. Pathology annotation remains the gold standard for tissue characterization due to high accuracy and expertise, yet it can be limited by subjectivity and considerable time requirements. Reference-free approaches, such as SNN clustering and STDeconvolve, recapitulated overall spatial organization as shown in pathology annotations, but the identified clusters contained multiple cell types rather than distinct individual ones (**Figure 3B,C** and **Supplementary Figure 5C**,D). These findings highlight the need for accurately mapping the spatial distribution of individual cell types within complex tissue architecture, which we addressed with our framework.

**Figure 3.**
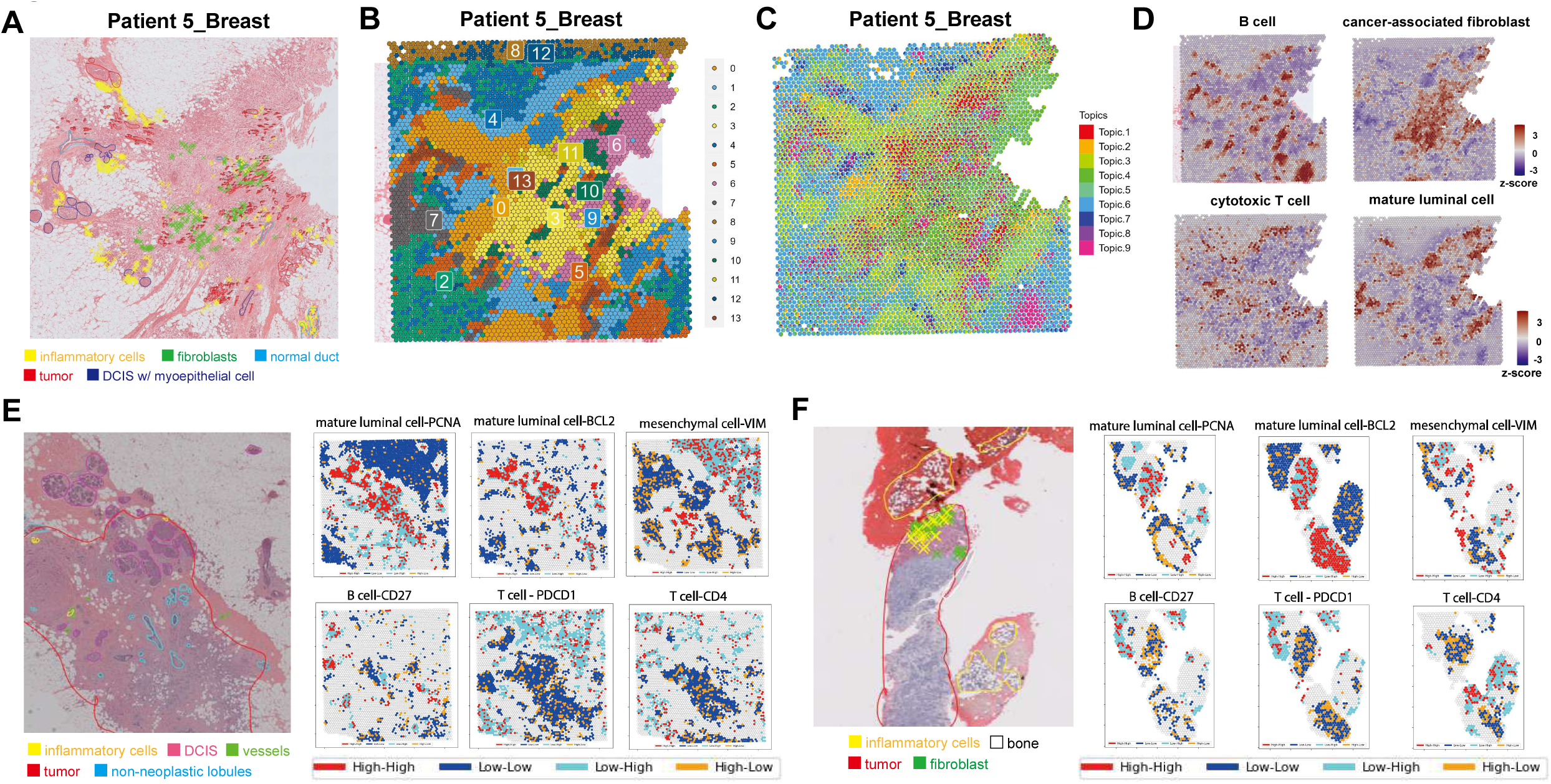
Deciphering spatial architecture of tumor microenvironment through cell type mapping. (A) Pathology annotations on primary breast tumor sample from Patient 5 (Patient 5_Breast). (B) Uniform manifold approximation and projection (UMAP) and spatial plot of UMAP clusters in Patient 5_Breast sample. (C) Mapping of deconvolved cell-type proportions on Visium data of Patient 5_Breast sample by STDeconvolve. (D) Spatial mapping of gene set coregulation analysis (GESECA) results for CellMarker 2.0 breast cancer-related cell types in Patient 5_Breast sample. Barcodes in red are those with high z-scores for the curated cell type, indicating that cell type member genes are collectively highly expressed in those regions. (E) Pathology annotations on an additional slide of primary tumor sample from Patient 5 submitted for Visium gene and protein expression analysis and visualization of local spatial autocorrelation based on bivariate Local Moran’s I of cell types and protein markers at *p* < 0.05. (F) Pathology annotations on additional slide of metastatic bone tumor sample from Patient 7 submitted for Visium gene and protein expression analysis and visualization of local spatial autocorrelation based on bivariate local Moran’s I of cell types and protein markers at *p* < 0.05.

To visualize the spatial distribution of cellular networks at the individual cell type level, we applied GESECA using a manually curated breast cancer cell marker database from CellMarker2.0. (**Figure 3D**). We validated our cell type mapping by visually comparing our results with the pathology annotations, observing close concordance. We also validated our approach using a more quantitative strategy, using Visium spatial gene and protein expression data that simultaneously capture gene and protein levels from tissue sections. We applied bivariate Local Moran’s I to GESECA results of cell types and their corresponding protein markers to evaluate the concordance between spatial distributions derived from GESECA mapping and protein-based mapping. We found extensive co-localization for cell types predominantly expressed in tumor (proliferative and structural markers) and significant co-absence (both absent) for less-abundant immune cell types in the tumor (**Figure 3E,F**). This congruency highlights the robustness of our approach for mapping spatial distribution of cell types using GESECA with manually curated cell type gene sets, reinforcing the initial step of our framework.

We next performed correlation analysis on the GESECA z-scores for breast cancer-related cell types from CellMarker 2.0. database, identified groups of cell types with distinct spatial correlation patterns (**Figure 4A**), and presented in partial heatmaps (**Figure 4B**). The partial heatmaps revealed two distinct groupings of cells, each showing positive intra-group spatial correlation: 1) immune cells (tumor-associated macrophages, cytotoxic T cells, CD4+ T cells), cancer-associated fibroblasts, and endothelial cells; and 2) proliferating cells (mature luminal cells, EMT cancer stem cells), basal cells, and B10 regulatory B cells. These two cell groups exhibited negative correlation with each other. However, as these spatial heatmap correlations do not guarantee co-localization or mutual exclusivity, visual comparisons of GESECA mapping results as well as Local Moran’s I application were followed. A visual comparison of cell types from the first group revealed similar spatial mapping within the tumors, with co-occurrence more clearly detected in primary tumors (**Figure 4C**). As predicted from the correlation pattern in the heatmap, mature luminal cells and B10 regulatory cells were closely mapped but in areas mutually exclusive from the immune cell within the first group. After visual estimations, we applied bivariate Local Moran’s I to quantitatively assess these spatial correlations and identify potential spatial coregulations indicative of cellular interactions affecting tumor progression. Primary and metastatic tumors had extensive co-localization and co-absence of macrophage and cytotoxic T cells, whereas in the same region, mutual exclusivity was observed between cytotoxic T cell and mature luminal cell (**Figure 4D**). This reflected a low frequency of proliferative cells in regions of high immune cell enrichment and vice versa. Interestingly, B10 regulatory cells showed extensive co-localization with mature luminal cells and mutual exclusivity with cytotoxic T cells, but such spatial correlation was lost in metastatic tumors (**Figure 4E**). To assess clinical relevance of these findings, we explored the potential role of B10 cells, a subset of regulatory B cells with high IL10 production. As a proxy for B10 cells, we examined *IL10* expression in breast tumor progression using publicly available datasets. High *IL10* levels in primary tumors were associated with shorter distant metastasis-free survival (hazard ratio: 1.44; 95% confidence interval: 1.07–1.94; *p* = 0.015) (**Figure 4F**). Additionally, cBioPortal analysis of the Metastatic Breast Cancer Project cohort showed that 54% of patients had mutations in *IL10* genes, considerably higher than other clinically relevant genomic alterations including *ESR1*, *PIK3CA*, and *FGFR1*, with most mutations being amplifications (**Figure 4G**). Overall, GIS-ROTA not only captured biologically meaningful spatial regulation of cell types but also uncovered a potential role of B10 regulatory cells in driving worse prognosis in metastatic breast cancer patients, consistent with supporting clinical data.

**Figure 4.**
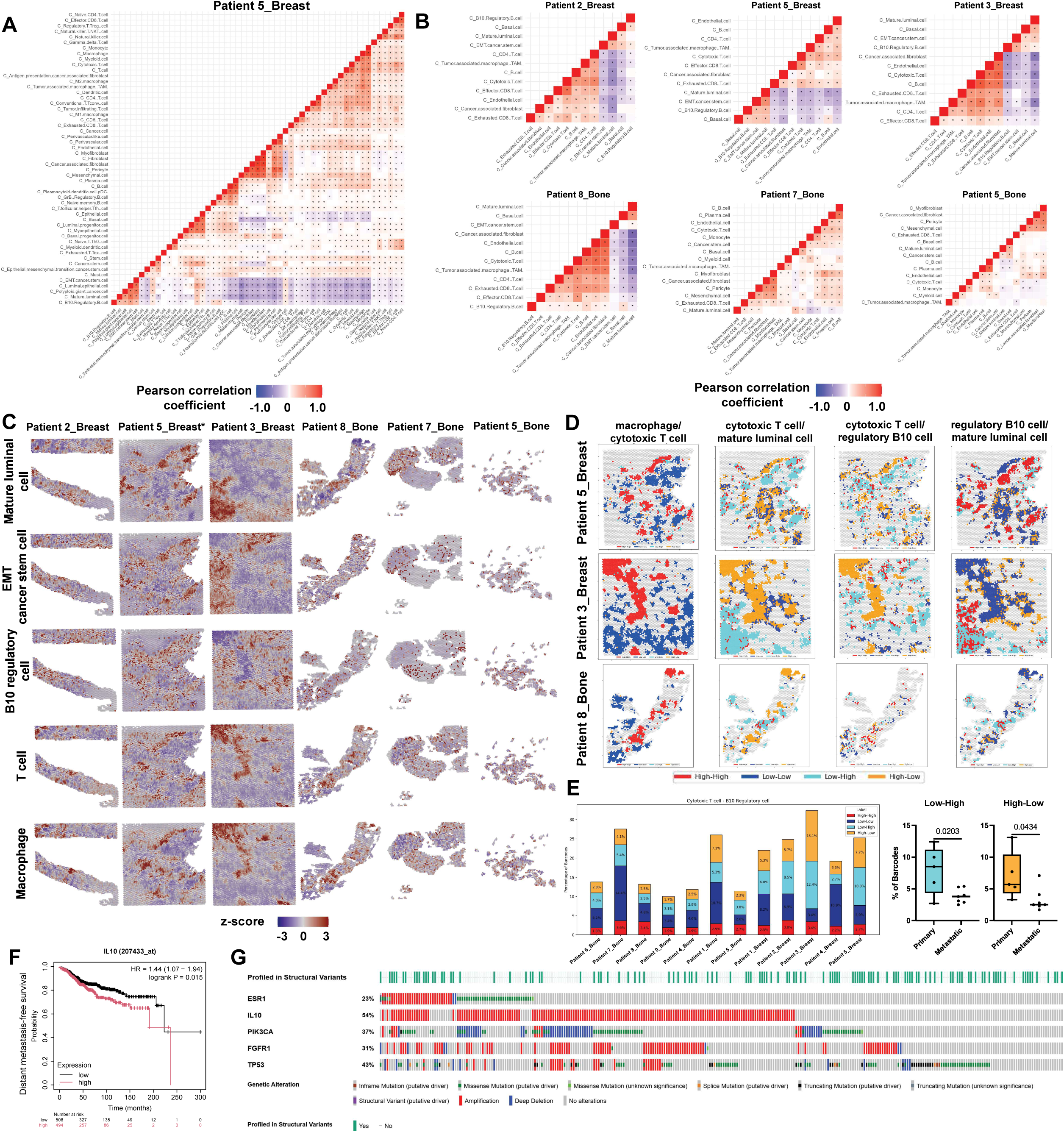
Identification of a potential immunomodulatory role for IL-10 in the ER+ breast tumor microenvironment. (A) Heatmap representing the correlation matrix between gene set coregulation analysis (GESECA) z-scores of breast cancer-related cell types in primary breast tumor sample from Patient 5. Grids with asterisks represent significant coefficients at *p* < 0.05. (B) Partial heatmaps of strong correlation in primary breast and metastatic bone tumors. (C) Spatial mapping of GESECA results for breast cancer-related cell types. Barcodes in red are those with high z-scores, indicating that cell type member genes are collectively highly expressed in those regions. (D) Visualization of local spatial autocorrelation based on bivariate local Moran’s I of macrophage, cytotoxic T cell, mature luminal cell, and regulatory B10 cell at *p* < 0.05. (E) Barplot displaying the proportion of barcodes assigned to each spatial cluster(left). Boxplot and p-values of the t-test comparing the percentage of barcodes for Low-High and High-Low cluster between primary breast and metastatic bone tumors (right). The percentage for each cluster is estimated relative to total number of barcodes per sample. (F) Distant metastasis-free survival curve of *IL10* level in estrogen receptor positive/HER2 negative breast cancer patient data plotted using KMPlotter online database. (G) cBioPortal data visualization and analysis showing genomic alteration frequency in *ESR1, IL10, PIK3CA, FGFR1*, and *TP53* in 379 breast cancer patient samples from the Metastatic Breast Cancer Project. *GESECA mapping of mature luminal cell for Patient 5_Breast sample is the same as shown in Figure 3D.

## Discussion

We developed GIS-ROTA, a novel GIS-augmented computational framework that begins with applying hypothesis-driven biological signatures to spatial data using GESECA. This biologically-informed data is subsequently reconstructed into an optimized structure for Local Moran’s I, a robust geospatial metrics that quantify spatial autocorrelation. This quantitative assessment of spatial co-regulation defines four spatial domains that highlight co-localization, co-absence, and mutual exclusivity between pathway or cell type gene sets. Unlike conventional approaches where SVGs drive spatial domain identification followed by post hoc interpretations, GIS-ROTA directly interrogates biologically meaningful spatial domains and identifies regions where specific pathways or cell types show coordinated spatial co-regulations. Our framework uncovered TME complexity that may highlight tumor prognosis and therapy response, from intra-tumor metabolic heterogeneity of endocrine-resistant cell populations to possible cellular interactions that drive immunomodulation in ER+ primary and metastatic breast tumors. We initially developed our framework using Visium data from a metastatic breast cancer xenograft mouse model and tested its generalizability on patient samples. As such, our framework is compatible with spatial -omics datasets of any tissue type as well as any curated gene sets of interest, providing a robust targeted approach to study tissue heterogeneity.

Previous studies leveraged unsupervised clustering, one of the standard Visium analytic approaches, to elucidate spatial distribution of cells within tissue microenvironments(35,36). In our Visium dataset of metastatic tumors, the UMAP plot revealed distinct transcriptional profiles between vehicle and fulvestrant-treated groups, corroborating findings from bulk- or single-cell RNAseq(37–39). However, we also observed some functional similarities between clusters, with overlapping up- or downregulation of gene sets across clusters. Such overlap might be attributed to density-based clustering, which may not fully account for spatial context(28). The TME is a heterogeneous ecosystem of cells whose functional relationships underlie varying therapy response across tumor regions(5,6), highlighting the necessity of spatial analysis. Identifying spatially variable genes, which might represent the spatial relationships of cells, is crucial to identify potential biomarkers with clinically meaningful signals associated with tumor progression(21,40). Spatial variation is often attributable to differences in cellular composition, functional dependencies, or cell–cell interactions(41). Cellular processes often affect groups of genes that share biological functions(42), and with higher effectiveness revealed by gene set analysis rather than single genes, we applied Local Moran’s I to results from GESECA, which examines spatial datasets at a gene set level. We found extensive co-localization between estrogen response and cholesterol homeostasis, which is supported by studies demonstrating dysregulated cholesterol homeostasis is associated with increased tumorigenicity and metastasis(43–46). We also identified mutual exclusivity between estrogen response and EMT, as demonstrated in breast cancer studies(47,48). Despite comparable findings with bulk or single-cell RNAseq, our hotspot analysis explicitly incorporates the spatial contexts among biological functions and conceptualizes those spatial relationships, allowing further downstream analyses on highlighted regions. Our approach also explores different pathways on an individual basis, avoiding the possible ambiguity from methods that consider statistical clustering based on entire sets of gene expression. With growing interest in applying computational geography methods to spatial transcriptomics data, previous studies leveraged GIS metrics to fully utilize the spatial context. Existing approaches include employing geographically weighted PCA(49), leveraging Moran’s I to detect regions enriched with spatially variable genes(19), and applying generalized linear framework(23). The key distinction between these methods and GIS-ROTA lies in the timing of biological interpretation. Conventional approaches determine biological relevance post hoc, whereas GIS-ROTA incorporates biological context prior to domain identification, eliminating the need for intermediate dimensionality reduction or additional interpretation. This allows researchers to directly examine their hypothesis, rather than discovering spatial patterns and subsequently assessing their biological relevance.

We expanded the applicability of GIS-ROTA by testing its generalizability to Visium dataset of primary and metastatic tumors from breast cancer patients. We found minimal overlap of SNN clusters mapping across patient samples, consistent with breast cancer studies reporting inter-tumor heterogeneity in RNA seq data from patients(50–52). Our bivariate Local Moran’s I analysis revealed comparable results to xenograft data analyses, such as co-localization of endocrine therapy response with metabolic pathways as well as mutual exclusivity with EMT. We found some additional spatial cross-correlations with estrogen response, including co-localization with cell-cycle pathways and mutual exclusivity with immune response pathways, due to fundamental developmental differences between mouse and human tumors(53) since we used immunocompromised mouse models, which fail to fully capture the intact TME of patient samples.

In addition to inter-tumor heterogeneity, we also observed intra-tumor heterogeneity as noted by some co-localized regions mapping to different tumor areas, varying by metabolic pathways. With the Visium dataset of patient tumor samples, we examined whether any cellular interaction might have contributed to such heterogeneity by applying GESECA to a breast cancer-related cell type database. Similarity of our transcript-based annotations, corresponding multiplex protein assay, and the pathologist’s annotations suggests the validity of our approach of GESECA-based cell type mapping. Further, our approach identified different subsets of immune cells, albeit less abundant in TME, suggesting the potential role of our framework to help pathologists produce high-quality annotations. In fact, by mapping and examining spatial correlation between different cancer-related cell types, we found several important cellular networks in tumors. Notably, we found mutual exclusivity between mature luminal cells and cytotoxic T cells in primary tumors. In the same area, we also found extensive co-localization of mature luminal cells with B10 regulatory cells. Immunotherapy has been a mainstay for cancer patients, yet low efficacy is reported with hormone receptor-positive metastatic breast cancer due to its nature of being “immunologically cold” with fewer infiltrating T cells(54,55). Findings from bivariate hotspot analyses on primary tumors suggested a possible role of estrogen receptor and IL10 signaling in dysregulating T cell activity in tumors, yet the role of IL10, a pleiotropic immune-regulatory cytokine(56), in breast tumor immunomodulation remains controversial(57,58) and warrants further validation in future studies.

Despite the robustness of our analytic approaches, there are limitations. One limitation is the low resolution, as each spot within a Visium capture area is 55 μm in diameter, with 1–10 cells per spot. While the cell type mapping from our approach was well validated with pathology annotation and Visium proteomics results, studies with higher-resolution data of multiple samples from the same tumor are still warranted to achieve more precise understanding of the cellular activities within their spatial context. We also acknowledge that all primary tumors had metastasized. Future studies exploring primary tumors that have not metastasized may uncover more substantial differences compared to those that have metastasized, thereby providing a clearer basis to compare spatial cellular relationships between primary and metastatic tumors.

Overall, GIS-ROTA pre-defines biological functions of interest with established gene sets, maps spatial enrichment of these functions, and identifies spatial relationships between biologically relevant features. Our approach directly produces interpretable results without requiring additional biological annotation, which differentiates from existing unsupervised approaches. Given the robust performance of our framework in hypothesis-driven spatial analysis and its potential generalizability to diseases beyond cancer, GIS-ROTA paves the way to advance development of personalized treatments.

## Methods

### Sample preparation

An animal study was previously conducted in accordance with NIH standards for the use and care of animals, with protocols approved by the University of Illinois at Urbana-Champaign (IACUC protocol #20158)(26). Four-week-old, ovariectomized, NOD SCID gamma immunodeficient female mice were received from Jackson Laboratory. After one week of acclimatization, mice were injected with MCF7-*ESR1*^Y537S^ cells (RRID:CVCL_0031) resuspended in 1% PBS via tail vein and randomized into vehicle and fulvestrant-treated groups. Fulvestrant, prepared in 10% dimethyl sulfoxide and 90% corn oil, was administered by intramuscular injection (100 mg/kg) twice per week for four weeks. After six weeks, mice were euthanized, and four fresh-frozen liver samples were collected for Visium library preparation (GEO database: GSE236681).

Fourteen primary and metastatic bone tumor biopsies—including three pairs of matching samples (Breast 6994, Bone 8713) (Breast 6997, Bone 8712) (Breast 8714, Bone 8715)—from breast cancer patients were collected from the Carle Foundation Hospital. Sample collection and processing were performed by a trained Carle professional in accordance with the “HT.300.003 – Preparation of Specimens for Gross Examination and Possible Submission to Carle Tissue Repository” policy and procedure. To briefly summarize, the tumor was initially fixed in formalin and then placed in a processor for ∼12 h. After processing, tissue was embedded in a block, cut on a microtome, and mounted on a glass slide for Visium library preparation. A certified pathologist manually annotated fibroblasts, immune cells, tumor, and other structural features within H&E images of tumor samples. Among 14 samples, one primary breast tumor and one metastatic bone tumor sample were submitted for Visium CytAssist gene and protein expression analysis, which allows simultaneous capture of protein and whole transcriptome expression. The remaining 12 samples were submitted for Visium gene expression analysis. Gene expression data are available in the GEO database (GSE298286).

### Library construction and sequence alignment

Visium libraries were prepared by the Carl R. Woese Institute for Genomic Biology Core Facility and the DNA Services laboratory of the Carver Biotechnology Center at the University of Illinois at Urbana-Champaign. Library preparation was performed according to 10X Genomics Visium fresh-frozen(59), CytAssist formalin-fixed paraffin-embedded gene expression(60), or gene and protein expression(61) protocol, depending on the tissue preparation method. A detailed explanation of library preparation has been published(26). Spatial gene expression data were processed and visualized using Spaceranger version 2.1.0 and Loupe Browser version 6.0. Barcoded reads were mapped to either human (GRCh38) or combined human and mouse reference genome (GRCh38 and GRCm39) from Ensembl Release 107 based on downstream analyses. The Visium gene and protein expression assay uses a 10X Genomics human immune cell profiling panel, which contains 35-plex CytAssist panel antibodies targeting immune cell profiling markers(62).

### Preprocessing and subsequent downstream analyses

Secondary analyses were performed using Seurat (version 3.0.0), a R package to analyze spatial RNAseq data(63). We initially applied a conventional graph-based clustering method to our spatially resolved dataset to identify cell populations through an unbiased approach. We performed unsupervised clustering based on the whole transcriptome to examine cell population groupings through a shared nearest neighbor (SNN) modularity optimization-based clustering algorithm with default parameters. Sequenced reads were mapped to the combined human and mouse reference genome, merged, and filtered based on the following criteria: 1) genes expressed in <10 barcodes, and 2) barcodes without any gene expression. The filtered matrix was used as input for SCTransform, which includes normalization, variance stabilization, and feature selection steps.

Pre-processed data proceeded to principal component analysis for dimensionality reduction, followed by SNN graph construction, cluster determination, and UMAP dimensional reduction. We visualized clusters by plotting them in UMAP space and overlaying them on histology images of samples. To identify molecular features of clusters, we performed differential expression analysis using the FindMarkers function and retrieved spatially variable genes per cluster(63). We then performed gene set enrichment analysis (GSEA) on these genes via fgsea R package (version 1.29.1)(29), using either human or mouse input identifiers from MSigDB curated gene sets. Since clustering was performed on reads mapped to the combined human and mouse reference, we initially assessed the enrichment percentage of human or mouse transcripts in each cluster and selected the curated gene set for GSEA accordingly(26). We also mapped the distribution of individual genes of interest using the SpatialFeaturePlot function from Seurat. When checking tumor-specific transcripts, we used the human reference (GRCh38)-mapped dataset.

### Gene set co-regulation analysis (GESECA)

GESECA estimates the contribution of gene sets to expression variability of the whole sample. We checked the degree of co-regulation of hallmark gene sets from Human MSigDB collections using the GESECA function with the human reference-mapped dataset as input. Principal component analysis on the barcode x gene matrix (rev.pca = TRUE) was initially performed for dimensionality reduction, and gene sets were selected from msigdbr R package (version 1.14.0)(64). Z-scores of different hallmark gene sets were estimated per barcode, indicating expression variance explained by a particular gene set. Z-scores per barcode were mapped using plotCoregulationProfileSpatial function, visualizing the distribution of different gene sets within each sample. All functions used for GESECA are available from the fgsea R package. We additionally applied GESECA to manually curated cell markers from the CellMarker 2.0 database when analyzing primary and metastatic tumor samples from breast cancer patients(65). This enabled mapping of different breast cancer-related cell types across samples. To select cell types that best matched our dataset, we filtered the cell marker database on the following criteria: species = human, tissue type = breast, cell type = cancer cell. Of note, we applied SCTransform to individual samples and thus used within-sample normalized datasets when examining cell type distribution. We focused on spatial and cellular variations specific to individual samples and preserving inherent cellular distribution patterns that might be unique to each sample.

### AnnData curation and hotspot identification using local Moran’s I

Hotspot analysis is a spatial analysis technique commonly used in geospatial studies to detect statistically significant clusters of high or low values(66–68). Spatial autocorrelation, one of the methods to detect hotspots, suggests tendency of gene expression to be more positively or negatively correlated at nearby locations than at distant locations, whereas spatially random genes fail to display such correlations. To identify potential clustering of spatially correlated genes within a tissue, we used local Moran’s I, a local measure of spatial association. The local Moran’s I statistic measures deviation of gene expression from the global average expression for a specific location (e.g., barcode) and compares to deviations in neighboring barcodes—typically within a defined spatial neighborhood, which would be the six surrounding barcodes in the Visium dataset(69). This enables identification of outliers whose direction of local deviation differs from that of their neighbors. The local Moran’s I is given by:

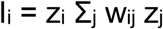

where *z* represents deviation from average gene expression across the sample for a given barcode *i* and its neighbor location *j*. *w_ij_* denotes the spatial weights assigned to each neighboring barcode, yet given the arrangement of neighboring barcodes in hexagonal packing with equal distances from the central barcode, w_ij_ = 1/6, resulting in a total weight sum of w_ij_ (Σ_j_ w_i_j) = 1.

To conceptualize the relationship among spatially associated genes in the Visium dataset, we developed a Python package that imports and reconstructs ST data into AnnData structure and applies a local spatial association method for hotspot analyses. We imported local spatial association methods from libpysal (version 4.8.1), a Python library for spatial data analysis. A pre-processed Visium output matrix comprises a two-dimensional matrix with barcodes and normalized gene expression values, which was integrated with barcode location (x and y) matrices to reconstruct an AnnData structure for each sample. Imported local spatial association methods (e.g., univariate and bivariate local Moran’s I) are then applied to the AnnData structure in a themeless manner, quantifying the degree of clustering of high- or low-gene expression regions. We identified contiguous barcodes with significant gene expression localization and tested the significance of spatial association using unadjusted *p* = 0.05(70). Barcodes with neighboring barcodes of similar gene expression levels show positive local Moran’s I index, whereas those with dissimilar values show negative index. A positive autocorrelation index indicates that a barcode has neighboring barcodes with similarly high or low expression of a particular gene: High–High or Low–Low spots; a negative index indicates dissimilar expression levels: High–Low or Low–High spots. Overall, four labels were generated and assigned to each barcode based on bivariate Moran’s I analysis results (red: High–High, co-expression of both genes; yellow: High–Low, high expression of first gene, low expression of second gene (mutual exclusivity); light blue: Low–High, low expression of first gene, high expression of second gene (mutual exclusivity); blue: Low–Low, co-absence of both genes). The resulting four classes of local spatial association can be further analyzed using other bioinformatics packages to detect biomarkers specific to each cluster group. We also reconstructed AnnData structure from the GESECA output matrix consisting of barcodes and z-scores of enriched pathways or cell types. The same principles were applied, where we leveraged univariate local Moran’s I to detect spatial clusters for an individual pathway and bivariate local Moran’s I for two different pathways.

### Hierarchical clustering and heatmap visualization of spatial correlation

Incorporation of bivariate hotspot analysis on these pathways into our framework allowed us to validate whether such positive (or negative) spatial correlation from the heatmap was due to extensive co-localization (or mutual exclusivity) rather than random dispersion with matching barcodes. Among 50 hallmark gene sets, we plotted the spatial heatmap for early estrogen response, associated with endocrine response, with all other hallmark gene sets. When exploring autocorrelations between cell types, however, we plotted heatmaps for individual samples and compared correlation coefficients of all available cell types within each sample.

### Unsupervised machine learning approach for cell-type deconvolution

In parallel to our guided approach of cell-type deconvolution via GESECA using curated cell type markers, we leveraged a reference-free, unsupervised approach to deconvolve our spatial dataset. We used STDeconvolve, a generative statistical model built on latent Dirichlet allocation(71). We used STDeconvolve to select the top 1000 genes likely to be informative of latent cell types, based on which latent Dirichlet allocation then inferred the transcriptional profile and proportional representation for each cell type per barcode. STDeconvolve also requires the number of cell types, *k*, to be set a priori, for which we selected lowest perplexity from the latent Dirichlet allocation model.

### Statistical analysis

GESECA generates results for multiple gene sets, and to avoid computational complexity we generated a spatial correlation heatmap to filter for gene sets with strong spatial associations. We first performed Pearson’s correlation analysis on z-scores per barcode across entire gene sets. A higher correlation coefficient from Pearson’s correlation analysis would suggest positive spatial correlation, whereas a lower coefficient would suggest negative spatial correlation. Hierarchical clustering was applied to these correlation coefficient values, which was then visualized as a heatmap. Based on the heatmap, we identified gene sets with specific patterns of spatial correlations, which were prioritized for hotspot analyses.

All statistical analyses of Visium datasets were performed using R software (version 4.3.2) and Python (version 3.12.4). Hierarchical clustering of correlation coefficient values was performed using Cluster 3.0 open-source clustering software, and the results were visualized in a heatmap using Java Treeview(72). Bar plots showing the proportions of barcodes for all four spatial clusters across samples were plotted using Python with the matplotlib library (version 3.10.3). Box plots comparing the percentage of barcodes for low–high and high–low spatial clusters were plotted using GraphPad Prism software (version 10.4.2), and *p*-values were calculated using Mann-Whitney U test.

## Acknowledgements

We acknowledge Drs. Alvaro Hernandez, Chris Wright, Jenny Drnevich, and Mayandi Sivaguru of the University of Illinois at Urbana-Champaign Roy J. Carver Biotechnology Center for their guidance. This work was supported by grants from the Personalized Nutrition Initiative (ZME, AS, and MGP), US Department of Agriculture National Institute of Food and Agriculture awards ILLU-698-924 and ILLU-698-331 (ZME) and Sylvia D. Stroup Scholar Award (ZME). Research reported in this publication was supported by a TiME T32 fellowship (JYY) and JPT fellowship (SA).

---

**Supplementary Figure 1.** (A) Spatial plot of SNN, k-means, BIRCH, FlowSOM clustering results on liver metastatic tumor samples from xenograft mouse model. (B) Uniform manifold approximation and projection (UMAP) plot labeled by individual samples (left) and SNN clusters in liver metastatic tumor samples (right). (C) Heatmap representing the correlation matrix between gene set coregulation analysis (GESECA) z-scores of hallmark early estrogen response and 179 KEGG gene sets from Molecular Signatures Database. (D) Visualization of local spatial auto-correlation based on univariate local Moran’s I at *p* < 0.05. (E) Visualization of local spatial cross-correlation based on bivariate local Moran’s I at *p* < 0.05.

**Supplementary Figure 2.** Uniform manifold approximation and projection (UMAP) plot labeled by individual samples (top) and clusters (bottom) defined from shared nearest neighbor (SNN) clustering (left), and spatial plot of SNN clustering results in primary breast and metastatic bone tumor samples (right).

**Supplementary Figure 3.** Heatmaps representing the correlation matrix between gene set coregulation analysis (GESECA) z-scores of hallmark pathways in primary breast and metastatic bone tumor samples. Grids with asterisks represent statistically significant correlation coefficient values at *p* < 0.05.

**Supplementary Figure 4.** Visualizations of local spatial autocorrelation based on bivariate local Moran’s I of hallmark early estrogen response (EER) with epithelial-mesenchymal transition (EMT), metabolic signaling, and cell cycle gene sets at *p* < 0.05. * Hotspot results for EER and EMT and those for EER and cholesterol homeostasis (CH) for primary breast tumors from Patient 2, 5, and 3, and metastatic bone tumors from Patient 9, 7, and 5 are the same as shown in Figure 2D. **BA: bile acid; FA: fatty acid; OXPHOS: oxidative phosphorylation.

**Supplementary Figure 5.** (B) Pathologist’s manual annotations on primary breast and metastatic bone tumor samples. (C) Spatial feature plot of fatty acid binding protein 4 (*FABP4*) and estrogen receptor target gene (*TFF1*), and spatial mapping of gene set coregulation analysis (GESECA) results for hallmark adipogenesis and early estrogen response gene sets in primary tumor sample from Patient 5 (Patient 5_Breast). (D) Spatial plot of the shared nearest neighbor (SNN) clustering results in primary breast and metastatic bone tumor samples. (E) Mapping of deconvolved cell-type proportions on Visium data by STDeconvolve. * Spatial mapping of GESECA results for hallmark early estrogen response gene set in Patient 5_Breast sample is the same as shown in Figure 2C.

