## Supplementary Figure 1 for "In silico reconstruction of primary and metastatic tumor architecture using GIS-augmented spatial transcriptomics"

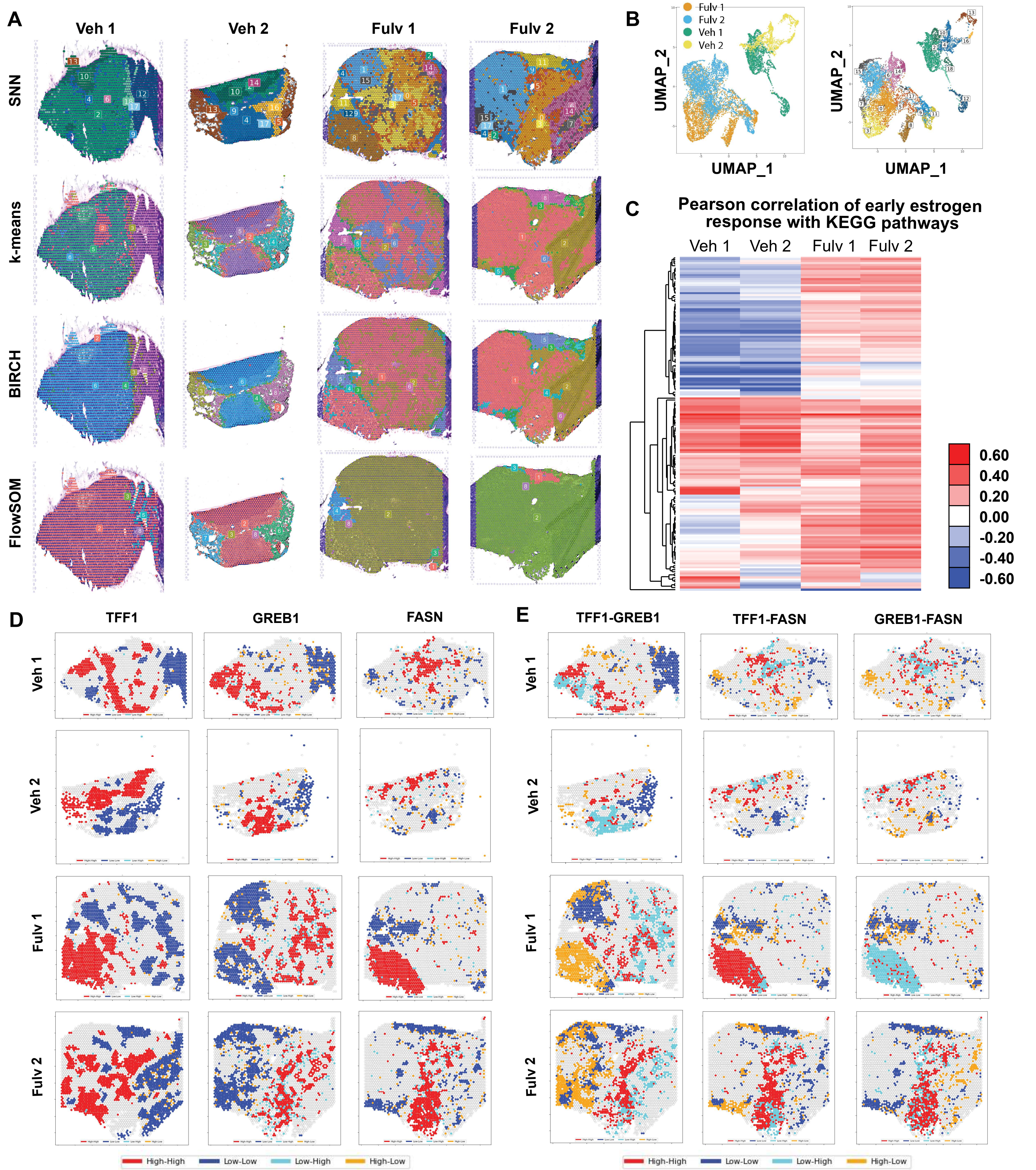


Supplementary Figure 1. (A) Spatial plot of SNN, k-means, BIRCH, FlowSOM clustering results on liver metastatic tumor samples from xenograft mouse model. (B) Uniform manifold approximation and projection (UMAP) plot labeled by individual samples (left) and SNN clusters in liver metastatic tumor samples (right). (C) Heatmap representing the correlation matrix between gene set coregulation analysis (GESECA) z-scores of hallmark early estrogen response and 179 KEGG gene sets from Molecular Signatures Database. (D) Visualization of local spatial auto-correlation based on univariate local Moran's I at $p < 0.05$. (E) Visualization of local spatial cross-correlation based on bivariate local Moran's I at $p < 0.05$.
