## Supplementary Figure 2 for "In silico reconstruction of primary and metastatic tumor architecture using GIS-augmented spatial transcriptomics"

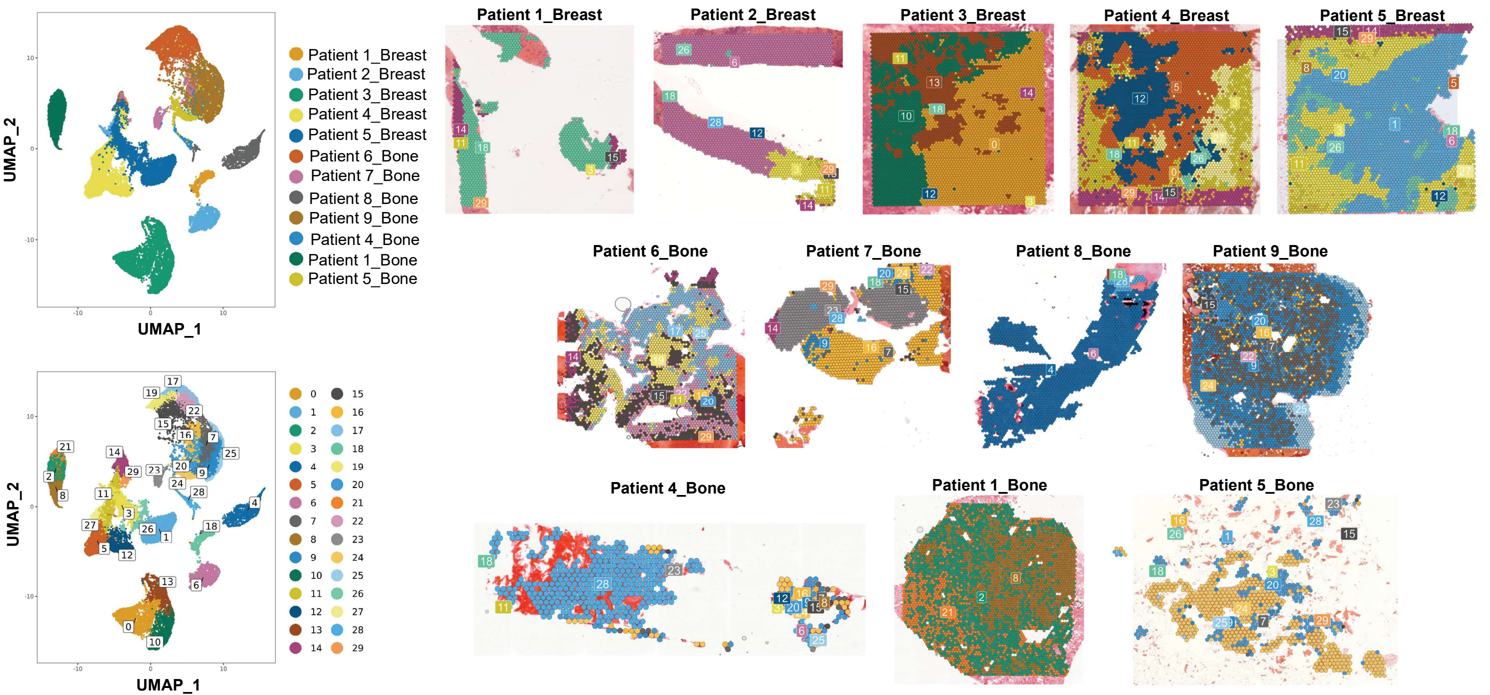


Supplementary Figure 2. Uniform manifold approximation and projection (UMAP) plot labeled by individual samples (top) and clusters (bottom) defined from the shared nearest neighbor (SNN) clustering (left), and spatial plot of SNN clustering results in primary breast and metastatic bone tumor samples (right).
