## Supplementary Figure 3 for "In silico reconstruction of primary and metastatic tumor architecture using GIS-augmented spatial transcriptomics"

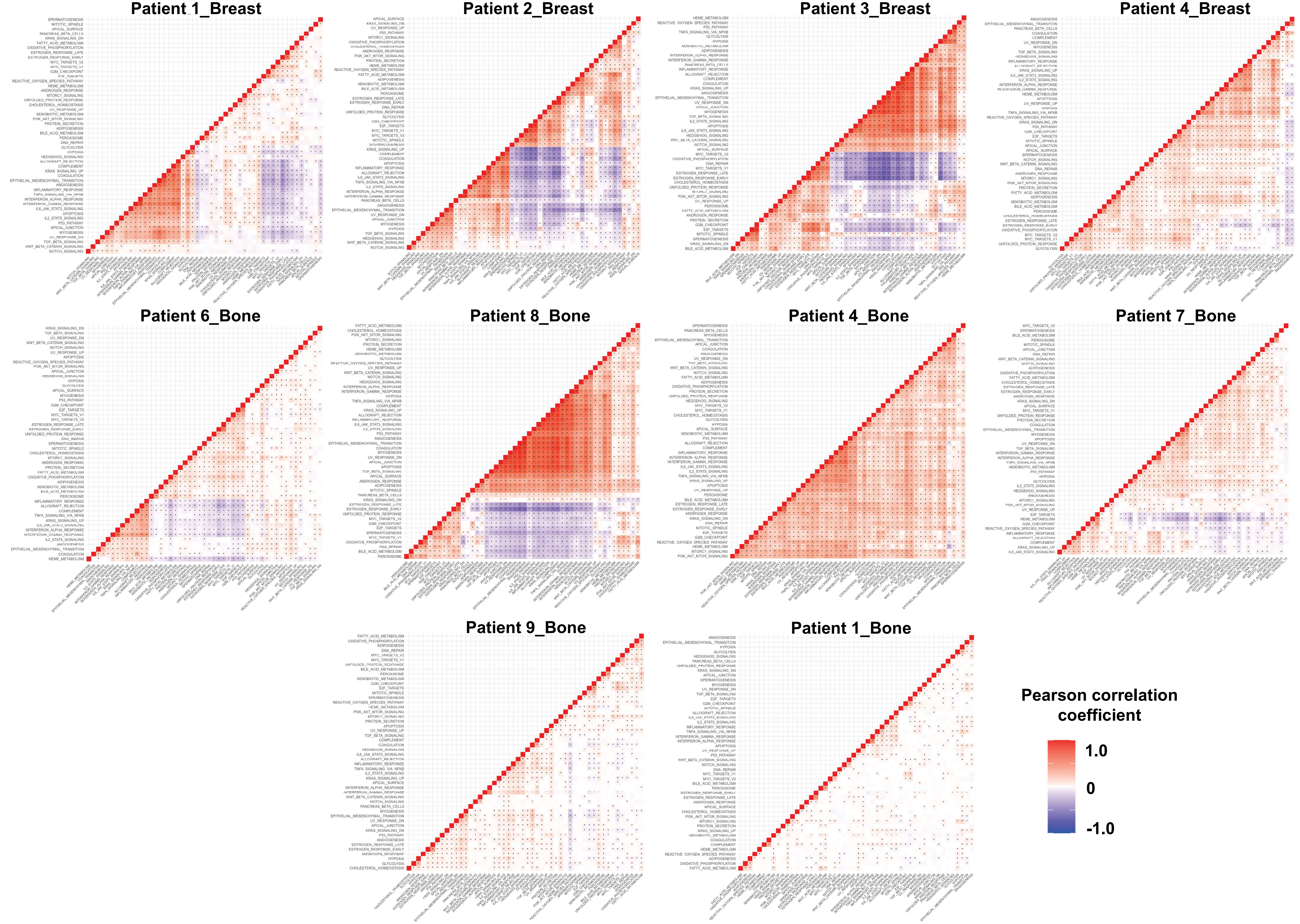


Supplementary Figure 3. Heatmaps representing the correlation matrix between gene set coregulation analysis (GESECA) z-scores of hallmark pathways in primary breast and metastatic bone tumor samples. Grids with asterisks represent statistically significant correlation coefficient values at $p < 0.05$.
