## Supplementary Figure 4 for "In silico reconstruction of primary and metastatic tumor architecture using GIS-augmented spatial transcriptomics"

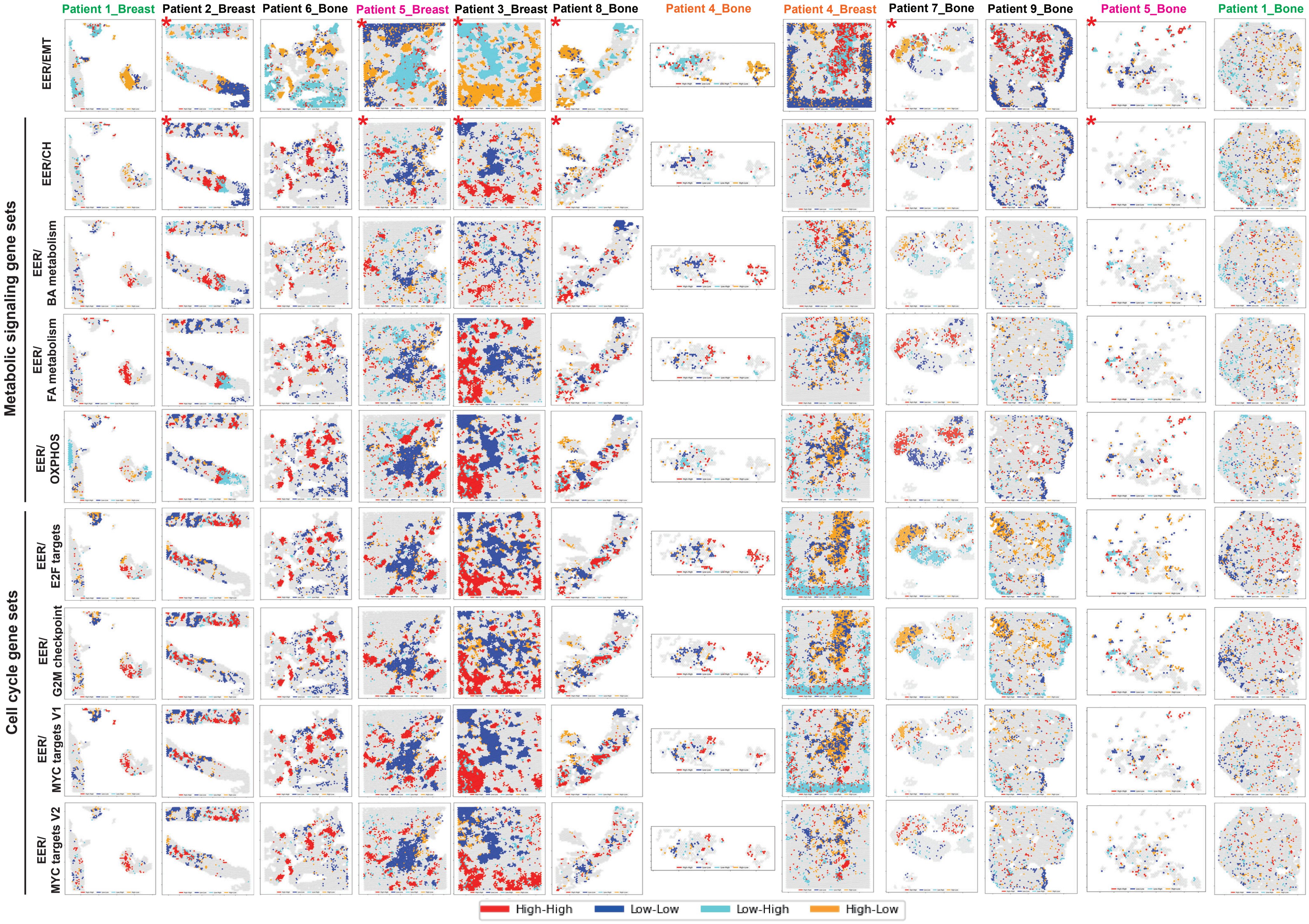


Supplementary Figure 4. Visualizations of local spatial autocorrelation based on bivariate local Moran's I of hallmark early estrogen response (EER) with epithelial-mesenchymal transition (EMT), metabolic signaling, and cell cycle gene sets at $p < 0.05$. * Hotspot results for EER and EMT and those for EER and cholesterol homeostasis (CH) for primary breast tumors from Patient 2, 5, and 3, and metastatic bone tumors from Patient 9, 7, and 5 are the same as shown in Figure 2D. **BA: bile acid; FA: fatty acid;
