## Supplementary Figure 5 for "In silico reconstruction of primary and metastatic tumor architecture using GIS-augmented spatial transcriptomics"

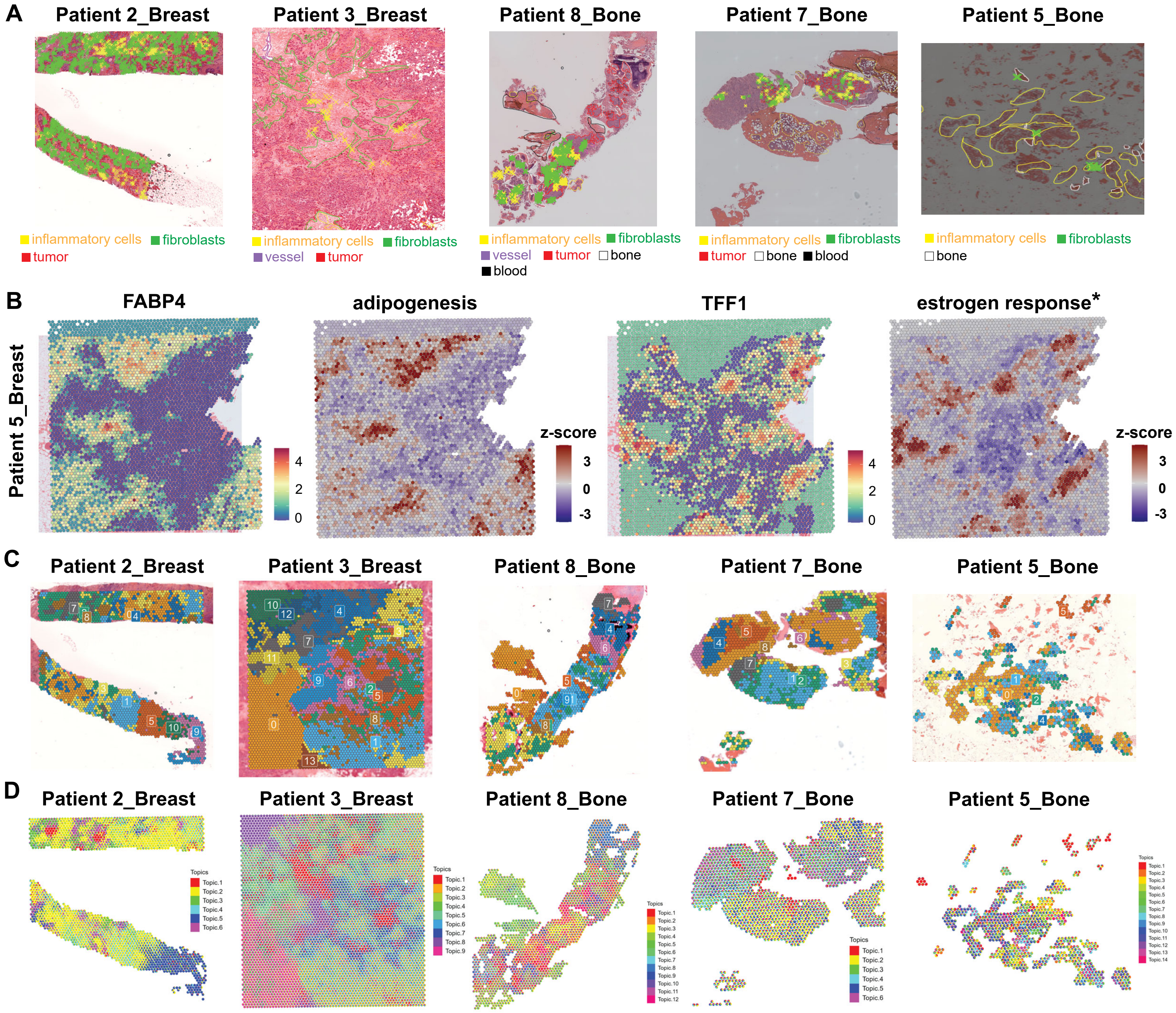


Supplementary Figure 5. (A) Pathologist's manual annotations on primary breast and metastatic bone tumor samples. (B) Spatial feature plot of fatty acid binding protein 4 (*FABP4*) and estrogen receptor target gene (*TFF1*), and spatial mapping of gene set coregulation analysis (GESECA) results for hallmark adipogenesis and early estrogen response gene sets in primary tumor sample from Patient 5 (Patient 5_Breast). (C) Spatial plot of the shared nearest neighbor (SNN) clustering results in primary breast and metastatic bone tumor samples. (D) Mapping of deconvolved cell-type proportions on Visium data by STDeconvolve. * Spatial mapping of GESECA results for hallmark early estrogen response gene set in Patient 5_Breast sample is the same as shown in Figure 2C.
