## Supplementary Table 1 for "In silico reconstruction of primary and metastatic tumor architecture using GIS-augmented spatial transcriptomics"

**Supplementary Table 1. Gene set enrichment analysis (GSEA) results on shared nearest neighbor (SNN) clusters enriched with human transcripts from metastatic tumors**

| Cluster 0 | pathway | pval | padj | ES | NES | nMoreExtreme | size | leadingEdge |
|---|---|---|---|---|---|---|---|---|
| 1 | HALLMARK_ESTROGEN_RESPONSE_EARLY | 0.000765 | 0.015723 | -0.31116 | -2.19623 | 0 | 35 | c("PRSS23", "TSKU", "MYC", "BHLHE40", "AREG", "SLC7A5", "KLK10", "SIAH2", "CXCL12", "STC2", "DLC1", "NBL1") |
| 2 | HALLMARK_ESTROGEN_RESPONSE_LATE | 0.000726 | 0.015723 | -0.34168 | -2.30228 | 0 | 32 | c("PRSS23", "AGR2", "GAL", "AREG", "SLC7A5", "KLK10", "SIAH2", "CXCL12", "SERPINA3", "NBL1") |
| 3 | HALLMARK_E2F_TARGETS | 0.000943 | 0.015723 | -0.43961 | -2.2325 | 1 | 18 | c("MYC", "PTTG1", "ANP32E", "CDKN3", "MAD2L1", "MYBL2", "H2AX", "BIRC5", "HMGA1", "SRSF1", "XPO1") |
| 4 | HALLMARK_G2M_CHECKPOINT | 0.001271 | 0.015893 | -0.40395 | -2.53839 | 1 | 27 | c("MT2A", "MYC", "SLC7A5", "PTTG1", "CDKN3", "MAD2L1", "MYBL2", "H2AX", "UBE2C", "BIRC5", "HMGA1", "BCL3", "SRSF1", "XPO1", "ODF2") |
| 5 | HALLMARK_UV_RESPONSE_DN | 0.010128 | 0.101275 | -0.47395 | -1.87171 | 26 | 11 | c("ID1", "MYC", "BHLHE40", "DLC1", "MGMT", "RXRA", "PMP22") |
| 6 | HALLMARK_COAGULATION | 0.013797 | 0.114979 | -0.57883 | -1.82519 | 43 | 7 | c("PRSS23", "FGG") |
| 7 | HALLMARK_CHOLESTEROL_HOMEOSTASIS | 0.036261 | 0.161182 | 0.471569 | 1.622501 | 269 | 12 | c("FASN", "CD9", "LSS", "SCD", "TP53INP1", "ABCA2", "TM7SF2", "ALCAM") |
| 8 | HALLMARK_DNA_REPAIR | 0.038684 | 0.161182 | 0.402155 | 1.616332 | 307 | 19 | c("NME3", "CCNO", "TAF1C", "POLD4", "EDF1", "SRSF6", "CANT1", "ADCY6", "COX17", "VPS28", "SUPT5H", "ERCC3", "POLR2J", "NT5C") |
| 9 | HALLMARK_MYOGENESIS | 0.038443 | 0.161182 | 0.387742 | 1.610021 | 311 | 21 | c("SPDEF", "MYO1C", "ITGB5", "SCD", "SYNGR2", "SH2B1", "GABARAPL2", "PLXNB2", "ATP6AP1", "EIF4A2", "GAA") |
| 10 | HALLMARK_APICAL_SURFACE | 0.030565 | 0.161182 | 0.766071 | 1.555557 | 183 | 3 | c("GATA3", "SULF2") |
| 11 | HALLMARK_MYC_TARGETS_V1 | 0.038123 | 0.161182 | -0.35596 | -1.59715 | 90 | 14 | c("MYC", "LDHA", "FBL", "MAD2L1", "NCBP2", "SRSF1", "XPO1") |
| 12 | HALLMARK_EPITHELIAL_MESENCHYMAL_TRANSITIO N | 0.031717 | 0.161182 | -0.34142 | -1.63796 | 69 | 16 | c("AREG", "FLNA", "PPIB", "CXCL12", "LGALS1", "PMP22", "QSOX1", "TIMP1", "SAT1", "COPA") |
| 13 | HALLMARK_ANDROGEN_RESPONSE | 0.053026 | 0.203947 | 0.464974 | 1.55007 | 388 | 11 | c("SPDEF", "ABHD2", "SCD", "TMPRSS2", "DHCR24", "KRT19", "APPBP2", "DBI") |
| 14 | HALLMARK_HYPOXIA | 0.061424 | 0.21698 | -0.30014 | -1.4765 | 131 | 17 | c("MT2A", "BHLHE40", "LXN", "SIAH2", "STC2", "LDHA") |
| 15 | HALLMARK_BILE_ACID_METABOLISM | 0.065094 | 0.21698 | 0.563577 | 1.492208 | 435 | 6 | c("DHCR24", "ABCA2", "GSTK1", "SULT2B1", "CROT") |
| 16 | HALLMARK_MYC_TARGETS_V2 | 0.073577 | 0.221847 | -0.82884 | -1.45589 | 317 | 2 | c("MYC", "SLC29A2") |
| 17 | HALLMARK_ANGIOGENESIS | 0.075428 | 0.221847 | -0.82711 | -1.45284 | 325 | 2 | c("FGFR1", "TIMP1") |
| 18 | HALLMARK_IL6_JAK_STAT3_SIGNALING | 0.115026 | 0.319517 | 0.600686 | 1.365697 | 727 | 4 | c("CD9", "STAT2", "TYK2") |
| 19 | HALLMARK_APICAL_JUNCTION | 0.188218 | 0.49531 | 0.314654 | 1.285084 | 1507 | 20 | c("EVL", "MDK", "CLDN4", "INPPL1", "PKD1", "CLDN7", "MYL12B", "JUP", "PIK3R3", "CDH1", "AKT2", "CTNND1", "SYMPK") |
| 20 | HALLMARK_NOTCH_SIGNALING | 0.212936 | 0.510875 | -0.45993 | -1.23688 | 743 | 5 | c("SKP1", "DTX2", "ARRB1", "HES1", "LFNG") |
| 21 | HALLMARK_IL2_STAT5_SIGNALING | 0.214568 | 0.510875 | 0.320838 | 1.245411 | 1684 | 17 | c("MUC1", "MYO1C", "SERPINB6", "SYNGR2", "ST3GAL4", "NCOA3") |
| 22 | HALLMARK_TNFA_SIGNALING_VIA_NFKB | 0.275222 | 0.616467 | -0.26402 | -1.1462 | 681 | 13 | c("MYC", "BHLHE40", "AREG") |
| 23 | HALLMARK_MTORC1_SIGNALING | 0.283575 | 0.616467 | 0.285801 | 1.167244 | 2271 | 20 | c("SYTL2", "CD9", "SLC9A3R1", "SCD", "PFKL", "DHCR24", "ACLY", "TM7SF2", "XBP1", "GGA2", "PIK3R3", "SERP1") |
| 24 | HALLMARK_ADIPOGENESIS | 0.347905 | 0.724803 | 0.354637 | 1.098071 | 2482 | 9 | c("PFKL", "DGAT1", "ADCY6", "ACLY", "MIGA2", "CD151", "DECR1", "PIM3", "LTC4S") |
| 25 | HALLMARK_PEROXISOME | 0.382514 | 0.743542 | 0.344308 | 1.066089 | 2729 | 9 | c("DHCR24", "FIS1", "CTBP1", "GSTK1", "SULT2B1", "ERCC3") |
| 26 | HALLMARK_KRAS_SIGNALING_UP | 0.386642 | 0.743542 | -0.33246 | -1.04833 | 1232 | 7 | c("SERPINA3", "TSPAN1", "SEMA3B", "RBM4", "AKT2", "JUP", "CROT") |
| 27 | HALLMARK_TGF_BETA_SIGNALING | 0.404236 | 0.748585 | -0.35198 | -1.03027 | 1335 | 6 | c("ID1", "ID3", "PPP1CA") |

| | | | | | | | | |
|---|---|---|---|---|---|---|---|---|
| 28 | HALLMARK_OXIDATIVE_PHOSPHORYLATION | 0.421513 | 0.752702 | 0.248632 | 1.032392 | 3420 | 21 | c("ACADVL", "ATP6V1G1", "ATP1B1", "RHOT2", "NDUFA1", "ATP6AP1", "TCIRG1", "COX7A2", "COX17") |
| 29 | HALLMARK_HEDGEHOG_SIGNALING | 0.55246 | 0.806689 | -0.43043 | -0.91289 | 2200 | 3 | c("SLIT1", "LDB1", "TLE3") |
| 30 | HALLMARK_INFLAMMATORY_RESPONSE | 0.51255 | 0.806689 | -0.44446 | -0.94264 | 2041 | 3 | c("MYC", "TIMP1") |
| 31 | HALLMARK_XENOBIOTIC_METABOLISM | 0.561314 | 0.806689 | 0.235814 | 0.915369 | 4407 | 17 | c("ESR1", "CSAD", "DCXR", "MCCC2", "SPINT2", "CROT", "FBP1", "JUP", "BLVRB", "IGFBP4", "MARCHF6", "BCAR1") |
| 32 | HALLMARK_GLYCOLYSIS | 0.488599 | 0.806689 | -0.19767 | -0.9724 | 1049 | 17 | c("STC2", "LDHA", "IDUA", "PYGB", "QSOX1", "CACNA1H", "NOL3", "SDC1", "HDLBP", "ELF3", "CHPF2", "AKR1A1", "GFUS", "CLDN3", "GMPPA", "BIK") |
| 33 | HALLMARK_HEME_METABOLISM | 0.537798 | 0.806689 | 0.316935 | 0.933923 | 3741 | 8 | c("SELENBP1", "RBM5") |
| 34 | HALLMARK_SPERMATOGENESIS | 0.564682 | 0.806689 | -0.33201 | -0.89287 | 1972 | 5 | c("CDKN3", "CHFR", "PEBP1", "SLC12A2") |
| 35 | HALLMARK_KRAS_SIGNALING_DN | 0.478414 | 0.806689 | -0.45913 | -0.97374 | 1905 | 3 | c("IDUA", "FGFR3", "LFNG") |
| 36 | HALLMARK_UNFOLDED_PROTEIN_RESPONSE | 0.582797 | 0.80944 | 0.245581 | 0.891627 | 4437 | 14 | c("ATF4", "WIPI1", "ARFGAP1", "FUS", "EIF4A2", "XBP1", "SERP1", "YWHAZ", "EDC4", "SEC31A") |
| 37 | HALLMARK_FATTY_ACID_METABOLISM | 0.600736 | 0.811805 | 0.224781 | 0.88655 | 4734 | 18 | c("FASN", "ACADVL", "ACSS1") |
| 38 | HALLMARK_PROTEIN_SECRETION | 0.668781 | 0.879975 | -0.1759 | -0.84389 | 1475 | 16 | c("ANP32E", "CTSC", "COPE", "GNAS", "ARFGAP3", "AP2M1", "SEC22B", "AP1G1", "SEC31A", "RAB5A", "CD63", "GBF1", "MON2", "ICA1", "ATP1A1", "STX16") |
| 39 | HALLMARK_MITOTIC_SPINDLE | 0.799163 | 0.880262 | 0.182095 | 0.718193 | 6298 | 18 | c("NUMA1", "RHOT2", "CTTN", "TUBGCP6", "KIF22", "FARP1", "ARFIP2", "EZR", "CYTH2", "CSNK1D", "BCAR1") |
| 40 | HALLMARK_WNT_BETA_CATENIN_SIGNALING | 0.729669 | 0.880262 | -0.36976 | -0.78421 | 2906 | 3 | MYC |
| 41 | HALLMARK_APOPTOSIS | 0.804253 | 0.880262 | 0.214618 | 0.715467 | 5899 | 11 | c("ERBB2", "RHOT2", "MADD", "BIK") |
| 42 | HALLMARK_INTERFERON_GAMMA_RESPONSE | 0.781222 | 0.880262 | 0.196881 | 0.731126 | 5998 | 15 | c("STAT2", "PSME2", "NCOA3", "TAPBP", "SPPL2A") |
| 43 | HALLMARK_PI3K_AKT_MTOR_SIGNALING | 0.801049 | 0.880262 | -0.17187 | -0.74612 | 1984 | 13 | c("VAV3", "PPP1CA", "AP2M1", "MKNK2", "SQSTM1", "UBE2D3", "ARPC3", "TSC2", "AKT1", "GRK2", "PIK3R3") |
| 44 | HALLMARK_UV_RESPONSE_UP | 0.705513 | 0.880262 | -0.2271 | -0.80768 | 2021 | 9 | c("GAL", "H2AX", "CYB5R1") |
| 45 | HALLMARK_ALLOGRAFT_REJECTION | 0.771402 | 0.880262 | -0.24001 | -0.75682 | 2459 | 7 | c("FLNA", "BCL3", "TIMP1", "IKBKB") |
| 46 | HALLMARK_PANCREAS_BETA_CELLS | 0.809841 | 0.880262 | 0.597222 | 0.796589 | 4015 | 1 | SRP14 |
| 47 | HALLMARK_COMPLEMENT | 0.864591 | 0.900616 | -0.17335 | -0.68457 | 2304 | 11 | c("CTSD", "CTSC", "ANXA5", "TIMP1", "PPP4C", "GNB2", "BRPF3") |
| 48 | HALLMARK_REACTIVE_OXYGEN_SPECIES_PATHWAY | 0.859535 | 0.900616 | 0.304048 | 0.691271 | 5439 | 4 | c("MGST1", "STK25", "LAMTOR5") |
| 49 | HALLMARK_INTERFERON_ALPHA_RESPONSE | 0.960628 | 0.960628 | 0.173819 | 0.538199 | 6855 | 9 | c("STAT2", "PSME2") |
| 50 | HALLMARK_P53_PATHWAY | 0.947674 | 0.960628 | -0.10685 | -0.62833 | 1629 | 24 | c("CTSD", "PVT1", "S100A10", "RPS12", "RXRA") |

| Cluster 1 | pathway | pval | padj | ES | NES | nMoreExtreme | size | leadingEdge |
|---|---|---|---|---|---|---|---|---|
| 1 | HALLMARK_ESTROGEN_RESPONSE_EARLY | 0.00041 | 0.010254 | -0.509535 | -3.125956 | 0 | 35 | c("PRSS23", "KLK10", "GREB1", "TFF3", "SIAH2", "TSKU", "AREG", "FAM102A", "IGF1R", "DLC1", "CXCL12", "CELSR2", "SCNN1A", "NBL1", "PDLIM3", "SEMA3B") |
| 2 | HALLMARK_ESTROGEN_RESPONSE_LATE | 0.000402 | 0.010254 | -0.573014 | -3.374598 | 0 | 32 | c("PRSS23", "KLK11", "AGR2", "KLK10", "TFF3", "SIAH2", "AREG", "FAM102A", "PDCD4", "CXCL12", "CELSR2", "SCNN1A", "NBL1", "PDLIM3", "GAL", "SEMA3B") |
| 3 | HALLMARK_MYC_TARGETS_V1 | 0.004972 | 0.042238 | 0.357706 | 1.889794 | 37 | 42 | c("RACK1", "RPL14", "RPL22", "NHP2", "RPL34", "NPM1", "RPS3", "RPL18", "RPS5", "RPLP0", "RPL6", "RPS6", "RPS2", "CNBP") |
| 4 | HALLMARK_UV_RESPONSE_DN | 0.003397 | 0.042238 | -0.644225 | -2.072939 | 11 | 8 | c("IGFBP5", "ID1", "IGF1R", "DLC1") |
| 5 | HALLMARK_COAGULATION | 0.005069 | 0.042238 | -0.576467 | -2.046802 | 16 | 10 | c("PRSS23", "FGG", "LAMP2", "GSN") |
| 6 | HALLMARK_KRAS_SIGNALING_DN | 0.002762 | 0.042238 | -0.723071 | -2.0388 | 9 | 6 | c("CALML5", "CELSR2", "LFNG", "SELENOP", "COPZ2") |
| 7 | HALLMARK_P53_PATHWAY | 0.013963 | 0.099736 | 0.393179 | 1.785662 | 101 | 26 | c("PVT1", "RACK1", "DDIT4", "RPS12", "RPL18", "IFI30", "WWP1", "DCXR", "RXRA", "RPL36", "EPS8L2") |
| 8 | HALLMARK_UNFOLDED_PROTEIN_RESPONSE | 0.039709 | 0.248182 | 0.442625 | 1.622746 | 272 | 14 | c("RPS14", "DDIT4", "NHP2", "NPM1", "SEC11A") |
| 9 | HALLMARK_ANDROGEN_RESPONSE | 0.066605 | 0.370028 | -0.400661 | -1.548814 | 215 | 12 | c("HPGD", "SELENOP", "AKT1", "TMPRSS2", "SPDEF", "UBE2I", "DBI", "PA2G4", "ZMIZ1") |
| 10 | HALLMARK_WNT_BETA_CATENIN_SIGNALING | 0.102516 | 0.465982 | -0.550514 | -1.433955 | 382 | 5 | c("DKK1", "MAML1", "JAG2", "NOTCH1", "LEF1") |
| 11 | HALLMARK_KRAS_SIGNALING_UP | 0.100107 | 0.465982 | -0.553082 | -1.440645 | 373 | 5 | c("APOD", "SEMA3B", "PRELID3B", "JUP", "RBM4") |
| 12 | HALLMARK_MITOTIC_SPINDLE | 0.221591 | 0.610999 | -0.305013 | -1.218346 | 701 | 13 | c("PREX1", "FLNA", "GSN", "ARF6", "ABR", "SHROOM1", "YWHAE", "ARFIP2", "RHOT2", "UXT", "NUMA1", "FARP1") |
| 13 | HALLMARK_TGF_BETA_SIGNALING | 0.1714 | 0.610999 | -0.627425 | -1.309296 | 693 | 3 | c("ID1", "UBE2D3") |
| 14 | HALLMARK_IL6_JAK_STAT3_SIGNALING | 0.22122 | 0.610999 | 0.890785 | 1.186871 | 1127 | 1 | CD9 |
| 15 | HALLMARK_MYOGENESIS | 0.238316 | 0.610999 | 0.296998 | 1.214543 | 1697 | 19 | c("CKB", "TNNT1", "ITGB5", "GABARAPL2", "SORBS3", "SYNGR2", "ITGB4", "SCD", "EIF4A2", "SPEG", "MB", "MYO1C", "COL6A2", "ATP6AP1") |
| 16 | HALLMARK_MTORC1_SIGNALING | 0.228524 | 0.610999 | 0.277389 | 1.228438 | 1659 | 24 | c("DDIT4", "SQSTM1", "GAPDH", "IFI30", "CD9", "SLC9A3R1", "SEC11A", "TM7SF2", "GPI") |
| 17 | HALLMARK_EPITHELIAL_MESENCHYMAL_TRANSITIO N | 0.187233 | 0.610999 | -0.344692 | -1.275279 | 612 | 11 | c("FLNA", "DKK1", "AREG", "CXCL12") |
| 18 | HALLMARK_REACTIVE_OXYGEN_SPECIES_PATHWAY | 0.2444 | 0.610999 | 0.43169 | 1.215962 | 1570 | 7 | c("FTL", "ATOX1", "PRDX2", "LAMTOR5") |
| 19 | HALLMARK_IL2_STAT5_SIGNALING | 0.183561 | 0.610999 | 0.385485 | 1.288574 | 1234 | 11 | c("PHLDA1", "MUC1", "CAPG", "COL6A1", "SYNGR2", "ST3GAL4") |
| 20 | HALLMARK_PEROXISOME | 0.204668 | 0.610999 | 0.425327 | 1.258493 | 1323 | 8 | c("CRABP2", "FIS1", "CNBP", "PRDX5") |
| 21 | HALLMARK_TNFA_SIGNALING_VIA_NFKB | 0.384777 | 0.716465 | 0.329818 | 1.062123 | 2557 | 10 | c("PHLDA1", "SQSTM1", "EIF1", "BCL3") |
| 22 | HALLMARK_HYPOXIA | 0.444208 | 0.716465 | 0.255839 | 1.00813 | 3136 | 17 | c("DDIT4", "GAPDH", "FBP1", "ALDOC", "SELENBP1", "GPI") |
| 23 | HALLMARK_APOPTOSIS | 0.435091 | 0.716465 | -0.260815 | -1.008216 | 1410 | 12 | c("LGALS3", "PDCD4", "GSN", "IFITM3") |
| 24 | HALLMARK_PROTEIN_SECRETION | 0.374833 | 0.716465 | -0.240277 | -1.047332 | 1119 | 16 | c("LAMP2", "TOM1L1", "AP3S1", "MON2", "BET1", "ARCN1", "ARF1", "RAB5A", "SEC31A", "AP2S1", "SEC22B", "COPE", "GNAS", "STX16", "CD63", "ERGIC3") |
| 25 | HALLMARK_APICAL_JUNCTION | 0.40512 | 0.716465 | 0.246586 | 1.043496 | 2911 | 21 | c("MDK", "SORBS3", "ITGB4", "RHOF", "CLDN7", "CERCAM", "GAMT", "SPEG", "HADH", "PTK2", "CLDN4", "BAIAP2", "THBS3", "JUP", "GNAI2", "PKD1", "PFN1", "MYL12B", "TIAL1") |
| 26 | HALLMARK_APICAL_SURFACE | 0.314137 | 0.716465 | 0.549582 | 1.118436 | 1870 | 3 | c("GATA3", "HSPB1") |
| 27 | HALLMARK_COMPLEMENT | 0.361637 | 0.716465 | -0.290495 | -1.074764 | 1183 | 11 | c("CTSD", "LGALS3", "LAMP2") |

| | | | | | | | | |
|---|---|---|---|---|---|---|---|---|
| 28 | HALLMARK_PI3K_AKT_MTOR_SIGNALING | 0.441258 | 0.716465 | -0.259073 | -1.001485 | 1430 | 12 | c("VAV3", "AKT1", "ACACA", "UBE2D3", "PIK3R3", "ARF1", "MAP2K3") |
| 29 | HALLMARK_ALLOGRAFT_REJECTION | 0.425888 | 0.716465 | 0.296717 | 1.023972 | 2878 | 12 | c("NPM1", "RPS3A", "RPL9", "CAPG", "BCL3") |
| 30 | HALLMARK_SPERMATOGENESIS | 0.431465 | 0.716465 | -0.482051 | -1.005933 | 1746 | 3 | c("PHKG2", "PIAS2", "PEBP1") |
| 31 | HALLMARK_PANCREAS_BETA_CELLS | 0.382304 | 0.716465 | 0.522807 | 1.063947 | 2276 | 3 | c("SEC11A", "SRP14") |
| 32 | HALLMARK_HEME_METABOLISM | 0.502147 | 0.784605 | -0.224954 | -0.95233 | 1519 | 15 | c("LAMP2", "SIDT2", "BCAM", "RBM5", "SLC11A2", "PPOX", "MAP2K3", "MGST3") |
| 33 | HALLMARK_G2M_CHECKPOINT | 0.575838 | 0.846821 | 0.33818 | 0.902201 | 3674 | 6 | c("HMGB3", "BCL3", "SNRPD1", "NUMA1") |
| 34 | HALLMARK_NOTCH_SIGNALING | 0.56105 | 0.846821 | -0.317222 | -0.894453 | 2030 | 6 | c("LFNG", "NOTCH3", "APH1A", "NOTCH1", "HES1") |
| 35 | HALLMARK_CHOLESTEROL_HOMEOSTASIS | 0.638087 | 0.911552 | 0.265356 | 0.854533 | 4241 | 10 | c("ALDOC", "CD9", "TM7SF2", "SCD", "S100A11") |
| 36 | HALLMARK_DNA_REPAIR | 0.698526 | 0.962103 | 0.197803 | 0.808895 | 4976 | 19 | c("NME4", "NME3", "EDF1", "VPS28", "ADCY6", "COX17", "POLD4", "NT5C", "DCTN4", "IMPDH2", "POLR2K", "POLR1D") |
| 37 | HALLMARK_ADIPOGENESIS | 0.923619 | 0.962103 | -0.11222 | -0.610349 | 2490 | 26 | c("PDCD4", "LPCAT3", "PRDX3", "ACADS", "UQCRQ", "SQOR", "ACLY", "AK2", "PIM3") |
| 38 | HALLMARK_INTERFERON_ALPHA_RESPONSE | 0.841383 | 0.962103 | -0.191374 | -0.679494 | 2821 | 10 | c("IFI27", "IFITM3", "MOV10", "PSME2", "BST2") |
| 39 | HALLMARK_INTERFERON_GAMMA_RESPONSE | 0.918876 | 0.962103 | -0.15126 | -0.604195 | 2910 | 13 | c("IFI27", "IFITM3", "RNF213") |
| 40 | HALLMARK_MYC_TARGETS_V2 | 0.862269 | 0.962103 | 0.251545 | 0.671076 | 5502 | 6 | c("NPM1", "CBX3", "IMP4") |
| 41 | HALLMARK_INFLAMMATORY_RESPONSE | 0.885501 | 0.962103 | 0.22735 | 0.640389 | 5691 | 7 | c("SLC7A2", "LY6E", "TIMP1", "HPN") |
| 42 | HALLMARK_XENOBIOTIC_METABOLISM | 0.84602 | 0.962103 | -0.146177 | -0.68851 | 2433 | 19 | c("AKR1C2", "HES6", "PGRMC1") |
| 43 | HALLMARK_FATTY_ACID_METABOLISM | 0.916809 | 0.962103 | -0.132312 | -0.608972 | 2677 | 18 | c("HPGD", "GLUL", "ACADS", "GRHPR", "PDHA1", "IDH3G", "ENO2", "S100A10", "ECH1", "ACAA2", "UBE2L6", "FASN", "HADH", "PSME1", "ALDOA") |
| 44 | HALLMARK_OXIDATIVE_PHOSPHORYLATION | 0.91047 | 0.962103 | 0.105129 | 0.610675 | 7260 | 57 | c("ATP5MC2", "OAT", "UQCRH", "NDUFA2", "ATP6V0E1", "TCIRG1", "UQCRB", "SLC25A6", "COX7C", "CPT1A", "GPI", "ATP5MG", "ATP6V1F", "ATP5PF", "COX6B1", "COX7A2L", "ATP5PO", "PHYH", "COX17", "CYB5A", "NDUFV2", "ATP6V1G1", "VDAC2", "RHOT2", "ATP5F1A", "ACAA2", "COX4I1", "NDUFAB1", "ATP6AP1", "ATP5MC1", "ATP5MF") |
| 45 | HALLMARK_GLYCOLYSIS | 0.897613 | 0.962103 | 0.155077 | 0.621464 | 6355 | 18 | c("DDIT4", "CLDN3", "B4GALT7", "ELF3", "ANKZF1", "ALDOA", "CYB5A", "ALDH7A1", "SPAG4", "PFKP", "AK4", "SLC16A3", "ENO2", "RARS1", "TPI1", "B4GALT2") |
| 46 | HALLMARK_UV_RESPONSE_UP | 0.744257 | 0.962103 | 0.215809 | 0.769549 | 5086 | 13 | c("SQSTM1", "DGAT1", "GRINA", "SULT1A1", "ATP6V1F", "ALDOA", "EIF2S3", "CLTB", "AP2S1", "DNAJB1", "ENO2") |
| 47 | HALLMARK_ANGIOGENESIS | 0.85799 | 0.962103 | -0.330769 | -0.690242 | 3473 | 3 | c("JAG2", "PTK2", "TIMP1") |
| 48 | HALLMARK_BILE_ACID_METABOLISM | 0.727737 | 0.962103 | 0.320428 | 0.790695 | 4559 | 5 | c("RXRA", "PRDX5", "PHYH") |
| 49 | HALLMARK_HEDGEHOG_SIGNALING | 0.947454 | 0.96679 | -0.525597 | -0.70261 | 4651 | 1 | LDB1 |
| 50 | HALLMARK_E2F_TARGETS | 0.994088 | 0.994088 | 0.15515 | 0.437019 | 6389 | 7 | c("HMGB3", "PNN", "LUC7L3", "MXD3", "PA2G4", "AK2", "PAN2") |

| Cluster 2 | pathway | pval | padj | ES | NES | nMoreExtreme | size | leadingEdge |
|---|---|---|---|---|---|---|---|---|
| 1 | HALLMARK_ESTROGEN_RESPONSE_EARLY | 0.000116 | 0.001462 | 0.491491 | 2.600226 | 0 | 62 | c("PRSS23", "KLK10", "TFF3", "GREB1", "TSKU", "BHLHE40", "NBL1", "SIAH2", "FAM102A", "IGF1R", "AREG", "CELSR2", "DLC1", "CBFA2T3", "SLC7A5", "CXCL12", "SLC2A1", "STC2", "SCNN1A", "PGR", "MYC", "XBP1", "FLNB", "PDLIM3", "B4GALT1", "FKBP5", "SCARB1", "RAB31", "ASB13", "CD44") |
| 2 | HALLMARK_ESTROGEN_RESPONSE_LATE | 0.000117 | 0.001462 | 0.523805 | 2.742312 | 0 | 60 | c("KLK11", "PRSS23", "AGR2", "KLK10", "TFF3", "NBL1", "SIAH2", "FAM102A", "AREG", "CELSR2", "SLC7A5", "COX6C", "FABP5", "CXCL12", "PDCD4", "TSPAN13", "SCNN1A", "PGR", "XBP1", "FLNB", "PDLIM3", "GAL", "ASCL1", "ID2", "FKBP5", "SCARB1", "RAB31", "CD44", "HOMER2") |
| 3 | HALLMARK_E2F_TARGETS | 0.00011 | 0.001462 | 0.390709 | 2.310975 | 0 | 104 | c("TUBB", "HMGB2", "RAD51C", "KPNA2", "NME1", "CKS1B", "MYC", "JPT1", "STMN1", "PLK1", "H2AZ1", "MYBL2", "RAN", "PAICS", "NOP56", "CHEK2", "CKS2", "TFRC", "CCNB2", "MTHFD2", "CDK4", "HMGA1", "BIRC5", "SSRP1", "AURKB", "UBE2S", "RRM2", "RPA3", "POLE4", "PTTG1", "MKI67", "PRDX4", "NOLC1", "SNRPB", "HNRNPD", "RANBP1", "H2AX", "ATAD2", "CDKN1A", "TOP2A", "TMPO", "ANP32E", "MCM7", "RAD21", "MCM4", "TP53", "CSE1L", "MCM3", "UBE2T", "PRKDC", "PCNA", "CDC20", "CTPS1", "TK1", "XRCC6", "PA2G4", "MAD2L1", "CDKN3", <br>"NASP", "CDCA3", "MCM6", "DCLRE1B", "DDX39A", "SRSF2", "ASF1B", "RACGAP1", "AURKA", "CDK1", "RNASEH2A", "SYNCRIP", "DEK", "CBX5", "TACC3", "RBBP7", "MCM2", "NCAPD2") |
| 4 | HALLMARK_GLYCOLYSIS | 0.000119 | 0.001462 | 0.456319 | 2.273575 | 0 | 49 | c("TFF3", "LDHA", "PKM", "PGK1", "STC2", "DDIT4", "HSPA5", "PGAM1", "ENO1", "STMN1", "MIF", "TPI1", "B4GALT1", "IER3", "BIK", "MET", "TXN", "FAM162A", "PFKP", "CD44", "PPIA", "PPFIA4", "ERO1A", "G6PD", "MDH2", "P4HA2") |
| 5 | HALLMARK_HYPOXIA | 0.000236 | 0.002311 | 0.426399 | 2.190148 | 1 | 55 | c("LXN", "MT2A", "BHLHE40", "SIAH2", "LDHA", "PGK1", "SLC2A1", "STC2", "PFKFB3", "DDIT4", "HSPA5", "ENO1", "MIF", "GAPDH", "GPI", "TPI1", "ANXA2", "IER3", "JUN", "FAM162A", "SCARB1", "PFKP", "TPD52", "PPFIA4", "ERO1A", "CDKN1A", "P4HA2", "DDIT3", "FOSL2") |

| | | | | | | | | |
|---|---|---|---|---|---|---|---|---|
| 6 | HALLMARK_MYC_TARGETS_V1 | 0.000538 | 0.004394 | 0.309963 | 1.897957 | 4 | 130 | c("SRM", "LDHA", "PGK1", "HSPE1", "FBL", "KPNA2", "NME1", "HSPD1", "MYC", "H2AZ1", "RAN", "NOP56", "HDGF", "SF3B3", "C1QBP", "CDK4", "PSMA7", "XPOT", "CCT5", "RPS2", "SNRPD1", "CCT3", "SNRPA1", "RPS10", "PPIA", "PRDX4", "HNRNPA2B1", "EIF3B", "CBX3", "NOLC1", "HNRNPD", "RANBP1", "MCM7", "MRPS18B", "SLC25A3", "YWHAQ", "EIF4A1", "HNRNPU", "SNRPG", "PSMB2", "MCM4", "HSP90AB1", "CCNA2", "NOP16", "ILF2", "PCNA", "CDC20", "AIMP2", "SSBP1", "CTPS1", "CCT2", "XRCC6", "KPNB1", "PA2G4", "LSM2", "SRSF3", "MAD2L1", "VDAC1", "TRIM28", "NPM1", "HPRT1", "SRSF7", "HNRNPR", "CCT4", "IMPDH2", "MCM6", "CCT7", "CAD", "TXNL4A", "BUB3", "HNRNPC", "SRSF2", "PSMD7", "KARS1", "SNRPA", "SYNCRIP", "ODC1", "DEK", "RSL1D1", "PSMA2", "IARS1", "MCM2") |
| 7 | HALLMARK_G2M_CHECKPOINT | 0.000781 | 0.005467 | 0.353571 | 2.031939 | 6 | 90 | c("MT2A", "HIF1A", "SLC7A5", "UBE2C", "NOTCH2", "KPNA2", "CKS1B", "MYC", "JPT1", "STMN1", "PLK1", "H2AZ1", "MYBL2", "NCL", "CKS2", "TPX2", "CCNB2", "CDK4", "HMGA1", "BIRC5", "HSPA8", "SLC7A1", "SNRPD1", "AURKB", "UBE2S", "PTTG1", "MKI67", "TROAP", "NOLC1", "UCK2", "HNRNPD", "H2AX", "TOP2A", "TMPO", "HNRNPU", "RAD21", "CCND1", "MCM3", "CCNA2", "H2AZ2", "CDC20", "DTYMK", "KPNB1", "MAD2L1", "CDKN3", "NUSAP1", "NASP", "GINS2", "NSD2", "MCM6", "DDX39A", "BUB3", "SRSF2", "RACGAP1", "AURKA", "CDK1", "SYNCRIP", "ODC1", "TACC3", "MCM2", "DKC1") |
| 8 | HALLMARK_UNFOLDED_PROTEIN_RESPONSE | 0.001483 | 0.008073 | 0.435069 | 2.012047 | 11 | 37 | c("SLC7A5", "STC2", "DDIT4", "EXOSC10", "HSPA5", "CKS1B", "XBP1", "NOP56", "HSP90B1", "CALR", "PDIA6", "MTHFD2", "XPOT", "LSM4", "NOLC1", "ERO1A", "H2AX", "HSPA9", "BANF1", "EIF4A1") |
| 9 | HALLMARK_UV_RESPONSE_DN | 0.001453 | 0.008073 | 0.527892 | 2.089627 | 10 | 22 | c("ID1", "IGFBP5", "BHLHE40", "IGF1R", "DLC1", "NOTCH2", "MYC", "ANXA2", "MET", "SLC7A1") |
| 10 | HALLMARK_CHOLESTEROL_HOMEOSTASIS | 0.009081 | 0.037081 | -0.433423 | -1.90617 | 24 | 16 | c("SCD", "FASN", "CLU", "CD9", "LSS", "SEMA3B", "NIBAN1", "SQLE", "TM7SF2") |
| 11 | HALLMARK_MTORC1_SIGNALING | 0.008709 | 0.037081 | 0.312798 | 1.753283 | 76 | 80 | c("IGFBP5", "BHLHE40", "LDHA", "CTSC", "PGK1", "SLC7A5", "PHGDH", "HSPE1", "SLC2A1", "DDIT4", "HSPA5", "HSPD1", "ENO1", "PLK1", "GAPDH", "XBP1", "GPI", "TPI1", "HSP90B1", "CALR", "TFRC", "MTHFD2", "CACYBP", "RRM2", "PPIA", "SHMT2", "ERO1A", "G6PD", "STIP1", "CDKN1A", "HSPA9") |
| 12 | HALLMARK_P53_PATHWAY | 0.008664 | 0.037081 | 0.412113 | 1.847132 | 68 | 33 | c("CTSD", "S100A10", "RAD51C", "DDIT4", "SLC3A2", "IER3", "JUN", "RPS27L", "FAM162A", "OSGIN1", "ZFP36L1", "CDKN1A", "HINT1", "BAX", "DDIT3", "IRAK1", "TP53", "TXNIP", "KLF4", "PCNA") |
| 13 | HALLMARK_COAGULATION | 0.026174 | 0.098655 | 0.488763 | 1.706836 | 188 | 15 | c("FGG", "PRSS23", "GSN") |
| 14 | HALLMARK_APOPTOSIS | 0.03566 | 0.124811 | 0.369066 | 1.640385 | 282 | 32 | c("LGALS3", "HMGB2", "PDCD4", "HSPB1", "GSN", "IER3", "BIK", "JUN", "CD44", "TNFRSF12A", "CDKN1A", "TOP2A", "BAX", "DDIT3", "CCND1", "ENO2", "TXNIP") |

| | | | | | | | | |
|---|---|---|---|---|---|---|---|---|
| 15 | HALLMARK_TNFA_SIGNALING_VIA_NFKB | 0.039646 | 0.127813 | 0.392878 | 1.620185 | 304 | 25 | c("BHLHE40", "AREG", "PFKFB3", "MYC", "B4GALT1", "IER3", "ID2", "JUN", "CD44", "PDLIM5", "CDKN1A", "FOSL2", "CCND1", "KLF4") |
| 16 | HALLMARK_ALLOGRAFT_REJECTION | 0.041735 | 0.127813 | 0.446939 | 1.623382 | 305 | 17 | c("HIF1A", "FLNA", "NME1", "RPL39", "TPD52", "EIF5A", "RPS19", "HLA-A") |
| 17 | HALLMARK_PROTEIN_SECRETION | 0.051486 | 0.148401 | 0.557425 | 1.563247 | 342 | 8 | c("CTSC", "BNIP3", "TPD52", "ANP32E", "KRT18", "SOD1", "PAM", "TMX1") |
| 18 | HALLMARK_MYC_TARGETS_V2 | 0.062122 | 0.16911 | 0.345039 | 1.533596 | 492 | 32 | c("SRM", "HSPE1", "HSPD1", "MYC", "PLK1", "NOP56", "CDK4", "CBX3", "NOLC1", "MRTO4", "PUS1", "MCM4", "NOP16", "AIMP2", "WDR74", "PA2G4", "NPM1") |
| 19 | HALLMARK_EPITHELIAL_MESENCHYMAL_TRANSITION | 0.072636 | 0.177958 | 0.361499 | 1.507675 | 562 | 26 | c("FLNA", "AREG", "CXCL12", "NOTCH2", "DKK1", "PPIB", "ID2", "JUN", "TPM4", "CD44", "TNFRSF12A", "MCM7", "TFPI2", "ENO2") |
| 20 | HALLMARK_INFLAMMATORY_RESPONSE | 0.06904 | 0.177958 | 0.479164 | 1.512902 | 480 | 11 | c("HIF1A", "MYC", "MET", "SLC7A1", "P2RX4", "CDKN1A", "ATP2A2") |
| 21 | HALLMARK_MYOGENESIS | 0.08694 | 0.202859 | -0.313747 | -1.477236 | 222 | 19 | c("ITGB5", "SCD", "CLU", "SORBS3", "MB", "CKB", "SPTAN1", "NQO1") |
| 22 | HALLMARK_TGF_BETA_SIGNALING | 0.102483 | 0.228258 | 0.452696 | 1.429332 | 713 | 11 | c("ID1", "ID3", "ID2", "RAB31") |
| 23 | HALLMARK_PI3K_AKT_MTOR_SIGNALING | 0.109929 | 0.234197 | 0.392163 | 1.42442 | 805 | 17 | c("VAV3", "SLC2A1", "CFL1", "HSP90B1", "CALR", "CDK4", "CDKN1A", "DDIT3", "PFN1", "CDK1", "PPP1CA", "TNFRSF1A", "PDK1", "RAC1") |
| 24 | HALLMARK_BILE_ACID_METABOLISM | 0.123408 | 0.251959 | -0.655464 | -1.374376 | 503 | 3 | c("SULT2B1", "RXRA") |
| 25 | HALLMARK_APICAL_JUNCTION | 0.135135 | 0.264865 | -0.283817 | -1.369009 | 339 | 20 | c("MDK", "EVL", "SORBS3", "PIK3R3", "JUP", "INSIG1", "RSU1", "ADAM15", "ARPC2", "TUBG1", "MYH9") |
| 26 | HALLMARK_KRAS_SIGNALING_DN | 0.164346 | 0.309729 | 0.418391 | 1.321018 | 1144 | 11 | c("CELSR2", "SPTBN2", "CALML5", "EDN2", "GP2", "LFNG", "FGFR3", "YBX2") |
| 27 | HALLMARK_ANDROGEN_RESPONSE | 0.1728 | 0.3136 | 0.306289 | 1.307928 | 1356 | 28 | c("HPGD", "B4GALT1", "MYL12A", "FKBP5", "SLC38A2", "HOMER2", "TPD52", "SMS", "STK39", "PDLIM5", "XRCC5", "CCND1", "CENPN", "AZGP1", "XRCC6", "PA2G4") |
| 28 | HALLMARK_COMPLEMENT | 0.190932 | 0.334131 | 0.384467 | 1.282798 | 1355 | 13 | c("LGALS3", "CTSD", "CTSC", "HSPA5") |
| 29 | HALLMARK_INTERFERON_GAMMA_RESPONSE | 0.205265 | 0.346826 | 0.347055 | 1.26058 | 1504 | 17 | c("MT2A", "HIF1A", "MTHFD2", "PFKP", "HLA-A", "CDKN1A", "PSMB2", "TXNIP") |
| 30 | HALLMARK_MITOTIC_SPINDLE | 0.282014 | 0.445764 | 0.246749 | 1.16725 | 2307 | 40 | c("FLNA", "NET1", "PREX1", "NOTCH2", "PLK1", "FLNB", "GSN", "TPX2", "CCNB2", "BIRC5") |
| 31 | HALLMARK_WNT_BETA_CATENIN_SIGNALING | 0.28158 | 0.445764 | 0.493105 | 1.16913 | 1760 | 5 | c("MYC", "DKK1", "TP53") |
| 32 | HALLMARK_FATTY_ACID_METABOLISM | 0.296696 | 0.454316 | 0.268573 | 1.159113 | 2334 | 29 | c("HPGD", "LDHA", "S100A10", "MIF", "H2AZ1", "GLUL", "SMS", "YWHAH", "MDH2", "CBR1", "ENO2", "HSP90AA1", "ALDOA") |
| 33 | HALLMARK_ANGIOGENESIS | 0.339588 | 0.489407 | -0.827511 | -1.105635 | 1682 | 1 | VEGFA |
| 34 | HALLMARK_KRAS_SIGNALING_UP | 0.33591 | 0.489407 | 0.367217 | 1.116076 | 2302 | 10 | c("TSPAN13", "ID2", "GADD45G", "ERO1A", "LCP1", "KLF4") |
| 35 | HALLMARK_PEROXISOME | 0.380165 | 0.532231 | -0.295265 | -1.049368 | 1195 | 10 | c("CRABP2", "SULT2B1") |
| 36 | HALLMARK_INTERFERON_ALPHA_RESPONSE | 0.45617 | 0.620898 | -0.34899 | -0.991115 | 1633 | 6 | c("IFI30", "UBE2L6", "PSMA3", "B2M") |

| | | | | | | | | |
|---|---|---|---|---|---|---|---|---|
| 37 | HALLMARK_OXIDATIVE_PHOSPHORYLATION | 0.469338 | 0.621556 | 0.190085 | 1.005642 | 4040 | 62 | c("LDHA", "COX6C", "GPI", "SLC25A5", "CYCS", "MRPL11", "TIMM10", "COX6A1", "COX8A", "MDH2", "HSPA9", "BAX", "SLC25A3", "NDUFB4", "COX5B", "ETFA", "ATP5ME", "VDAC1", "TIMM13", "NDUFA1", "NDUFB8", "ATP5F1B", "MRPS12", "CS", "ISCU", "ETFB", "NDUFAB1", "ATP5F1C", "PHB2", "NDUFB6", "NDUFC1", "PRDX3", "COX7B", "ATP5MF", "UQCR11", "ATP5F1E", "ECH1", "ATP5PB", "SDHA", "ATP5PO", "UQCRFS1", "MGST3", "TIMM50", "COX5A", "MRPL15", "GRPEL1", "UQCRQ", "NDUFA4", "CYC1", "LRPPRC", "TOMM22", "TIMM8B") |
| 38 | HALLMARK_HEDGEHOG_SIGNALING | 0.485553 | 0.626108 | -0.46137 | -0.9674 | 1982 | 3 | c("VEGFA", "MYH9", "NRCAM") |
| 39 | HALLMARK_IL6_JAK_STAT3_SIGNALING | 0.526791 | 0.661866 | 0.371896 | 0.942888 | 3381 | 6 | c("JUN", "CD44", "TNFRSF12A") |
| 40 | HALLMARK_APICAL_SURFACE | 0.568176 | 0.67904 | -0.379955 | -0.905937 | 2220 | 4 | GATA3 |
| 41 | HALLMARK_REACTIVE_OXYGEN_SPECIES_PATHWAY | 0.563984 | 0.67904 | 0.266822 | 0.913387 | 4036 | 14 | c("TXN", "PFKP", "PRDX4", "G6PD", "FTL", "NDUFB4", "SOD1", "PDLIM1", "GSR", "JUNB", "PRDX1") |
| 42 | HALLMARK_XENOBIOTIC_METABOLISM | 0.680061 | 0.793404 | 0.187748 | 0.819718 | 5381 | 30 | c("MT2A", "AKR1C2", "DDT", "PGD", "ID2", "MTHFD1", "SHMT2", "PDLIM5", "CBR1", "ATP2A2", "HPRT1", "KARS1", "UGDH", "HES6", "AHCY", "TNFRSF1A", "ARPP19", "LONP1", "PTGES3", "PGRMC1", "GSR") |
| 43 | HALLMARK_SPERMATOGENESIS | 0.777811 | 0.886343 | 0.252046 | 0.73578 | 5236 | 9 | c("CCNB2", "CDKN3", "AURKA", "YBX2", "CDK1", "PSMG1") |
| 44 | HALLMARK_IL2_STAT5_SIGNALING | 0.834331 | 0.929141 | 0.158847 | 0.678316 | 6551 | 28 | c("BHLHE40", "IGF1R", "MYC", "XBP1", "CD44", "P2RX4", "UCK2", "PUS1") |
| 45 | HALLMARK_DNA_REPAIR | 0.912871 | 0.96396 | -0.105536 | -0.630905 | 1843 | 34 | c("BCAM", "NME3", "POLD4", "ADCY6", "COX17", "POLR2K", "ALYREF", "GPX4", "POLR2E", "POLR2I", "RFC4", "APRT", "SAC3D1") |
| 46 | HALLMARK_NOTCH_SIGNALING | 0.899844 | 0.96396 | 0.245389 | 0.622148 | 5776 | 6 | c("NOTCH2", "CCND1", "LFNG") |
| 47 | HALLMARK_UV_RESPONSE_UP | 0.924615 | 0.96396 | 0.134463 | 0.574189 | 7260 | 28 | c("PDLIM3", "GAL", "TFRC", "STIP1", "H2AX", "HNRNPU", "PPIF", "ENO2", "SLC6A8", "ALDOA", "PDAP1", "BTG1", "AMD1", "FEN1", "CEBPG", "POLR2H", "JUNB", "GGH", "FURIN", "GRPEL1", "PSMC3", "DNAJA1", "MRPL23", "EIF5") |
| 48 | HALLMARK_ADIPOGENESIS | 0.98113 | 0.996551 | 0.095688 | 0.458381 | 8058 | 42 | c("PDCD4", "PFKFB3", "DDT", "VEGFB", "SCARB1", "MTCH2", "COX6A1", "COX8A", "MDH2", "ALDOA", "YWHAG", "SOD1", "RTN3", "UBC", "CD151", "CS", "ETFB", "DNAJC15", "NDUFAB1", "PRDX3", "COX7B", "UQCR11", "ECH1", "ATP5PO", "MGST3", "MRPL15", "GRPEL1", "SLC1A5", "UQCRQ", "CYC1", "GPAT4") |
| 49 | HALLMARK_HEME_METABOLISM | 0.996551 | 0.996551 | 0.101281 | 0.394689 | 7511 | 21 | c("SLC2A1", "HDGF", "TFRC", "P4HA2", "SLC6A8", "GLRX5", "ABCB6", "TOP1", "BNIP3L", "NFE2L1", "UBAC1", "SEC14L1", "BLVRA", "EIF2AK1", "MGST3") |

| Cluster 3 | pathway | pval | padj | ES | NES | nMoreExtreme | size | leadingEdge |
|---|---|---|---|---|---|---|---|---|
| 1 | HALLMARK_HYPOXIA | 0.001091 | 0.04902 | -0.379819 | -2.361745 | 0 | 25 | c("MT2A", "SIAH2", "IER3", "BHLHE40", "LXN", "STC2", "LDHA", "CA12", "PGK1", "SLC2A1", "HEXA") |
| 2 | HALLMARK_ESTROGEN_RESPONSE_EARLY | 0.001961 | 0.04902 | -0.405254 | -3.079086 | 0 | 36 | c("PRSS23", "SIAH2", "SLC7A5", "TSKU", "SLC7A2", "BHLHE40", "MYC", "GFRA1", "CXCL12", "STC2", "CBFA2T3", "AREG", "CA12", "DLC1", "SLC2A1") |
| 3 | HALLMARK_ESTROGEN_RESPONSE_LATE | 0.006818 | 0.097201 | -0.272804 | -2.22459 | 2 | 40 | c("PRSS23", "AGR2", "SIAH2", "GAL", "SLC7A5", "FABP5", "CXCL12", "COX6C", "AREG", "CA12") |
| 4 | HALLMARK_E2F_TARGETS | 0.007776 | 0.097201 | -0.374273 | -2.019084 | 9 | 19 | c("HMGB2", "MYC", "MYBL2", "TUBB", "ANP32E", "NME1", "HMGA1", "PAICS", "AURKB") |
| 5 | HALLMARK_APOPTOSIS | 0.026087 | 0.26087 | 0.436856 | 1.668592 | 236 | 25 | c("CLU", "MGMT", "LGALS3", "CCNA1", "BIK", "SAT1", "BCL2L1", "CDKN1A", "BTG3", "APP", "ERBB2", "FDXR", "IFITM3", "DNAJA1", "TIMP1") |
| 6 | HALLMARK_CHOLESTEROL_HOMEOSTASIS | 0.041891 | 0.330654 | 0.495525 | 1.606342 | 349 | 14 | c("CLU", "SQLE", "LGALS3", "ALCAM", "NIBAN1", "MAL2", "PCYT2", "TM7SF2", "TP53INP1") |
| 7 | HALLMARK_MYC_TARGETS_V1 | 0.052905 | 0.330654 | -0.251058 | -1.531482 | 50 | 24 | c("MYC", "LDHA", "FBL", "NME1", "PGK1", "PSMD8", "RPL6", "TCP1", "TUFM", "RPL18", "PPM1G", "COX5A") |
| 8 | HALLMARK_KRAS_SIGNALING_DN | 0.048426 | 0.330654 | 0.704382 | 1.51235 | 322 | 4 | c("CALML5", "CCNA1", "LFNG") |
| 9 | HALLMARK_G2M_CHECKPOINT | 0.10578 | 0.510287 | -0.222278 | -1.382144 | 96 | 25 | c("MT2A", "SLC7A5", "MYC", "MYBL2", "HMGA1", "AURKB") |
| 10 | HALLMARK_PROTEIN_SECRETION | 0.138045 | 0.510287 | -0.275939 | -1.310607 | 208 | 15 | c("ANP32E", "ARFIP1", "NAPA", "ICA1", "TMED10", "AP2M1", "GNAS", "CLTC", "GBF1", "COPB2", "LAMP2", "RAB5A") |
| 11 | HALLMARK_APICAL_JUNCTION | 0.153086 | 0.510287 | 0.394654 | 1.354224 | 1316 | 17 | c("EVL", "MDK", "PIK3R3", "MYL12B", "CLDN4", "ACTN1", "INSIG1", "CDH1", "CLDN7", "TUBG1") |
| 12 | HALLMARK_APICAL_SURFACE | 0.144865 | 0.510287 | 0.766805 | 1.298871 | 844 | 2 | c("APP", "SULF2") |
| 13 | HALLMARK_UNFOLDED_PROTEIN_RESPONSE | 0.104978 | 0.510287 | -0.326019 | -1.394396 | 193 | 12 | c("SLC7A5", "STC2", "CNOT2", "EDC4", "BANF1", "DCP2", "ATP6V0D1") |
| 14 | HALLMARK_COAGULATION | 0.123482 | 0.510287 | -0.272114 | -1.350964 | 182 | 16 | c("PRSS23", "FGG") |
| 15 | HALLMARK_KRAS_SIGNALING_UP | 0.137609 | 0.510287 | 0.480153 | 1.365882 | 1074 | 9 | c("PCSK1N", "SLPI", "TSPAN1") |
| 16 | HALLMARK_NOTCH_SIGNALING | 0.242112 | 0.682504 | 0.467589 | 1.222397 | 1810 | 7 | c("PSENEN", "LFNG", "HES1", "DTX2", "SKP1", "ARRB1") |
| 17 | HALLMARK_HEDGEHOG_SIGNALING | 0.218722 | 0.682504 | -0.494342 | -1.214301 | 728 | 4 | c("TLE3", "ADGRG1", "CELSR1", "TLE1") |
| 18 | HALLMARK_COMPLEMENT | 0.245701 | 0.682504 | 0.338595 | 1.228084 | 2171 | 21 | c("CLU", "LGALS3", "CALM1", "CALM3", "HSPA1A", "S100A13") |
| 19 | HALLMARK_IL6_JAK_STAT3_SIGNALING | 0.275851 | 0.725923 | -0.416567 | -1.151108 | 834 | 5 | c("STAM2", "CD9", "IRF9", "STAT1", "IL13RA1") |
| 20 | HALLMARK_ADIPOGENESIS | 0.294712 | 0.736781 | -0.206121 | -1.111957 | 378 | 19 | c("DRAM2", "UQCR11", "ABCB8", "ARL4A", "UQCR10", "LPCAT3", "CCNG2", "DGAT1", "CD151", "REEP5", "REEP6", "DHRS7B", "SCP2", "POR", "UBC", "APLP2", "BCL6", "DECR1") |
| 21 | HALLMARK_INFLAMMATORY_RESPONSE | 0.321797 | 0.766184 | -0.233459 | -1.083377 | 529 | 14 | c("SLC7A2", "MYC", "PSEN1", "SELENOS", "NFKBIA", "TNFSF10", "BST2", "C5AR1") |
| 22 | HALLMARK_SPERMATOGENESIS | 0.368523 | 0.837552 | 0.305602 | 1.094873 | 3242 | 20 | c("PCSK1N", "NPHP1", "MLF1", "CCNA1", "SLC12A2", "STRBP", "IFT88") |
| 23 | HALLMARK_ANDROGEN_RESPONSE | 0.4048 | 0.880001 | 0.304388 | 1.058022 | 3507 | 18 | c("SAT1", "TMPRSS2", "ABHD2", "ACTN1", "DBI", "B2M", "DHCR24", "INSIG1") |
| 24 | HALLMARK_TNFA_SIGNALING_VIA_NFKB | 0.804824 | 0.933199 | 0.18631 | 0.702668 | 7273 | 24 | c("SAT1", "MARCKS", "ACKR3", "CDKN1A", "BTG3", "HES1", "PPP1R15A", "EFNA1", "BCL6", "LITAF") |
| 25 | HALLMARK_MITOTIC_SPINDLE | 0.671937 | 0.933199 | -0.125363 | -0.837758 | 509 | 29 | c("FLNA", "RAB3GAP1", "NEK2", "ARHGAP27", "MYH10", "GSN", "CDC42EP4", "KIF3B", "SMC4", "MYO1E", "TBCD", "CDK1", "NET1") |

| | | | | | | | | |
|---|---|---|---|---|---|---|---|---|
| 26 | HALLMARK_WNT_BETA_CATENIN_SIGNALING | 0.662952 | 0.933199 | 0.512682 | 0.868418 | 3866 | 2 | DKK1 |
| 27 | HALLMARK_TGF_BETA_SIGNALING | 0.591125 | 0.933199 | -0.272568 | -0.884863 | 1491 | 7 | c("ID1", "SMURF1", "KLF10") |
| 28 | HALLMARK_DNA_REPAIR | 0.499888 | 0.933199 | 0.265397 | 0.976878 | 4461 | 22 | c("CCNO", "CETN2", "ERCC1") |
| 29 | HALLMARK_MYOGENESIS | 0.745298 | 0.933199 | 0.201685 | 0.760657 | 6735 | 24 | c("CLU", "MB", "ATP6AP1", "CDKN1A", "PLXNB2", "AGRN", "APP") |
| 30 | HALLMARK_INTERFERON_ALPHA_RESPONSE | 0.650471 | 0.933199 | 0.245794 | 0.84342 | 5595 | 17 | c("LGALS3BP", "IFI27", "HLA-C", "B2M", "IFITM3", "ISG15", "IFITM1", "LY6E", "CD47", "IRF9", "PSME2", "ELF1", "OAS1") |
| 31 | HALLMARK_INTERFERON_GAMMA_RESPONSE | 0.839879 | 0.933199 | 0.167148 | 0.66228 | 7762 | 29 | c("LGALS3BP", "HLA-B", "IFI27", "VAMP8", "CDKN1A", "B2M", "SRI", "SPPL2A", "IFITM3", "ISG15", "TAPBP") |
| 32 | HALLMARK_MTORC1_SIGNALING | 0.519193 | 0.933199 | 0.249596 | 0.962423 | 4733 | 26 | c("SQLE", "PIK3R3", "CDKN1A", "NIBAN1", "SYTL2", "DHCR24", "INSIG1", "TM7SF2", "PPP1R15A", "FDXR", "SORD", "TUBG1") |
| 33 | HALLMARK_MYC_TARGETS_V2 | 0.682193 | 0.933199 | 0.430923 | 0.838152 | 4305 | 3 | c("PLK4", "SORD") |
| 34 | HALLMARK_OXIDATIVE_PHOSPHORYLATION | 0.588598 | 0.933199 | -0.125639 | -0.903854 | 381 | 32 | c("LDHA", "COX6C", "ATP6V0E1", "COX7C", "CYB5A", "UQCR11", "COX5A", "UQCR10", "COX6B1", "OAT", "HADHB", "ATP5F1E", "ATP6V0C", "ATP5MG", "NDUFB1", "ACAA1", "NDUFA3", "SURF1", "NDUFA1", "ATP6V1D", "UQCRB", "SLC25A4") |
| 35 | HALLMARK_GLYCOLYSIS | 0.812217 | 0.933199 | 0.191672 | 0.695195 | 7179 | 21 | c("TFF3", "BIK", "SDC1", "AGRN", "CLDN3") |
| 36 | HALLMARK_REACTIVE_OXYGEN_SPECIES_PATHWAY | 0.800802 | 0.933199 | 0.27512 | 0.719235 | 5989 | 7 | c("CDKN2D", "MGST1", "IPCEF1") |
| 37 | HALLMARK_P53_PATHWAY | 0.612621 | 0.933199 | 0.213312 | 0.887917 | 5814 | 36 | c("H1-2", "SAT1", "H2AJ", "SDC1", "VAMP8", "PERP", "CDKN1A", "PLXNB2", "CCP110", "APP", "MXD4", "PPP1R15A", "FDXR") |
| 38 | HALLMARK_UV_RESPONSE_UP | 0.725682 | 0.933199 | 0.216975 | 0.777352 | 6385 | 20 | c("PPP1R2", "SELENOW", "CYB5R1", "BTG3", "AQP3", "BSG", "DNAJA1", "PSMC3") |
| 39 | HALLMARK_UV_RESPONSE_DN | 0.823885 | 0.933199 | 0.186104 | 0.685017 | 7353 | 22 | c("MGMT", "PIK3R3", "SRI", "INSIG1", "ERBB2", "ANXA2", "RUNX1", "NFIB", "MIOS", "PIAS3", "APBB2") |
| 40 | HALLMARK_ANGIOGENESIS | 0.757871 | 0.933199 | 0.359142 | 0.7711 | 5054 | 4 | c("APP", "TIMP1", "ITGAV", "LRPAP1") |
| 41 | HALLMARK_HEME_METABOLISM | 0.455882 | 0.933199 | -0.154496 | -0.98086 | 402 | 26 | c("SLC2A1", "RANBP10", "CAST", "PIGQ", "CIR1", "CTSB", "GAPVD1", "PICALM", "BLVRA", "VEZF1", "PSMD9", "LRP10", "BCAM", "IGSF3", "LAMP2", "PPOX", "GDE1", "HAGH", "RBM5", "ADIPOR1", "BSG", "AQP3", "UCP2", "SELENBP1", "NUDT4", "YPEL5") |
| 42 | HALLMARK_BILE_ACID_METABOLISM | 0.732074 | 0.933199 | 0.254994 | 0.768835 | 5890 | 11 | c("EFHC1", "IDH2", "DHCR24", "FDXR", "ABCA2", "SULT2B1", "SCP2") |
| 43 | HALLMARK_PEROXISOME | 0.740091 | 0.933199 | 0.221932 | 0.76154 | 6366 | 17 | c("ERCC1", "IDH2", "DHCR24", "DHRS3", "FABP6", "MVP", "SULT2B1", "FIS1", "PABPC1", "PEX2", "SCP2", "SLC25A4") |
| 44 | HALLMARK_ALLOGRAFT_REJECTION | 0.482143 | 0.933199 | -0.224417 | -0.959839 | 890 | 12 | c("FLNA", "EIF5A", "NME1", "PTPN6", "HLA-A", "INHBB", "CD47", "STAT1", "HLA-E", "TAPBP", "TIMP1", "B2M") |
| 45 | HALLMARK_PANCREAS_BETA_CELLS | 0.662509 | 0.933199 | -0.483046 | -0.847196 | 2761 | 2 | c("SRP14", "SPCS1") |
| 46 | HALLMARK_EPITHELIAL_MESENCHYMAL_TRANSITIO N | 0.930302 | 0.962438 | 0.158932 | 0.552431 | 8061 | 18 | c("DKK1", "SAT1", "SDC1", "BASP1") |
| 47 | HALLMARK_XENOBIOTIC_METABOLISM | 0.943189 | 0.962438 | 0.13687 | 0.527758 | 8599 | 26 | c("HES6", "PTGES3", "AKR1C2", "SPINT2", "ESR1") |
| 48 | HALLMARK_FATTY_ACID_METABOLISM | 0.909185 | 0.962438 | 0.167758 | 0.583108 | 7878 | 18 | c("HSP90AA1", "ACADVL", "HSPH1", "DHCR24", "DECR1", "ACSS1") |
| 49 | HALLMARK_IL2_STAT5_SIGNALING | 0.930568 | 0.962438 | 0.152539 | 0.546499 | 8188 | 20 | c("MUC1", "ALCAM", "BCL2L1", "DRC1") |
| 50 | HALLMARK_PI3K_AKT_MTOR_SIGNALING | 0.999387 | 0.999387 | 0.112597 | 0.348619 | 8148 | 12 | c("PIK3R3", "CDKN1A", "ARPC3", "MAPK10", "CAB39", "CLTC", "PIN1", "AP2M1", "TSC2") |

| Cluster 3 | pathway | pval | padj | ES | NES | nMoreExtreme | size | leadingEdge |
|---|---|---|---|---|---|---|---|---|
| 1 | HALLMARK_G2M_CHECKPOINT | 0.000219 | 0.002244 | 0.402487 | 2.344857 | 0 | 59 | c("MT2A", "MYC", "SLC7A5", "TNPO2", "UBE2C", "CCND1", "DDX39A", "STMN1", "AURKB", "CKS1B", "PTTG1", "TROAP", "PLK1", "TOP2A", "SRSF10", "HMGA1", "MYBL2", "H2AX", "ILF3", "CKS2", "HSPA8", "UBE2S", "NOLC1", "CDK4", "TPX2", "MKI67", "GINS2", "ODC1", "MAD2L1", "AURKA", "KPNA2", "UCK2", "PRC1", "CCNA2", "MCM2", "TRA2B", "BUB3", "CCNB2", "NUSAP1", "JPT1") |
| 2 | HALLMARK_ESTROGEN_RESPONSE_EARLY | 0.000229 | 0.002244 | 0.41016 | 2.540657 | 0 | 75 | c("PRSS23", "SIAH2", "GREB1", "CXCL12", "AREG", "CA12", "CELSR2", "MYC", "IGF1R", "DLC1", "BHLHE40", "HSPB8", "PDZK1", "TSKU", "KRT18", "SLC7A5", "TPBG", "GLA", "SLC2A1", "SCARB1", "PDLIM3", "STC2", "BCL2", "NPY1R", "CBFA2T3", "CCND1", "ELOVL5", "MYBL1", "FLNB", "FKBP5", "FAM102A", "RBBP8", "GJA1", "SLC7A2", "KCNK15", "TPD52L1", "NRIP1", "CCN5", "KRT8", "AMFR", "THSD4", "FDFT1", "MREG", "TGIF2") |
| 3 | HALLMARK_ESTROGEN_RESPONSE_LATE | 0.000222 | 0.002244 | 0.434185 | 2.598454 | 0 | 65 | c("PRSS23", "AGR2", "SIAH2", "CXCL12", "AREG", "PDCD4", "CA12", "CELSR2", "IGSF1", "HSPB8", "PDZK1", "GAL", "SLC7A5", "TPBG", "GLA", "SCARB1", "PDLIM3", "BCL2", "NPY1R", "CCND1", "SERPINA3", "RNASEH2A", "ELOVL5", "COX6C", "FLNB", "FKBP5", "FAM102A", "TOP2A", "HPRT1", "RBBP8", "TPD52L1", "NRIP1", "CCN5") |
| 4 | HALLMARK_E2F_TARGETS | 0.000219 | 0.002244 | 0.502804 | 2.890636 | 0 | 56 | c("MYC", "RAD51C", "HMGB2", "ANP32E", "RNASEH2A", "DDX39A", "STMN1", "UBE2T", "TUBB", "NOP56", "AURKB", "CKS1B", "PTTG1", "PLK1", "TOP2A", "HMGA1", "PAICS", "MYBL2", "NME1", "H2AX", "ILF3", "CKS2", "CDKN1A", "UBE2S", "TFRC", "CDCA3", "DCTPP1", "NOLC1", "DNMT1", "CDK4", "MTHFD2", "MKI67", "MAD2L1", "AURKA", "MCM7", "RRM2", "KPNA2", "RAN", "PRDX4", "MCM2", "TRA2B", "CCNB2", "CHEK2", "JPT1", "TMPO", "NCAPD2", "RAD21", "POLD2", "CTPS1", "BIRC5", "ESPL1") |
| 5 | HALLMARK_MYC_TARGETS_V1 | 0.000221 | 0.002244 | 0.457967 | 2.573158 | 0 | 52 | c("MYC", "FBL", "SRM", "NOP56", "CAD", "XPOT", "HPRT1", "C1QBP", "LDHA", "NME1", "TYMS", "HDGF", "PGK1", "YWHAQ", "NOLC1", "CDK4", "ODC1", "MAD2L1", "IARS1", "LSM7", "DDX21", "SRSF7", "NOP16", "MCM7", "KPNA2", "IMPDH2", "HSPE1", "RAN", "CCNA2", "RPS5", "PRDX4", "MCM2", "TRA2B", "BUB3", "SET", "CCT2", "HNRNPA3", "SSBP1", "SF3B3", "HSPD1") |
| 6 | HALLMARK_INTERFERON_ALPHA_RESPONSE | 0.000613 | 0.004384 | 0.606665 | 2.206905 | 2 | 14 | c("IFI27", "IFITM1", "BST2", "ISG15", "IFITM3", "TXNIP") |
| 7 | HALLMARK_UNFOLDED_PROTEIN_RESPONSE | 0.000626 | 0.004384 | 0.538088 | 2.314083 | 2 | 22 | c("HSPA5", "SLC7A5", "STC2", "HYOU1", "NOP56", "CALR", "CKS1B", "HSP90B1", "XPOT", "PDIA6", "H2AX", "IFIT1", "KHSRP", "NOLC1", "MTHFD2", "IARS1") |
| 8 | HALLMARK_HYPOXIA | 0.00128 | 0.007841 | 0.419156 | 2.145979 | 5 | 37 | c("SIAH2", "MT2A", "LXN", "CA12", "BHLHE40", "IER3", "HSPA5", "TPBG", "SLC2A1", "ALDOA", "SCARB1", "STC2", "BCL2") |
| 9 | HALLMARK_CHOLESTEROL_HOMEOSTASIS | 0.002155 | 0.011734 | -0.601271 | -2.090555 | 10 | 13 | c("SCD", "CD9", "FASN", "CLU", "TM7SF2", "LSS", "TP53INP1", "SQLE") |
| 10 | HALLMARK_INTERFERON_GAMMA_RESPONSE | 0.002505 | 0.012276 | 0.459279 | 1.975158 | 11 | 22 | c("IFI27", "MT2A", "BST2", "ISG15", "IFITM3", "TXNIP") |

| | | | | | | | |
|---|---|---|---|---|---|---|---|
| 11 | HALLMARK_INFLAMMATORY_RESPONSE | 0.006942 | 0.028345 | 0.541408 | 1.916337 | 33 | 13 | c("IFITM1", "BST2", "MYC", "TPBG", "P2RX4", "EDN1", "SLC7A2", "CDKN1A") |
| 12 | HALLMARK_UV_RESPONSE_DN | 0.006776 | 0.028345 | 0.482114 | 1.927729 | 32 | 18 | c("ID1", "IGFBP5", "MYC", "IGF1R", "DLC1", "BHLHE40") |
| 13 | HALLMARK_MYC_TARGETS_V2 | 0.010496 | 0.039562 | 0.483899 | 1.848905 | 50 | 16 | c("MYC", "SRM", "FARSA", "NOP56", "PLK1", "DCTPP1", "NOLC1", "CDK4", "NOP16", "HSPE1", "PUS1", "HSPD1", "RCL1") |
| 14 | HALLMARK_APOPTOSIS | 0.065409 | 0.219343 | 0.337768 | 1.513875 | 311 | 25 | c("PDCD4", "IER3", "IFITM3", "TXNIP", "KRT18", "HMGB2", "CCND1", "BAX") |
| 15 | HALLMARK_FATTY_ACID_METABOLISM | 0.067146 | 0.219343 | 0.381844 | 1.526801 | 326 | 18 | c("HPGD", "S100A10", "ALDOA", "ELOVL5", "LDHA", "OSTC", "LGALS1", "GLUL", "CBR1", "MIF", "ODC1") |
| 16 | HALLMARK_MITOTIC_SPINDLE | 0.121941 | 0.324279 | 0.333294 | 1.378875 | 582 | 20 | c("FLNA", "PALLD", "FLNB", "PLK1", "TOP2A", "TPX2", "AURKA", "PRC1", "CCNB2", "NUSAP1", "DST", "ARHGDIA", "PIF1", "BIRC5", "ESPL1") |
| 17 | HALLMARK_ANDROGEN_RESPONSE | 0.132359 | 0.324279 | -0.337529 | -1.374553 | 687 | 21 | c("SCD", "SAT1", "TMPRSS2", "SPDEF", "INSIG1", "ACSL3", "DBI") |
| 18 | HALLMARK_MYOGENESIS | 0.131236 | 0.324279 | -0.33283 | -1.377125 | 683 | 22 | c("SCD", "CLU", "SPDEF", "SYNGR2", "MB", "ITGB5", "ATP6AP1", "MYO1C", "PLXNB2", "ERBB3") |
| 19 | HALLMARK_APICAL_JUNCTION | 0.111263 | 0.324279 | -0.370191 | -1.428195 | 570 | 18 | c("MDK", "EVL", "CLDN4", "INSIG1", "PIK3R3") |
| 20 | HALLMARK_APICAL_SURFACE | 0.126836 | 0.324279 | -0.683015 | -1.359341 | 638 | 3 | c("GATA3", "SULF2", "LYPD3") |
| 21 | HALLMARK_ADIPOGENESIS | 0.147228 | 0.327917 | 0.334047 | 1.335685 | 716 | 18 | c("PDCD4", "HSPB8", "ALDOA", "SCARB1", "PFKFB3", "DDT") |
| 22 | HALLMARK_MTORC1_SIGNALING | 0.142294 | 0.327917 | 0.224977 | 1.293401 | 649 | 56 | c("IGFBP5", "BHLHE40", "HSPA5", "SLC7A5", "GLA", "PHGDH", "SLC2A1", "ALDOA", "ELOVL5", "DDX39A", "CALR", "HSP90B1", "PLK1", "HPRT1", "LDHA", "PSMG1", "PPA1", "CDKN1A", "TFRC", "ATP5MC1", "PGK1", "MTHFD2", "AURKA", "RRM2", "SDF2L1", "CACYBP", "HSPE1", "WARS1", "BTG2", "MCM2", "DDIT3", "ATP2A2", "HSPD1", "G6PD") |
| 23 | HALLMARK_IL6_JAK_STAT3_SIGNALING | 0.181963 | 0.346775 | -0.73992 | -1.25715 | 911 | 2 | CD9 |
| 24 | HALLMARK_PROTEIN_SECRETION | 0.184003 | 0.346775 | 0.417207 | 1.286016 | 910 | 9 | c("KRT18", "GLA", "ANP32E", "TPD52", "SOD1", "DST", "BNIP3") |
| 25 | HALLMARK_P53_PATHWAY | 0.181139 | 0.346775 | 0.247313 | 1.266184 | 848 | 37 | c("CTSD", "S100A10", "IER3", "RAD51C", "TXNIP", "RPL36", "BAX", "RPS27L") |
| 26 | HALLMARK_BILE_ACID_METABOLISM | 0.164673 | 0.346775 | -0.508741 | -1.325929 | 825 | 6 | c("SULT2B1", "DHCR24", "IDH2", "EFHC1", "RXRA") |
| 27 | HALLMARK_EPITHELIAL_MESENCHYMAL_TRANSITION | 0.198786 | 0.36076 | 0.269874 | 1.246795 | 949 | 27 | c("FLNA", "CXCL12", "AREG", "TPM4", "MGP") |
| 28 | HALLMARK_WNT_BETA_CATENIN_SIGNALING | 0.213537 | 0.373691 | 0.621253 | 1.238067 | 1059 | 3 | c("MYC", "JAG2", "KAT2A") |
| 29 | HALLMARK_DNA_REPAIR | 0.234906 | 0.39691 | -0.310369 | -1.220941 | 1209 | 19 | c("CCNO", "NME3", "NME4", "POLD4", "CANT1", "EDF1") |
| 30 | HALLMARK_ALLOGRAFT_REJECTION | 0.295122 | 0.482033 | 0.299376 | 1.14387 | 1433 | 16 | c("FLNA", "RPS19", "NME1", "B2M", "CD47", "TPD52", "RPL39", "WARS1") |
| 31 | HALLMARK_TNFA_SIGNALING_VIA_NFKB | 0.427424 | 0.551153 | 0.218329 | 1.01845 | 2022 | 28 | c("AREG", "MYC", "BHLHE40", "IER3", "CCND1") |
| 32 | HALLMARK_TGF_BETA_SIGNALING | 0.417239 | 0.551153 | 0.351713 | 1.034723 | 2066 | 8 | c("ID1", "ID3", "SMAD6", "JUNB", "ID2") |
| 33 | HALLMARK_PI3K_AKT_MTOR_SIGNALING | 0.398281 | 0.551153 | 0.305347 | 1.04985 | 1945 | 12 | c("SLC2A1", "VAV3", "CALR", "HSP90B1", "CDKN1A", "CDK4") |
| 34 | HALLMARK_GLYCOLYSIS | 0.417191 | 0.551153 | 0.226045 | 1.027966 | 1989 | 26 | c("IER3", "HSPA5", "TPBG", "ALDOA", "STC2", "STMN1", "LDHA", "PPFIA4", "PGK1", "MIF", "AURKA", "ENO2", "SOD1", "FAM162A", "G6PD", "IRS2", "CENPA") |

| | | | | | | | | |
|---|---|---|---|---|---|---|---|---|
| 35 | HALLMARK_REACTIVE_OXYGEN_SPECIES_PATHWAY | 0.367401 | 0.551153 | 0.346626 | 1.068453 | 1818 | 9 | c("PRDX2", "GPX4", "JUNB", "PRDX4", "SOD1", "G6PD") |
| 36 | HALLMARK_HEME_METABOLISM | 0.382267 | 0.551153 | -0.285153 | -1.061497 | 1965 | 16 | c("SELENBP1", "AQP3") |
| 37 | HALLMARK_COAGULATION | 0.370469 | 0.551153 | 0.335146 | 1.073328 | 1808 | 10 | PRSS23 |
| 38 | HALLMARK_IL2_STAT5_SIGNALING | 0.387776 | 0.551153 | 0.237903 | 1.05331 | 1858 | 24 | c("MYC", "IGF1R", "BHLHE40", "IFITM3", "BCL2", "MAPKAPK2", "P2RX4", "PTRH2", "AHNAK", "GPX4") |
| 39 | HALLMARK_HEDGEHOG_SIGNALING | 0.440343 | 0.553252 | -0.607985 | -1.032988 | 2206 | 2 | c("TLE3", "SLIT1") |
| 40 | HALLMARK_COMPLEMENT | 0.589947 | 0.722685 | 0.266396 | 0.886253 | 2898 | 11 | c("CTSD", "HSPA5") |
| 41 | HALLMARK_NOTCH_SIGNALING | 0.693581 | 0.816326 | -0.308361 | -0.803678 | 3478 | 6 | c("PSENEN", "LFNG", "ARRB1", "HES1") |
| 42 | HALLMARK_OXIDATIVE_PHOSPHORYLATION | 0.721803 | 0.816326 | 0.17787 | 0.79721 | 3442 | 25 | c("SLC25A6", "BAX", "COX6C", "MRPL34", "LDHA", "SLC25A5", "GPX4", "ATP5MC1", "CS", "ATP5F1D", "NDUFA7", "MGST3") |
| 43 | HALLMARK_UV_RESPONSE_UP | 0.749687 | 0.816326 | 0.180482 | 0.776174 | 3590 | 22 | c("GAL", "ALDOA", "PDLIM3", "H2AX", "TFRC", "DDX21", "EIF5", "JUNB", "BTG2", "ENO2", "FOS") |
| 44 | HALLMARK_ANGIOGENESIS | 0.710113 | 0.816326 | 0.408107 | 0.813297 | 3524 | 3 | c("COL3A1", "JAG2") |
| 45 | HALLMARK_KRAS_SIGNALING_UP | 0.748748 | 0.816326 | -0.313338 | -0.759875 | 3736 | 5 | c("TSPAN1", "JUP", "APOD") |
| 46 | HALLMARK_KRAS_SIGNALING_DN | 0.793504 | 0.845255 | 0.260168 | 0.728456 | 3957 | 7 | c("CELSR2", "EDN1") |
| 47 | HALLMARK_XENOBIOTIC_METABOLISM | 0.814092 | 0.848734 | -0.160922 | -0.712642 | 4251 | 27 | c("FBP1", "ESR1", "SPINT2", "HES6", "BLVRB", "RAP1GAP", "JUP", "IGFBP4", "BCAR1", "TMBIM6", "NQO1", "CSAD") |
| 48 | HALLMARK_PEROXISOME | 0.83405 | 0.851426 | -0.201456 | -0.679044 | 4266 | 12 | c("SULT2B1", "CRABP2", "DHCR24", "IDH2", "CTBP1") |
| 49 | HALLMARK_SPERMATOGENESIS | 0.956513 | 0.956513 | 0.224974 | 0.552439 | 4794 | 5 | c("PSMG1", "AURKA", "CCNB2", "CDKN3") |

| Cluster 8 | pathway | pval | padj | ES | NES | nMoreExtreme | size | leadingEdge |
|---|---|---|---|---|---|---|---|---|
| 1 | HALLMARK_ANDROGEN_RESPONSE | 0.000221 | 0.005532 | 0.509566 | 2.144249 | 1 | 31 | c("SCD", "SAT1", "ACSL3", "ABHD2", "INSIG1", "DBI", "DHCR24", "KRT8", "ZBTB10", "TMPRSS2", "TMEM50A", "IDI1", "HSD17B14", "KRT19", "SPDEF", "ZMIZ1", "ITGAV") |
| 2 | HALLMARK_EPITHELIAL_MESENCHYMAL_TRANSITION | 0.000221 | 0.005532 | 0.512545 | 2.156788 | 1 | 31 | c("TIMP3", "SDC4", "SAT1", "ITGB5", "BASP1", "SDC1", "IGFBP4", "PMP22", "QSOX1", "RHOB", "CD59", "FUCA1", "TIMP1", "SGCG", "IGFBP2", "ITGB1", "ITGAV") |
| 3 | HALLMARK_MYC_TARGETS_V2 | 0.000333 | 0.005544 | -0.86562 | -2.53773 | 0 | 6 | c("HSPE1", "MYC", "MYBBP1A", "NOP56", "PPAN", "UNG") |
| 4 | HALLMARK_G2M_CHECKPOINT | 0.000612 | 0.00668 | -0.52561 | -2.70974 | 0 | 18 | c("SNRPD1", "DDX39A", "MYC", "HSPA8", "PTTG1", "UBE2S", "ILF3", "PRMT5", "POLE", "H2AZ1", "NASP", "CDC20") |
| 5 | HALLMARK_E2F_TARGETS | 0.000668 | 0.00668 | -0.87127 | -4.72372 | 0 | 20 | c("MCM7", "DDX39A", "MYC", "NOP56", "PTTG1", "UBE2S", "ILF3", "NME1", "SSRP1", "TP53", "RAD51C", "POLE", "H2AZ1", "TUBB", "HMGB2", "NASP", "CDC20", "POP7", "ASF1A", "UNG") |
| 6 | HALLMARK_MYC_TARGETS_V1 | 0.000922 | 0.00768 | -0.47851 | -3.06432 | 0 | 28 | c("LDHA", "MCM7", "HSPE1", "SNRPD1", "LSM7", "RPL22", "MYC", "NOP56", "RPS10", "PGK1", "NME1", "H2AZ1", "FBL", "CDC20", "MRPL9", "PABPC4") |
| 7 | HALLMARK_MYOGENESIS | 0.002389 | 0.017066 | 0.449249 | 1.966575 | 21 | 36 | c("APOD", "CLU", "SCD", "BAG1", "ITGB5", "ATP6AP1", "MB", "SYNGR2", "SGCG", "CRAT", "SPDEF", "ITGB1", "CKB", "MEF2D", "GAA", "ERBB3", "GABARAPL2") |
| 8 | HALLMARK_CHOLESTEROL_HOMEOSTASIS | 0.003084 | 0.019275 | 0.452813 | 1.92035 | 27 | 32 | c("CLU", "SCD", "SEMA3B", "LSS", "CD9", "FASN", "DHCR7", "SQLE", "LDLR", "TM7SF2", "CYP51A1", "ANXA5", "IDI1") |
| 9 | HALLMARK_HEME_METABOLISM | 0.005864 | 0.03258 | 0.421958 | 1.847109 | 53 | 36 | c("AQP3", "PDZK1IP1", "LRP10", "BSG", "HEBP1", "LAMP2", "BCAM", "DCAF10", "ADIPOR1", "SIDT2", "IGSF3", "CTSB", "RAP1GAP", "EIF2AK1", "GDE1") |
| 10 | HALLMARK_KRAS_SIGNALING_UP | 0.007929 | 0.039644 | 0.517734 | 1.837324 | 65 | 17 | c("APOD", "TSPAN1", "SEMA3B", "JUP", "PCSK1N", "FUCA1") |
| 11 | HALLMARK_COAGULATION | 0.009323 | 0.042379 | 0.452517 | 1.802426 | 81 | 25 | c("TIMP3", "CLU", "CD9", "LAMP2", "CSRP1", "CTSH", "ARF4", "TIMP1", "S100A13", "MMP15", "MSRB2", "CTSB") |
| 12 | HALLMARK_MTORC1_SIGNALING | 0.012495 | 0.048444 | 0.365873 | 1.715637 | 117 | 48 | c("SCD", "ACSL3", "CD9", "DHCR7", "SYTL2", "INSIG1", "IFI30", "SQLE", "DHCR24", "RPN1", "QDPR", "LDLR", "SLC9A3R1", "TM7SF2", "CYP51A1", "IDI1", "XBP1") |
| 13 | HALLMARK_UV_RESPONSE_UP | 0.012596 | 0.048444 | 0.390067 | 1.74223 | 116 | 39 | c("AQP3", "RET", "CYB5R1", "BSG", "CDKN2B", "RPN1", "ATP6V1F", "TMBIM6", "RHOB", "EPCAM", "SPR", "GRINA", "IGFBP2", "OLFM1", "CLTB", "PPT1", "CREG1", "FURIN") |
| 14 | HALLMARK_ESTROGEN_RESPONSE_EARLY | 0.014173 | 0.050617 | 0.322417 | 1.642545 | 137 | 73 | c("AQP3", "BAG1", "RET", "SULT2B1", "SEMA3B", "ABHD2", "CLIC3", "FASN", "IGFBP4", "RETREG1", "DHCR7", "SLC7A2", "INHBB", "SLC39A6", "TFF1", "RHOD", "ITPK1", "GFRA1", "KRT8", "SH3BP5", "SLC9A3R1", "XBP1", "CANT1", "TOB1", "KRT19", "OLFM1") |
| 15 | HALLMARK_COMPLEMENT | 0.022554 | 0.07518 | 0.383513 | 1.664475 | 206 | 35 | c("CLU", "USP14", "LAMP2", "CSRP1", "CTSH", "CD59", "TIMP1", "S100A13", "MMP15", "CTSB", "ANXA5", "GATA3", "GNAI2", "PFN1", "PPP4C", "CD46", "PRKCD", "MSRB1", "ANG", "LGMN") |

| | | | | | | | | |
|---|---|---|---|---|---|---|---|---|
| 16 | HALLMARK_ESTROGEN_RESPONSE_LATE | 0.027588 | 0.086214 | 0.313048 | 1.570656 | 266 | 67 | c("GPER1", "BAG1", "RET", "SULT2B1", "CDH1", "SEMA3B", "CD9", "ABHD2", "CLIC3", "IGFBP4", "DHCR7", "MDK", "TFF1", "LSR", "ISG20", "ITPK1", "PERP") |
| 17 | HALLMARK_XENOBIOTIC_METABOLISM | 0.033696 | 0.099105 | 0.357146 | 1.595186 | 312 | 39 | c("BCAR1", "FBP1", "COMT", "IGFBP4", "SPINT2", "JUP", "TMBIM6", "AOX1", "NINJ1", "RAP1GAP", "PGRMC1", "SLC1A5", "IDH1", "SAR1B", "DDAH2", "ESR1", "ACP2", "ECH1", "PINK1", "POR", "NPC1", "PGD", "BLVRB", "DHRS7", "ACO2") |
| 18 | HALLMARK_APOPTOSIS | 0.042929 | 0.119247 | 0.361787 | 1.570185 | 393 | 35 | c("TIMP3", "CLU", "SAT1", "ERBB2", "ISG20", "HSPB1", "RHOB", "TIMP1", "TSPO") |
| 19 | HALLMARK_APICAL_JUNCTION | 0.046937 | 0.123519 | 0.345752 | 1.544297 | 435 | 39 | c("CDH1", "NECTIN2", "MDK", "INSIG1", "JUP", "CERCAM", "CLDN4", "EVL", "CRAT", "ITGB1", "GAMT", "GNAI2", "MVD", "PFN1", "ITGA3", "CLDN7", "MYH9", "MYL12B") |
| 20 | HALLMARK_TNFA_SIGNALING_VIA_NFKB | 0.081543 | 0.19047 | 0.343678 | 1.46862 | 743 | 33 | c("SDC4", "SAT1", "LITAF", "ACKR3", "RHOB", "NINJ1", "LDLR", "ZBTB10", "BCL3", "EFNA1") |
| 21 | HALLMARK_ADIPOGENESIS | 0.081543 | 0.19047 | 0.295191 | 1.42821 | 777 | 56 | c("APLP2", "DHCR7", "SLC25A1", "REEP5", "QDPR", "TOB1", "CRAT", "SLC1A5", "UCP2", "IDH1", "DNAJC15", "COX6A1", "SCP2", "GHITM", "ECHS1", "ADIPOR2", "SDHC", "ECH1", "POR", "ACLY", "TST", "JAGN1", "DHRS7", "ATP1B3", "DHRS7B", "CCNG2", "ACO2", "SLC66A3", "RNF11", "PPP1R15B", "RMDN3", "ACOX1", "LPCAT3", "CMBL", "PIM3", "SULT1A1", "REEP6", "NDUFA5", "DNAJB9", "RTN3", "RETSAT") |
| 22 | HALLMARK_UV_RESPONSE_DN | 0.083807 | 0.19047 | -0.26047 | -1.44127 | 117 | 21 | c("ID1", "IGF1R", "BHLHE40", "MYC", "DLC1") |
| 23 | HALLMARK_PEROXISOME | 0.121811 | 0.253772 | 0.34495 | 1.38756 | 1078 | 26 | c("CRABP2", "SULT2B1", "DHCR24", "TSPO", "IDI1", "ECI2", "CRAT", "ISOC1", "IDH1", "IDH2", "PABPC1", "SCP2") |
| 24 | HALLMARK_KRAS_SIGNALING_DN | 0.117487 | 0.253772 | 0.453995 | 1.39974 | 906 | 11 | c("LYPD3", "IGFBP2", "GAMT", "CPB1", "GP2", "MFSD6") |
| 25 | HALLMARK_APICAL_SURFACE | 0.16603 | 0.33206 | 0.414852 | 1.317708 | 1303 | 12 | c("LYPD3", "GSTM3", "HSPB1", "GATA3") |
| 26 | HALLMARK_HEDGEHOG_SIGNALING | 0.174543 | 0.335659 | 0.590944 | 1.290819 | 1144 | 4 | c("SLIT1", "CELSR1", "MYH9") |
| 27 | HALLMARK_SPERMATOGENESIS | 0.192648 | 0.356756 | 0.536621 | 1.264336 | 1304 | 5 | c("GSTM3", "PCSK1N") |
| 28 | HALLMARK_INTERFERON_ALPHA_RESPONSE | 0.201171 | 0.359235 | 0.400618 | 1.272495 | 1579 | 12 | c("IFI30", "LY6E", "ISG20", "HLA-C", "IFITM2") |
| 29 | HALLMARK_HYPOXIA | 0.288358 | 0.497169 | 0.275707 | 1.169257 | 2617 | 32 | c("SDC4", "SULT2B1", "FBP1", "SLC25A1", "ISG20", "KDELR3", "ACKR3", "EFNA1", "GAA", "NEDD4L", "HEXA", "MYH9") |
| 30 | HALLMARK_P53_PATHWAY | 0.309253 | 0.498824 | 0.241164 | 1.147278 | 2940 | 51 | c("SAT1", "SDC1", "MXD4", "INHBB", "IFI30", "VAMP8", "CDKN2B", "PERP", "NINJ1", "FUCA1") |
| 31 | HALLMARK_ANGIOGENESIS | 0.309271 | 0.498824 | 0.483078 | 1.138183 | 2094 | 5 | c("TIMP1", "LRPAP1", "ITGAV", "FGFR1", "APP") |
| 32 | HALLMARK_MITOTIC_SPINDLE | 0.400802 | 0.626253 | -0.18901 | -1.02477 | 599 | 20 | c("FLNA", "PREX1", "TAOK2", "ARFGEF1", "CAPZB", "MARK4", "ARL8A", "RANBP9", "SEPTIN9", "RALBP1", "ACTN4", "WASL", "FARP1", "EZR", "RHOF", "CDC42EP4") |
| 33 | HALLMARK_INFLAMMATORY_RESPONSE | 0.43777 | 0.663288 | 0.28921 | 1.02634 | 3643 | 17 | c("SLC7A2", "LY6E", "LDLR", "TIMP1") |
| 34 | HALLMARK_NOTCH_SIGNALING | 0.480183 | 0.683705 | 0.451779 | 0.986837 | 3149 | 4 | c("ARRB1", "HES1", "APH1A", "DTX2") |
| 35 | HALLMARK_UNFOLDED_PROTEIN_RESPONSE | 0.484872 | 0.683705 | 0.247609 | 0.996008 | 4294 | 26 | c("EIF4EBP1", "KDELR3", "BAG3", "HERPUD1", "XBP1", "ALDH18A1", "WIPI1", "SPCS1") |

| | | | | | | | | |
|---|---|---|---|---|---|---|---|---|
| 36 | HALLMARK_IL2_STAT5_SIGNALING | 0.492268 | 0.683705 | 0.22287 | 0.987659 | 4551 | 38 | c("ST3GAL4", "RHOB", "TWSG1", "SOCS2", "SYNGR2", "PLEC", "XBP1", "COL6A1", "SERPINB6", "SLC1A5", "ITGAV", "BMPR2", "FURIN", "BCL2L1", "IGF2R", "HOPX", "IRF6", "CKAP4", "BATF") |
| 37 | HALLMARK_INTERFERON_GAMMA_RESPONSE | 0.554316 | 0.749076 | 0.226823 | 0.938694 | 4969 | 29 | c("IFI30", "HLA-DRB1", "VAMP8", "LY6E", "ISG20", "IFITM2", "ISOC1", "LGALS3BP", "VAMP5", "TAPBP", "HLA-B", "MVP", "RAPGEF6") |
| 38 | HALLMARK_PROTEIN_SECRETION | 0.661346 | 0.831544 | 0.188467 | 0.851765 | 6172 | 41 | c("CD63", "LAMP2", "ARF1", "NAPA", "ARCN1", "TMED10", "PPT1", "COPB2", "SEC24D", "COPE", "IGF2R", "COPB1", "RAB2A", "LMAN1", "TSG101", "SCRN1", "ATP6V1H", "SEC31A", "CLTA", "ADAM10", "SEC22B", "SCAMP3", "GLA", "SCAMP1", "KRT18", "RER1") |
| 39 | HALLMARK_GLYCOLYSIS | 0.640936 | 0.831544 | 0.186463 | 0.874352 | 6052 | 48 | c("SDC1", "CHPF", "FUT8", "QSOX1", "ISG20", "KDELR3", "CLDN3") |
| 40 | HALLMARK_ALLOGRAFT_REJECTION | 0.665235 | 0.831544 | 0.245104 | 0.835846 | 5420 | 15 | c("INHBB", "TIMP1", "BCL3", "HLA-E", "DEGS1", "TAPBP", "RPS9", "AKT1", "UBE2N", "CCND3") |
| 41 | HALLMARK_PI3K_AKT_MTOR_SIGNALING | 0.720517 | 0.857758 | -0.137 | -0.81078 | 891 | 24 | c("VAV3", "DDIT3", "MYD88", "MAPKAP1", "CAB39", "GRK2", "YWHAB", "PLA2G12A", "RIPK1", "AP2M1", "PAK4", "ITPR2", "MAPK1", "CFL1", "AKT1S1", "UBE2N", "AKT1", "ACTR2", "RALB", "PIK3R3", "UBE2D3", "PFN1", "ARPC3") |
| 42 | HALLMARK_BILE_ACID_METABOLISM | 0.716202 | 0.857758 | 0.199243 | 0.793606 | 6298 | 25 | c("SULT2B1", "DHCR24", "IDI1", "ISOC1", "IDH1", "IDH2", "SCP2", "NPC1", "ABCA2", "PEX11A", "SLC35B2", "RXRA", "ABCA3") |
| 43 | HALLMARK_WNT_BETA_CATENIN_SIGNALING | 0.743185 | 0.864168 | -0.28843 | -0.77489 | 2398 | 5 | c("MYC", "TP53") |
| 44 | HALLMARK_REACTIVE_OXYGEN_SPECIES_PATHWAY | 0.760947 | 0.864712 | 0.250288 | 0.749323 | 5786 | 10 | c("MGST1", "SELENOS", "PTPA", "TXNRD1") |
| 45 | HALLMARK_IL6_JAK_STAT3_SIGNALING | 0.807698 | 0.897442 | 0.255944 | 0.710039 | 5938 | 8 | c("CD9", "IL13RA1") |
| 46 | HALLMARK_TGF_BETA_SIGNALING | 0.891398 | 0.915867 | 0.181227 | 0.618016 | 7263 | 15 | c("CDH1", "BMPR2", "FURIN", "UBE2D3", "NCOR2", "XIAP", "SMAD3", "ENG") |
| 47 | HALLMARK_DNA_REPAIR | 0.847059 | 0.915867 | -0.11299 | -0.70954 | 935 | 27 | c("CCNO", "DDB2", "NME1", "SSRP1", "TP53") |
| 48 | HALLMARK_FATTY_ACID_METABOLISM | 0.872241 | 0.915867 | 0.141626 | 0.650973 | 8178 | 44 | c("FASN", "AUH", "DHCR24", "IDI1", "ECI2", "CRAT", "ERP29", "SERINC1", "IDH1", "ALAD", "ECI1", "ECHS1", "ADIPOR2", "SDHC", "ECH1") |
| 49 | HALLMARK_PANCREAS_BETA_CELLS | 0.897549 | 0.915867 | 0.271124 | 0.638796 | 6079 | 5 | c("SPCS1", "CHGA", "SRP9", "STXBP1") |
| 50 | HALLMARK_OXIDATIVE_PHOSPHORYLATION | 0.981979 | 0.981979 | 0.091223 | 0.44767 | 9426 | 60 | c("ATP6AP1", "ATP6V0C", "ATP6V1F", "ATP6V0B", "CYB5R3", "ATP6V1G1", "NDUFC2", "IDH1", "IDH2", "ATP6V0E1", "COX6A1", "OAT", "NDUFA4", "ECI1", "ATP5PD", "ECHS1", "SDHC", "ECH1") |

| Cluster 10 | pathway | pval | padj | ES | NES | nMoreExtreme | size | leadingEdge |
|---|---|---|---|---|---|---|---|---|
| 1 | HALLMARK_G2M_CHECKPOINT | 0.000152 | 0.002541 | 0.466684 | 2.831758 | 0 | 72 | c("HMGA1", "SLC7A5", "MT2A", "MYBL2", "MYC", "H2AX", "NCL", "CKS1B", "PLK1", "UBE2C", "HSPA8", "KPNA2", "BIRC5", "JPT1", "TPX2", "RAD23B", "CDK4", "MKI67", "TMPO", "CCNA2", "CKS2", "MAD2L1", "SLC7A1", "MCM2", "PTTG1", "CDC20", "LMNB1", "GINS2", "NOLC1", "MCM6", "ODC1", "RAD21", "TRA2B", "CCNB2", "CDKN3", "DTYMK", "CCND1", "SYNCRIP", "HNRNPD", "TOP2A", "GSPT1", "TACC3", "UBE2S", "UCK2", "PRC1", "G3BP1", "RACGAP1", "DBF4") |
| 2 | HALLMARK_E2F_TARGETS | 0.000148 | 0.002541 | 0.499727 | 3.200136 | 0 | 90 | c("HMGA1", "MYBL2", "MYC", "TUBB", "H2AX", "CKS1B", "HMGB2", "PLK1", "KPNA2", "ANP32E", "PRDX4", "BIRC5", "JPT1", "PAICS", "NME1", "MTHFD2", "XRCC6", "RAN", "DCTPP1", "CDK4", "NOP56", "MKI67", "TMPO", "TP53", "TK1", "DEK", "RRM2", "CKS2", "MAD2L1", "MCM2", "EIF2S1", "PTTG1", "MCM4", "CDC20", "LMNB1", "NOLC1", "RAD51C", "MCM6", "RAD21", "TRA2B", "PCNA", "RFC3", "CCNB2", "CDKN3", "RPA3", "NCAPD2", "MCM7", "SYNCRIP", "HNRNPD", "TOP2A", "SNRPB", "PRPS1", "ASF1B", "GSPT1", "NAP1L1", "TACC3", "RANBP1", □ "UBE2S", "PA2G4", "TBRG4") |
| 3 | HALLMARK_MYC_TARGETS_V1 | 0.000145 | 0.002541 | 0.486482 | 3.338953 | 0 | 124 | c("MYC", "FBL", "LDHA", "PGK1", "HSPD1", "KPNA2", "HDGF", "CCT3", "PRDX4", "PSMA7", "CCT7", "CCT2", "C1QBP", "NME1", "HPRT1", "SET", "CCT5", "CYC1", "KARS1", "CCT4", "XRCC6", "SLC25A3", "RAN", "RAD23B", "RSL1D1", "CDK4", "NOP56", "RPS2", "YWHAQ", "DDX21", "EIF3B", "GLO1", "DEK", "SNRPD2", "CCNA2", "IMPDH2", "MAD2L1", "MCM2", "EIF1AX", "EIF2S1", "EEF1B2", "SERBP1", "PSMB3", "MCM4", "CDC20", "SRSF3", "PPIA", "SNRPA1", "NOLC1", "PHB2", "MCM6", "HNRNPC", "ODC1", "HNRNPR", "CANX", "TRA2B", "SF3B3", "PCNA", □ "EIF4H", "AIMP2", "PSMD7", "ABCE1", "HNRNPA1", "MCM7", "SRM", "TRIM28", "MRPL23", "SYNCRIP", "TUFM", "POLE3", "HNRNPD", "NOP16", "VDAC1", "FAM120A", "GSPT1", "RPS5", "HSP90AB1", "NAP1L1", "RANBP1", "PA2G4", "UBA2", "ILF2", "G3BP1", "XPOT", "IARS1", "ETF1", "MRPL9", "PABPC4") |
| 4 | HALLMARK_MYC_TARGETS_V2 | 0.000648 | 0.008096 | 0.469195 | 2.176704 | 3 | 29 | c("MYC", "HSPD1", "PLK1", "DCTPP1", "CDK4", "NOP56", "MCM4", "NOLC1", "AIMP2", "WDR43", "EXOSC5", "SRM", "TMEM97", "NOP16", "MRTO4", "PA2G4", "TBRG4", "SLC19A1", "PES1", "PUS1", "IPO4", "WDR74", "NPM1", "HSPE1") |

| | | | | | | | | |
|---|---|---|---|---|---|---|---|---|
| 5 | HALLMARK_ESTROGEN_RESPONSE_LATE | 0.00091 | 0.009103 | 0.333287 | 2.032751 | 5 | 73 | c("PRSS23", "AGR2", "GAL", "SERPINA3", "SIAH2", "SLC7A5", "KLK10", "COX6C", "CCN5", "AREG", "HPRT1", "ETFB", "FAM102A", "FKBP5", "PDLIM3", "NBL1", "HOMER2", "SCARB1", "KLK11", "CDC20", "GINS2", "ASS1", "FABP5", "FKBP4", "TFF1", "ELOVL5", "CCND1", "CA12", "PDCD4", "EMP2", "TOP2A", "MEST", "CXCL12", "NRIP1", "CD44", "TSPAN13", "NXT1", "PPIF", "TFPI2", "TPD52L1", "RAB31") |
| 6 | HALLMARK_OXIDATIVE_PHOSPHORYLATION | 0.002306 | 0.019213 | 0.32065 | 1.916818 | 14 | 68 | c("LDHA", "SLC25A5", "COX6C", "COX8A", "MDH2", "ETFB", "CYC1", "ATP5MC3", "SLC25A3", "NDUFB7", "CYCS", "MRPL11", "TIMM8B", "CS", "MRPS12", "GPI", "ATP5F1D", "PHB2", "TIMM10", "ATP5ME", "HSD17B10", "UQCRFS1", "UQCRC1", "HSPA9", "ATP5F1B", "TIMM50", "FH", "ATP5MC1", "VDAC1", "LRPPRC", "ATP5F1C", "ETFA", "CYB5R3", "COX7B", "NDUFA4", "COX6A1", "DLD", "MRPS11", "NDUFB8", "SDHD", "GPX4", "NDUFB4", "SDHC", "MRPS15") |
| 7 | HALLMARK_MYOGENESIS | 0.00287 | 0.020499 | -0.413285 | -2.040708 | 10 | 27 | c("SCD", "ITGB5", "GABARAPL2", "CLU", "ATP6AP1", "MB", "SPDEF", "CKB", "SYNGR2", "SH2B1", "MYO1C", "GAA", "SORBS3", "PLXNB2", "EIF4A2") |
| 8 | HALLMARK_GLYCOLYSIS | 0.004127 | 0.025794 | 0.353639 | 1.883002 | 25 | 45 | c("LDHA", "PGK1", "ENO1", "MIF", "G6PD", "HSPA5", "PGAM1", "MDH2", "TXN", "B4GALT1", "PFKP", "PPIA", "SOD1", "ARPP19", "IER3", "STC2", "TPI1", "FKBP4", "PKM", "PRPS1", "NDUFV3", "P4HA1", "CD44", "FAM162A", "DLD", "CLN6", "STMN1", "ERO1A") |
| 9 | HALLMARK_HYPOXIA | 0.004865 | 0.027028 | 0.337278 | 1.854758 | 30 | 50 | c("SIAH2", "MT2A", "LDHA", "PGK1", "ENO1", "MIF", "CCN5", "HSPA5", "LXN", "ERRFI1", "SCARB1", "SLC2A1", "AKAP12", "PFKP", "GPI", "GAPDH", "BHLHE40", "IER3", "JUN", "STC2", "TPI1", "ANXA2") |
| 10 | HALLMARK_ESTROGEN_RESPONSE_EARLY | 0.00716 | 0.032547 | 0.290659 | 1.758755 | 46 | 71 | c("PRSS23", "TSKU", "SIAH2", "SLC7A5", "MYC", "KRT8", "KLK10", "KRT18", "CCN5", "AREG", "CBFA2T3", "FAM102A", "FKBP5", "PDLIM3", "NBL1", "B4GALT1", "SCARB1", "SLC2A1", "GREB1", "TGIF2", "IGF1R", "BHLHE40", "DLC1", "STC2", "FKBP4", "TFF1", "KCNK15", "ELOVL5", "CCND1", "CA12") |
| 11 | HALLMARK_UNFOLDED_PROTEIN_RESPONSE | 0.006814 | 0.032547 | 0.36377 | 1.887454 | 42 | 41 | c("SLC7A5", "H2AX", "CKS1B", "CALR", "LSM4", "HSPA5", "PDIA6", "MTHFD2", "NOP56", "HYOU1", "EIF2S1", "HSP90B1", "NOLC1", "EIF4A3", "STC2", "HSPA9", "EXOSC5", "GEMIN4") |
| 12 | HALLMARK_CHOLESTEROL_HOMEOSTASIS | 0.009066 | 0.037774 | -0.417677 | -1.888979 | 35 | 21 | c("CD9", "SCD", "CLU", "FASN", "TM7SF2", "TP53INP1", "SQLE", "SEMA3B", "LSS", "ABCA2") |
| 13 | HALLMARK_MTORC1_SIGNALING | 0.011864 | 0.045631 | 0.261557 | 1.682775 | 79 | 92 | c("SLC7A5", "LDHA", "PGK1", "ENO1", "HSPD1", "PLK1", "CALR", "CACYBP", "PHGDH", "G6PD", "HSPA5", "MTHFD2", "HPRT1", "SLC1A5", "TOMM40", "RRM2", "IGFBP5", "MCM2", "SLC2A1", "MCM4", "PPIA", "HSP90B1", "GPI", "GAPDH", "BHLHE40", "GMPS", "ATP2A2", "CANX", "PPA1", "TPI1", "SHMT2", "HSPA9") |

| | | | | | | | | |
|---|---|---|---|---|---|---|---|---|
| 14 | HALLMARK_ALLOGRAFT_REJECTION | 0.020837 | 0.074417 | 0.398491 | 1.740294 | 127 | 24 | c("FLNA", "EIF5A", "RPL39", "RPS19", "NME1", "RPS9", "HLA-A", "ABCE1", "STAT1", "B2M", "HIF1A", "UBE2N", "MRPL3", "AARS1", "TPD52", "NPM1") |
| 15 | HALLMARK_ADIPOGENESIS | 0.032407 | 0.108022 | 0.3087 | 1.623754 | 203 | 43 | c("COX8A", "MDH2", "ETFB", "CYC1", "SLC1A5", "NDUFB7", "YWHAG", "CS", "SCARB1", "UBC", "SOD1", "CHCHD10", "UQCRC1", "PDCD4", "TKT", "RTN3", "DDT", "SLC19A1", "COX7B", "COX6A1", "DLD", "STOM", "GPX4", "HSPB8", "GBE1", "SDHC", "DHCR7", "UBQLN1", "ALDOA", "NDUFS3") |
| 16 | HALLMARK_FATTY_ACID_METABOLISM | 0.03902 | 0.121936 | 0.340336 | 1.613857 | 241 | 31 | c("LDHA", "LGALS1", "MIF", "HPGD", "MDH2", "S100A10", "HSP90AA1", "CBR1", "ODC1", "OSTC", "HSD17B10", "ELOVL5", "YWHAH", "NTHL1", "FH") |
| 17 | HALLMARK_INTERFERON_ALPHA_RESPONSE | 0.056702 | 0.16677 | 0.391553 | 1.546132 | 337 | 18 | c("IFI27", "ISG15", "IFITM1", "IFITM3", "ADAR", "B2M", "PSMA3", "HERC6", "EIF2AK2") |
| 18 | HALLMARK_REACTIVE_OXYGEN_SPECIES_PATHWAY | 0.068264 | 0.189622 | 0.446291 | 1.510652 | 395 | 12 | c("G6PD", "PRDX4", "TXN", "PFKP", "SOD1", "GSR", "JUNB", "GPX4", "NDUFB4") |
| 19 | HALLMARK_APOPTOSIS | 0.07659 | 0.201553 | 0.297415 | 1.469914 | 477 | 35 | c("HMGB2", "KRT18", "HSPB1", "SOD1", "TNFRSF12A", "IER3", "LMNA", "CTNNB1", "JUN", "BID", "CCND1", "IFITM3", "PDCD4", "TOP2A", "GSR", "DNAJA1", "DFFA", "CD44", "ETF1", "LGALS3") |
| 20 | HALLMARK_UV_RESPONSE_UP | 0.084251 | 0.210627 | 0.286861 | 1.441586 | 527 | 37 | c("GAL", "H2AX", "DDX21", "PDLIM3", "FKBP4", "BID", "MRPL23", "POLE3", "DNAJA1", "JUNB", "RPN1", "GGH", "PPIF", "EIF5", "SLC6A8", "SPR", "TMBIM6", "TARS1", "FEN1", "PSMC3", "TFRC", "CEBPG", "ALDOA", "CYB5B", "AMD1", "STIP1") |
| 21 | HALLMARK_MITOTIC_SPINDLE | 0.09371 | 0.212977 | 0.292934 | 1.434737 | 583 | 34 | c("FLNA", "PLK1", "BIRC5", "TPX2", "LMNB1", "CDC42EP1", "CCNB2", "GEMIN4", "ARHGDIA", "MAPRE1", "TOP2A", "DYNLL2", "LRPPRC", "CDC42", "PRC1", "PALLD", "RACGAP1") |
| 22 | HALLMARK_WNT_BETA_CATENIN_SIGNALING | 0.093475 | 0.212977 | 0.584117 | 1.415857 | 519 | 5 | c("MYC", "TP53", "CTNNB1") |
| 23 | HALLMARK_INTERFERON_GAMMA_RESPONSE | 0.110559 | 0.235259 | 0.282675 | 1.397064 | 689 | 35 | c("IFI27", "ISG15", "MT2A", "MTHFD2", "OAS3", "PFKP", "HLA-A", "IFITM3", "STAT1", "ADAR", "B2M", "PSMA3", "HIF1A", "HERC6", "EIF2AK2", "IFIT1", "OAS2", "NAMPT", "PSMB10", "PSMB2", "PSMA2", "ISG20", "LYSMD2", "HELZ2", "SRI", "PLSCR1", "PNP", "TRIM14", "CDKN1A", "BTG1") |
| 24 | HALLMARK_PI3K_AKT_MTOR_SIGNALING | 0.112924 | 0.235259 | 0.329673 | 1.397393 | 684 | 22 | c("CFL1", "CALR", "CDK4", "YWHAB", "PFN1", "SLC2A1", "VAV3", "HSP90B1", "ARHGDIA", "RAC1") |
| 25 | HALLMARK_EPITHELIAL_MESENCHYMAL_TRANSITION | 0.138534 | 0.277067 | 0.286289 | 1.343152 | 855 | 30 | c("FLNA", "PPIB", "LGALS1", "TPM4", "AREG", "TNFRSF12A", "JUN", "COLGALT1", "MCM7", "MEST", "CXCL12", "COL3A1", "CD44", "TFPI2", "SLC6A8") |
| 26 | HALLMARK_PROTEIN_SECRETION | 0.175286 | 0.337088 | 0.364173 | 1.305148 | 1026 | 14 | c("KRT18", "ANP32E", "TMX1", "BNIP3", "SOD1", "TMED2") |
| 27 | HALLMARK_INFLAMMATORY_RESPONSE | 0.220003 | 0.407414 | 0.346478 | 1.241734 | 1288 | 14 | c("MYC", "IFITM1", "SLC7A1", "P2RX4", "ATP2A2", "HIF1A", "EIF2AK2", "KLF6", "ADRM1", "NAMPT") |
| 28 | HALLMARK_UV_RESPONSE_DN | 0.260527 | 0.465226 | 0.30091 | 1.188207 | 1552 | 18 | c("MYC", "ID1", "IGFBP5", "SLC7A1", "IGF1R", "BHLHE40", "DLC1", "ANXA2") |
| 29 | HALLMARK_APICAL_JUNCTION | 0.308014 | 0.501217 | -0.234832 | -1.123997 | 1179 | 25 | c("EVL", "MDK", "CLDN4", "PIK3R3", "INSIG1", "PKD1") |
| 30 | HALLMARK_APICAL_SURFACE | 0.310754 | 0.501217 | -0.489949 | -1.125005 | 1412 | 4 | c("GATA3", "SULF2") |
| 31 | HALLMARK_BILE_ACID_METABOLISM | 0.300951 | 0.501217 | -0.366584 | -1.128067 | 1297 | 8 | c("SULT2B1", "EFHC1", "ABCA2", "RXRA") |

| | | | | | | | | |
|---|---|---|---|---|---|---|---|---|
| 32 | HALLMARK_XENOBIOTIC_METABOLISM | 0.369171 | 0.56172 | 0.231687 | 1.07485 | 2279 | 29 | c("MT2A", "HPRT1", "KARS1", "SLC1A5", "GSTO1", "ARPP19", "CBR1", "ATP2A2", "SHMT2", "ELOVL5", "GSR", "TMEM97", "DDT", "PGD") |
| 33 | HALLMARK_KRAS_SIGNALING_UP | 0.370735 | 0.56172 | 0.353967 | 1.074706 | 2107 | 9 | c("SERPINA3", "LCP1", "AKAP12") |
| 34 | HALLMARK_COAGULATION | 0.385745 | 0.567272 | -0.277422 | -1.047111 | 1601 | 13 | c("CD9", "CRIP2", "CLU") |
| 35 | HALLMARK_KRAS_SIGNALING_DN | 0.444901 | 0.635574 | -0.499165 | -1.013681 | 2054 | 3 | c("EDN1", "MACROH2A2", "SPTBN2") |
| 36 | HALLMARK_NOTCH_SIGNALING | 0.470376 | 0.635644 | -0.388532 | -0.978616 | 2087 | 5 | c("PSENEN", "HES1", "ARRB1") |
| 37 | HALLMARK_ANDROGEN_RESPONSE | 0.466228 | 0.635644 | 0.206701 | 0.993452 | 2905 | 32 | c("KRT8", "HPGD", "XRCC6", "FKBP5", "XRCC5", "HOMER2", "B4GALT1", "AKAP12", "MYL12A", "ELOVL5", "CCND1", "GSR", "PA2G4", "B2M", "SMS", "TPD52", "SLC38A2", "ADRM1", "SPCS3", "VAPA", "SRF", "STK39") |
| 38 | HALLMARK_TGF_BETA_SIGNALING | 0.507154 | 0.667308 | 0.283239 | 0.958737 | 2941 | 12 | c("ID1", "CTNNB1", "RHOA", "JUNB", "RAB31", "THBS1", "FKBP1A", "ID3", "CDK9", "XIAP", "SPTBN1") |
| 39 | HALLMARK_TNFA_SIGNALING_VIA_NFKB | 0.523047 | 0.670573 | 0.211175 | 0.946447 | 3233 | 26 | c("MYC", "AREG", "B4GALT1", "PHLDA2", "BHLHE40", "IER3", "JUN", "CCND1", "JUNB", "KLF2", "CD44") |
| 40 | HALLMARK_COMPLEMENT | 0.583547 | 0.729433 | 0.249216 | 0.893159 | 3418 | 14 | c("CTSD", "HSPA5", "PFN1") |
| 41 | HALLMARK_P53_PATHWAY | 0.623987 | 0.76096 | 0.166777 | 0.877241 | 3927 | 43 | c("CTSD", "S100A10", "IRAK1", "SLC3A2", "TP53", "RPL36", "RAD51C", "IER3", "JUN", "PCNA", "RAP2B", "OSGIN1") |
| 42 | HALLMARK_DNA_REPAIR | 0.656305 | 0.763145 | 0.171635 | 0.848274 | 4095 | 35 | c("NME1", "HPRT1", "ARL6IP1", "ALYREF", "TP53", "IMPDH2", "APRT", "TMED2", "PCNA", "RFC3", "RPA3", "POLR1D") |
| 43 | HALLMARK_HEDGEHOG_SIGNALING | 0.641722 | 0.763145 | -0.505562 | -0.874117 | 3069 | 2 | c("TLE3", "ETS2") |
| 44 | HALLMARK_IL6_JAK_STAT3_SIGNALING | 0.700417 | 0.778241 | -0.24745 | -0.803806 | 3022 | 9 | c("CD9", "STAT2", "TYK2") |
| 45 | HALLMARK_HEME_METABOLISM | 0.700076 | 0.778241 | -0.178122 | -0.80557 | 2779 | 21 | c("SELENBP1", "AQP3", "RBM5") |
| 46 | HALLMARK_SPERMATOGENESIS | 0.733927 | 0.797747 | -0.205372 | -0.775162 | 3047 | 13 | c("PHKG2", "SLC12A2", "PCSK1N", "CHFR", "TALDO1", "BUB1", "PSMG1", "AURKA", "KIF2C", "DBF4", "RPL39L", "CDKN3", "CCNB2") |
| 47 | HALLMARK_IL2_STAT5_SIGNALING | 0.857788 | 0.893445 | 0.141238 | 0.669743 | 5319 | 31 | c("MYC", "SLC1A5", "MAPKAPK2", "P2RX4", "IGF1R", "GSTO1", "BHLHE40", "ODC1", "IFITM3", "P4HA1", "UCK2", "CD44") |
| 48 | HALLMARK_PEROXISOME | 0.875576 | 0.893445 | 0.178991 | 0.641482 | 5129 | 14 | c("SOD1", "ELOVL5", "TOP2A", "YWHAH", "PRDX5", "CLN6", "PABPC1", "CRABP1", "CTPS1", "TSPO") |
| 49 | HALLMARK_PANCREAS_BETA_CELLS | 0.873955 | 0.893445 | -0.395439 | -0.683715 | 4180 | 2 | c("SRP14", "SRPRB") |
| 50 | HALLMARK_ANGIOGENESIS | 0.936133 | 0.936133 | -0.291041 | -0.591031 | 4323 | 3 | c("TIMP1", "S100A4", "COL3A1") |

| Cluster 15 | pathway | pval | padj | ES | NES | nMoreExtreme | size | leadingEdge |
|---|---|---|---|---|---|---|---|---|
| 1 | HALLMARK_ESTROGEN_RESPONSE_EARLY | 0.000403 | 0.009666 | -0.436809 | -2.490842 | 0 | 32 | c("PRSS23", "GREB1", "KLK10", "MLPH", "SIAH2", "IGF1R", "CELSR2", "TSKU", "AREG", "CXCL12", "SCNN1A", "CCND1", "DLC1") |
| 2 | HALLMARK_ESTROGEN_RESPONSE_LATE | 0.000394 | 0.009666 | -0.475012 | -2.639079 | 0 | 30 | c("PRSS23", "KLK11", "AGR2", "KLK10", "SIAH2", "PDCD4", "CELSR2", "AREG", "PLXNB1", "CXCL12", "SCNN1A", "CCND1") |
| 3 | HALLMARK_UV_RESPONSE_DN | 0.003624 | 0.051017 | -0.659295 | -2.00335 | 12 | 7 | c("IGFBP5", "ID1", "IGF1R") |
| 4 | HALLMARK_COAGULATION | 0.004251 | 0.051017 | -0.535764 | -1.989043 | 13 | 11 | c("PRSS23", "FGG") |
| 5 | HALLMARK_MITOTIC_SPINDLE | 0.02976 | 0.25157 | -0.451575 | -1.676486 | 97 | 11 | c("PREX1", "FLNA", "RHOT2", "TUBGCP6", "NET1", "FLNB", "SPTBN1", "CDC42", "PCGF5", "RHOF") |
| 6 | HALLMARK_APOPTOSIS | 0.04128 | 0.25157 | -0.344236 | -1.613036 | 119 | 19 | c("LGALS3", "TXNIP", "PDCD4", "RHOT2", "BIK", "CCND1", "BCAP31", "CTNNB1", "HSPB1", "BID", "SQSTM1", "GSR", "GPX4", "VDAC2", "LEF1", "DAP3") |
| 7 | HALLMARK_MYC_TARGETS_V1 | 0.034266 | 0.25157 | 0.293421 | 1.585412 | 272 | 54 | c("RPLP0", "RACK1", "RPS5", "RPS2", "RPL18", "EEF1B2", "RPS6", "HNRNPA1", "RPS10", "CANX", "NPM1", "NHP2", "RPS3", "RPL6", "GLO1", "SNRPD2", "RPL34", "YWHAE", "PRDX4", "PABPC1", "RPL14", "RSL1D1", "G3BP1", "NAP1L1", "PSMB3", "SLC25A3", "SET", "MRPL23", "PHB2", "CYC1", "ERH", "CCT2", "CLNS1A", "EIF1AX", "EIF4H") |
| 8 | HALLMARK_PEROXISOME | 0.041928 | 0.25157 | 0.597669 | 1.563294 | 266 | 6 | c("CRABP2", "TSPO", "PRDX5", "PABPC1") |
| 9 | HALLMARK_MYOGENESIS | 0.094451 | 0.453367 | 0.516241 | 1.428169 | 605 | 7 | c("TNNT1", "CKB", "ITGB5", "MB", "AK1", "ITGB4") |
| 10 | HALLMARK_SPERMATOGENESIS | 0.086894 | 0.453367 | -0.697203 | -1.452797 | 361 | 3 | c("PHKG2", "TALDO1", "RPL39L") |
| 11 | HALLMARK_WNT_BETA_CATENIN_SIGNALING | 0.104997 | 0.458171 | -0.59993 | -1.421769 | 415 | 4 | c("DKK1", "CSNK1E", "CTNNB1") |
| 12 | HALLMARK_P53_PATHWAY | 0.156762 | 0.53747 | 0.320667 | 1.321654 | 1128 | 22 | c("PVT1", "RPL36", "RACK1", "RPL18", "IFI30", "CCNG1") |
| 13 | HALLMARK_ALLOGRAFT_REJECTION | 0.146939 | 0.53747 | 0.378096 | 1.336546 | 1007 | 14 | c("RPS9", "RPS19", "RPL9", "NPM1", "CAPG", "DEGS1", "RPS3A", "UBE2N") |
| 14 | HALLMARK_KRAS_SIGNALING_DN | 0.151101 | 0.53747 | -0.441696 | -1.342149 | 541 | 7 | c("CALML5", "CELSR2", "LFNG") |
| 15 | HALLMARK_TGF_BETA_SIGNALING | 0.194169 | 0.621342 | -0.443907 | -1.256179 | 705 | 6 | c("ID1", "CTNNB1", "SPTBN1", "CDK9", "TGIF1", "HDAC1") |
| 16 | HALLMARK_G2M_CHECKPOINT | 0.271341 | 0.772515 | -0.299644 | -1.161469 | 889 | 12 | c("ILF3", "SNRPD1", "CCND1", "DDX39A", "CDK4", "MCM6") |
| 17 | HALLMARK_NOTCH_SIGNALING | 0.273599 | 0.772515 | -0.497574 | -1.179196 | 1083 | 4 | c("LFNG", "CCND1", "FBXW11") |
| 18 | HALLMARK_OXIDATIVE_PHOSPHORYLATION | 0.347214 | 0.925903 | 0.188633 | 1.097171 | 2859 | 72 | c("NDUFA2", "COX8A", "OAT", "ATP5MC3", "ATP6V0E1", "ATP5F1E", "NDUFB1", "ATP5PD", "NDUFA4", "NDUFB4", "ECI1", "NDUFA7", "SDHC", "CYB5A", "COX7A2L", "COX6A1", "ACAT1", "COX7B", "ATP5F1C", "SLC25A6", "FH", "NDUFB7", "ATP5F1D", "NDUFC1", "NDUFS7", "ATP6V1F", "SLC25A3", "HSD17B10", "COX6B1", "MGST3", "PHB2", "CYC1", "NDUFB5", "ACAA2", "CYB5R3", "NDUFC2", "ATP5MG", "POLR2F", "UQCRH", "COX7C", "UQCR10", "CYCS", "SDHD", "TOMM22", "TIMM8B", "ATP5MC2", "MDH2", "VDAC2") |
| 19 | HALLMARK_TNFA_SIGNALING_VIA_NFKB | 0.880745 | 0.983157 | -0.184429 | -0.653765 | 2931 | 10 | c("AREG", "CCND1", "LITAF", "SQSTM1", "TGIF1", "SAT1", "DRAM1", "TNIP1", "ACKR3") |
| 20 | HALLMARK_HYPOXIA | 0.772317 | 0.983157 | 0.201391 | 0.746046 | 5389 | 16 | c("FBP1", "MIF", "ALDOC", "PRDX5", "GAPDH", "KDELR3", "ACKR3", "ALDOA") |
| 21 | HALLMARK_CHOLESTEROL_HOMEOSTASIS | 0.778766 | 0.983157 | 0.255603 | 0.742193 | 5075 | 8 | c("DHCR7", "ALDOC", "CYP51A1", "S100A11") |

| | | | | | | | | |
|---|---|---|---|---|---|---|---|---|
| 22 | HALLMARK_DNA_REPAIR | 0.748882 | 0.983157 | -0.166503 | -0.780209 | 2176 | 19 | c("CCNO", "SRSF6", "TAF1C", "BCAP31", "GTF2H5", "POLR1H", "GPX4", "BCAM", "DCTN4", "MPC2", "DAD1", "POLR2F", "NME4", "POLR1D", "ALYREF", "AK1", "MPG", "APRT", "TMED2") |
| 23 | HALLMARK_ADIPOGENESIS | 0.861526 | 0.983157 | 0.139752 | 0.667985 | 6513 | 35 | c("COX8A", "UCP2", "DHCR7", "SDHC", "COX6A1", "TOB1", "LTC4S", "COX7B", "LPCAT3", "SULT1A1", "NDUFB7", "ALDOA", "MGST3", "CYC1", "ACAA2", "CHCHD10", "UQCR10", "MDH2", "UQCR11", "ATP1B3", "TST", "HADH", "SUCLG1", "GPX4", "PPP1R15B", "CMPK1", "CD302", "ACO2", "RTN3", "SCP2", "COQ5", "BCL2L13", "TALDO1", "ETFB") |
| 24 | HALLMARK_ANDROGEN_RESPONSE | 0.876708 | 0.983157 | -0.175173 | -0.650335 | 2886 | 11 | c("HPGD", "CCND1", "SLC38A2", "TMEM50A") |
| 25 | HALLMARK_PROTEIN_SECRETION | 0.871526 | 0.983157 | 0.189823 | 0.651977 | 5894 | 13 | c("TMED2", "SEC24D", "ARF1", "CD63", "GNAS", "TMED10", "CLTA", "NAPA", "COPE", "RAB2A", "TMX1", "COPB1") |
| 26 | HALLMARK_INTERFERON_ALPHA_RESPONSE | 0.492519 | 0.983157 | 0.352959 | 0.976455 | 3159 | 7 | c("IFI30", "LY6E", "BST2") |
| 27 | HALLMARK_INTERFERON_GAMMA_RESPONSE | 0.824862 | 0.983157 | 0.213217 | 0.693672 | 5533 | 11 | c("IFI30", "LY6E", "BST2") |
| 28 | HALLMARK_APICAL_JUNCTION | 0.708749 | 0.983157 | 0.245063 | 0.797279 | 4754 | 11 | c("MDK", "GAMT", "PFN1") |
| 29 | HALLMARK_APICAL_SURFACE | 0.823398 | 0.983157 | 0.36537 | 0.734164 | 4806 | 3 | GATA3 |
| 30 | HALLMARK_COMPLEMENT | 0.564118 | 0.983157 | 0.272114 | 0.911751 | 3791 | 12 | c("GATA3", "CALM1", "ATOX1", "PFN1") |
| 31 | HALLMARK_UNFOLDED_PROTEIN_RESPONSE | 0.529902 | 0.983157 | 0.280168 | 0.938739 | 3561 | 12 | c("RPS14", "NPM1", "NHP2", "H2AX") |
| 32 | HALLMARK_PI3K_AKT_MTOR_SIGNALING | 0.683447 | 0.983157 | -0.2032 | -0.818171 | 2212 | 13 | c("VAV3", "CDK4", "ACTR3", "SQSTM1", "SFN", "PAK4", "RAC1", "CFL1", "YWHAB", "MAPK9", "ARF1", "UBE2N") |
| 33 | HALLMARK_E2F_TARGETS | 0.471705 | 0.983157 | -0.213307 | -0.97658 | 1391 | 18 | c("PNN", "ILF3", "CHEK2", "DDX39A", "CDK4", "MCM6", "RAN", "HMGA1", "NAA38", "XRCC6", "CENPM", "DCTPP1", "CKS2", "HMGB3", "TK1", "NAP1L1", "PRDX4", "H2AX") |
| 34 | HALLMARK_EPITHELIAL_MESENCHYMAL_TRANSITIO N | 0.706932 | 0.983157 | 0.238308 | 0.798482 | 4751 | 12 | c("PPIB", "LGALS1", "CAPG", "IGFBP2", "ITGB5", "TIMP1", "SAT1") |
| 35 | HALLMARK_INFLAMMATORY_RESPONSE | 0.493719 | 0.983157 | 0.372467 | 0.974245 | 3143 | 6 | c("LY6E", "BST2", "SLC7A2", "TIMP1") |
| 36 | HALLMARK_XENOBIOTIC_METABOLISM | 0.575253 | 0.983157 | 0.250909 | 0.908156 | 3974 | 15 | c("FBP1", "COMT", "TMBIM6", "BLVRB", "CYB5A", "DDAH2", "PYCR1", "ALDH9A1", "GSTO1") |
| 37 | HALLMARK_GLYCOLYSIS | 0.625663 | 0.983157 | 0.198188 | 0.865103 | 4602 | 26 | c("MIF", "CLDN3", "SDHC", "CYB5A", "SLC16A3", "TXN", "KDELR3", "ALDOA", "SRD5A3", "ALDH9A1", "B4GALT7", "NDUFV3", "SOX9", "AK4") |
| 38 | HALLMARK_REACTIVE_OXYGEN_SPECIES_PATHWAY | 0.493556 | 0.983157 | 0.335089 | 0.972996 | 3216 | 8 | c("ATOX1", "NDUFB4", "PRDX4", "TXN") |
| 39 | HALLMARK_UV_RESPONSE_UP | 0.393561 | 0.983157 | 0.274359 | 1.056598 | 2774 | 18 | c("RPN1", "TMBIM6", "H2AX", "CDC5L", "SULT1A1", "IGFBP2", "ALDOA", "ATP6V1F", "CREG1", "MRPL23", "SPR") |
| 40 | HALLMARK_ANGIOGENESIS | 0.761108 | 0.983157 | 0.616939 | 0.824048 | 3819 | 1 | TIMP1 |
| 41 | HALLMARK_HEME_METABOLISM | 0.697719 | 0.983157 | 0.247701 | 0.805861 | 4680 | 11 | c("UCP2", "BLVRB", "TCEA1", "HEBP1", "MGST3", "CTSB", "BCAM", "ADIPOR1", "LRP10", "GMPS") |
| 42 | HALLMARK_BILE_ACID_METABOLISM | 0.787077 | 0.983157 | 0.30476 | 0.745696 | 4908 | 5 | c("PRDX5", "PXMP2", "ALDH9A1") |
| 43 | HALLMARK_KRAS_SIGNALING_UP | 0.817095 | 0.983157 | 0.590953 | 0.789338 | 4100 | 1 | SOX9 |
| 44 | HALLMARK_MTORC1_SIGNALING | 0.951475 | 0.988045 | 0.124461 | 0.543278 | 6999 | 26 | c("IFI30", "RPN1", "CCNG1", "CANX", "DHCR7", "CYP51A1", "TCEA1") |
| 45 | HALLMARK_MYC_TARGETS_V2 | 0.916456 | 0.988045 | -0.263832 | -0.625253 | 3630 | 4 | c("SRM", "CDK4") |

| | | | | | | | | |
|---|---|---|---|---|---|---|---|---|
| 46 | HALLMARK_FATTY_ACID_METABOLISM | 0.984759 | 0.988045 | 0.107188 | 0.460077 | 7171 | 25 | c("LGALS1", "MIF", "ECI1", "SDHC", "OSTC", "LTC4S", "FH", "ALDOA", "ALDH9A1", "HSD17B10", "ACAA2", "S100A10") |
| 47 | HALLMARK_IL2_STAT5_SIGNALING | 0.986783 | 0.988045 | -0.139794 | -0.495542 | 3284 | 10 | c("IGF1R", "NCOA3") |
| 48 | HALLMARK_PANCREAS_BETA_CELLS | 0.988045 | 0.988045 | 0.507218 | 0.677493 | 4958 | 1 | SRP9 |
