## Supplementary Table 2 for "In silico reconstruction of primary and metastatic tumor architecture using GIS-augmented spatial transcriptomics"

**Supplementary Table 2. Summary of the characteristics of tumor samples collected for Visium analysis**

| Sample ID | DV200 | Dyslipidemia before Mets | Dyslipidemia after Mets | Age at Mets Dx | Living status | Survival after Mets Dx | Metastatic sites | ER Status | PR Status | HER2 Status | Treatment before Mets Dx | Treatment after Mets Dx |
|---|---|---|---|---|---|---|---|---|---|---|---|---|
| Patient1_Breast | 14 | No | No | 57 | alive | - | bone, lymph node, respiratory | Positive | Positive | Negative | Not available | Hormonal therapy (CDK4/6 inhibitor) |
| Patient2_Breast | 30 | No | No | NA | alive | - | bone | Positive | Positive | Negative | Not available | Hormonal therapy (AI, CDK4/6 inhibitors) |
| Patient3_Breast | 36 | Yes | No | 61 | alive | - | respiratory | Positive | Positive | Negative | Chemotherapy (Paclitaxel, Cyclophosphamide) + doxorubicin | Hormonal therapy (CDK4/6 inhibitor) |
| Patient4_Breast | 35 | Yes | No | 74 | alive | - | bone | Positive | Positive | Positive | Hormonal therapy (AI), Chemotherapy (Docetaxel, Cyclophosphamide) | Hormonal therapy (Fulvestrant, CDK4/6 inhibitors), Chemotherapy (Paclitaxel) (2nd line), Chemotherapy (Eribulin) (3rd line), HER2-directed treatment (Enhertu) (4th line), Hormonal therapy (Fulvestrant) and Chemotherapy (Fluorouracil) (5th line) |
| Patient5_Breast | 31 | Yes | No | NA | alive | - | bone | Positive | Positive | Negative | Not available | Hormonal therapy (AI, CDK4/6 inhibitors) |
| Patient6_Bone | 25 | No | No | 61 | - | 1390 days | bone, liver | Positive | Positive | Negative | Hormonal therapy (Tamoxifen), Chemotherapy (Docetaxel, Cyclophosphamide) | Hormonal therapy (Fulvestrant, CDK4/6 inhibitors) |
| Patient7_Bone | 33 | No | No | 82 | - | 1317 days | bone | Positive | Positive | Negative | Hormonal therapy (Fulvestrant, CDK4/6 inhibitors), Chemotherapy (Docetaxel, Cyclophosphamide) | Hormonal therapy (Fulvestrant, CDK4/6 inhibitors) |
| Patient8_Bone | 36 | Yes | Yes | 75 | alive | - | bone | Positive | Positive | Negative | Not available | Hormonal therapy (Fulvestrant, CDK4/6 inhibitors) |
| Patient9_Bone | 25 | Yes | No | 61 | alive | - | bone | Positive | Positive | Negative | Hormonal therapy (Tamoxifen, AI), Chemotherapy (Paclitaxel, Cyclophosphamide) + doxorubicin | Hormonal therapy (Fulvestrant, CDK4/6 inhibitors) |
| Patient4_Bone | 24 | Yes | No | 74 | alive | - | bone | Positive | Positive | Positive | Hormonal therapy (AI), Chemotherapy (Docetaxel, Cyclophosphamide) | Hormonal therapy (Fulvestrant, CDK4/6 inhibitors), Chemotherapy (Paclitaxel) (2nd line), Chemotherapy (Eribulin) (3rd line), HER2-directed treatment (Enhertu) (4th line), Hormonal therapy (Fulvestrant) and Chemotherapy (Fluorouracil) (5th line) |
| Patient1_Bone | 21 | No | No | 57 | alive | - | bone, lymph node, respiratory | Positive | Positive | Negative | Not available | Hormonal therapy (CDK4/6 inhibitor) |
| Patient5_Bone | 40 | Yes | No | NA | alive | - | bone | Positive | Positive | Negative | Not available | Hormonal therapy (AI, CDK4/6 inhibitors) |

* Matching pairs of primary and metastatic bone samples are color-coded
