## Supplementary Table 3 for "In silico reconstruction of primary and metastatic tumor architecture using GIS-augmented spatial transcriptomics"

**Supplementary Table 3. Gene set enrichment analysis (GSEA) results on shared nearest neighbor (SNN) clusters from primary breast and metastatic tumors**

| Cluster 0 | pathway | pval | padj | ES | NES | nMoreExtreme | size | leadingEdge |
|---|---|---|---|---|---|---|---|---|
| 1 | HALLMARK_ESTROGEN_RESPONSE_EARLY | 0.000239 | 0.001707 | 0.539991 | 2.949784 | 0 | 100 | c("HSPB8", "IL17RB", "PDZK1", "MAPT", "GREB1", "RHOBTB3", "RAPGEFL1", "SLC9A3R1", "PGR", "ADCY1", "KCNK15", "SCNN1A", "SEC14L2", "PAPSS2", "ELOVL2", "ABCA3", "SEMA3B", "STC2", "RARA", "KDM4B", "FRK", "CA12", "RASGRP1", "AMFR", "GFRA1", "SLC7A2", "ESRP2", "PDLIM3", "GAB2", "PRSS23", "DHRS2", "FAM102A", "FASN", "KRT8", "ITPK1", "MED13L", "TOB1", "FHL2", "WFS1", "RHOD", "AKAP1", "CANT1", "FLNB", "CELSR1", "KRT19", "THSD4", "OVOL2", "PTGES", "SLC7A5", "DYNLT3", "SYNGR1", "ASB13", "BHLHE40", "NRIP1", "MYC") |
| 2 | HALLMARK_ESTROGEN_RESPONSE_LATE | 0.000237 | 0.001707 | 0.384764 | 2.067985 | 0 | 92 | c("SERPINA1", "HSPB8", "IL17RB", "PDZK1", "SERPINA3", "MAPT", "CPE", "RAPGEFL1", "SLC9A3R1", "PGR", "SCNN1A", "PAPSS2", "CDH1", "LSR", "ABCA3", "SEMA3B", "FRK", "CA12", "SERPINA5", "DLG5", "AMFR", "LLGL2", "CD9", "PDLIM3", "PRSS23", "DHRS2", "DCXR", "FAM102A", "ITPK1", "FGFR3", "TOB1", "COX6C", "WFS1", "FLNB") |
| 3 | HALLMARK_COMPLEMENT | 0.000186 | 0.001707 | -0.572451 | -2.556239 | 0 | 55 | c("S100A9", "APOC1", "CTSS", "C1QA", "PIM1", "PLAT", "C3", "LTF", "FCER1G", "CTSD", "CTSL", "LGMN", "C1QC", "C1S", "LIPA", "EHD1", "SERPING1", "C1R", "CTSB", "PLAUR", "LAP3", "CD36", "LRP1", "PRKCD", "CALM3", "PFN1", "GNAI3") |
| 4 | HALLMARK_EPITHELIAL_MESENCHYMAL_TRANSITION | 0.000175 | 0.001707 | -0.477908 | -2.354987 | 0 | 87 | c("FUCA1", "ELN", "FBLN2", "DAB2", "JUN", "GPC1", "WIPF1", "CAPG", "DCN", "IL32", "FBLN1", "EMP3", "IGFBP2", "PLAUR", "NOTCH2", "LRP1", "SFRP4", "SAT1", "TPM4", "VIM", "MXRA5", "PFN2", "TPM1", "LOXL1", "CALD1", "PDGFRB", "MMP2", "LAMC1", "PCOLCE", "DPYSL3", "DST", "IGFBP3", "MYL9", "VCAN", "FSTL1", "ACTA2", "LUM", "THY1", "ITGB5", "CDH11", "FBN1", "COL16A1", "COL4A2", "RHOB", "HTRA1", "TPM2", "SPP1", "QSOX1", "FAP") |
| 5 | HALLMARK_HEME_METABOLISM | 0.000182 | 0.001707 | -0.6079 | -2.869773 | 0 | 72 | c("HBB", "HBD", "SLC4A1", "SNCA", "AHSP", "CAT", "SLC25A37", "MPP1", "ACP5", "GYPC", "C3", "UCP2", "MAP2K3", "UBAC1", "NR3C1", "TRAK2", "CIR1", "BACH1", "FOXO3", "UROD", "TFRC", "CCND3", "RNF19A", "CTSB", "TNS1") |
| 6 | HALLMARK_ALLOGRAFT_REJECTION | 0.0002 | 0.001707 | -0.592309 | -2.282651 | 0 | 30 | c("CD74", "CTSS", "STAB1", "SPI1", "HCLS1", "CD4", "ITGB2", "CAPG", "IFNGR2", "CCND3", "STAT1", "ETS1", "SRGN", "TAPBP", "PSMB10", "B2M", "MMP9") |
| 7 | HALLMARK_KRAS_SIGNALING_UP | 0.00019 | 0.001707 | -0.562467 | -2.407205 | 0 | 46 | c("FUCA1", "CTSS", "PLAT", "GYPC", "FCER1G", "IKZF1", "APOD", "GPNMB", "GLRX", "CXCR4", "ITGB2", "NIN", "SPARCL1", "PECAM1", "LAPTM5", "ITGBL1", "ENG", "ETS1", "ANKH", "LCP1", "PLAUR", "WDR33", "MAFB") |
| 8 | HALLMARK_INTERFERON_GAMMA_RESPONSE | 0.00076 | 0.004223 | -0.502618 | -2.117459 | 3 | 43 | c("CD74", "TNFAIP2", "PIM1", "IFI30", "IFI27", "SAMHD1", "TXNIP", "C1S", "SERPING1", "STAT1", "C1R", "TAPBP", "LAP3", "ARID5B", "EIF4E3", "PSMB10", "MYD88", "B2M", "SOD2", "NFKBIA", "STAT2") |
| 9 | HALLMARK_COAGULATION | 0.000752 | 0.004223 | -0.477273 | -2.069693 | 3 | 48 | c("S100A1", "APOC1", "C1QA", "PLAT", "C3", "CFD", "LGMN", "C1S", "CAPN2", "GSN", "A2M", "PECAM1", "SERPING1", "ANXA1", "C1R", "CTSB") |
| 10 | HALLMARK_APOPTOSIS | 0.002018 | 0.009171 | -0.430184 | -1.962174 | 10 | 60 | c("HMOX1", "GPX3", "CD14", "PLAT", "JUN", "GSR", "CFLAR", "TXNIP", "GSN", "DCN", "DPYD", "ANXA1", "ANKH", "IGF2R", "SLC20A1", "SAT1", "BNIP3L", "BTG2", "PDGFRB", "MMP2", "SOD2") |
| 11 | HALLMARK_P53_PATHWAY | 0.001858 | 0.009171 | -0.432576 | -1.931637 | 9 | 55 | c("HMOX1", "FUCA1", "IFI30", "ABCC5", "JUN", "S100A4", "CTSD", "RAP2B", "TXNIP", "RXRA", "FOXO3", "STOM", "CCND3", "GM2A") |
| 12 | HALLMARK_REACTIVE_OXYGEN_SPECIES_PATHWAY | 0.002218 | 0.009241 | -0.54687 | -2.021009 | 10 | 25 | c("MPO", "GPX3", "CAT", "LSP1", "GLRX", "GSR", "TXNRD1") |
| 13 | HALLMARK_HYPOXIA | 0.0033 | 0.012691 | -0.403595 | -1.872157 | 17 | 66 | c("DTNA", "HMOX1", "PIM1", "BCL2", "JUN", "S100A4", "GPC1", "GLRX", "CXCR4", "NR3C1", "FOXO3", "DCN", "ETS1", "SDC3", "CAVIN1", "PLAUR", "PNRC1", "WSB1", "BNIP3L", "ZFP36", "NEDD4L", "IGFBP3", "FOS", "HK1", "HEXA", "PAM", "IDS", "SDC2", "SLC2A1", "FBP1", "CITED2") |
| 14 | HALLMARK_MYC_TARGETS_V1 | 0.004088 | 0.0146 | 0.339167 | 1.746669 | 17 | 76 | c("NME1", "CLNS1A", "TOMM70", "GOT2", "HSP90AB1", "IMPDH2", "PPIA", "LSM7", "CBX3", "HNRNPC", "GNL3", "CANX", "VDAC3", "XPOT", "CCT3", "ETF1", "ERH", "ILF2", "APEX1", "HNRNPU", "PSMD3", "MYC", "SNRPA", "EIF4E", "PSMC4", "SF3B3", "SNRPD2", "NOP16", "PRPF31", "TUFM", "VDAC1", "PSMA2", "PRDX4", "POLD2", "EXOSC7", "EIF3B", "NHP2", "HPRT1", "UBA2", "RSL1D1", "CYC1", "UBE2E1", "RNPS1", "SRSF1", "HNRNPD", "SSBP1", "EIF2S1", "NDUFAB1", "SMARCC1", "CUL1", "PRPS2", "HSPD1") |

| | | | | | | | | |
|---|---|---|---|---|---|---|---|---|
| 15 | HALLMARK_TNFA_SIGNALING_VIA_NFKB | 0.006765 | 0.022551 | -0.376127 | -1.743028 | 36 | 65 | c("TNFAIP2", "JUN", "KLF9", "PLPP3", "MAP2K3", "CFLAR", "B4GALT5", "DUSP4", "EHD1", "MARCKS", "IFNGR2", "FOSB", "PLAUR", "SMAD3", "SGK1", "PNRC1", "KLF2", "SAT1", "ZFP36", "BTG2", "LITAF", "NINJ1", "IER5", "SOD2", "NFKBIA", "EGR1", "FOS") |
| 16 | HALLMARK_OXIDATIVE_PHOSPHORYLATION | 0.00791 | 0.024718 | 0.292598 | 1.585476 | 32 | 97 | c("PHYH", "ATP1B1", "NDUFC2", "NDUFC1", "COX6A1", "TOMM70", "GOT2", "FH", "COX6C", "NDUFA4", "COX7C", "NDUFB1", "NDUFA5", "TIMM9", "COX4I1", "NQO2", "VDAC3", "COX7B", "ATP5F1C", "UQCR11", "GRPEL1", "SLC25A4", "NDUFA7", "UQCRC1", "ATP5PD", "ATP5ME", "NDUFA2", "UQCRB", "MGST3", "HSD17B10", "TIMM13", "ALDH6A1", "MDH2", "NDUFB2", "SDHA", "NDUFA3", "NDUFS7", "ATP6V1F", "ATP5F1B", "ATP5MF", "IDH2", "VDAC1", "NDUFS4", "ATP5MC2", "ATP6V1C1", "UQCRQ", "NDUFA1", "NDUFB7", "AIFM1", "UQCRC2", "UQCR10", "ATP6V0E1", "ECI1", "ATP6V1E1", "CYC1", "OXA1L", "COX6B1", "IDH3B", "TIMM50", "TIMM17A", "OGDH", "COX17") |
| 17 | HALLMARK_UNFOLDED_PROTEIN_RESPONSE | 0.018143 | 0.053363 | 0.361247 | 1.614515 | 85 | 43 | c("STC2", "LSM1", "EIF4EBP1", "EIF4A3", "WFS1", "EIF4G1", "BAG3", "XPOT", "SLC7A5", "EDC4", "SPCS1", "EIF4E", "ATF6", "KIF5B", "NOP14", "XBP1", "SEC31A", "VEGFA", "KHSRP", "YWHAZ", "ATF4", "SERP1", "ATP6V0D1", "NHP2", "SRPRB", "GOSR2") |
| 18 | HALLMARK_IL6_JAK_STAT3_SIGNALING | 0.020565 | 0.057124 | -0.461589 | -1.705847 | 101 | 25 | c("HMOX1", "CD14", "PIM1", "JUN", "IL1R1", "A2M", "IFNGR2", "STAT1") |
| 19 | HALLMARK_IL2_STAT5_SIGNALING | 0.026599 | 0.069997 | -0.368697 | -1.588603 | 140 | 47 | c("S100A1", "PIM1", "BCL2", "CTSZ", "NFKBIZ", "CAPG", "CCND3", "ENPP1", "LRIG1", "IGF2R", "AHR", "ANXA4", "SNX14", "PLEC", "BMPR2", "AHNAK", "CD81", "RHOB", "EMP1", "RNH1", "SLC1A5", "SPP1") |
| 20 | HALLMARK_INTERFERON_ALPHA_RESPONSE | 0.041002 | 0.097624 | -0.455687 | -1.602962 | 197 | 21 | c("CD74", "IFI30", "IFI27", "TXNIP", "C1S") |
| 21 | HALLMARK_UV_RESPONSE_DN | 0.039141 | 0.097624 | -0.360531 | -1.527186 | 205 | 44 | c("ADD3", "DAB2", "PLPP3", "AKT3", "NR3C1", "TGFBR2", "RXRA", "SMAD3", "PRDM2", "ATP2B4", "NOTCH2", "ANXA4", "RBPMS") |
| 22 | HALLMARK_INFLAMMATORY_RESPONSE | 0.07458 | 0.169499 | -0.370107 | -1.44857 | 376 | 32 | c("CYBB", "STAB1", "CD14", "IL1R1", "IFNGR2", "EMP3", "TAPBP", "PLAUR", "AHR", "GNAI3", "BTG2") |
| 23 | HALLMARK_ADIPOGENESIS | 0.098966 | 0.215143 | -0.28034 | -1.342516 | 554 | 75 | c("ALDH2", "APOE", "GPX3", "RAB34", "CAT", "C3", "UCP2", "FABP4", "SPARCL1", "STOM", "VEGFB", "TALDO1", "CAVIN1", "LAMA4", "CD36", "PHLDB1", "SCP2") |
| 24 | HALLMARK_MITOTIC_SPINDLE | 0.112924 | 0.235258 | -0.282667 | -1.315848 | 615 | 67 | c("SHROOM1", "ABR", "CNTRL", "NIN", "RAB3GAP1", "PCM1", "GSN", "MARCKS", "EPB41L2", "TRIO", "CKAP5", "NOTCH2", "WASF2", "ABL1", "VCL", "ARHGEF2", "ARHGAP29") |
| 25 | HALLMARK_NOTCH_SIGNALING | 0.14502 | 0.290039 | -0.501492 | -1.361327 | 660 | 9 | c("ARRB1", "LFNG", "NOTCH2") |
| 26 | HALLMARK_MTORC1_SIGNALING | 0.155993 | 0.299987 | -0.264648 | -1.249352 | 854 | 72 | c("CORO1A", "ADD3", "IFI30", "LGMN", "GLRX", "MAP2K3", "GSR", "CXCR4", "ITGB2", "TXNRD1", "TFRC") |
| 27 | HALLMARK_CHOLESTEROL_HOMEOSTASIS | 0.176517 | 0.326884 | 0.326087 | 1.252563 | 889 | 25 | c("SEMA3B", "LSS", "CD9", "CYP51A1", "FASN", "ALCAM", "TRIB3", "ETHE1", "CTNNB1", "PMVK", "GUSB", "ACTG1") |
| 28 | HALLMARK_XENOBIOTIC_METABOLISM | 0.256648 | 0.458301 | -0.248054 | -1.146689 | 1408 | 64 | c("HMOX1", "ALDH2", "APOE", "CAT", "GSR", "IL1R1", "CNDP2", "FBLN1", "PGD", "PDK4", "CD36") |
| 29 | HALLMARK_MYOGENESIS | 0.270681 | 0.466691 | -0.258758 | -1.142909 | 1442 | 52 | c("DTNA", "GPX3", "SORBS3", "LSP1", "CFD", "APOD", "GSN") |
| 30 | HALLMARK_KRAS_SIGNALING_DN | 0.292358 | 0.487264 | 0.304691 | 1.13315 | 1514 | 22 | c("LYPD3", "GPRC5C", "HSD11B2", "IDUA", "SKIL", "FGFR3", "CCDC106", "CLSTN3", "NR4A2", "SLC25A23", "EFHD1") |
| 31 | HALLMARK_WNT_BETA_CATENIN_SIGNALING | 0.35666 | 0.565304 | 0.44001 | 1.092878 | 1951 | 7 | c("CTNNB1", "MYC", "NCOR2", "CSNK1E", "CUL1", "NCSTN") |
| 32 | HALLMARK_TGF_BETA_SIGNALING | 0.367443 | 0.565304 | -0.294015 | -1.073232 | 1798 | 24 | c("CDK9", "IFNGR2", "LTBP2", "ENG", "SMAD3", "SKI", "SLC20A1", "BMPR2") |
| 33 | HALLMARK_UV_RESPONSE_UP | 0.373101 | 0.565304 | -0.239697 | -1.058718 | 1988 | 52 | c("HMOX1", "GPX3", "ARRB2", "UROD", "TFRC", "CCND3", "FOSB", "IGFBP2", "MGAT1", "PRKCD", "BTG2", "SOD2", "NFKBIA", "FOS", "NXF1", "RHOB") |
| 34 | HALLMARK_PANCREAS_BETA_CELLS | 0.420573 | 0.61849 | -0.431044 | -1.024612 | 1892 | 6 | c("AKT3", "MAFB") |
| 35 | HALLMARK_HEDGEHOG_SIGNALING | 0.435624 | 0.622319 | 0.361512 | 1.023656 | 2364 | 10 | c("ADGRG1", "CELSR1", "TLE3", "NRP1", "ETS2", "VEGFA") |
| 36 | HALLMARK_ANDROGEN_RESPONSE | 0.574387 | 0.717984 | -0.218218 | -0.924357 | 3022 | 44 | c("BMPR1B", "GSR", "CCND3", "ANKH", "SGK1", "ARID5B", "SAT1", "TMEM50A", "DBI", "B2M", "GNAI3") |
| 37 | HALLMARK_APICAL_JUNCTION | 0.560989 | 0.717984 | -0.198693 | -0.93568 | 3085 | 71 | c("SORBS3", "AKT3", "PECAM1", "TSPAN4", "EPB41L2", "SDC3", "CNN2", "RAC2", "ADAM15", "VCL", "CD99", "PFN1", "MMP9", "PARVA", "MMP2", "MDK", "CD276", "CTNND1", "MYL9", "EVL", "VCAN", "THY1", "CDH11", "FBN1", "COL16A1") |
| 38 | HALLMARK_PI3K_AKT_MTOR_SIGNALING | 0.517294 | 0.717984 | -0.227424 | -0.958104 | 2721 | 43 | c("ITPR2", "MAP2K3", "CXCR4", "GRK2", "ACTR3", "CAB39", "PFN1", "MYD88", "SMAD2", "STAT2", "CFL1") |
| 39 | HALLMARK_MYC_TARGETS_V2 | 0.560158 | 0.717984 | 0.28125 | 0.92181 | 2974 | 15 | c("CBX3", "GNL3", "UNG", "MYC", "NOC4L", "NOP16", "SORD", "FARSA", "HSPD1", "DCTPP1", "TBRG4") |

| | | | | | | | | |
|---|---|---|---|---|---|---|---|---|
| 40 | HALLMARK_GLYCOLYSIS | 0.547677 | 0.717984 | 0.192307 | 0.949307 | 2486 | 63 | c("ARTN", "CACNA1H", "IDUA", "STC2", "NOL3", "GOT2", "PPIA", "FUT8", "ALDH7A1", "AGL", "MIF", "CLDN3", "PFKP") |
| 41 | HALLMARK_PROTEIN_SECRETION | 0.64816 | 0.771619 | 0.192601 | 0.88121 | 3046 | 47 | c("TPD52", "TOM1L1", "TMED2", "AP1G1", "RAB2A", "ARF1", "AP2S1", "COPB2", "SEC31A", "ARFIP1", "USO1", "CLCN3", "TMED10", "SCAMP3", "RAB22A", "LAMP2", "RAB5A", "STAM", "VPS45", "GOSR2", "ARFGEF2", "NAPA", "SOD1", "COPE", "AP2M1", "ERGIC3", "OCRL", "CLTC") |
| 42 | HALLMARK_APICAL_SURFACE | 0.636127 | 0.771619 | 0.296682 | 0.870131 | 3422 | 11 | c("LYPD3", "SULF2", "EPHB4", "HSPB1") |
| 43 | HALLMARK_E2F_TARGETS | 0.70479 | 0.819523 | 0.189522 | 0.851841 | 3339 | 44 | c("NME1", "JPT1", "CDKN2A", "NUDT21", "TK1", "POP7", "UNG", "MYC", "SNRPB", "CTCF", "MYBL2", "PRDX4", "POLD2", "RAD50", "ILF3", "TUBB", "SLBP", "TMPO", "SRSF1", "DDX39A", "HNRNPD", "EIF2S1", "PRKDC", "DCTPP1", "NBN", "TBRG4", "MCM3", "HMGB2") |
| 44 | HALLMARK_SPERMATOGENESIS | 0.731718 | 0.831498 | -0.267448 | -0.773435 | 3381 | 11 | c("SLC12A2", "TALDO1") |
| 45 | HALLMARK_ANGIOGENESIS | 0.780967 | 0.867741 | 0.238447 | 0.760766 | 4135 | 14 | c("SERPINA5", "FGFR1", "STC1", "NRP1", "VEGFA") |
| 46 | HALLMARK_DNA_REPAIR | 0.821891 | 0.874352 | 0.166506 | 0.784912 | 3806 | 53 | c("NME1", "IMPDH2", "NUDT21", "CANT1", "TMED2", "POLR2H", "NT5C", "DAD1", "BRF2", "GMPR2", "NME4", "TAF1C", "POLR2C", "POLR2I", "APRT", "SDCBP", "GTF2F1", "POLR3C", "SUPT5H", "RAE1", "POLB", "BCAM", "HPRT1", "SEC61A1") |
| 47 | HALLMARK_BILE_ACID_METABOLISM | 0.81246 | 0.874352 | -0.222262 | -0.718457 | 3820 | 16 | c("CAT", "RXRA") |
| 48 | HALLMARK_FATTY_ACID_METABOLISM | 0.883516 | 0.901546 | 0.160691 | 0.712657 | 4201 | 42 | c("RAP1GDS1", "FASN", "PRDX6", "FH", "MIF", "TDO2", "UGDH", "APEX1", "GCDH", "HSD17B10", "MDH2", "SDHA", "HSP90AA1", "ELOVL5") |
| 49 | HALLMARK_PEROXISOME | 0.880948 | 0.901546 | -0.177163 | -0.672266 | 4387 | 28 | c("CAT", "ABCC5") |
| 50 | HALLMARK_G2M_CHECKPOINT | 0.989541 | 0.989541 | 0.117321 | 0.539652 | 4635 | 48 | c("JPT1", "UCK2", "BCL3", "TLE3", "SLC7A5", "MNAT1", "NUMA1", "HNRNPU", "MYC", "KIF5B", "CTCF", "CHMP1A", "SLC38A1", "MYBL2", "TNPO2", "ILF3", "ATF5", "UBE2C", "TMPO", "LIG3", "SRSF1", "DDX39A", "HNRNPD", "SMARCC1", "CUL1", "CCND1") |

| Cluster 1 | pathway | pval | padj | ES | NES | nMoreExtreme | size | leadingEdge |
|---|---|---|---|---|---|---|---|---|
| 1 | HALLMARK_EPITHELIAL_MESENCHYMAL_TRANSITION | 0.000125 | 0.006249 | 0.431919 | 2.052476 | 0 | 126 | c("AREG", "MATN3", "ADAM12", "ACTA2", "CDH11", "LOXL1", "CADM1", "FAP", "PDLIM4", "MFAP5", "VCAN", "SDC1", "IGFBP3", "TPM4", "MXRA5", "HTRA1", "PFN2", "TAGLN", "COL12A1", "THY1", "JUN", "THBS2", "TNFRSF12A", "DPYSL3", "CTHRC1", "NID2", "COL16A1", "COL5A2", "FBN1", "POSTN", "CCN2", "COL5A1", "ELN", "MYL9", "SNAI2", "LOX", "MMP14", "INHBA", "CALD1", "TPM2", "NNMT", "LAMA2", "GADD45B", "SFRP4", "RHOB", "COL8A2", "LRRC15", "ITGB5", "SERPINH1", "COL11A1", "BMP1", "LAMC1", "MGP", "PRRX1", "FSTL3", "MMP2", "COMP", "EFEMP2", "EDIL3", "CAP2", "CD44", "LUM", "BASP1", "COL3A1", "GADD45A", "SLIT3", "LRP1", "GAS1", "FBLN1", "CCN1", "TNFAIP3", "ECM2", "PLOD2", "WIPF1", "TIMP1", "DCN", "COL6A3", "MYLK", "PDGFRB", "FLNA", "PMEPA1", "COPA", "BGN", "COL6A2", "FBLN2") |
| 2 | HALLMARK_ESTROGEN_RESPONSE_EARLY | 0.000896 | 0.008965 | -0.394105 | -2.027525 | 1 | 90 | c("GFRA1", "ADCY1", "RET", "BLVRB", "RHOBTB3", "SEC14L2", "GAB2", "ABHD2", "SIAH2", "CCND1", "SCNN1A", "BAG1", "SLC39A6", "FAM102A", "MAST4", "SLC2A1", "KDM4B", "ITPK1", "RHOD", "PRSS23", "HSPB8", "AMFR", "MED13L", "FLNB") |
| 3 | HALLMARK_ESTROGEN_RESPONSE_LATE | 0.000893 | 0.008965 | -0.402596 | -2.063829 | 1 | 88 | c("SERPINA1", "LTF", "RET", "BLVRB", "COX6C", "CDH1", "SERPINA3", "ABHD2", "SIAH2", "CCND1", "SCNN1A", "BAG1", "SLC29A1", "FAM102A", "CD9", "ITPK1", "PRSS23", "SCUBE2", "HSPB8", "AMFR", "FLNB", "DNAJC1", "LSR", "AGR2", "TOP2A", "SLC9A3R1", "DCXR", "CPE", "PLAAT3", "SEMA3B") |
| 4 | HALLMARK_MTORC1_SIGNALING | 0.000774 | 0.008965 | -0.398796 | -1.869216 | 1 | 57 | c("SCD", "GGA2", "CORO1A", "SQSTM1", "IFI30", "SLC2A1", "CD9", "SYTL2", "TFRC", "ITGB2", "SERP1", "LTA4H", "GSK3B", "TXNRD1", "M6PR", "ACACA", "SLC9A3R1", "CALR", "P4HA1", "LGMN", "GSR", "NUPR1", "PIK3R3") |
| 5 | HALLMARK_HEME_METABOLISM | 0.000397 | 0.008965 | -0.570801 | -2.716038 | 0 | 61 | c("HBB", "HBD", "SLC4A1", "BLVRB", "SNCA", "ACP5", "FBXO7", "GYPC", "RAD23A", "SLC2A1", "MPP1", "TFRC", "SLC25A37", "PSMD9", "USP15", "FBXO9", "HAGH", "PRDX2", "TRAK2", "CCND3", "NCOA4", "UBAC1", "BSG", "CTSB", "HEBP1", "KHNYN") |
| 6 | HALLMARK_ANDROGEN_RESPONSE | 0.007715 | 0.064291 | -0.407911 | -1.765654 | 20 | 41 | c("STEAP4", "SCD", "ABHD2", "CCND1", "BMPR1B", "SAT1", "INPP4B", "CCND3", "NCOA4", "AKT1", "SELENOP", "GSR", "ELL2", "SPDEF", "MAF", "B2M") |
| 7 | HALLMARK_INFLAMMATORY_RESPONSE | 0.013599 | 0.097135 | 0.430019 | 1.657408 | 98 | 41 | c("PDE4B", "EREG", "NFKBIA", "RGS1", "BTG2", "OLR1", "PCDH7", "PDPN", "AHR", "ITGB8", "MMP14", "INHBA", "DCBLD2", "SLC7A2", "AXL") |
| 8 | HALLMARK_COMPLEMENT | 0.018692 | 0.116822 | -0.335229 | -1.578776 | 47 | 58 | c("SERPINA1", "LTF", "PLAT", "CTSS", "CTSL", "USP15", "LTA4H", "PRKCD", "CTSD", "HSPA1A", "LGALS3", "LIPA", "PLAUR", "LGMN", "CTSB", "PFN1", "FCER1G", "S100A9", "GRB2", "GNAI2", "PIM1") |
| 9 | HALLMARK_OXIDATIVE_PHOSPHORYLATION | 0.021568 | 0.119821 | -0.309383 | -1.540823 | 51 | 75 | c("COX6C", "NDUFC2", "COX6B1", "COX8A", "TCIRG1", "NDUFB2", "NDUFC1", "ATP5MF", "ATP6V0B", "UQCRC1", "COX4I1", "BAX", "COX7C", "NDUFB6", "NDUFA1", "ACAA1", "ATP6V0C", "ATP6V1F", "NDUFS7", "ATP5F1B", "UQCR11", "ETFB", "NDUFS6", "NDUFA4", "ATP6AP1", "MGST3", "NDUFA3", "FH", "ECHS1", "ATP5MC2", "NDUFV1", "COX6A1", "NDUFB5", "OXA1L") |
| 10 | HALLMARK_PI3K_AKT_MTOR_SIGNALING | 0.030668 | 0.139401 | -0.366325 | -1.575536 | 83 | 40 | c("SQSTM1", "SLC2A1", "ARPC3", "GRK2", "GSK3B", "PPP1CA", "ACACA", "CALR", "MYD88", "AKT1", "PIK3R3", "DUSP3", "PFN1", "MKNK2", "GRB2") |
| 11 | HALLMARK_UV_RESPONSE_UP | 0.028947 | 0.139401 | -0.346884 | -1.552766 | 76 | 47 | c("RET", "HMOX1", "SQSTM1", "HNRNPU", "FOSB", "TFRC", "NR4A1", "GRINA", "SOD2", "ARRB2", "PRKCD", "CCND3", "BSG", "BTG1", "ACAA1", "ATP6V1F", "POLR2H", "PPT1", "GPX3", "UROD", "CLTB") |
| 12 | HALLMARK_REACTIVE_OXYGEN_SPECIES_PATHWAY | 0.04233 | 0.176374 | -0.470974 | -1.598933 | 132 | 17 | c("MPO", "SOD2", "TXNRD1", "PRDX2", "FTL", "GSR", "GPX3") |
| 13 | HALLMARK_GLYCOLYSIS | 0.086816 | 0.333909 | 0.325191 | 1.349547 | 645 | 59 | c("EGLN3", "HOMER1", "CITED2", "SOX9", "TFF3", "VCAN", "ELF3", "SDC1", "IGFBP3", "GPC4", "COL5A1", "PAM", "TPST1", "CHPF", "IER3", "B4GALT2", "KIF2A", "CD44", "STC1", "EXT1", "DDIT4", "NDUFV3", "CYB5A", "ALDH9A1", "PLOD2", "SDC2", "PMM2", "DCN", "IL13RA1") |
| 14 | HALLMARK_XENOBIOTIC_METABOLISM | 0.098061 | 0.341407 | -0.299754 | -1.336767 | 262 | 46 | c("BLVRB", "HMOX1", "APOE", "COMT", "DHRS7", "PSMB10", "FBP1", "DCXR", "BCAR1", "GSR", "MCCC2", "PGD", "SPINT2") |
| 15 | HALLMARK_ALLOGRAFT_REJECTION | 0.102422 | 0.341407 | -0.338285 | -1.360602 | 295 | 31 | c("MMP9", "CTSS", "HCLS1", "ITGB2", "SPI1", "BCL3", "PSMB10", "SRGN", "TAPBP", "CCND3", "AKT1", "STAB1", "CD47", "B2M") |
| 16 | HALLMARK_IL6_JAK_STAT3_SIGNALING | 0.184874 | 0.577731 | -0.325907 | -1.245561 | 549 | 25 | c("HMOX1", "MAP3K8", "CD9", "CD14", "MYD88", "LTBR", "SOCS3", "GRB2", "TYK2", "PIM1") |

| | | | | | | | | |
|---|---|---|---|---|---|---|---|---|
| 17 | HALLMARK_CHOLESTEROL_HOMEOSTASIS | 0.201345 | 0.59219 | -0.321211 | -1.227614 | 598 | 25 | c("SCD", "ALCAM", "CD9", "CTNNB1", "LGALS3", "PLAUR", "LGMN", "SEMA3B") |
| 18 | HALLMARK_UV_RESPONSE_DN | 0.241758 | 0.65597 | 0.273374 | 1.164038 | 1825 | 66 | c("AKT3", "NFIB", "RBPMS", "CITED2", "ERBB2", "DDAH1", "TOGARAM1", "CDC42BPA", "ATXN1", "DLC1", "COL5A2", "ATP2B4", "SNAI2", "RUNX1", "DMAC2L", "ANXA4", "COL11A1", "LAMC1", "SRI", "IGFBP5", "CAP2", "COL3A1", "CCN1", "SDC2", "PDGFRB", "FHL2", "SCAF8", "LPAR1", "BHLHE40", "PHF3", "APBB2", "COL1A1", "COL1A2", "ABCC1", "MIOS", "MGMT") |
| 19 | HALLMARK_BILE_ACID_METABOLISM | 0.249269 | 0.65597 | 0.392879 | 1.193512 | 1703 | 16 | c("DIO2", "EFHC1", "PEX11A", "ATXN1", "PEX19") |
| 20 | HALLMARK_MITOTIC_SPINDLE | 0.336727 | 0.701514 | 0.250789 | 1.084769 | 2546 | 72 | c("LLGL1", "PCM1", "PALLD", "KIFAP3", "CDC42BPA", "ARHGAP29", "TRIO", "NIN", "BCL2L11", "KIF3B", "EPB41L2", "RASAL2", "FARP1", "TIAM1", "ARHGEF12", "ARHGEF2", "MYO1E", "NCK2", "CDK5RAP2", "ALMS1", "SHROOM1", "MYH10", "SYNPO", "ABL1", "SOS1", "ARHGEF11", "ARF6", "EZR", "MYH9") |
| 21 | HALLMARK_TGF_BETA_SIGNALING | 0.324361 | 0.701514 | -0.279111 | -1.095139 | 963 | 28 | c("CDH1", "CTNNB1", "PPM1A", "FNTA", "PPP1CA") |
| 22 | HALLMARK_INTERFERON_ALPHA_RESPONSE | 0.322001 | 0.701514 | -0.318122 | -1.115226 | 984 | 19 | c("IFITM2", "IFI30", "PSME1", "LY6E", "CD47", "TRIM25", "B2M") |
| 23 | HALLMARK_APICAL_JUNCTION | 0.281701 | 0.701514 | 0.261678 | 1.128419 | 2125 | 71 | c("AKT3", "CDH11", "NECTIN4", "VCAN", "THY1", "PTK2", "MDK", "COL16A1", "FBN1", "MYL9", "NLGN2", "CLDN4", "EPB41L2", "CNN2", "CTNND1", "PARVA", "NECTIN2", "WNK4", "BMP1", "THBS3", "ACTG1", "MYH10", "MMP2", "RSU1", "SHC1", "MAP3K20", "EVL", "MYH9", "ADAM15", "PCDH1", "VCL", "LIMA1", "TSPAN4", "JUP", "TIAL1", "ACTB", "WASL", "INPPL1", "YWHAH", "CERCAM", "TAOK2") |
| 24 | HALLMARK_P53_PATHWAY | 0.299755 | 0.701514 | -0.226111 | -1.099016 | 734 | 68 | c("TM4SF1", "HMOX1", "STOM", "RAP2B", "IFI30", "FUCA1", "SAT1", "CTSD", "S100A4", "ABCC5", "CCND3", "DCXR", "IER5", "BAX", "BTG1", "NUPR1", "RXRA", "MKNK2", "MDM2") |
| 25 | HALLMARK_KRAS_SIGNALING_DN | 0.36706 | 0.734119 | -0.296182 | -1.072367 | 1120 | 21 | c("CPB1", "GPRC5C", "BMPR1B") |
| 26 | HALLMARK_NOTCH_SIGNALING | 0.420485 | 0.758354 | -0.341782 | -1.024733 | 1403 | 11 | c("CCND1", "ARRB1", "LFNG") |
| 27 | HALLMARK_ADIPOGENESIS | 0.395072 | 0.758354 | -0.216852 | -1.031843 | 993 | 61 | c("G3BP2", "TKT", "GPAT4", "APOE", "STOM", "COX8A", "DHRS7", "HSPB8", "CMBL", "UQCRC1", "TALDO1", "PIM3", "GPX3", "UQCR11", "ETFB") |
| 28 | HALLMARK_MYC_TARGETS_V2 | 0.424678 | 0.758354 | -0.340606 | -1.021206 | 1417 | 11 | c("GNL3", "PA2G4", "CBX3", "MCM4", "IMP4", "CDK4", "SRM", "NOP56", "DCTPP1", "WDR43", "DDX18") |
| 29 | HALLMARK_KRAS_SIGNALING_UP | 0.451327 | 0.778151 | 0.243294 | 1.013847 | 3365 | 60 | c("PEG3", "EREG", "MMP11", "SOX9", "SPON1", "IGFBP3", "MMD", "DUSP6", "ITGBL1", "CFB", "NIN", "INHBA", "SPARCL1", "SDCCAG8", "DCBLD2") |
| 30 | HALLMARK_COAGULATION | 0.526355 | 0.877259 | -0.213104 | -0.959879 | 1397 | 48 | c("SERPINA1", "MMP9", "CFD", "PLAT", "CD9", "PRSS23", "LTA4H") |
| 31 | HALLMARK_TNFA_SIGNALING_VIA_NFKB | 0.599131 | 0.911824 | 0.211257 | 0.920268 | 4547 | 75 | c("PDE4B", "EGR3", "AREG", "SLC16A6", "NFKBIA", "BTG2", "OLR1", "ATF3", "SPSB1", "JUN", "EGR1", "INHBA", "EFNA1", "GADD45B", "RHOB", "NFIL3", "IER3", "B4GALT5", "CD44", "NFE2L2", "KLF6", "IER2", "GADD45A", "KLF9", "CCN1", "TNFAIP3", "PFKFB3", "ZFP36") |
| 32 | HALLMARK_ANGIOGENESIS | 0.572479 | 0.911824 | 0.287054 | 0.924964 | 3968 | 20 | c("OLR1", "VCAN", "PTK2", "COL5A2", "POSTN", "APP", "LUM", "STC1", "COL3A1") |
| 33 | HALLMARK_IL2_STAT5_SIGNALING | 0.62004 | 0.911824 | 0.21941 | 0.901382 | 4609 | 56 | c("S100A1", "ENPP1", "CISH", "AHR", "SPRED2", "F2RL2", "ADAM19", "WLS", "LRIG1", "GADD45B", "RHOB", "NFIL3", "TIAM1", "HIPK2", "ANXA4", "MYO1E", "PLEC", "CD44", "ST3GAL4", "SNX9", "KLF6") |
| 34 | HALLMARK_PANCREAS_BETA_CELLS | 0.616419 | 0.911824 | 0.423057 | 0.896713 | 3851 | 5 | c("AKT3", "SRP9", "VDR") |
| 35 | HALLMARK_HYPOXIA | 0.697717 | 0.918098 | 0.198131 | 0.860726 | 5287 | 74 | c("CITED2", "ATF3", "IGFBP3", "JUN", "GPC4", "KLF7", "CAVIN1", "CCN2", "COL5A1", "LOX", "EFNA1", "PAM", "NFIL3", "AKAP12", "TGFB3", "TES", "IER3", "ERRFI1", "FOXO3", "STC1", "KLF6", "EXT1", "CCN1", "DDIT4", "TNFAIP3", "PFKFB3", "MYH9", "ZFP36", "NDST1", "SDC2") |
| 36 | HALLMARK_WNT_BETA_CATENIN_SIGNALING | 0.716925 | 0.918098 | -0.272786 | -0.791844 | 2405 | 10 | c("CTNNB1", "NCOR2", "CUL1", "CSNK1E", "TP53", "MAML1", "CCND2", "NCSTN", "JAG1", "ADAM17") |
| 37 | HALLMARK_APOPTOSIS | 0.720138 | 0.918098 | 0.198341 | 0.841073 | 5413 | 65 | c("TGFB2", "EGR3", "EREG", "BTG2", "ERBB2", "ATF3", "JUN", "PTK2", "TNFRSF12A", "PDCD4", "CREBBP", "BCL2L11") |
| 38 | HALLMARK_INTERFERON_GAMMA_RESPONSE | 0.704185 | 0.918098 | -0.195651 | -0.837384 | 1951 | 39 | c("IFITM2", "IFI30", "PSME1", "SOD2", "PSMB10", "TAPBP", "MYD88", "LY6E", "BTG1", "SOCS3", "TRIM25", "B2M", "PIM1") |

| | | | | | | | | |
|---|---|---|---|---|---|---|---|---|
| 39 | HALLMARK_MYC_TARGETS_V1 | 0.734479 | 0.918098 | -0.178214 | -0.84207 | 1880 | 59 | c("CLNS1A", "GNL3", "HNRNPU", "PCBP1", "CNBP", "BUB3", "HNRNPD", "SF3B3", "IMPDH2", "PA2G4", "PPIA", "CBX3", "HNRNPC", "CUL1", "VDAC3", "POLD2", "CYC1", "CANX", "FAM120A", "EEF1B2", "RAD23B", "DUT", "SMARCC1", "MCM4", "RACK1", "SET", "SRSF2", "NCBP2", "SNRPA", "POLE3", "SNRPD2", "CDK4", "HNRNPR", "SERBP1", "SRSF3", "TUFM", "FBL", "SRM", "ILF2", "NDUFAB1", "NOP56", "SRPK1", "DDX21", "PPM1G", "EIF4G2", "HNRNPA3", "YWHAE", "KPNB1", "PSMD1", "DDX18", "RNPS1", "TFDP1", "NHP2", "HDGF", "RUVBL2", "GLO1", □ "CCT3", "YWHAQ") |
| 40 | HALLMARK_FATTY_ACID_METABOLISM | 0.659237 | 0.918098 | -0.215526 | -0.856197 | 1951 | 29 | c("BMPR1B", "PSME1", "ACAA1", "UROD", "UGDH", "FH", "ECHS1", "HSP90AA1", "CD36", "HSD17B10", "IDH3B", "HSD17B11", "SERINC1", "MIF", "HSPH1", "ELOVL5", "PDHB", "ACADVL", "LGALS1", "PTPRG", "UBE2L6", "YWHAH", "ACADM", "S100A10", "ALDH9A1", "SMS", "DECR1", "D2HGDH", "DHCR24") |
| 41 | HALLMARK_DNA_REPAIR | 0.796184 | 0.925796 | 0.198871 | 0.774179 | 5800 | 43 | c("CCNO", "PDE4B") |
| 42 | HALLMARK_MYOGENESIS | 0.779556 | 0.925796 | 0.193832 | 0.796301 | 5795 | 56 | c("TPD52L1", "ADAM12", "WWTR1", "IGFBP3", "TAGLN", "EPHB3", "CNN3", "AEBP1", "TPM2", "PDLIM7", "SVIL", "LAMA2", "NAV2", "GADD45B", "ITGB5", "PTGIS", "APP", "IGFBP7", "SORBS1") |
| 43 | HALLMARK_SPERMATOGENESIS | 0.775955 | 0.925796 | 0.286706 | 0.761569 | 5156 | 10 | c("NPHP1", "IFT88", "SLC12A2") |
| 44 | HALLMARK_PEROXISOME | 0.818634 | 0.930266 | 0.209822 | 0.744723 | 5763 | 29 | c("CRABP1", "CADM1", "PEX11A", "ATXN1", "CRABP2", "SEMA3C") |
| 45 | HALLMARK_G2M_CHECKPOINT | 0.988141 | 0.997935 | -0.117798 | -0.533404 | 2582 | 50 | c("CCND1", "HNRNPU", "RBM14", "BCL3", "TOP2A", "BUB3") |
| 46 | HALLMARK_PROTEIN_SECRETION | 0.961618 | 0.997935 | -0.140677 | -0.571307 | 2805 | 32 | c("ANP32E", "MON2", "CD63", "M6PR", "PPT1", "RER1", "TOM1L1", "CLTA", "TPD52", "COPB2") |
| 47 | HALLMARK_APICAL_SURFACE | 0.942193 | 0.997935 | -0.198435 | -0.576017 | 3161 | 10 | GSTM3 |
| 48 | HALLMARK_HEDGEHOG_SIGNALING | 0.898391 | 0.997935 | -0.217313 | -0.630816 | 3014 | 10 | c("CELSR1", "NRP1", "ADGRG1") |
| 49 | HALLMARK_UNFOLDED_PROTEIN_RESPONSE | 0.997935 | 0.997935 | 0.107913 | 0.413427 | 7247 | 40 | c("ATF3", "PAIP1", "WIPI1", "CKS1B", "PARN", "NHP2", "SHC1", "ERN1", "DDIT4", "CNOT6", "DCP2", "HSPA9", "SRPRA", "NOP56", "MTREX", "SLC1A4", "PDIA6", "ARFGAP1", "DCTN1") |
| 50 | HALLMARK_E2F_TARGETS | 0.935767 | 0.997935 | -0.148538 | -0.623203 | 2665 | 36 | c("NUDT21", "TFRC", "ANP32E", "HMGB2", "TOP2A", "PNN", "HNRNPD") |

| Cluster 2 | pathway | pval | padj | ES | NES | nMoreExtreme | size | leadingEdge |
|---|---|---|---|---|---|---|---|---|
| 1 | HALLMARK_ESTROGEN_RESPONSE_EARLY | 0.0001 | 0.001272 | -0.682992 | -2.071535 | 0 | 76 | c("RET", "SEC14L2", "PGR", "MYB", "KRT19", "STC2", "HSPB8", "GFRA1", "THSD4", "MUC1", "SLC7A2", "TFF1", "PRSS23", "MAPT", "ELF3", "NBL1", "SEMA3B", "TFF3", "LRIG1", "ADCY1", "CCND1", "KRT8", "FHL2", "GREB1", "RHOBTB3", "CXCL12", "GJA1", "CA12", "SLC39A6", "FLNB", "IGFBP4", "HES1", "PLAAT3", "RHOD", "SCNN1A", "TTC39A", "FOS", "MAST4", "FASN", "IGF1R", "GAB2", "SLC9A3R1", "MLPH", "SIAH2", "TOB1", "XBP1", "FAM102A") |
| 2 | HALLMARK_ESTROGEN_RESPONSE_LATE | 0.000101 | 0.001272 | -0.63143 | -1.911138 | 0 | 73 | c("RET", "SERPINA3", "PGR", "MYB", "KRT19", "AGR2", "CXCL14", "HSPB8", "TFF1", "PRSS23", "MAPT", "CPE", "PRLR", "NBL1", "SCUBE2", "SEMA3B", "LTF", "TFF3", "CCND1", "CXCL12", "CA12", "FLNB", "MDK", "IGFBP4", "PERP", "CDH1", "PLAAT3", "SCNN1A", "TSPAN13", "TOP2A", "FOS", "DLG5", "SLC9A3R1", "SIAH2", "TOB1", "XBP1", "FAM102A", "COX6C", "LLGL2", "ELOVL5", "UGDH", "PDCD4", "BCL2", "SERPINA1", "DNAJC1", "ADD3", "LSR", "ABHD2", "NRIP1") |
| 3 | HALLMARK_MYOGENESIS | 0.000102 | 0.001272 | -0.599884 | -1.763853 | 0 | 49 | c("APOD", "STC2", "COL6A3", "HSPB8", "AEBP1", "SPDEF", "TNNT1", "CNN3", "ADAM12", "COL3A1", "PVALB", "CLU", "COL6A2", "COL1A1", "IGFBP7", "IGFBP3", "ERBB3", "COL4A2", "TPM2", "TAGLN") |
| 4 | HALLMARK_EPITHELIAL_MESENCHYMAL_TRANSITION | 0.0001 | 0.001272 | -0.701835 | -2.172814 | 0 | 108 | c("POSTN", "COL11A1", "COL5A2", "FSTL1", "COL6A3", "COMP", "FBLN1", "BGN", "CCN2", "CDH11", "THBS2", "COL12A1", "SFRP4", "DCN", "THBS1", "LUM", "COL5A1", "LOXL1", "TNC", "FBLN2", "MGP", "THY1", "NNMT", "CCN1", "ADAM12", "CTHRC1", "CALD1", "ACTA2", "TIMP3", "ELN", "MXRA5", "DPYSL3", "GEM", "PRRX1", "COL4A1", "LOX", "COL3A1", "COL16A1", "PCOLCE", "CXCL12", "LOXL2", "FBN1", "PDGFRB", "MYL9", "GJA1", "SDC1", "COL6A2", "COL1A1", "FN1", "HTRA1", "COL1A2", "GPC1", "IGFBP3", "IGFBP4", "COL4A2", "INHBA", "FAP", "IGFBP2", "TPM2", "RHOB", "TAGLN", "PMEPA1", "LRRC15", "TPM1") |
| 5 | HALLMARK_BILE_ACID_METABOLISM | 0.000932 | 0.00932 | 0.580301 | 2.254213 | 0 | 17 | c("CYP27A1", "ABCA1", "NPC1", "AQP9", "SLC23A2", "HSD17B4", "HSD3B7", "ALDH1A1", "ACSL1", "SLC29A1", "RXRA") |
| 6 | HALLMARK_INFLAMMATORY_RESPONSE | 0.003344 | 0.020903 | 0.39908 | 2.006374 | 0 | 39 | c("MARCO", "CXCL8", "CCL2", "PLAUR", "CD48", "ABCA1", "MSR1", "AQP9", "ADM", "CYBB", "EMP3", "PIK3R5", "CD82", "CLEC5A", "TNFRSF1B", "CSF1", "ICAM1", "LYN", "CD14") |
| 7 | HALLMARK_UV_RESPONSE_DN | 0.00305 | 0.020903 | -0.529606 | -1.560254 | 29 | 50 | c("COL11A1", "COL5A2", "INPP4B", "IGFBP5", "EFEMP1", "CCN1", "RBPMS", "FHL2", "COL3A1", "PDGFRB", "GJA1", "ERBB2", "COL1A1", "COL1A2", "APBB2", "DUSP1", "IGF1R", "PIK3R3", "AKT3") |
| 8 | HALLMARK_ALLOGRAFT_REJECTION | 0.002681 | 0.020903 | 0.407502 | 2.006485 | 0 | 35 | c("CCL2", "CAPG", "CTSS", "CCR1", "ITGB2", "SPI1", "BCAT1", "CD4", "FGR", "MMP9", "IRF8", "C2", "CSF1", "B2M", "CD74", "ICAM1", "LYN", "TAPBP", "PSMB10", "IFNGR1", "SRGN") |
| 9 | HALLMARK_IL6_JAK_STAT3_SIGNALING | 0.005057 | 0.028094 | 0.441399 | 1.921047 | 3 | 23 | c("HMOX1", "CCR1", "TNFRSF21", "PIK3R5", "TNFRSF1B", "CSF1", "CD36", "MYD88", "CD14", "IFNGR1", "STAT2", "GRB2", "PIM1", "CSF3R", "STAT1") |
| 10 | HALLMARK_COAGULATION | 0.010194 | 0.050968 | -0.510585 | -1.495678 | 99 | 47 | c("PLAT", "COMP", "MMP11", "THBS1", "PRSS23", "CRIP2", "TIMP3", "CFB", "C1R", "FBN1", "CLU", "FN1", "HTRA1", "S100A1", "HPN", "C3", "C1S", "SERPING1", "S100A13", "SPARC", "MMP2") |
| 11 | HALLMARK_NOTCH_SIGNALING | 0.020773 | 0.094423 | -0.718241 | -1.553758 | 164 | 8 | c("CCND1", "NOTCH3", "HES1") |
| 12 | HALLMARK_P53_PATHWAY | 0.026144 | 0.108932 | 0.258859 | 1.419386 | 3 | 54 | c("HMOX1", "IFI30", "CTSD", "GM2A", "FUCA1", "RAP2B", "NUPR1", "UPP1", "CEBPA", "SAT1", "TCN2") |
| 13 | HALLMARK_ANDROGEN_RESPONSE | 0.035248 | 0.128149 | -0.500445 | -1.42921 | 339 | 36 | c("AZGP1", "KRT19", "BMPR1B", "INPP4B", "SPDEF", "CCND1", "KRT8", "TPD52", "STEAP4", "PMEPA1", "UAP1", "SELENOP", "ELOVL5") |
| 14 | HALLMARK_KRAS_SIGNALING_UP | 0.035882 | 0.128149 | -0.478366 | -1.401299 | 351 | 47 | c("SERPINA3", "APOD", "PLAT", "MMP11", "SPARCL1", "CPE", "SEMA3B", "CFB", "PRRX1", "ITGBL1", "IGFBP3", "JUP", "TSPAN13", "INHBA") |
| 15 | HALLMARK_ANGIOGENESIS | 0.050079 | 0.166929 | -0.574794 | -1.445281 | 444 | 16 | c("POSTN", "COL5A2", "FSTL1", "STC1", "LUM", "COL3A1") |
| 16 | HALLMARK_XENOBIOTIC_METABOLISM | 0.058824 | 0.183824 | 0.239896 | 1.315407 | 8 | 54 | c("HMOX1", "APOE", "CYP27A1", "CES1", "UPP1", "BCAT1", "NPC1", "PGD", "AQP9", "POR", "IRF8", "ACP2", "SLC6A6", "CNDP2", "PTGDS", "FBP1", "ABCC3", "CD36", "BLVRB") |
| 17 | HALLMARK_KRAS_SIGNALING_DN | 0.089557 | 0.263404 | -0.553688 | -1.373346 | 788 | 15 | c("CPB1", "BMPR1B", "EFHD1", "GPRC5C", "IGFBP2", "SELENOP", "IDUA", "BTG2") |
| 18 | HALLMARK_HYPOXIA | 0.104876 | 0.291322 | -0.416281 | -1.25356 | 1040 | 67 | c("STC2", "BGN", "CCN2", "STC1", "DCN", "COL5A1", "CCN1", "TGFB3", "CAVIN1", "LOX", "TPD52", "CA12", "GPC1", "IGFBP3", "SELENBP1", "EFNA1", "DUSP1", "FOS", "SIAH2", "DTNA", "CCNG2", "MT2A", "BCL2", "PAM", "PNRC1", "DDIT4") |

| | | | | | | | | |
|---|---|---|---|---|---|---|---|---|
| 19 | HALLMARK_APICAL_JUNCTION | 0.122957 | 0.307392 | -0.423706 | -1.254317 | 1210 | 54 | c("PARVA", "CDH11", "THY1", "COL16A1", "FBN1", "MYL9", "MDK", "JUP", "CDH1", "CLDN4", "PCDH1", "LIMA1", "PIK3R3", "AKT3", "CERCAM", "CLDN7", "MMP2", "ACTG1", "VCAN", "EPB41L2", "SHC1") |
| 20 | HALLMARK_APICAL_SURFACE | 0.12054 | 0.307392 | -0.584384 | -1.335899 | 999 | 10 | c("GSTM3", "THY1", "GATA3") |
| 21 | HALLMARK_GLYCOLYSIS | 0.152149 | 0.362261 | -0.425525 | -1.24258 | 1489 | 45 | c("STC2", "STC1", "DCN", "COL5A1", "ELF3", "TFF3", "SDC1", "GPC1", "IGFBP3", "FUT8", "CHPF", "EGLN3", "AGRN", "IDUA", "VCAN", "PAM", "DDIT4") |
| 22 | HALLMARK_TGF_BETA_SIGNALING | 0.171835 | 0.390533 | -0.465727 | -1.257956 | 1595 | 24 | c("THBS1", "LTBP2", "CDH1", "PMEPA1", "CTNNB1", "FNTA", "ID3", "SMAD3", "SKIL", "SERPINE1", "BMPR2", "ARID4B", "JUNB") |
| 23 | HALLMARK_SPERMATOGENESIS | 0.183706 | 0.399361 | -0.625899 | -1.255995 | 1370 | 6 | c("GSTM3", "SLC12A2") |
| 24 | HALLMARK_APOPTOSIS | 0.22685 | 0.472604 | -0.390322 | -1.164234 | 2243 | 59 | c("PLAT", "BGN", "DCN", "LUM", "TIMP3", "CCND1", "PDGFRB", "CLU", "ERBB2", "ERBB3", "RHOB", "TOP2A", "IFITM3", "NEDD9", "CTNNB1", "MMP2", "PDCD4", "BTG2", "HMGB2", "ANKH", "RARA", "TIMP1", "APP", "SPTAN1") |
| 25 | HALLMARK_MTORC1_SIGNALING | 0.248366 | 0.496732 | 0.209274 | 1.147498 | 37 | 54 | c("IFI30", "ITGB2", "LGMN", "NUPR1", "HK2", "BCAT1", "TXNRD1", "GLRX", "M6PR", "TOMM40", "ENO1", "SLC2A1", "SLC6A6", "SQSTM1", "LTA4H", "GLA") |
| 26 | HALLMARK_REACTIVE_OXYGEN_SPECIES_PATHWAY | 0.263797 | 0.507302 | 0.276785 | 1.132119 | 238 | 20 | c("FTL", "SOD2", "LSP1", "TXNRD1", "GLRX", "ATOX1", "MBP", "GPX4", "PRNP", "SRXN1", "ABCC1") |
| 27 | HALLMARK_INTERFERON_GAMMA_RESPONSE | 0.318182 | 0.589226 | 0.209556 | 1.086534 | 69 | 44 | c("IFI30", "CCL2", "IL18BP", "UPP1", "SOD2", "IRF5", "IRF8", "TNFAIP2", "B2M", "CD74", "ICAM1", "MYD88", "LAP3", "SAMHD1", "OAS3", "TAPBP") |
| 28 | HALLMARK_MITOTIC_SPINDLE | 0.386322 | 0.689861 | -0.366303 | -1.077048 | 3795 | 49 | c("PALLD", "FLNB", "NET1", "ARHGAP29", "SHROOM1", "TOP2A", "NEDD9", "ARHGEF12", "ARFIP2", "CTTN", "SMC4", "DST", "NUMA1", "EPB41L2", "SPTAN1", "CYTH2", "FARP1", "STAU1", "EZR", "YWHAE", "WASL", "GSN", "MYH9", "RANBP9") |
| 29 | HALLMARK_PANCREAS_BETA_CELLS | 0.42235 | 0.72819 | -0.525053 | -1.053627 | 3151 | 6 | c("SYT13", "AKT3") |
| 30 | HALLMARK_TNFA_SIGNALING_VIA_NFKB | 0.456404 | 0.760673 | -0.341114 | -1.030384 | 4532 | 70 | c("DUSP4", "TNC", "CCN1", "CCND1", "GEM", "PLK2", "FOSB", "NR4A1", "EGR1", "HES1", "INHBA", "EFNA1", "RHOB", "DUSP1", "PMEPA1", "FOS") |
| 31 | HALLMARK_G2M_CHECKPOINT | 0.51381 | 0.802828 | -0.356508 | -0.994869 | 4873 | 30 | c("CCND1", "SLC12A2", "TOP2A", "SLC38A1", "MT2A", "SMC4", "HNRNPU", "NUMA1", "SMAD3", "MYC", "BUB3", "BCL3", "SMARCC1", "CUL3", "ABL1", "RBM14", "HNRNPD", "YTHDC1", "NCL", "CDKN1B") |
| 32 | HALLMARK_HEDGEHOG_SIGNALING | 0.50328 | 0.802828 | -0.446572 | -0.992653 | 4065 | 9 | c("THY1", "ADGRG1", "CELSR1", "MYH9", "VEGFA", "LDB1", "DPYSL2", "RASA1") |
| 33 | HALLMARK_WNT_BETA_CATENIN_SIGNALING | 0.561381 | 0.825561 | -0.503937 | -0.960269 | 4014 | 5 | c("CTNNB1", "CSNK1E", "MYC") |
| 34 | HALLMARK_UV_RESPONSE_UP | 0.550554 | 0.825561 | -0.339821 | -0.972559 | 5319 | 37 | c("RET", "EPCAM", "FOSB", "NR4A1", "IGFBP2", "RHOB", "FOS") |
| 35 | HALLMARK_INTERFERON_ALPHA_RESPONSE | 0.615894 | 0.832289 | 0.220695 | 0.902699 | 557 | 20 | c("IFI30", "TMEM140", "CSF1", "B2M", "CD74", "LAP3", "OAS1", "STAT2", "IFI35", "TAP1") |
| 36 | HALLMARK_E2F_TARGETS | 0.606123 | 0.832289 | -0.345914 | -0.925978 | 5582 | 23 | c("TOP2A", "SMC4", "RBBP7", "HMGB2", "MYC", "MCM7", "ANP32E", "PRKDC", "NUDT21", "POLD2", "NME1", "HNRNPD", "TUBB", "CDKN1B", "LUC7L3", "SSRP1", "ILF3") |
| 37 | HALLMARK_HEME_METABOLISM | 0.597256 | 0.832289 | -0.32301 | -0.95277 | 5876 | 51 | c("HBB", "HBD", "SLC4A1", "BCAM", "SELENBP1", "PRDX2", "C3", "FBXO9", "ARHGEF12", "SLC25A37", "BTG2", "FBXO7") |
| 38 | HALLMARK_ADIPOGENESIS | 0.757419 | 0.946774 | -0.284766 | -0.84626 | 7477 | 56 | c("HSPB8", "SPARCL1", "CAVIN1", "COL4A1", "LAMA4", "FABP4", "C3", "TOB1", "PHLDB1", "CMBL", "CCNG2", "PDCD4", "GPAT4") |
| 39 | HALLMARK_PI3K_AKT_MTOR_SIGNALING | 0.751402 | 0.946774 | 0.179145 | 0.839018 | 401 | 29 | c("SLC2A1", "SQSTM1", "DUSP3", "MYD88", "ACTR3", "CXCR4", "CFL1", "PFN1", "ARPC3", "ARHGDIA", "STAT2", "GRB2", "MAPKAP1", "CALR", "CDK4") |
| 40 | HALLMARK_MYC_TARGETS_V1 | 0.723343 | 0.946774 | -0.29505 | -0.857616 | 7061 | 43 | c("CLNS1A", "IMPDH2", "CBX3", "HSP90AB1", "HNRNPU", "GNL3", "MYC", "HSPD1", "MCM7", "CCT3", "SNRPD2", "BUB3", "YWHAE", "ILF2", "RNPS1", "NHP2", "SMARCC1", "RACK1", "POLD2", "NME1", "CANX", "HNRNPD", "PCBP1", "VDAC3", "HNRNPC", "LSM7", "HDGF", "PPIA", "SF3B3", "EEF1B2") |
| 41 | HALLMARK_FATTY_ACID_METABOLISM | 0.783216 | 0.955142 | -0.287221 | -0.784928 | 7344 | 26 | c("BMPR1B", "FASN", "PRDX6", "ELOVL5", "UGDH", "FH", "MIF", "ECI1", "DHCR24") |
| 42 | HALLMARK_COMPLEMENT | 0.84679 | 0.962262 | -0.260311 | -0.779718 | 8389 | 62 | c("PLAT", "LTF", "MMP13", "GATA3", "CFB", "C1R", "CLU", "FN1", "COL4A2", "C3", "C1S", "SERPING1") |
| 43 | HALLMARK_MYC_TARGETS_V2 | 0.825845 | 0.962262 | -0.327641 | -0.728289 | 6671 | 9 | c("CBX3", "GNL3", "MYC", "HSPD1") |
| 44 | HALLMARK_IL2_STAT5_SIGNALING | 0.832485 | 0.962262 | -0.262653 | -0.772284 | 8179 | 49 | c("MUC1", "LRIG1", "S100A1", "ENPP1", "RHOB", "COL6A1", "IGF1R", "IFITM3", "XBP1", "ITGA6", "NFKBIZ", "MYO1C", "CKAP4", "BCL2", "ITGAV", "ECM1", "MYC", "BMPR2", "PHLDA1", "ALCAM") |
| 45 | HALLMARK_CHOLESTEROL_HOMEOSTASIS | 0.878205 | 0.975783 | -0.257723 | -0.668544 | 7945 | 19 | c("SEMA3B", "CLU", "FASN", "CTNNB1", "ACTG1", "PNRC1") |
| 46 | HALLMARK_DNA_REPAIR | 0.942161 | 0.999186 | -0.213637 | -0.590245 | 8893 | 28 | c("BCAM", "IMPDH2", "CANT1", "POLR2H", "BRF2", "NME4", "NME3", "NELFCD", "TMED2", "NUDT21", "NME1", "POLR2I") |

| | | | | | | | | |
|---|---|---|---|---|---|---|---|---|
| 47 | HALLMARK_PROTEIN_SECRETION | 0.990415 | 0.999186 | 0.121111 | 0.605965 | 309 | 38 | c("ABCA1", "IGF2R", "M6PR", "CD63", "PPT1", "GLA", "STX7", "CLTA", "ATP6V1H", "SNX2", "CTSC", "STX12", "SEC22B", "VAMP3", "LAMP2") |
| 48 | HALLMARK_UNFOLDED_PROTEIN_RESPONSE | 0.988576 | 0.999186 | -0.164795 | -0.469173 | 9518 | 35 | c("STC2", "XBP1", "DDIT4", "EEF2", "SHC1", "DNAJA4", "LSM1", "FUS", "NHP2", "BANF1", "VEGFA", "SPCS1", "SEC31A", "EIF4A2", "EIF4G1") |
| 49 | HALLMARK_OXIDATIVE_PHOSPHORYLATION | 0.999186 | 0.999186 | -0.121973 | -0.358162 | 9816 | 48 | c("MAOB", "PDK4", "COX6C", "ATP1B1", "COX6B1", "NDUFC2", "FH", "ECI1", "COX6A1") |
| 50 | HALLMARK_PEROXISOME | 0.971966 | 0.999186 | -0.198751 | -0.520482 | 8840 | 20 | c("CRABP2", "TOP2A", "ELOVL5") |

| Cluster 3 | pathway | pval | padj | ES | NES | nMoreExtreme | size | leadingEdge |
|---|---|---|---|---|---|---|---|---|
| 1 | HALLMARK_ESTROGEN_RESPONSE_EARLY | 0.000141 | 0.003558 | -0.648388 | -2.72491 | 0 | 75 | c("ADCY1", "RET", "SEC14L2", "GFRA1", "ABHD2", "SCNN1A", "SLC39A6", "SLC2A1", "SLC9A3R1", "PGR", "HSPB8", "PRSS23", "STC2", "KRT8", "GAB2", "FLNB", "MUC1", "CELSR1", "GREB1", "CLDN7", "MAPT", "SIAH2", "RHOBTB3", "TFF1", "KDM4B", "CA12", "BLVRB", "THSD4", "CCND1", "FAM102A", "RHOD", "KRT19", "IGF1R", "MYB", "GJA1", "SEMA3B", "RARA", "CANT1", "BAG1", "MLPH") |
| 2 | HALLMARK_ESTROGEN_RESPONSE_LATE | 0.000142 | 0.003558 | -0.614249 | -2.544477 | 0 | 70 | c("S100A9", "LTF", "RET", "CDH1", "SERPINA1", "SERPINA3", "COX6C", "ABHD2", "SCNN1A", "AGR2", "SLC9A3R1", "PGR", "HSPB8", "PRSS23", "CD9", "TOP2A", "LSR", "FLNB", "MAPT", "TSPAN13", "SIAH2", "TFF1", "SCUBE2", "CA12", "BLVRB", "CCND1", "FAM102A", "LLGL2", "DNAJC1", "KRT19", "MYB", "SEMA3B", "DCXR", "BAG1", "PAPSS2", "DLG5", "MAPK13") |
| 3 | HALLMARK_ADIPOGENESIS | 0.000329 | 0.005479 | 0.55894 | 2.658733 | 0 | 67 | c("FABP4", "ADIPOQ", "LPL", "LIPE", "GPAM", "ITIH5", "ITGA7", "GPX3", "ANGPTL4", "CAVIN2", "CD36", "PEMT", "ITSN1", "FZD4", "PPARG", "ALDH2", "SORBS1", "LAMA4", "CAVIN1", "SNCG", "LIFR", "PHLDB1", "C3", "PFKFB3") |
| 4 | HALLMARK_HEME_METABOLISM | 0.002395 | 0.029936 | -0.488856 | -1.867011 | 15 | 46 | c("HBB", "SLC4A1", "HBD", "SLC2A1", "SNCA", "ACP5", "SLC25A37", "TFRC", "BLVRB", "FBXO7", "FBXO9", "HAGH", "PRDX2", "BCAM", "UBAC1") |
| 5 | HALLMARK_MYOGENESIS | 0.005096 | 0.050955 | 0.371368 | 1.688462 | 15 | 56 | c("FHL1", "CRYAB", "APOD", "ITGA7", "GPX3", "CD36", "BIN1", "CFD", "SCD", "SORBS1", "PTGIS", "ACSL1", "IGF1", "IGFBP7", "GSN", "COX7A1", "COL4A2", "TAGLN", "SOD3") |
| 6 | HALLMARK_UV_RESPONSE_DN | 0.017977 | 0.128406 | 0.353842 | 1.545502 | 58 | 47 | c("CAV1", "PPARG", "PRKAR2B", "DLC1", "FBLN5", "CELF2", "TGFBR2", "MAP1B", "DUSP1", "DAB2", "RBPMS", "CCN1", "NFIB", "PTEN", "NRP1", "LAMC1", "PTPRM", "ATP2B4", "EFEMP1", "ADD3", "NR3C1", "SNAI2", "AKT3", "TJP1", "MGLL", "PDGFRB", "CITED2", "PLPP3", "NEK7", "SERPINE1") |
| 7 | HALLMARK_BILE_ACID_METABOLISM | 0.015506 | 0.128406 | 0.540126 | 1.761443 | 60 | 16 | c("LIPE", "ABCA8", "ABCA6", "ACSL1", "RETSAT", "HSD17B11", "PEX19", "ALDH1A1", "OPTN", "ABCA1", "CAT") |
| 8 | HALLMARK_KRAS_SIGNALING_UP | 0.032907 | 0.205667 | 0.337427 | 1.473805 | 107 | 47 | c("RBP4", "APOD", "G0S2", "ANGPTL4", "EPB41L3", "F13A1", "SPRY2", "ADGRL4", "AKAP12", "MMD", "TNFAIP3", "ADGRA2", "ENG", "ETS1", "SPARCL1", "KLF4", "MAFB", "NRP1", "PECAM1") |
| 9 | HALLMARK_MYC_TARGETS_V1 | 0.054799 | 0.304438 | -0.417244 | -1.486486 | 353 | 34 | c("CLNS1A", "GNL3", "HNRNPU", "BUB3", "HSP90AB1", "HDGF", "IMPDH2", "CBX3", "HSPE1", "NME1", "VDAC3", "POLD2", "PSMD3", "ILF2", "NDUFAB1", "TXNL4A", "PPIA", "CANX", "LSM7", "PCBP1", "HNRNPA2B1", "PA2G4", "DHX15", "HNRNPC", "NOP56", "SMARCC1", "TUFM", "CCT3") |
| 10 | HALLMARK_ANDROGEN_RESPONSE | 0.070274 | 0.351368 | -0.407853 | -1.443117 | 451 | 33 | c("BMPR1B", "ABHD2", "SPDEF", "KRT8", "INPP4B", "TPD52", "AZGP1", "STEAP4", "CCND1", "KRT19", "GSR") |
| 11 | HALLMARK_MTORC1_SIGNALING | 0.077916 | 0.354165 | -0.36225 | -1.398892 | 524 | 48 | c("SLC2A1", "SLC9A3R1", "CD9", "IFI30", "TFRC", "GGA2", "CORO1A", "SYTL2", "GSR", "STC1", "XBP1", "P4HA1", "NUPR1", "CALR", "BHLHE40", "DHCR24", "MAP2K3", "SQSTM1", "GSK3B", "SERP1", "HSPE1", "LTA4H") |
| 12 | HALLMARK_DNA_REPAIR | 0.098606 | 0.410857 | -0.468629 | -1.415687 | 593 | 18 | c("BRF2", "CANT1", "BCAM", "NME4", "NUDT21", "IMPDH2", "POLR2H", "NELFCD") |
| 13 | HALLMARK_INTERFERON_GAMMA_RESPONSE | 0.125984 | 0.414467 | 0.331026 | 1.312771 | 447 | 32 | c("TXNIP", "ARID5B", "TNFAIP3", "SAMHD1", "MT2A", "C1R", "FGL2", "C1S", "SERPING1", "APOL6", "PML", "SOCS3", "NFKBIA", "IRF8", "CIITA") |
| 14 | HALLMARK_UNFOLDED_PROTEIN_RESPONSE | 0.111305 | 0.414467 | -0.415683 | -1.373361 | 700 | 25 | c("STC2", "LSM1", "DNAJA4", "XBP1", "EIF4EBP1", "BANF1", "CALR", "VEGFA", "SPCS1", "SERP1", "LSM4", "EIF4A3", "FUS", "ATP6V0D1") |
| 15 | HALLMARK_E2F_TARGETS | 0.132629 | 0.414467 | -0.441542 | -1.351557 | 803 | 19 | c("TOP2A", "TFRC", "ANP32E", "RBBP7", "NUDT21", "SNRPB", "HMGB2", "NME1", "SMC4", "POLD2") |
| 16 | HALLMARK_SPERMATOGENESIS | 0.12581 | 0.414467 | -0.628022 | -1.382652 | 698 | 6 | c("GSTM3", "STRBP", "SLC12A2", "TALDO1", "VDAC3") |
| 17 | HALLMARK_TNFA_SIGNALING_VIA_NFKB | 0.150729 | 0.418693 | 0.262078 | 1.220266 | 464 | 61 | c("G0S2", "KLF9", "PER1", "TNFAIP3", "IRS2", "PFKFB3", "NFIL3", "ZFP36", "DUSP1", "PNRC1", "CCN1", "KLF4", "TRIP10", "JUN", "GEM", "CEBPD", "ETS2", "SOCS3", "NFKBIA", "JAG1", "KLF6", "NFE2L2", "GADD45B", "MCL1", "RHOB", "NFAT5", "ABCA1", "PLPP3", "KLF2", "EGR1") |
| 18 | HALLMARK_APICAL_SURFACE | 0.14381 | 0.418693 | -0.561309 | -1.354074 | 818 | 8 | c("PLAUR", "GSTM3", "SULF2", "GATA3") |
| 19 | HALLMARK_XENOBIOTIC_METABOLISM | 0.170595 | 0.448935 | 0.290737 | 1.237973 | 578 | 42 | c("PDK4", "RBP4", "CD36", "PEMT", "ALDH2", "IL1R1", "ADH5", "IGF1", "MT2A", "RETSAT") |
| 20 | HALLMARK_HYPOXIA | 0.202003 | 0.505008 | 0.244236 | 1.167777 | 604 | 69 | c("ANGPTL4", "GPC3", "SRPX", "CAV1", "CCN2", "CAVIN1", "AKAP12", "TNFAIP3", "IRS2", "PFKFB3", "NFIL3", "DDIT4", "ZFP36", "MT2A", "DUSP1", "PNRC1", "CCN1", "ETS1", "ERRFI1", "JUN") |

| | | | | | | | | |
|---|---|---|---|---|---|---|---|---|
| 21 | HALLMARK_COMPLEMENT | 0.285503 | 0.594798 | -0.293047 | -1.13606 | 1929 | 49 | c("S100A9", "LTF", "MMP13", "SERPINA1", "PLAT", "PLAUR", "HSPA1A", "GATA3", "CTSD", "CTSL", "FN1") |
| 22 | HALLMARK_COAGULATION | 0.283941 | 0.594798 | -0.310999 | -1.137325 | 1852 | 38 | c("MMP9", "SERPINA1", "PRSS23", "CD9", "PLAT", "COMP", "MMP11") |
| 23 | HALLMARK_KRAS_SIGNALING_DN | 0.268217 | 0.594798 | -0.464774 | -1.191776 | 1545 | 10 | c("CPB1", "BMPR1B", "GPRC5C") |
| 24 | HALLMARK_PANCREAS_BETA_CELLS | 0.284462 | 0.594798 | -0.609367 | -1.185821 | 1524 | 4 | SYT13 |
| 25 | HALLMARK_ALLOGRAFT_REJECTION | 0.297727 | 0.595455 | -0.363497 | -1.142902 | 1833 | 21 | c("MMP9", "CAPG", "INHBA", "TPD52", "TAPBP", "CTSS", "BCL3") |
| 26 | HALLMARK_IL6_JAK_STAT3_SIGNALING | 0.368528 | 0.708708 | 0.311566 | 1.072562 | 1451 | 19 | c("CD36", "IL1R1", "TNFRSF21", "A2M", "LEPR", "JUN", "IFNGR1", "SOCS3") |
| 27 | HALLMARK_FATTY_ACID_METABOLISM | 0.397347 | 0.735828 | 0.267103 | 1.042993 | 1437 | 30 | c("G0S2", "CD36", "ACSL1", "RETSAT", "PTPRG", "HSDL2", "HSD17B11", "DECR1", "ALDH1A1", "ACOX1", "DLD", "ACADVL", "MGLL", "LGALS1", "ECHS1", "EPHX1", "SERINC1") |
| 28 | HALLMARK_MYC_TARGETS_V2 | 0.429086 | 0.766224 | -0.471712 | -1.03852 | 2383 | 6 | c("GNL3", "CBX3", "HSPE1") |
| 29 | HALLMARK_MITOTIC_SPINDLE | 0.470282 | 0.783803 | 0.22708 | 0.996414 | 1534 | 48 | c("SPTBN1", "ITSN1", "BIN1", "SYNPO", "GSN", "EPB41L2", "ABL1", "ARHGEF7", "DST", "AKAP13", "ARHGAP29", "MYO1E", "TRIO", "PKD2", "ROCK1", "HOOK3", "WASF2", "KLC1", "VCL", "NIN", "KIF1B", "UXT") |
| 30 | HALLMARK_UV_RESPONSE_UP | 0.456653 | 0.783803 | -0.30495 | -1.007514 | 2875 | 25 | c("RET", "HMOX1", "EPCAM", "TFRC", "HNRNPU") |
| 31 | HALLMARK_CHOLESTEROL_HOMEOSTASIS | 0.488287 | 0.78756 | 0.276886 | 0.980722 | 1875 | 21 | c("LPL", "PPARG", "SCD", "NFIL3", "PNRC1", "ERRFI1", "ANTXR2", "JAG1") |
| 32 | HALLMARK_APICAL_JUNCTION | 0.621921 | 0.939908 | 0.205166 | 0.912242 | 2019 | 50 | c("CD209", "VWF", "SLIT2", "ARHGEF6", "CLDN5", "EPB41L2", "MYL9", "LIMA1", "PTEN", "MAP3K20", "PECAM1", "NLGN2", "PARVA", "SDC3", "CD34", "AKT3", "TJP1", "MSN", "SIRPA", "SHC1", "VCAN", "THY1", "VCL", "ITGB1", "FBN1", "NF2") |
| 33 | HALLMARK_PI3K_AKT_MTOR_SIGNALING | 0.616386 | 0.939908 | -0.26842 | -0.886823 | 3881 | 25 | c("SLC2A1", "PAK4", "CALR", "PPP1CA", "MAP2K3", "SQSTM1", "ARPC3", "GSK3B", "MYD88", "PFN1") |
| 34 | HALLMARK_GLYCOLYSIS | 0.639138 | 0.939908 | -0.236209 | -0.87966 | 4209 | 41 | c("STC2", "FUT8", "SDC1", "MIF", "STC1", "TGFBI", "PKM", "GPC1", "P4HA1", "CHPF", "TFF3", "VEGFA", "IER3", "IDUA", "QSOX1", "COL5A1", "TALDO1", "AGRN", "COPB2", "ENO1", "GFPT1") |
| 35 | HALLMARK_INTERFERON_ALPHA_RESPONSE | 0.6714 | 0.947255 | 0.268104 | 0.840162 | 2694 | 14 | c("TXNIP", "C1S", "NCOA7", "LPAR6") |
| 36 | HALLMARK_EPITHELIAL_MESENCHYMAL_TRANSITION | 0.733189 | 0.947255 | 0.174688 | 0.875833 | 2027 | 88 | c("SLIT3", "SFRP1", "CCN2", "SLIT2", "TNFAIP3", "FBLN5", "LRP1", "NNMT", "MYL9", "CXCL12", "COL4A2", "TAGLN", "DAB2", "CCN1", "LAMA2", "JUN", "ACTA2", "COL4A1", "GEM", "LAMC1", "DST", "CALD1", "IGFBP4", "DCN", "TPM2", "WIPF1", "TPM4", "SNAI2", "VIM", "GADD45B", "ADAM12", "PRRX1", "SFRP4", "RHOB", "PDGFRB", "LGALS1", "VCAN", "THY1", "ITGB1", "SERPINE1", "COL6A2", "THBS2", "TIMP1", "FBN1", "HTRA1", "MMP2", "GAS1", "BASP1", "DPYSL3", "IGFBP3", "FBLN2", "IL32") |
| 37 | HALLMARK_INFLAMMATORY_RESPONSE | 0.776749 | 0.947255 | 0.208966 | 0.781231 | 2852 | 26 | c("GPC3", "STAB1", "IL1R1", "RGS1", "AHR", "NFKBIA", "KLF6", "AXL", "ABCA1", "MSR1", "CD14", "SERPINE1", "KIF1B", "TIMP1") |
| 38 | HALLMARK_OXIDATIVE_PHOSPHORYLATION | 0.765032 | 0.947255 | -0.203182 | -0.809687 | 5241 | 57 | c("COX6C", "NDUFC2", "ATP1B1", "COX6B1", "ATP6V0B", "COX8A", "NDUFC1", "TCIRG1", "NDUFB2", "FH", "ATP6AP1", "NDUFS6", "ATP6V0C", "IDH2", "COX7C", "UQCRC1", "ECI1", "VDAC3", "ATP5MF", "UQCR11", "NDUFA1", "ATP5MC2", "NDUFAB1", "NDUFB6", "RHOT2", "NDUFA2", "COX6A1", "ATP5PD", "ATP6V1F", "ACAA1", "NDUFA3", "MGST3", "NDUFV1") |
| 39 | HALLMARK_REACTIVE_OXYGEN_SPECIES_PATHWAY | 0.70971 | 0.947255 | 0.249786 | 0.814593 | 2791 | 16 | c("GPX3", "MGST1") |
| 40 | HALLMARK_ANGIOGENESIS | 0.690017 | 0.947255 | 0.26611 | 0.817398 | 2826 | 13 | c("LPL", "TNFRSF21", "NRP1", "JAG1") |
| 41 | HALLMARK_IL2_STAT5_SIGNALING | 0.750924 | 0.947255 | 0.187685 | 0.834513 | 2438 | 50 | c("ITIH5", "DHRS3", "TNFRSF21", "NFIL3", "FGL2", "PLPP1", "NRP1", "WLS", "IFNGR1", "HIPK2", "AHR", "EMP1", "MYO1C", "PRNP", "SWAP70", "KLF6", "MYO1E", "GADD45B", "IRF8", "FAH", "RHOB", "NDRG1") |
| 42 | HALLMARK_PEROXISOME | 0.797537 | 0.949448 | 0.212554 | 0.743821 | 3107 | 20 | c("ACSL1", "ITGB1BP1", "DHRS3", "RETSAT", "HSD17B11", "ALDH1A1", "ACOX1") |
| 43 | HALLMARK_WNT_BETA_CATENIN_SIGNALING | 0.862964 | 0.976286 | -0.314456 | -0.655802 | 4722 | 5 | CTNNB1 |
| 44 | HALLMARK_TGF_BETA_SIGNALING | 0.929066 | 0.976286 | -0.194057 | -0.594007 | 5631 | 19 | CDH1 |
| 45 | HALLMARK_G2M_CHECKPOINT | 0.950302 | 0.976286 | -0.172924 | -0.571317 | 5984 | 25 | c("TOP2A", "CCND1", "HNRNPU", "BUB3", "BCL3", "SLC12A2", "RBM14", "SMC4") |
| 46 | HALLMARK_APOPTOSIS | 0.934615 | 0.976286 | -0.167154 | -0.651599 | 6317 | 51 | c("HMOX1", "TOP2A", "PLAT", "CCND1", "ERBB3", "CTNNB1", "RARA", "GSR") |
| 47 | HALLMARK_NOTCH_SIGNALING | 0.960316 | 0.976286 | -0.219599 | -0.529749 | 5468 | 8 | c("CCND1", "LFNG") |
| 48 | HALLMARK_PROTEIN_SECRETION | 0.976286 | 0.976286 | -0.153792 | -0.496809 | 6051 | 23 | c("TPD52", "ANP32E", "TOM1L1", "M6PR", "COPB2", "MON2", "ARF1", "PPT1", "NAPA", "CD63") |
| 49 | HALLMARK_HEDGEHOG_SIGNALING | 0.918753 | 0.976286 | -0.237446 | -0.592006 | 5246 | 9 | c("CELSR1", "ADGRG1", "VEGFA") |

| | | | | | | | | |
|---|---|---|---|---|---|---|---|---|
| 50 | HALLMARK_P53_PATHWAY | 0.917262 | 0.976286 | -0.172868 | -0.664062 | 6163 | 47 | c("HMOX1", "IFI30", "TM4SF1", "SDC1", "DCXR", "CTSD", "NUPR1", "RAP2B", "EPS8L2", "TRAF4", "IER3", "TOB1", "HEXIM1", "SLC3A2", "VAMP8", "FUCA1", "S100A4") |

| Cluster 4 | pathway | pval | padj | ES | NES | nMoreExtreme | size | leadingEdge |
|---|---|---|---|---|---|---|---|---|
| 1 | HALLMARK_TNFA_SIGNALING_VIA_NFKB | 0.000585 | 0.004611 | -0.608979 | -3.205411 | 0 | 76 | c("FOS", "BTG2", "CCN1", "GEM", "DUSP1", "NR4A1", "EGR1", "FOSB", "PHLDA1", "ZFP36", "TGIF1", "CD44", "CEBPD", "MAP3K8", "IER3", "PLAU", "SGK1", "JUN", "CEBPB", "KLF6", "BTG1", "IER2", "NFKBIA", "JUNB", "MCL1", "BHLHE40", "SOD2", "SOCS3", "DUSP4", "BCL3", "B4GALT5", "EHD1", "PLAUR", "FOSL2", "ID2", "HES1", "PLPP3", "KLF2", "SAT1", "NINJ1") |
| 2 | HALLMARK_HYPOXIA | 0.0006 | 0.004611 | -0.393587 | -2.089919 | 0 | 80 | c("CA12", "FOS", "CCN1", "HMOX1", "DUSP1", "ZFP36", "MAP3K1", "STC1", "ANXA2", "IER3", "JUN", "KLF6", "BTG1", "SLC2A1", "BHLHE40", "MIF", "PIM1", "PLAUR", "CCN2", "CITED2", "ENO1", "FOSL2", "LOX", "GAA", "SDC3", "COL5A1") |
| 3 | HALLMARK_IL6_JAK_STAT3_SIGNALING | 0.000378 | 0.004611 | -0.640624 | -2.519075 | 0 | 23 | c("HMOX1", "CD36", "CD44", "MAP3K8", "JUN", "TNFRSF12A", "CD9", "SOCS3", "PIM1", "STAT3", "TNFRSF1A", "IL1R1", "IL13RA1", "PTPN1") |
| 4 | HALLMARK_ESTROGEN_RESPONSE_LATE | 0.000639 | 0.004611 | -0.406355 | -2.213153 | 0 | 94 | c("SERPINA3", "LTF", "TFF1", "S100A9", "PRLR", "CA12", "FOS", "SERPINA1", "PERP", "ZFP36", "PAPSS2", "HSPB8", "CHPT1", "CD44", "SGK1", "UGDH", "CD9", "NBL1", "BLVRB", "CXCL12", "FARP1", "PDCD4", "MDK", "ELOVL5", "KRT19", "MAPT", "ID2", "SEMA3B", "RAB31", "MYB", "ITPK1", "CDH1", "PLAAT3", "IDH2", "ETFB") |
| 5 | HALLMARK_COMPLEMENT | 0.00051 | 0.004611 | -0.518988 | -2.5394 | 0 | 56 | c("LTF", "S100A9", "SERPINA1", "LGALS3", "CD55", "CD36", "C1S", "CEBPB", "CTSD", "C1R", "LGMN", "CTSL", "APOC1", "CTSB", "MMP14", "PIM1", "EHD1", "PLAUR", "C3", "CTSS", "TIMP2") |
| 6 | HALLMARK_EPITHELIAL_MESENCHYMAL_TRANSITION | 0.000646 | 0.004611 | -0.578406 | -3.159838 | 0 | 96 | c("ADAM12", "CCN1", "ELN", "GEM", "MGP", "CAPG", "CTHRC1", "CD44", "MMP2", "THBS2", "JUN", "POSTN", "TNFRSF12A", "FBLN1", "FAP", "SDC1", "CXCL12", "MMP14", "FBLN2", "COL3A1", "IGFBP2", "PLAUR", "VCAN", "CCN2", "ACTA2", "COL6A3", "NNMT", "LGALS1", "COL5A2", "DPYSL3", "HTRA1", "MYL9", "ID2", "LOX", "COL11A1", "LOXL1", "SFRP4", "SPP1", "COL6A2", "COL5A1", "LRP1", "COL1A2", "FBN1", "TAGLN", "COL16A1", "COL1A1", "SAT1", "DCN", "TPM2", "FUCA1", "LOXL2", "ITGB5", "DAB2", "CALU", "TPM1", "SERPINH1", "FLNA", "PRRX1", "MXRA5", "VIM", "SERPINE1", "EMP3", "PMEPA1", "ITGB1") |
| 7 | HALLMARK_P53_PATHWAY | 0.000548 | 0.004611 | -0.437376 | -2.242179 | 0 | 67 | c("NUPR1", "FOS", "BTG2", "HMOX1", "PERP", "S100A10", "IFI30", "IER3", "JUN", "RAP2B", "EPHX1", "CTSD", "BTG1", "ZFP36L1", "SDC1", "SLC3A2", "TM4SF1") |
| 8 | HALLMARK_REACTIVE_OXYGEN_SPECIES_PATHWAY | 0.00107 | 0.00535 | -0.632691 | -2.350544 | 2 | 19 | c("MPO", "NQO1", "JUNB", "SOD2", "FTL", "LSP1", "CAT", "GPX4", "GPX3", "EGLN2", "ATOX1") |
| 9 | HALLMARK_UV_RESPONSE_UP | 0.001021 | 0.00535 | -0.375088 | -1.835297 | 1 | 56 | c("FOS", "BTG2", "HMOX1", "NR4A1", "FOSB", "TFRC", "EPHX1", "BTG1", "NFKBIA", "JUNB", "SOD2", "MMP14", "UROD", "IGFBP2") |
| 10 | HALLMARK_COAGULATION | 0.000944 | 0.00535 | -0.458974 | -2.135825 | 1 | 45 | c("SERPINA1", "MMP11", "C1S", "MMP2", "PLAU", "C1R", "CD9", "LGMN", "CTSK", "APOC1", "CTSB", "MMP14", "C3", "HTRA1", "CFD", "LRP1", "FBN1", "CAPN2", "ANXA1") |
| 11 | HALLMARK_HEME_METABOLISM | 0.001217 | 0.00553 | -0.350136 | -1.862874 | 1 | 81 | c("SLC4A1", "BTG2", "HBD", "SLC25A37", "TFRC", "SNCA", "GLRX5", "SLC2A1", "BLVRB", "ELL2", "CTSB", "MPP1", "UROD", "SLC30A1", "UCP2", "C3", "CAT", "HDGF", "HEBP1", "MGST3", "GYPC", "UBAC1", "HBB", "TNS1", "PGLS", "ADIPOR1", "CCND3", "CIR1", "FBXO7", "MAP2K3", "FOXO3") |
| 12 | HALLMARK_ESTROGEN_RESPONSE_EARLY | 0.001962 | 0.008175 | -0.325505 | -1.780415 | 2 | 97 | c("TFF1", "SEC14L2", "CA12", "CELSR1", "FOS", "GAB2", "PAPSS2", "HSPB8", "CHPT1", "CD44", "KRT8", "FHL2", "SLC2A1", "BHLHE40", "NBL1", "BLVRB", "RHOBTB3", "CXCL12", "FARP1", "FASN", "ELOVL5", "KRT19", "UGCG", "LRIG1", "MAPT", "SH3BP5", "SEMA3B", "RAB31", "HES1", "MYB", "ITPK1") |
| 13 | HALLMARK_APOPTOSIS | 0.003331 | 0.012813 | -0.340045 | -1.769695 | 5 | 71 | c("LGALS3", "BTG2", "HMOX1", "CD44", "MMP2", "IER3", "NEDD9", "JUN", "TNFRSF12A", "MCL1", "SOD2", "PDCD4", "TSPO", "DAP3", "TIMP2", "GPX4", "GPX3", "SAT1", "PPT1", "DCN", "DPYD", "ANXA1", "IGF2R", "APP", "LMNA") |
| 14 | HALLMARK_MTORC1_SIGNALING | 0.004386 | 0.015664 | -0.324463 | -1.680706 | 7 | 69 | c("EGLN3", "NUPR1", "BTG2", "IFI30", "STC1", "TFRC", "CORO1A", "CD9", "LGMN", "SLC2A1", "BHLHE40", "ELOVL5", "ENO1", "M6PR", "ITGB2", "PSMB5", "PIK3R3") |
| 15 | HALLMARK_XENOBIOTIC_METABOLISM | 0.005578 | 0.018594 | -0.356427 | -1.725408 | 10 | 54 | c("HMOX1", "NQO1", "PAPSS2", "CD36", "PDK4", "UGDH", "EPHX1", "FBLN1", "APOE", "BLVRB", "ELOVL5", "PGD", "CAT", "ID2", "TNFRSF1A", "ALDH2", "PGRMC1", "DDT", "NINJ1", "IL1R1", "DHRS7") |
| 16 | HALLMARK_BILE_ACID_METABOLISM | 0.008115 | 0.025358 | 0.517391 | 1.719984 | 59 | 24 | c("SERPINA6", "RBP1", "ABCA2", "ABCA5", "ISOC1", "SLC29A1", "ABCA3", "AR", "EPHX2", "EFHC1", "PFKM", "FADS2") |

| | | | | | | | | |
|---|---|---|---|---|---|---|---|---|
| 17 | HALLMARK_UV_RESPONSE_DN | 0.015735 | 0.04628 | -0.299493 | -1.529961 | 28 | 66 | c("INPP4B", "EFEMP1", "CCN1", "DUSP1", "RUNX1", "ANXA2", "RBPMS", "FHL2", "BHLHE40", "COL3A1", "CITED2", "COL5A2", "COL11A1", "PLPP3", "COL1A2", "COL1A1", "PIK3R3", "ATP2B4", "DAB2", "AKT3", "MTA1") |
| 18 | HALLMARK_CHOLESTEROL_HOMEOSTASIS | 0.028493 | 0.079148 | -0.403386 | -1.638651 | 72 | 26 | c("LGALS3", "S100A11", "TNFRSF12A", "CD9", "LGMN", "FASN", "PLAUR", "SEMA3B") |
| 19 | HALLMARK_INTERFERON_GAMMA_RESPONSE | 0.03286 | 0.086473 | -0.343532 | -1.539943 | 73 | 39 | c("IFI30", "ARID5B", "C1S", "BTG1", "NFKBIA", "C1R", "SOD2", "SOCS3", "PIM1", "STAT3") |
| 20 | HALLMARK_ALLOGRAFT_REJECTION | 0.041667 | 0.104167 | -0.36355 | -1.529099 | 102 | 30 | c("CAPG", "BCL3", "SPI1", "SRGN", "CTSS", "ITGB2", "NME1", "CCND3", "HCLS1", "CD47", "FLNA") |
| 21 | HALLMARK_ANGIOGENESIS | 0.044266 | 0.105396 | -0.454746 | -1.591122 | 126 | 16 | c("STC1", "POSTN", "COL3A1", "VCAN", "COL5A2", "SPP1") |
| 22 | HALLMARK_KRAS_SIGNALING_UP | 0.054251 | 0.123299 | -0.279432 | -1.390694 | 103 | 60 | c("SERPINA3", "APOD", "MMP11", "MAP3K1", "GPNMB", "PLAU", "IKZF1", "PLAUR", "SPARCL1", "MAFB", "CTSS", "ID2", "SEMA3B", "GYPC", "SPP1", "ITGB2", "NIN", "FUCA1", "CPE") |
| 23 | HALLMARK_INFLAMMATORY_RESPONSE | 0.101119 | 0.219824 | -0.324694 | -1.355762 | 252 | 29 | c("BTG2", "CD55", "KLF6", "NFKBIA", "MMP14", "PLAUR") |
| 24 | HALLMARK_FATTY_ACID_METABOLISM | 0.11101 | 0.228448 | -0.286731 | -1.312782 | 242 | 42 | c("CD36", "S100A10", "UGDH", "EPHX1", "FASN", "UROD", "MIF", "ELOVL5", "LGALS1", "GLUL", "APEX1", "HSD17B4", "FH") |
| 25 | HALLMARK_GLYCOLYSIS | 0.114224 | 0.228448 | -0.24884 | -1.263332 | 211 | 65 | c("EGLN3", "PYGL", "CD44", "STC1", "IER3", "SDC1", "MIF", "VCAN", "FUT8", "CITED2", "ENO1", "SDC3", "COL5A1", "DCN", "CHPF", "DDIT4", "PGLS", "PAM", "IL13RA1") |
| 26 | HALLMARK_WNT_BETA_CATENIN_SIGNALING | 0.157169 | 0.302249 | 0.456215 | 1.307985 | 1105 | 14 | c("LEF1", "HDAC5", "HEY2", "HDAC2", "CTNNB1", "KAT2A", "AXIN1", "TP53", "JAG1") |
| 27 | HALLMARK_KRAS_SIGNALING_DN | 0.163379 | 0.302553 | 0.413353 | 1.291378 | 1175 | 19 | c("CPB1", "BMPR1B", "RGS11", "LFNG", "THNSL2", "MFSD6", "FGFR3") |
| 28 | HALLMARK_ADIPOGENESIS | 0.242092 | 0.417401 | -0.210876 | -1.121951 | 397 | 81 | c("CD36", "HSPB8", "APOE", "PDCD4", "UCP2", "SPARCL1", "STOM", "C3", "FABP4", "CAT", "GPX4", "UQCR10", "GPX3", "MGST3", "ALDH2", "DDT", "TKT", "ETFB", "VEGFB", "DHRS7", "GPAT4", "PFKFB3", "CCNG2", "COX8A") |
| 29 | HALLMARK_APICAL_SURFACE | 0.233916 | 0.417401 | -0.378014 | -1.20584 | 708 | 12 | c("SULF2", "GSTM3", "PLAUR") |
| 30 | HALLMARK_SPERMATOGENESIS | 0.256582 | 0.427636 | 0.392295 | 1.187844 | 1841 | 17 | c("GAD1", "ACE", "PIAS2", "MAP7", "LPIN1", "STRBP", "BRAF", "SLC12A2", "PRKAR2A", "IFT88") |
| 31 | HALLMARK_G2M_CHECKPOINT | 0.276291 | 0.44563 | 0.299203 | 1.149887 | 2177 | 45 | c("GINS2", "SMC2", "CDC25B", "RAD21", "XPO1", "SMAD3", "MARCKS", "SS18", "PDS5B", "YTHDC1", "CUL3", "SLC12A2", "KPNB1", "CCND1", "TOP2A", "RBM14", "SNRPD1", "CUL4A", "NCL", "PAFAH1B1", "SFPQ", "HNRNPD", "HUS1", "UPF1", "PURA", "DR1", "RAD23B", "ATRX", "MCM2", "HSPA8", "SLC38A1") |
| 32 | HALLMARK_PANCREAS_BETA_CELLS | 0.36369 | 0.568266 | 0.450629 | 1.094121 | 2463 | 8 | c("SYT13", "STXBP1") |
| 33 | HALLMARK_IL2_STAT5_SIGNALING | 0.449817 | 0.681542 | 0.259412 | 1.018038 | 3571 | 50 | c("APLP1", "IKZF2", "IGF1R", "BCL2", "NFKBIZ", "IRF6", "MAPKAPK2", "SLC39A8") |
| 34 | HALLMARK_ANDROGEN_RESPONSE | 0.497337 | 0.731377 | -0.2084 | -0.979076 | 1026 | 47 | c("INPP4B", "ARID5B", "SGK1", "KRT8", "ELL2", "ELOVL5", "KRT19", "SLC38A2", "ACTN1", "SAT1", "UAP1", "MAF", "CCND3") |
| 35 | HALLMARK_OXIDATIVE_PHOSPHORYLATION | 0.539759 | 0.771084 | -0.180142 | -0.963582 | 895 | 83 | c("MAOB", "NDUFC2", "ATP1B1", "PDK4", "TCIRG1", "GPX4", "UQCR10", "MGST3", "OXA1L", "SLC25A6", "IDH2", "NDUFA2", "ETFB", "FH", "NDUFS6", "ATP5F1E", "ATP5PD", "COX8A", "ATP5MG") |
| 36 | HALLMARK_MYOGENESIS | 0.566715 | 0.787104 | 0.221429 | 0.941243 | 4705 | 77 | c("GNAO1", "PGAM2", "DTNA", "FKBP1B", "MAPRE3", "CACNA1H", "AKT2", "CAMK2B", "HDAC5", "KCNH2", "DAPK2", "LPIN1", "TSC2", "TNNT1", "AK1", "COL4A2") |
| 37 | HALLMARK_INTERFERON_ALPHA_RESPONSE | 0.608773 | 0.822666 | -0.237872 | -0.883733 | 1706 | 19 | c("IFI30", "C1S") |
| 38 | HALLMARK_NOTCH_SIGNALING | 0.670635 | 0.882415 | 0.309402 | 0.848631 | 4674 | 12 | c("LFNG", "KAT2A", "NOTCH3", "ARRB1", "JAG1", "CCND1", "FBXW11") |
| 39 | HALLMARK_HEDGEHOG_SIGNALING | 0.70146 | 0.899308 | 0.292658 | 0.821902 | 4898 | 13 | c("UNC5C", "HEY2", "DPYSL2", "ACHE", "LDB1") |
| 40 | HALLMARK_E2F_TARGETS | 0.739856 | 0.912053 | 0.212822 | 0.821891 | 5852 | 46 | c("RBBP7", "CDC25B", "RAD21", "RAD50", "SMC3", "XPO1", "CBX5", "CCP110", "NUP205", "DUT", "MSH2", "PDS5B", "CNOT9", "ORC2", "WDR90", "TP53", "PAN2", "TOP2A", "DCTPP1", "NAA38", "HNRNPD", "HUS1") |
| 41 | HALLMARK_MYC_TARGETS_V2 | 0.747883 | 0.912053 | -0.267536 | -0.769391 | 2384 | 9 | c("PA2G4", "FARSA", "MCM4", "GNL3", "HK2", "SORD", "HSPE1", "DCTPP1", "HSPD1") |
| 42 | HALLMARK_PROTEIN_SECRETION | 0.844575 | 0.99038 | 0.193507 | 0.735914 | 6623 | 43 | c("ICA1", "TSG101", "ARFGEF1", "LMAN1", "AP2B1", "SCAMP1", "VPS4B", "ARFIP1", "TOM1L1", "RAB2A", "ATP6V1H", "RAB14", "YIPF6", "STX16", "NAPA", "ADAM10", "CLCN3") |
| 43 | HALLMARK_PEROXISOME | 0.851726 | 0.99038 | 0.198552 | 0.722146 | 6536 | 35 | c("SERPINA6", "ABCB9", "ISOC1", "SLC25A4", "EPHX2", "VPS4B", "ABCC5", "MSH2", "ERCC3", "HRAS", "SCP2", "ACSL1", "CRAT", "PEX2", "TOP2A") |

| | | | | | | | | |
|---|---|---|---|---|---|---|---|---|
| 44 | HALLMARK_MITOTIC_SPINDLE | 0.92171 | 0.999826 | -0.139147 | -0.72416 | 1659 | 71 | c("ARF6", "NEDD9", "FARP1", "PREX1", "SHROOM1", "NIN", "PALLD", "FLNA", "TRIO", "SEPTIN9", "NET1", "RANBP9", "CDC42BPA", "ARL8A", "KLC1", "SUN2", "GSN", "PKD2", "CCDC88A", "PXN", "CSNK1D", "TUBGCP2", "RASA1", "EZR", "ARHGAP29", "SMC1A", "KIF5B", "ARHGEF2", "ARHGAP4", "RAPGEF6", "NUSAP1", "DYNLL2", "WASF2", "NUMA1", "CD2AP", "SPTAN1", "ABR", "EPB41L2", "KIF3B", "DOCK4", "MYH10", "ATG4B", "UXT", "NF1", "LRPPRC", "PAFAH1B1", "PCM1", "CNTROB", "FLNB", "TOP2A", "ARHGEF7", "EPB41", "ARHGEF12", "YWHAE", "TAOK2", "CKAP5", "RABGAP1", "CTTN", "RICTOR", "SPTBN1", "LATS1", "CYTH2", "ROCK1", "RHOT2", "MARCKS", "SMC3", "RAPGEF5", "PCGF5", "RAB3GAP1", "ARFIP2", "ARFGEF1") |
| 45 | HALLMARK_TGF_BETA_SIGNALING | 0.891287 | 0.999826 | -0.162774 | -0.679663 | 2229 | 29 | c("TGIF1", "JUNB", "ID2", "RAB31", "CDH1") |
| 46 | HALLMARK_DNA_REPAIR | 0.964533 | 0.999826 | 0.150783 | 0.587415 | 7668 | 48 | c("TSG101", "AK1", "ADCY6", "DUT", "MPG", "POLR1D", "GTF2H5", "ERCC5", "ARL6IP1", "NELFB", "ERCC3", "DGUOK", "BCAM", "COX17", "NME3", "CANT1", "POLB", "TP53", "NUDT9") |
| 47 | HALLMARK_APICAL_JUNCTION | 0.973262 | 0.999826 | -0.122358 | -0.649363 | 1637 | 79 | c("MMP2", "MDK", "ZYX", "VCAN", "CNN2", "MYL9", "CERCAM", "CTNNA1", "SDC3", "CDH1", "ACTN1", "FBN1", "COL16A1", "PIK3R3", "CD99", "AKT3", "THBS3") |
| 48 | HALLMARK_PI3K_AKT_MTOR_SIGNALING | 0.958036 | 0.999826 | -0.136604 | -0.616649 | 2145 | 40 | c("VAV3", "SLC2A1", "TNFRSF1A", "PIK3R3", "ARPC3", "MAP2K3", "SQSTM1", "PAK4", "MYD88", "CDKN1B", "STAT2") |
| 49 | HALLMARK_MYC_TARGETS_V1 | 0.979829 | 0.999826 | 0.137334 | 0.556396 | 7917 | 59 | c("CCT4", "TXNL4A", "HSP90AB1", "XPO1", "HDAC2", "DUT", "COPS5", "SRSF3", "HSPD1", "EIF4G2", "SET", "KPNB1", "HDDC2", "ORC2", "YWHAE", "SSBP1", "CYC1", "SNRPD1", "PSMA4", "HNRNPD", "HSPE1", "YWHAQ", "IFRD1", "CNBP", "NDUFAB1", "IMPDH2", "EIF3J", "RAD23B", "PSMD3", "MCM2", "RRM1", "EIF4A1", "CUL1", "DHX15", "PSMB2", "GNL3", "SMARCC1", "RNPS1", "HNRNPA2B1", "DEK") |
| 50 | HALLMARK_UNFOLDED_PROTEIN_RESPONSE | 0.999874 | 0.999874 | 0.077643 | 0.302479 | 7949 | 48 | c("TUBB2A", "DNAJA4", "YWHAZ", "DCTN1", "CXXC1", "FUS", "PARN", "HSPA5", "LSM4", "EIF4A2", "SPCS1", "CNOT4", "DNAJC3", "ALDH18A1", "BAG3", "IMP3", "CNOT6", "MTREX", "EIF4A1", "TATDN2", "HSPA9", "SRPRB", "KHSRP", "EXOSC10", "MTHFD2", "VEGFA", "CALR", "KIF5B") |

| Cluster 5 | pathway | pval | padj | ES | NES | nMoreExtreme | size | leadingEdge |
|---|---|---|---|---|---|---|---|---|
| 1 | HALLMARK_TNFA_SIGNALING_VIA_NFKB | 0.00012 | 0.002999 | 0.526699 | 2.430503 | 0 | 87 | c("FOSB", "FOS", "NR4A1", "NR4A3", "EGR2", "DUSP4", "EGR1", "DUSP1", "IER2", "SERPINE1", "NR4A2", "TNFAIP6", "GEM", "JUNB", "JUN", "GFPT2", "CDKN1A", "MAFF", "SDC4", "BTG2", "TRIB1", "GADD45B", "KLF2", "IER3", "KLF4", "EGR3", "ZFP36", "JAG1", "TGIF1", "SOCS3", "SPHK1", "PPP1R15A", "CCN1", "GADD45A", "KLF9", "PER1", "ZC3H12A", "TRIP10", "KLF6", "RHOB", "TIPARP", "NINJ1", "PDLIM5", "HES1") |
| 2 | HALLMARK_EPITHELIAL_MESENCHYMAL_TRANSITIO N | 0.000115 | 0.002999 | 0.470616 | 2.314681 | 0 | 127 | c("ELN", "ABI3BP", "LAMA2", "LAMA3", "COL16A1", "SFRP4", "CTHRC1", "SERPINE1", "COL6A3", "GPC1", "TGFBR3", "GEM", "FBLN1", "COL3A1", "JUN", "GREM1", "FBLN2", "MMP2", "COL1A1", "FBLN5", "MXRA5", "COL1A2", "TPM1", "HTRA1", "CAP2", "PDGFRB", "SDC4", "FAS", "FSTL3", "FAP", "MMP14", "DPYSL3", "GADD45B", "FSTL1", "PCOLCE", "SLIT2", "COL5A2", "MYL9", "POSTN", "BMP1", "ITGB5", "SPARC", "DCN", "FBN1", "PRRX1", "ITGAV", "LAMC1", "CCN1", "COL6A2", "GADD45A", "IGFBP2", "THBS2", "EDIL3", "ECM2", "SLIT3", "CCN2", □ "RHOB", "TNFRSF12A", "IGFBP4", "FERMT2", "CALD1", "COL5A1", "SERPINE2") |
| 3 | HALLMARK_ESTROGEN_RESPONSE_LATE | 0.000559 | 0.006991 | -0.422651 | -2.254442 | 0 | 73 | c("S100A9", "SERPINA1", "PGR", "LTF", "TOP2A", "CDH1", "RET", "SLC9A3R1", "COX6C", "PRSS23", "TFF3", "MAPT", "UGDH", "LLGL2", "ABHD2", "BLVRB", "DCXR", "PAPSS2", "TOB1", "PRLR", "CCND1", "XBP1", "KRT19", "IDH2", "RABEP1") |
| 4 | HALLMARK_HEME_METABOLISM | 0.000508 | 0.006991 | -0.474562 | -2.382358 | 0 | 57 | c("HBD", "HBB", "SLC4A1", "ACP5", "SLC2A1", "SNCA", "CTSB", "SLC25A37", "MPP1", "TFRC", "BLVRB", "VEZF1", "NCOA4", "HAGH", "ELL2", "NFE2L1") |
| 5 | HALLMARK_P53_PATHWAY | 0.00132 | 0.013202 | 0.3864 | 1.776994 | 10 | 85 | c("FOS", "INHBB", "JUN", "ZMAT3", "PLK3", "CDKN1A", "AEN", "FAS", "BTG2", "TSPYL2", "DDB2", "RETSAT", "IER3", "PHLDA3", "KLF4", "KIF13B", "PITPNC1", "SPHK1", "POLH", "CD82", "SLC35D1", "CCND2", "PPP1R15A", "TAX1BP3", "SESN1", "GADD45A", "MXD4", "PERP", "STOM", "SLC19A2", "NOTCH1", "ISCU", "NINJ1", "TM7SF3", "XPC", "BLCAP", "CSRNP2", "CCNG1", "PLK2", "ZBTB16", "TXNIP", "PTPN14", "PLXNB2", "RNF19B") |
| 6 | HALLMARK_COMPLEMENT | 0.002577 | 0.021478 | -0.390876 | -1.973067 | 4 | 59 | c("S100A9", "SERPINA1", "LTF", "MMP13", "APOC1", "CTSB", "CTSS", "CTSD", "CTSL", "LIPA", "CD55", "FCER1G", "CD36", "LTA4H", "GRB2", "PPP4C", "PLAT") |
| 7 | HALLMARK_ESTROGEN_RESPONSE_EARLY | 0.004292 | 0.027628 | -0.306836 | -1.719159 | 6 | 89 | c("PGR", "ADCY1", "RET", "SLC39A6", "RHOBTB3", "SLC9A3R1", "SLC2A1", "FASN", "SLC7A2", "PRSS23", "GAB2", "TFF3", "MAPT", "ABHD2", "BLVRB", "CANT1", "PAPSS2", "CLDN7", "GJA1", "TOB1", "AKAP1", "CCND1", "XBP1", "KRT19") |
| 8 | HALLMARK_MTORC1_SIGNALING | 0.00442 | 0.027628 | -0.390759 | -1.877856 | 8 | 49 | c("SCD", "IFI30", "SLC9A3R1", "SLC2A1", "CORO1A", "TFRC", "ITGB2", "NUPR1", "EGLN3", "CANX", "SLC1A5", "ACACA", "ENO1", "XBP1", "DDIT4", "LTA4H", "SQSTM1", "PSMD13", "PPIA") |
| 9 | HALLMARK_UV_RESPONSE_DN | 0.008393 | 0.046629 | 0.354078 | 1.596419 | 68 | 76 | c("DUSP1", "SERPINE1", "TGFBR3", "COL3A1", "COL1A1", "FBLN5", "COL1A2", "CAP2", "PDGFRB", "LPAR1", "KCNMA1", "PTPRM", "ID1", "DLC1", "COL5A2", "CAV1", "LAMC1", "CCN1", "TJP1", "RBPMS", "MTA1", "PDLIM5", "ADD3", "PIK3R3", "NR1D2", "DDAH1", "MGMT", "CDKN1B", "RUNX1", "NOTCH2", "PRDM2", "NRP1", "APBB2", "MIOS", "SNAI2", "TGFBR2", "MAP1B", "SMAD7", "ATXN1") |
| 10 | HALLMARK_MYOGENESIS | 0.010675 | 0.053374 | 0.360581 | 1.593782 | 86 | 68 | c("PPP1R3C", "LAMA2", "COL6A3", "IGF1", "COL3A1", "COL1A1", "CDKN1A", "SSPN", "PTGIS", "EFS", "AGRN", "ABLIM1", "SOD3", "GADD45B", "GSN", "PVALB", "PDLIM7", "ITGB5", "SPHK1", "SPARC", "COL6A2", "COX7A1", "NOTCH1", "WWTR1", "AEBP1", "COL15A1", "IGFBP7", "SPTAN1", "SORBS3", "ITGB1", "PLXNB2", "SVIL", "SMTN", "PTP4A3", "CLU", "PFKM", "APP") |
| 11 | HALLMARK_ALLOGRAFT_REJECTION | 0.030155 | 0.137069 | -0.360507 | -1.593525 | 67 | 36 | c("MMP9", "CAPG", "CTSS", "ITGB2", "SPI1", "SRGN", "NME1", "TPD52") |
| 12 | HALLMARK_HYPOXIA | 0.037855 | 0.157729 | 0.318367 | 1.44247 | 311 | 78 | c("FOS", "PPP1R3C", "DUSP1", "SERPINE1", "GPC1", "TGFB3", "JUN", "CDKN1A", "AKAP12", "CAVIN3", "MAFF", "SDC4", "IER3", "KDELR3", "ZFP36", "PAM", "CAV1", "TPST2", "DCN", "PPP1R15A", "CCN1", "KLF6", "CCN2", "HDLBP", "COL5A1", "TIPARP", "SDC3", "P4HA2", "PDGFB", "XPNPEP1", "NDST1", "STC1", "CAVIN1", "CDKN1B") |
| 13 | HALLMARK_E2F_TARGETS | 0.043385 | 0.158028 | -0.391686 | -1.591593 | 100 | 27 | c("TOP2A", "HMGB2", "TFRC", "ANP32E", "NME1", "SMC4", "SRSF2", "TUBB") |
| 14 | HALLMARK_REACTIVE_OXYGEN_SPECIES_PATHWA Y | 0.044248 | 0.158028 | -0.50054 | -1.624044 | 114 | 13 | c("MPO", "FTL", "NQO1", "GPX4", "PRDX2") |
| 15 | HALLMARK_INFLAMMATORY_RESPONSE | 0.060844 | 0.202813 | 0.370502 | 1.436124 | 472 | 37 | c("SERPINE1", "TNFAIP6", "CDKN1A", "LPAR1", "BTG2", "MMP14", "SPHK1", "CD82", "RNF144B", "SCN1B", "KLF6", "AXL", "HPN", "PCDH7") |

| | | | | | | | | |
|---|---|---|---|---|---|---|---|---|
| 16 | HALLMARK_NOTCH_SIGNALING | 0.09358 | 0.292437 | 0.484797 | 1.400666 | 687 | 12 | c("FZD1", "NOTCH3", "JAG1", "NOTCH1", "HES1", "ARRB1", "NOTCH2", "TCF7L2") |
| 17 | HALLMARK_BILE_ACID_METABOLISM | 0.195096 | 0.541934 | 0.379257 | 1.249271 | 1471 | 19 | c("CYP7B1", "RETSAT", "ALDH1A1", "DHCR24", "ABCD3", "PRDX5", "PFKM", "ATXN1", "AR", "SCP2", "ISOC1", "HSD17B11", "RXRA", "ABCA2", "ALDH9A1", "HSD17B4", "SLC23A2") |
| 18 | HALLMARK_KRAS_SIGNALING_UP | 0.187008 | 0.541934 | 0.290721 | 1.228129 | 1496 | 54 | c("SPARCL1", "GFPT2", "AKAP12", "GADD45G", "ITGBL1", "ADGRA2", "TRIB2", "TRIB1", "MAFB", "KLF4", "GUCY1A1", "SPON1", "CCND2", "PRRX1", "PPP1R15A", "SCN1B", "PLVAP", "CFH", "ANO1", "SPRY2") |
| 19 | HALLMARK_WNT_BETA_CATENIN_SIGNALING | 0.254878 | 0.620384 | 0.421784 | 1.186053 | 1867 | 11 | c("FZD1", "JAG1", "CCND2", "NOTCH1") |
| 20 | HALLMARK_TGF_BETA_SIGNALING | 0.290247 | 0.620384 | 0.302434 | 1.149084 | 2240 | 34 | c("SERPINE1", "JUNB", "ID3", "LTBP2", "ID1", "TGIF1", "SKI", "PPP1R15A", "TJP1", "BMPR2", "WWTR1", "KLF10", "SMURF1", "SMAD7", "HIPK2", "SKIL", "ID2", "PMEPA1", "PPM1A", "ENG", "THBS1", "CDK9") |
| 21 | HALLMARK_APOPTOSIS | 0.242866 | 0.620384 | 0.263139 | 1.166286 | 1982 | 69 | c("TGFBR3", "JUN", "MMP2", "CDKN1A", "IGFBP6", "PDGFRB", "FAS", "BTG2", "GADD45B", "GSN", "RETSAT", "IER3", "EGR3", "CAV1", "DCN", "CCND2", "GADD45A", "RHOB", "TNFRSF12A", "SPTAN1", "LMNA", "TXNIP", "MGMT", "CDKN1B", "BCL2L2", "RELA", "TNFSF10", "CLU", "PDCD4", "SMAD7", "APP", "LUM", "ANKH", "ATF3", "VDAC2", "PEA15", "PTK2", "TIMP2") |
| 22 | HALLMARK_ANDROGEN_RESPONSE | 0.290015 | 0.620384 | -0.229784 | -1.118798 | 577 | 52 | c("BMPR1B", "SCD", "ABHD2", "TPD52", "AZGP1", "NCOA4", "CCND1", "SPDEF", "B2M", "KRT19", "ELL2", "SMS") |
| 23 | HALLMARK_XENOBIOTIC_METABOLISM | 0.271874 | 0.620384 | 0.27608 | 1.153201 | 2171 | 51 | c("PTGES", "SERPINE1", "IGF1", "FBLN1", "CSAD", "SLC12A4", "NDRG2", "FAS", "RETSAT", "SLC35D1", "DHRS1", "ALDH2", "IGFBP4", "DHRS7", "NINJ1", "PDLIM5") |
| 24 | HALLMARK_COAGULATION | 0.297784 | 0.620384 | 0.271154 | 1.132622 | 2378 | 51 | c("SERPINE1", "MMP2", "F2RL2", "HTRA1", "MAFF", "GNG12", "MMP14", "GSN", "C1S", "CSRP1", "BMP1", "SPARC", "FBN1", "CFH", "SERPING1", "WDR1", "C1R", "HPN", "A2M", "ISCU", "CAPN2", "CTSK", "LRP1", "PDGFB", "C3", "CTSO", "VWF") |
| 25 | HALLMARK_ADIPOGENESIS | 0.340226 | 0.657041 | 0.241303 | 1.087955 | 2796 | 76 | c("CYP4B1", "SPARCL1", "QDPR", "SLC5A6", "SSPN", "FZD4", "RETSAT", "REEP6", "PTGER3", "PHLDB1", "GADD45A", "PIM3", "ALDH2", "STOM", "SLC27A1", "DHRS7", "VEGFB", "COL15A1", "UCK1", "C3", "SUCLG1", "CAVIN1", "RNF11", "LAMA4", "PDCD4", "COX8A", "PPP1R15B", "CMPK1", "MAP4K3", "PREB", "LPCAT3", "UCP2", "SCP2", "MGLL") |
| 26 | HALLMARK_MYC_TARGETS_V1 | 0.362643 | 0.657041 | -0.220937 | -1.070802 | 729 | 51 | c("CLNS1A", "HSP90AB1", "NME1", "CANX", "VDAC3", "PSMD3", "SRSF2", "EIF4A1", "PPIA", "IMPDH2", "SMARCC1", "EEF1B2", "SF3B3", "NOP56", "HNRNPA3", "CYC1", "RACK1", "SERBP1", "RAD23B", "EIF2S1", "HNRNPD", "HNRNPA2B1", "EIF4H", "SET", "ERH", "APEX1", "FBL", "NCBP2", "DEK", "PSMD1", "USP1", "XRCC6", "HSPE1", "PWP1", "CDK4", "EIF3D", "PA2G4", "YWHAQ", "PHB2", "TRA2B", "HNRNPU", "GNL3", "HNRNPR", "SRM", "POLE3") |
| 27 | HALLMARK_MYC_TARGETS_V2 | 0.394224 | 0.657041 | -0.418336 | -1.042947 | 1173 | 6 | c("NOP56", "HSPE1", "CDK4", "PA2G4", "GNL3", "SRM") |
| 28 | HALLMARK_UV_RESPONSE_UP | 0.377237 | 0.657041 | 0.263313 | 1.073264 | 2992 | 46 | c("FOSB", "FOS", "NR4A1", "JUNB", "OLFM1", "BTG2", "MMP14") |
| 29 | HALLMARK_ANGIOGENESIS | 0.387001 | 0.657041 | 0.353569 | 1.069198 | 2869 | 14 | c("COL3A1", "FSTL1", "JAG1", "COL5A2", "POSTN", "CCND2", "ITGAV", "VCAN", "STC1", "NRP1", "APP", "LUM") |
| 30 | HALLMARK_PANCREAS_BETA_CELLS | 0.347884 | 0.657041 | -0.438893 | -1.094196 | 1035 | 6 | SYT13 |
| 31 | HALLMARK_APICAL_JUNCTION | 0.435957 | 0.703156 | 0.227597 | 1.02616 | 3583 | 76 | c("LAMA3", "COL16A1", "MMP2", "LIMA1", "CD276", "JAM3", "CERCAM", "CD34", "CD99", "SLIT2", "TMEM8B", "MYL9", "BMP1", "LDLRAP1", "FBN1", "TJP1", "PCDH1", "FSCN1", "CLDN5", "VCL", "RASA1", "SDC3", "VCAN", "PIK3R3", "SORBS3", "ITGB1", "VWF", "CTNND1", "PKD1", "RRAS", "EVL", "MYH10") |
| 32 | HALLMARK_MITOTIC_SPINDLE | 0.478729 | 0.723787 | 0.223857 | 0.997937 | 3915 | 71 | c("SHROOM1", "CDC42EP1", "GSN", "PALLD", "ABL1", "TIAM1", "MYO1E", "PKD2", "ARHGAP29", "FSCN1", "CLASP1", "BCR", "VCL", "RASA1", "FLNA", "PDLIM5", "TRIO", "FARP1", "SPTAN1", "CYTH2", "SYNPO", "TUBGCP6", "ATG4B", "CAPZB", "NOTCH2") |
| 33 | HALLMARK_FATTY_ACID_METABOLISM | 0.472622 | 0.723787 | -0.209173 | -0.994703 | 983 | 47 | c("BMPR1B", "FASN", "UGDH", "GLUL", "HSP90AA1") |
| 34 | HALLMARK_KRAS_SIGNALING_DN | 0.492175 | 0.723787 | 0.29648 | 0.988857 | 3710 | 20 | c("NR4A2", "TCF7L1", "TFAP2B", "BTG2", "COPZ2", "CPB1", "IGFBP2") |
| 35 | HALLMARK_GLYCOLYSIS | 0.519089 | 0.741555 | 0.232929 | 0.972957 | 4146 | 51 | c("GPC1", "GLCE", "AGRN", "IER3", "KDELR3", "PAM", "DCN", "HDLBP", "ELF3", "COL5A1", "B4GALT2", "CHPF", "SDC3", "P4HA2", "VCAN") |
| 36 | HALLMARK_APICAL_SURFACE | 0.580262 | 0.80592 | 0.357349 | 0.915462 | 4156 | 8 | c("TMEM8B", "GSTM3", "APP", "GAS1", "EPHB4") |
| 37 | HALLMARK_CHOLESTEROL_HOMEOSTASIS | 0.65032 | 0.8129 | -0.213899 | -0.859753 | 1524 | 26 | c("SCD", "FASN", "ACTG1") |

| | | | | | | | | |
|---|---|---|---|---|---|---|---|---|
| 38 | HALLMARK_G2M_CHECKPOINT | 0.621028 | 0.8129 | -0.194758 | -0.890715 | 1328 | 41 | c("TOP2A", "KIF5B", "CCND1", "HSPA8", "SMC4", "SRSF2", "MT2A", "SMARCC1", "SLC12A2", "BCL3", "NCL", "CUL3", "RAD23B", "PRPF4B", "HNRNPD", "MARCKS", "SS18", "TENT4A", "PURA", "UPF1", "SMAD3", "CDK4", "ATRX", "NUMA1", "SLC38A1", "YTHDC1", "EWSR1", "STAG1", "FOXN3", "TRA2B", "HNRNPU", "MAP3K20", "PML", "ARID4A", "SFPQ", "RASAL2", "NOTCH2", "BUB3", "CDKN1B") |
| 39 | HALLMARK_INTERFERON_GAMMA_RESPONSE | 0.61457 | 0.8129 | 0.222996 | 0.908932 | 4875 | 46 | c("TNFAIP6", "CDKN1A", "SSPN", "FAS", "C1S", "SOCS3", "CFH", "JAK2", "SERPING1", "C1R", "AUTS2", "CMTR1", "TXNIP", "APOL6", "TNFSF10", "ARID5B") |
| 40 | HALLMARK_IL2_STAT5_SIGNALING | 0.640225 | 0.8129 | 0.211267 | 0.892479 | 5124 | 54 | c("F2RL2", "PLPP1", "PLAGL1", "NFKBIZ", "MAFF", "COL6A1", "GADD45B", "TIAM1", "MYO1E", "CCND2", "ITGAV", "BMPR2", "KLF6", "ST3GAL4", "RHOB") |
| 41 | HALLMARK_IL6_JAK_STAT3_SIGNALING | 0.692338 | 0.844314 | 0.237513 | 0.840405 | 5294 | 25 | c("JUN", "LEPR", "FAS", "SOCS3", "TNFRSF12A", "A2M", "PDGFC", "OSMR", "TNFRSF1A", "IL1R1", "IL13RA1") |
| 42 | HALLMARK_DNA_REPAIR | 0.960832 | 0.993976 | -0.126862 | -0.597949 | 1986 | 46 | c("BRF2", "CANT1", "NME1", "POLD4", "NME4", "GPX4", "IMPDH2", "GUK1", "HCLS1", "SUPT5H", "DCTN4", "EIF1B", "STX3", "VPS28", "ADRM1", "POLR2I", "SEC61A1", "POLR2G", "POLR2K", "POLR2E", "NFX1", "NCBP2", "POLR2A", "BCAM", "SMAD5", "ITPA", "ADCY6", "DGUOK", "RBX1", "NME3", "CSTF3", "DDB1", "DAD1", "AK1", "SURF1", "SSRP1", "POLB", "ERCC3", "ELOA", "ERCC1", "AAAS", "TK2", "POLR2F") |
| 43 | HALLMARK_PROTEIN_SECRETION | 0.944595 | 0.993976 | 0.155423 | 0.594032 | 7296 | 35 | c("ATP1A1", "SSPN", "GALC", "PAM", "GBF1", "CAV2", "STX12", "VAMP3", "MON2", "DST", "ICA1", "ARFGAP3", "SEC22B", "RER1", "AP2M1", "CLTA") |
| 44 | HALLMARK_INTERFERON_ALPHA_RESPONSE | 0.982818 | 0.993976 | -0.118481 | -0.481441 | 2287 | 27 | c("IFI30", "IFITM2", "B2M", "CD74", "LAP3", "IRF9", "NUB1", "PSME1", "LY6E", "TAP1", "IFI35", "UBE2L6", "CNP", "PARP14", "PSMB8", "STAT2", "IFI44", "IRF1", "TRIM26", "CD47", "NCOA7", "MVB12A", "LPAR6", "TXNIP", "CMTR1") |
| 45 | HALLMARK_HEDGEHOG_SIGNALING | 0.87421 | 0.993976 | 0.246398 | 0.673227 | 6358 | 10 | c("RASA1", "DPYSL2", "NRP1", "PML", "CELSR1", "ETS2", "MYH9", "THY1", "LDB1") |
| 46 | HALLMARK_UNFOLDED_PROTEIN_RESPONSE | 0.927108 | 0.993976 | -0.138911 | -0.642823 | 1945 | 43 | c("LSM1", "EIF4EBP1", "KIF5B", "XBP1", "DDIT4", "EIF4A3", "LSM4", "DNAJA4", "EIF4A1", "ATP6V0D1", "YWHAZ", "NOP56", "CNOT2", "CEBPB", "EIF2S1", "STC2", "GOSR2", "ATF6", "SEC31A", "WFS1", "SHC1", "FUS", "ATF4", "BANF1", "SRPRB", "TATDN2", "DCTN1", "WIPI1", "ARFGAP1", "EDEM1", "BAG3", "YIF1A", "ATF3", "SEC11A", "MTREX", "EXOSC10", "PREB", "EEF2") |
| 47 | HALLMARK_PI3K_AKT_MTOR_SIGNALING | 0.993976 | 0.993976 | -0.098086 | -0.408215 | 2309 | 29 | c("SLC2A1", "ACACA", "SQSTM1", "GRB2") |
| 48 | HALLMARK_OXIDATIVE_PHOSPHORYLATION | 0.992252 | 0.993976 | -0.099348 | -0.52854 | 1792 | 72 | c("COX6C", "NDUFC2", "COX6A1", "ATP5PD", "TCIRG1", "UQCRC1", "VDAC3", "IDH2", "GPX4", "FH", "NDUFA4", "MAOB", "UQCRB") |
| 49 | HALLMARK_PEROXISOME | 0.931937 | 0.993976 | -0.142102 | -0.602204 | 2135 | 31 | TOP2A |
| 50 | HALLMARK_SPERMATOGENESIS | 0.980761 | 0.993976 | 0.179814 | 0.505636 | 7187 | 11 | c("JAM3", "MAST2") |

| Cluster 6 | pathway | pval | padj | ES | NES | nMoreExtreme | size | leadingEdge |
|---|---|---|---|---|---|---|---|---|
| 1 | HALLMARK_TNFA_SIGNALING_VIA_NFKB | 0.000325 | 0.003264 | -0.4422 | -2.194481 | 0 | 80 | c("FOS", "NR4A1", "FOSB", "DUSP1", "MAP3K8", "EGR1", "SERPINE1", "PHLDA1", "CCN1", "BTG2", "ZFP36", "IER2", "TNC", "VEGFA", "SQSTM1", "JUN", "BTG1", "SOCS3", "JUNB", "NFKBIA", "KLF9", "EFNA1", "MCL1", "SDC4", "CEBPB", "MAP2K3", "MYC", "CEBPD", "B4GALT5") |
| 2 | HALLMARK_ESTROGEN_RESPONSE_EARLY | 0.000326 | 0.003264 | -0.381657 | -1.944792 | 0 | 90 | c("HSPB8", "FOS", "RET", "PAPSS2", "RHOBTB3", "SLC2A1", "GAB2", "ABHD2", "SLC7A2", "BLVRB", "PGR", "TOB1", "ADCY1", "AKAP1", "FHL2", "MINDY1", "GFRA1", "ELF3", "SCNN1A", "SLC9A3R1", "ITPK1", "TTC39A", "STC2", "RARA") |
| 3 | HALLMARK_INTERFERON_ALPHA_RESPONSE | 0.000148 | 0.003264 | 0.543111 | 2.279895 | 0 | 50 | c("IFI27", "IFI44L", "ISG15", "IFIT2", "MX1", "IFI44", "TRIM14", "IFIT3", "CD74", "OAS1", "SAMD9", "EPSTI1", "CSF1", "PSMB9", "PARP14", "IRF7", "SAMD9L", "B2M", "PARP9", "IRF9", "IFITM1", "SLC25A28", "ELF1", "LPAR6", "UBA7", "MOV10", "TAP1", "PLSCR1", "IFIH1") |
| 4 | HALLMARK_INTERFERON_GAMMA_RESPONSE | 0.000145 | 0.003264 | 0.514244 | 2.377817 | 0 | 79 | c("IFI27", "IFI44L", "ISG15", "CCL5", "XAF1", "IFIT1", "IFIT2", "STAT1", "MX1", "IFI44", "MX2", "LCP2", "TRIM14", "OAS2", "IFIT3", "TNFSF10", "CD74", "EPSTI1", "OAS3", "PSMB9", "FGL2", "PARP14", "IRF8", "IRF7", "SAMD9L", "CIITA", "B2M", "JAK2", "IRF9", "IL10RA", "SAMHD1", "SLC25A28", "CFH", "CASP7", "ST8SIA4", "TNFAIP3", "SERPING1", "VCAM1", "TAP1", "PLSCR1", "IFIH1", "MTHFD2", "PSMB2", "PML", "APOL6", "IFITM3") |
| 5 | HALLMARK_HEME_METABOLISM | 0.000314 | 0.003264 | -0.481246 | -2.256835 | 0 | 61 | c("HBD", "HBB", "SLC4A1", "SLC2A1", "BTG2", "SNCA", "BLVRB", "TFRC", "SLC25A37", "BCAM", "HDGF", "SELENBP1", "CIR1", "CAT", "MGST3", "MPP1", "PRDX2", "GLRX5", "HEBP1", "ADIPOR1", "MAP2K3", "RNF19A") |
| 6 | HALLMARK_OXIDATIVE_PHOSPHORYLATION | 0.00096 | 0.007997 | -0.39528 | -1.889181 | 2 | 68 | c("NDUFC2", "MAOB", "PDK4", "ATP1B1", "NDUFA1", "COX4I1", "NDUFS4", "NDUFS2", "VDAC3", "MGST3", "ECH1", "COX7C", "IDH3B", "UQCRB", "NDUFB5", "NDUFS6", "NDUFC1", "NDUFB6", "ATP5PD", "NDUFA2") |
| 7 | HALLMARK_ALLOGRAFT_REJECTION | 0.001623 | 0.011596 | 0.430545 | 1.80736 | 10 | 50 | c("CCL5", "FYB1", "STAT1", "IGSF6", "GPR65", "LCP2", "CD74", "CSF1", "THY1", "IRF8", "IRF7", "B2M", "GALNT1", "STAB1", "JAK2", "IL2RG", "CCND2", "ST8SIA4", "TAP1", "TRAF2", "CSK") |
| 8 | HALLMARK_HYPOXIA | 0.002265 | 0.014159 | -0.371995 | -1.81233 | 6 | 73 | c("FOS", "DUSP1", "SERPINE1", "CCN1", "SLC2A1", "CITED2", "ZFP36", "HMOX1", "STC1", "SELENBP1", "STC2", "VEGFA", "FBP1", "JUN", "BTG1", "EFNA1", "P4HA1", "SDC4", "MIF") |
| 9 | HALLMARK_UV_RESPONSE_UP | 0.00279 | 0.015499 | -0.44206 | -1.94449 | 8 | 47 | c("FOS", "NR4A1", "FOSB", "RET", "BTG2", "TFRC", "HMOX1", "EPHX1", "SQSTM1", "BTG1", "JUNB", "NFKBIA", "SELENOW", "PRPF3") |
| 10 | HALLMARK_ESTROGEN_RESPONSE_LATE | 0.005518 | 0.027588 | -0.34193 | -1.703327 | 16 | 81 | c("HSPB8", "S100A9", "FOS", "RET", "PAPSS2", "PRLR", "ABHD2", "BLVRB", "ZFP36", "PGR", "TOB1", "CPE", "SCNN1A", "SLC9A3R1", "ITPK1") |
| 11 | HALLMARK_KRAS_SIGNALING_UP | 0.011898 | 0.054081 | 0.367607 | 1.603339 | 80 | 59 | c("GALNT3", "MAP3K1", "GUCY1A1", "LAPTM5", "IRF8", "PLAT", "C3AR1", "TSPAN1", "FCER1G", "CD37", "ITGBL1", "LY96", "ST6GAL1", "IL10RA", "IL2RG", "PLAU", "CFH", "CSF2RA", "CCND2", "TNFAIP3", "TMEM176B", "CCSER2", "TRIB2", "CBR4", "MMP11") |
| 12 | HALLMARK_REACTIVE_OXYGEN_SPECIES_PATHWAY | 0.018192 | 0.075802 | -0.480746 | -1.733952 | 61 | 22 | c("MPO", "PRDX6", "EGLN2", "NDUFS2", "CAT", "SOD1", "JUNB", "PRDX2", "NQO1") |
| 13 | HALLMARK_CHOLESTEROL_HOMEOSTASIS | 0.028839 | 0.110919 | 0.433388 | 1.563701 | 190 | 27 | c("FADS2", "SCD", "CHKA", "FASN", "SREBF2", "FDFT1", "CXCL16", "GPX8", "ERRFI1", "PLSCR1", "TP53INP1", "CPEB2", "LDLR", "LGMN", "STX5", "PNRC1", "JAG1", "CTNNB1", "NIBAN1") |
| 14 | HALLMARK_XENOBIOTIC_METABOLISM | 0.037198 | 0.132849 | -0.344942 | -1.539253 | 119 | 50 | c("SERPINE1", "PAPSS2", "PDK4", "BLVRB", "HMOX1", "CD36", "EPHX1", "ALDH9A1", "CFB", "CAT", "FBP1", "ECH1", "ATP2A2") |
| 15 | HALLMARK_G2M_CHECKPOINT | 0.041488 | 0.138294 | 0.360379 | 1.467576 | 280 | 44 | c("RBM14", "CCND1", "LMNB1", "MARCKS", "NUSAP1", "BUB3", "SLC38A1", "TMPO", "NUP98", "MKI67", "TOP2A", "MAP3K20", "PML", "UBE2C", "SMC1A", "KIF22", "EWSR1", "NSD2", "DR1", "TGFB1") |
| 16 | HALLMARK_PANCREAS_BETA_CELLS | 0.065988 | 0.206214 | -0.691939 | -1.559232 | 239 | 5 | c("SYT13", "SRP9") |

| | | | | | | | | |
|---|---|---|---|---|---|---|---|---|
| 17 | HALLMARK_MITOTIC_SPINDLE | 0.081276 | 0.239048 | 0.287847 | 1.333963 | 562 | 80 | c("FGD6", "SAC3D1", "DOCK4", "CTTN", "LMNB1", "MARCKS", "FLNB", "NUSAP1", "DYNC1H1", "SPTAN1", "RABGAP1", "LATS1", "ARFIP2", "EZR", "PALLD", "TOP2A", "CAPZB", "FSCN1", "FARP1", "CDK5RAP2", "SYNPO", "ECT2", "SMC1A", "PCM1", "KIF22", "PDLIM5", "RALBP1", "SOS1", "EPB41", "MYH9", "ROCK1", "ARHGEF7", "YWHAE", "PAFAH1B1", "MYO9B", "SUN2", "WASF2", "EPB41L2", "SHROOM1", "DOCK2", "CKAP5", "KLC1", "SMC4", "RHOT2", "CNTRL", "STK38L", "DST", "AKAP13", "TAOK2", "ABL1", "NOTCH2", "MYH10", "TUBGCP6", "FLNA", "VCL", "PREX1") |
| 18 | HALLMARK_UV_RESPONSE_DN | 0.13181 | 0.366138 | -0.27761 | -1.295628 | 420 | 59 | c("DUSP1", "SERPINE1", "CCN1", "CITED2", "IGFBP5", "FHL2") |
| 19 | HALLMARK_COMPLEMENT | 0.156195 | 0.411041 | 0.276604 | 1.243516 | 1073 | 68 | c("RCE1", "CCL5", "C3", "DOCK4", "LCP2", "C1QC", "C1QA", "APOC1", "CTSH", "PSMB9", "IRF7", "CTSC", "PLAT", "USP14", "FCER1G", "JAK2", "CFH", "GNB4", "CASP7", "TNFAIP3", "SERPING1", "SH2B3", "PRCP", "PLSCR1", "DOCK10", "PPP4C", "CTSD", "RHOG", "DYRK2", "C2", "RASGRP1", "C1R", "LGMN", "IRF1", "LRP1", "C1S", "GRB2", "ZEB1", "LAP3", "TIMP2") |
| 20 | HALLMARK_EPITHELIAL_MESENCHYMAL_TRANSITIO N | 0.18647 | 0.466175 | 0.240858 | 1.184315 | 1300 | 110 | c("LRRC15", "MFAP5", "LUM", "COL8A2", "MXRA5", "LOXL1", "TGM2", "SDC1", "FBLN2", "SFRP4", "IL32", "EFEMP2", "COMP", "THY1", "GLIPR1", "NID2", "ECM2", "GREM1", "FSTL1", "COL12A1", "GJA1", "COL5A3", "TNFAIP3", "VCAM1", "FBN1", "ITGB5", "COL11A1", "WIPF1", "DPYSL3", "FBLN1", "DAB2", "MYLK", "PMEPA1", "EDIL3", "SNAI2", "COL5A2", "COL16A1", "BGN", "TGFB1", "LRP1", "PRRX1", "CALD1", "PDGFRB", "EMP3", "SERPINE2", "MCM7", "LAMC1", "INHBA", "BASP1", "MATN3", "FAP", "IGFBP3", "GPC1", "ACTA2", "NNMT", "DCN", "TPM1", "CDH11", "SPARC") |
| 21 | HALLMARK_IL2_STAT5_SIGNALING | 0.23758 | 0.565668 | 0.26313 | 1.163509 | 1625 | 63 | c("SLC29A2", "LRIG1", "GPR65", "TNFSF10", "TGM2", "CSF1", "FGL2", "IRF8", "CD81", "DENND5A", "IL10RA", "SWAP70", "SNX14", "GLIPR2", "CCND2", "RNH1", "MYO1C", "GSTO1", "PLSCR1", "SPRED2", "SNX9", "IFITM3") |
| 22 | HALLMARK_GLYCOLYSIS | 0.280461 | 0.63686 | -0.250718 | -1.133773 | 899 | 53 | c("EGLN3", "CITED2", "STC1", "ELF3", "ALDH9A1", "STC2", "VEGFA", "SOD1", "IDUA") |
| 23 | HALLMARK_KRAS_SIGNALING_DN | 0.292956 | 0.63686 | 0.375589 | 1.151093 | 1912 | 15 | c("IFI44L", "SPTBN2", "MX1", "BMPR1B", "THNSL2", "SKIL", "CELSR2", "SYNPO") |
| 24 | HALLMARK_APOPTOSIS | 0.321078 | 0.642155 | -0.231673 | -1.096777 | 1012 | 64 | c("BTG2", "NEDD9", "HMOX1", "CLU", "HSPB1", "DAP3", "RARA", "SQSTM1", "SOD1", "EMP1", "JUN", "LMNA", "MCL1", "ERBB2", "BCL2L1") |
| 25 | HALLMARK_ADIPOGENESIS | 0.314981 | 0.642155 | -0.230756 | -1.09921 | 983 | 67 | c("HSPB8", "TOB1", "GPAT4", "CD36", "FABP4", "CMBL", "CAT", "MGST3", "SOD1", "ECH1") |
| 26 | HALLMARK_MYOGENESIS | 0.345335 | 0.664106 | -0.230551 | -1.076002 | 1102 | 59 | c("PVALB", "HSPB8", "CLU", "CD36", "STC2", "MEF2D", "APOD") |
| 27 | HALLMARK_APICAL_JUNCTION | 0.407941 | 0.718124 | 0.231407 | 1.040327 | 2804 | 68 | c("FYB1", "CRAT", "ITGB4", "MDK", "EVL", "THY1", "YWHAH", "PIK3R3", "CAP1", "ARPC2", "PARVA", "MSN", "VCAM1", "FBN1", "HRAS", "FSCN1", "MAP3K20", "INSIG1", "MYH9", "COL16A1", "TJP1", "NF2", "ACTB", "TIAL1", "NECTIN2", "CD99", "GNAI2", "CDH11", "EPB41L2", "SIRPA", "CERCAM", "PECAM1", "SORBS3", "PKD1", "CTNND1", "MYL9", "MMP2", "TAOK2", "MYH10", "HADH", "ARHGEF6", "BMP1", "RAC2", "VCL") |
| 28 | HALLMARK_FATTY_ACID_METABOLISM | 0.426676 | 0.718124 | 0.251336 | 1.030125 | 2894 | 45 | c("CRAT", "UGDH", "FASN", "BMPR1B", "GSTZ1", "REEP6", "ALDH1A1", "YWHAH", "ELOVL5") |
| 29 | HALLMARK_COAGULATION | 0.430874 | 0.718124 | -0.230388 | -1.013407 | 1389 | 47 | c("S100A1", "SERPINE1", "S100A13", "CLU", "MMP9") |
| 30 | HALLMARK_BILE_ACID_METABOLISM | 0.425182 | 0.718124 | 0.307464 | 1.036033 | 2795 | 21 | c("FADS2", "SLC22A18", "DIO2", "ALDH1A1", "SCP2", "ATXN1", "AR", "DHCR24", "PRDX5", "ABCA3") |
| 31 | HALLMARK_HEDGEHOG_SIGNALING | 0.465074 | 0.729882 | 0.371508 | 1.00363 | 3035 | 10 | c("CELSR1", "THY1", "DPYSL2", "TLE1", "PML", "MYH9", "NRP2") |
| 32 | HALLMARK_UNFOLDED_PROTEIN_RESPONSE | 0.476306 | 0.729882 | -0.242621 | -0.983281 | 1567 | 34 | c("LSM1", "DNAJA4", "EIF4EBP1", "STC2", "VEGFA") |
| 33 | HALLMARK_MYC_TARGETS_V1 | 0.481722 | 0.729882 | -0.217257 | -0.979672 | 1554 | 52 | c("CLNS1A", "HSP90AB1", "CBX3", "HDGF", "NME1", "VDAC3", "IMPDH2", "CCT3") |
| 34 | HALLMARK_IL6_JAK_STAT3_SIGNALING | 0.51757 | 0.739386 | -0.240993 | -0.950586 | 1737 | 30 | c("MAP3K8", "HMOX1", "CD36", "CD9", "JUN", "SOCS3") |
| 35 | HALLMARK_P53_PATHWAY | 0.511453 | 0.739386 | -0.206937 | -0.970445 | 1629 | 61 | c("FOS", "BTG2", "TOB1", "HMOX1", "TM4SF1", "EPHX1") |
| 36 | HALLMARK_ANDROGEN_RESPONSE | 0.578629 | 0.762987 | 0.229816 | 0.928331 | 3925 | 42 | c("SCD", "GUCY1A1", "BMPR1B", "CCND1", "ELOVL5", "B2M", "FKBP5", "GPD1L", "XRCC6", "PMEPA1", "INSIG1", "PDLIM5", "AZGP1", "DHCR24", "UBE2J1", "AKT1", "ZMIZ1", "SELENOP") |
| 37 | HALLMARK_MTORC1_SIGNALING | 0.570375 | 0.762987 | -0.196598 | -0.933797 | 1778 | 66 | c("EGLN3", "SLC2A1", "BTG2", "TFRC", "STC1", "IGFBP5", "SLC9A3R1", "SQSTM1", "ATP2A2", "CD9", "NUPR1", "P4HA1", "LTA4H", "MAP2K3") |

| | | | | | | | | |
|---|---|---|---|---|---|---|---|---|
| 38 | HALLMARK_INFLAMMATORY_RESPONSE | 0.61039 | 0.762987 | 0.21853 | 0.917357 | 4135 | 50 | c("CCL5", "LCP2", "MSR1", "TNFSF10", "CSF1", "IRF7", "C3AR1", "STAB1", "IFITM1", "IL10RA", "CD14", "ITGB8", "AXL", "RHOG", "LDLR", "CYBB", "RASGRP1", "IFNGR2", "IL1R1") |
| 39 | HALLMARK_ANGIOGENESIS | 0.599423 | 0.762987 | -0.269386 | -0.888935 | 2076 | 16 | c("STC1", "VEGFA", "FGFR1", "LRPAP1") |
| 40 | HALLMARK_PEROXISOME | 0.591662 | 0.762987 | -0.229855 | -0.899813 | 2000 | 29 | c("CRABP2", "ALDH9A1", "CAT", "SOD1", "ECH1") |
| 41 | HALLMARK_MYC_TARGETS_V2 | 0.656045 | 0.800055 | 0.375962 | 0.862541 | 4199 | 6 | c("SLC29A2", "DCTPP1") |
| 42 | HALLMARK_WNT_BETA_CATENIN_SIGNALING | 0.685911 | 0.816561 | 0.273794 | 0.839114 | 4478 | 15 | c("LEF1", "CCND2", "FZD1", "AXIN1", "JAG1", "CTNNB1", "TP53", "CSNK1E", "MAML1", "NUMB", "NCOR2") |
| 43 | HALLMARK_DNA_REPAIR | 0.737632 | 0.857711 | 0.206 | 0.820445 | 4964 | 40 | c("SAC3D1", "EDF1", "ZWINT", "POLD4", "TYMS", "POLR2G", "TSG101", "GTF2H1", "SURF1", "DDB2", "TAF10", "NELFB") |
| 44 | HALLMARK_PI3K_AKT_MTOR_SIGNALING | 0.805365 | 0.915188 | 0.199802 | 0.761925 | 5403 | 34 | c("PPP1CA", "GRK2", "ACACA", "PIK3R3", "IL2RG", "CFL1", "TRAF2", "HRAS", "MKNK2") |
| 45 | HALLMARK_TGF_BETA_SIGNALING | 0.883837 | 0.92305 | 0.185225 | 0.677742 | 5850 | 29 | c("PPP1CA", "RAB31", "TGFBR1", "SKI", "SKIL", "PMEPA1", "IFNGR2", "TGFB1", "TJP1", "HDAC1", "CTNNB1", "LTBP2", "SMURF2") |
| 46 | HALLMARK_NOTCH_SIGNALING | 0.895196 | 0.92305 | 0.226048 | 0.646221 | 5850 | 12 | c("CCND1", "FZD1", "TCF7L2", "JAG1", "SAP30", "NOTCH2", "NOTCH3", "CUL1", "RBX1", "NOTCH1") |
| 47 | HALLMARK_PROTEIN_SECRETION | 0.904589 | 0.92305 | -0.161307 | -0.65864 | 2995 | 35 | c("ANP32E", "SOD1", "CD63", "ERGIC3", "LMAN1", "TOM1L1", "VPS45", "SNX2", "ARF1", "STX12", "SNAP23", "COPB2", "SEC31A", "STX7", "RAB14", "AP2M1", "NAPA", "YKT6", "VPS4B", "PPT1", "MAPK1", "ICA1", "SCRN1", "PAM", "STX16", "ABCA1", "DST", "SEC24D") |
| 48 | HALLMARK_APICAL_SURFACE | 0.845777 | 0.92305 | 0.244503 | 0.698978 | 5527 | 12 | c("THY1", "IL2RG", "ATP8B1", "ADIPOR2", "SULF2", "GSTM3", "LYN") |
| 49 | HALLMARK_E2F_TARGETS | 0.862736 | 0.92305 | 0.174013 | 0.727684 | 5838 | 49 | c("ATAD2", "RRM2", "LMNB1", "RPA1", "TMPO", "TUBB", "NCAPD2", "MKI67", "ASF1B", "TOP2A", "MTHFD2", "XRCC6", "SMC1A", "KIF22", "TK1", "NBN", "DNMT1", "HELLS", "MCM7", "DCTPP1", "MXD3") |
| 50 | HALLMARK_SPERMATOGENESIS | 0.923807 | 0.923807 | 0.212514 | 0.607531 | 6037 | 12 | c("HSPA2", "MAST2", "STAM2", "NF2", "PHKG2", "GSTM3", "STRBP") |

| Cluster 7 | pathway | pval | padj | ES | NES | nMoreExtreme | size | leadingEdge |
|---|---|---|---|---|---|---|---|---|
| 1 | HALLMARK_HYPOXIA | 0.0001 | 0.000818 | -0.552303 | -1.767214 | 0 | 60 | c("SERPINE1", "STC1", "CCN1", "MIF", "DCN", "GPC1", "BGN", "CCN2", "STC2", "JUN", "IGFBP3", "COL5A1", "ANXA2", "CA12", "EFNA1", "BHLHE40", "TGFBI", "HMOX1", "TPD52", "SDC4", "FOS", "TGFB3", "DTNA", "FBP1", "SDC2", "LOX", "CAVIN1", "MT2A", "DUSP1") |
| 2 | HALLMARK_ESTROGEN_RESPONSE_EARLY | 1.00E-04 | 0.000818 | -0.553796 | -1.803766 | 0 | 76 | c("ADCY1", "GFRA1", "HSPB8", "PGR", "NBL1", "RET", "SLC7A2", "GREB1", "CELSR1", "GJA1", "FLNB", "MAPT", "SEC14L2", "STC2", "PRSS23", "TFF1", "MLPH", "ELF3", "SCNN1A", "FARP1", "CA12", "BHLHE40", "CCND1", "SLC39A6", "HES1", "KRT8", "SEMA3B", "THSD4", "LRIG1", "FOS", "CLDN7", "TOB1", "FASN", "KRT19", "RHOBTB3", "FAM102A", "TFF3", "TTC39A", "PLAAT3", "PAPSS2", "RHOD", "NRIP1", "GAB2", "MYOF", "SLC9A3R1", "IGFBP4", "B4GALT1", "MUC1", "ABHD2", "RARA", "MAST4") |
| 3 | HALLMARK_ESTROGEN_RESPONSE_LATE | 0.0001 | 0.000818 | -0.526036 | -1.702937 | 0 | 70 | c("CXCL14", "SERPINA3", "HSPB8", "AGR2", "PRLR", "SCUBE2", "PGR", "NBL1", "RET", "CD9", "FLNB", "CPE", "UGDH", "MAPT", "DLG5", "PRSS23", "TFF1", "SCNN1A", "FARP1", "CA12", "COX6C", "CCND1", "SEMA3B", "CDH1", "FOS", "LLGL2", "TOB1", "MDK", "LSR", "SERPINA1", "KRT19", "FAM102A", "TFF3", "PLAAT3", "PAPSS2", "NRIP1", "SGK1", "MYOF", "SLC9A3R1", "IGFBP4", "RABEP1", "DNAJC1", "ABHD2") |
| 4 | HALLMARK_EPITHELIAL_MESENCHYMAL_TRANSITIO N | 1.00E-04 | 0.000818 | -0.627318 | -2.087534 | 0 | 107 | c("MMP2", "SPP1", "HTRA1", "ADAM12", "COMP", "SERPINE1", "COL5A2", "FSTL1", "COL11A1", "ACTA2", "COL4A1", "ELN", "LUM", "SFRP4", "MMP14", "FBLN1", "FBLN2", "FAP", "CCN1", "TIMP1", "TIMP3", "POSTN", "GJA1", "THBS2", "DCN", "GPC1", "TAGLN", "BGN", "CCN2", "PDGFRB", "COL12A1", "MXRA5", "LOXL1", "CDH11", "LOXL2", "ITGB5", "JUN", "THY1", "IGFBP3", "PCOLCE", "COL5A1", "ECM1", "LRRC15", "CALD1", "TGFBI", "PRRX1", "SDC4", "CTHRC1", "SDC1", "ITGAV", "TPM4", "DPYSL3", "MYL9", "COL16A1", "COL4A2", "LRP1", "FBN1", "TNC", "LOX", "LAMC1", "DST", "IGFBP2", "COL1A1", "COL3A1", "SERPINH1", "DAB2", "NNMT", "GEM", "RHOB", "PMP22", "COL1A2", "ITGB1", "CAPG", "PFN2", "FUCA1", "PMEPA1", "THBS1", "CALU", "IGFBP4") |
| 5 | HALLMARK_UV_RESPONSE_DN | 0.0001 | 0.000818 | -0.549853 | -1.714728 | 0 | 45 | c("SERPINE1", "COL5A2", "COL11A1", "CCN1", "NRP1", "GJA1", "PDGFRB", "ERBB2", "IGFBP5", "ANXA2", "RBPMS", "BHLHE40", "EFEMP1", "INPP4B", "SDC2", "LAMC1", "DUSP1", "AKT3", "COL1A1", "COL3A1", "DAB2", "PLPP3", "APBB2", "PMP22", "COL1A2") |
| 6 | HALLMARK_COAGULATION | 0.0001 | 0.000818 | -0.677304 | -2.112187 | 0 | 45 | c("MMP2", "HTRA1", "APOC1", "MMP11", "COMP", "SERPINE1", "C3", "C1S", "PLAT", "CTSK", "MMP14", "S100A1", "CD9", "TIMP1", "TIMP3", "SERPING1", "C1R", "C1QA", "PRSS23", "CLU", "A2M", "S100A13", "CFB", "LRP1", "CRIP2", "FBN1", "LGMN", "SERPINA1", "CAPN2") |
| 7 | HALLMARK_COMPLEMENT | 0.0003 | 0.001841 | -0.536725 | -1.693073 | 2 | 51 | c("APOC1", "SERPINE1", "C3", "C1S", "C1QC", "PLAT", "MMP14", "TIMP1", "SERPING1", "C1R", "C1QA", "CTSD", "CLU", "S100A13", "LIPA", "CFB", "COL4A2", "LRP1", "MMP13", "LGMN", "SERPINA1", "GATA3", "TIMP2", "CSRP1", "CTSL", "ANXA5") |
| 8 | HALLMARK_KRAS_SIGNALING_UP | 0.000301 | 0.001841 | -0.53959 | -1.672573 | 2 | 42 | c("GPNMB", "SPP1", "MMP11", "SERPINA3", "SPARCL1", "APOD", "PLAT", "NRP1", "CPE", "IGFBP3", "PRRX1", "ITGBL1", "SEMA3B", "CFB", "JUP", "TMEM176B") |
| 9 | HALLMARK_MYOGENESIS | 0.0004 | 0.00218 | -0.528403 | -1.657667 | 3 | 48 | c("ADAM12", "HSPB8", "APOD", "IGFBP7", "CNN3", "PVALB", "TAGLN", "STC2", "ITGB5", "IGFBP3", "BHLHE40", "CLU", "SPDEF", "DTNA", "COL4A2", "PLXNB2", "NQO1", "TNNT1", "ERBB3", "SCD", "AEBP1", "ATP6AP1", "GPX3", "MYO1C", "COL1A1", "COL3A1") |
| 10 | HALLMARK_ANGIOGENESIS | 0.000732 | 0.003586 | -0.67401 | -1.80239 | 6 | 15 | c("SPP1", "COL5A2", "FSTL1", "LUM", "STC1", "NRP1", "TIMP1", "POSTN", "ITGAV", "FGFR1", "COL3A1") |
| 11 | HALLMARK_APICAL_JUNCTION | 0.003701 | 0.016487 | -0.467702 | -1.491581 | 36 | 57 | c("MMP2", "PARVA", "PCDH1", "CDH11", "THY1", "TGFBI", "ACTG1", "NECTIN2", "CLDN4", "CDH1", "CD276", "MYL9", "COL16A1", "CLDN7", "JUP", "FBN1", "EVL", "MDK", "LIMA1", "AKT3", "SDC3", "SHC1", "CTNND1", "CERCAM", "ITGB1", "EPB41L2", "B4GALT1", "SORBS3", "PIK3R3", "THBS3", "CTNNA1", "TSPAN4", "MSN", "VCAN", "EXOC4", "WASL") |

| | | | | | | | | |
|---|---|---|---|---|---|---|---|---|
| 12 | HALLMARK_E2F_TARGETS | 0.006667 | 0.027222 | 0.392463 | 2.140482 | 0 | 24 | c("ANP32E", "TFRC", "SPC24", "MXD3", "MYBL2", "PLK1", "LUC7L3", "HMGB2", "TACC3", "LBR", "MYC", "PNN") |
| 13 | HALLMARK_MYC_TARGETS_V2 | 0.01312 | 0.049451 | 0.585414 | 1.765133 | 17 | 7 | c("PLK1", "MYC", "NOP56", "GNL3", "CBX3", "SRM", "HSPE1") |
| 14 | HALLMARK_KRAS_SIGNALING_DN | 0.019243 | 0.067351 | -0.620297 | -1.559821 | 177 | 11 | c("CPB1", "BMPR1B", "SELENOP", "EFHD1", "IGFBP2", "GPRC5C", "SKIL", "SGK1") |
| 15 | HALLMARK_CHOLESTEROL_HOMEOSTASIS | 0.02586 | 0.084477 | -0.542659 | -1.502339 | 250 | 18 | c("CD9", "ACTG1", "CLU", "SEMA3B", "FASN", "LGMN", "SCD", "S100A11", "ANXA5", "CTNNB1", "LGALS3", "ALCAM") |
| 16 | HALLMARK_APOPTOSIS | 0.033307 | 0.102002 | -0.42012 | -1.341499 | 332 | 58 | c("MMP2", "LUM", "PLAT", "IFITM3", "TIMP1", "TIMP3", "DCN", "BGN", "PDGFRB", "ERBB2", "JUN", "CCND1", "CLU", "PEA15", "HMOX1", "ERBB3", "TIMP2", "GPX3", "EMP1", "CTNNB1", "RHOB", "LMNA", "GPX4", "PPT1", "LGALS3", "SPTAN1", "RARA", "GSN") |
| 17 | HALLMARK_ANDROGEN_RESPONSE | 0.041792 | 0.107779 | -0.442636 | -1.3662 | 416 | 40 | c("BMPR1B", "AZGP1", "SELENOP", "CCND1", "KRT8", "TPD52", "SPDEF", "ITGAV", "INPP4B", "B2M", "KRT19", "SCD", "DHCR24", "DBI", "SGK1", "PMEPA1", "ACSL3", "B4GALT1", "ABHD2", "UAP1") |
| 18 | HALLMARK_PI3K_AKT_MTOR_SIGNALING | 0.04 | 0.107779 | 0.25655 | 1.470459 | 3 | 27 | c("MAP2K3", "SLC2A1", "SLA", "RAC1", "SMAD2", "PLA2G12A", "TNFRSF1A", "ARHGDIA", "MAPK1", "AKT1", "PAK4", "ARPC3", "GSK3B", "ITPR2", "STAT2", "UBE2D3", "GRB2", "CAB39", "PPP1CA", "ACACA", "SQSTM1", "ARF1", "DUSP3", "VAV3", "PIK3R3", "CFL1", "CALR") |
| 19 | HALLMARK_IL2_STAT5_SIGNALING | 0.041417 | 0.107779 | -0.423897 | -1.339736 | 413 | 52 | c("SPP1", "CD81", "S100A1", "ENPP1", "IFITM3", "NRP1", "PHLDA1", "ECM1", "BHLHE40", "LRIG1", "AHR", "ITGAV", "MYO1C", "EMP1", "MAPKAPK2", "RHOB", "CYFIP1", "CAPG", "GPX4", "ANXA4", "MUC1", "ALCAM", "CTSZ", "BMPR2", "IGF1R", "AHCY", "PLEC", "CD44", "SYNGR2", "MAP3K8", "AHNAK") |
| 20 | HALLMARK_INTERFERON_ALPHA_RESPONSE | 0.058836 | 0.144147 | -0.504205 | -1.40827 | 572 | 19 | c("C1S", "IFITM3", "IFI30", "IFI27", "CD74", "B2M", "IFITM2", "LGALS3BP") |
| 21 | HALLMARK_GLYCOLYSIS | 0.0626 | 0.146067 | -0.42019 | -1.31266 | 624 | 46 | c("STC1", "MIF", "DCN", "GPC1", "STC2", "IGFBP3", "ELF3", "COL5A1", "TGFBI", "SDC1", "CHPF", "SDC2", "TFF3", "EGLN3", "PKM", "SDC3", "DDIT4", "IL13RA1", "PAM", "PPIA", "B4GALT1", "QSOX1", "AGRN", "HDLBP", "CD44", "COPB2", "VCAN") |
| 22 | HALLMARK_TNFA_SIGNALING_VIA_NFKB | 0.0814 | 0.17629 | -0.39056 | -1.257537 | 813 | 65 | c("SERPINE1", "NR4A1", "EGR1", "CCN1", "PHLDA1", "JUN", "FOSB", "EFNA1", "BHLHE40", "CCND1", "DUSP4", "HES1", "PLK2", "SDC4", "FOS", "NFKBIA", "TNC", "EIF1", "DUSP1", "PLPP3", "SOCS3", "GEM", "RHOB", "SGK1", "PMEPA1", "MARCKS", "B4GALT1", "KLF9", "FOSL2", "CD44", "PLAU", "JUNB", "NINJ1", "MAP3K8") |
| 23 | HALLMARK_INTERFERON_GAMMA_RESPONSE | 0.082748 | 0.17629 | -0.429019 | -1.310111 | 824 | 36 | c("C1S", "IFITM3", "SERPING1", "C1R", "IFI30", "IFI27", "CFB", "CD74", "B2M", "NFKBIA", "MT2A", "IFITM2", "LGALS3BP", "SOCS3", "STAT3", "BST2", "LY6E") |
| 24 | HALLMARK_P53_PATHWAY | 0.093512 | 0.190921 | -0.40654 | -1.267801 | 933 | 45 | c("NUPR1", "CD81", "TM4SF1", "JUN", "S100A10", "IFI30", "CTSD", "HMOX1", "PLK2", "SDC1", "FOS", "TOB1", "PLXNB2", "EPHX1", "ZFP36L1") |
| 25 | HALLMARK_APICAL_SURFACE | 0.114896 | 0.216534 | -0.573528 | -1.338094 | 1011 | 8 | c("SULF2", "GSTM3", "THY1", "GATA3", "B4GALT1", "APP") |
| 26 | HALLMARK_XENOBIOTIC_METABOLISM | 0.113837 | 0.216534 | -0.399363 | -1.245422 | 1136 | 45 | c("SERPINE1", "FBLN1", "UGDH", "PDK4", "HMOX1", "ESR1", "CFB", "JUP", "TMEM176B", "EPHX1", "FBP1", "NQO1", "MT2A", "ALDH2", "APOE", "PAPSS2", "SPINT2", "IGFBP4", "BCAR1", "AHCY", "NINJ1", "ATP2A2", "PGRMC1", "MAN1A1", "MCCC2", "DHRS7", "ALDH9A1") |
| 27 | HALLMARK_SPERMATOGENESIS | 0.134435 | 0.243975 | -0.668714 | -1.288989 | 1032 | 4 | c("GSTM3", "SLC12A2", "PEBP1") |
| 28 | HALLMARK_NOTCH_SIGNALING | 0.161538 | 0.272944 | -0.532205 | -1.275442 | 1448 | 9 | c("NOTCH3", "CCND1", "HES1") |
| 29 | HALLMARK_PANCREAS_BETA_CELLS | 0.160487 | 0.272944 | -0.565578 | -1.273836 | 1384 | 7 | c("SYT13", "AKT3", "SPCS1", "MAFB") |
| 30 | HALLMARK_FATTY_ACID_METABOLISM | 0.207841 | 0.339473 | -0.408502 | -1.198582 | 2056 | 26 | c("BMPR1B", "MIF", "UGDH", "S100A10", "FASN", "EPHX1", "PCBD1", "DHCR24") |
| 31 | HALLMARK_TGF_BETA_SIGNALING | 0.273517 | 0.422673 | -0.390237 | -1.144991 | 2706 | 26 | c("SERPINE1", "LTBP2", "CDH1", "CTNNB1", "SKIL", "PMEPA1", "THBS1", "ID3", "BMPR2", "ENG", "JUNB", "RAB31", "RHOA", "NCOR2", "HIPK2", "FNTA", "ARID4B", "PPP1CA") |
| 32 | HALLMARK_HEDGEHOG_SIGNALING | 0.276031 | 0.422673 | -0.484776 | -1.161777 | 2475 | 9 | c("CELSR1", "NRP1", "THY1") |
| 33 | HALLMARK_MTORC1_SIGNALING | 0.334467 | 0.496633 | -0.342688 | -1.085175 | 3342 | 53 | c("NUPR1", "STC1", "CD9", "IGFBP5", "CANX", "IFI30", "BHLHE40", "LGMN", "SCD", "DHCR24", "EGLN3", "SYTL2", "SERPINH1", "USO1", "DDIT4", "CALR", "SLC9A3R1", "PPIA", "ACSL3", "PIK3R3", "PSMB5") |
| 34 | HALLMARK_INFLAMMATORY_RESPONSE | 0.345002 | 0.497209 | -0.376234 | -1.097113 | 3409 | 25 | c("SERPINE1", "MMP14", "SLC7A2", "TIMP1", "AHR", "NFKBIA", "STAB1") |
| 35 | HALLMARK_WNT_BETA_CATENIN_SIGNALING | 0.388098 | 0.528245 | -0.519444 | -1.065977 | 3136 | 5 | c("CSNK1E", "CTNNB1", "NCSTN", "NCOR2") |

| | | | | | | | | |
|---|---|---|---|---|---|---|---|---|
| 36 | HALLMARK_ADIPOGENESIS | 0.385754 | 0.528245 | -0.334772 | -1.058054 | 3855 | 52 | c("COL4A1", "SPARCL1", "C3", "HSPB8", "FABP4", "PHLDB1", "TOB1", "CAVIN1", "COX6A1", "CD151", "ALDH2", "APOE", "LAMA4", "GPX3", "VEGFB", "CMBL", "GPX4", "GPAT4", "CCNG2", "UQCRQ", "UQCR11", "NDUFAB1", "UBC", "PDCD4", "DHRS7", "SCP2", "SDHB", "APLP2", "RAB34", "REEP5", "UQCRC1", "ESYT1", "SOD1", "NDUFB7", "CYC1", "UQCR10", "COX8A") |
| 37 | HALLMARK_UV_RESPONSE_UP | 0.411606 | 0.545099 | -0.339958 | -1.049284 | 4106 | 40 | c("NR4A1", "MMP14", "RET", "FOSB", "HMOX1", "FOS", "NFKBIA", "EPHX1", "IGFBP2", "GPX3", "EPCAM", "RHOB", "CREG1", "SELENOW", "PPT1", "ATP6V1F") |
| 38 | HALLMARK_PROTEIN_SECRETION | 0.491013 | 0.633149 | -0.331024 | -1.004131 | 4889 | 34 | c("TPD52", "DST", "TMED2", "TOM1L1", "USO1", "PAM", "SEC31A", "PPT1", "ARFGEF2", "LAMP2", "COPB2", "COPE", "ERGIC3", "RAB2A", "NAPA", "VPS45", "ARF1", "SNX2", "CD63", "ARCN1", "YKT6") |
| 39 | HALLMARK_IL6_JAK_STAT3_SIGNALING | 0.595979 | 0.748794 | -0.329571 | -0.935692 | 5839 | 21 | c("CD9", "JUN", "A2M", "HMOX1", "SOCS3", "IL13RA1", "STAT3", "CD44", "MAP3K8", "CD14", "STAT1", "IL1R1") |
| 40 | HALLMARK_MITOTIC_SPINDLE | 0.679672 | 0.832598 | -0.285844 | -0.910195 | 6793 | 56 | c("PALLD", "FLNB", "SHROOM1", "FARP1", "DST", "ABL1", "ARHGAP29", "NET1", "CTTN", "TRIO", "EPB41L2", "MARCKS", "YWHAE", "DYNLL2", "SPTAN1", "FLNA", "GSN", "NOTCH2", "BCAR1", "ARFIP2", "CAPZB", "WASL", "ARHGEF12", "EZR", "PREX1", "CYTH2", "MYH9", "KIF5B", "STAU1", "ACTN4", "RASA1", "ARL8A", "NEDD9") |
| 41 | HALLMARK_UNFOLDED_PROTEIN_RESPONSE | 0.758478 | 0.884891 | -0.278843 | -0.836993 | 7536 | 31 | c("STC2", "YWHAZ", "SHC1", "SPCS1", "EIF4G1", "DDIT4", "CALR", "SEC31A", "EIF4A2", "LSM1", "ATP6V0D1", "ATF4", "EIF4A3", "DCTN1", "VEGFA", "KIF5B", "EIF4EBP1", "NHP2", "LSM4", "XBP1", "EEF2", "HERPUD1") |
| 42 | HALLMARK_ALLOGRAFT_REJECTION | 0.756211 | 0.884891 | -0.281328 | -0.829561 | 7487 | 27 | c("TIMP1", "THY1", "TPD52", "CD74", "B2M", "STAB1") |
| 43 | HALLMARK_OXIDATIVE_PHOSPHORYLATION | 0.8432 | 0.951721 | -0.253322 | -0.818522 | 8431 | 68 | c("ATP1B1", "MAOB", "PDK4", "COX6C", "NDUFC2", "NDUFA4", "COX6A1", "ATP5F1B", "ATP6AP1", "NDUFB2", "GPX4", "COX7C", "ATP6V0E1", "ATP6V1F", "ATP6V0C", "UQCRQ", "UQCRB", "COX7A2", "UQCR11", "NDUFAB1", "FH", "ATP6V0B", "SDHB", "COX4I1", "ATP5MC2", "ATP5ME", "NDUFS4", "NDUFA1", "OXA1L", "UQCRC1", "ATP5F1E", "GLUD1", "CYB5R3", "ISCU", "HADHA", "NDUFB7", "CYB5A", "ATP5F1D", "CYC1", "UQCR10", "COX8A", "ECI1", "ATP5MG", "ETFB", "ATP5PD") |
| 44 | HALLMARK_BILE_ACID_METABOLISM | 0.868966 | 0.951721 | 0.177221 | 0.722588 | 503 | 13 | c("KLF1", "CAT") |
| 45 | HALLMARK_PEROXISOME | 0.87403 | 0.951721 | -0.254451 | -0.696272 | 8443 | 17 | c("CRABP2", "SEMA3C", "DHCR24") |
| 46 | HALLMARK_DNA_REPAIR | 0.972834 | 0.984237 | -0.182895 | -0.53931 | 9632 | 27 | c("BCAM", "TMED2", "GPX4", "SEC61A1", "BRF2", "NME1", "IMPDH2", "CANT1", "APRT", "POLR2H", "POLR2I", "SUPT5H", "POLD4", "NUDT21", "NCBP2", "ERCC1", "RBX1", "NME3", "ADRM1") |
| 47 | HALLMARK_G2M_CHECKPOINT | 0.978683 | 0.984237 | -0.180311 | -0.542936 | 9732 | 32 | c("CCND1", "SS18", "MT2A", "ABL1", "SLC12A2", "HSPA8", "MARCKS", "NOTCH2", "BCL3", "EWSR1", "KIF5B", "SLC38A1", "NCL", "CUL3") |
| 48 | HALLMARK_MYC_TARGETS_V1 | 0.979375 | 0.984237 | -0.182666 | -0.569649 | 9781 | 45 | c("CANX", "CLNS1A", "CCT3", "HSP90AB1", "PPIA", "YWHAE", "PCBP1", "NME1", "NDUFAB1", "XRCC6", "PSMD3", "SNRPD2", "RNPS1", "IMPDH2", "ILF2", "RACK1", "CNBP", "HNRNPC", "PSMA7", "NHP2", "CYC1", "HNRNPD", "FAM120A", "TUFM", "HSPE1", "POLD2", "SRM", "HNRNPU", "NCBP2") |
| 49 | HALLMARK_REACTIVE_OXYGEN_SPECIES_PATHWAY | 0.984237 | 0.984237 | -0.177 | -0.49002 | 9552 | 18 | c("NQO1", "GPX3", "GPX4", "JUNB", "ATOX1", "SOD2", "FTL", "PTPA", "SOD1", "EGLN2") |
| 50 | HALLMARK_HEME_METABOLISM | NA | NA | 0.520257 | NA | 0 | 76 | c("SPTB", "AHSP", "TFRC", "SPTA1", "SLC4A1", "KLF1", "OSBP2", "HMBS", "HBD", "UBAC1", "NFE2", "KEL", "CA1", "TMCC2", "TRAK2", "SLC25A37", "TFDP2", "DMTN", "UROD", "NNT", "CAT", "NARF", "HAGH", "CPOX", "ANK1", "PPOX", "RBM38", "KLF3", "MAP2K3", "CA2", "PIGQ", "FBXO7", "ALAD", "SNCA", "ALAS2", "PRDX2", "GLRX5", "KAT2B", "CCND3", "SLC2A1", "LMO2", "GYPC", "HBB", "BPGM", "BLVRB", "HDGF", "RBM5") |

| Cluster 8 | pathway | pval | padj | ES | NES | nMoreExtreme | size | leadingEdge |
|---|---|---|---|---|---|---|---|---|
| 1 | HALLMARK_ESTROGEN_RESPONSE_EARLY | 1.00E-04 | 0.001667 | -0.636994 | -1.881613 | 0 | 77 | c("RET", "LRIG1", "GFRA1", "TFF1", "SLC7A2", "MAPT", "STC2", "SEC14L2", "HSPB8", "PGR", "PRSS23", "CA12", "CCND1", "SCNN1A", "RHOD", "KRT8", "NBL1", "THSD4", "MYB", "FHL2", "TFF3", "ADCY1", "ELF3", "GJA1", "RHOBTB3", "FLNB", "SLC39A6", "KRT19", "PLAAT3", "FAM102A", "CXCL12", "GREB1", "FASN", "HES1", "MUC1", "CLDN7", "SLC9A3R1", "TTC39A", "SEMA3B", "FOS", "TOB1", "IGFBP4", "MAST4", "GAB2", "IGF1R", "MLPH", "SIAH2", "MED13L", "ELOVL5", "MINDY1", "ABHD2", "BCL2", "FARP1") |
| 2 | HALLMARK_ESTROGEN_RESPONSE_LATE | 0.0001 | 0.001667 | -0.624363 | -1.831683 | 0 | 69 | c("RET", "SERPINA3", "AGR2", "TFF1", "MAPT", "CPE", "HSPB8", "PGR", "PRSS23", "SCUBE2", "CXCL14", "CA12", "CCND1", "SCNN1A", "LTF", "PRLR", "CDH1", "NBL1", "MYB", "TFF3", "MDK", "FLNB", "KRT19", "PLAAT3", "FAM102A", "CXCL12", "SLC9A3R1", "COX6C", "DLG5", "TSPAN13", "SEMA3B", "FOS", "TOB1", "IGFBP4", "LLGL2", "SIAH2", "PDCD4", "TOP2A", "UGDH", "ELOVL5", "ATP2B4", "ABHD2", "BCL2", "FARP1", "SERPINA1", "DNAJC1", "ADD3", "NRIP1") |
| 3 | HALLMARK_EPITHELIAL_MESENCHYMAL_TRANSITION | 1.00E-04 | 0.001667 | -0.608853 | -1.829284 | 0 | 107 | c("FSTL1", "CCN1", "THBS1", "SFRP4", "ELN", "LOXL2", "THBS2", "COL5A2", "FBLN2", "COL12A1", "FAP", "COMP", "LUM", "COL11A1", "THY1", "MXRA5", "FBLN1", "CCN2", "HTRA1", "POSTN", "TNC", "BGN", "COL5A1", "PMEPA1", "ACTA2", "GPC1", "FBN1", "LOXL1", "CALD1", "PRRX1", "LRRC15", "TIMP3", "GJA1", "MYL9", "SDC1", "ADAM12", "CDH11", "COL4A1", "CTHRC1", "GEM", "PDGFRB", "IGFBP3", "DCN", "CXCL12", "INHBA", "NNMT", "TPM2", "DPYSL3", "TAGLN", "PCOLCE", "COL16A1", "TPM1", "IGFBP2", "LOX", "IGFBP4", "COL4A2", "MMP2", □ "RHOB", "COL3A1", "COL6A3") |
| 4 | HALLMARK_MYOGENESIS | 0.002407 | 0.030084 | -0.540095 | -1.538012 | 23 | 44 | c("APOD", "AEBP1", "STC2", "HSPB8", "CLU", "SPDEF", "PVALB", "TNNT1", "CNN3", "ADAM12", "IGFBP7", "IGFBP3", "ERBB3", "TPM2", "TAGLN", "COL4A2", "COL3A1", "COL6A3", "DTNA", "COL1A1", "COL6A2", "AGRN", "APP", "MEF2D", "GSN", "MYO1C", "SPTAN1", "BHLHE40", "MYH9", "SPARC") |
| 5 | HALLMARK_COAGULATION | 0.003317 | 0.033169 | -0.55099 | -1.548806 | 32 | 38 | c("THBS1", "PLAT", "PRSS23", "CLU", "COMP", "HTRA1", "MMP11", "FBN1", "CRIP2", "S100A1", "TIMP3", "CFB", "C1R", "SERPING1", "C3", "S100A13", "MMP2", "C1S", "SERPINA1", "FN1", "TIMP1", "GSN", "SERPINE1") |
| 6 | HALLMARK_NOTCH_SIGNALING | 0.014699 | 0.097196 | -0.759987 | -1.54635 | 119 | 6 | c("CCND1", "HES1", "NOTCH3") |
| 7 | HALLMARK_UV_RESPONSE_DN | 0.015551 | 0.097196 | -0.497645 | -1.413732 | 154 | 43 | c("INPP4B", "CCN1", "EFEMP1", "COL5A2", "COL11A1", "IGFBP5", "FHL2", "GJA1", "PDGFRB", "PIK3R3", "ERBB2", "APBB2", "DUSP1", "IGF1R", "COL3A1", "AKT3", "RBPMS", "COL1A1", "COL1A2", "ATP2B4", "ADD3", "LAMC1", "SERPINE1") |
| 8 | HALLMARK_KRAS_SIGNALING_UP | 0.012967 | 0.097196 | -0.522233 | -1.462256 | 128 | 36 | c("SERPINA3", "APOD", "CPE", "PLAT", "MMP11", "JUP", "SPARCL1", "PRRX1", "CFB", "ITGBL1", "IGFBP3", "INHBA", "TSPAN13", "SEMA3B", "GYPC") |
| 9 | HALLMARK_ANDROGEN_RESPONSE | 0.0313 | 0.173892 | -0.513065 | -1.414952 | 309 | 31 | c("INPP4B", "BMPR1B", "AZGP1", "SPDEF", "TPD52", "CCND1", "PMEPA1", "KRT8", "KRT19", "STEAP4", "SELENOP", "UAP1", "ELOVL5", "ABHD2") |
| 10 | HALLMARK_BILE_ACID_METABOLISM | 0.036723 | 0.180852 | 0.394737 | 1.551802 | 25 | 13 | c("CYP27A1", "ABCA1", "HSD17B4", "SLC23A2", "ALDH1A1", "OPTN") |
| 11 | HALLMARK_KRAS_SIGNALING_DN | 0.039788 | 0.180852 | -0.613109 | -1.460195 | 366 | 12 | c("CPB1", "BMPR1B", "EFHD1", "GPRC5C", "IGFBP2", "SELENOP", "BTG2", "IDUA", "SKIL") |
| 12 | HALLMARK_ALLOGRAFT_REJECTION | 0.047393 | 0.197472 | 0.287069 | 1.43622 | 9 | 23 | c("CAPG", "CCL2", "CTSS", "SPI1", "ITGB2", "FGR", "MMP9", "CD4", "IRF8", "TAPBP", "CD74", "B2M") |
| 13 | HALLMARK_ANGIOGENESIS | 0.056288 | 0.216491 | -0.565527 | -1.40819 | 533 | 15 | c("FSTL1", "COL5A2", "LUM", "POSTN", "STC1", "FGFR1", "COL3A1", "APP", "VCAN", "TIMP1") |
| 14 | HALLMARK_HYPOXIA | 0.064632 | 0.222222 | -0.435764 | -1.269653 | 645 | 61 | c("CCN1", "STC2", "CA12", "TPD52", "CCN2", "BGN", "TGFB3", "COL5A1", "GPC1", "STC1", "IGFBP3", "SELENBP1", "DCN", "EFNA1", "CCNG2", "DUSP1", "FOS", "LOX", "CAVIN1", "PAM", "DTNA", "SIAH2", "MT2A", "BCL2", "PNRC1", "MIF") |
| 15 | HALLMARK_P53_PATHWAY | 0.066667 | 0.222222 | 0.213597 | 1.435756 | 1 | 45 | c("HMOX1", "IFI30", "GM2A", "CTSD", "FUCA1", "RAP2B", "NUPR1", "SAT1", "IRAK1", "UPP1", "CEBPA") |
| 16 | HALLMARK_APICAL_JUNCTION | 0.080188 | 0.241741 | -0.438585 | -1.264609 | 800 | 52 | c("THY1", "CDH1", "FBN1", "JUP", "PARVA", "MYL9", "MDK", "CDH11", "CLDN4", "LIMA1", "PIK3R3", "CLDN7", "PCDH1", "COL16A1", "CERCAM", "MMP2", "AKT3") |
| 17 | HALLMARK_INFLAMMATORY_RESPONSE | 0.082192 | 0.241741 | 0.246321 | 1.349631 | 11 | 27 | c("CXCL8", "MARCO", "CCL2", "PLAUR", "ABCA1", "CD14", "EMP3", "MSR1", "CYBB", "TAPBP") |
| 18 | HALLMARK_IL6_JAK_STAT3_SIGNALING | 0.120388 | 0.334412 | 0.311907 | 1.294975 | 61 | 15 | c("HMOX1", "CD14", "MYD88", "CD9", "JUN", "CD44", "TYK2", "TNFRSF1A", "A2M", "IL1R1", "IL13RA1", "MAP3K8", "TNFRSF12A", "SOCS3", "STAT3") |
| 19 | HALLMARK_APICAL_SURFACE | 0.163957 | 0.431465 | -0.564676 | -1.264132 | 1441 | 9 | c("GSTM3", "THY1", "GATA3", "SULF2", "APP") |

| | | | | | | | | |
|---|---|---|---|---|---|---|---|---|
| 20 | HALLMARK_APOPTOSIS | 0.24577 | 0.589508 | -0.393925 | -1.135838 | 2454 | 52 | c("PLAT", "CLU", "LUM", "CCND1", "BGN", "TIMP3", "PDGFRB", "DCN", "ERBB3", "ERBB2", "CTNNB1", "MMP2", "RHOB", "IFITM3", "PDCD4", "TOP2A", "BTG2", "NEDD9", "APP", "TIMP1", "HMGB2", "GSN", "RARA", "TXNIP", "ANKH", "SPTAN1") |
| 21 | HALLMARK_GLYCOLYSIS | 0.247593 | 0.589508 | -0.401996 | -1.145651 | 2468 | 45 | c("STC2", "COL5A1", "GPC1", "STC1", "TFF3", "ELF3", "SDC1", "FUT8", "IGFBP3", "DCN", "PAM", "CHPF", "AGRN", "EGLN3", "MIF", "GFPT1", "VCAN") |
| 22 | HALLMARK_PANCREAS_BETA_CELLS | 0.351567 | 0.799017 | -0.562853 | -1.095193 | 2769 | 5 | c("SYT13", "AKT3", "SRP9", "SPCS1") |
| 23 | HALLMARK_TGF_BETA_SIGNALING | 0.439766 | 0.956013 | -0.398968 | -1.04807 | 4281 | 21 | c("THBS1", "CDH1", "PMEPA1", "CTNNB1", "LTBP2", "FNTA", "ID3", "SERPINE1", "SKIL") |
| 24 | HALLMARK_HEDGEHOG_SIGNALING | 0.461285 | 0.96101 | -0.458701 | -1.026888 | 4056 | 9 | c("THY1", "ADGRG1", "MYH9", "CELSR1", "LDB1", "DPYSL2", "VEGFA") |
| 25 | HALLMARK_MITOTIC_SPINDLE | 0.491941 | 0.961303 | -0.350392 | -1.010314 | 4913 | 52 | c("NET1", "SHROOM1", "FLNB", "PALLD", "ARHGAP29", "CTTN", "ARHGEF12", "TOP2A", "NEDD9", "ARFIP2", "SMC4", "NUMA1", "FARP1", "EPB41L2", "GSN", "CYTH2", "STAU1", "DST", "SPTAN1", "RANBP9", "MYH9") |
| 26 | HALLMARK_SPERMATOGENESIS | 0.499878 | 0.961303 | -0.487762 | -0.992453 | 4080 | 6 | c("GSTM3", "SLC12A2") |
| 27 | HALLMARK_TNFA_SIGNALING_VIA_NFKB | 0.605003 | 1 | -0.326831 | -0.954555 | 6046 | 63 | c("CCN1", "CCND1", "NR4A1", "TNC", "PMEPA1", "DUSP4", "EGR1", "FOSB", "PLK2", "GEM", "INHBA", "HES1", "EFNA1", "DUSP1", "FOS", "RHOB") |
| 28 | HALLMARK_CHOLESTEROL_HOMEOSTASIS | 0.704617 | 1 | -0.335384 | -0.852881 | 6745 | 17 | c("CLU", "FASN", "SEMA3B", "CTNNB1", "ACTG1", "PNRC1") |
| 29 | HALLMARK_WNT_BETA_CATENIN_SIGNALING | 0.619622 | 1 | -0.469432 | -0.913416 | 4881 | 5 | c("CTNNB1", "CSNK1E", "MYC") |
| 30 | HALLMARK_DNA_REPAIR | 0.96639 | 1 | -0.205494 | -0.560875 | 9545 | 28 | c("BCAM", "IMPDH2", "CANT1", "NUDT21", "NME3", "POLR2H", "BRF2", "NME4", "NME1", "NELFCD") |
| 31 | HALLMARK_G2M_CHECKPOINT | 0.748735 | 1 | -0.307361 | -0.83891 | 7395 | 28 | c("CCND1", "SLC12A2", "TOP2A", "SLC38A1", "MT2A", "SMC4", "NUMA1", "MYC", "HNRNPU", "BUB3", "SMAD3", "SMARCC1", "HNRNPD", "RBM14", "KIF5B", "CDKN1B", "ABL1", "BCL3", "ATRX", "TOP1", "NCL", "SS18") |
| 32 | HALLMARK_ADIPOGENESIS | 0.786416 | 1 | -0.294217 | -0.844806 | 7849 | 50 | c("HSPB8", "PHLDB1", "SPARCL1", "COL4A1", "CMBL", "CCNG2", "C3", "TOB1", "LAMA4", "CAVIN1", "PDCD4", "GPAT4", "FABP4") |
| 33 | HALLMARK_PROTEIN_SECRETION | 1 | 1 | 0.103249 | 0.60728 | 97 | 31 | c("IGF2R", "M6PR", "ABCA1", "CD63", "PPT1") |
| 34 | HALLMARK_INTERFERON_ALPHA_RESPONSE | 0.971799 | 1 | -0.204938 | -0.521158 | 9303 | 17 | c("IFITM3", "C1S", "IFITM2", "IFI27", "TXNIP") |
| 35 | HALLMARK_INTERFERON_GAMMA_RESPONSE | 0.966734 | 1 | -0.209277 | -0.585093 | 9618 | 35 | c("CFB", "C1R", "SERPING1", "IFITM3", "C1S", "IFITM2", "MT2A", "STAT3", "IFI27", "TXNIP", "SOCS3", "ARID5B") |
| 36 | HALLMARK_COMPLEMENT | 0.622928 | 1 | -0.332247 | -0.937779 | 6199 | 40 | c("MMP13", "PLAT", "CLU", "LTF", "GATA3", "CFB", "C1R", "SERPING1", "C3", "COL4A2", "S100A13", "C1S") |
| 37 | HALLMARK_UNFOLDED_PROTEIN_RESPONSE | 0.997583 | 1 | -0.135629 | -0.37677 | 9904 | 33 | c("STC2", "DNAJA4", "EEF2", "XBP1", "DDIT4", "SHC1", "LSM1", "BANF1", "FUS", "SPCS1", "SEC31A", "VEGFA", "NHP2") |
| 38 | HALLMARK_PI3K_AKT_MTOR_SIGNALING | 0.92891 | 1 | 0.144444 | 0.722663 | 195 | 23 | c("DUSP3", "SLC2A1", "SQSTM1", "MYD88", "ARHGDIA", "CFL1") |
| 39 | HALLMARK_MTORC1_SIGNALING | 0.987275 | 1 | -0.186501 | -0.534306 | 9852 | 48 | c("IGFBP5", "STC1", "SYTL2", "PIK3R3", "SLC9A3R1", "SERPINH1", "BTG2", "ELOVL5", "EGLN3", "ADD3", "ACACA", "XBP1", "DDIT4", "BHLHE40", "PSMB5", "SLC1A5", "DHCR24") |
| 40 | HALLMARK_E2F_TARGETS | 0.785149 | 1 | -0.303039 | -0.79607 | 7644 | 21 | c("TOP2A", "SMC4", "HMGB2", "RBBP7", "NUDT21", "MCM7", "MYC", "POLD2", "NME1", "TUBB", "HNRNPD", "ANP32E", "CDKN1B", "PRKDC", "SSRP1", "LUC7L3", "ILF3", "PNN", "SNRPB") |
| 41 | HALLMARK_MYC_TARGETS_V1 | 0.797833 | 1 | -0.29078 | -0.826059 | 7951 | 43 | c("CLNS1A", "HSP90AB1", "IMPDH2", "GNL3", "CBX3", "ILF2", "CCT3", "MCM7", "MYC", "RACK1", "PCBP1", "SNRPD2", "HNRNPU", "POLD2", "YWHAE", "BUB3", "RNPS1", "NME1", "NHP2", "SMARCC1", "VDAC3", "HDGF", "PPIA", "HNRNPD") |
| 42 | HALLMARK_MYC_TARGETS_V2 | 0.831793 | 1 | -0.348461 | -0.737099 | 7021 | 7 | c("GNL3", "CBX3", "MYC") |
| 43 | HALLMARK_XENOBIOTIC_METABOLISM | 0.791319 | 1 | -0.29347 | -0.828331 | 7875 | 40 | c("FBLN1", "JUP", "ESR1", "CFB", "PDK4", "IGFBP4", "SPINT2", "MCCC2", "UGDH", "ELOVL5", "MT2A", "CSAD", "SERPINE1") |
| 44 | HALLMARK_FATTY_ACID_METABOLISM | 0.853557 | 1 | -0.273768 | -0.740458 | 8398 | 26 | c("BMPR1B", "FASN", "PRDX6", "UGDH", "ELOVL5", "MIF", "FH", "ECI1", "DHCR24") |
| 45 | HALLMARK_OXIDATIVE_PHOSPHORYLATION | 1 | 1 | -0.095893 | -0.276892 | 9988 | 53 | c("MAOB", "PDK4", "COX6C", "ATP1B1", "COX6B1", "NDUFC2", "COX6A1", "FH", "ECI1", "IDH2", "UQCRQ", "VDAC3", "ATP5F1B", "ISCU", "CYB5R3", "NDUFC1", "IDH3B", "NDUFS2", "NDUFA4") |
| 46 | HALLMARK_REACTIVE_OXYGEN_SPECIES_PATHWAY | 0.967307 | 1 | -0.210503 | -0.535308 | 9260 | 17 | c("MPO", "PRDX2", "PRDX6") |
| 47 | HALLMARK_UV_RESPONSE_UP | 0.577443 | 1 | -0.348055 | -0.959878 | 5718 | 31 | c("RET", "EPCAM", "NR4A1", "FOSB", "IGFBP2", "FOS", "RHOB") |
| 48 | HALLMARK_HEME_METABOLISM | 0.752854 | 1 | -0.301458 | -0.867214 | 7517 | 51 | c("HBB", "SLC4A1", "BCAM", "HBD", "PRDX2", "SELENBP1", "C3", "GYPC", "SLC25A37", "ARHGEF12", "BTG2", "FBXO7", "HAGH", "GDE1", "FBXO9", "EIF2AK1", "FOXO3", "LRP10", "VEZF1", "ADD1") |

| | | | | | | | | |
|---|---|---|---|---|---|---|---|---|
| 49 | HALLMARK_IL2_STAT5_SIGNALING | 0.565047 | 1 | -0.342339 | -0.970261 | 5628 | 42 | c("LRIG1", "S100A1", "MUC1", "ENPP1", "NFKBIZ", "IGF1R", "RHOB", "IFITM3", "ITGA6", "COL6A1", "BCL2", "MYO1C", "ECM1", "CKAP4", "XBP1", "MAPKAPK2", "ITGAV", "BHLHE40", "PHLDA1", "MYC", "ALCAM", "SLC1A5") |
| 50 | HALLMARK_PEROXISOME | 0.949797 | 1 | -0.222215 | -0.570568 | 9137 | 18 | c("CRABP2", "TOP2A", "ELOVL5", "IDH2", "DHCR24", "FIS1", "SMARCC1", "LONP2") |

| Cluster 9 | pathway | pval | padj | ES | NES | nMoreExtreme | size | leadingEdge |
|---|---|---|---|---|---|---|---|---|
| 1 | HALLMARK_HYPOXIA | 0.000101 | 0.000735 | -0.565491 | -1.901091 | 0 | 63 | c("STC2", "BGN", "DCN", "CCN1", "MIF", "COL5A1", "ANXA2", "CCN2", "TGFBI", "JUN", "LOX", "CA12", "BHLHE40", "STC1", "SERPINE1", "IGFBP3", "HMOX1", "CAVIN1", "EFNA1", "TPD52", "SDC2", "TGFB3", "GPC1", "SDC4", "FOS", "DTNA", "DUSP1", "DDIT4", "SDC3", "FBP1", "MT2A", "PAM") |
| 2 | HALLMARK_ESTROGEN_RESPONSE_EARLY | 0.000101 | 0.000735 | -0.60544 | -2.074702 | 0 | 79 | c("NBL1", "STC2", "KRT19", "MAPT", "PGR", "HSPB8", "TFF1", "PRSS23", "ADCY1", "SLC39A6", "MLPH", "SLC7A2", "KRT8", "SEC14L2", "CA12", "BHLHE40", "PLAAT3", "IGFBP4", "GFRA1", "THSD4", "ELF3", "CCND1", "GJA1", "MUC1", "FLNB", "HES1", "GREB1", "SCNN1A", "RET", "CLDN7", "FASN", "RHOD", "CELSR1", "FARP1", "FAM102A", "SEMA3B", "PAPSS2", "FOS", "TTC39A", "TOB1", "MYOF", "TFF3", "RHOBTB3", "XBP1") |
| 3 | HALLMARK_ESTROGEN_RESPONSE_LATE | 0.000101 | 0.000735 | -0.571395 | -1.948356 | 0 | 74 | c("NBL1", "PRLR", "KRT19", "MAPT", "PGR", "CXCL14", "HSPB8", "DLG5", "TFF1", "SERPINA3", "PRSS23", "AGR2", "CD9", "SCUBE2", "CA12", "PLAAT3", "IGFBP4", "CPE", "COX6C", "CCND1", "FLNB", "SCNN1A", "RET", "UGDH", "LSR", "MDK", "FARP1", "FAM102A", "SEMA3B", "PAPSS2", "CDH1", "FOS", "TOB1", "MYOF", "TFF3", "SERPINA1", "XBP1", "LLGL2") |
| 4 | HALLMARK_MYOGENESIS | 0.000103 | 0.000735 | -0.609604 | -1.994226 | 0 | 48 | c("STC2", "ADAM12", "HSPB8", "COL6A2", "APOD", "COL6A3", "PVALB", "SPDEF", "IGFBP7", "NQO1", "BHLHE40", "COL1A1", "TAGLN", "COL3A1", "CNN3", "CLU", "SCD", "IGFBP3", "ITGB5", "ERBB3", "TNNT1", "COL4A2", "AEBP1", "GPX3", "DTNA", "SPARC", "ATP6AP1", "SORBS3", "MYO1C", "PLXNB2", "ITGB1", "SPTAN1", "APP") |
| 5 | HALLMARK_EPITHELIAL_MESENCHYMAL_TRANSITIO N | 0.0001 | 0.000735 | -0.730446 | -2.556521 | 0 | 107 | c("SPP1", "POSTN", "SFRP4", "BGN", "HTRA1", "TIMP3", "FAP", "MMP2", "DCN", "COL5A2", "LUM", "ADAM12", "COMP", "THY1", "FBLN1", "ACTA2", "THBS2", "CCN1", "COL6A2", "CDH11", "COL6A3", "MMP14", "PRRX1", "COL5A1", "COL12A1", "FSTL1", "ELN", "CCN2", "COL16A1", "TGFBI", "TPM4", "CTHRC1", "DPYSL3", "FBLN2", "JUN", "LOX", "TNC", "IGFBP4", "LOXL1", "COL4A1", "SERPINE1", "COL11A1", "COL1A1", "TAGLN", "COL3A1", "LOXL2", "MXRA5", "TIMP1", "GJA1", "RHOB", "LRRC15", "IGFBP3", "GEM", "CALD1", "COL1A2", "PCOLCE", "LAMC1", "ITGB5", "MGP", "PFN2", "LRP1", "FBN1", "DAB2", "MYL9", "SDC1", "FN1", "COL4A2", "PMP22", "ITGAV", "PDGFRB", "LGALS1", "GPC1", "SDC4", "SERPINH1", "PMEPA1", "FUCA1", "SPARC", "NNMT", "IGFBP2") |
| 6 | HALLMARK_UV_RESPONSE_DN | 0.000103 | 0.000735 | -0.600479 | -1.956027 | 0 | 46 | c("COL5A2", "CCN1", "INPP4B", "EFEMP1", "ANXA2", "IGFBP5", "BHLHE40", "RBPMS", "SERPINE1", "COL11A1", "COL1A1", "COL3A1", "GJA1", "COL1A2", "LAMC1", "ERBB2", "NRP1", "SDC2", "DAB2", "PMP22", "PDGFRB", "DUSP1", "PLPP3", "APBB2", "AKT3", "ANXA4", "PIK3R3") |
| 7 | HALLMARK_COAGULATION | 0.000103 | 0.000735 | -0.711719 | -2.307937 | 0 | 44 | c("S100A1", "HTRA1", "TIMP3", "MMP2", "APOC1", "COMP", "C3", "MMP11", "C1R", "PLAT", "PRSS23", "C1S", "MMP14", "CD9", "CTSK", "SERPINE1", "A2M", "SERPING1", "CLU", "TIMP1", "CFB", "C1QA", "CRIP2", "LGMN", "LRP1", "S100A13", "FBN1", "FN1", "CTSB", "SPARC", "CAPN2", "SERPINA1", "CSRP1", "THBS1") |
| 8 | HALLMARK_APOPTOSIS | 0.001425 | 0.008465 | -0.500281 | -1.660529 | 13 | 55 | c("BGN", "TIMP3", "MMP2", "DCN", "LUM", "IFITM3", "PLAT", "JUN", "CCND1", "CLU", "TIMP1", "RHOB", "HMOX1", "ERBB2", "ERBB3", "EMP1", "PDGFRB", "GPX3", "PEA15", "HSPB1", "TIMP2", "CTNNB1", "LMNA", "SPTAN1", "GPX4", "APP", "RARA", "DAP", "GSN", "LGALS3", "PPT1", "CD44") |
| 9 | HALLMARK_APICAL_JUNCTION | 0.001524 | 0.008465 | -0.492831 | -1.643511 | 14 | 58 | c("MMP2", "THY1", "CDH11", "COL16A1", "TGFBI", "CD276", "PCDH1", "PARVA", "FBN1", "CLDN4", "CLDN7", "NECTIN2", "MDK", "MYL9", "ACTG1", "CDH1", "LIMA1", "CTNND1", "EVL", "JUP", "SDC3", "SORBS3", "ITGB1", "AKT3", "PIK3R3", "EPB41L2", "CERCAM", "TSPAN4", "CTNNA1", "VCAN", "SHC1", "WASL") |
| 10 | HALLMARK_KRAS_SIGNALING_UP | 0.001744 | 0.008722 | -0.506674 | -1.650463 | 16 | 46 | c("SPP1", "GPNMB", "MMP11", "SERPINA3", "PLAT", "APOD", "PRRX1", "SPARCL1", "CPE", "TMEM176B", "CFB", "IGFBP3", "NRP1", "ITGBL1", "EMP1", "SEMA3B", "FUCA1", "JUP", "MAFB") |
| 11 | HALLMARK_COMPLEMENT | 0.00193 | 0.008773 | -0.487143 | -1.624541 | 18 | 58 | c("APOC1", "C3", "C1R", "PLAT", "C1S", "MMP14", "C1QC", "GATA3", "SERPINE1", "LIPA", "SERPING1", "CLU", "TIMP1", "CFB", "C1QA", "CTSD", "LGMN", "LRP1", "S100A13", "FN1", "COL4A2", "CTSB", "TIMP2", "ANXA5", "SERPINA1", "CSRP1") |

| | | | | | | | | |
|---|---|---|---|---|---|---|---|---|
| 12 | HALLMARK_E2F_TARGETS | 0.002841 | 0.011837 | 0.622228 | 3.360468 | 0 | 40 | c("TFRC", "SPC24", "ANP32E", "HMMR", "MXD3", "PLK1", "MYBL2", "CENPM", "LBR", "HMGB2", "RRM2", "TACC3", "MKI67", "DCK", "HELLS", "EZH2", "LUC7L3", "SMC4", "PCNA", "DDX39A", "DNMT1", "SRSF2", "LIG1", "ASF1B", "TUBG1", "SRSF1", "TOP2A", "PPP1R8", "PNN", "MYC", "MCM4") |
| 13 | HALLMARK_CHOLESTEROL_HOMEOSTASIS | 0.003485 | 0.01271 | -0.624253 | -1.727434 | 30 | 17 | c("CD9", "CLU", "SCD", "LGMN", "FASN", "SEMA3B", "ACTG1", "ANXA5", "S100A11", "CTNNB1", "ALCAM", "LGALS3", "PMVK") |
| 14 | HALLMARK_G2M_CHECKPOINT | 0.003559 | 0.01271 | 0.348278 | 1.916458 | 0 | 45 | c("E2F2", "HMMR", "PLK1", "MYBL2", "KIF20B", "PRC1", "LBR", "TACC3", "MKI67", "TFDP1", "E2F4", "TPX2", "NUSAP1", "KMT5A", "EZH2", "SMC4", "SLC7A5", "DDX39A", "SRSF2") |
| 15 | HALLMARK_ANGIOGENESIS | 0.004237 | 0.014124 | -0.616553 | -1.727774 | 37 | 18 | c("SPP1", "POSTN", "COL5A2", "LUM", "FSTL1", "STC1", "COL3A1", "TIMP1", "NRP1", "ITGAV") |
| 16 | HALLMARK_ANDROGEN_RESPONSE | 0.008705 | 0.027202 | -0.486722 | -1.555992 | 83 | 40 | c("KRT19", "INPP4B", "SPDEF", "KRT8", "BMPR1B", "CCND1", "SELENOP", "SCD", "TPD52", "AZGP1", "ITGAV", "B2M", "PMEPA1", "DHCR24", "DBI", "SGK1", "UAP1", "ACSL3", "SMS", "ABHD2", "MAF") |
| 17 | HALLMARK_IL2_STAT5_SIGNALING | 0.010187 | 0.029963 | -0.456457 | -1.511975 | 99 | 54 | c("SPP1", "S100A1", "IFITM3", "COL6A1", "PHLDA1", "ENPP1", "BHLHE40", "CD81", "MUC1", "RHOB", "NRP1", "EMP1", "ITGAV", "ECM1", "XBP1", "MYO1C", "CYFIP1", "ALCAM", "AHNAK", "GPX4", "ANXA4", "LRIG1", "AHR", "BMPR2", "CAPG", "CTSZ", "PLEC", "SYNGR2", "IGF1R", "MAP3K8", "CD44", "SERPINB6", "MAPKAPK2", "HIPK2") |
| 18 | HALLMARK_TNFA_SIGNALING_VIA_NFKB | 0.011437 | 0.03177 | -0.439454 | -1.481098 | 112 | 65 | c("NR4A1", "CCN1", "PHLDA1", "EGR1", "JUN", "TNC", "BHLHE40", "SERPINE1", "PLK2", "CCND1", "RHOB", "HES1", "GEM", "DUSP4", "EFNA1", "FOSB", "SDC4", "FOS", "PMEPA1", "SOCS3", "EIF1", "DUSP1", "BCL3", "PLPP3", "NFKBIA", "SGK1") |
| 19 | HALLMARK_KRAS_SIGNALING_DN | 0.020775 | 0.054671 | -0.609391 | -1.584998 | 177 | 13 | c("CPB1", "EFHD1", "BMPR1B", "SELENOP", "IGFBP2", "GPRC5C", "SKIL", "SGK1") |
| 20 | HALLMARK_APICAL_SURFACE | 0.025415 | 0.063538 | -0.651036 | -1.547147 | 204 | 9 | c("THY1", "GATA3", "SULF2", "GSTM3", "HSPB1", "APP") |
| 21 | HALLMARK_GLYCOLYSIS | 0.035773 | 0.081302 | -0.4301 | -1.415648 | 349 | 50 | c("STC2", "DCN", "MIF", "COL5A1", "TGFBI", "STC1", "ELF3", "IGFBP3", "SDC2", "SDC1", "GPC1", "CHPF", "TFF3", "EGLN3", "DDIT4", "SDC3", "IL13RA1", "PAM", "PKM", "PPIA", "AGRN", "VCAN", "PLOD1", "CD44", "PGLS", "IDUA", "GFPT1", "ALDH9A1") |
| 22 | HALLMARK_HEME_METABOLISM | 0.034483 | 0.081302 | 0.617177 | 3.982802 | 0 | 92 | c("SPTB", "AHSP", "SLC4A1", "CA1", "SPTA1", "KEL", "KLF1", "HBD", "HMBS", "E2F2", "NFE2", "TMCC2", "OSBP2", "ANK1", "DMTN", "TFRC", "RBM38", "SLC25A37", "CA2", "CPOX", "TRIM58", "UBAC1", "SNCA", "ERMAP", "ALAS2", "UROD", "TRAK2", "CAT", "KAT2B", "PPOX", "TFDP2", "NARF", "HAGH", "FBXO7", "KLF3", "MAP2K3", "NNT", "LMO2", "HTRA2", "GYPC", "CCND3", "PRDX2", "RANBP10", "BPGM", "ALAD", "GCLC", "PIGQ", "USP15", "FECH", "SLC2A1", "SLC6A8", "TENT5C", "SLC25A38", "GLRX5", "FN3K", "ABCB6", "XPO7", "RAD23A", □<br>"KDM7A", "BLVRB", "SLC66A2", "HBB", "FBXO9", "TOP1", "HDGF", "UCP2") |
| 23 | HALLMARK_P53_PATHWAY | 0.041141 | 0.089437 | -0.428678 | -1.402355 | 400 | 48 | c("NUPR1", "TM4SF1", "JUN", "IFI30", "S100A10", "CD81", "PLK2", "EPS8L2", "HMOX1", "CTSD", "SDC1", "FOS", "FUCA1", "TOB1", "DDIT4", "EPHX1", "ZFP36L1", "PLXNB2") |
| 24 | HALLMARK_INTERFERON_ALPHA_RESPONSE | 0.051676 | 0.107659 | -0.515048 | -1.45909 | 466 | 19 | c("IFITM3", "C1S", "IFI30", "IFI27", "B2M", "IFITM2", "LGALS3BP", "CD74", "LY6E") |
| 25 | HALLMARK_NOTCH_SIGNALING | 0.06557 | 0.131139 | -0.618511 | -1.427416 | 517 | 8 | c("NOTCH3", "CCND1", "HES1") |
| 26 | HALLMARK_INTERFERON_GAMMA_RESPONSE | 0.078342 | 0.150658 | -0.423022 | -1.352352 | 755 | 40 | c("IFITM3", "C1R", "C1S", "IFI30", "SERPING1", "CFB", "IFI27", "B2M", "SOCS3", "IFITM2", "MT2A", "LGALS3BP", "NFKBIA", "CD74", "STAT3", "LY6E") |
| 27 | HALLMARK_ADIPOGENESIS | 0.081798 | 0.150962 | -0.403678 | -1.325402 | 798 | 49 | c("HSPB8", "C3", "SPARCL1", "APOE", "PHLDB1", "COL4A1", "CAVIN1", "FABP4", "GPX3", "TOB1", "CD151", "VEGFB", "LAMA4", "COX6A1", "GPX4", "RAB34", "CMBL", "ALDH2", "GPAT4", "UQCRQ", "CCNG2", "APLP2", "DHRS7", "NDUFAB1", "UBC", "UQCR11", "SCP2", "UQCR10", "PDCD4", "REEP5") |
| 28 | HALLMARK_XENOBIOTIC_METABOLISM | 0.084539 | 0.150962 | -0.40389 | -1.321263 | 823 | 48 | c("FBLN1", "NQO1", "IGFBP4", "APOE", "SERPINE1", "TMEM176B", "CFB", "ESR1", "HMOX1", "PDK4", "UGDH", "PAPSS2", "SPINT2", "JUP", "FBP1", "MT2A", "EPHX1", "PGRMC1", "BCAR1") |
| 29 | HALLMARK_MYC_TARGETS_V2 | 0.10213 | 0.176085 | 0.494042 | 1.40594 | 234 | 7 | c("PLK1", "MYC", "MCM4", "PA2G4", "CBX3", "SRM", "HSPE1") |
| 30 | HALLMARK_TGF_BETA_SIGNALING | 0.159207 | 0.256785 | -0.430778 | -1.27425 | 1468 | 24 | c("SERPINE1", "LTBP2", "CDH1", "PMEPA1", "CTNNB1", "SKIL", "THBS1", "BMPR2", "ID3", "ENG", "RHOA", "HIPK2", "JUNB") |

| | | | | | | | | |
|---|---|---|---|---|---|---|---|---|
| 31 | HALLMARK_ALLOGRAFT_REJECTION | 0.156069 | 0.256785 | 0.240886 | 1.244289 | 80 | 32 | c("ACHE", "ELANE", "NCF4", "FGR", "SRGN", "PRKCB", "CCND3", "LTB", "HCLS1", "WAS") |
| 32 | HALLMARK_PANCREAS_BETA_CELLS | 0.200364 | 0.313068 | -0.558738 | -1.242666 | 1542 | 7 | c("SYT13", "MAFB", "AKT3", "SPCS1") |
| 33 | HALLMARK_PI3K_AKT_MTOR_SIGNALING | 0.228614 | 0.346384 | 0.238668 | 1.163742 | 154 | 27 | c("MAP2K3", "PRKCB", "SLA", "SLC2A1", "GRK2", "CXCR4", "MYD88", "PFN1", "ARPC3", "PAK4", "AKT1", "ARHGDIA", "UBE2D3", "ITPR2", "PPP1CA", "GRB2", "GSK3B", "STAT2", "CAB39", "ACACA", "SQSTM1", "VAV3", "CFL1", "CALR", "ARF1", "PIK3R3", "DUSP3") |
| 34 | HALLMARK_REACTIVE_OXYGEN_SPECIES_PATHWAY | 0.248549 | 0.365514 | 0.274673 | 1.15792 | 256 | 18 | c("MPO", "CAT", "CDKN2D", "PRDX2", "MSRA", "GCLC") |
| 35 | HALLMARK_FATTY_ACID_METABOLISM | 0.258958 | 0.36994 | -0.387537 | -1.176016 | 2420 | 28 | c("MIF", "S100A10", "BMPR1B", "UGDH", "FASN", "LGALS1", "GLUL", "PCBD1", "DHCR24", "EPHX1") |
| 36 | HALLMARK_SPERMATOGENESIS | 0.323133 | 0.448795 | -0.591945 | -1.118267 | 2227 | 4 | c("GSTM3", "SLC12A2", "PEBP1") |
| 37 | HALLMARK_BILE_ACID_METABOLISM | 0.359242 | 0.485463 | 0.291525 | 1.055304 | 549 | 12 | c("KLF1", "CAT") |
| 38 | HALLMARK_PROTEIN_SECRETION | 0.457917 | 0.602523 | -0.329667 | -1.027267 | 4357 | 33 | c("TPD52", "TOM1L1", "TMED2", "PAM", "DST", "SEC31A", "ARF1", "USO1", "RAB2A", "ARFGEF2", "PPT1", "ERGIC3", "CD63", "LAMP2", "COPE", "COPB2", "VPS45") |
| 39 | HALLMARK_UV_RESPONSE_UP | 0.476565 | 0.610981 | -0.317513 | -1.017607 | 4605 | 41 | c("NR4A1", "MMP14", "RHOB", "HMOX1", "RET", "FOSB", "GPX3", "ATP6V1F", "FOS", "IGFBP2", "EPCAM", "EPHX1", "NFKBIA") |
| 40 | HALLMARK_UNFOLDED_PROTEIN_RESPONSE | 0.505853 | 0.632316 | -0.321502 | -0.997104 | 4796 | 32 | c("STC2", "DDIT4", "XBP1", "YWHAZ", "SPCS1", "SEC31A", "CALR", "SHC1", "EIF4EBP1", "EIF4G1", "EIF4A2", "NHP2", "ATF4", "KIF5B", "EEF2", "LSM1", "EIF4A3", "DCTN1", "VEGFA", "LSM4", "ATP6V0D1") |
| 41 | HALLMARK_HEDGEHOG_SIGNALING | 0.554557 | 0.673744 | -0.388518 | -0.948109 | 4563 | 10 | c("THY1", "NRP1", "CELSR1", "DPYSL2", "ADGRG1", "MYH9") |
| 42 | HALLMARK_MTORC1_SIGNALING | 0.573468 | 0.673744 | -0.286571 | -0.963407 | 5662 | 63 | c("NUPR1", "CD9", "IGFBP5", "BHLHE40", "IFI30", "STC1", "SCD", "LGMN", "SERPINH1", "DHCR24", "CANX", "SYTL2", "EGLN3", "DDIT4", "XBP1", "PIK3R3", "ACSL3", "PPIA", "USO1", "ATP2A2", "CALR", "SLC9A3R1") |
| 43 | HALLMARK_INFLAMMATORY_RESPONSE | 0.57942 | 0.673744 | -0.311426 | -0.945049 | 5416 | 28 | c("MMP14", "SLC7A2", "SERPINE1", "TIMP1", "NFKBIA", "STAB1", "AHR", "LY6E", "ATP2A2", "BST2", "IL1R1", "CD14") |
| 44 | HALLMARK_IL6_JAK_STAT3_SIGNALING | 0.666162 | 0.757002 | -0.294512 | -0.876102 | 6161 | 25 | c("CD9", "JUN", "A2M", "HMOX1", "SOCS3", "IL13RA1", "STAT3", "MAP3K8", "CD44", "IL1R1", "CD14", "TNFRSF12A") |
| 45 | HALLMARK_OXIDATIVE_PHOSPHORYLATION | 0.682058 | 0.757842 | -0.267749 | -0.894944 | 6720 | 59 | c("MAOB", "ATP1B1", "COX6C", "PDK4", "ATP6V1F", "ATP6AP1", "COX6A1", "NDUFC2", "GPX4", "COX7C", "NDUFA4", "ATP6V0E1", "ATP5F1B", "NDUFB2", "UQCRQ", "UQCRB", "FH", "ATP6V0C", "NDUFAB1", "UQCR11", "ECI1", "NDUFS4", "UQCR10", "ATP5F1D", "ATP5F1E", "ATP5MC2", "CYB5A", "COX4I1") |
| 46 | HALLMARK_DNA_REPAIR | 0.708777 | 0.770409 | 0.178373 | 0.849366 | 532 | 25 | c("PNP", "HCLS1", "PCNA", "LIG1", "GUK1", "NME4", "NELFCD", "DCTN4", "ERCC1", "POLR2H", "ADRM1", "APRT", "NUDT21", "SUPT5H", "POLR2I", "POLD4", "NME3", "CANT1", "IMPDH2", "BRF2", "NME1", "SEC61A1", "GPX4", "BCAM", "TMED2") |
| 47 | HALLMARK_PEROXISOME | 0.772993 | 0.822333 | -0.279906 | -0.774556 | 6874 | 17 | c("CRABP2", "SEMA3C", "DHCR24") |
| 48 | HALLMARK_WNT_BETA_CATENIN_SIGNALING | 0.808075 | 0.841745 | -0.376038 | -0.758165 | 5843 | 5 | c("CTNNB1", "CSNK1E", "NCOR2", "NCSTN") |
| 49 | HALLMARK_MITOTIC_SPINDLE | 0.905095 | 0.923566 | -0.208476 | -0.700287 | 8935 | 62 | c("PALLD", "FLNB", "SHROOM1", "FARP1", "ARHGAP29", "SPTAN1", "DST", "CTTN", "NET1", "BCAR1", "DYNLL2", "EPB41L2", "FLNA", "TRIO", "GSN", "YWHAE", "MARCKS", "EZR", "WASL", "ABL1", "ARFIP2", "MYH9", "PREX1", "NEDD9", "CAPZB", "CYTH2", "KIF5B", "SEPTIN9", "AKAP13", "NOTCH2", "ARHGEF12", "NUMA1") |
| 50 | HALLMARK_MYC_TARGETS_V1 | 0.950718 | 0.950718 | -0.186292 | -0.599288 | 9201 | 42 | c("CANX", "CCT3", "CLNS1A", "HSP90AB1", "PPIA", "YWHAE", "SNRPD2", "HNRNPC", "NHP2", "NME1", "ILF2", "RNPS1", "NDUFAB1", "IMPDH2", "PCBP1", "RACK1", "PSMA7", "PSMD3", "CNBP", "HSPE1", "FAM120A", "POLD2", "SRM", "APEX1", "TXNL4A", "TUFM", "BUB3", "HNRNPU", "LSM7") |

| Cluster 10 | pathway | pval | padj | ES | NES | nMoreExtreme | size | leadingEdge |
|---|---|---|---|---|---|---|---|---|
| 1 | HALLMARK_COMPLEMENT | 0.00017 | 0.002157 | -0.544895 | -2.365338 | 0 | 60 | c("S100A9", "APOC1", "CTSS", "C1QA", "C3", "C1QC", "FCER1G", "LIPA", "LGMN", "CTSD", "CTSL", "PLAT", "CD36", "C1S", "PIM1", "EHD1", "LTF", "SERPING1", "LRP1", "CTSB", "STX4", "LAP3", "PLAUR", "C1R", "LTA4H") |
| 2 | HALLMARK_EPITHELIAL_MESENCHYMAL_TRANSITIO N | 0.000158 | 0.002157 | -0.443867 | -2.10716 | 0 | 96 | c("ELN", "FUCA1", "FBLN2", "SFRP4", "DAB2", "JUN", "WIPF1", "PFN2", "IL32", "GPC1", "DCN", "FBLN1", "LRP1", "COL16A1", "EMP3", "LOXL1", "NOTCH2", "MXRA5", "CALD1", "ACTA2", "PLAUR", "VCAN", "CDH11", "FBN1", "FSTL1", "DPYSL3", "TPM2", "VIM", "PCOLCE", "MYL9", "MMP2", "COLGALT1", "FAP", "LUM", "PDGFRB", "TAGLN", "THY1", "COL4A2", "NNMT", "IGFBP3", "HTRA1", "LAMC1", "ID2", "CCN2", "TPM4", "PRRX1", "CAPG", "SPP1", "LRRC15", "DST", "TNFRSF12A", "COL4A1", "SAT1", "COL1A1", "COL1A2", "SPARC") |
| 3 | HALLMARK_HEME_METABOLISM | 0.000167 | 0.002157 | -0.562721 | -2.501141 | 0 | 68 | c("HBB", "HBD", "SLC4A1", "SNCA", "C3", "GYPC", "SLC25A37", "MAP2K3", "UCP2", "TRAK2", "ACP5", "BACH1", "UROD", "RNF19A", "UBAC1", "MPP1", "CCND3", "TFRC", "CAT", "TNS1", "FOXO3", "RBM5", "CTSB") |
| 4 | HALLMARK_KRAS_SIGNALING_UP | 0.000173 | 0.002157 | -0.501434 | -2.123574 | 0 | 53 | c("CTSS", "IKZF1", "FUCA1", "ITGB2", "GYPC", "FCER1G", "CXCR4", "GPNMB", "PLAT", "LAPTM5", "APOD", "GLRX", "ITGBL1", "PECAM1", "ETS1", "ENG", "ANKH", "MAFB", "NIN", "SPARCL1", "PLAUR", "MMP9") |
| 5 | HALLMARK_ESTROGEN_RESPONSE_EARLY | 0.000279 | 0.002326 | 0.475891 | 2.551946 | 0 | 100 | c("GAB2", "DHRS2", "ANXA9", "RHOBTB3", "RASGRP1", "PGR", "SCNN1A", "SEC14L2", "PRSS23", "HSPB8", "ELOVL2", "TSKU", "PAPSS2", "MAPT", "SLC27A2", "KRT8", "STC2", "MED13L", "SEMA3B", "SIAH2", "CA12", "TOB1", "GREB1", "FRK", "GFRA1", "RHOD", "SLC9A3R1", "FAM102A", "KRT19", "CANT1", "WFS1", "MUC1", "AMFR", "XBP1", "DYNLT3", "GJA1", "KLF4", "RARA", "SYBU", "HES1", "ESRP2", "BHLHE40", "CLDN7", "FLNB", "SFN", "ITPK1", "ADCY1", "PDZK1", "SLC1A4", "FHL2", "ELOVL5", "NRIP1", "FKBP5", "OPN3", "KDM4B", "CD44", □ "GLA") |
| 6 | HALLMARK_ESTROGEN_RESPONSE_LATE | 0.000272 | 0.002326 | 0.359823 | 1.91771 | 0 | 96 | c("SERPINA5", "DHRS2", "CDH1", "ANXA9", "PGR", "SERPINA3", "SCNN1A", "PRSS23", "HSPB8", "SORD", "PAPSS2", "MAPT", "SLC27A2", "CHST8", "SERPINA1", "IDH2", "SEMA3B", "SIAH2", "COX6C", "CA12", "TOB1", "CPE", "CD9", "FRK", "PRKAR2B", "LSR", "SLC9A3R1", "UGDH", "PKP3", "FAM102A", "KRT19", "AGR2", "WFS1", "AMFR", "XBP1", "DLG5", "DYNLT3", "BATF", "GINS2", "KLF4", "LLGL2", "FLNB", "SFN", "ITPK1", "PDZK1", "SLC1A4", "ELOVL5", "NRIP1", "FKBP5", "PRLR", "OPN3", "CD44", "GLA") |
| 7 | HALLMARK_ALLOGRAFT_REJECTION | 0.000564 | 0.004031 | -0.603741 | -2.173348 | 2 | 27 | c("CTSS", "SPI1", "HCLS1", "CD74", "ITGB2", "STAB1", "SRGN", "CCND3", "ETS1", "PSMB10", "IFNGR2", "TAPBP", "MMP9") |
| 8 | HALLMARK_MYC_TARGETS_V1 | 0.001859 | 0.010326 | 0.335121 | 1.74937 | 6 | 86 | c("CLNS1A", "GOT2", "GLO1", "MCM5", "RSL1D1", "TOMM70", "PRPS2", "MCM2", "CANX", "HSPD1", "HSP90AB1", "HNRNPC", "PSMB3", "XRCC6", "IMPDH2", "RRM1", "NME1", "ERH", "CBX3", "ILF2", "SNRPD2", "PPIA", "VDAC3", "PSMC4", "RAD23B", "VDAC1", "PSMD3", "PCBP1", "POLD2", "HNRNPU", "PSMA4", "PSMA2", "FBL", "EEF1B2", "ETF1", "YWHAE", "EIF2S1", "GNL3", "UBE2E1", "LSM7", "EIF4E", "NHP2", "EIF3D", "NDUFAB1", "PCNA", "UBA2", "PPM1G", "APEX1", "CCT3", "PRDX4", "EIF4G2", "SRSF3", "SNRPA", "SF3B3", "RACK1", "PWP1", □ "HDGF", "TUFM", "DEK", "PSMB2") |
| 9 | HALLMARK_COAGULATION | 0.001741 | 0.010326 | -0.449392 | -1.891923 | 9 | 52 | c("APOC1", "C1QA", "C3", "S100A1", "LGMN", "PLAT", "C1S", "A2M", "GSN", "PECAM1", "CFD", "SERPING1", "LRP1", "CTSB", "CTSK", "C1R", "FBN1", "MMP9", "LTA4H") |
| 10 | HALLMARK_OXIDATIVE_PHOSPHORYLATION | 0.004428 | 0.022142 | 0.300642 | 1.610706 | 15 | 99 | c("NDUFA6", "NDUFC2", "GOT2", "NDUFB1", "IDH2", "FH", "COX6A1", "COX6C", "ATP6V1E1", "TOMM70", "MGST3", "PHYH", "TIMM17A", "TIMM9", "COX4I1", "UQCR10", "ATP5MC3", "ATP5MC2", "PDHX", "ATP1B1", "COX7C", "VDAC3", "COX7B", "NDUFA2", "NDUFA7", "VDAC1", "TOMM22", "UQCRC1", "COX7A2L", "ATP5F1C", "ATP5F1B", "NDUFA3", "HSD17B10", "NDUFC1", "NQO2", "ATP5PD", "NDUFA4", "ECI1", "OXA1L", "NDUFS7", "UQCRQ", "COX17", "HSPA9", "NDUFB7", "UQCR11", "ATP6V0E1", "MDH2", "NDUFS4", "COX6B1", "NDUFA1", "ATP5ME", "SLC25A4", □ "ATP5F1E") |

| | | | | | | | | |
|---|---|---|---|---|---|---|---|---|
| 11 | HALLMARK_UNFOLDED_PROTEIN_RESPONSE | 0.006119 | 0.027814 | 0.380203 | 1.751512 | 25 | 50 | c("LSM1", "STC2", "SERP1", "EIF4EBP1", "HSPA5", "WFS1", "XBP1", "EIF4A3", "EIF4G1", "HERPUD1", "SPCS1", "SLC1A4", "KDELR3", "PAIP1", "KIF5B", "ATF4", "VEGFA", "CALR", "SEC31A", "HSPA9", "PDIA6", "EIF2S1", "BAG3", "KHSRP", "LSM4", "SRPRB", "EIF4E", "SEC11A", "NHP2", "HYOU1", "ATF6", "EIF4A2", "SHC1") |
| 12 | HALLMARK_INTERFERON_GAMMA_RESPONSE | 0.007266 | 0.030273 | -0.437734 | -1.762295 | 40 | 43 | c("IFI30", "CD74", "SAMHD1", "C1S", "PIM1", "IFI27", "TNFAIP2", "TXNIP", "SERPING1", "ARID5B", "PSMB10", "LAP3", "TAPBP", "MYD88", "C1R", "SOD2", "STAT2") |
| 13 | HALLMARK_ADIPOGENESIS | 0.012702 | 0.048854 | -0.359834 | -1.619598 | 76 | 72 | c("FABP4", "APOE", "C3", "ALDH2", "GPX3", "RAB34", "UCP2", "CD36", "STOM", "VEGFB", "CAVIN1", "CAT", "LAMA4", "PHLDB1", "SPARCL1") |
| 14 | HALLMARK_E2F_TARGETS | 0.020735 | 0.074054 | 0.336989 | 1.560583 | 87 | 51 | c("TK1", "MCM5", "NUDT21", "MCM2", "CDKN2A", "UBR7", "XRCC6", "UBE2T", "MCM3", "NME1", "CTCF", "PHF5A", "LMNB1", "MYBL2", "UNG", "SNRPB", "CDKN1A", "TMPO", "LIG1", "ASF1B", "POLD2", "TOP2A", "RFC2") |
| 15 | HALLMARK_IL6_JAK_STAT3_SIGNALING | 0.030403 | 0.101343 | -0.488512 | -1.648982 | 156 | 21 | c("CD14", "JUN", "CD36", "PIM1", "HMOX1", "A2M", "IL1R1", "IFNGR2", "MYD88") |
| 16 | HALLMARK_HYPOXIA | 0.03932 | 0.122876 | -0.319555 | -1.470108 | 242 | 80 | c("DTNA", "BCL2", "CXCR4", "JUN", "PIM1", "HMOX1", "SDC3", "GLRX", "CAVIN1", "GPC1", "DCN", "ETS1", "FOXO3", "FOS", "WSB1", "PAM", "PNRC1", "PLAUR", "NEDD4L", "ZFP36", "PFKFB3", "DUSP1", "S100A4") |
| 17 | HALLMARK_TNFA_SIGNALING_VIA_NFKB | 0.045439 | 0.133645 | -0.325065 | -1.457608 | 273 | 71 | c("DUSP4", "B4GALT5", "KLF9", "PLPP3", "MAP2K3", "JUN", "FOSB", "CFLAR", "SGK1", "EHD1", "TNFAIP2", "MARCKS", "FOS", "PNRC1", "IFNGR2", "PLAUR", "ZFP36", "PFKFB3", "DUSP1", "SOD2", "BTG2") |
| 18 | HALLMARK_FATTY_ACID_METABOLISM | 0.060831 | 0.168976 | 0.314941 | 1.410761 | 261 | 45 | c("TDO2", "PRDX6", "MIF", "FH", "S100A10", "UGDH", "EPHX1", "GLUL", "RAP1GDS1", "ELOVL5", "ECI2", "HSD17B10", "ALDH3A2", "HSP90AA1", "ECI1", "NCAPH2", "SUCLG2", "ADSL", "MDH2", "PCBD1", "HADH", "PSME1", "ALDH9A1", "ADIPOR2", "BLVRA", "GPD2", "APEX1", "ENO2", "FASN", "SLC22A5", "RETSAT", "GRHPR") |
| 19 | HALLMARK_REACTIVE_OXYGEN_SPECIES_PATHWAY | 0.065087 | 0.17128 | -0.415723 | -1.496521 | 345 | 27 | c("MPO", "GPX3", "LSP1", "GLRX", "CAT", "GSR", "TXNRD1", "FTL", "SOD2") |
| 20 | HALLMARK_DNA_REPAIR | 0.085335 | 0.207074 | 0.299045 | 1.356036 | 365 | 47 | c("BRF2", "MPC2", "NUDT21", "GTF2A2", "CANT1", "TMED2", "IMPDH2", "DAD1", "NME1", "POLR2C", "POLB", "LIG1", "APRT", "RFC2", "GTF2F1", "BCAM", "CETN2", "NME4", "POLR3C", "ARL6IP1", "POLR2H", "COX17", "SEC61A1", "GMPR2") |
| 21 | HALLMARK_P53_PATHWAY | 0.086971 | 0.207074 | -0.310154 | -1.364549 | 517 | 64 | c("IFI30", "FUCA1", "ABCC5", "RXRA", "JUN", "CTSD", "RAP2B", "HMOX1", "STOM", "CCND3", "TXNIP", "FOXO3", "FOS", "FBXW7", "S100A4", "BTG2", "MXD4") |
| 22 | HALLMARK_APOPTOSIS | 0.095302 | 0.216597 | -0.311046 | -1.356225 | 563 | 61 | c("GPX3", "CD14", "JUN", "PLAT", "HMOX1", "CFLAR", "GSN", "DCN", "IGF2R", "TXNIP", "ANKH", "DPYD", "GSR", "SOD2", "BTG2", "RHOT2", "MMP2", "LUM", "ANXA1", "PDGFRB") |
| 23 | HALLMARK_INTERFERON_ALPHA_RESPONSE | 0.122824 | 0.267009 | -0.419938 | -1.379639 | 627 | 19 | c("IFI30", "CD74", "C1S", "IFI27", "TXNIP", "LAP3", "STAT2") |
| 24 | HALLMARK_IL2_STAT5_SIGNALING | 0.130775 | 0.272447 | -0.312845 | -1.311163 | 752 | 51 | c("BCL2", "NFKBIZ", "S100A1", "CTSZ", "PIM1", "AHR", "CCND3", "PLEC", "HIPK2", "IGF2R") |
| 25 | HALLMARK_NOTCH_SIGNALING | 0.147503 | 0.273743 | -0.538059 | -1.360394 | 693 | 8 | c("ARRB1", "LFNG", "NOTCH2") |
| 26 | HALLMARK_MYC_TARGETS_V2 | 0.147821 | 0.273743 | 0.439837 | 1.324002 | 752 | 12 | c("SORD", "MCM5", "HSPD1", "CBX3", "UNG") |
| 27 | HALLMARK_INFLAMMATORY_RESPONSE | 0.144912 | 0.273743 | -0.334671 | -1.308703 | 805 | 37 | c("CYBB", "STAB1", "CD14", "AHR", "IL1R1", "EMP3", "IFNGR2", "PLAUR", "TAPBP", "BTG2") |
| 28 | HALLMARK_UV_RESPONSE_DN | 0.159728 | 0.285229 | -0.305452 | -1.26927 | 916 | 49 | c("TGFBR2", "RXRA", "PLPP3", "DAB2", "ADD3", "AKT3", "PRDM2", "RBPMS") |
| 29 | HALLMARK_PROTEIN_SECRETION | 0.172073 | 0.296678 | 0.26759 | 1.218125 | 731 | 48 | c("TPD52", "ARF1", "TOM1L1", "TMED2", "AP2S1", "TMED10", "RAB5A", "AP1G1", "COPB2", "MAPK1", "ARFGAP3", "PPT1", "COPE", "GLA", "SCAMP3", "RAB22A", "LAMP2", "COG2", "SEC31A", "CLTC", "RAB2A", "COPB1", "YIPF6", "TSG101", "CLCN3", "USO1", "ARCN1", "SOD1", "ARFIP1") |
| 30 | HALLMARK_CHOLESTEROL_HOMEOSTASIS | 0.182465 | 0.304108 | 0.328314 | 1.240171 | 871 | 24 | c("SEMA3B", "CD9", "CLU", "ALCAM", "LDLR", "CYP51A1", "ACTG1", "PMVK", "CTNNB1") |
| 31 | HALLMARK_G2M_CHECKPOINT | 0.221489 | 0.357241 | 0.252685 | 1.170178 | 939 | 51 | c("MCM5", "UCK2", "MCM2", "BCL3", "NUSAP1", "GINS2", "HOXC10", "MCM3", "CTCF", "LMNB1", "MYBL2", "TPX2", "MT2A", "TMPO", "RAD23B", "MNAT1", "TLE3", "HNRNPU", "TOP2A", "CCND1", "KIF5B", "NCL", "SMC4") |
| 32 | HALLMARK_APICAL_JUNCTION | 0.274445 | 0.415826 | -0.2504 | -1.1304 | 1669 | 73 | c("AKT3", "SORBS3", "EPB41L2", "SDC3", "CNN2", "PECAM1", "EVL", "COL16A1", "STX4", "RAC2", "VCAN", "CDH11", "FBN1", "MMP9", "TIAL1", "MYL9", "ZYX", "MMP2", "TSPAN4", "ADAM15", "PBX2", "PARVA", "CD99", "THY1", "LIMA1", "MAPK13", "INPPL1", "AKT2", "PFN1", "CERCAM") |

| | | | | | | | | |
|---|---|---|---|---|---|---|---|---|
| 33 | HALLMARK_GLYCOLYSIS | 0.269779 | 0.415826 | 0.230224 | 1.118259 | 1097 | 62 | c("GOT2", "FUT8", "MIF", "STC2", "STC1", "GFPT1", "ALDH7A1", "HSPA5", "GMPPA", "COPB2", "HAX1", "PPIA", "AGL", "PSMC4", "KDELR3", "CD44", "BPNT1", "IL13RA1", "CITED2", "GALK2", "PGLS", "PLOD2", "VEGFA", "CHPF", "IER3", "COG2", "MDH2", "ARPP19", "ALDH9A1", "ELF3", "FAM162A", "TGFBI", "P4HA2", "CLN6", "IDUA", "ENO2", "PKM", "HDLBP") |
| 34 | HALLMARK_MITOTIC_SPINDLE | 0.305019 | 0.435742 | -0.250447 | -1.110085 | 1816 | 67 | c("SHROOM1", "CNTRL", "EPB41L2", "ABR", "TRIO", "GSN", "PCM1", "MARCKS", "ABL1", "NIN", "NOTCH2", "RAB3GAP1", "WASF2", "CTTN") |
| 35 | HALLMARK_MYOGENESIS | 0.301773 | 0.435742 | -0.26201 | -1.114024 | 1752 | 54 | c("GPX3", "DTNA", "LSP1", "CD36", "SORBS3", "APOD", "GSN", "CFD", "IGFBP7") |
| 36 | HALLMARK_UV_RESPONSE_UP | 0.317645 | 0.441174 | -0.264009 | -1.106489 | 1828 | 51 | c("GPX3", "ARRB2", "HMOX1", "UROD", "FOSB", "CCND3", "TFRC", "MGAT1", "NXF1", "FOS", "SOD2", "BTG2") |
| 37 | HALLMARK_XENOBIOTIC_METABOLISM | 0.361935 | 0.489102 | -0.244067 | -1.066595 | 2146 | 62 | c("APOE", "ALDH2", "CD36", "HMOX1", "PDK4", "CNDP2", "MAN1A1", "CAT", "FBLN1", "IL1R1", "PGD", "PSMB10", "GSR") |
| 38 | HALLMARK_ANGIOGENESIS | 0.397622 | 0.523187 | 0.315181 | 1.044276 | 1972 | 16 | c("SERPINA5", "THBD", "STC1") |
| 39 | HALLMARK_PEROXISOME | 0.471597 | 0.604612 | 0.238051 | 0.998626 | 2116 | 35 | c("CRABP2", "SLC27A2", "HSD11B2", "IDH2", "ABCD3", "ELOVL5", "ECI2", "GNPAT", "SIAH1", "TOP2A", "ALDH9A1", "PEX11B", "SLC25A4", "ISOC1", "FDPS", "CLN6", "SLC35B2", "SEMA3C", "LONP2", "SOD1", "PRDX1", "RETSAT") |
| 40 | HALLMARK_KRAS_SIGNALING_DN | 0.521632 | 0.65204 | 0.268407 | 0.957171 | 2543 | 20 | c("LYPD3", "GPRC5C", "EFHD1", "HSD11B2", "NR4A2", "SPTBN2", "GAMT", "IGFBP2") |
| 41 | HALLMARK_MTORC1_SIGNALING | 0.537388 | 0.655351 | -0.204503 | -0.953379 | 3348 | 87 | c("CORO1A", "IFI30", "ITGB2", "CXCR4", "MAP2K3", "LGMN", "ADD3", "GLRX", "TFRC") |
| 42 | HALLMARK_TGF_BETA_SIGNALING | 0.576 | 0.685714 | -0.256245 | -0.90369 | 3023 | 25 | c("LTBP2", "SKI", "HIPK2", "ENG", "IFNGR2") |
| 43 | HALLMARK_PANCREAS_BETA_CELLS | 0.618054 | 0.718668 | -0.358352 | -0.867394 | 2902 | 7 | c("AKT3", "MAFB") |
| 44 | HALLMARK_SPERMATOGENESIS | 0.645124 | 0.733096 | 0.319884 | 0.866699 | 3353 | 9 | c("GSTM3", "VDAC3", "CSNK2A2", "MAP7", "ZC3H14", "HSPA2") |
| 45 | HALLMARK_WNT_BETA_CATENIN_SIGNALING | 0.666179 | 0.740199 | 0.426305 | 0.868566 | 3645 | 4 | c("CTNNB1", "CSNK1E", "NCSTN", "NCOR2") |
| 46 | HALLMARK_PI3K_AKT_MTOR_SIGNALING | 0.724674 | 0.787689 | -0.205339 | -0.817874 | 4055 | 41 | c("ITPR2", "CXCR4", "MAP2K3", "GRK2", "MYD88", "SMAD2", "STAT2") |
| 47 | HALLMARK_ANDROGEN_RESPONSE | 0.778207 | 0.811206 | 0.184056 | 0.803975 | 3427 | 41 | c("SORD", "SPDEF", "TPD52", "KRT8", "UAP1", "TSC22D1", "KRT19", "XRCC6", "ACSL3", "ELOVL5", "FKBP5") |
| 48 | HALLMARK_BILE_ACID_METABOLISM | 0.778757 | 0.811206 | 0.216271 | 0.771249 | 3797 | 20 | c("SLC27A2", "IDH2", "PHYH", "ABCD3", "GNPAT", "GCLM", "ALDH9A1", "PEX19", "ISOC1", "SLC35B2", "LONP2", "SOD1", "RETSAT") |
| 49 | HALLMARK_APICAL_SURFACE | 0.81331 | 0.81331 | 0.24243 | 0.729767 | 4142 | 12 | c("LYPD3", "GSTM3", "SULF2", "HSPB1", "ADIPOR2", "ATP8B1", "CRYBG1", "B4GALT1") |
| 50 | HALLMARK_HEDGEHOG_SIGNALING | 0.800658 | 0.81331 | -0.25713 | -0.720964 | 3891 | 11 | c("DPYSL2", "LDB1", "NRP1", "THY1") |

| Cluster 11 | pathway | pval | padj | ES | NES | nMoreExtreme | size | leadingEdge |
|---|---|---|---|---|---|---|---|---|
| 1 | HALLMARK_ESTROGEN_RESPONSE_EARLY | 0.0001 | 0.002502 | -0.545771 | -2.112961 | 0 | 73 | c("RET", "PGR", "ADCY1", "SLC2A1", "SLC39A6", "SCNN1A", "TFF3", "ABHD2", "SLC9A3R1", "TTC39A", "CLDN7", "SIAH2", "MAPT", "HSPB8", "GAB2", "SLC7A2", "MYB", "GREB1", "CANT1", "SEC14L2", "TFF1", "GJA1", "FLNB", "RHOD", "CA12", "RHOBTB3", "RAB31", "IGF1R", "PRSS23", "ELF3", "PAPSS2", "MAST4", "THSD4", "GFRA1", "TOB1", "SEMA3B", "MINDY1", "CELSR1") |
| 2 | HALLMARK_ESTROGEN_RESPONSE_LATE | 0.0001 | 0.002502 | -0.618131 | -2.381405 | 0 | 70 | c("LTF", "RET", "PGR", "S100A9", "SERPINA1", "CDH1", "TOP2A", "PRLR", "SCNN1A", "LLGL2", "TFF3", "SERPINA3", "ABHD2", "SLC9A3R1", "SIAH2", "MAPT", "UGDH", "HSPB8", "COX6C", "AGR2", "MYB", "DCXR", "DLG5", "SCUBE2", "TFF1", "LSR", "TSPAN13", "SGK1", "FLNB", "MAPK13", "CA12", "RABEP1", "RAB31", "IDH2", "PRSS23", "PAPSS2", "TOB1", "SEMA3B") |
| 3 | HALLMARK_COMPLEMENT | 0.00161 | 0.026841 | -0.494899 | -1.79019 | 15 | 42 | c("LTF", "APOC1", "S100A9", "SERPINA1", "PLAT", "PLAUR", "FCER1G", "CTSD", "HSPA1A", "CTSS", "LIPA", "PIM1", "CFB", "CTSL", "CD55", "GRB2", "USP15", "PPP4C", "LTA4H", "ATOX1") |
| 4 | HALLMARK_KRAS_SIGNALING_UP | 0.002816 | 0.035204 | -0.478574 | -1.727342 | 27 | 41 | c("MMP9", "SERPINA3", "SPP1", "PLAT", "IKZF1", "MMP11", "PLAUR", "ITGB2", "FCER1G", "GPNMB", "PLAU", "JUP", "CTSS", "TSPAN13", "CFB", "SEMA3B", "IGFBP3", "LCP1", "LAPTM5", "MAP3K1", "INHBA", "ANKH", "AKT2") |
| 5 | HALLMARK_COAGULATION | 0.005886 | 0.058859 | -0.492622 | -1.708607 | 57 | 32 | c("MMP9", "APOC1", "SERPINA1", "PLAT", "S100A1", "MMP11", "PLAU", "COMP", "CFB") |
| 6 | HALLMARK_HEME_METABOLISM | 0.013514 | 0.112613 | -0.404598 | -1.527277 | 134 | 59 | c("HBB", "SLC4A1", "HBD", "SNCA", "SLC2A1", "SLC25A37", "ACP5", "MPP1", "CIR1", "ELL2", "TFRC", "HDGF", "UBAC1", "FBXO7", "HAGH", "EIF2AK1", "BSG", "MAP2K3", "USP15", "NCOA4", "TRAK2", "GLRX5", "FBXO9", "RAD23A", "VEZF1") |
| 7 | HALLMARK_ALLOGRAFT_REJECTION | 0.02978 | 0.212716 | -0.465303 | -1.556345 | 289 | 26 | c("MMP9", "CAPG", "TPD52", "NME1", "ITGB2", "CTSS", "SPI1", "HCLS1", "PSMB10", "SRGN", "BCL3", "INHBA") |
| 8 | HALLMARK_XENOBIOTIC_METABOLISM | 0.06035 | 0.335278 | -0.392649 | -1.41721 | 599 | 41 | c("HMOX1", "UGDH", "ESR1", "DCXR", "JUP", "SLC1A5", "CFB", "NQO1", "APOE", "MCCC2", "PAPSS2", "FBP1", "PGD", "SPINT2", "PSMB10", "CNDP2") |
| 9 | HALLMARK_KRAS_SIGNALING_DN | 0.053887 | 0.335278 | -0.547145 | -1.512144 | 478 | 11 | c("BMPR1B", "GPRC5C", "SGK1", "EFHD1", "IDUA") |
| 10 | HALLMARK_PANCREAS_BETA_CELLS | 0.074953 | 0.374765 | -0.624121 | -1.454033 | 596 | 6 | SYT13 |
| 11 | HALLMARK_MTORC1_SIGNALING | 0.119193 | 0.530371 | -0.349694 | -1.295829 | 1187 | 50 | c("SLC2A1", "EGLN3", "SLC9A3R1", "IFI30", "ITGB2", "CANX", "CORO1A", "TFRC", "SLC1A5", "GGA2", "SYTL2", "STC1", "ACSL3", "MAP2K3", "ENO1", "CD9", "ACACA", "PPIA", "LTA4H", "SERP1", "CXCR4", "M6PR", "TXNRD1", "GSK3B", "GSR", "NUPR1", "CCT6A", "PSMD13", "P4HA1", "USO1", "YKT6", "BHLHE40", "ACTR3") |
| 12 | HALLMARK_EPITHELIAL_MESENCHYMAL_TRANSITION | 0.127289 | 0.530371 | -0.318067 | -1.242358 | 1271 | 80 | c("COL11A1", "CAPG", "TNC", "SPP1", "SDC1", "LOXL2", "LRRC15", "PLAUR", "TGFBI", "GJA1", "LOXL1", "VEGFA", "COMP", "ADAM12", "IGFBP3", "QSOX1", "MCM7", "ECM1", "COL12A1", "MXRA5", "CDH11", "TNFRSF12A") |
| 13 | HALLMARK_ANDROGEN_RESPONSE | 0.151887 | 0.564833 | -0.353714 | -1.272463 | 1508 | 40 | c("BMPR1B", "TPD52", "ABHD2", "SPDEF", "ELL2", "SGK1", "SRP19", "SMS", "INPP4B", "ACSL3", "STEAP4", "NCOA4", "AZGP1", "ANKH", "UAP1", "LMAN1", "ACTN1", "UBE2I") |
| 14 | HALLMARK_HEDGEHOG_SIGNALING | 0.178729 | 0.564833 | -0.527954 | -1.288431 | 1461 | 7 | c("VEGFA", "ADGRG1", "CELSR1", "LDB1", "THY1") |
| 15 | HALLMARK_BILE_ACID_METABOLISM | 0.180747 | 0.564833 | 0.300689 | 1.238408 | 183 | 12 | c("ALDH9A1", "CAT", "RXRA", "SCP2", "OPTN", "GSTK1", "SLC29A1", "LONP2", "DHCR24", "SOD1", "HSD17B4") |
| 16 | HALLMARK_SPERMATOGENESIS | 0.168989 | 0.564833 | -0.558221 | -1.300503 | 1345 | 6 | c("SLC12A2", "STRBP", "TALDO1", "VDAC3", "GSTM3") |
| 17 | HALLMARK_GLYCOLYSIS | 0.206914 | 0.60857 | -0.330926 | -1.208644 | 2058 | 45 | c("EGLN3", "TFF3", "SDC1", "TGFBI", "VEGFA", "FUT8", "MIF", "ELF3", "IDUA", "STC1", "IGFBP3", "QSOX1", "COPB2", "ENO1", "PPIA", "RBCK1", "TALDO1", "CXCR4", "PKM", "SDC2", "PLOD1", "IER3") |
| 18 | HALLMARK_E2F_TARGETS | 0.250412 | 0.69559 | -0.361183 | -1.197904 | 2428 | 25 | c("TOP2A", "HMGB2", "SMC4", "NME1", "ANP32E", "TFRC", "RBBP7", "SRSF2", "MCM7") |
| 19 | HALLMARK_ADIPOGENESIS | 0.285714 | 0.75188 | 0.144772 | 1.13473 | 11 | 47 | c("ADIPOQ", "FABP4", "GPX3", "LPL", "CD36", "ALDH2", "C3", "ETFB", "DHRS7", "YWHAG") |
| 20 | HALLMARK_TNFA_SIGNALING_VIA_NFKB | 0.388889 | 0.769395 | 0.133489 | 1.182037 | 6 | 58 | c("G0S2", "FOS", "FOSB", "SOCS3", "ZFP36", "DUSP1", "NR4A1", "RHOB", "EGR1", "MCL1", "KLF2", "KLF6", "GEM", "CEBPB") |
| 21 | HALLMARK_HYPOXIA | 0.466867 | 0.769395 | -0.269138 | -1.019135 | 4663 | 61 | c("SLC2A1", "TPD52", "HMOX1", "SIAH2", "DTNA", "PLAUR", "TGFBI", "PIM1", "VEGFA", "CA12", "MIF", "STC1", "IGFBP3", "FBP1", "ENO1", "EFNA1", "CCNG2") |
| 22 | HALLMARK_WNT_BETA_CATENIN_SIGNALING | 0.350542 | 0.769395 | 0.398034 | 1.058946 | 807 | 5 | c("NCSTN", "CSNK1E", "NCOR2", "MYC", "CTNNB1") |
| 23 | HALLMARK_IL6_JAK_STAT3_SIGNALING | 0.452484 | 0.769395 | -0.32565 | -1.02392 | 4289 | 19 | c("HMOX1", "PIM1", "PTPN1", "MAP3K8", "CD9", "TNFRSF12A", "GRB2", "MYD88") |

| | | | | | | | | |
|---|---|---|---|---|---|---|---|---|
| 24 | HALLMARK_DNA_REPAIR | 0.348285 | 0.769395 | -0.318432 | -1.104446 | 3431 | 32 | c("NME1", "BRF2", "CANT1", "HCLS1", "POLD4", "NME4", "NELFCD", "IMPDH2", "NUDT21", "APRT", "POLR2I", "BCAM", "POLR1D", "DCTN4", "ERCC1", "POLR2H", "SEC61A1", "RBX1", "TMED2", "GUK1", "VPS28", "CSTF3", "NME3", "EIF1B", "SUPT5H") |
| 25 | HALLMARK_NOTCH_SIGNALING | 0.467412 | 0.769395 | 0.289862 | 0.973271 | 752 | 8 | c("NOTCH2", "CCND1", "ARRB1", "APH1A", "PSENEN", "RBX1", "LFNG", "HES1") |
| 26 | HALLMARK_APICAL_JUNCTION | 0.366108 | 0.769395 | -0.293756 | -1.08533 | 3648 | 49 | c("MMP9", "CDH1", "CLDN7", "CLDN4", "RAC2", "TGFBI", "JUP", "MAPK13", "INPPL1", "CDH11", "AKT2", "NECTIN2", "ACTG1", "THY1", "SYMPK", "ACTN1", "VASP", "THBS3", "MDK") |
| 27 | HALLMARK_APICAL_SURFACE | 0.483863 | 0.769395 | -0.406511 | -0.992058 | 3957 | 7 | c("PLAUR", "SULF2", "THY1", "GSTM3", "FLOT2") |
| 28 | HALLMARK_UNFOLDED_PROTEIN_RESPONSE | 0.47808 | 0.769395 | -0.290719 | -1.008329 | 4710 | 32 | c("EIF4EBP1", "LSM1", "DNAJA4", "VEGFA", "EIF4A3", "SRPRA", "NHP2", "SPCS1", "EIF4G1", "SERP1", "ATP6V0D1", "KIF5B", "FUS", "GOSR2", "EIF4A2", "BANF1", "NOP56", "LSM4", "HERPUD1", "YWHAZ", "SEC31A", "CALR", "EDEM1") |
| 29 | HALLMARK_MYC_TARGETS_V1 | 0.42761 | 0.769395 | -0.284918 | -1.04685 | 4258 | 47 | c("CLNS1A", "NME1", "CANX", "HSP90AB1", "HDGF", "HNRNPC", "SMARCC1", "SNRPD2", "SRSF2", "MCM7", "PPIA", "BUB3", "PSMA7", "NHP2", "IMPDH2", "POLD2", "VDAC3", "CBX3", "GNL3", "LSM7", "ILF2", "CYC1", "SF3B3", "EIF3B", "NOP56", "MYC", "EIF4H", "TXNL4A", "PSMD3", "YWHAE", "RNPS1", "CCT3", "DHX15", "EEF1B2") |
| 30 | HALLMARK_FATTY_ACID_METABOLISM | 0.42268 | 0.769395 | -0.315938 | -1.047846 | 4099 | 25 | c("BMPR1B", "UGDH", "FH", "ECI1", "MIF", "SMS") |
| 31 | HALLMARK_ANGIOGENESIS | 0.341607 | 0.769395 | -0.396598 | -1.120212 | 3068 | 12 | c("SPP1", "VEGFA", "LRPAP1", "STC1") |
| 32 | HALLMARK_IL2_STAT5_SIGNALING | 0.492413 | 0.769395 | -0.274848 | -1.003829 | 4899 | 45 | c("CAPG", "SPP1", "S100A1", "PHLDA1", "ALCAM", "PIM1", "SLC1A5", "ENPP1", "IGF1R", "MAP3K8", "ECM1", "CKAP4", "ITGA6") |
| 33 | HALLMARK_P53_PATHWAY | 0.521074 | 0.789507 | -0.269601 | -0.979177 | 5179 | 43 | c("HMOX1", "IFI30", "SDC1", "DCXR", "CTSD", "HEXIM1", "IP6K2", "EPS8L2", "ABCC5", "TOB1") |
| 34 | HALLMARK_PROTEIN_SECRETION | 0.541625 | 0.792087 | -0.272798 | -0.964166 | 5360 | 36 | c("TPD52", "ANP32E", "TOM1L1", "ARFGEF2", "COPB2", "STX16", "VPS45", "IGF2R", "ARF1", "LMAN1", "M6PR", "MAPK1", "COPB1", "GOSR2", "SNX2", "TMED2", "USO1", "YKT6", "PPT1", "RAB2A", "COPE", "ARCN1", "LAMP2", "RER1", "SOD1", "SEC31A") |
| 35 | HALLMARK_PEROXISOME | 0.554461 | 0.792087 | -0.298297 | -0.947957 | 5288 | 20 | c("TOP2A", "CRABP2", "IDH2", "ABCC5", "SMARCC1") |
| 36 | HALLMARK_PI3K_AKT_MTOR_SIGNALING | 0.611134 | 0.848797 | -0.264975 | -0.910336 | 6004 | 30 | c("SLC2A1", "VAV3", "MAP2K3", "ARPC3", "ACACA", "GRB2", "PPP1CA", "ITPR2", "MYD88", "CXCR4", "PLA2G12A", "ARF1", "MAPK1", "GSK3B", "PAK4", "GRK2", "SMAD2", "ACTR3") |
| 37 | HALLMARK_MYC_TARGETS_V2 | 0.649835 | 0.855047 | 0.272504 | 0.855295 | 1183 | 7 | c("HSPE1", "PA2G4", "SRM", "MYC", "NOP56", "GNL3", "CBX3") |
| 38 | HALLMARK_REACTIVE_OXYGEN_SPECIES_PATHWAY | 0.645068 | 0.855047 | -0.286581 | -0.869683 | 6022 | 16 | c("MPO", "NQO1", "ATOX1", "TXNRD1", "GSR", "NDUFS2", "SBNO2", "LSP1", "PTPA", "PRDX2", "EGLN2", "SOD1") |
| 39 | HALLMARK_CHOLESTEROL_HOMEOSTASIS | 0.687214 | 0.859018 | -0.284105 | -0.83509 | 6309 | 14 | c("PLAUR", "ALCAM", "SEMA3B", "CD9", "TNFRSF12A", "ACTG1") |
| 40 | HALLMARK_G2M_CHECKPOINT | 0.670591 | 0.859018 | -0.250119 | -0.86751 | 6607 | 32 | c("TOP2A", "SMC4", "SLC12A2", "SLC38A1", "SMARCC1", "SRSF2", "BUB3", "BCL3", "TOP1", "MARCKS", "RBM14", "CUL4A", "KIF5B", "YTHDC1", "PRPF4B", "MYC", "ILF3", "SS18", "CDKN1B", "HSPA8", "CUL3", "HNRNPU", "MT2A", "HNRNPD", "ATRX") |
| 41 | HALLMARK_MYOGENESIS | 0.739477 | 0.883575 | -0.23082 | -0.82284 | 7325 | 38 | c("TNNT1", "SPDEF", "DTNA", "HSPB8", "NQO1", "PVALB", "ADAM12", "IGFBP3", "ERBB3", "SYNGR2", "AKT2", "BAG1", "PDLIM7") |
| 42 | HALLMARK_INTERFERON_GAMMA_RESPONSE | 0.746456 | 0.883575 | -0.232219 | -0.813918 | 7371 | 34 | c("IFI30", "PIM1", "CFB", "PTPN1", "PSMB10", "NFKBIA", "RBCK1", "IFI27", "SRI", "MYD88", "IFITM2", "RNF213") |
| 43 | HALLMARK_UV_RESPONSE_UP | 0.759875 | 0.883575 | -0.226956 | -0.804948 | 7521 | 37 | c("RET", "HMOX1", "EPCAM") |
| 44 | HALLMARK_MITOTIC_SPINDLE | 0.849123 | 0.94347 | -0.200166 | -0.750622 | 8469 | 56 | c("TOP2A", "SMC4", "PREX1", "EZR", "ARFIP2", "FLNB", "ARF6", "NEDD9", "ARL8A", "ARHGAP4", "SEPTIN9", "CNTRL", "NET1", "ABR", "RHOT2", "MARCKS", "CD2AP", "PCM1", "STAU1", "KIF5B", "CKAP5", "DYNLL2") |
| 45 | HALLMARK_APOPTOSIS | 0.844788 | 0.94347 | -0.201295 | -0.743717 | 8419 | 49 | c("TOP2A", "HMGB2", "HMOX1", "PLAT", "ERBB2", "SLC20A1", "NEDD9", "ERBB3", "TNFRSF12A", "RARA", "DAP3", "ANKH", "RHOT2") |
| 46 | HALLMARK_INFLAMMATORY_RESPONSE | 0.877216 | 0.953496 | -0.19851 | -0.658383 | 8508 | 25 | c("PLAUR", "SLC7A2", "CYBB", "CD55", "NFKBIA", "SRI", "INHBA") |
| 47 | HALLMARK_TGF_BETA_SIGNALING | 0.977374 | 0.977374 | -0.145252 | -0.478495 | 9459 | 24 | c("CDH1", "SLC20A1", "RAB31", "PPM1A", "PPP1CA", "SKIL", "CTNNB1", "FNTA") |
| 48 | HALLMARK_INTERFERON_ALPHA_RESPONSE | 0.976052 | 0.977374 | -0.160382 | -0.480118 | 9047 | 15 | c("IFI30", "ELF1", "IFI27", "IFITM2") |

| | | | | | | | | |
|---|---|---|---|---|---|---|---|---|
| 49 | HALLMARK_OXIDATIVE_PHOSPHORYLATION | 0.948749 | 0.977374 | -0.164824 | -0.630361 | 9477 | 66 | c("COX6C", "NDUFC2", "MAOB", "TCIRG1", "FH", "ECI1", "ATP1B1", "IDH2", "COX6A1", "ATP5PD", "ATP6V0C", "NDUFV1", "UQCRQ", "ATP6V0B", "NDUFB7", "VDAC3", "RHOT2", "ATP5MF", "NDUFC1", "UQCR10", "NDUFA2", "NDUFS2", "NDUFS7", "ATP5F1B", "ATP5MG", "CYC1", "UQCRC1", "ATP6V0E1", "ATP6AP1", "ATP5F1E", "ACAA1", "GLUD1", "NDUFB2", "NDUFS4", "LRPPRC") |
| 50 | HALLMARK_UV_RESPONSE_DN | 0.965451 | 0.977374 | -0.149717 | -0.531004 | 9556 | 37 | c("COL11A1", "GJA1", "ERBB2", "IGF1R", "INPP4B") |

| Cluster 12 | pathway | pval | padj | ES | NES | nMoreExtreme | size | leadingEdge |
|---|---|---|---|---|---|---|---|---|
| 1 | HALLMARK_P53_PATHWAY | 0.00011 | 0.005505 | 0.40918 | 1.865987 | 0 | 86 | c("FOS", "ZBTB16", "JAG2", "ZMAT3", "F2R", "INHBB", "CDKN1A", "DDB2", "GADD45A", "PLK3", "TSPYL2", "AEN", "PITPNC1", "TXNIP", "PHLDA3", "PTPN14", "POLH", "PERP", "PPP1R15A", "FAS", "KLF4", "XPC", "SESN1", "SLC35D1", "NDRG1", "RETSAT", "BTG2", "VWA5A", "JUN", "KIF13B", "FBXW7", "CTSF", "BLCAP", "CSRNP2", "TM7SF3", "ELP1", "CCND2", "ITGB4", "ISCU", "MXD4", "SLC19A2", "KRT17", "CD82", "CCNG1") |
| 2 | HALLMARK_TNFA_SIGNALING_VIA_NFKB | 0.000773 | 0.011891 | 0.386653 | 1.75278 | 6 | 83 | c("NR4A1", "FOS", "NR4A3", "FOSB", "EGR1", "IER2", "CDKN1A", "DUSP1", "MAFF", "SDC4", "GADD45A", "JUNB", "DUSP4", "PER1", "KLF9", "JAG1", "NR4A2", "TRIP10", "PPP1R15A", "KLF4", "GEM", "BTG2", "JUN", "ZFP36", "KLF2", "SOCS3", "GADD45B", "NFKB2", "ZC3H12A", "NFKB1", "PNRC1", "IRS2", "KLF10", "NINJ1", "TGIF1", "ETS2", "PDLIM5", "TRIB1", "IER5", "CEBPD") |
| 3 | HALLMARK_ESTROGEN_RESPONSE_EARLY | 0.001189 | 0.011891 | -0.350178 | -2.165748 | 0 | 92 | c("PGR", "ADCY1", "RET", "SLC39A6", "RHOBTB3", "SLC9A3R1", "HSPB8", "FASN", "SLC2A1", "GAB2", "PRSS23", "MAPT", "SLC7A2", "TFF3", "GJA1", "PAPSS2", "CANT1", "KRT8", "BLVRB", "CCND1", "CLDN7", "MAST4", "KRT19", "SCNN1A", "TOB1", "RARA", "SIAH2", "RAB31", "AKAP1", "BAG1") |
| 4 | HALLMARK_ESTROGEN_RESPONSE_LATE | 0.00099 | 0.011891 | -0.424882 | -2.484185 | 0 | 75 | c("S100A9", "TOP2A", "SERPINA1", "PGR", "RET", "CDH1", "COX6C", "SLC9A3R1", "HSPB8", "PRSS23", "MAPT", "PRLR", "LTF", "TFF3", "UGDH", "PAPSS2", "LLGL2", "BLVRB", "CCND1", "DCXR", "KRT19", "SCNN1A", "TOB1", "SIAH2", "IDH2", "RAB31", "BAG1") |
| 5 | HALLMARK_HEME_METABOLISM | 0.00092 | 0.011891 | -0.414063 | -2.342938 | 0 | 65 | c("HBB", "HBD", "SLC4A1", "SNCA", "ACP5", "CTSB", "SLC2A1", "TFRC", "MPP1", "SLC25A37", "BLVRB") |
| 6 | HALLMARK_COMPLEMENT | 0.005892 | 0.049102 | -0.316283 | -1.711535 | 6 | 55 | c("S100A9", "APOC1", "SERPINA1", "CTSB", "CTSD", "LTF", "CTSS", "PLAUR", "FN1", "CD55", "FCER1G") |
| 7 | HALLMARK_COAGULATION | 0.017733 | 0.126666 | -0.313048 | -1.64167 | 22 | 47 | c("MMP9", "APOC1", "SERPINA1", "MMP11", "S100A1", "CTSB", "PRSS23", "FN1", "PLAU") |
| 8 | HALLMARK_MTORC1_SIGNALING | 0.021176 | 0.132353 | -0.300855 | -1.590193 | 26 | 48 | c("SCD", "IFI30", "SLC9A3R1", "SLC2A1", "TFRC", "CORO1A", "EGLN3", "NUPR1", "ITGB2", "ENO1", "SQSTM1", "ACACA") |
| 9 | HALLMARK_MYOGENESIS | 0.029821 | 0.16567 | 0.33873 | 1.481896 | 265 | 67 | c("FXYD1", "SOD3", "LAMA2", "IGF1", "APOD", "MYH11", "GSN", "ABLIM1", "SSPN", "EFS", "CDKN1A", "AGRN", "COX7A1", "SVIL", "PTP4A3", "PTGIS", "COL15A1", "GPX3", "IGFBP7", "GADD45B", "PICK1", "PRNP", "ITGB4", "CRYAB") |
| 10 | HALLMARK_E2F_TARGETS | 0.038437 | 0.192184 | -0.325261 | -1.547702 | 59 | 33 | c("TOP2A", "HMGB2", "TFRC", "NME1", "ANP32E", "SMC4") |
| 11 | HALLMARK_UV_RESPONSE_DN | 0.061114 | 0.277792 | 0.305338 | 1.382336 | 553 | 82 | c("FBLN5", "TGFBR3", "KIT", "ACVR2A", "ID1", "ADGRL2", "DUSP1", "CAV1", "CAP2", "PTPRM", "NR1D2", "LPAR1", "PLCB4", "PDGFRB", "ADD3", "DLC1", "KCNMA1", "MAP1B", "TGFBR2", "TJP1", "TOGARAM1", "PIAS3", "RBPMS", "NFKB1", "COL3A1", "MTA1", "ATXN1", "PRDM2", "TFPI", "PDLIM5", "ATRN", "NOTCH2", "NRP1", "CCN1", "APBB2", "NFIB", "MIOS", "PIK3R3", "LAMC1", "MGMT", "NR3C1", "CDKN1B", "YTHDC1", "DBP", "PTEN", "ICA1", "COL1A2", "PMP22") |
| 12 | HALLMARK_BILE_ACID_METABOLISM | 0.103578 | 0.431575 | 0.392768 | 1.367668 | 853 | 23 | c("ABCA6", "ALDH1A1", "ABCA8", "EPHX2", "RETSAT", "HSD17B11", "PFKM", "DHCR24", "PRDX5", "ATXN1", "AR", "ISOC1", "SLC23A2", "SCP2", "ABCD3", "ALDH9A1", "ABCA2") |
| 13 | HALLMARK_EPITHELIAL_MESENCHYMAL_TRANSITIO N | 0.121457 | 0.433775 | 0.269431 | 1.269871 | 1126 | 107 | c("FBLN5", "LAMA2", "ELN", "LAMA3", "FBLN2", "SFRP1", "FBLN1", "ABI3BP", "TGFBR3", "SLIT3", "DCN", "SLIT2", "SDC4", "GADD45A", "SFRP4", "CAP2", "GPC1", "IGFBP4", "SERPINE2", "FSTL3", "FAS", "COL16A1", "PDGFRB", "DST", "GEM", "JUN", "MMP2", "ECM2") |
| 14 | HALLMARK_KRAS_SIGNALING_DN | 0.118966 | 0.433775 | 0.400054 | 1.344769 | 970 | 20 | c("ZBTB16", "KRT5", "TCF7L1", "TFAP2B", "NR4A2", "TENT5C", "BTG2", "IGFBP2", "SYNPO", "SELENOP") |
| 15 | HALLMARK_ALLOGRAFT_REJECTION | 0.130828 | 0.436094 | -0.261828 | -1.311763 | 187 | 39 | c("MMP9", "CAPG", "NME1", "CTSS", "SPI1", "ITGB2", "TPD52") |
| 16 | HALLMARK_HYPOXIA | 0.143174 | 0.447419 | 0.280266 | 1.256965 | 1290 | 78 | c("FOS", "EGFR", "DCN", "AKAP12", "SRPX", "CDKN1A", "DUSP1", "MAFF", "SDC4", "CAV1", "GPC1", "CAVIN3", "TPST2", "PDK1", "PPP1R15A", "TGFB3", "NDRG1", "JUN", "ZFP36", "PAM", "PDGFB", "GRHPR", "PNRC1", "STC1", "IRS2", "CAVIN1", "KLHL24", "WSB1", "PRDX5", "IDS", "NDST1") |
| 17 | HALLMARK_REACTIVE_OXYGEN_SPECIES_PATHWA Y | 0.178273 | 0.524332 | -0.363616 | -1.303239 | 383 | 14 | c("MPO", "FTL", "GPX4", "PRDX2") |

| | | | | | | | |
|---|---|---|---|---|---|---|---|
| 18 | HALLMARK_INTERFERON_GAMMA_RESPONSE | 0.190796 | 0.529989 | 0.296382 | 1.233092 | 1674 | 51 | c("SSPN", "C1S", "C1R", "CDKN1A", "CFH", "SERPING1", "JAK2", "TXNIP", "ARID5B", "FAS", "SOCS3", "TNFSF10", "CMTR1", "NFKB1", "CIITA", "SLC25A28", "AUTS2", "FGL2", "APOL6", "RIPK1", "EIF4E3", "ISOC1") |
| 19 | HALLMARK_NOTCH_SIGNALING | 0.219897 | 0.555872 | 0.393743 | 1.225869 | 1743 | 15 | c("FZD7", "FZD1", "NOTCH3", "JAG1", "TCF7L2", "ARRB1", "NOTCH2", "NOTCH1", "HES1", "APH1A") |
| 20 | HALLMARK_XENOBIOTIC_METABOLISM | 0.222349 | 0.555872 | 0.288385 | 1.199819 | 1951 | 51 | c("NDRG2", "IGF1", "FBLN1", "PTGES", "PTGDS", "ENTPD5", "CSAD", "IGFBP4", "SLC12A4", "DHRS1", "FAS", "SLC35D1", "RETSAT", "ALDH2") |
| 21 | HALLMARK_WNT_BETA_CATENIN_SIGNALING | 0.23486 | 0.559191 | 0.417561 | 1.210966 | 1814 | 12 | c("JAG2", "FZD1", "JAG1", "CCND2", "CSNK1E", "NOTCH1") |
| 22 | HALLMARK_IL2_STAT5_SIGNALING | 0.331556 | 0.753535 | -0.196104 | -1.09349 | 372 | 61 | c("SPP1", "CAPG", "PHLDA1", "S100A1", "CTSZ", "ECM1", "ENPP1") |
| 23 | HALLMARK_KRAS_SIGNALING_UP | 0.354681 | 0.771047 | 0.256446 | 1.08773 | 3128 | 56 | c("APOD", "SPRY2", "SPARCL1", "AKAP12", "ADGRA2", "CFH", "ADGRL4", "GUCY1A1", "PRDM1", "GADD45G", "PPP1R15A", "ITGBL1", "KLF4", "VWA5A", "ST6GAL1", "PECAM1", "TRIB2", "CCND2", "PLVAP", "F13A1", "ANO1", "MTMR10") |
| 24 | HALLMARK_CHOLESTEROL_HOMEOSTASIS | 0.38172 | 0.795251 | -0.240692 | -1.05403 | 638 | 26 | c("SCD", "FASN", "ACTG1", "PLAUR") |
| 25 | HALLMARK_APOPTOSIS | 0.447175 | 0.798527 | 0.230703 | 1.027208 | 4020 | 75 | c("IGFBP6", "GSN", "TGFBR3", "DCN", "F2R", "CDKN1A", "GADD45A", "CAV1", "TXNIP", "FAS", "PDGFRB", "RETSAT", "BTG2", "JUN", "MMP2", "GPX3", "TNFSF10", "CCND2", "GADD45B") |
| 26 | HALLMARK_ANDROGEN_RESPONSE | 0.445644 | 0.798527 | -0.19113 | -1.002313 | 577 | 47 | c("BMPR1B", "SCD", "KRT8", "TPD52", "CCND1", "KRT19", "SPDEF") |
| 27 | HALLMARK_HEDGEHOG_SIGNALING | 0.443856 | 0.798527 | 0.383132 | 1.024154 | 3351 | 9 | c("TLE1", "DPYSL2", "ETS2", "RASA1", "NRP1") |
| 28 | HALLMARK_ANGIOGENESIS | 0.401578 | 0.798527 | -0.289178 | -1.036446 | 864 | 14 | SPP1 |
| 29 | HALLMARK_PANCREAS_BETA_CELLS | 0.484808 | 0.835876 | 0.382765 | 0.988475 | 3637 | 8 | c("FOXO1", "STXBP1", "LMO2", "AKT3", "SEC11A", "SRPRB") |
| 30 | HALLMARK_TGF_BETA_SIGNALING | 0.517889 | 0.857945 | 0.255491 | 0.974987 | 4385 | 34 | c("ID1", "JUNB", "ID3", "PPP1R15A", "LTBP2", "TJP1", "SMURF1", "SKI", "BMPR2", "KLF10", "WWTR1", "TGIF1") |
| 31 | HALLMARK_MYC_TARGETS_V1 | 0.549085 | 0.857945 | -0.175827 | -0.946045 | 659 | 53 | c("CLNS1A", "NME1", "HSP90AB1", "CANX", "VDAC3", "PSMD3", "NOP56", "LSM7", "SET", "RNPS1", "APEX1", "CDK4", "SRSF3", "CYC1", "HNRNPA2B1", "CUL1", "SF3B3", "EIF4H", "HNRNPA3", "EIF2S1", "EEF1B2", "RACK1", "HNRNPD", "NCBP2", "HSPE1", "YWHAQ", "PA2G4", "ERH", "BUB3", "USP1", "PRPF31", "FBL", "XRCC6", "SRM", "DEK", "HNRNPR", "GNL3", "FAM120A", "HNRNPU", "POLE3", "PWP1", "EXOSC7", "PRDX4", "TRA2B", "UBE2L3", "PHB2", "PCBP1", "ETF1", "SF3A1", "EIF3D", "CCT4", "SNRPA1", "CNBP") |
| 32 | HALLMARK_SPERMATOGENESIS | 0.542639 | 0.857945 | -0.287027 | -0.914624 | 1278 | 10 | c("TALDO1", "VDAC3", "SLC12A2", "STRBP", "AGFG1", "TSN", "NF2", "PEBP1", "MAST2", "GSTM3") |
| 33 | HALLMARK_ADIPOGENESIS | 0.594473 | 0.870495 | 0.204443 | 0.93232 | 5398 | 86 | c("CYP4B1", "SPARCL1", "SSPN", "C3", "GADD45A", "FZD4", "ITIH5", "EPHX2", "REEP6", "DNAJC15", "LIFR", "QDPR", "RETSAT", "ALDH2", "SLC5A6", "SLC27A1", "COL15A1", "GPX3", "DHRS7") |
| 34 | HALLMARK_FATTY_ACID_METABOLISM | 0.616808 | 0.870495 | -0.171511 | -0.89943 | 799 | 47 | c("BMPR1B", "FASN", "UGDH", "GLUL", "HSP90AA1") |
| 35 | HALLMARK_UV_RESPONSE_UP | 0.601725 | 0.870495 | -0.178112 | -0.911657 | 836 | 42 | c("RET", "HMOX1", "TFRC", "EPCAM", "ARRB2", "SQSTM1") |
| 36 | HALLMARK_PEROXISOME | 0.626756 | 0.870495 | 0.233945 | 0.892764 | 5307 | 34 | c("ALDH1A1", "EPHX2", "DHRS3", "RETSAT", "CNBP", "HSD17B11", "MSH2", "DHCR24", "PRDX5", "DLG4", "ACAA1", "ATXN1", "ISOC1", "SLC23A2", "PEX2", "FDPS", "SCP2", "ABCD3", "ERCC1", "ECH1", "ALDH9A1") |
| 37 | HALLMARK_APICAL_SURFACE | 0.720471 | 0.97361 | 0.301667 | 0.80639 | 5440 | 9 | c("SRPX", "TMEM8B", "GAS1", "GSTM3") |
| 38 | HALLMARK_IL6_JAK_STAT3_SIGNALING | 0.755246 | 0.993744 | 0.226128 | 0.787407 | 6226 | 23 | c("LEPR", "FAS", "A2M", "JUN", "SOCS3") |
| 39 | HALLMARK_MITOTIC_SPINDLE | 0.806071 | 0.99797 | 0.175719 | 0.785783 | 7248 | 77 | c("SHROOM1", "GSN", "SORBS2", "CDC42EP1", "ARHGAP10", "TIAM1", "DST", "ABL1", "ARHGAP29", "ALMS1", "SYNPO", "CLASP1", "TUBGCP6", "PKD2", "ATG4B", "PDLIM5", "RASA1", "FARP1", "NOTCH2", "MYO1E", "NCK2", "PALLD", "NCK1", "ARHGEF7", "WASF2", "RASAL2", "SPTBN1", "RICTOR", "TUBGCP3", "SUN2", "RABGAP1", "LATS1", "SPTAN1", "KLC1", "MAP3K11", "MYH10", "CYTH2", "TLK1", "TRIO", "RHOT2", "NUMA1", "EPB41L2", "WASL", "FLNA", "ARHGEF12", "ACTN4", "KIF3B", "KIF1B", "KIFAP3", "TUBGCP2") |
| 40 | HALLMARK_DNA_REPAIR | 0.967585 | 0.99797 | -0.108582 | -0.574377 | 1193 | 50 | c("NME1", "CANT1", "BRF2", "NME4", "GPX4", "POLD4", "GTF2F1", "VPS28", "BCAP31", "POLR2I", "ITPA", "NUDT21", "ADRM1", "POLR2G", "STX3", "POLR2A", "CSTF3", "DGUOK", "DDB1", "SMAD5", "SUPT5H", "EIF1B", "POLR1D", "POLR2E", "NCBP2", "POLR2K", "DAD1", "AK1", "RBX1", "SEC61A1", "NME3", "BCAM") |
| 41 | HALLMARK_G2M_CHECKPOINT | 0.852187 | 0.99797 | -0.138178 | -0.718435 | 1129 | 45 | c("TOP2A", "SMC4", "CCND1", "HSPA8", "KIF5B") |
| 42 | HALLMARK_PROTEIN_SECRETION | 0.803701 | 0.99797 | 0.195435 | 0.763553 | 6861 | 38 | c("EGFR", "SSPN", "CAV2", "ATP1A1", "GALC", "DST") |

| | | | | | | | | |
|---|---|---|---|---|---|---|---|---|
| 43 | HALLMARK_INTERFERON_ALPHA_RESPONSE | 0.99797 | 0.99797 | 0.104387 | 0.389308 | 8356 | 31 | c("C1S", "TXNIP", "LPAR6", "TENT5A", "CMTR1", "SLC25A28", "NCOA7", "MVB12A", "IFI44", "TRIM26", "CD47", "GBP2", "PSMB8", "CASP8", "LAP3", "CSF1", "IFI35", "IRF1", "IFITM3", "STAT2", "PSME1", "LY6E", "NUB1", "PARP14", "IRF9", "ELF1", "CD74") |
| 44 | HALLMARK_APICAL_JUNCTION | 0.913256 | 0.99797 | 0.152037 | 0.676949 | 8211 | 75 | c("LAMA3", "CD34", "EGFR", "SLIT2", "CLDN5", "LDLRAP1", "TMEM8B", "COL16A1", "LIMA1", "VWF", "MMP2", "PCDH1", "PECAM1", "PKD1") |
| 45 | HALLMARK_UNFOLDED_PROTEIN_RESPONSE | 0.989206 | 0.99797 | -0.091085 | -0.477661 | 1282 | 47 | c("LSM1", "EIF4EBP1", "DNAJA4", "KIF5B", "ATP6V0D1", "EIF4A3", "YWHAZ", "ERO1A", "SEC31A", "NOP56", "PDIA6", "CEBPB", "SPCS1", "CNOT6", "STC2", "ATF6", "WFS1", "GOSR2", "EIF4A2", "EIF2S1", "KHSRP", "BANF1", "FUS", "HYOU1", "YIF1A", "SRPRA", "SHC1", "PREB", "BAG3", "TATDN2", "ATF4", "DNAJC3", "SRPRB", "DCTN1", "SEC11A", "EEF2", "WIPI1", "SLC1A4", "EDEM1", "ATF3", "MTREX", "EXOSC10", "ALDH18A1", "HERPUD1") |
| 46 | HALLMARK_PI3K_AKT_MTOR_SIGNALING | 0.848362 | 0.99797 | 0.19003 | 0.714379 | 7121 | 32 | c("EGFR", "CDKN1A", "TIAM1", "PDK1", "PPP2R1B", "RALB", "PAK4", "RIPK1", "PIK3R3", "TNFRSF1A", "NCK1", "CDKN1B", "PTEN", "MKNK2") |
| 47 | HALLMARK_MYC_TARGETS_V2 | 0.90625 | 0.99797 | 0.238385 | 0.637231 | 6843 | 9 | c("RCL1", "PES1", "IMP4", "GNL3", "SRM", "PA2G4", "HSPE1") |
| 48 | HALLMARK_INFLAMMATORY_RESPONSE | 0.94714 | 0.99797 | 0.151999 | 0.590076 | 8080 | 37 | c("ACVR2A", "CDKN1A", "LPAR1", "BTG2", "RNF144B", "TNFSF10", "NFKB1", "CD82", "AHR", "RELA", "AXL", "SCN1B", "OSMR", "CD14", "TNFRSF1B", "HPN", "STAB1", "KIF1B", "SERPINE1", "GNAI3", "CSF1", "KLF6", "IRF1", "GABBR1", "TAPBP", "IL1R1") |
| 49 | HALLMARK_OXIDATIVE_PHOSPHORYLATION | 0.94129 | 0.99797 | -0.114009 | -0.660418 | 977 | 72 | c("COX6C", "NDUFC2", "COX6A1", "TCIRG1", "ATP5PD", "MAOB", "IDH2", "GPX4", "VDAC3", "UQCRC1", "NDUFA4", "OXA1L", "HSD17B10", "NDUFC1", "NDUFA3", "COX7A2", "NDUFS7", "ECI1", "CYC1", "COX7A2L", "ATP5MF", "DECR1", "ATP5MC2", "NDUFB5", "COX7C", "NDUFS4", "SDHA", "UQCRC2", "OPA1", "IDH3B", "ACADVL", "NDUFA1", "COX6B1", "IDH3G", "ACADSB", "SDHB", "PDK4", "ETFB", "NDUFS1", "ATP5F1D", "ECHS1", "PDHB", "VDAC2", "ATP5PF", "NDUFS6", "RHOT2", "HADHA", "ECH1", "NDUFB6", "COX8A", "NDUFA8", "COX11", "NDUFA9", "NDUFB8", "BAX", "CYB5R3", "GLUD1", "ACO2", "POLR2F", "TIMM10", "SUCLG1", "PHB2", "ETFA", "SURF1", "ACAA1", "CPT1A", "MFN2", "ISCU", "SLC25A6", "PDP1", "RETSAT") |
| 50 | HALLMARK_GLYCOLYSIS | 0.966013 | 0.99797 | 0.13418 | 0.556034 | 8469 | 50 | c("EGFR", "DCN", "GLCE", "GPC1", "AGRN") |

| Cluster 13 | pathway | pval | padj | ES | NES | nMoreExtreme | size | leadingEdge |
|---|---|---|---|---|---|---|---|---|
| 1 | HALLMARK_HYPOXIA | 0.000142 | 0.001873 | 0.485553 | 2.17858 | 0 | 94 | c("CCN5", "TGFBI", "LOX", "CCN1", "DDIT4", "F3", "SERPINE1", "BGN", "TMEM45A", "ADM", "P4HA1", "COL5A1", "CCN2", "VEGFA", "P4HA2", "MT2A", "ACKR3", "NDRG1", "SDC2", "PAM", "ENO2", "IGFBP3", "CAV1", "MYH9", "KDELR3", "ENO1", "LXN", "SRPX", "PDGFB", "PFKP", "GBE1", "FOSL2", "MIF", "CAVIN3", "ANGPTL4", "ERO1A", "FAM162A", "ILVBL", "CAVIN1", "BTG1", "DUSP1", "KLF6", "ERRFI1") |
| 2 | HALLMARK_EPITHELIAL_MESENCHYMAL_TRANSITION | 0.000137 | 0.001873 | 0.718058 | 3.431147 | 0 | 136 | c("LOXL2", "COL11A1", "SPOCK1", "TGFBI", "GAS1", "THBS1", "FN1", "EDIL3", "LOX", "NID2", "CCN1", "PLOD2", "ITGA5", "COL5A3", "COL7A1", "SERPINE1", "TIMP3", "BGN", "GPX7", "PLOD1", "P3H1", "COL12A1", "NNMT", "THBS2", "COL5A2", "COMP", "COL4A1", "INHBA", "SERPINH1", "FERMT2", "THY1", "COL5A1", "PRRX1", "CCN2", "VEGFA", "ITGB1", "PCOLCE", "COL6A2", "LRRC15", "COL8A2", "CDH11", "ADAM12", "PMEPA1", "FSTL1", "POSTN", "LUM", "VCAM1", "CADM1", "PDGFRB", "COL4A2", "NT5E", "GREM1", "ENO2", "FMOD", "COL3A1", "ITGAV", "CTHRC1", "IGFBP3", "TNC", "ECM2", "SNAI2", "DPYSL3", "GJA1", "TAGLN", "TPM2", "SDC1", "BMP1", "COL1A2", "EFEMP2", "MMP14", "MFAP5", "MYLK", "FAP", "COL1A1", "COL6A3", "FBN1", "CALD1", "SPARC", "MYL9", "ACTA2", "FLNA", "HTRA1", "GEM", "CALU", "BASP1", "TGFB1", "MXRA5", "TIMP1", "MMP2", "PPIB", "TPM4") |
| 3 | HALLMARK_UV_RESPONSE_DN | 0.000148 | 0.001873 | 0.531086 | 2.09763 | 0 | 48 | c("COL11A1", "CCN1", "F3", "SERPINE1", "COL5A2", "SDC2", "PDGFRB", "TFPI", "COL3A1", "SNAI2", "GJA1", "COL1A2", "COL1A1", "CAV1", "PDLIM5", "LTBP1", "EFEMP1", "NRP1", "MAP1B", "LAMC1", "PMP22", "DUSP1", "NEK7", "TJP1", "CDC42BPA", "IGFBP5", "PTPRM", "PTEN", "LPAR1", "ATXN1") |
| 4 | HALLMARK_ANGIOGENESIS | 0.00015 | 0.001873 | 0.630991 | 2.051521 | 0 | 21 | c("COL5A2", "VEGFA", "FSTL1", "POSTN", "LUM", "COL3A1", "ITGAV", "TIMP1", "FGFR1", "NRP1", "VCAN", "SPP1", "CCND2") |
| 5 | HALLMARK_GLYCOLYSIS | 0.000295 | 0.002578 | 0.47975 | 1.996087 | 1 | 63 | c("TGFBI", "PLOD2", "DDIT4", "PLOD1", "P4HA1", "COL5A1", "VEGFA", "P4HA2", "SDC2", "NT5E", "PAM", "ENO2", "IGFBP3", "SDC1", "CHPF", "KDELR3", "ENO1", "SLC16A3", "PFKP", "TPST1", "MIF", "ANGPTL4", "ERO1A", "VCAN", "FAM162A") |
| 6 | HALLMARK_HEME_METABOLISM | 0.000309 | 0.002578 | -0.61163 | -2.603495 | 0 | 53 | c("HBB", "HBD", "SLC4A1", "SNCA", "UCP2", "BTG2", "SLC25A37", "ACP5", "TFRC", "MAP2K3", "CAT", "UROD", "UBAC1", "MPP1", "CCND3", "RNF19A", "CIR1", "HAGH", "NCOA4", "C3", "RBM5") |
| 7 | HALLMARK_MYOGENESIS | 0.000741 | 0.005296 | 0.457921 | 1.885547 | 4 | 59 | c("COL15A1", "FABP3", "FST", "AEBP1", "ITGB1", "COL6A2", "ADAM12", "COL4A2", "COL3A1", "IGFBP3", "TAGLN", "TPM2", "MYLK", "COL1A1", "COL6A3", "MYH9", "SPARC", "CNN3", "SPHK1", "TGFB1", "PDLIM7", "NQO1", "HSPB8", "WWTR1", "ITGB5", "COX7A1", "MB", "SMTN", "CDKN1A", "PRNP", "IGFBP7", "NOTCH1", "CLU", "PTP4A3") |
| 8 | HALLMARK_ESTROGEN_RESPONSE_LATE | 0.003095 | 0.017195 | -0.415196 | -1.845923 | 9 | 63 | c("CXCL14", "S100A9", "LTF", "SCUBE2", "MYB", "BCL2", "TFF1", "FOS", "TFF3", "MAPK13", "TSPAN13", "RET", "PDCD4") |
| 9 | HALLMARK_KRAS_SIGNALING_UP | 0.00281 | 0.017195 | 0.433261 | 1.759093 | 18 | 55 | c("TMEM158", "RGS16", "SPON1", "GFPT2", "INHBA", "PLAU", "PLVAP", "PRRX1", "GNG11", "MMP11", "TFPI", "IGFBP3", "TMEM176A", "NRP1", "ANGPTL4", "ERO1A", "TMEM176B", "ENG", "PRDM1", "CPE", "SPP1", "CCND2", "CFH", "EMP1", "ETS1") |
| 10 | HALLMARK_APICAL_JUNCTION | 0.003688 | 0.018439 | 0.41526 | 1.738052 | 24 | 65 | c("TGFBI", "VWF", "FSCN1", "THY1", "ITGB1", "CDH11", "VCAM1", "CERCAM", "BMP1", "PARVA", "FBN1", "MYH9", "MPZL1", "MYL9", "CD99", "MMP2", "ACTN1", "VCAN", "ACTG1", "RSU1", "MSN", "CDH1", "VCL", "NLGN2", "ACTB", "TJP1", "RRAS", "CNN2", "ADAM9", "CD34", "CTNNA1", "COL16A1", "CD276", "PTEN", "JUP") |
| 11 | HALLMARK_COAGULATION | 0.005613 | 0.025514 | 0.424855 | 1.714132 | 37 | 53 | c("THBS1", "FN1", "F3", "SERPINE1", "TIMP3", "COMP", "VWF", "PLAU", "MMP11", "C1R", "PRSS23", "BMP1", "MMP14", "SERPING1", "FBN1", "SPARC", "HTRA1", "PDGFB", "TIMP1", "MMP2", "CTSK", "WDR1", "CFH", "C1S", "LRP1", "THBD", "GNG12", "ADAM9", "ARF4", "CLU") |
| 12 | HALLMARK_INFLAMMATORY_RESPONSE | 0.006393 | 0.026638 | 0.458563 | 1.711371 | 42 | 37 | c("PDPN", "RGS16", "ITGA5", "F3", "CD55", "SERPINE1", "ADM", "INHBA", "OSMR", "MMP14", "SPHK1", "AXL", "RGS1", "TIMP1", "KLF6", "IL1R1", "PCDH7", "CDKN1A") |
| 13 | HALLMARK_TGF_BETA_SIGNALING | 0.041885 | 0.153404 | 0.439869 | 1.518504 | 279 | 27 | c("THBS1", "SERPINE1", "PMEPA1", "TGFB1", "ID3", "ENG", "SMAD1", "WWTR1", "CDH1", "CTNNB1", "TJP1", "TGIF1", "FNTA", "NCOR2", "RAB31") |
| 14 | HALLMARK_HEDGEHOG_SIGNALING | 0.042953 | 0.153404 | 0.562881 | 1.52752 | 287 | 11 | c("THY1", "VEGFA", "NRP2", "MYH9", "NRP1", "ADGRG1") |
| 15 | HALLMARK_UV_RESPONSE_UP | 0.068237 | 0.227456 | -0.365119 | -1.456272 | 225 | 39 | c("FOSB", "BTG2", "FOS", "TFRC", "PRKCD", "UROD", "CCND3", "ARRB2", "RET", "ACAA1") |

| | | | | | | | | |
|---|---|---|---|---|---|---|---|---|
| 16 | HALLMARK_E2F_TARGETS | 0.090579 | 0.283058 | -0.416235 | -1.443828 | 298 | 23 | c("TFRC", "ANP32E", "MCM7", "RBBP7", "HMGB2", "PA2G4", "LUC7L3", "SRSF2", "TOP2A", "PNN", "PRKDC") |
| 17 | HALLMARK_REACTIVE_OXYGEN_SPECIES_PATHWAY | 0.109427 | 0.321844 | -0.464929 | -1.435479 | 360 | 15 | c("MPO", "CAT", "GPX3", "LSP1", "TXNRD1", "GSR", "PRDX2") |
| 18 | HALLMARK_APOPTOSIS | 0.121861 | 0.338503 | 0.322099 | 1.299551 | 824 | 53 | c("TIMP3", "BGN", "LUM", "PDGFRB", "ENO2", "IFITM3", "CAV1", "PAK1", "NEDD9", "TIMP1", "MMP2", "TIMP2", "CCND2", "DAP", "EMP1", "CTNNB1", "ANKH", "CDKN1A", "GADD45A", "DCN", "CLU", "CD44", "DDIT3", "RELA", "BNIP3L", "PEA15", "ANXA1", "LMNA", "GADD45B") |
| 19 | HALLMARK_APICAL_SURFACE | 0.14079 | 0.351976 | 0.489185 | 1.327526 | 943 | 11 | c("GAS1", "THY1", "SRPX", "SULF2", "LYPD3", "PLAUR") |
| 20 | HALLMARK_KRAS_SIGNALING_DN | 0.137553 | 0.351976 | -0.47151 | -1.404023 | 451 | 13 | c("CPB1", "BTG2", "LFNG") |
| 21 | HALLMARK_COMPLEMENT | 0.173148 | 0.412257 | -0.285358 | -1.235365 | 560 | 57 | c("S100A9", "LTF", "CTSS", "APOC1", "PRKCD", "EHD1", "PIM1", "LIPA", "PLAT", "C1QA", "CD36", "LTA4H", "C3", "LAP3", "C1QC") |
| 22 | HALLMARK_MYC_TARGETS_V2 | 0.190409 | 0.413932 | -0.553156 | -1.313216 | 670 | 6 | c("PA2G4", "HSPE1", "MYC", "GNL3", "HK2", "CBX3") |
| 23 | HALLMARK_BILE_ACID_METABOLISM | 0.184436 | 0.413932 | -0.45164 | -1.314242 | 601 | 12 | c("CAT", "DHCR24", "RXRA", "SLC29A1", "HSD17B4") |
| 24 | HALLMARK_TNFA_SIGNALING_VIA_NFKB | 0.217071 | 0.452231 | 0.271234 | 1.174709 | 1502 | 77 | c("CCN1", "F3", "SERPINE1", "GFPT2", "INHBA", "PLAU", "VEGFA", "KYNU", "ACKR3", "PMEPA1", "TNC", "RCAN1", "SPHK1", "GEM", "SPSB1", "PDLIM5", "EIF1", "FOSL2", "BTG1", "DUSP1", "KLF6", "MARCKS", "TSC22D1", "CDKN1A", "GADD45A", "PLAUR", "PHLDA1", "TGIF1", "CEBPB", "ETS2", "CD44", "RELA", "CEBPD", "ABCA1", "PNRC1", "JAG1", "SOCS3", "GADD45B", "PER1") |
| 25 | HALLMARK_CHOLESTEROL_HOMEOSTASIS | 0.269754 | 0.481703 | 0.355734 | 1.171616 | 1805 | 22 | c("GPX8", "ACTG1", "ERRFI1", "TRIB3", "ATF5", "CTNNB1", "ANTXR2", "PLAUR", "CLU", "PNRC1", "JAG1") |
| 26 | HALLMARK_WNT_BETA_CATENIN_SIGNALING | 0.265474 | 0.481703 | 0.433882 | 1.177449 | 1779 | 11 | c("FZD1", "CCND2", "CTNNB1", "NOTCH1", "NCOR2", "JAG1") |
| 27 | HALLMARK_ALLOGRAFT_REJECTION | 0.25279 | 0.481703 | -0.320136 | -1.172964 | 837 | 28 | c("CTSS", "SPI1", "TAPBP", "CD74", "HCLS1", "CCND3", "CAPG") |
| 28 | HALLMARK_SPERMATOGENESIS | 0.246036 | 0.481703 | -0.488442 | -1.223817 | 868 | 7 | c("SLC12A2", "TALDO1") |
| 29 | HALLMARK_PROTEIN_SECRETION | 0.331741 | 0.552901 | 0.29402 | 1.101966 | 2221 | 38 | c("PAM", "CAV2", "ARFGAP3", "SEC31A", "SEC24D", "TMED10", "TMED2", "USO1", "AP2M1", "RAB2A", "LMAN1", "SNX2", "ABCA1", "ARCN1", "LAMP2", "AP2S1", "RAB22A", "RAB5A", "MAPK1", "ARF1", "CLTC", "COPB2") |
| 30 | HALLMARK_IL2_STAT5_SIGNALING | 0.325987 | 0.552901 | 0.270022 | 1.09632 | 2203 | 55 | c("RGS16", "P4HA1", "COL6A1", "NDRG1", "NT5E", "ITGAV", "SPRY4", "IFITM3", "NRP1", "TWSG1", "ITGA6", "PRAF2", "SPP1", "KLF6", "CCND2", "EMP1", "CKAP4", "ECM1", "PRNP", "PLAGL1") |
| 31 | HALLMARK_PEROXISOME | 0.353666 | 0.570429 | -0.320595 | -1.090492 | 1176 | 21 | c("CAT", "DHCR24", "ABCC5", "ACAA1", "SEMA3C", "HSD17B4") |
| 32 | HALLMARK_ESTROGEN_RESPONSE_EARLY | 0.418575 | 0.654024 | -0.224618 | -1.020966 | 1315 | 69 | c("MYB", "BCL2", "TFF1", "SLC39A6", "FOS", "TFF3", "GFRA1", "TTC39A", "RET", "THSD4", "MAST4", "LRIG1", "SLC7A2", "MLPH", "UGCG", "CELSR1", "FASN") |
| 33 | HALLMARK_P53_PATHWAY | 0.464497 | 0.703783 | 0.242305 | 1.002939 | 3139 | 61 | c("RGS16", "DDIT4", "NDRG1", "SDC1", "TM4SF1", "KRT17", "S100A10", "SPHK1", "TGFB1", "LDHB", "FAM162A", "BTG1", "CCND2", "TRIB3", "TSC22D1", "CDKN1A", "GADD45A", "TAX1BP3", "TSPYL2", "ZFP36L1", "UPP1", "NOTCH1", "PTPN14") |
| 34 | HALLMARK_UNFOLDED_PROTEIN_RESPONSE | 0.499556 | 0.716232 | 0.250396 | 0.979648 | 3375 | 46 | c("DDIT4", "VEGFA", "KDELR3", "PDIA5", "ERO1A", "EIF4EBP1", "WFS1", "SEC31A", "TSPYL2", "LSM1", "ATF4", "HSPA5", "CEBPB", "EIF4A3", "EIF4A2", "PDIA6", "CALR", "DNAJC3", "SHC1", "ALDH18A1", "KHSRP", "YWHAZ", "HERPUD1", "CNOT2", "EEF2", "SEC11A", "HSPA9", "EIF2S1", "YIF1A", "STC2", "SRPRB", "EIF4G1", "MTHFD2", "GOSR2", "SPCS1", "KIF5B", "SRPRA", "SERP1") |
| 35 | HALLMARK_PI3K_AKT_MTOR_SIGNALING | 0.515687 | 0.716232 | -0.251694 | -0.951188 | 1692 | 32 | c("ITPR2", "MAP2K3", "GRK2", "VAV3", "MKNK2", "MYD88", "PIK3R3", "ACACA", "CXCR4") |
| 36 | HALLMARK_XENOBIOTIC_METABOLISM | 0.504173 | 0.716232 | -0.231295 | -0.964978 | 1630 | 49 | c("ESR1", "ALDH2", "APOE", "CAT", "PGD", "CSAD", "PDK4", "CD36", "FBP1", "CNDP2", "MCCC2", "GSR", "CYB5A", "PSMB10", "NINJ1") |
| 37 | HALLMARK_ADIPOGENESIS | 0.57963 | 0.783283 | -0.213651 | -0.924934 | 1877 | 57 | c("UCP2", "FABP4", "ALDH2", "APOE", "CAT", "TALDO1", "PDCD4", "CD36", "GPX3", "C3") |
| 38 | HALLMARK_MTORC1_SIGNALING | 0.625812 | 0.823437 | -0.202429 | -0.899982 | 2021 | 63 | c("CORO1A", "BTG2", "IFI30", "TFRC", "MAP2K3", "DHCR24", "SCD", "LTA4H", "TXNRD1", "GGA2", "PIK3R3", "HSPE1", "GSR", "ITGB2", "ACACA") |
| 39 | HALLMARK_NOTCH_SIGNALING | 0.668702 | 0.849636 | 0.299272 | 0.851205 | 4490 | 13 | c("FZD1", "FZD7", "NOTCH3", "NOTCH1") |
| 40 | HALLMARK_INTERFERON_GAMMA_RESPONSE | 0.679708 | 0.849636 | 0.225999 | 0.857659 | 4568 | 40 | c("MT2A", "VCAM1", "C1R", "IFITM3", "SERPING1", "IFITM2", "PFKP", "BTG1", "CFH", "VAMP5", "C1S", "CDKN1A", "UPP1") |
| 41 | HALLMARK_IL6_JAK_STAT3_SIGNALING | 0.743324 | 0.906493 | 0.247271 | 0.794403 | 4954 | 20 | c("OSMR", "TGFB1", "PDGFC", "IL1R1", "STAT3", "TNFRSF1A", "CD44", "SOCS3", "TNFRSF21", "CD9", "A2M", "MAP3K8", "GRB2") |
| 42 | HALLMARK_FATTY_ACID_METABOLISM | 0.777562 | 0.925669 | 0.224889 | 0.776355 | 5197 | 27 | c("ENO2", "TDO2", "S100A10", "MIF", "LGALS1", "PTPRG", "PRDX6", "UGDH", "EPHX1", "GLUL", "FH", "YWHAH", "GRHPR") |

| | | | | | | | |
|---|---|---|---|---|---|---|---|
| 43 | HALLMARK_G2M_CHECKPOINT | 0.859909 | 0.943511 | -0.191392 | -0.685189 | 2841 | 26 | c("SLC12A2", "RBM14", "SLC38A1", "BUB3", "SRSF2") |
| 44 | HALLMARK_ANDROGEN_RESPONSE | 0.839462 | 0.943511 | 0.196134 | 0.739207 | 5615 | 39 | c("NDRG1", "PMEPA1", "ITGAV", "MYL12A", "PDLIM5", "ACTN1", "SLC38A2", "KRT8", "TSC22D1", "ANKH", "SEC24D", "KRT19") |
| 45 | HALLMARK_INTERFERON_ALPHA_RESPONSE | 0.884051 | 0.943511 | 0.211691 | 0.661514 | 5908 | 18 | c("IFITM3", "IFITM2", "GBP2") |
| 46 | HALLMARK_MYC_TARGETS_V1 | 0.90577 | 0.943511 | -0.16063 | -0.701511 | 2950 | 59 | c("MCM7", "BUB3", "HNRNPA2B1", "PA2G4", "HSPE1", "SRSF2", "TXNL4A", "HDGF", "DHX15", "NHP2", "HNRNPA3", "VDAC3", "LSM7", "CUL1", "YWHAQ", "SRSF3", "HNRNPD", "MYC", "NME1", "ILF2", "PWP1", "CNBP", "PSMC4", "SF3B3", "HNRNPU", "FBL", "GNL3", "EIF4G2", "APEX1", "PPIA", "SET", "CBX3", "EIF2S1", "HSP90AB1", "UBA2", "DEK", "PSMD3", "ERH", "RACK1", "VDAC1", "XRCC6", "EEF1B2", "PSMB2", "PRPF31", "YWHAE", "PRPS2", "IMPDH2", "SNRPD2", "EIF3D", "GOT2", "TOMM70", "HNRNPC", "RAD23B", "PCBP1", "PRDX4", "GLO1", "UBE2E1", "CANX") |
| 47 | HALLMARK_OXIDATIVE_PHOSPHORYLATION | 0.835995 | 0.943511 | -0.171583 | -0.761967 | 2721 | 62 | c("MAOB", "PDK4", "ACAA1", "NDUFV1", "NDUFV2", "ATP6V0B", "CYB5A", "TCIRG1", "ATP6V0C", "UQCR10", "VDAC3", "NDUFB7", "NDUFB6", "OPA1", "ATP6V1F", "NDUFB5", "NDUFA3", "ISCU", "NDUFB2", "ATP5F1E", "MDH2", "NDUFA4", "POLR2F", "ATP5F1B", "HADHA", "ATP5MG", "IDH3G", "NDUFC1", "UQCR11", "ATP5MC2", "UQCRC2", "ATP5PD", "HSPA9", "COX6B1", "VDAC1", "COX6A1", "IDH2", "NDUFA1", "ATP5ME", "UQCRC1", "ATP6V0E1", "NDUFB1", "NDUFS4", "OXA1L", "GOT2", "COX7A2L", "NQO2", "ATP1B1", "FH", "COX7C", "PHYH", "TOMM70", "CYB5R3", <br>"TIMM17A", "UQCRB", "ATP5F1C", "ATP6V1E1", "GPI", "MGST3", "COX4I1") |
| 48 | HALLMARK_PANCREAS_BETA_CELLS | 0.889954 | 0.943511 | 0.277112 | 0.654133 | 5757 | 7 | c("FOXO1", "MAFB", "SEC11A", "SRPRB", "SPCS1", "AKT3", "SRP9") |
| 49 | HALLMARK_MITOTIC_SPINDLE | 0.967012 | 0.967012 | 0.146161 | 0.604985 | 6536 | 61 | c("FSCN1", "PALLD", "SYNPO", "CAPZB", "MYH9", "FLNA", "PDLIM5", "NEDD9") |
| 50 | HALLMARK_DNA_REPAIR | 0.964459 | 0.967012 | -0.148226 | -0.524705 | 3174 | 25 | c("CSTF3", "HCLS1", "NME3", "DDB1", "NELFCD", "BCAM") |

| Cluster 14 | pathway | pval | padj | ES | NES | nMoreExtr | size | leadingEdge |
|---|---|---|---|---|---|---|---|---|
| 1 | HALLMARK_KRAS_SIGNALING_UP | 0.004328 | 0.21638 | -0.34077 | -1.905 | 39 | 42 | c("MMP9", "CXCR4", "GPNMB", "IKZF1", "SERPINA3", "TMEM176B", "LCP1", "ITGB2", "PLAUR", "PLAT", "TSPAN13", "CTSS", "GYPC", "ANKH", "CPE", "PLAU", "SEMA3B", "MAFB", "ITGBL1", "FCER1G", "NRP1", "LAPTM5", "SPARCL1", "IGFBP3") |
| 2 | HALLMARK_HEME_METABOLISM | 0.009842 | 0.246048 | -0.28684 | -1.77212 | 93 | 61 | c("HBB", "HBD", "SLC4A1", "SNCA", "NCOA4", "FBXO9", "SEC14L1", "SLC25A37", "ACP5", "SLC2A1", "TFRC", "FOXO3", "CIR1", "GYPC", "MPP1", "TNS1", "TRAK2", "KHNYN", "USP15", "VEZF1", "FBXO7", "EIF2AK1", "ELL2", "CAT", "LAMP2", "PGLS", "UROD", "RAD23A", "ARHGEF12", "LRP10", "HAGH", "BSG", "BACH1", "CCND3", "TOP1", "GLRX5", "HDGF", "PSMD9", "SELENBP1", "BLVRB", "MAP2K3", "UBAC1", "NFE2L1", "RBM5", "UCP2", "PICALM", "RNF19A", "BTG2") |
| 3 | HALLMARK_WNT_BETA_CATENIN_SIGNALING | 0.055099 | 0.597185 | -0.59186 | -1.51627 | 368 | 5 | c("MYC", "CTNNB1", "NCSTN") |
| 4 | HALLMARK_ESTROGEN_RESPONSE_EARLY | 0.059718 | 0.597185 | -0.23303 | -1.49064 | 576 | 70 | c("PAPSS2", "GFRA1", "RHOBTB3", "B4GALT1", "MYC", "HSPB8", "RET", "RARA", "SLC2A1", "GAB2", "ADCY1", "KDM4B", "SEC14L2", "FAM102A", "AMFR", "PGR", "SH3BP5", "THSD4", "CELSR1", "CANT1", "ABHD2", "SLC9A3R1", "MAPT", "RHOD", "ADD3", "SEMA3B") |
| 5 | HALLMARK_KRAS_SIGNALING_DN | 0.048166 | 0.597185 | -0.43825 | -1.58455 | 368 | 12 | c("GPRC5C", "BMPR1B", "KMT2D", "LFNG", "CPB1", "SGK1", "EFHD1", "SKIL") |
| 6 | HALLMARK_ESTROGEN_RESPONSE_LATE | 0.119257 | 0.662542 | -0.21509 | -1.36344 | 1149 | 67 | c("CDH1", "PAPSS2", "SERPINA1", "TOP2A", "S100A9", "SCUBE2", "LTF", "HSPB8", "RET", "SERPINA3", "TSPAN13", "FAM102A", "AMFR", "PGR", "MAPK13", "CPE", "ABHD2", "SLC9A3R1", "MAPT", "ADD3", "LLGL2", "SGK1", "SEMA3B", "AGR2") |
| 7 | HALLMARK_APICAL_SURFACE | 0.108236 | 0.662542 | -0.45407 | -1.40421 | 777 | 8 | c("B4GALT1", "PLAUR", "GSTM3") |
| 8 | HALLMARK_MTORC1_SIGNALING | 0.090321 | 0.662542 | -0.24351 | -1.4283 | 850 | 50 | c("PSMB5", "CXCR4", "CORO1A", "PSMD13", "SLC2A1", "ITGB2", "STC1", "TFRC", "ACTR3", "TXNRD1", "ATP2A2", "IFI30", "P4HA1", "SYTL2", "SLC9A3R1", "ADD3", "GGA2", "SLC1A5", "GSK3B", "PSMC4", "CANX") |
| 9 | HALLMARK_IL2_STAT5_SIGNALING | 0.116413 | 0.662542 | -0.24299 | -1.38859 | 1087 | 45 | c("PIM1", "MYC", "ANXA4", "ITGA6", "NFKBIZ", "CAPG", "P4HA1", "CKAP4", "HIPK2", "S100A1", "PHLDA1", "SLC1A5", "IGF2R", "NRP1", "MAP3K8", "BCL2", "ITGAV", "BCL2L1", "RNH1", "CYFIP1", "EMP1", "CCND3", "RABGAP1L", "SNX14", "IGF1R") |
| 10 | HALLMARK_COMPLEMENT | 0.167372 | 0.721859 | -0.21864 | -1.29795 | 1582 | 52 | c("PRKCD", "SERPINA1", "GRB2", "S100A9", "PIM1", "APOC1", "LTF", "PLAUR", "LIPA", "PLAT", "HSPA1A", "CTSS", "CD55", "USP15") |
| 11 | HALLMARK_E2F_TARGETS | 0.146214 | 0.721859 | -0.28108 | -1.34206 | 1262 | 26 | c("HMGB2", "TOP2A", "MYC", "TFRC", "ANP32E", "RBBP7", "NUDT21", "SMC4", "POLD2", "NME1", "HNRNPD", "LUC7L3", "NOP56", "MCM7", "NAA38", "SNRPB", "CDKN1B", "STAG1", "SSRP1", "PA2G4") |
| 12 | HALLMARK_FATTY_ACID_METABOLISM | 0.187683 | 0.721859 | 0.194296 | 1.234833 | 255 | 26 | c("LGALS1", "ECH1", "MIF", "ALDH9A1", "APEX1", "ELOVL5", "HSP90AA1", "ACAA1", "HSD17B4", "ACADVL", "SERINC1", "DHCR24", "PCBD1", "PDHA1", "PSME1", "ECHS1", "ECI1", "SMS", "UGDH") |
| 13 | HALLMARK_BILE_ACID_METABOLISM | 0.174712 | 0.721859 | 0.290323 | 1.257729 | 408 | 12 | c("ALDH9A1", "GSTK1", "HSD17B4", "RXRA", "DHCR24", "OPTN", "IDH2", "LONP2", "SLC29A1", "SOD1", "SCP2", "CAT") |
| 14 | HALLMARK_NOTCH_SIGNALING | 0.239288 | 0.753665 | -0.39061 | -1.20796 | 1719 | 8 | c("PSENEN", "LFNG", "NOTCH2", "APH1A", "RBX1", "NOTCH3", "ARRB1") |
| 15 | HALLMARK_UNFOLDED_PROTEIN_RESPONSE | 0.234481 | 0.753665 | -0.24318 | -1.21684 | 2069 | 30 | c("DCTN1", "ATP6V0D1", "EIF4A3", "EIF4A2", "EIF4EBP1", "BANF1", "GOSR2", "KIF5B", "LSM1", "HERPUD1", "SRPRA", "DNAJA4", "LSM4", "SEC31A", "NOP56", "EIF4G1", "FUS", "CEBPB") |
| 16 | HALLMARK_GLYCOLYSIS | 0.241173 | 0.753665 | -0.21137 | -1.20791 | 2253 | 45 | c("FUT8", "CXCR4", "B4GALT1", "CITED2", "TGFBI", "RBCK1", "STC1", "SDC1", "P4HA1", "PLOD1", "TALDO1", "GFPT1", "PSMC4", "SDC3", "PGLS", "IGFBP3", "AKR1A1", "ENO1", "SOD1", "ARPP19", "PYGL", "TFF3", "PKM", "IER3", "QSOX1", "GPC1", "IL13RA1", "SDC2", "COPB2", "EGLN3", "CHPF", "ELF3", "PPIA", "IDUA", "PYGB") |
| 17 | HALLMARK_G2M_CHECKPOINT | 0.269312 | 0.772471 | -0.22672 | -1.17425 | 2415 | 33 | c("TOP2A", "ABL1", "MYC", "SMAD3", "CUL3", "YTHDC1", "KIF5B", "SMC4", "BCL3", "MT2A", "PAFAH1B1", "SLC38A1", "SMARCC1", "NOTCH2", "HNRNPD", "CUL4A", "TOP1", "RBM14", "SLC12A2", "CDKN1B", "MARCKS", "STAG1") |
| 18 | HALLMARK_ANDROGEN_RESPONSE | 0.27809 | 0.772471 | -0.21897 | -1.16919 | 2526 | 36 | c("NCOA4", "B4GALT1", "STEAP4", "BMPR1B", "SRP19", "ANKH", "ABHD2", "ELL2", "SGK1", "SLC38A2", "UAP1", "ITGAV", "INPP4B", "UBE2I", "TPD52", "SCD", "ADRM1", "ACSL3", "GNAI3", "CCND3") |

| | | | | | | | | |
|---|---|---|---|---|---|---|---|---|
| 19 | HALLMARK_DNA_REPAIR | 0.322818 | 0.849521 | -0.21662 | -1.12192 | 2895 | 33 | c("NME3", "BRF2", "POLR1D", "POLR2I", "EIF1B", "HCLS1", "DDB1", "ERCC1", "VPS28", "CANT1", "NUDT21") |
| 20 | HALLMARK_PROTEIN_SECRETION | 0.352702 | 0.881754 | -0.2055 | -1.09727 | 3204 | 36 | c("MAPK1", "TOM1L1", "ARFGEF2", "ANP32E", "GOSR2", "RER1", "MON2", "STX16", "STX7", "IGF2R", "LAMP2", "M6PR", "RAB2A", "TPD52", "ARF1", "SEC31A", "SOD1", "VPS45", "PPT1", "TMED2", "NAPA") |
| 21 | HALLMARK_XENOBIOTIC_METABOLISM | 0.384229 | 0.883838 | -0.18693 | -1.06823 | 3590 | 45 | c("PAPSS2", "TMEM176B", "HMOX1", "ATP2A2", "CD36", "CSAD", "SLC1A5", "CAT", "MT2A", "TNFRSF1A", "DHRS7", "COMT", "EPHX1", "DDAH2", "BCAR1", "ARPP19", "NQO1", "PGRMC1", "PDK4", "SERPINE1", "MCCC2", "MAN1A1", "PSMB10", "BLVRB", "DCXR", "ESR1", "PGD", "UGDH", "CNDP2") |
| 22 | HALLMARK_OXIDATIVE_PHOSPHORYLATION | 0.388889 | 0.883838 | 0.09746 | 1.029396 | 125 | 72 | c("GPX4", "ECH1", "ATP5MC2", "CYB5R3", "ETFB", "GLUD1", "NDUFV2", "ATP6V1F", "CYB5A", "OXA1L", "COX4I1", "TIMM13", "ATP5MG", "BAX", "ACAA1", "COX7A2", "COX8A", "NDUFA4", "ACADVL", "ATP6V0B", "ISCU", "NDUFB7", "NDUFS2", "COX7C", "NDUFA3", "UQCR10", "PDHA1", "ATP6AP1", "ATP5F1E", "NDUFV1", "COX6C", "NDUFS6", "COX7A2L", "ECHS1", "NDUFB6", "MGST3", "HADHA", "ATP5F1D", "IDH2", "NDUFAB1", "NDUFS4", "ECI1", "NDUFA2", "ATP5PD", "NDUFB2") |
| 23 | HALLMARK_EPITHELIAL_MESENCHYMAL_TRANSITIO N | 0.440789 | 0.918311 | 0.079027 | 0.997277 | 66 | 98 | c("TIMP1", "TPM1", "CD44", "HTRA1", "TAGLN", "COL5A2", "COL5A1", "COL3A1", "COL1A2", "SPARC", "LGALS1", "COL6A3", "COL6A2", "COL1A1", "FN1", "MGP", "SPP1", "CALD1", "CTHRC1", "MYL9", "TPM2", "IGFBP2", "TPM4", "DST", "JUN", "PFN2", "CCN1", "CCN2", "FBLN1", "FLNA", "ECM1", "FAP", "ITGB5", "FBN1", "PMEPA1", "MMP14", "CXCL12", "VEGFA", "LRRC15", "SDC4", "COL4A1", "LRP1", "ID2", "CALU", "GEM", "COL4A2", "LOX", "SERPINH1", "DPYSL3", "THBS2", "INHBA", "ITGA5", "TNFRSF12A", "GJA1", "COPA") |
| 24 | HALLMARK_ALLOGRAFT_REJECTION | 0.440092 | 0.918311 | -0.20923 | -1.01241 | 3830 | 27 | c("MMP9", "ITGB2", "SRGN", "SPI1", "HCLS1", "CTSS", "CAPG", "NME1", "BCL3", "IFNGR2", "TPD52", "IKBKB") |
| 25 | HALLMARK_TNFA_SIGNALING_VIA_NFKB | 0.568345 | 0.931162 | 0.092291 | 0.907627 | 236 | 63 | c("BHLHE40", "CD44", "JUNB", "MCL1", "PNRC1", "TGIF1", "EGR1", "IER2", "FOS", "LITAF", "DUSP1", "JUN", "CCN1", "NFE2L2", "SOD2", "NR4A1", "PMEPA1", "NINJ1", "HES1", "VEGFA", "CEBPD", "KLF9", "SDC4", "SOCS3", "EIF1", "KLF2", "ID2", "IER5", "GEM", "INHBA", "EHD1", "EFNA1", "PFKFB3", "ZFP36", "FOSL2", "TNIP1", "CFLAR") |
| 26 | HALLMARK_HYPOXIA | 0.692332 | 0.931162 | -0.13313 | -0.83005 | 6635 | 63 | c("HK1", "PIM1", "CXCR4", "DTNA", "CITED2", "TGFBI", "SLC2A1", "STC1", "PLAUR", "HMOX1", "FOXO3") |
| 27 | HALLMARK_CHOLESTEROL_HOMEOSTASIS | 0.899139 | 0.931162 | -0.15288 | -0.62994 | 7309 | 17 | c("CTNNB1", "PLAUR", "SEMA3B", "PMVK", "ANXA5", "SCD", "LGMN", "ATXN2", "LGALS3", "TNFRSF12A", "ALCAM") |
| 28 | HALLMARK_MITOTIC_SPINDLE | 0.557608 | 0.931162 | -0.1526 | -0.93449 | 5313 | 59 | c("PREX1", "LRPPRC", "TOP2A", "NEDD9", "CKAP5", "ABL1") |
| 29 | HALLMARK_TGF_BETA_SIGNALING | 0.778768 | 0.931162 | -0.1556 | -0.74294 | 6726 | 26 | c("CDH1", "FNTA", "SMAD3", "CTNNB1", "ARID4B", "HIPK2", "SKI", "SKIL", "THBS1", "IFNGR2", "PPP1CA", "SERPINE1", "ID3", "NCOR2") |
| 30 | HALLMARK_IL6_JAK_STAT3_SIGNALING | 0.550118 | 0.931162 | -0.20154 | -0.92229 | 4675 | 23 | c("GRB2", "PIM1", "LTBR", "HMOX1", "CD36", "STAT2", "MAP3K8", "STAT3", "TNFRSF1A", "IFNGR2", "PTPN1") |
| 31 | HALLMARK_APOPTOSIS | 0.840499 | 0.931162 | -0.11597 | -0.69222 | 7956 | 53 | c("HMGB2", "TOP2A", "NEDD9", "RARA", "CTNNB1", "PLAT", "HMOX1") |
| 32 | HALLMARK_ADIPOGENESIS | 0.759432 | 0.931162 | -0.12741 | -0.77408 | 7225 | 57 | c("COX6A1", "HSPB8", "FABP4", "UQCRQ", "UBC", "TKT", "GPAT4", "G3BP2", "YWHAG", "RAB34", "CD36", "SLC1A5", "TALDO1", "UQCRC1", "CAT", "SPARCL1", "CAVIN1", "CCNG2", "PHLDB1", "SCP2", "DHRS7", "PIM3", "LAMA4", "SOD1", "SDHB", "PPP1R15B", "CYC1", "UQCR11", "BCL2L13") |
| 33 | HALLMARK_MYOGENESIS | 0.835017 | 0.931162 | 0.083522 | 0.719258 | 495 | 48 | c("APP", "BHLHE40", "CLU", "TAGLN", "SYNGR2", "IGFBP7", "AEBP1", "COL3A1", "SPARC", "COL6A3", "COL6A2", "COL1A1", "CNN3", "TPM2", "CFD", "MYH9", "GSN", "AGRN", "APOD", "PLXNB2", "ITGB5", "SPDEF", "SPTAN1", "SORBS3", "MEF2D", "COL4A2", "ATP6AP1", "MYO1C", "SH2B1", "FLII", "BAG1", "ITGB1") |
| 34 | HALLMARK_INTERFERON_ALPHA_RESPONSE | 0.874508 | 0.931162 | 0.130735 | 0.693958 | 1553 | 18 | c("LGALS3BP", "ADAR", "B2M", "CD74", "TXNIP", "IFITM3", "C1S", "ELF1", "CD47", "LY6E", "PSME1", "IFITM2") |

| | | | | | | | | |
|---|---|---|---|---|---|---|---|---|
| 35 | HALLMARK_INTERFERON_GAMMA_RESPONSE | 0.667849 | 0.931162 | 0.10918 | 0.857236 | 564 | 39 | c("LGALS3BP", "ADAR", "B2M", "CD74", "TXNIP", "RNF213", "SAMHD1", "TAPBP", "IFITM3", "C1S", "SOD2", "SERPING1", "CFB", "SOCS3", "LY6E", "PSME1", "MYD88", "ARID5B", "C1R", "IFITM2", "STAT1", "EIF4E3", "SRI", "VAMP8", "LAP3", "PSMB10", "BST2", "BTG1", "TNFAIP2", "PTPN1", "TRIM25", "STAT3", "MT2A", "NFKBIA") |
| 36 | HALLMARK_APICAL_JUNCTION | 0.813804 | 0.931162 | -0.11921 | -0.71848 | 7722 | 55 | c("CDH1", "INPPL1", "THBS3", "MMP9", "B4GALT1") |
| 37 | HALLMARK_HEDGEHOG_SIGNALING | 0.64761 | 0.931162 | -0.26075 | -0.84625 | 4768 | 9 | c("ADGRG1", "CELSR1", "RASA1", "NRP1", "LDB1") |
| 38 | HALLMARK_PI3K_AKT_MTOR_SIGNALING | 0.619648 | 0.931162 | -0.17256 | -0.87353 | 5499 | 31 | c("MAPK1", "GRB2", "CXCR4", "SLC2A1", "ACTR3", "PAK4", "STAT2", "GSK3B", "PLA2G12A", "ITPR2", "TNFRSF1A", "VAV3", "ARF1", "RAC1", "PPP1CA", "CAB39", "CDKN1B", "MAP2K3", "SMAD2") |
| 39 | HALLMARK_MYC_TARGETS_V1 | 0.931162 | 0.931162 | -0.0987 | -0.58315 | 8805 | 51 | c("MYC", "CLNS1A", "LSM7", "VDAC3", "GNL3", "ILF2", "POLD2", "SNRPD2", "NME1", "PSMC4", "CANX", "SMARCC1", "HNRNPD", "RNPS1", "YWHAE", "NOP56", "MCM7", "TUFM", "EIF3B", "HDGF", "CYC1", "CNBP", "PA2G4", "SF3B3", "NDUFAB1", "CBX3", "PPIA", "PSMA7", "IMPDH2", "EIF4A1", "EEF1B2", "EIF4H", "NCBP2", "HNRNPC", "SRSF2", "HNRNPU", "PSMD3", "HNRNPA3", "HSP90AB1", "BUB3") |
| 40 | HALLMARK_MYC_TARGETS_V2 | 0.844065 | 0.931162 | -0.23528 | -0.68972 | 5937 | 7 | c("MYC", "GNL3") |
| 41 | HALLMARK_INFLAMMATORY_RESPONSE | 0.898909 | 0.931162 | -0.12921 | -0.6252 | 7824 | 27 | c("MYC", "CYBB", "PLAUR", "ATP2A2", "CD55", "ATP2C1", "NFKBIA", "IFNGR2", "ADRM1", "GNAI3", "STAB1", "SERPINE1", "BST2", "EMP3", "KLF6", "BTG2", "SRI") |
| 42 | HALLMARK_REACTIVE_OXYGEN_SPECIES_PATHWAY | 0.836249 | 0.931162 | -0.17232 | -0.69388 | 6740 | 16 | c("MPO", "TXNRD1", "CAT", "EGLN2", "LSP1", "SOD1", "NQO1", "PTPA", "GPX3", "SBNO2", "GSR", "NDUFS2", "ATOX1") |
| 43 | HALLMARK_P53_PATHWAY | 0.790419 | 0.931162 | -0.12655 | -0.73875 | 7440 | 49 | c("EPS8L2", "IP6K2", "ABCC5", "RAP2B", "SDC1", "HMOX1", "FOXO3", "IFI30", "PLK2", "FBXW7") |
| 44 | HALLMARK_UV_RESPONSE_UP | 0.552764 | 0.931162 | -0.17236 | -0.92756 | 5038 | 37 | c("PRKCD", "RET", "TFRC", "HMOX1", "MGAT1", "NXF1", "EPCAM", "CREG1", "NFKBIA", "UROD", "ARRB2", "EPHX1", "BSG", "FOSB", "BTG1", "CCND3", "PPT1", "CLTB", "PRPF3", "YKT6", "GPX3", "EIF5", "WIZ", "BTG2") |
| 45 | HALLMARK_UV_RESPONSE_DN | 0.697982 | 0.931162 | -0.14222 | -0.81748 | 6537 | 46 | c("MYC", "CITED2", "ANXA4", "SMAD3", "LAMC1", "EFEMP1", "YTHDC1", "ATP2C1", "ADD3", "NRP1", "RBPMS", "DAB2", "PMP22", "PDGFRB", "COL11A1", "INPP4B", "NOTCH2") |
| 46 | HALLMARK_ANGIOGENESIS | 0.538165 | 0.931162 | -0.24847 | -0.92725 | 4194 | 13 | c("STC1", "FGFR1", "NRP1", "ITGAV", "FSTL1") |
| 47 | HALLMARK_COAGULATION | 0.911392 | 0.931162 | -0.10839 | -0.60594 | 8423 | 42 | c("SERPINA1", "MMP9", "APOC1") |
| 48 | HALLMARK_PEROXISOME | 0.921905 | 0.931162 | 0.110398 | 0.650436 | 1451 | 22 | c("ECH1", "ALDH9A1", "ELOVL5", "GSTK1", "SOD2", "CRABP2", "ACAA1", "HSD17B4") |
| 49 | HALLMARK_SPERMATOGENESIS | 0.550704 | 0.931162 | -0.33565 | -0.92283 | 3795 | 6 | c("VDAC3", "GSTM3", "TALDO1") |
| 50 | HALLMARK_PANCREAS_BETA_CELLS | 0.803631 | 0.931162 | 0.262959 | 0.738894 | 2655 | 5 | c("SRP9", "SEC11A", "SPCS1") |

| Cluster 15 | pathway | pval | padj | ES | NES | nMoreExtreme | size | leadingEdge |
|---|---|---|---|---|---|---|---|---|
| 1 | HALLMARK_ESTROGEN_RESPONSE_EARLY | 0.0001 | 0.001012 | -0.458517 | -1.987588 | 0 | 68 | c("NBL1", "GJA1", "SEC14L2", "ADCY1", "PGR", "MAPT", "SCNN1A", "NRIP1", "HSPB8", "RHOD", "RHOBTB3", "CXCL12", "FHL2", "GREB1", "CA12", "SEMA3B", "PAPSS2", "FOS", "RET", "SLC7A2", "FARP1", "MAST4", "ELOVL5", "BCL2", "RAB31", "B4GALT1", "ELF3", "GAB2", "MED13L", "LRIG1", "ABHD2", "SLC2A1", "CELSR1", "TTC39A", "SH3BP5", "IGF1R", "KDM4B", "MYOF", "FAM102A", "PLAAT3", "MYC", "THSD4", "AKAP1") |
| 2 | HALLMARK_COMPLEMENT | 0.000101 | 0.001012 | -0.516327 | -2.154112 | 0 | 51 | c("C1QA", "SERPINE1", "C1S", "C1QC", "APOC1", "COL4A2", "MMP14", "SERPING1", "C3", "LGMN", "LGALS3", "LIPA", "PLAT", "CD36", "C1R", "CTSD", "CTSL", "LTF") |
| 3 | HALLMARK_EPITHELIAL_MESENCHYMAL_TRANSITIO N | 1.00E-04 | 0.001012 | -0.668912 | -3.021339 | 0 | 99 | c("SFRP4", "COMP", "FSTL1", "COL16A1", "ADAM12", "SERPINE1", "FAP", "LRRC15", "GEM", "ACTA2", "CCN2", "THBS2", "THY1", "IGFBP3", "ELN", "LAMC1", "CDH11", "GJA1", "COL4A2", "MMP14", "COL12A1", "FBN1", "INHBA", "DAB2", "FUCA1", "COL4A1", "LOXL1", "CCN1", "FBLN2", "PRRX1", "CTHRC1", "DST", "PMEPA1", "CALD1", "NNMT", "LOX", "LOXL2", "PCOLCE", "FBLN1", "DPYSL3", "COL11A1", "FLNA", "PDGFRB", "SERPINH1", "JUN", "MXRA5", "CXCL12", "MYL9", "PFN2", "SDC1", "ITGB5", "TPM2", "TNC") |
| 4 | HALLMARK_COAGULATION | 0.000101 | 0.001012 | -0.625861 | -2.523659 | 0 | 41 | c("COMP", "C1QA", "SERPINE1", "C1S", "MMP11", "CTSK", "A2M", "APOC1", "MMP14", "FBN1", "PLAU", "GSN", "SERPING1", "C3", "S100A1", "LGMN", "PLAT", "C1R", "MMP9", "CRIP2") |
| 5 | HALLMARK_KRAS_SIGNALING_UP | 0.000101 | 0.001012 | -0.552679 | -2.236048 | 0 | 42 | c("APOD", "GPNMB", "MMP11", "SPARCL1", "IGFBP3", "NRP1", "PLAU", "INHBA", "FUCA1", "ITGBL1", "PRRX1", "PLAT", "MMP9", "CPE", "SERPINA3", "EMP1", "MAFB", "SEMA3B", "ENG", "TMEM176B") |
| 6 | HALLMARK_UV_RESPONSE_DN | 0.000504 | 0.004202 | -0.468708 | -1.918994 | 4 | 45 | c("SERPINE1", "PLPP3", "NRP1", "LAMC1", "GJA1", "SDC2", "DAB2", "CCN1", "COL11A1", "RBPMS", "PDGFRB", "AKT3", "EFEMP1", "FHL2", "ATP2B4", "RUNX1", "CDKN1B", "ANXA4", "APBB2", "PMP22", "TGFBR2", "PIK3R3", "CITED2") |
| 7 | HALLMARK_TNFA_SIGNALING_VIA_NFKB | 0.000801 | 0.005651 | -0.421177 | -1.807489 | 7 | 63 | c("SERPINE1", "FOSB", "GEM", "PHLDA1", "PLPP3", "PLAU", "INHBA", "EGR1", "CCN1", "PMEPA1", "KLF9", "PFKFB3", "NR4A1", "MARCKS", "JUN", "DUSP4", "PLK2", "TNC", "SOCS3", "FOSL2", "FOS") |
| 8 | HALLMARK_APICAL_JUNCTION | 0.000904 | 0.005651 | -0.43766 | -1.841117 | 8 | 54 | c("COL16A1", "THY1", "CDH11", "FBN1", "SDC3", "PARVA", "AKT3", "MMP9", "CD276", "CDH1", "MYL9", "SHC1", "MDK", "THBS3", "INPPL1", "CERCAM", "TSPAN4", "TGFBI", "LIMA1", "SORBS3", "B4GALT1", "PIK3R3", "CTNNA1") |
| 9 | HALLMARK_HYPOXIA | 0.001502 | 0.008347 | -0.413486 | -1.77448 | 14 | 63 | c("SERPINE1", "CCN2", "HMOX1", "IGFBP3", "SDC2", "CAVIN1", "CCN1", "PFKFB3", "LOX", "PAM", "SDC3", "MT2A", "JUN", "STC1", "TGFB3", "DDIT4", "CA12", "FOSL2", "FOS", "TPD52", "TGFBI", "BCL2", "CDKN1B", "CITED2", "PLAUR", "SLC2A1", "HEXA", "IER3", "SDC4", "GPC1", "NEDD4L") |
| 10 | HALLMARK_ESTROGEN_RESPONSE_LATE | 0.00841 | 0.04205 | -0.371065 | -1.604983 | 83 | 67 | c("CXCL14", "NBL1", "PGR", "MAPT", "SCNN1A", "NRIP1", "HSPB8", "CXCL12", "CDH1", "CPE", "SERPINA3", "LTF", "CA12", "MDK", "SEMA3B", "PAPSS2", "PRLR", "FOS", "RET", "ATP2B4", "FARP1", "ELOVL5", "BCL2", "IDH2", "RAB31", "SCUBE2", "SERPINA1", "ABHD2", "SGK1", "SLC29A1", "MYOF", "FAM102A", "PLAAT3") |
| 11 | HALLMARK_MYOGENESIS | 0.012214 | 0.055516 | -0.398919 | -1.627719 | 120 | 44 | c("APOD", "ADAM12", "PVALB", "IGFBP3", "COL4A2", "GPX3", "GSN", "HSPB8", "CD36", "CNN3", "ITGB5", "TPM2") |
| 12 | HALLMARK_TGF_BETA_SIGNALING | 0.024507 | 0.095115 | -0.436491 | -1.610766 | 235 | 26 | c("SERPINE1", "LTBP2", "ID3", "PMEPA1", "CDH1", "FNTA", "ENG", "HIPK2", "RAB31", "SKI", "THBS1", "BMPR2") |
| 13 | HALLMARK_IL2_STAT5_SIGNALING | 0.02473 | 0.095115 | -0.380047 | -1.550713 | 244 | 44 | c("ITGA6", "PHLDA1", "NRP1", "S100A1", "EMP1", "CD81", "AHR", "NFKBIZ", "ITGAV", "HIPK2", "BCL2", "ANXA4", "CAPG", "CYFIP1", "BMPR2", "LRIG1", "MAPKAPK2", "MAP3K8", "IGF1R", "MYC", "CKAP4", "KLF6", "SNX14", "PLEC", "SERPINB6", "MYO1C", "PIM1", "ENPP1") |
| 14 | HALLMARK_P53_PATHWAY | 0.040354 | 0.144122 | -0.356115 | -1.476779 | 400 | 49 | c("HMOX1", "FUCA1", "CTSD", "JUN", "SDC1", "DDIT4", "PLK2", "RAP2B", "CD81", "S100A10", "FOS", "HEXIM1", "EPHX1", "BAX", "EI24", "IFI30", "IP6K2", "IER3", "STOM", "SEC61A1", "EPS8L2", "BTG2", "MDM2", "MXD4", "ISCU", "PLXNB2", "SLC3A2", "IER5", "TAX1BP3", "BTG1") |

| | | | | | | | | |
|---|---|---|---|---|---|---|---|---|
| 15 | HALLMARK_ADIPOGENESIS | 0.054724 | 0.147109 | -0.33685 | -1.42117 | 544 | 55 | c("FABP4", "SPARCL1", "LAMA4", "GPX3", "CAVIN1", "COL4A1", "C3", "HSPB8", "PFKFB3", "CD36", "PHLDB1", "RAB34", "CMBL", "ALDH2", "GPAT4", "DHRS7", "VEGFB", "PIM3", "SCP2", "STOM", "DDT", "UBC", "TALDO1", "CAT", "UQCR10", "UQCRQ", "CYC1") |
| 16 | HALLMARK_APICAL_SURFACE | 0.051821 | 0.147109 | -0.594513 | -1.51842 | 413 | 7 | c("THY1", "SULF2", "B4GALT1", "PLAUR") |
| 17 | HALLMARK_MYC_TARGETS_V1 | 0.046154 | 0.147109 | 0.174969 | 1.483382 | 2 | 49 | c("RACK1", "BUB3", "TXNL4A", "CNBP", "DHX15", "EIF4A1", "CBX3", "NOP56", "HNRNPU", "HNRNPD", "SMARCC1", "APEX1", "VDAC3", "TUFM", "PPIA", "HSP90AB1", "YWHAE", "FAM120A", "ILF2", "HDGF", "EIF4H", "HNRNPA3", "IMPDH2", "CLNS1A", "EEF1B2", "NME1", "RNPS1", "CCT3", "MCM7", "HNRNPC", "SRM", "PSMD3", "POLD2", "NDUFAB1", "SF3B3", "SNRPD2", "PA2G4", "EIF3B", "SRSF2", "PSMA7", "PCBP1", "CYC1", "GNL3", "MYC", "NCBP2", "NHP2", "HSPE1") |
| 18 | HALLMARK_GLYCOLYSIS | 0.047192 | 0.147109 | -0.358201 | -1.466554 | 467 | 45 | c("IGFBP3", "EGLN3", "SDC2", "PAM", "SDC3", "STC1", "SDC1", "DDIT4", "CHPF", "PLOD1", "TGFBI", "AKR1A1", "B4GALT1", "ELF3", "CITED2", "GFPT1", "IER3", "PKM", "CYB5A", "GPC1", "TALDO1", "HDLBP", "MIF", "PYGB") |
| 19 | HALLMARK_KRAS_SIGNALING_DN | 0.055901 | 0.147109 | -0.501774 | -1.520297 | 493 | 12 | c("CPB1", "BMPR1B", "SELENOP", "GPRC5C", "IGFBP2", "SGK1", "BTG2", "SKIL") |
| 20 | HALLMARK_XENOBIOTIC_METABOLISM | 0.076636 | 0.189614 | -0.340545 | -1.394266 | 759 | 45 | c("SERPINE1", "HMOX1", "PDK4", "CD36", "FBLN1", "MT2A", "PAPSS2", "MAN1A1", "BCAR1", "TMEM176B", "ALDH2", "ELOVL5", "EPHX1", "DHRS7", "PGRMC1", "NQO1", "COMT", "CYB5A", "DDT", "CNDP2", "CAT", "DDAH2", "UGDH") |
| 21 | HALLMARK_UV_RESPONSE_UP | 0.079638 | 0.189614 | -0.356004 | -1.408817 | 782 | 37 | c("FOSB", "HMOX1", "GPX3", "MMP14", "NR4A1", "UROD", "FOS", "RET", "CREG1", "CLTB", "PPT1", "IGFBP2", "EPHX1") |
| 22 | HALLMARK_INFLAMMATORY_RESPONSE | 0.089616 | 0.203672 | -0.381868 | -1.409192 | 862 | 26 | c("SERPINE1", "MMP14", "INHBA", "STAB1", "AHR", "ITGA5", "SLC7A2", "CYBB", "PLAUR", "NFKBIA", "SRI", "BST2", "MYC", "BTG2", "GNAI3", "KLF6") |
| 23 | HALLMARK_ANDROGEN_RESPONSE | 0.095466 | 0.207535 | -0.35286 | -1.382784 | 936 | 35 | c("BMPR1B", "PMEPA1", "SELENOP", "MAF", "ELL2", "ITGAV", "TPD52", "ELOVL5", "STEAP4", "B4GALT1", "ACSL3", "DBI", "ANKH", "ABHD2", "ARID5B", "SGK1", "GNAI3", "TMEM50A", "SMS", "INPP4B") |
| 24 | HALLMARK_DNA_REPAIR | 0.131148 | 0.273224 | 0.188771 | 1.26215 | 31 | 32 | c("DDB1", "TAF10", "NELFCD", "HCLS1", "POLD4", "EIF1B", "POLR2H", "BCAM", "NME4", "IMPDH2", "POLR2A", "APRT", "NME1", "GUK1", "ADRM1", "POLR2I", "VPS28", "CSTF3", "NME3", "RBX1", "CANT1", "DCTN4", "SUPT5H", "POLR1D", "NUDT21", "SSRP1", "SEC61A1", "BRF2", "ERCC1", "NCBP2", "TMED2") |
| 25 | HALLMARK_ANGIOGENESIS | 0.139764 | 0.279527 | -0.425081 | -1.344774 | 1264 | 14 | c("FSTL1", "NRP1", "STC1", "ITGAV") |
| 26 | HALLMARK_HEDGEHOG_SIGNALING | 0.159199 | 0.306152 | -0.473385 | -1.313584 | 1335 | 9 | c("THY1", "NRP1", "DPYSL2", "ADGRG1", "CELSR1") |
| 27 | HALLMARK_SPERMATOGENESIS | 0.165617 | 0.306697 | 0.433011 | 1.275427 | 367 | 6 | c("SLC12A2", "GSTM3", "VDAC3", "PEBP1", "STRBP") |
| 28 | HALLMARK_APOPTOSIS | 0.191708 | 0.342335 | -0.297139 | -1.232208 | 1904 | 49 | c("HMOX1", "GPX3", "GSN", "LGALS3", "PLAT", "PDGFRB", "JUN", "EMP1", "TNFRSF12A", "ANXA1", "PPT1", "CDKN1B", "BAX", "ANKH") |
| 29 | HALLMARK_IL6_JAK_STAT3_SIGNALING | 0.264453 | 0.440755 | -0.33021 | -1.184734 | 2524 | 23 | c("A2M", "HMOX1", "CD36", "JUN", "SOCS3", "TNFRSF12A") |
| 30 | HALLMARK_ALLOGRAFT_REJECTION | 0.257681 | 0.440755 | -0.325695 | -1.190002 | 2473 | 25 | c("THY1", "INHBA", "FLNA", "STAB1", "MMP9") |
| 31 | HALLMARK_CHOLESTEROL_HOMEOSTASIS | 0.284502 | 0.458873 | -0.355537 | -1.167683 | 2613 | 16 | c("LGMN", "LGALS3", "SEMA3B", "TNFRSF12A", "ANXA5", "PLAUR") |
| 32 | HALLMARK_WNT_BETA_CATENIN_SIGNALING | 0.386297 | 0.568083 | -0.466226 | -1.066345 | 2880 | 5 | c("CSNK1E", "MYC", "NCOR2", "NCSTN", "CTNNB1") |
| 33 | HALLMARK_PEROXISOME | 0.384306 | 0.568083 | 0.184762 | 1.04053 | 190 | 22 | c("TOP2A", "CNBP", "SOD2", "ALDH9A1", "SMARCC1", "TSPO", "ECH1", "GSTK1", "LONP2", "CRABP2", "ACAA1", "FIS1") |
| 34 | HALLMARK_PANCREAS_BETA_CELLS | 0.376106 | 0.568083 | -0.470125 | -1.075262 | 2804 | 5 | c("AKT3", "MAFB") |
| 35 | HALLMARK_NOTCH_SIGNALING | 0.414063 | 0.591518 | 0.300767 | 1.013701 | 741 | 8 | c("LFNG", "NOTCH2", "APH1A", "HES1", "ARRB1", "PSENEN", "RBX1") |
| 36 | HALLMARK_FATTY_ACID_METABOLISM | 0.470717 | 0.653773 | -0.273237 | -1.008318 | 4532 | 26 | c("BMPR1B", "CD36", "S100A10", "UROD", "IDH3B", "ELOVL5", "EPHX1", "HSD17B4", "UGDH", "SMS", "MIF") |
| 37 | HALLMARK_PROTEIN_SECRETION | 0.594218 | 0.770654 | -0.233783 | -0.920291 | 5836 | 36 | c("DST", "PAM", "TPD52", "PPT1", "ANP32E", "TMED2", "MAPK1", "RAB14", "NAPA", "RER1", "M6PR", "RAB2A", "CLTA", "GOSR2", "LAMP2", "CD63", "COPB2", "LMAN1", "TOM1L1", "SEC31A", "STX7", "VPS45", "ARFGEF2", "SEC22B", "SOD1", "SNX2", "USO1", "STX16", "COPE") |
| 38 | HALLMARK_REACTIVE_OXYGEN_SPECIES_PATHWAY | 0.60111 | 0.770654 | -0.273225 | -0.897346 | 5522 | 16 | c("GPX3", "MPO") |
| 39 | HALLMARK_BILE_ACID_METABOLISM | 0.587756 | 0.770654 | -0.297065 | -0.900059 | 5193 | 12 | c("IDH2", "OPTN", "HSD17B4", "SCP2", "SLC29A1", "CAT") |
| 40 | HALLMARK_MTORC1_SIGNALING | 0.66479 | 0.830988 | -0.211426 | -0.882066 | 6613 | 51 | c("EGLN3", "LGMN", "SERPINH1", "STC1", "CANX", "DDIT4", "ELOVL5", "PIK3R3", "ACSL3", "IFI30", "HSPE1", "SLC2A1", "ACACA", "SYTL2", "ACTR3", "PSMB5", "BTG2", "PSMD13", "SERP1", "ADD3", "ITGB2", "M6PR", "TFRC", "ATP2A2", "SCD", "CCT6A", "GGA2", "LTA4H") |

| | | | | | | | |
|---|---|---|---|---|---|---|---|
| 41 | HALLMARK_INTERFERON_GAMMA_RESPONSE | 0.791395 | 0.879328 | -0.193335 | -0.765084 | 7780 | 37 c("C1S", "SERPING1", "C1R", "MT2A", "SOCS3", "IFI27", "IFI30", "ARID5B", "NFKBIA", "SRI", "BST2", "STAT2", "PIM1", "BTG1", "TNFAIP2", "RBCK1") |
| 42 | HALLMARK_UNFOLDED_PROTEIN_RESPONSE | 0.779258 | 0.879328 | -0.199602 | -0.769047 | 7603 | 32 c("SHC1", "DDIT4", "ATP6V0D1", "EIF4EBP1", "NHP2", "DCTN1", "EIF4G1", "BANF1", "DNAJA4", "SERP1", "EIF4A3", "GOSR2", "SRPRA", "ATF4", "HERPUD1", "SEC31A", "LSM1", "VEGFA", "EDEM1", "CALR") |
| 43 | HALLMARK_E2F_TARGETS | 0.736694 | 0.879328 | -0.218131 | -0.79699 | 7072 | 25 c("CDKN1B", "ANP32E", "HMGB2", "STAG1", "TUBB", "MYC", "RBBP7", "SSRP1", "NUDT21", "SMC4", "TFRC", "SRSF2", "PA2G4", "SNRPB", "POLD2", "PRKDC", "MCM7") |
| 44 | HALLMARK_MYC_TARGETS_V2 | 0.743022 | 0.879328 | -0.30708 | -0.784301 | 5935 | 7 c("HSPE1", "MYC", "GNL3", "PA2G4", "SRM") |
| 45 | HALLMARK_HEME_METABOLISM | 0.773265 | 0.879328 | -0.188629 | -0.803174 | 7710 | 59 c("C3", "ACP5", "CIR1", "SLC4A1", "ELL2", "UROD", "CAST", "PSMD9", "UBAC1", "MPP1", "TNS1", "NFE2L1") |
| 46 | HALLMARK_INTERFERON_ALPHA_RESPONSE | 0.896495 | 0.974452 | -0.190385 | -0.625276 | 8236 | 16 C1S |
| 47 | HALLMARK_MITOTIC_SPINDLE | 0.937368 | 0.976425 | -0.144417 | -0.610725 | 9338 | 56 c("GSN", "DST", "MARCKS", "FLNA", "PALLD", "SHROOM1", "ARHGAP29", "BCAR1", "FARP1", "KLC1", "NIN", "ARHGEF12", "SPTAN1", "ABR", "ARHGEF2", "TRIO", "SMC4", "WASL", "RAB3GAP1", "NUMA1", "PCM1", "RASA1", "ABL1", "CLIP1", "PREX1", "ARHGDIA", "ARF6", "ACTN4", "CKAP5", "ARFIP2", "WASF2") |
| 48 | HALLMARK_G2M_CHECKPOINT | 0.928425 | 0.976425 | -0.153922 | -0.596537 | 9079 | 33 c("MARCKS", "MT2A", "CDKN1B", "PRPF4B", "SS18", "SMAD3", "YTHDC1", "STAG1", "MYC", "SMC4", "NUMA1", "SRSF2", "ABL1", "NCL", "EWSR1", "CUL3", "RBM14", "BCL3", "CUL4A", "KIF5B", "SLC38A1") |
| 49 | HALLMARK_PI3K_AKT_MTOR_SIGNALING | 0.970588 | 0.990396 | -0.13131 | -0.505924 | 9470 | 32 c("VAV3", "CDKN1B", "PIK3R3", "SLC2A1", "SMAD2", "ACACA", "MAPK1", "ACTR3", "ARPC3", "STAT2", "PLA2G12A", "ARHGDIA", "GRB2", "CAB39", "ITPR2", "CALR") |
| 50 | HALLMARK_OXIDATIVE_PHOSPHORYLATION | 0.997698 | 0.997698 | -0.089212 | -0.388091 | 9969 | 70 c("PDK4", "MAOB", "CYB5R3", "TIMM13", "IDH3B", "ATP1B1", "IDH2", "NDUFS7", "BAX", "COX7A2L", "CYB5A", "COX4I1", "SLC25A6", "UQCR10", "ATP5PD", "NDUFB6", "UQCRQ", "ISCU", "CYC1", "COX7A2", "UQCRB", "ATP6V0C", "ATP5F1D", "HADHA", "NDUFC1", "ECHS1", "ATP5F1B", "NDUFAB1", "UQCR11", "NDUFS6", "NDUFA4", "COX7C", "NDUFS4", "RHOT2", "NDUFB8", "ATP5MG", "FH", "ATP6AP1", "NDUFB2", "NDUFA2", "ATP6V0E1", "NDUFV2", "NDUFB7", "COX6C", "NDUFS2", "NDUFA1", "NDUFB5", "OXA1L", "COX6A1", "UQCRC1", "NDUFA3", "MGST3", □ "VDAC3", "ETFB", "COX8A", "ACAA1", "PDHA1", "ACADVL", "NDUFC2", "ECH1", "LRPPRC", "ATP5F1E", "ATP6V0B", "ECI1", "ATP5MC2", "COX6B1", "ATP5MF", "ATP6V1F") |

| Cluster 16 | pathway | pval | padj | ES | NES | nMoreExtreme | size | leadingEdge |
|---|---|---|---|---|---|---|---|---|
| 1 | HALLMARK_ESTROGEN_RESPONSE_EARLY | 0.0003 | 0.005 | -0.443576 | -1.635142 | 2 | 74 | c("HSPB8", "PGR", "MAPT", "GREB1", "GJA1", "PAPSS2", "NBL1", "RHOBTB3", "CA12", "ADCY1", "TOB1", "SCNN1A", "SEC14L2", "BHLHE40", "ELF3", "LRIG1", "SLC7A2", "MED13L", "FARP1", "GFRA1", "TFF1", "FLNB", "CELSR1", "PRSS23", "PLAAT3", "SEMA3B", "CD44", "RHOD", "GAB2", "ABHD2", "SLC39A6", "FOS", "KDM4B", "STC2", "FAM102A", "THSD4", "SIAH2", "AMFR", "RAB31", "IGF1R", "CHPT1", "TTC39A", "RARA", "NCOR2", "NRIP1", "ELOVL5", "CANT1", "SLC9A3R1", "FASN", "ADD3") |
| 2 | HALLMARK_EPITHELIAL_MESENCHYMAL_TRANSITION | 1.00E-04 | 0.005 | -0.553107 | -2.098788 | 0 | 107 | c("ADAM12", "MXRA5", "FAP", "ACTA2", "SERPINE1", "CCN1", "THY1", "LOX", "FBLN2", "ITGB1", "DPYSL3", "COL5A2", "TGFBI", "SFRP4", "FBLN1", "MMP14", "LOXL2", "THBS2", "NNMT", "GJA1", "PRRX1", "JUN", "CTHRC1", "COL16A1", "COL11A1", "IGFBP3", "CDH11", "HTRA1", "CCN2", "MYL9", "GEM", "SPP1", "DAB2", "TAGLN", "LOXL1", "PCOLCE", "LAMC1", "ITGB5", "COL12A1", "LRRC15", "DST", "LRP1", "TNC", "COL5A1", "MMP2", "ITGAV", "TPM4", "ELN", "PDGFRB", "CALD1", "THBS1", "PFN2", "LUM", "TIMP3", "ITGA5", "CD44", "COPA", "CAPG", "FSTL1", "PMEPA1") |
| 3 | HALLMARK_COAGULATION | 0.0002 | 0.005 | -0.542398 | -1.887776 | 1 | 43 | c("APOC1", "SERPINE1", "MMP11", "S100A1", "MMP14", "HTRA1", "SERPING1", "CTSK", "LRP1", "ANXA1", "MMP2", "S100A13", "C1R", "THBS1", "LGMN", "CSRP1", "PRSS23", "LAMP2", "TIMP3", "CLU", "C1QA", "CD9", "C3", "CRIP2", "C1S", "CFB", "COMP", "SERPINA1", "FBN1", "GSN") |
| 4 | HALLMARK_KRAS_SIGNALING_UP | 0.001004 | 0.012545 | -0.492054 | -1.691777 | 9 | 39 | c("SPARCL1", "SERPINA3", "APOD", "MMP11", "EMP1", "GPNMB", "PRRX1", "CPE", "IGFBP3", "SPP1", "TMEM176B", "LAPTM5", "FCER1G", "MAFB", "SEMA3B", "AKT2", "NRP1", "ENG", "CFB", "FUCA1", "ITGB2", "PLAU", "ETS1", "MAP3K1", "TSPAN13") |
| 5 | HALLMARK_ESTROGEN_RESPONSE_LATE | 0.0025 | 0.022536 | -0.421845 | -1.543013 | 24 | 68 | c("HSPB8", "SERPINA3", "PGR", "CDH1", "MAPT", "COX6C", "PAPSS2", "DLG5", "PRLR", "CPE", "NBL1", "CA12", "TOB1", "SCNN1A", "UGDH", "LLGL2", "FARP1", "TFF1", "FLNB", "PRSS23", "PLAAT3", "SEMA3B", "CD44", "ABHD2", "CD9", "AGR2", "FOS", "FAM102A", "SGK1", "SERPINA1", "SIAH2", "AMFR", "RAB31", "CXCL14", "CHPT1", "IDH2", "RABEP1", "NCOR2", "TSPAN13", "NRIP1", "ELOVL5") |
| 6 | HALLMARK_UV_RESPONSE_DN | 0.002704 | 0.022536 | -0.453435 | -1.592287 | 26 | 46 | c("SERPINE1", "CCN1", "COL5A2", "SDC2", "GJA1", "EFEMP1", "COL11A1", "INPP4B", "DUSP1", "SRI", "DAB2", "ERBB2", "LAMC1", "RBPMS", "BHLHE40", "ANXA4", "PDGFRB", "PLPP3", "ANXA2", "NRP1", "PMP22", "AKT3", "TGFBR2", "IGF1R", "APBB2", "COL3A1", "NEK7", "RUNX1", "ADD3", "CITED2", "IGFBP5") |
| 7 | HALLMARK_HYPOXIA | 0.0038 | 0.027146 | -0.424137 | -1.539802 | 37 | 63 | c("SERPINE1", "CCN1", "LOX", "TPD52", "TGFBI", "SDC2", "JUN", "MIF", "IGFBP3", "CCN2", "DUSP1", "CA12", "EFNA1", "CAVIN1", "BHLHE40", "DTNA", "CCNG2", "COL5A1", "ANXA2", "MT2A", "KLF6", "P4HA1", "FOS", "HDLBP", "STC2", "PAM", "SDC4", "SIAH2", "TGFB3", "FOSL2", "ETS1", "STC1", "FBP1", "MAP3K1", "HMOX1", "BGN", "VEGFA", "CITED2", "PNRC1") |
| 8 | HALLMARK_TNFA_SIGNALING_VIA_NFKB | 0.0045 | 0.028128 | -0.420874 | -1.530037 | 44 | 64 | c("PHLDA1", "SERPINE1", "NR4A1", "CCN1", "EGR1", "FOSB", "JUN", "PLK2", "DUSP4", "DUSP1", "GEM", "MAP3K8", "EFNA1", "BHLHE40", "NFKBIA", "TNC", "EIF1", "PLPP3", "NFE2L2", "JUNB", "KLF6", "CD44", "SOCS3", "PMEPA1", "FOS", "KLF9", "SGK1", "SDC4") |
| 9 | HALLMARK_MYOGENESIS | 0.007009 | 0.035046 | -0.436477 | -1.540001 | 69 | 48 | c("ADAM12", "HSPB8", "ITGB1", "APOD", "IGFBP3", "PVALB", "GPX3", "SCD", "TAGLN", "NQO1", "ITGB5", "BHLHE40", "ERBB3", "DTNA", "SPDEF", "IGFBP7", "CLU", "AKT2", "AEBP1", "STC2", "CNN3", "MYO1C", "GSN", "ATP6AP1", "MEF2D", "COL3A1", "SYNGR2", "SPTAN1", "AGRN", "EIF4A2") |
| 10 | HALLMARK_COMPLEMENT | 0.006909 | 0.035046 | -0.43701 | -1.541884 | 68 | 48 | c("APOC1", "SERPINE1", "MMP14", "CTSD", "LIPA", "SERPING1", "CD46", "LRP1", "S100A13", "ANXA5", "C1R", "FCER1G", "LGMN", "CSRP1", "GNAI2", "LAMP2", "CLU", "LGALS3", "C1QA", "C3", "HSPA1A", "C1S", "CFB", "SERPINA1", "CTSL", "ATOX1", "C1QC", "CD55") |
| 11 | HALLMARK_IL2_STAT5_SIGNALING | 0.008211 | 0.037321 | -0.433493 | -1.529474 | 81 | 48 | c("PHLDA1", "EMP1", "S100A1", "IFITM3", "MAP3K8", "SPP1", "BHLHE40", "LRIG1", "ANXA4", "ITGAV", "RNH1", "KLF6", "CD44", "NFKBIZ", "P4HA1", "NRP1", "CAPG", "MAPKAPK2", "CD81", "AHR", "ENPP1", "SNX14", "IGF1R", "SERPINB6", "MYO1C") |
| 12 | HALLMARK_E2F_TARGETS | 0.027682 | 0.11534 | 0.307099 | 1.60778 | 7 | 20 | c("ANP32E", "PLK1", "TFRC", "SPC24", "PPP1R8", "HMGB2") |

| | | | | | | | | |
|---|---|---|---|---|---|---|---|---|
| 13 | HALLMARK_TGF_BETA_SIGNALING | 0.044309 | 0.170419 | -0.448329 | -1.445787 | 435 | 25 | c("SERPINE1", "CTNNB1", "CDH1", "SKIL", "THBS1", "JUNB", "ARID4B", "PMEPA1", "SKI", "ENG", "RAB31", "LTBP2", "ID3", "FNTA", "NCOR2", "BMPR2", "PPP1CA", "RHOA") |
| 14 | HALLMARK_GLYCOLYSIS | 0.072294 | 0.258193 | -0.379628 | -1.329327 | 721 | 45 | c("TGFBI", "SDC2", "MIF", "IGFBP3", "PPIA", "IDUA", "EGLN3", "ELF3", "COL5A1", "CD44", "P4HA1", "COPB2", "HDLBP", "QSOX1", "STC2", "PGLS", "PAM", "GFPT1", "PKM", "FUT8", "SDC1", "STC1", "AGRN", "VEGFA", "CITED2", "SOD1", "DCN", "PLOD1", "ARPP19", "CHPF", "IL13RA1", "B4GALT1") |
| 15 | HALLMARK_CHOLESTEROL_HOMEOSTASIS | 0.086087 | 0.278973 | -0.460253 | -1.381919 | 825 | 17 | c("CTNNB1", "ACTG1", "SCD", "ANXA5", "LGMN", "CLU", "LGALS3", "SEMA3B", "CD9") |
| 16 | HALLMARK_HEME_METABOLISM | 0.090909 | 0.278973 | 0.271995 | 1.91956 | 0 | 52 | c("KLF1", "SLC4A1", "TFRC", "UBAC1", "SPTB", "AHSP", "MPP1", "ANK1", "MAP2K3", "RBM38", "HBD", "ALAS2", "CCND3", "CAT", "UROD", "SNCA", "SLC25A37", "SLC2A1", "HMBS", "RAD23A") |
| 17 | HALLMARK_PEROXISOME | 0.094851 | 0.278973 | 0.278283 | 1.348469 | 34 | 18 | c("TSPO", "CAT", "SMARCC1", "FIS1", "CNBP", "ERCC1", "ALDH9A1", "TOP2A", "SCP2", "DHCR24", "SOD2", "SEMA3C", "HSD17B4", "LONP2", "SOD1", "ELOVL5", "IDH2", "CRABP2") |
| 18 | HALLMARK_ADIPOGENESIS | 0.112312 | 0.280781 | -0.355877 | -1.263939 | 1121 | 51 | c("SPARCL1", "HSPB8", "FABP4", "ALDH2", "GPX3", "TOB1", "CAVIN1", "PHLDB1", "COX6A1", "CCNG2", "CD151", "CMBL", "UBC", "CMPK1", "C3", "CYC1", "UQCR11", "DHRS7", "LAMA4", "GPAT4", "UQCRQ", "VEGFB") |
| 19 | HALLMARK_XENOBIOTIC_METABOLISM | 0.107945 | 0.280781 | -0.371126 | -1.284075 | 1075 | 41 | c("SERPINE1", "FBLN1", "PAPSS2", "ALDH2", "NQO1", "UGDH", "PDK4", "TMEM176B", "BCAR1", "MT2A", "ESR1", "CFB", "DHRS7", "ATP2A2", "NINJ1", "FBP1", "ELOVL5", "HMOX1", "EPHX1", "SPINT2", "PGRMC1", "IL1R1", "CD36", "ARPP19", "JUP") |
| 20 | HALLMARK_PANCREAS_BETA_CELLS | 0.10288 | 0.280781 | -0.628199 | -1.357446 | 810 | 5 | c("MAFB", "SPCS1", "AKT3", "SRP9", "SEC11A") |
| 21 | HALLMARK_WNT_BETA_CATENIN_SIGNALING | 0.174299 | 0.391676 | -0.582864 | -1.259484 | 1373 | 5 | c("CTNNB1", "CSNK1E", "NCSTN", "NCOR2") |
| 22 | HALLMARK_BILE_ACID_METABOLISM | 0.17374 | 0.391676 | 0.311682 | 1.239723 | 130 | 12 | c("KLF1", "RXRA", "CAT", "OPTN", "SLC29A1", "ALDH9A1", "SCP2", "DHCR24", "HSD17B4", "LONP2") |
| 23 | HALLMARK_KRAS_SIGNALING_DN | 0.180171 | 0.391676 | -0.474461 | -1.258273 | 1620 | 10 | c("CPB1", "IDUA", "EFHD1", "SKIL") |
| 24 | HALLMARK_INTERFERON_GAMMA_RESPONSE | 0.192312 | 0.400649 | -0.359221 | -1.211791 | 1910 | 34 | c("SRI", "IFITM3", "SERPING1", "NFKBIA", "IFI30", "IFI27", "STAT1", "IFITM2", "C1R", "MT2A", "B2M", "SOCS3", "C1S", "CFB", "SAMHD1", "ARID5B") |
| 25 | HALLMARK_APICAL_JUNCTION | 0.211727 | 0.41196 | -0.326158 | -1.16689 | 2115 | 55 | c("THY1", "ITGB1", "TGFBI", "CDH1", "COL16A1", "CDH11", "MYL9", "ACTG1", "LIMA1", "MMP2", "CD276", "EVL", "GNAI2", "PARVA", "AKT2", "THBS3", "CLDN4", "PCDH1", "AKT3", "EXOC4", "FBN1", "WASL", "CD99", "CTNND1", "SHC1", "SYMPK", "TSPAN4", "RASA1", "PECAM1", "JUP", "CLDN7", "MSN", "B4GALT1", "NECTIN2") |
| 26 | HALLMARK_SPERMATOGENESIS | 0.214219 | 0.41196 | -0.599975 | -1.211698 | 1611 | 4 | c("SLC12A2", "GSTM3") |
| 27 | HALLMARK_INFLAMMATORY_RESPONSE | 0.238081 | 0.44089 | -0.370354 | -1.184908 | 2336 | 24 | c("SERPINE1", "MMP14", "SRI", "NFKBIA", "STAB1", "SLC7A2", "ITGA5", "KLF6", "AHR", "CD55", "CD14", "ATP2A2") |
| 28 | HALLMARK_PI3K_AKT_MTOR_SIGNALING | 0.290076 | 0.500132 | 0.179695 | 1.085939 | 37 | 28 | c("ARPC3", "MAP2K3", "SLC2A1", "CXCR4", "PIK3R3", "CDKN1B", "PLA2G12A", "SQSTM1", "PAK4", "RAC1", "AKT1", "STAT2", "GRK2", "VAV3", "ARHGDIA", "ACTR3", "SMAD2", "GRB2", "UBE2D3", "GSK3B", "CALR", "PPP1CA", "ACACA", "MAPK1", "CFL1", "CAB39", "ARF1", "DUSP3") |
| 29 | HALLMARK_MYC_TARGETS_V2 | 0.28618 | 0.500132 | 0.366148 | 1.118387 | 438 | 7 | c("PLK1", "NOP56", "HSPE1", "MYC", "CBX3", "SRM", "GNL3") |
| 30 | HALLMARK_APICAL_SURFACE | 0.300981 | 0.501635 | -0.457134 | -1.137277 | 2607 | 8 | c("THY1", "GSTM3", "SULF2") |
| 31 | HALLMARK_PROTEIN_SECRETION | 0.344425 | 0.555525 | -0.321932 | -1.094813 | 3425 | 36 | c("TPD52", "VPS45", "TMED2", "DST", "USO1", "PPT1", "RAB2A", "LAMP2", "ARF1", "COPB2", "SNX2", "PAM", "NAPA", "SEC31A") |
| 32 | HALLMARK_ANDROGEN_RESPONSE | 0.382427 | 0.597542 | -0.313081 | -1.06825 | 3803 | 37 | c("TPD52", "INPP4B", "SCD", "SPDEF", "ITGAV", "B2M", "ACSL3", "ABHD2", "PMEPA1", "STEAP4", "SGK1", "SELENOP", "ARID5B", "SMS", "UAP1", "ELOVL5", "BMPR1B") |
| 33 | HALLMARK_HEDGEHOG_SIGNALING | 0.396713 | 0.60108 | -0.410505 | -1.059125 | 3523 | 9 | c("THY1", "CELSR1", "ADGRG1", "NRP1") |
| 34 | HALLMARK_UNFOLDED_PROTEIN_RESPONSE | 0.44179 | 0.631129 | -0.309203 | -1.034934 | 4382 | 32 | c("ATF4", "EIF4G1", "YWHAZ", "SPCS1", "STC2", "FUS", "SEC31A", "EIF4A3", "EIF4EBP1", "SHC1", "SEC11A", "EIF4A2", "HERPUD1", "VEGFA", "LSM1", "ATP6V0D1", "KIF5B") |
| 35 | HALLMARK_UV_RESPONSE_UP | 0.440032 | 0.631129 | -0.302102 | -1.030789 | 4376 | 37 | c("NR4A1", "MMP14", "FOSB", "GPX3", "NFKBIA", "CREG1", "PPT1", "ATP6V1F", "JUNB", "HNRNPU") |

| | | | | | | | | |
|---|---|---|---|---|---|---|---|---|
| 36 | HALLMARK_APOPTOSIS | 0.454427 | 0.631149 | -0.28462 | -1.022064 | 4541 | 57 | c("EMP1", "CTNNB1", "JUN", "GPX3", "IFITM3", "ERBB2", "ERBB3", "ANXA1", "MMP2", "PDGFRB", "PPT1", "LUM", "TIMP3", "CLU", "LGALS3", "CD44", "CREBBP", "GSN", "CD14", "RARA", "TXNIP", "HMOX1", "SPTAN1", "BGN", "SOD1", "PEA15", "DCN", "PLAT") |
| 37 | HALLMARK_IL6_JAK_STAT3_SIGNALING | 0.503621 | 0.639834 | -0.323827 | -0.991908 | 4867 | 19 | c("JUN", "MAP3K8", "STAT1", "CD44", "SOCS3", "CD9", "CD14", "HMOX1", "STAT3", "IL1R1", "CD36", "IL13RA1", "A2M", "IFNGR2", "TNFRSF12A", "GRB2") |
| 38 | HALLMARK_NOTCH_SIGNALING | 0.524664 | 0.639834 | 0.286935 | 0.938726 | 701 | 8 | c("CCND1", "NOTCH2", "ARRB1", "HES1", "PSENEN", "LFNG", "APH1A", "NOTCH3") |
| 39 | HALLMARK_MYC_TARGETS_V1 | 0.515224 | 0.639834 | -0.284036 | -0.991498 | 5143 | 44 | c("PPIA", "SNRPD2", "YWHAE", "HSP90AB1", "CANX", "HNRNPC", "NME1", "CCT3", "PSMD3", "CLNS1A", "HNRNPU", "IMPDH2") |
| 40 | HALLMARK_P53_PATHWAY | 0.505308 | 0.639834 | -0.283955 | -0.997138 | 5044 | 46 | c("S100A10", "TM4SF1", "JUN", "PLK2", "CTSD", "TOB1", "IFI30", "SEC61A1", "NUPR1", "MDM2", "FOS", "CD81", "EPS8L2", "ZFP36L1", "FUCA1", "SDC1") |
| 41 | HALLMARK_ALLOGRAFT_REJECTION | 0.474531 | 0.639834 | -0.316628 | -1.013017 | 4657 | 24 | c("THY1", "TPD52", "STAB1", "STAT1", "NME1", "B2M", "CAPG") |
| 42 | HALLMARK_ANGIOGENESIS | 0.544554 | 0.648279 | -0.318778 | -0.957137 | 5224 | 17 | c("COL5A2", "SPP1", "ITGAV", "LUM", "NRP1", "FSTL1", "STC1", "COL3A1", "POSTN", "VEGFA") |
| 43 | HALLMARK_MTORC1_SIGNALING | 0.648784 | 0.7544 | -0.257221 | -0.915311 | 6481 | 52 | c("PPIA", "EGLN3", "SCD", "BHLHE40", "IFI30", "CANX", "USO1", "NUPR1", "LGMN", "ACSL3", "P4HA1", "CD9", "ITGB2", "ATP2A2", "PSMB5", "ACACA", "STC1", "ELOVL5", "SEC11A", "SERPINH1", "SLC9A3R1", "SYTL2", "ADD3", "IGFBP5") |
| 44 | HALLMARK_INTERFERON_ALPHA_RESPONSE | 0.786882 | 0.894184 | -0.2533 | -0.775879 | 7605 | 19 | c("IFITM3", "IFI30", "IFI27", "IFITM2", "B2M", "C1S") |
| 45 | HALLMARK_FATTY_ACID_METABOLISM | 0.860258 | 0.955842 | -0.220995 | -0.72175 | 8482 | 27 | c("S100A10", "MIF", "UGDH", "FH", "ECI1", "SMS", "ELOVL5", "BMPR1B", "EPHX1", "FASN", "CD36") |
| 46 | HALLMARK_MITOTIC_SPINDLE | 0.89934 | 0.976601 | -0.203956 | -0.731005 | 8987 | 56 | c("SHROOM1", "ARHGAP29", "PALLD", "YWHAE", "DST", "NET1", "FARP1", "DYNLL2", "FLNB", "BCAR1", "CAPZB", "EZR", "PCM1", "WASL", "ARHGEF12", "GSN", "ABL1") |
| 47 | HALLMARK_REACTIVE_OXYGEN_SPECIES_PATHWAY | 0.918005 | 0.976601 | -0.216855 | -0.632639 | 8687 | 15 | c("GPX3", "NQO1", "JUNB", "ATOX1", "PTPA", "SOD1") |
| 48 | HALLMARK_DNA_REPAIR | 0.991693 | 0.997082 | -0.133734 | -0.439362 | 9788 | 28 | c("TMED2", "SEC61A1", "NME1", "IMPDH2") |
| 49 | HALLMARK_G2M_CHECKPOINT | 0.997082 | 0.997082 | -0.119869 | -0.404363 | 9907 | 34 | c("SLC38A1", "SLC12A2", "MT2A", "HNRNPU", "ABL1", "HNRNPD", "NCL", "CUL3", "HSPA8", "SS18", "KIF5B", "PRPF4B") |
| 50 | HALLMARK_OXIDATIVE_PHOSPHORYLATION | 0.9663 | 0.997082 | -0.174561 | -0.63554 | 9662 | 65 | c("COX6C", "ATP1B1", "PDK4", "COX6A1", "ATP6V1F", "ATP6V0E1", "MAOB", "CYC1", "UQCR11", "FH", "IDH2", "ATP6AP1", "UQCRQ", "COX4I1", "CYB5R3", "ECI1", "ATP6V0C", "COX7C", "HADHA", "TIMM13", "ATP5MF", "NDUFB2", "UQCR10", "GLUD1", "ATP5MC2", "NDUFB8", "NDUFC2", "ATP5F1E", "NDUFAB1", "UQCRB", "TCIRG1", "BAX", "NDUFA4", "ATP5F1B", "NDUFB7", "ISCU", "NDUFA3") |

| Cluster 17 | pathway | pval | padj | ES | NES | nMoreExtreme | size | leadingEdge |
|---|---|---|---|---|---|---|---|---|
| 1 | HALLMARK_EPITHELIAL_MESENCHYMAL_TRANSITION | 0.0001 | 0.002536 | -0.767804 | -2.073165 | 0 | 105 | c("COL5A1", "COL11A1", "CDH11", "ELN", "GJA1", "THY1", "COL16A1", "FBLN2", "MMP2", "FBN1", "CCN2", "COL4A2", "COL12A1", "FAP", "COMP", "COL5A2", "ACTA2", "CCN1", "FSTL1", "PRRX1", "INHBA", "SFRP4", "COL4A1", "BGN", "DPYSL3", "THBS2", "LUM", "LAMC1", "POSTN", "DCN", "LRRC15", "GEM", "PDGFRB", "LOX", "CALD1", "MMP14", "NNMT", "ADAM12", "LOXL2", "HTRA1", "CTHRC1", "IGFBP3", "SERPINE1", "CXCL12", "LOXL1", "TAGLN", "TPM2", "COL1A1", "COL3A1", "PCOLCE", "COL1A2", "COL6A3", "COL6A2", "DAB2", "DST", "TIMP3", "FBLN1", "FN1", "PMEPA1", "MYL9", "MXRA5", "LRP1", "SPARC", "TPM1", "JUN", "TNC", "FUCA1", "PMP22", "PFN2", "TGFBI", "SPP1", "TNFRSF12A", "SDC1", "TIMP1", "SERPINH1", "IGFBP4", "PLOD1", "ITGA5") |
| 2 | HALLMARK_COAGULATION | 0.000101 | 0.002536 | -0.745854 | -1.918517 | 0 | 41 | c("C1S", "MMP2", "FBN1", "COMP", "A2M", "CTSK", "MMP14", "MMP11", "HTRA1", "C3", "SERPINE1", "C1R", "PLAU", "APOC1", "SERPING1", "S100A1", "C1QA", "GSN", "CRIP2", "TIMP3", "LGMN", "FN1", "MMP9", "LRP1", "SPARC", "CAPN2", "CTSB", "PRSS23", "TIMP1") |
| 3 | HALLMARK_UV_RESPONSE_DN | 0.000203 | 0.003385 | -0.675507 | -1.734519 | 1 | 40 | c("COL11A1", "GJA1", "COL5A2", "CCN1", "LAMC1", "PDGFRB", "SERPINE1", "EFEMP1", "COL1A1", "NRP1", "COL3A1", "AKT3", "COL1A2", "PLPP3", "DAB2", "FHL2", "ANXA4", "SDC2", "CDKN1B", "RBPMS", "PMP22", "RUNX1") |
| 4 | HALLMARK_MYOGENESIS | 0.000305 | 0.003808 | -0.656706 | -1.686244 | 2 | 40 | c("AEBP1", "IGFBP7", "COL4A2", "APOD", "ADAM12", "IGFBP3", "TAGLN", "TPM2", "COL1A1", "PVALB", "COL3A1", "COL6A3", "COL6A2", "GPX3", "GSN", "SORBS3", "SPARC", "CD36", "HSPB8", "CNN3", "PDLIM7") |
| 5 | HALLMARK_COMPLEMENT | 0.001008 | 0.010077 | -0.610425 | -1.589205 | 9 | 49 | c("C1S", "COL4A2", "MMP14", "C3", "SERPINE1", "C1R", "APOC1", "SERPING1", "LTF", "C1QA", "C1QC", "CTSL", "LGMN", "FN1", "LIPA", "LRP1", "CTSD", "TIMP2", "CTSB", "CD36", "LGALS3") |
| 6 | HALLMARK_SPERMATOGENESIS | 0.001687 | 0.014062 | 0.864084 | 1.899952 | 4 | 4 | c("SLC12A2", "GSTM3", "HSPA2") |
| 7 | HALLMARK_KRAS_SIGNALING_UP | 0.002649 | 0.018923 | -0.617162 | -1.578182 | 25 | 38 | c("PRRX1", "INHBA", "SPARCL1", "APOD", "MMP11", "IGFBP3", "PLAU", "GPNMB", "NRP1", "TMEM176B", "SERPINA3", "MMP9", "EMP1", "MAFB", "ENG", "CPE", "FUCA1", "SPP1", "ANKH", "PLAT") |
| 8 | HALLMARK_APOPTOSIS | 0.012237 | 0.067984 | -0.564165 | -1.459489 | 120 | 44 | c("MMP2", "BGN", "LUM", "DCN", "PDGFRB", "HMOX1", "GPX3", "GSN", "TIMP3", "EMP1", "TIMP2", "CDKN1B", "JUN", "LGALS3", "TNFRSF12A", "TIMP1", "ANKH", "PLAT", "BTG2") |
| 9 | HALLMARK_ANGIOGENESIS | 0.011385 | 0.067984 | -0.69678 | -1.569123 | 101 | 14 | c("COL5A2", "FSTL1", "LUM", "POSTN", "NRP1", "COL3A1", "SPP1", "TIMP1", "ITGAV") |
| 10 | HALLMARK_HYPOXIA | 0.014766 | 0.073832 | -0.536959 | -1.407476 | 146 | 56 | c("COL5A1", "CCN2", "CCN1", "BGN", "CAVIN1", "DCN", "LOX", "IGFBP3", "SERPINE1", "HMOX1", "PAM", "SDC2", "FOS", "TGFB3", "CDKN1B", "JUN", "TGFBI", "SDC3", "DDIT4", "SLC2A1", "ANXA2", "HEXA", "ZFP36", "MT2A", "PFKFB3", "FOSL2") |
| 11 | HALLMARK_HEME_METABOLISM | 0.017544 | 0.079745 | 0.380696 | 2.032314 | 0 | 52 | c("HBB", "ALAS2", "TENT5C", "EPB41", "TRIM58", "HEBP1", "SNCA", "GYPC", "SLC25A37", "BNIP3L", "HBD", "ADIPOR1", "FECH", "RIOK3", "RBM38", "FBXO7", "MBOAT2", "BPGM", "MAP2K3", "USP15", "MXI1") |
| 12 | HALLMARK_MYC_TARGETS_V2 | 0.019237 | 0.080155 | 0.77655 | 1.707481 | 56 | 4 | c("SLC19A1", "WDR74", "MCM4") |
| 13 | HALLMARK_APICAL_JUNCTION | 0.027972 | 0.107585 | -0.548145 | -1.412473 | 275 | 42 | c("CDH11", "THY1", "COL16A1", "MMP2", "FBN1", "AKT3", "SORBS3", "MDK", "MMP9", "MYL9", "CD276", "CDH1", "CERCAM", "TGFBI", "SDC3", "CD99", "PARVA") |
| 14 | HALLMARK_TNFA_SIGNALING_VIA_NFKB | 0.054931 | 0.183102 | -0.498926 | -1.31108 | 546 | 59 | c("CCN1", "INHBA", "GEM", "SERPINE1", "PLAU", "FOSB", "NR4A1", "PHLDA1", "PLPP3", "EGR1", "KLF9", "MARCKS", "PMEPA1", "MAP3K8", "SGK1", "FOS", "JUN", "TNC", "BTG2", "SOCS3", "ID2", "DUSP4", "SQSTM1", "ZFP36", "PFKFB3", "FOSL2", "B4GALT1", "DUSP1", "JUNB", "PLAUR") |
| 15 | HALLMARK_ESTROGEN_RESPONSE_LATE | 0.054127 | 0.183102 | -0.500134 | -1.313164 | 538 | 58 | c("CXCL14", "NBL1", "CXCL12", "PGR", "PAPSS2", "LTF", "SERPINA3", "MAPT", "MDK", "CDH1", "SGK1", "ABHD2", "FOS", "SCNN1A", "CPE", "PRSS23", "NRIP1", "HSPB8", "IGFBP4", "ELOVL5", "PRLR", "ID2", "RAB31", "SERPINA1", "MYOF", "SEMA3B", "ATP2B4", "ZFP36", "FARP1", "IDH2", "FAM102A", "CA12", "SCUBE2", "BCL2") |

| | | | | | | | | |
|---|---|---|---|---|---|---|---|---|
| 16 | HALLMARK_ESTROGEN_RESPONSE_EARLY | 0.077896 | 0.243425 | -0.485699 | -1.278466 | 775 | 60 | c("GJA1", "ADCY1", "NBL1", "CXCL12", "PGR", "SEC14L2", "PAPSS2", "MAPT", "FHL2", "RHOBTB3", "RHOD", "ABHD2", "FOS", "SCNN1A", "SLC7A2", "PRSS23", "NRIP1", "HSPB8", "IGFBP4", "ELOVL5", "GAB2", "IGF1R", "SLC2A1", "RAB31", "GREB1", "MYOF", "SEMA3B", "MAST4", "B4GALT1") |
| 17 | HALLMARK_ADIPOGENESIS | 0.0892 | 0.262353 | -0.517667 | -1.317639 | 872 | 36 | c("FABP4", "COL4A1", "CAVIN1", "SPARCL1", "LAMA4", "C3", "PHLDB1", "APOE", "GPX3", "CD36", "HSPB8", "STOM") |
| 18 | HALLMARK_PEROXISOME | 0.120161 | 0.333781 | 0.386396 | 1.350272 | 148 | 12 | c("ALB", "PRDX5", "TOP2A", "ABCC5", "ACSL1") |
| 19 | HALLMARK_DNA_REPAIR | 0.164404 | 0.396726 | 0.311126 | 1.260643 | 108 | 19 | c("DDB1", "ARL6IP1", "POLR2G", "EIF1B", "POLD4", "BCAM", "NME3", "SEC61A1", "POLR2A", "NCBP2", "DCTN4", "SUPT5H", "APRT", "TMED2", "ERCC1", "NUDT21", "POLR1D") |
| 20 | HALLMARK_HEDGEHOG_SIGNALING | 0.166625 | 0.396726 | -0.651236 | -1.276762 | 1330 | 7 | c("THY1", "NRP1", "DPYSL2") |
| 21 | HALLMARK_PANCREAS_BETA_CELLS | 0.157702 | 0.396726 | -0.780986 | -1.268455 | 1031 | 3 | c("AKT3", "MAFB") |
| 22 | HALLMARK_TGF_BETA_SIGNALING | 0.176339 | 0.400771 | -0.524427 | -1.259596 | 1658 | 21 | c("LTBP2", "SERPINE1", "ID3", "PMEPA1", "CDH1", "ENG", "ID2", "FNTA", "RAB31", "THBS1", "JUNB", "SKIL", "BMPR2") |
| 23 | HALLMARK_MYC_TARGETS_V1 | 0.186263 | 0.40492 | 0.323509 | 1.232667 | 159 | 16 | c("BUB3", "RRM1", "MCM4", "APEX1", "PSMA7", "RACK1", "EIF4H", "HDGF", "NCBP2", "HNRNPA3", "MYC", "VDAC3", "NHP2", "EEF1B2", "LSM7", "CANX") |
| 24 | HALLMARK_XENOBIOTIC_METABOLISM | 0.209224 | 0.418447 | -0.476493 | -1.204032 | 2036 | 33 | c("SERPINE1", "HMOX1", "PDK4", "PAPSS2", "TMEM176B", "APOE", "FBLN1", "CD36", "IGFBP4", "ELOVL5", "CYB5A", "ID2", "CAT", "BCAR1", "MT2A", "NQO1", "DDT", "MAN1A1") |
| 25 | HALLMARK_GLYCOLYSIS | 0.20547 | 0.418447 | -0.4669 | -1.196698 | 2020 | 39 | c("COL5A1", "DCN", "EGLN3", "IGFBP3", "PAM", "SDC2", "CHPF", "TGFBI", "SDC3", "SDC1", "PLOD1", "DDIT4", "CYB5A") |
| 26 | HALLMARK_G2M_CHECKPOINT | 0.319005 | 0.61347 | 0.245898 | 1.092596 | 140 | 25 | c("SLC12A2", "INCENP", "TOP2A", "BUB3", "LMNB1", "SLC38A1", "HSPA8", "CCND1", "CUL3") |
| 27 | HALLMARK_ANDROGEN_RESPONSE | 0.410869 | 0.708394 | -0.424461 | -1.067621 | 3991 | 31 | c("BMPR1B", "MAF", "PMEPA1", "SGK1", "ABHD2", "SELENOP", "ANKH", "ELOVL5", "ITGAV", "ELL2", "STEAP4", "ARID5B", "B4GALT1") |
| 28 | HALLMARK_REACTIVE_OXYGEN_SPECIES_PATHWAY | 0.400473 | 0.708394 | -0.484715 | -1.077815 | 3552 | 13 | c("MPO", "GPX3", "CAT", "NQO1", "JUNB", "GPX4", "FTL") |
| 29 | HALLMARK_IL2_STAT5_SIGNALING | 0.390812 | 0.708394 | -0.415528 | -1.072892 | 3861 | 43 | c("NRP1", "S100A1", "PHLDA1", "COL6A1", "ANXA4", "EMP1", "MAP3K8", "SPP1", "CTSZ", "ITGA6", "IGF1R", "ITGAV", "MAPKAPK2", "GPX4", "NFKBIZ", "BMPR2", "BCL2", "CAPG", "AHR", "PIM1", "CKAP4", "CD44", "PLEC", "KLF6", "BHLHE40", "CD81") |
| 30 | HALLMARK_P53_PATHWAY | 0.427814 | 0.713023 | -0.412343 | -1.052863 | 4195 | 37 | c("HMOX1", "CTSD", "S100A10", "FOS", "JUN", "FUCA1", "RAP2B", "SDC1", "STOM", "BTG2", "DDIT4", "ZFP36L1", "IFI30", "NUPR1", "PLXNB2", "PLK2", "TAX1BP3", "IER5", "ISCU", "CD81") |
| 31 | HALLMARK_IL6_JAK_STAT3_SIGNALING | 0.458155 | 0.717571 | -0.434381 | -1.037028 | 4291 | 20 | c("A2M", "HMOX1", "MAP3K8", "JUN", "CD36", "TNFRSF12A", "SOCS3") |
| 32 | HALLMARK_BILE_ACID_METABOLISM | 0.459246 | 0.717571 | 0.316354 | 0.972617 | 754 | 9 | c("AQP9", "PRDX5", "ACSL1", "GSTK1", "HSD17B4") |
| 33 | HALLMARK_APICAL_SURFACE | 0.521157 | 0.74825 | -0.505356 | -0.990761 | 4162 | 7 | c("THY1", "SULF2", "B4GALT1", "PLAUR") |
| 34 | HALLMARK_UV_RESPONSE_UP | 0.523775 | 0.74825 | -0.39595 | -0.995909 | 5088 | 31 | c("MMP14", "HMOX1", "FOSB", "NR4A1", "GPX3", "FOS", "UROD", "BTG2", "SQSTM1") |
| 35 | HALLMARK_ALLOGRAFT_REJECTION | 0.523145 | 0.74825 | -0.409788 | -0.990231 | 4949 | 22 | c("THY1", "INHBA", "MMP9", "STAB1", "TIMP1") |
| 36 | HALLMARK_FATTY_ACID_METABOLISM | 0.616815 | 0.856688 | -0.383135 | -0.920234 | 5802 | 21 | c("BMPR1B", "S100A10", "CD36", "UROD", "ELOVL5") |
| 37 | HALLMARK_NOTCH_SIGNALING | 0.693302 | 0.888925 | -0.563673 | -0.848774 | 4129 | 2 | NOTCH3 |
| 38 | HALLMARK_E2F_TARGETS | 0.693361 | 0.888925 | 0.219022 | 0.823365 | 657 | 15 | c("TOP2A", "LMNB1", "ASF1B", "MCM4", "SNRPB") |
| 39 | HALLMARK_INFLAMMATORY_RESPONSE | 0.660457 | 0.888925 | -0.360236 | -0.898268 | 6381 | 29 | c("INHBA", "MMP14", "SERPINE1", "STAB1", "SLC7A2", "TIMP1", "ITGA5", "BTG2") |
| 40 | HALLMARK_UNFOLDED_PROTEIN_RESPONSE | 0.737724 | 0.922155 | -0.34315 | -0.819227 | 6910 | 20 | c("DDIT4", "SHC1", "HERPUD1", "DNAJA4", "STC2", "SRPRA", "VEGFA", "XBP1", "BANF1", "ATF4", "SERP1", "DCTN1", "NHP2", "EIF4EBP1", "EDEM1", "SEC31A", "ATP6V0D1") |
| 41 | HALLMARK_CHOLESTEROL_HOMEOSTASIS | 0.810118 | 0.960221 | -0.32558 | -0.742979 | 7333 | 15 | c("LGMN", "LGALS3", "TNFRSF12A", "SEMA3B", "PLAUR", "ANXA5", "ACTG1", "CD9") |
| 42 | HALLMARK_INTERFERON_ALPHA_RESPONSE | 0.82579 | 0.960221 | -0.31094 | -0.725793 | 7607 | 17 | C1S |
| 43 | HALLMARK_KRAS_SIGNALING_DN | 0.813015 | 0.960221 | -0.349816 | -0.740663 | 6908 | 10 | c("BMPR1B", "SGK1", "SELENOP", "BTG2") |
| 44 | HALLMARK_WNT_BETA_CATENIN_SIGNALING | 0.919315 | 0.98448 | -0.389266 | -0.632235 | 6015 | 3 | c("CSNK1E", "NCOR2", "MYC") |
| 45 | HALLMARK_PROTEIN_SECRETION | 0.889624 | 0.98448 | -0.274571 | -0.655502 | 8333 | 20 | c("PAM", "DST", "ANP32E", "PPT1", "M6PR", "LMAN1", "TPD52", "TMED2", "ARCN1") |
| 46 | HALLMARK_INTERFERON_GAMMA_RESPONSE | 0.894311 | 0.98448 | -0.270436 | -0.687481 | 8740 | 35 | c("C1S", "C1R", "SERPING1", "IFI27", "SOCS3", "CD74", "ARID5B", "IFI30", "MT2A") |

| | | | | | | | | |
|---|---|---|---|---|---|---|---|---|
| 47 | HALLMARK_MTORC1_SIGNALING | 0.925411 | 0.98448 | -0.251825 | -0.640981 | 9056 | 36 | c("EGLN3", "LGMN", "SERPINH1", "ELOVL5", "BTG2", "DDIT4", "SLC2A1", "SQSTM1", "IFI30", "NUPR1", "CANX", "M6PR", "CD9", "TFRC", "BHLHE40", "SYTL2", "ADD3") |
| 48 | HALLMARK_MITOTIC_SPINDLE | 0.97987 | 0.997942 | -0.193773 | -0.496654 | 9637 | 39 | c("GSN", "DST", "PALLD", "MARCKS", "BCAR1", "FLNA", "PCM1", "FARP1", "ARHGAP29", "SHROOM1") |
| 49 | HALLMARK_PI3K_AKT_MTOR_SIGNALING | 0.986428 | 0.997942 | -0.184475 | -0.434522 | 9157 | 18 | c("CDKN1B", "VAV3", "SLC2A1", "SQSTM1", "DUSP3") |
| 50 | HALLMARK_OXIDATIVE_PHOSPHORYLATION | 0.997942 | 0.997942 | -0.146884 | -0.369448 | 9695 | 31 | c("PDK4", "MAOB", "CYB5A", "ATP1B1", "IDH2", "CYB5R3", "GPX4", "NDUFS7", "SLC25A6", "ISCU", "TIMM13") |

| Cluster 18 | pathway | pval | padj | ES | NES | nMoreExtreme | size | leadingEdge |
|---|---|---|---|---|---|---|---|---|
| 1 | HALLMARK_TNFA_SIGNALING_VIA_NFKB | 0.000747 | 0.008375 | -0.488171 | -2.804461 | 0 | 61 | c("NR4A1", "PHLDA1", "FOSB", "DUSP1", "GEM", "EGR1", "BTG1", "CCN1", "SAT1", "FOS", "MAP3K8", "BTG2", "SGK1", "KLF2", "KLF6", "JUN", "PLPP3", "ZFP36", "MAP2K3", "MYC", "SERPINE1", "DUSP4", "SQSTM1", "CEBPB") |
| 2 | HALLMARK_ESTROGEN_RESPONSE_LATE | 0.000838 | 0.008375 | -0.330817 | -1.955778 | 0 | 71 | c("S100A9", "TFF1", "TFF3", "SERPINA1", "HSPB8", "SERPINA3", "CXCL14", "FOS", "BLVRB", "AGR2", "PRLR", "CPE", "SGK1", "CDH1", "AMFR", "ZFP36") |
| 3 | HALLMARK_COMPLEMENT | 0.000654 | 0.008375 | -0.490438 | -2.612782 | 0 | 46 | c("S100A9", "CD36", "SERPINA1", "APOC1", "CTSL", "CTSS", "LGALS3", "PLAT", "LGMN", "CLU", "CTSB", "PIM1", "SERPINE1", "PFN1", "LTA4H", "CTSD", "CEBPB") |
| 4 | HALLMARK_HEME_METABOLISM | 0.000688 | 0.008375 | -0.483295 | -2.640437 | 0 | 51 | c("HBB", "HBD", "SLC4A1", "SNCA", "ACP5", "TFRC", "MPP1", "CAT", "BLVRB", "BTG2", "SLC25A37", "GYPC", "CTSB", "MAP2K3") |
| 5 | HALLMARK_PANCREAS_BETA_CELLS | 0.000287 | 0.008375 | 0.893701 | 2.029126 | 1 | 5 | c("PCSK1", "INSM1") |
| 6 | HALLMARK_HYPOXIA | 0.001428 | 0.008922 | -0.351072 | -1.990165 | 1 | 57 | c("STC2", "HMOX1", "DUSP1", "BTG1", "CCN1", "CXCR4", "FOS", "CCN2", "KLF6", "PIM1", "JUN", "ZFP36", "SERPINE1", "CITED2", "BGN", "CDKN1B", "CCNG2", "IDS") |
| 7 | HALLMARK_REACTIVE_OXYGEN_SPECIES_PATHWAY | 0.001408 | 0.008922 | -0.551239 | -2.164567 | 2 | 17 | c("MPO", "NQO1", "FTL", "CAT", "GPX3", "LSP1") |
| 8 | HALLMARK_KRAS_SIGNALING_UP | 0.001199 | 0.008922 | -0.396289 | -2.01165 | 1 | 38 | c("MMP9", "SERPINA3", "CTSS", "GPNMB", "PLAT", "SPARCL1", "CXCR4", "CPE", "GYPC", "LCP1", "IKZF1", "PECAM1", "ITGB2", "EMP1") |
| 9 | HALLMARK_IL6_JAK_STAT3_SIGNALING | 0.003605 | 0.018023 | -0.475074 | -2.062359 | 6 | 23 | c("HMOX1", "CD36", "MAP3K8", "PIM1", "JUN", "A2M", "IL1R1", "IFNGR2", "IFNGR1", "IL13RA1", "IRF9", "STAT2", "STAT1", "IFNAR1", "PTPN1", "STAT3") |
| 10 | HALLMARK_ADIPOGENESIS | 0.003273 | 0.018023 | -0.300147 | -1.76549 | 3 | 69 | c("FABP4", "CD36", "APOE", "HSPB8", "SPARCL1", "CAT", "UQCRQ", "TALDO1", "GPX3", "UBC", "UQCR10", "MGST3", "ALDH2", "ECH1", "CCNG2", "C3", "SCP2", "DHRS7", "COL15A1", "MYLK", "REEP5", "RMDN3") |
| 11 | HALLMARK_MYOGENESIS | 0.004019 | 0.018267 | -0.344449 | -1.866516 | 5 | 49 | c("PVALB", "STC2", "NQO1", "CD36", "HSPB8", "SCD", "GPX3", "CLU", "LSP1") |
| 12 | HALLMARK_XENOBIOTIC_METABOLISM | 0.004743 | 0.019761 | -0.340323 | -1.851994 | 6 | 50 | c("PDK4", "HMOX1", "NQO1", "CD36", "APOE", "CAT", "BLVRB", "EPHX1", "SERPINE1", "IL1R1", "UGDH", "ALDH2", "ECH1") |
| 13 | HALLMARK_APOPTOSIS | 0.011348 | 0.037825 | -0.30105 | -1.693988 | 15 | 56 | c("HMOX1", "LGALS3", "PLAT", "SAT1", "BTG2", "GPX3", "ANXA1", "CLU", "JUN", "TXNIP", "EMP1", "SQSTM1", "MMP2", "BGN", "IFITM3", "CDKN1B", "APP", "TIMP2") |
| 14 | HALLMARK_UV_RESPONSE_UP | 0.011046 | 0.037825 | -0.331429 | -1.761675 | 16 | 45 | c("HMOX1", "NR4A1", "FOSB", "BTG1", "TFRC", "FOS", "BTG2", "GPX3", "EPHX1", "ARRB2") |
| 15 | HALLMARK_COAGULATION | 0.011196 | 0.037825 | -0.370234 | -1.835396 | 18 | 35 | c("MMP9", "SERPINA1", "APOC1", "PLAT", "LGMN", "ANXA1", "CLU", "CTSB", "SERPINE1", "A2M", "PECAM1", "LTA4H", "MMP2") |
| 16 | HALLMARK_OXIDATIVE_PHOSPHORYLATION | 0.028369 | 0.088652 | -0.275098 | -1.547956 | 39 | 56 | c("PDK4", "MAOB", "NDUFC2", "UQCRQ", "ATP5F1B", "COX4I1", "NDUFC1", "UQCR10", "MGST3", "ATP5F1E", "VDAC3", "ECH1", "ATP6V0E1", "NDUFA2") |
| 17 | HALLMARK_ESTROGEN_RESPONSE_EARLY | 0.030528 | 0.089789 | 0.310237 | 1.481859 | 270 | 83 | c("GJA1", "SLC1A1", "REEP1", "CELSR1", "PODXL", "SLC39A6", "SIAH2", "FRK", "LRIG1", "SLC19A2", "CELSR2", "RAB17", "CCN5", "BCL2", "ABCA3", "OVOL2", "OLFM1", "SEMA3B", "MINDY1", "MREG", "ABAT", "ADCY1", "OPN3", "MUC1", "TJP3", "UGCG", "SLC7A2") |
| 18 | HALLMARK_ANDROGEN_RESPONSE | 0.06878 | 0.191055 | -0.296493 | -1.489892 | 114 | 37 | c("STEAP4", "SCD", "SAT1", "SGK1", "SELENOP", "SLC38A2", "SMS", "MAF", "ACTN1", "ACSL3", "ITGAV", "SPDEF") |
| 19 | HALLMARK_CHOLESTEROL_HOMEOSTASIS | 0.076808 | 0.202126 | -0.344018 | -1.458052 | 153 | 21 | c("LGALS3", "SCD", "LGMN", "CLU", "ACTG1", "ALCAM", "ECH1") |
| 20 | HALLMARK_ALLOGRAFT_REJECTION | 0.099043 | 0.247608 | -0.295063 | -1.405923 | 175 | 31 | c("MMP9", "SRGN", "CTSS", "SPI1", "ITGB2", "HCLS1", "CD47") |
| 21 | HALLMARK_MTORC1_SIGNALING | 0.111349 | 0.253066 | -0.230317 | -1.305628 | 155 | 57 | c("TFRC", "CXCR4", "CORO1A", "SCD", "LGMN", "BTG2", "IGFBP5", "NUPR1", "MAP2K3", "LTA4H", "ITGB2", "SQSTM1") |
| 22 | HALLMARK_P53_PATHWAY | 0.11127 | 0.253066 | -0.227777 | -1.302799 | 155 | 59 | c("HMOX1", "BTG1", "SAT1", "FOS", "BTG2", "EPHX1", "TM4SF1", "JUN", "NUPR1", "ABCC5", "RACK1", "TXNIP", "CTSD") |
| 23 | HALLMARK_MYC_TARGETS_V1 | 0.187793 | 0.408247 | -0.205079 | -1.200241 | 239 | 66 | c("CLNS1A", "MYC", "RACK1", "EEF1B2", "VDAC3", "EIF4A1", "PSMA7", "PCBP1", "MCM4", "PA2G4", "SRM", "EIF4G2", "SRSF3", "PRPF31", "GNL3", "TUFM", "POLE3", "ILF2", "SMARCC1", "HNRNPD", "BUB3", "NOP16", "PWP1", "EIF3B", "UBA2", "DHX15", "RAD23B", "SRPK1", "RRM1", "RNPS1", "DEK", "NCBP2", "TRA2B", "PSMD1", "IMPDH2", "PSMD3", "HSPD1", "POLD2", "SSBP1", "FAM120A", "SET", "UBE2E1", "NOP56", "HDGF", "EIF3J", "CBX3", "HNRNPA2B1", "SRSF2", "HDAC2", "MCM7", "SRSF1", "USP1", "CUL1", "HNRNPU", "DUT", "EIF4H", □ "CCT3", "KPNB1", "HSPE1", "HNRNPR", "TXNL4A") |
| 24 | HALLMARK_KRAS_SIGNALING_DN | 0.209908 | 0.437308 | 0.370713 | 1.245315 | 1660 | 18 | c("BMPR1B", "THNSL2", "RGS11", "CELSR2", "CLSTN3", "IGFBP2", "MTHFR", "IFI44L", "CCDC106", "MFSD6", "GAMT") |

| | | | | | | | |
|---|---|---|---|---|---|---|---|
| 25 | HALLMARK_FATTY_ACID_METABOLISM | 0.363986 | 0.727971 | 0.269023 | 1.087769 | 3031 | 37 c("REEP6", "GPD2", "BMPR1B", "BLVRA", "D2HGDH", "ENO2", "YWHAH", "DHCR24", "GLUL", "UROS", "CRAT", "ADIPOR2") |
| 26 | HALLMARK_APICAL_SURFACE | 0.393251 | 0.728243 | 0.345742 | 1.059603 | 3029 | 13 c("EPHB4", "TMEM8B", "SULF2") |
| 27 | HALLMARK_IL2_STAT5_SIGNALING | 0.380485 | 0.728243 | 0.25537 | 1.074881 | 3216 | 44 c("TNFRSF18", "ECM1", "LRIG1", "ENPP1", "RABGAP1L", "BCL2", "S100A1", "NOP2", "PHTF2", "MUC1", "TNFSF10") |
| 28 | HALLMARK_EPITHELIAL_MESENCHYMAL_TRANSITION | 0.448718 | 0.801282 | -0.167736 | -1.006336 | 524 | 76 c("GEM", "SFRP4", "MGP", "CCN1", "SAT1", "CCN2", "JUN", "QSOX1", "SERPINE1", "VIM", "PMP22", "MMP2", "BGN", "WIPF1") |
| 29 | HALLMARK_WNT_BETA_CATENIN_SIGNALING | 0.536097 | 0.866081 | 0.317501 | 0.947969 | 4098 | 12 c("AXIN1", "KAT2A", "FZD1", "MAML1", "CUL1", "NOTCH1", "HDAC2", "NCSTN", "CSNK1E", "NUMB") |
| 30 | HALLMARK_TGF_BETA_SIGNALING | 0.545016 | 0.866081 | -0.207719 | -0.939275 | 1016 | 26 c("CDH1", "SERPINE1", "LTBP2", "IFNGR2", "UBE2D3", "SKI", "HDAC1", "SKIL", "PPM1A", "NCOR2", "SLC20A1", "CDK9", "BMPR2", "TRIM33", "KLF10", "PMEPA1", "MAP3K7", "RAB31", "SMAD7", "HIPK2", "TGFBR1", "SMURF2", "SMURF1", "TJP1", "SMAD3", "ARID4B") |
| 31 | HALLMARK_NOTCH_SIGNALING | 0.555906 | 0.866081 | 0.319191 | 0.929552 | 4230 | 11 c("KAT2A", "SAP30", "NOTCH2", "FZD1", "CUL1", "NOTCH1") |
| 32 | HALLMARK_APICAL_JUNCTION | 0.588147 | 0.866081 | 0.203917 | 0.92838 | 5120 | 64 c("AMH", "AMIGO2", "PIK3R3", "TMEM8B", "EVL", "THBS3", "SGCE") |
| 33 | HALLMARK_ANGIOGENESIS | 0.580404 | 0.866081 | -0.294745 | -0.875036 | 1551 | 8 c("APP", "ITGAV", "NRP1", "FGFR1", "FSTL1", "PTK2", "STC1", "VEGFA") |
| 34 | HALLMARK_SPERMATOGENESIS | 0.588935 | 0.866081 | 0.303406 | 0.905886 | 4502 | 12 c("SPATA6", "IFT88", "PHKG2", "MLLT10") |
| 35 | HALLMARK_UNFOLDED_PROTEIN_RESPONSE | 0.632838 | 0.885681 | -0.167264 | -0.88257 | 978 | 44 c("STC2", "LSM1", "ATP6V0D1", "EIF4EBP1", "HERPUD1", "CEBPB", "EIF4A1", "EEF2", "ATF4", "LSM4", "SPCS1", "XBP1", "KIF5B", "SERP1", "MTHFD2", "EIF4G1", "DCTN1", "YIF1A", "FUS", "PREB", "CNOT2", "ARFGAP1", "CXXC1", "EDEM1", "NOP56", "DNAJC3", "ATF6", "PARN", "ALDH18A1", "HYOU1", "EIF4A2", "SHC1", "NOP14", "GOSR2", "SRPRA", "VEGFA", "TATDN2", "SLC30A5", "CNOT4", "MTREX", "EXOSC10", "ERN1", "IMP3") |
| 36 | HALLMARK_MYC_TARGETS_V2 | 0.63769 | 0.885681 | 0.254275 | 0.868944 | 5068 | 19 c("PPRC1", "NOP2", "PPAN", "AIMP2", "TCOF1", "MPHOSPH10") |
| 37 | HALLMARK_HEDGEHOG_SIGNALING | 0.669331 | 0.904502 | 0.297502 | 0.840203 | 5023 | 10 c("CELSR1", "VEGFA") |
| 38 | HALLMARK_INFLAMMATORY_RESPONSE | 0.752257 | 0.924141 | -0.162249 | -0.789385 | 1332 | 33 c("BTG2", "KLF6", "SRI", "MYC", "SERPINE1", "IL1R1") |
| 39 | HALLMARK_GLYCOLYSIS | 0.757796 | 0.924141 | -0.140516 | -0.807765 | 1044 | 60 c("STC2", "TFF3", "CXCR4", "TALDO1", "QSOX1", "CITED2") |
| 40 | HALLMARK_UV_RESPONSE_DN | 0.709614 | 0.924141 | 0.187208 | 0.840669 | 6118 | 60 c("GJA1", "CDON", "PIK3R3", "ICA1", "TOGARAM1", "COL11A1", "CDK13", "DBP", "NOTCH2", "LPAR1", "CDC42BPA", "MTA1", "SMAD3", "INPP4B", "DLG1", "ERBB2", "PIAS3", "TJP1", "SCAF8") |
| 41 | HALLMARK_BILE_ACID_METABOLISM | 0.754938 | 0.924141 | 0.226187 | 0.772956 | 6000 | 19 c("PEX1", "ABCA3", "HSD3B7", "EFHC1", "DHCR24", "ABCA2", "PEX13", "GNPAT") |
| 42 | HALLMARK_E2F_TARGETS | 0.818836 | 0.974805 | -0.141521 | -0.753945 | 1251 | 46 c("TFRC", "NUDT21", "MYC", "CDKN1B", "SNRPB", "HELLS", "SLBP", "DNMT1", "MCM4", "PA2G4", "CDC25B", "NUP107", "MTHFD2", "TUBB", "HNRNPD", "DCTPP1", "DEK", "TRA2B", "NUP153", "POLD2", "STAG1", "SMC3", "NOP56", "SMC1A", "NBN", "SSRP1", "CNOT9", "ILF3", "PNN", "NUP205", "SRSF2", "MCM7", "KIF22", "SRSF1", "USP1", "DUT", "LUC7L3", "TBRG4", "CBX5") |
| 43 | HALLMARK_MITOTIC_SPINDLE | 0.973795 | 0.995053 | 0.11991 | 0.56117 | 8583 | 74 c("SORBS2", "CDC42EP1", "ARHGAP29", "FGD6", "LATS1", "DST", "CYTH2", "ALMS1", "NOTCH2", "CDC42BPA", "TUBGCP6", "KIF1B", "DLG1", "TAOK2", "CSNK1D", "ARHGEF12", "WASL", "NUMA1", "RHOT2", "ARL8A", "RASAL2", "KIFAP3", "PDLIM5", "TUBGCP2", "RABGAP1", "KIF22", "FARP1", "PPP4R2", "DOCK4", "TUBGCP3", "SMC1A", "ARHGEF11", "HDAC6", "BCAR1", "RASA1", "ARHGDIA", "SMC3", "ARHGEF2", "ARFIP2", "MID1IP1", "RALBP1", "MAPRE1", "KLC1", "SPTAN1", "ROCK1", "RAB3GAP1", "RASA2", "TLK1", "HOOK3", "CD2AP", "TRIO", "LRPPRC"□ ) |
| 44 | HALLMARK_DNA_REPAIR | 0.992031 | 0.995053 | 0.107236 | 0.443933 | 8340 | 41 c("POM121", "GTF2H5", "DGCR8", "POLL", "ADCY6", "NME3", "CSTF3", "SURF1", "ELOA", "ELL", "DUT", "RALA", "NFX1", "POLR1D", "BCAM", "SSRP1", "CANT1", "ARL6IP1", "POLR2H", "NME4", "SMAD5", "CCNO", "IMPDH2", "NCBP2", "STX3", "AAAS", "TMED2", "GTF2F1") |
| 45 | HALLMARK_G2M_CHECKPOINT | 0.946242 | 0.995053 | 0.135568 | 0.598735 | 8131 | 55 c("HOXC10", "CUL5", "NSD2", "SAP30", "NOTCH2", "MAP3K20", "HUS1", "SMAD3", "LIG3", "NUMA1", "MTF2", "SFPQ", "RASAL2", "ATRX", "KPNB1", "HNRNPU", "CUL1", "PURA", "SRSF1", "KIF22", "ARID4A", "SRSF2", "ILF3", "YTHDC1", "TENT4A", "SMC1A", "TNPO2", "HSPA8", "STAG1", "PRPF4B", "NUP98", "EWSR1", "CUL3", "TRA2B", "CUL4A", "UPF1", "SS18", "RAD23B", "ATF5", "DR1", "RBM14", "BUB3", "HNRNPD", "ABL1", "SMARCC1") |

| | | | | | | | | |
|---|---|---|---|---|---|---|---|---|
| 46 | HALLMARK_PROTEIN_SECRETION | 0.945618 | 0.995053 | 0.143986 | 0.582195 | 7876 | 37 | c("SCRN1", "COG2", "ICA1", "GBF1", "DST", "SCAMP3", "KIF1B", "GALC", "SEC24D", "GOSR2", "VPS45", "YKT6", "ARCN1", "STX12", "STX16", "MON2", "RER1", "ADAM10", "ARFGAP3", "TOM1L1", "ARFIP1", "USO1", "TMED2") |
| 47 | HALLMARK_INTERFERON_ALPHA_RESPONSE | 0.948706 | 0.995053 | 0.142503 | 0.569735 | 7878 | 35 | c("MOV10", "TRIM26", "TRIM14", "SAMD9L", "IFI44L", "SLC25A28", "RNF31", "IFI44", "OAS1", "NUB1", "CNP", "LGALS3BP", "IFI35", "TAP1", "ADAR", "EIF2AK2", "CMTR1", "CASP8", "CSF1", "PSMB8", "PARP14", "LY6E", "ISG15", "ELF1") |
| 48 | HALLMARK_INTERFERON_GAMMA_RESPONSE | 0.995053 | 0.995053 | -0.083581 | -0.46131 | 1407 | 53 | c("BTG1", "SRI", "PIM1", "TXNIP", "SAMHD1", "IFITM3", "PML", "BST2", "PSME1", "CIITA", "IFI27", "TAPBP", "IRF9", "STAT2", "STAT1", "MTHFD2", "OGFR", "PTPN1", "TRIM25", "STAT3", "SP110", "BPGM", "PTPN6", "RBCK1", "NFKB1", "ISG15", "LY6E", "PARP14", "PSMB8", "CASP8", "CMTR1", "EIF2AK2", "OAS3", "ADAR", "CASP4", "TAP1", "ZNFX1", "IFI35", "LGALS3BP", "IFI44", "RNF31", "SLC25A28", "RIPK1", "SPPL2A", "IFI44L", "RNF213", "CFB", "SAMD9L", "TRIM14", "TRIM26", "TNFSF10") |
| 49 | HALLMARK_PI3K_AKT_MTOR_SIGNALING | 0.989783 | 0.995053 | -0.0934 | -0.491078 | 1549 | 43 | c("CXCR4", "MAP2K3", "PFN1", "GRK2", "SQSTM1", "CDKN1B", "CFL1") |
| 50 | HALLMARK_PEROXISOME | 0.881903 | 0.995053 | -0.145615 | -0.673345 | 1612 | 28 | c("CAT", "ABCC5", "ECH1", "SCP2", "ALDH9A1", "HRAS", "ECI2", "ITGB1BP1", "VPS4B", "PRDX1", "HSD17B4", "SMARCC1", "PEX2", "ERCC1", "ACAA1", "GSTK1", "SEMA3C", "ACOX1", "PEX11B", "GNPAT", "PEX13", "FIS1", "CRAT", "FDPS", "DHCR24", "YWHAH", "HSD3B7", "CRABP2") |

| Cluster 19 | pathway | pval | padj | ES | NES | nMoreExtreme | size | leadingEdge |
|---|---|---|---|---|---|---|---|---|
| 1 | HALLMARK_TNFA_SIGNALING_VIA_NFKB | 0.000109 | 0.000706 | -0.657083 | -2.089182 | 0 | 53 | c("CCN1", "INHBA", "SERPINE1", "MARCKS", "PLPP3", "NR4A1", "EGR1", "GEM", "FOSB", "PHLDA1", "KLF9", "SGK1", "FOSL2", "PLAU", "FOS", "PMEPA1", "SOCS3", "JUN", "NFE2L2", "DUSP1", "PFKFB3", "TNC", "MAP3K8", "PLAUR", "B4GALT1", "BTG2", "ZFP36", "SQSTM1", "BTG1", "JUNB", "PNRC1", "ID2", "CEBPD", "CD44", "SAT1") |
| 2 | HALLMARK_HYPOXIA | 0.000107 | 0.000706 | -0.625094 | -2.013878 | 0 | 60 | c("COL5A1", "CCN2", "CAVIN1", "IGFBP3", "CCN1", "SERPINE1", "LOX", "HMOX1", "DCN", "PAM", "BGN", "SDC3", "CDKN1B", "TGFBI", "SDC2", "TGFB3", "FOSL2", "FOS", "JUN", "PIM1", "DUSP1", "PFKFB3", "PLAUR", "DDIT4", "MT2A", "ANXA2", "ZFP36") |
| 3 | HALLMARK_MYOGENESIS | 0.000111 | 0.000706 | -0.662265 | -2.060624 | 0 | 45 | c("COL4A2", "IGFBP3", "AEBP1", "IGFBP7", "APOD", "ADAM12", "PVALB", "TAGLN", "GSN", "COL1A1", "COL3A1", "CD36", "TPM2", "COL6A3", "SPARC", "COL6A2", "GPX3", "SORBS3", "HSPB8") |
| 4 | HALLMARK_COMPLEMENT | 0.00011 | 0.000706 | -0.700259 | -2.184945 | 0 | 46 | c("COL4A2", "C3", "MMP14", "C1S", "SERPINE1", "APOC1", "C1QC", "LGALS3", "LTF", "SERPING1", "C1R", "LRP1", "CD36", "TIMP2", "C1QA", "FN1", "LGMN", "SERPINA1", "CTSD", "CTSL", "LIPA", "CTSB", "PIM1", "PLAUR", "TIMP1") |
| 5 | HALLMARK_EPITHELIAL_MESENCHYMAL_TRANSITION | 0.000102 | 0.000706 | -0.803456 | -2.72009 | 0 | 106 | c("COL5A2", "THBS2", "COMP", "COL4A1", "ACTA2", "COL4A2", "FSTL1", "COL11A1", "COL12A1", "CDH11", "COL5A1", "CALD1", "COL16A1", "FBN1", "CCN2", "IGFBP3", "DPYSL3", "LOXL2", "SFRP4", "CCN1", "LAMC1", "THY1", "MMP14", "ADAM12", "MMP2", "INHBA", "LUM", "SERPINE1", "GJA1", "FBLN2", "LOX", "CXCL12", "PRRX1", "CTHRC1", "FAP", "NNMT", "PDGFRB", "HTRA1", "TAGLN", "ELN", "DCN", "BGN", "LOXL1", "COL1A1", "POSTN", "DAB2", "COL3A1", "MYL9", "LRRC15", "COL1A2", "GEM", "LRP1", "TPM2", "MXRA5", "PCOLCE", "TGFBI", "FN1", "COL6A3", "SPARC", "FBLN1", "COL6A2", "TIMP3", "DST", "ITGA5", "SPP1", "TPM1", "PMEPA1", "JUN", "FUCA1", "ITGAV", "VIM", "VCAN", "TNC", "PMP22", "SERPINH1", "PLAUR", "PLOD1", "TIMP1", "FLNA") |
| 6 | HALLMARK_UV_RESPONSE_DN | 0.00011 | 0.000706 | -0.653611 | -2.057275 | 0 | 49 | c("COL5A2", "COL11A1", "CCN1", "LAMC1", "SERPINE1", "GJA1", "PDGFRB", "NRP1", "EFEMP1", "PLPP3", "COL1A1", "DAB2", "COL3A1", "AKT3", "COL1A2", "CDKN1B", "FHL2", "SDC2", "ANXA4", "TGFBR2", "DUSP1", "PMP22", "ANXA2", "RUNX1", "IGF1R", "ADD3", "RBPMS") |
| 7 | HALLMARK_COAGULATION | 0.000113 | 0.000706 | -0.824954 | -2.518601 | 0 | 39 | c("COMP", "FBN1", "C3", "MMP14", "C1S", "MMP2", "A2M", "SERPINE1", "APOC1", "MMP11", "HTRA1", "CTSK", "GSN", "S100A1", "SERPING1", "MMP9", "CRIP2", "C1R", "LRP1", "C1QA", "FN1", "SPARC", "TIMP3", "LGMN", "SERPINA1", "PLAU", "CAPN2", "CTSB", "ANXA1") |
| 8 | HALLMARK_KRAS_SIGNALING_UP | 0.000113 | 0.000706 | -0.677587 | -2.059782 | 0 | 38 | c("IGFBP3", "APOD", "INHBA", "PRRX1", "GPNMB", "TMEM176B", "MMP11", "SPARCL1", "NRP1", "MMP9", "ANKH", "MAFB", "PLAU", "SPP1", "FUCA1", "SERPINA3", "CPE", "ENG", "PLAUR", "EMP1", "ITGB2", "IKZF1", "FCER1G", "PECAM1", "ID2") |
| 9 | HALLMARK_ANGIOGENESIS | 0.00014 | 0.000778 | -0.815768 | -1.961277 | 0 | 12 | c("COL5A2", "FSTL1", "LUM", "NRP1", "POSTN", "COL3A1", "SPP1", "ITGAV", "VCAN", "TIMP1") |
| 10 | HALLMARK_SPERMATOGENESIS | 0.00052 | 0.002599 | 0.826537 | 2.022267 | 1 | 6 | c("SLC12A2", "GSTM3", "CDK1", "HSPA2") |
| 11 | HALLMARK_G2M_CHECKPOINT | 0.00091 | 0.004136 | 0.444254 | 2.099308 | 0 | 40 | c("SLC12A2", "INCENP", "WRN", "TOP2A", "AURKB", "CDK1", "BUB3", "E2F1", "LMNB1", "POLE", "SAP30", "UBE2C", "SNRPD1", "NSD2", "SLC38A1", "TPX2", "HOXC10", "TACC3", "SMARCC1", "HNRNPU", "NOTCH2", "CKS1B", "CCND1", "HSPA8", "MYBL2", "KPNB1") |
| 12 | HALLMARK_E2F_TARGETS | 0.001058 | 0.004409 | 0.4493 | 2.215182 | 0 | 46 | c("TOP2A", "AURKB", "CDK1", "LMNB1", "CSE1L", "ASF1B", "POLE", "MCM4", "POLD1", "HELLS", "RPA1", "POLD2", "CBX5", "TACC3", "NOP56", "TP53", "SNRPB", "DNMT1", "EIF2S1", "CKS1B", "TK1", "MYBL2", "NUP205", "SLBP", "NAA38", "NUP107", "DDX39A", "PCNA", "TUBG1", "HNRNPD") |
| 13 | HALLMARK_APICAL_JUNCTION | 0.001434 | 0.005517 | -0.577815 | -1.806649 | 12 | 47 | c("CDH11", "COL16A1", "FBN1", "THY1", "MMP2", "SDC3", "MMP9", "AKT3", "MYL9", "TGFBI", "CD99", "SORBS3", "CDH1", "CD276", "PARVA", "VCAN", "CERCAM", "B4GALT1", "MDK", "CNN2") |
| 14 | HALLMARK_APOPTOSIS | 0.003554 | 0.012422 | -0.525354 | -1.683259 | 32 | 56 | c("MMP2", "LUM", "HMOX1", "PDGFRB", "GSN", "DCN", "BGN", "LGALS3", "CDKN1B", "ANKH", "TIMP2", "TIMP3", "GPX3", "JUN", "ANXA1", "TIMP1", "EMP1", "BTG2", "TNFRSF12A", "PPT1", "SQSTM1", "DPYD", "CD44", "SAT1", "DAP") |

| | | | | | | | | |
|---|---|---|---|---|---|---|---|---|
| 15 | HALLMARK_MYC_TARGETS_V1 | 0.003727 | 0.012422 | 0.370849 | 1.872529 | 2 | 52 | c("BUB3", "C1QBP", "RRM1", "MCM4", "HSPD1", "SNRPD1", "HPRT1", "ERH", "HDAC2", "POLD2", "TYMS", "NOP56", "SMARCC1", "HNRNPU", "CBX3", "EIF2S1", "PWP1", "SNRPA", "IMPDH2", "KPNB1", "VDAC1", "PCNA", "CNBP", "NDUFAB1", "HNRNPD", "TOMM70", "EIF4A1", "SNRPD2", "PPIA", "TXNL4A", "GNL3", "NME1", "MCM7", "DEK", "YWHAE", "TUFM") |
| 16 | HALLMARK_MYC_TARGETS_V2 | 0.015152 | 0.045985 | 0.565133 | 1.741463 | 44 | 11 | c("WDR74", "SLC19A1", "MCM4", "HSPD1", "NOP56", "CBX3", "SORD", "FARSA") |
| 17 | HALLMARK_PEROXISOME | 0.015635 | 0.045985 | 0.396609 | 1.683052 | 24 | 28 | c("ALB", "TOP2A", "PRDX5", "SLC25A4", "ABCD3", "SMARCC1", "NUDT19", "HRAS", "SOD1", "CRABP2", "PEX11B", "LONP2", "ABCC5", "DHCR24", "ECI2", "CNBP") |
| 18 | HALLMARK_DNA_REPAIR | 0.02085 | 0.057917 | 0.354249 | 1.621829 | 25 | 36 | c("DDB1", "POLR2G", "ARL6IP1", "ZWINT", "POLD1", "UMPS", "ADCY6", "HPRT1", "TYMS", "GTF2H5", "TP53") |
| 19 | HALLMARK_TGF_BETA_SIGNALING | 0.023498 | 0.061838 | -0.588872 | -1.610493 | 186 | 21 | c("SERPINE1", "LTBP2", "ID3", "RAB31", "PMEPA1", "CDH1", "ENG", "SKI", "JUNB", "THBS1", "ID2", "FNTA", "BMPR2", "SLC20A1", "SKIL") |
| 20 | HALLMARK_OXIDATIVE_PHOSPHORYLATION | 0.028078 | 0.070194 | 0.275007 | 1.487501 | 12 | 74 | c("NDUFS8", "OAT", "NDUFV1", "GLUD1", "ALDH6A1", "COX8A", "ATP5PF", "AIFM1", "AFG3L2", "SLC25A4", "NDUFB2", "VDAC2", "COX6B1", "TCIRG1", "MDH2", "ACADSB", "ATP5MF", "GPI", "UQCRC1", "NDUFS6", "HSPA9", "ATP5F1B", "SLC25A11", "ECHS1", "RHOT1", "FH", "UQCRQ", "TIMM50", "ATP6AP1", "VDAC1", "NDUFV2", "ATP6V0B", "ATP6V1F", "NDUFAB1", "ATP5MC2", "TOMM70", "ETFB", "DLD", "NDUFS2", "NDUFC2", "ATP5F1A") |
| 21 | HALLMARK_ADIPOGENESIS | 0.033713 | 0.079265 | -0.462076 | -1.485133 | 313 | 58 | c("COL4A1", "FABP4", "C3", "CAVIN1", "LAMA4", "SPARCL1", "APOE", "STOM", "PHLDB1", "CD36", "GPX3", "HSPB8") |
| 22 | HALLMARK_INFLAMMATORY_RESPONSE | 0.034877 | 0.079265 | -0.549797 | -1.555151 | 286 | 25 | c("MMP14", "INHBA", "SERPINE1", "ITGA5", "STAB1", "PLAUR", "SLC7A2", "TIMP1", "BTG2", "CYBB", "CD55") |
| 23 | HALLMARK_HEDGEHOG_SIGNALING | 0.044686 | 0.097144 | -0.746365 | -1.512273 | 274 | 6 | c("THY1", "NRP1", "DPYSL2") |
| 24 | HALLMARK_IL6_JAK_STAT3_SIGNALING | 0.059961 | 0.119923 | -0.562979 | -1.491689 | 464 | 18 | c("A2M", "HMOX1", "CD36", "SOCS3", "JUN", "PIM1", "MAP3K8", "TNFRSF12A", "IL13RA1", "CD44") |
| 25 | HALLMARK_ALLOGRAFT_REJECTION | 0.058651 | 0.119923 | -0.530996 | -1.490519 | 479 | 24 | c("THY1", "INHBA", "MMP9", "STAB1", "CD74", "TIMP1", "FLNA", "ITGB2") |
| 26 | HALLMARK_ESTROGEN_RESPONSE_EARLY | 0.064787 | 0.12459 | -0.416799 | -1.374155 | 617 | 74 | c("ADCY1", "GJA1", "CXCL12", "NBL1", "PAPSS2", "SEC14L2", "PGR", "MAPT", "FHL2", "ELOVL5", "RAB31", "HSPB8", "RHOBTB3", "ABHD2", "FOS", "NRIP1", "RHOD", "SLC7A2", "B4GALT1", "GAB2", "SCNN1A", "GREB1", "PRSS23", "IGF1R", "IGFBP4", "ADD3") |
| 27 | HALLMARK_ESTROGEN_RESPONSE_LATE | 0.095293 | 0.176469 | -0.401751 | -1.324543 | 908 | 74 | c("CXCL14", "CXCL12", "NBL1", "LTF", "PAPSS2", "PGR", "MAPT", "SERPINA1", "ELOVL5", "SGK1", "RAB31", "HSPB8", "ABHD2", "FOS", "CDH1", "SERPINA3", "NRIP1", "CPE", "MDK", "ZFP36", "SCNN1A", "PRSS23", "IGFBP4", "ADD3", "MYOF", "FARP1", "ID2") |
| 28 | HALLMARK_NOTCH_SIGNALING | 0.128235 | 0.22899 | 0.506431 | 1.392775 | 445 | 8 | c("SAP30", "HES1", "NOTCH2", "CCND1", "APH1A") |
| 29 | HALLMARK_GLYCOLYSIS | 0.148295 | 0.255681 | -0.413063 | -1.291521 | 1343 | 47 | c("COL5A1", "IGFBP3", "DCN", "PAM", "EGLN3", "SDC3", "TGFBI", "SDC2", "CHPF", "VCAN", "DDIT4", "B4GALT1", "PLOD1") |
| 30 | HALLMARK_XENOBIOTIC_METABOLISM | 0.168886 | 0.272397 | -0.411021 | -1.275318 | 1521 | 44 | c("SERPINE1", "PDK4", "TMEM176B", "HMOX1", "PAPSS2", "APOE", "CD36", "FBLN1", "ELOVL5") |
| 31 | HALLMARK_IL2_STAT5_SIGNALING | 0.165641 | 0.272397 | -0.416925 | -1.281608 | 1479 | 41 | c("NRP1", "S100A1", "COL6A1", "PHLDA1", "SPP1", "ANXA4", "CTSZ", "PIM1", "ITGAV", "ITGA6", "MAP3K8", "EMP1", "CKAP4", "NFKBIZ", "IGF1R", "MAPKAPK2", "CD44", "PLEC", "GPX4", "BHLHE40", "BMPR2", "KLF6", "AHR", "SERPINB6", "BCL2") |
| 32 | HALLMARK_BILE_ACID_METABOLISM | 0.189324 | 0.286855 | 0.358301 | 1.264665 | 453 | 16 | c("PXMP2", "PRDX5", "AR", "PEX1", "ABCD3", "SOD1", "LONP2", "DHCR24") |
| 33 | HALLMARK_PANCREAS_BETA_CELLS | 0.187577 | 0.286855 | -0.710072 | -1.304195 | 1059 | 4 | c("AKT3", "MAFB") |
| 34 | HALLMARK_ANDROGEN_RESPONSE | 0.242166 | 0.356126 | -0.394812 | -1.20871 | 2155 | 40 | c("BMPR1B", "ANKH", "MAF", "ELOVL5", "SGK1", "ABHD2", "PMEPA1", "STEAP4", "ITGAV", "SELENOP", "B4GALT1", "ELL2", "ARID5B") |
| 35 | HALLMARK_UV_RESPONSE_UP | 0.259238 | 0.370341 | -0.38955 | -1.1926 | 2307 | 40 | c("MMP14", "HMOX1", "NR4A1", "FOSB", "GPX3", "FOS", "BTG2", "MGAT1", "UROD", "PPT1", "SQSTM1", "BTG1", "JUNB") |
| 36 | HALLMARK_REACTIVE_OXYGEN_SPECIES_PATHWAY | 0.273144 | 0.379366 | -0.472128 | -1.195517 | 2026 | 15 | c("MPO", "GPX3", "CAT", "LSP1", "FTL", "JUNB") |
| 37 | HALLMARK_INTERFERON_GAMMA_RESPONSE | 0.299689 | 0.403125 | -0.387244 | -1.157424 | 2597 | 34 | c("C1S", "SERPING1", "C1R", "SOCS3", "CD74", "PIM1", "IFI30", "MT2A", "IFI27", "ARID5B", "BTG1") |
| 38 | HALLMARK_APICAL_SURFACE | 0.306375 | 0.403125 | -0.520599 | -1.171041 | 2056 | 9 | c("THY1", "SULF2", "PLAUR", "B4GALT1") |

| | | | | | | | | |
|---|---|---|---|---|---|---|---|---|
| 39 | HALLMARK_P53_PATHWAY | 0.344147 | 0.441213 | -0.355118 | -1.110344 | 3118 | 47 | c("HMOX1", "STOM", "S100A10", "RAP2B", "CTSD", "FOS", "JUN", "FUCA1", "IFI30", "DDIT4", "ZFP36L1", "BTG2", "BTG1", "PLXNB2", "SDC1", "S100A4", "SAT1", "NUPR1", "HEXIM1") |
| 40 | HALLMARK_MTORC1_SIGNALING | 0.383249 | 0.479061 | 0.205967 | 1.044065 | 301 | 53 | c("INSIG1", "GSR", "IGFBP5", "PPA1", "SLC1A5", "MCM4", "HSPD1", "ACLY", "EBP", "LDLR", "TRIB3", "RPA1", "HPRT1") |
| 41 | HALLMARK_HEME_METABOLISM | 0.412199 | 0.502681 | -0.332956 | -1.058627 | 3797 | 53 | c("C3", "SLC4A1", "ACP5", "CAST", "CTSB", "TNS1", "CIR1", "UBAC1", "CAT", "ELL2", "MPP1", "BTG2", "SLC30A1", "UROD") |
| 42 | HALLMARK_CHOLESTEROL_HOMEOSTASIS | 0.491749 | 0.571801 | 0.239561 | 0.970079 | 893 | 24 | c("MAL2", "STX5", "CHKA", "EBP", "LDLR", "TRIB3", "SREBF2", "ALCAM", "LSS") |
| 43 | HALLMARK_PI3K_AKT_MTOR_SIGNALING | 0.482176 | 0.571801 | 0.231919 | 0.984174 | 770 | 28 | c("CDK1", "E2F1", "PPP1CA", "GRK2", "TRIB3", "ARF1", "ACACA", "HRAS", "GSK3B", "MAP3K7", "PFN1", "PAK4", "RAF1", "AKT1") |
| 44 | HALLMARK_FATTY_ACID_METABOLISM | 0.618229 | 0.702533 | -0.305236 | -0.892127 | 5249 | 30 | c("BMPR1B", "CD36", "S100A10", "ELOVL5") |
| 45 | HALLMARK_INTERFERON_ALPHA_RESPONSE | 0.758258 | 0.825508 | -0.281878 | -0.764108 | 5990 | 20 | c("C1S", "CD74", "IFI30", "IFI27") |
| 46 | HALLMARK_KRAS_SIGNALING_DN | 0.759467 | 0.825508 | -0.311635 | -0.761419 | 5474 | 13 | c("BMPR1B", "SGK1", "SELENOP", "BTG2") |
| 47 | HALLMARK_PROTEIN_SECRETION | 0.803754 | 0.855057 | -0.26679 | -0.71656 | 6294 | 19 | c("PAM", "DST") |
| 48 | HALLMARK_MITOTIC_SPINDLE | 0.913658 | 0.941787 | -0.201716 | -0.629393 | 8274 | 46 | c("GSN", "MARCKS", "DST", "PALLD", "FLNA", "PCM1", "FARP1", "PREX1", "SUN2", "MYH9", "ARHGAP29", "CNTRL", "BCAR1", "AKAP13") |
| 49 | HALLMARK_WNT_BETA_CATENIN_SIGNALING | 0.922952 | 0.941787 | 0.229349 | 0.597166 | 3401 | 7 | c("HDAC2", "TP53", "NUMB", "NCSTN") |
| 50 | HALLMARK_UNFOLDED_PROTEIN_RESPONSE | 0.979054 | 0.979054 | -0.163059 | -0.46764 | 8179 | 27 | c("DDIT4", "HERPUD1", "SHC1", "DCTN1", "DNAJA4", "EDEM1", "SRPRA", "SERP1", "ATF4", "CEBPB", "ATP6V0D1", "CALR", "WFS1", "EEF2", "SPCS1", "LSM1", "ATF6", "EIF4A2", "EIF4A1", "KHSRP", "LSM4", "SRPRB", "FUS", "CKS1B", "EIF2S1", "HSPA9", "NOP56") |

| Cluster 20 | pathway | pval | padj | ES | NES | nMoreExtreme | size | leadingEdge |
|---|---|---|---|---|---|---|---|---|
| 1 | HALLMARK_ESTROGEN_RESPONSE_EARLY | 1.00E-04 | 0.001004 | -0.509453 | -1.972219 | 0 | 73 | c("SEC14L2", "SLC39A6", "FLNB", "HES1", "PGR", "ADCY1", "TFF1", "HSPB8", "MAPT", "CA12", "STC2", "RET", "FARP1", "BHLHE40", "GFRA1", "PRSS23", "GAB2", "FASN", "SLC7A2", "ELF3", "LRIG1", "CCND1", "GREB1", "SCNN1A", "NBL1", "RHOBTB3", "CLDN7", "PLAAT3", "SLC9A3R1", "ABHD2", "SEMA3B", "THSD4", "FAM102A", "KRT8", "TFF3", "MLPH", "TOB1", "FOS", "IGF1R", "MYOF", "PAPSS2", "CELSR1", "RHOD", "GJA1") |
| 2 | HALLMARK_ESTROGEN_RESPONSE_LATE | 0.0001 | 0.001004 | -0.507062 | -1.955634 | 0 | 70 | c("DLG5", "FLNB", "SERPINA3", "CD9", "UGDH", "CDH1", "PGR", "COX6C", "TFF1", "HSPB8", "CXCL14", "MAPT", "SCUBE2", "CA12", "CPE", "RET", "FARP1", "AGR2", "PRLR", "PRSS23", "CCND1", "SCNN1A", "NBL1", "SERPINA1", "PLAAT3", "SLC9A3R1", "ABHD2", "MDK", "SEMA3B", "LSR", "FAM102A") |
| 3 | HALLMARK_MYOGENESIS | 0.0001 | 0.001004 | -0.512172 | -1.892399 | 0 | 48 | c("ADAM12", "HSPB8", "APOD", "TAGLN", "SCD", "STC2", "PVALB", "BHLHE40", "CNN3", "CLU", "IGFBP7", "SPDEF", "GSN", "AEBP1", "MYO1C", "TNNT1", "ITGB1", "ATP6AP1", "COL4A2", "ITGB5", "SPTAN1", "COL3A1", "IGFBP3", "SORBS3", "DTNA", "EIF4A2", "NQO1") |
| 4 | HALLMARK_EPITHELIAL_MESENCHYMAL_TRANSITION | 1.00E-04 | 0.001004 | -0.488707 | -1.963819 | 0 | 108 | c("COL5A2", "ADAM12", "COMP", "CCN1", "HTRA1", "THY1", "TAGLN", "CCN2", "FBLN1", "SFRP4", "TIMP3", "ELN", "THBS2", "TPM4", "COL11A1", "FLNA", "MXRA5", "LOXL2", "TIMP1", "MMP2", "ACTA2", "JUN", "SERPINE1", "COL12A1", "TGFBI", "COL4A1", "MYL9", "LRP1", "LUM", "FBLN2", "CTHRC1", "POSTN", "COL16A1", "CALD1", "SPP1", "THBS1", "CDH11", "ITGB1", "FSTL1", "LAMC1", "COL5A1", "PMEPA1", "MMP14", "BGN", "PDGFRB", "COL4A2", "GEM", "ITGB5", "CAPG", "LOXL1", "FAP", "SDC4", "COL3A1", "LOX", "DCN", "IGFBP3", "ECM1", "MGP", "DPYSL3", "DAB2", "GJA1", "TNC", "LRRC15", "ITGAV") |
| 5 | HALLMARK_COAGULATION | 0.0001 | 0.001004 | -0.619766 | -2.257819 | 0 | 43 | c("MMP11", "C3", "CD9", "APOC1", "COMP", "HTRA1", "C1S", "TIMP3", "CFB", "PRSS23", "C1R", "TIMP1", "MMP2", "CLU", "SERPINE1", "CAPN2", "GSN", "S100A1", "LRP1", "CRIP2", "SERPINA1", "THBS1", "C1QA", "PLAT", "MMP14", "A2M", "SERPING1") |
| 6 | HALLMARK_COMPLEMENT | 0.0002 | 0.00167 | -0.491281 | -1.829289 | 1 | 51 | c("C3", "APOC1", "CTSD", "C1S", "CFB", "C1R", "TIMP1", "C1QC", "CLU", "SERPINE1", "CD46", "CTSL", "LIPA", "LRP1", "TIMP2", "SERPINA1", "C1QA", "PLAT", "MMP14", "COL4A2", "SERPING1", "ANXA5", "GNAI2", "LGALS3", "LGMN", "FCER1G") |
| 7 | HALLMARK_HYPOXIA | 0.0005 | 0.003574 | -0.44402 | -1.685342 | 4 | 60 | c("MIF", "CCN1", "CCN2", "CA12", "STC2", "HMOX1", "BHLHE40", "ANXA2", "STC1", "JUN", "SERPINE1", "TGFB3", "TGFBI", "PAM", "MT2A", "TPD52", "COL5A1", "BGN", "SDC4", "LOX", "DCN", "FOS", "IGFBP3", "EFNA1", "DTNA", "DUSP1", "DDIT4", "MYH9", "FOSL2", "SDC2", "SDC3", "CCNG2", "PNRC1", "MAP3K1", "HDLBP", "CAVIN1", "SIAH2", "BCL2", "FBP1", "GAA") |
| 8 | HALLMARK_ANGIOGENESIS | 0.001708 | 0.010676 | -0.615568 | -1.820941 | 15 | 14 | c("COL5A2", "NRP1", "STC1", "TIMP1", "LUM", "POSTN", "SPP1", "FSTL1", "FGFR1", "COL3A1") |
| 9 | HALLMARK_MYC_TARGETS_V2 | 0.009687 | 0.053814 | 0.662687 | 1.877054 | 16 | 6 | c("PA2G4", "HSPE1", "SRM", "CBX3", "NOP56", "GNL3") |
| 10 | HALLMARK_UV_RESPONSE_DN | 0.011331 | 0.056653 | -0.417649 | -1.530432 | 112 | 45 | c("COL5A2", "CCN1", "NRP1", "COL11A1", "BHLHE40", "ANXA2", "SERPINE1", "EFEMP1", "INPP4B", "ERBB2", "LAMC1", "RBPMS", "IGFBP5", "PDGFRB", "COL3A1", "IGF1R", "DUSP1", "DAB2", "ANXA4", "GJA1", "AKT3", "SDC2", "PLPP3", "PIK3R3") |
| 11 | HALLMARK_KRAS_SIGNALING_UP | 0.018692 | 0.084962 | -0.416777 | -1.503378 | 185 | 40 | c("MMP11", "SERPINA3", "APOD", "GPNMB", "SPARCL1", "NRP1", "CPE", "CFB", "JUP", "SPP1", "SEMA3B", "PLAT", "IGFBP3", "FCER1G", "ENG", "ITGB2", "FUCA1", "PRRX1", "MAP3K1", "MAFB", "EMP1", "INHBA", "PLAUR", "LAPTM5", "PLAU") |
| 12 | HALLMARK_CHOLESTEROL_HOMEOSTASIS | 0.02425 | 0.101042 | -0.496868 | -1.555807 | 231 | 18 | c("CD9", "ACTG1", "SCD", "FASN", "CLU", "SEMA3B", "ANXA5", "LGALS3", "CTNNB1", "LGMN", "S100A11") |
| 13 | HALLMARK_ANDROGEN_RESPONSE | 0.027239 | 0.104765 | -0.408157 | -1.466803 | 270 | 39 | c("BMPR1B", "AZGP1", "SCD", "SPDEF", "CCND1", "INPP4B", "SELENOP", "TPD52", "B2M", "DHCR24", "ABHD2", "PMEPA1", "DBI", "KRT8", "STEAP4") |
| 14 | HALLMARK_APICAL_JUNCTION | 0.03333 | 0.119036 | -0.373647 | -1.400852 | 332 | 54 | c("CDH1", "PARVA", "THY1", "ACTG1", "MMP2", "TGFBI", "MYL9", "CLDN7", "WASL", "EVL", "JUP", "COL16A1", "CTNND1", "CDH11", "ITGB1", "MDK", "GNAI2", "SORBS3", "NECTIN2", "TSPAN4", "MYH9", "AKT3", "PIK3R3", "SYMPK", "LIMA1", "THBS3", "SDC3", "CTNNA1", "CLDN4", "ACTN4", "PCDH1", "FBN1", "CERCAM", "MSN", "VCAN", "ACTN1", "B4GALT1", "CD276", "MAPK13", "AKT2") |

| | | | | | | | | |
|---|---|---|---|---|---|---|---|---|
| 15 | HALLMARK_APICAL_SURFACE | 0.038046 | 0.120662 | -0.597688 | -1.527647 | 330 | 8 | c("GSTM3", "SULF2", "THY1") |
| 16 | HALLMARK_GLYCOLYSIS | 0.038612 | 0.120662 | -0.387457 | -1.415679 | 384 | 44 | c("MIF", "STC2", "STC1", "ELF3", "TGFBI", "PAM", "PKM", "SOD1", "PGLS", "COL5A1", "TFF3", "FUT8", "DCN", "IGFBP3", "PPIA", "IL13RA1", "DDIT4", "COPB2", "CHPF", "SDC1", "CYB5A", "SDC2", "SDC3", "HDLBP", "IDUA", "QSOX1", "EGLN3", "PLOD1", "GPC1", "GFPT1", "IER3", "VCAN", "VEGFA", "B4GALT1", "CD44", "ALDH9A1") |
| 17 | HALLMARK_XENOBIOTIC_METABOLISM | 0.052352 | 0.153976 | -0.377616 | -1.379724 | 521 | 44 | c("UGDH", "FBLN1", "CFB", "HMOX1", "MAN1A1", "SERPINE1", "PDK4", "MT2A", "MCCC2", "ESR1", "JUP", "SPINT2", "ATP2A2", "APOE", "DCXR", "ALDH2", "NQO1", "PAPSS2", "COMT", "CYB5A", "EPHX1") |
| 18 | HALLMARK_APOPTOSIS | 0.064464 | 0.178228 | -0.351027 | -1.327552 | 643 | 58 | c("IFITM3", "TIMP3", "HMOX1", "TIMP1", "MMP2", "JUN", "CLU", "CCND1", "GSN", "SOD1", "TIMP2", "LUM", "ERBB2", "PEA15", "PLAT", "BGN", "PDGFRB", "SPTAN1", "LGALS3", "DCN", "CTNNB1", "PPT1", "EMP1", "GPX4", "DAP3", "LMNA", "CREBBP", "APP", "DAP", "ERBB3", "SOD2", "RARA", "IER3", "CD14", "PDCD4", "BNIP3L", "SQSTM1", "GPX3", "CD44", "TXNIP") |
| 19 | HALLMARK_INTERFERON_GAMMA_RESPONSE | 0.070428 | 0.178228 | -0.386476 | -1.368153 | 698 | 35 | c("IFITM3", "C1S", "CFB", "IFI30", "C1R", "MT2A", "STAT3", "IFI27", "B2M", "CD74", "NFKBIA", "SERPING1", "IFITM2") |
| 20 | HALLMARK_HEDGEHOG_SIGNALING | 0.071291 | 0.178228 | -0.54156 | -1.429028 | 630 | 9 | c("THY1", "NRP1", "DPYSL2", "ADGRG1", "CELSR1", "MYH9") |
| 21 | HALLMARK_IL2_STAT5_SIGNALING | 0.082615 | 0.196702 | -0.357803 | -1.319031 | 823 | 47 | c("IFITM3", "NRP1", "CD81", "BHLHE40", "LRIG1", "S100A1", "MYO1C", "PHLDA1", "AHR", "SPP1", "CAPG", "IGF1R", "CTSZ", "ECM1", "MAPKAPK2", "ANXA4", "ITGAV", "MAP3K8", "SERPINB6", "ENPP1", "PLEC", "EMP1", "ALCAM", "GPX4", "BCL2", "CYFIP1", "NFKBIZ") |
| 22 | HALLMARK_INTERFERON_ALPHA_RESPONSE | 0.090624 | 0.205964 | -0.441576 | -1.382673 | 866 | 18 | c("IFITM3", "C1S", "IFI30", "IFI27", "B2M", "CD74", "IFITM2", "LGALS3BP") |
| 23 | HALLMARK_P53_PATHWAY | 0.10836 | 0.235565 | -0.349213 | -1.283858 | 1080 | 46 | c("NUPR1", "TM4SF1", "CTSD", "CD81", "HMOX1", "IFI30", "JUN", "S100A10", "HEXIM1", "PLK2", "TOB1", "FOS", "MDM2", "DCXR", "ZFP36L1", "DDIT4", "PLXNB2", "EPS8L2", "SDC1", "FUCA1", "EPHX1", "RACK1") |
| 24 | HALLMARK_TNFA_SIGNALING_VIA_NFKB | 0.23187 | 0.483062 | -0.302774 | -1.15096 | 2317 | 61 | c("HES1", "CCN1", "NR4A1", "BHLHE40", "JUN", "SERPINE1", "CCND1", "EGR1", "EIF1", "PHLDA1", "PMEPA1", "NFKBIA", "GEM", "FOSB", "SDC4", "PLK2", "TGIF1", "FOS", "EFNA1", "DUSP1", "TNC", "MAP3K8", "FOSL2", "PLPP3", "DUSP4", "MARCKS", "PNRC1", "SOCS3", "JUNB", "INHBA", "PLAUR", "PLAU", "SOD2", "IER3", "VEGFA", "SQSTM1", "B4GALT1", "CD44", "KLF9", "BCL3") |
| 25 | HALLMARK_ALLOGRAFT_REJECTION | 0.247998 | 0.495995 | -0.362278 | -1.186031 | 2414 | 22 | c("THY1", "FLNA", "TIMP1", "TPD52", "B2M", "CD74", "CAPG") |
| 26 | HALLMARK_MITOTIC_SPINDLE | 0.326794 | 0.545056 | -0.293175 | -1.09915 | 3264 | 54 | c("FLNB", "FLNA", "FARP1", "PALLD", "GSN", "SHROOM1", "WASL", "ARL8A", "SPTAN1", "CTTN", "KIF5B", "ARHGEF12", "NUMA1", "YWHAE", "EZR", "MYH9", "DYNLL2", "CAPZB", "MARCKS", "NET1", "PCM1", "PREX1") |
| 27 | HALLMARK_TGF_BETA_SIGNALING | 0.321668 | 0.545056 | -0.33145 | -1.117033 | 3161 | 26 | c("CDH1", "SERPINE1", "THBS1", "PMEPA1", "TGIF1", "CTNNB1", "ENG", "SKIL", "RHOA", "FNTA", "LTBP2", "JUNB", "NCOR2", "BMPR2", "PPP1CA") |
| 28 | HALLMARK_MTORC1_SIGNALING | 0.30928 | 0.545056 | -0.299225 | -1.108281 | 3085 | 49 | c("CD9", "NUPR1", "SCD", "BHLHE40", "IFI30", "STC1", "SYTL2", "CANX", "SLC9A3R1", "DHCR24", "IGFBP5", "ATP2A2", "GSK3B", "LGMN", "PPIA", "DDIT4", "ITGB2", "PSMB5", "PIK3R3", "CALR", "ADD3", "USO1", "SERPINH1", "EGLN3") |
| 29 | HALLMARK_FATTY_ACID_METABOLISM | 0.327213 | 0.545056 | -0.33278 | -1.114346 | 3207 | 25 | c("BMPR1B", "UGDH", "MIF", "FASN", "DHCR24", "S100A10", "PCBD1", "HSP90AA1", "FH") |
| 30 | HALLMARK_UV_RESPONSE_UP | 0.337935 | 0.545056 | -0.310884 | -1.100551 | 3353 | 35 | c("NR4A1", "HMOX1", "RET", "EPCAM", "MMP14", "NFKBIA", "FOSB", "FOS", "CREG1", "SELENOW", "PPT1", "POLR2H", "EPHX1", "JUNB", "IGFBP2", "ATP6V1F", "SOD2", "GRINA", "HNRNPU", "SQSTM1", "GPX3", "EIF5", "RHOB") |
| 31 | HALLMARK_KRAS_SIGNALING_DN | 0.333222 | 0.545056 | -0.411676 | -1.11763 | 2992 | 10 | c("BMPR1B", "EFHD1", "SELENOP", "SKIL", "IDUA", "IGFBP2") |
| 32 | HALLMARK_WNT_BETA_CATENIN_SIGNALING | 0.389065 | 0.589492 | -0.518887 | -1.066009 | 2952 | 4 | c("CTNNB1", "NCOR2", "CSNK1E", "NCSTN") |
| 33 | HALLMARK_ADIPOGENESIS | 0.385124 | 0.589492 | -0.284101 | -1.062699 | 3846 | 53 | c("C3", "HSPB8", "SPARCL1", "UQCR11", "COL4A1", "GPAT4", "SOD1", "PHLDB1", "VEGFB", "CD151", "CMBL", "APOE", "APLP2", "TOB1", "COX6A1", "ALDH2", "NDUFAB1", "CCNG2", "UBC", "GPX4", "CAVIN1", "REEP5", "UQCRQ", "UQCRC1") |
| 34 | HALLMARK_PI3K_AKT_MTOR_SIGNALING | 0.4375 | 0.631229 | 0.168093 | 1.006033 | 55 | 29 | c("GRK2", "CDKN1B", "PFN1", "PLA2G12A", "ACTR3", "ACACA", "SMAD2", "TNFRSF1A", "ARHGDIA", "RAC1", "ARPC3", "GRB2", "PAK4", "ITPR2", "MKNK2", "MAPK1", "AKT1", "UBE2D3", "SQSTM1", "STAT2", "PPP1CA", "CALR", "PIK3R3", "DUSP3", "CAB39", "VAV3", "ARF1", "GSK3B", "CFL1") |
| 35 | HALLMARK_E2F_TARGETS | 0.44186 | 0.631229 | 0.19293 | 0.991694 | 151 | 20 | c("ANP32E", "TFRC", "HMGB2", "CDKN1B", "NAA38", "STAG1", "TOP2A", "SSRP1", "PA2G4", "ILF3", "MCM7", "SNRPB", "NOP56", "NUDT21", "HNRNPD", "POLD2", "PRKDC", "RBBP7", "NME1") |

| | | | | | | | | |
|---|---|---|---|---|---|---|---|---|
| 36 | HALLMARK_PROTEIN_SECRETION | 0.472443 | 0.644641 | -0.286444 | -1.01403 | 4688 | 35 | c("TMED2", "SEC31A", "PAM", "SOD1", "COPE", "TPD52", "ARF1", "NAPA", "RAB2A", "COPB2", "PPT1", "ARFGEF2", "CD63", "TOM1L1", "ERGIC3") |
| 37 | HALLMARK_SPERMATOGENESIS | 0.477034 | 0.644641 | -0.452183 | -0.996782 | 3769 | 5 | c("GSTM3", "SLC12A2", "PEBP1") |
| 38 | HALLMARK_INFLAMMATORY_RESPONSE | 0.522787 | 0.687878 | -0.289891 | -0.976974 | 5138 | 26 | c("TIMP1", "SLC7A2", "SERPINE1", "AHR", "MMP14", "NFKBIA", "ATP2A2") |
| 39 | HALLMARK_NOTCH_SIGNALING | 0.539939 | 0.692229 | -0.36131 | -0.953398 | 4778 | 9 | c("HES1", "CCND1", "NOTCH3") |
| 40 | HALLMARK_BILE_ACID_METABOLISM | 0.60521 | 0.756513 | -0.33286 | -0.903657 | 5435 | 10 | c("SOD1", "DHCR24", "LONP2", "HSD17B4", "ALDH9A1", "SCP2", "OPTN", "SLC29A1") |
| 41 | HALLMARK_IL6_JAK_STAT3_SIGNALING | 0.631574 | 0.770212 | -0.276879 | -0.896311 | 6124 | 21 | c("CD9", "HMOX1", "JUN", "STAT3", "A2M") |
| 42 | HALLMARK_MYC_TARGETS_V1 | 0.683377 | 0.813544 | -0.241098 | -0.890819 | 6815 | 48 | c("HSP90AB1", "CANX", "CLNS1A", "CCT3", "PPIA", "SNRPD2", "YWHAE", "PSMD3", "ILF2", "NDUFAB1", "TUFM", "IMPDH2", "SF3B3", "CNBP", "RACK1", "FAM120A", "NME1", "HNRNPC", "PCBP1", "PSMA7", "RNPS1", "POLD2", "HNRNPA2B1", "NHP2", "HNRNPU", "APEX1", "HNRNPD") |
| 43 | HALLMARK_G2M_CHECKPOINT | 0.768841 | 0.817916 | -0.235879 | -0.80513 | 7579 | 28 | c("CCND1", "NCL", "MT2A", "HSPA8", "KIF5B", "SS18", "NUMA1", "SLC12A2") |
| 44 | HALLMARK_UNFOLDED_PROTEIN_RESPONSE | 0.759424 | 0.817916 | -0.233605 | -0.816198 | 7534 | 32 | c("STC2", "SEC31A", "YWHAZ", "EIF4G1", "ATP6V0D1", "EIF4A2", "KIF5B", "LSM4", "DDIT4", "ATF4", "CALR", "LSM1") |
| 45 | HALLMARK_OXIDATIVE_PHOSPHORYLATION | 0.7275 | 0.817916 | -0.227485 | -0.875795 | 7274 | 69 | c("ATP1B1", "COX6C", "ATP5F1B", "UQCR11", "MAOB", "PDK4", "NDUFC2", "ATP6V0C", "ATP6AP1", "ATP5F1D", "NDUFB2", "UQCRB", "NDUFA4", "COX6A1", "COX7C", "FH", "NDUFA1", "NDUFS4", "NDUFAB1", "CYB5A", "CYB5R3", "NDUFC1", "ATP5MC2", "COX7A2", "GPX4", "UQCRQ", "UQCRC1", "ATP6V0B", "ATP6V1F") |
| 46 | HALLMARK_REACTIVE_OXYGEN_SPECIES_PATHWAY | 0.747626 | 0.817916 | -0.261914 | -0.799283 | 7085 | 16 | c("SOD1", "NQO1", "GPX4", "JUNB", "EGLN2", "FTL", "SOD2", "NDUFS2", "ATOX1", "GPX3", "TXNRD1", "GSR", "PTPA") |
| 47 | HALLMARK_PANCREAS_BETA_CELLS | 0.736935 | 0.817916 | -0.367103 | -0.809235 | 5823 | 5 | c("AKT3", "MAFB", "SPCS1") |
| 48 | HALLMARK_PEROXISOME | 0.793426 | 0.826486 | -0.240379 | -0.761554 | 7627 | 19 | c("CRABP2", "SOD1", "DHCR24", "SEMA3C", "CNBP", "LONP2", "HSD17B4", "SOD2", "ALDH9A1", "SCP2", "ELOVL5", "FIS1") |
| 49 | HALLMARK_DNA_REPAIR | 0.966788 | 0.986518 | -0.160817 | -0.558766 | 9576 | 31 | c("BCAM", "TMED2") |
| 50 | HALLMARK_HEME_METABOLISM | 0.998397 | 0.998397 | -0.114768 | -0.427338 | 9965 | 51 | c("C3", "BCAM") |

| Cluster 21 | pathway | pval | padj | ES | NES | nMoreExtreme | size | leadingEdge |
|---|---|---|---|---|---|---|---|---|
| 1 | HALLMARK_ESTROGEN_RESPONSE_EARLY | 0.000108 | 0.001139 | -0.716802 | -2.562917 | 0 | 78 | c("NBL1", "STC2", "KRT19", "PGR", "SEC14L2", "MUC1", "RET", "TFF3", "PRSS23", "CCND1", "CXCL12", "ELF3", "GFRA1", "TFF1", "MAPT", "KRT8", "ADCY1", "SLC7A2", "GJA1", "IGFBP4", "SCNN1A", "GREB1", "RHOBTB3", "MAST4", "CA12", "TTC39A", "HSPB8", "FASN", "PLAAT3", "SLC39A6", "THSD4", "RHOD", "FHL2", "MYB", "LRIG1", "GAB2", "FLNB", "XBP1", "SEMA3B", "HES1", "IGF1R", "MLPH", "FOS", "CLDN7", "FAM102A", "SLC9A3R1", "MED13L", "TOB1", "SIAH2") |
| 2 | HALLMARK_ESTROGEN_RESPONSE_LATE | 0.000109 | 0.001139 | -0.637459 | -2.254908 | 0 | 71 | c("CXCL14", "NBL1", "KRT19", "SERPINA3", "PGR", "RET", "TFF3", "PRSS23", "LTF", "CCND1", "TSPAN13", "CXCL12", "AGR2", "TFF1", "MAPT", "PRLR", "SCUBE2", "CPE", "IGFBP4", "SCNN1A", "CA12", "CDH1", "HSPB8", "PLAAT3", "MYB", "MDK", "DLG5", "FLNB", "XBP1", "TOP2A", "SEMA3B", "FOS", "FAM102A", "SLC9A3R1", "TOB1", "COX6C", "SIAH2", "LLGL2", "PDCD4", "UGDH", "ELOVL5", "LSR", "NRIP1", "DNAJC1") |
| 3 | HALLMARK_MYOGENESIS | 0.000114 | 0.001139 | -0.620746 | -2.09063 | 0 | 50 | c("COL6A3", "COL3A1", "COL1A1", "STC2", "AEBP1", "TNNT1", "APOD", "ADAM12", "PVALB", "IGFBP3", "CLU", "CNN3", "COL6A2", "SPDEF", "COL4A2", "ERBB3", "HSPB8", "TPM2", "IGFBP7", "TAGLN") |
| 4 | HALLMARK_EPITHELIAL_MESENCHYMAL_TRANSITIO N | 0.000104 | 0.001139 | -0.724418 | -2.686332 | 0 | 111 | c("COL5A1", "BGN", "MGP", "COL6A3", "POSTN", "COL3A1", "COL5A2", "COL1A1", "COL1A2", "THBS1", "MXRA5", "DCN", "COL11A1", "CCN1", "LOX", "CCN2", "LUM", "COMP", "THY1", "SFRP4", "COL16A1", "THBS2", "FSTL1", "TNC", "CDH11", "FBLN1", "PDGFRB", "PRRX1", "FN1", "CXCL12", "COL12A1", "NNMT", "FBLN2", "FBN1", "CTHRC1", "LOXL2", "SDC1", "ADAM12", "ELN", "PCOLCE", "CALD1", "GPC1", "IGFBP3", "GJA1", "COL6A2", "IGFBP4", "MYL9", "ACTA2", "FAP", "HTRA1", "LRRC15", "COL4A1", "COL4A2", "TIMP3", "IGFBP2", "TPM2", □ "GEM", "LOXL1", "INHBA", "PMEPA1", "RHOB", "TAGLN") |
| 5 | HALLMARK_UV_RESPONSE_DN | 0.000113 | 0.001139 | -0.565652 | -1.91433 | 0 | 51 | c("COL3A1", "COL5A2", "COL1A1", "COL1A2", "COL11A1", "CCN1", "INPP4B", "PDGFRB", "IGFBP5", "GJA1", "EFEMP1", "FHL2", "ERBB2", "PIK3R3", "RBPMS", "DUSP1", "AKT3", "APBB2", "IGF1R") |
| 6 | HALLMARK_IL6_JAK_STAT3_SIGNALING | 0.000534 | 0.004449 | 0.510217 | 2.102666 | 0 | 30 | c("HMOX1", "CCR1", "TNFRSF21", "PIK3R5", "CSF1", "TNFRSF1B", "CD36", "CD14", "IFNGR1", "IL17RA", "MYD88", "IL2RG", "CSF3R", "ITGB3", "TGFB1", "PIM1", "STAT2", "IRF1", "GRB2", "STAT1") |
| 7 | HALLMARK_INFLAMMATORY_RESPONSE | 0.000952 | 0.005291 | 0.48185 | 2.349185 | 0 | 57 | c("MARCO", "CCL2", "CXCL8", "CCRL2", "AQP9", "PLAUR", "CD48", "ABCA1", "PTAFR", "MSR1", "CMKLR1", "CLEC5A", "CYBB", "PIK3R5", "CD82", "SLC31A2", "ADM", "FPR1", "C5AR1", "IL10RA", "CSF1", "EMP3", "TNFRSF1B", "LYN", "ICAM1", "CD14", "C3AR1", "PTPRE", "NMI", "CSF3R", "ITGB3", "TAPBP", "ATP2A2", "LCP2", "ADRM1", "RHOG", "SLC11A2", "STAB1") |
| 8 | HALLMARK_BILE_ACID_METABOLISM | 0.000813 | 0.005291 | 0.629896 | 2.264827 | 1 | 19 | c("CYP27A1", "AQP9", "ABCA1", "NPC1", "HSD3B7", "SLC23A2", "ABCD1", "ALDH1A1", "ACSL1", "HSD17B4") |
| 9 | HALLMARK_ALLOGRAFT_REJECTION | 0.000849 | 0.005291 | 0.424095 | 2.017191 | 0 | 51 | c("CCR1", "CCL2", "CAPG", "ITGB2", "CTSS", "CD4", "BCAT1", "FCGR2B", "FGR", "SPI1", "C2", "IRF8", "MMP9", "CSF1", "WAS", "LYN", "ICAM1", "IFNGR1", "IL2RG", "ELF4", "B2M", "CD74", "PSMB10", "TAPBP", "SRGN", "TGFB1", "LCP2", "HCLS1", "STAB1", "NCK1", "MTIF2", "TAP2", "NCF4", "STAT1", "TAP1") |
| 10 | HALLMARK_COAGULATION | 0.003781 | 0.018903 | -0.506208 | -1.695436 | 32 | 48 | c("THBS1", "COMP", "PLAT", "PRSS23", "FN1", "MMP11", "CFB", "FBN1", "CLU", "S100A1", "CRIP2", "HTRA1", "TIMP3", "C1R", "C1S", "SERPING1", "C3", "S100A13", "SPARC") |
| 11 | HALLMARK_ANGIOGENESIS | 0.007967 | 0.036211 | -0.624643 | -1.719254 | 58 | 17 | c("POSTN", "COL3A1", "COL5A2", "LUM", "FSTL1", "STC1", "FGFR1") |
| 12 | HALLMARK_INTERFERON_GAMMA_RESPONSE | 0.009479 | 0.039494 | 0.301983 | 1.553484 | 7 | 70 | c("IFI30", "CCL2", "IL18BP", "UPP1", "IRF5", "CMKLR1", "SOD2", "PSME2", "IRF8", "FPR1", "IL10RA", "OAS3", "ICAM1", "SAMHD1", "MYD88", "LAP3", "TNFAIP2", "B2M", "IFIH1", "CD74", "CIITA", "PSMB10", "NMI", "TAPBP", "PIM1", "ZNFX1", "LCP2", "IFIT3", "IFI35", "OAS2", "PSMB8", "CASP4", "VAMP8", "STAT2", "TRIM14", "IRF2", "IRF1", "STAT1", "TAP1", "APOL6", "FGL2", "PTPN6", "MX1", "NLRC5", "UBE2L6", "CASP8") |

| | | | | | | | | |
|---|---|---|---|---|---|---|---|---|
| 13 | HALLMARK_COMPLEMENT | 0.011283 | 0.043398 | 0.275707 | 1.462634 | 7 | 79 | c("PLA2G7", "APOC1", "CTSL", "LIPA", "CTSD", "CTSS", "FDX1", "CTSB", "LGMN", "LGALS3", "FCER1G", "PLEK", "PLAUR", "CPM", "ME1", "C2", "PIK3R5", "WAS", "CTSH", "LYN", "CD36", "LTA4H", "ITGAM", "ANXA5", "CEBPB", "LAP3", "CTSC", "ATOX1", "GNB4", "KYNU", "ADAM9", "PIM1", "CALM3", "SRC", "GNAI2", "C1QA", "LCP2", "PRKCD", "RHOG", "SH2B3", "STX4", "CASP4", "LAMP2", "DOCK4", "IRF2", "IRF1", "GRB2", "PFN1", "PRCP") |
| 14 | HALLMARK_APICAL_JUNCTION | 0.01916 | 0.068427 | -0.447238 | -1.53349 | 170 | 56 | c("THY1", "COL16A1", "CDH11", "PARVA", "FBN1", "MYL9", "CLDN4", "JUP", "CDH1", "PCDH1", "MDK", "PIK3R3", "AKT3", "LIMA1", "CLDN7", "CERCAM", "MMP2", "VCAN") |
| 15 | HALLMARK_HYPOXIA | 0.0285 | 0.094999 | -0.414824 | -1.463879 | 260 | 70 | c("COL5A1", "BGN", "STC2", "DCN", "CCN1", "LOX", "CCN2", "GPC1", "IGFBP3", "EFNA1", "CA12", "STC1", "SELENBP1", "CAVIN1", "TGFB3", "TPD52", "DUSP1", "FOS", "SIAH2", "DTNA", "CCNG2", "MT2A") |
| 16 | HALLMARK_ANDROGEN_RESPONSE | 0.042855 | 0.126388 | -0.449812 | -1.470101 | 364 | 41 | c("KRT19", "AZGP1", "INPP4B", "CCND1", "KRT8", "BMPR1B", "SPDEF", "PMEPA1", "TPD52", "STEAP4", "UAP1", "SELENOP") |
| 17 | HALLMARK_INTERFERON_ALPHA_RESPONSE | 0.043658 | 0.126388 | 0.34338 | 1.463369 | 73 | 34 | c("IFI30", "CCRL2", "TMEM140", "PSME2", "CSF1", "OAS1", "LAP3", "B2M", "IFIH1", "CD74", "NMI", "IFIT3", "IFI35", "PSMB8", "STAT2", "TRIM14", "IRF2", "IRF1", "TAP1", "MX1", "UBE2L6", "CASP8") |
| 18 | HALLMARK_P53_PATHWAY | 0.0455 | 0.126388 | 0.273981 | 1.353474 | 45 | 60 | c("HMOX1", "IFI30", "GM2A", "CTSD", "FUCA1", "TCN2", "UPP1", "CEBPA", "RAP2B", "DRAM1", "NUPR1", "SAT1", "CD82", "VWA5A", "IRAK1") |
| 19 | HALLMARK_PI3K_AKT_MTOR_SIGNALING | 0.068905 | 0.181328 | 0.318861 | 1.376148 | 116 | 35 | c("MKNK1", "DUSP3", "SLC2A1", "RIT1", "MYD88", "SQSTM1", "CXCR4", "IL2RG", "ACTR3", "RALB", "MAPKAP1", "CFL1", "CDK4", "ARHGDIA", "NCK1", "STAT2", "GRB2", "PFN1", "SLA", "ARPC3", "CALR") |
| 20 | HALLMARK_TGF_BETA_SIGNALING | 0.081494 | 0.203734 | -0.46998 | -1.416852 | 645 | 26 | c("THBS1", "CDH1", "PMEPA1", "LTBP2", "CTNNB1", "ID3", "FNTA", "SKIL", "SMAD3", "BMPR2", "SERPINE1", "JUNB") |
| 21 | HALLMARK_XENOBIOTIC_METABOLISM | 0.106897 | 0.254516 | 0.263387 | 1.260952 | 123 | 52 | c("HMOX1", "CYP27A1", "APOE", "CES1", "BCAT1", "UPP1", "AQP9", "NPC1", "PGD", "POR", "ACP2", "IRF8", "PTGDS", "SLC6A6", "FBP1", "CD36", "ABCC3", "CNDP2") |
| 22 | HALLMARK_MTORC1_SIGNALING | 0.118712 | 0.269801 | 0.248136 | 1.23505 | 117 | 61 | c("IFI30", "ITGB2", "LGMN", "BCAT1", "HK2", "GLRX", "NUPR1", "ME1", "TXNRD1", "TOMM40", "M6PR", "G6PD", "SLC6A6", "GLA", "ENO1", "LTA4H", "SLC2A1", "RIT1", "CTSC", "SQSTM1", "CXCR4", "ACTR3") |
| 23 | HALLMARK_APICAL_SURFACE | 0.135751 | 0.293586 | -0.559762 | -1.345701 | 916 | 10 | c("GSTM3", "THY1", "GATA3") |
| 24 | HALLMARK_GLYCOLYSIS | 0.145592 | 0.293586 | -0.373664 | -1.274863 | 1292 | 54 | c("COL5A1", "STC2", "DCN", "TFF3", "ELF3", "SDC1", "GPC1", "IGFBP3", "STC1", "FUT8", "CHPF", "EGLN3", "VCAN", "IDUA", "DDIT4", "AGRN") |
| 25 | HALLMARK_REACTIVE_OXYGEN_SPECIES_PATHWAY | 0.146793 | 0.293586 | 0.338102 | 1.289563 | 324 | 23 | c("GLRX", "FTL", "TXNRD1", "SOD2", "LSP1", "G6PD", "MBP", "SRXN1", "ATOX1", "MGST1", "PRNP", "PRDX1", "ABCC1") |
| 26 | HALLMARK_APOPTOSIS | 0.168923 | 0.324853 | -0.349467 | -1.233239 | 1546 | 70 | c("BGN", "DCN", "LUM", "PDGFRB", "PLAT", "CCND1", "CLU", "TIMP3", "ERBB3", "RHOB", "ERBB2", "TOP2A", "CTNNB1", "IFITM3", "BTG2", "MMP2", "CREBBP", "PDCD4") |
| 27 | HALLMARK_NOTCH_SIGNALING | 0.195743 | 0.362486 | -0.605329 | -1.263645 | 1222 | 6 | c("CCND1", "NOTCH3", "HES1") |
| 28 | HALLMARK_KRAS_SIGNALING_UP | 0.247856 | 0.440357 | -0.340199 | -1.17274 | 2224 | 58 | c("SERPINA3", "PLAT", "SPARCL1", "PRRX1", "APOD", "MMP11", "TSPAN13", "CFB", "CPE", "IGFBP3", "JUP", "INHBA") |
| 29 | HALLMARK_KRAS_SIGNALING_DN | 0.255407 | 0.440357 | -0.476243 | -1.20469 | 1794 | 12 | c("EFHD1", "BMPR1B", "IGFBP2", "SELENOP", "BTG2", "IDUA", "SKIL") |
| 30 | HALLMARK_HEDGEHOG_SIGNALING | 0.335081 | 0.558468 | -0.496994 | -1.126719 | 2172 | 8 | c("THY1", "ADGRG1", "LDB1", "VEGFA", "MYH9", "CELSR1") |
| 31 | HALLMARK_PEROXISOME | 0.357032 | 0.575858 | 0.259059 | 1.057469 | 687 | 29 | c("SOD2", "HSD3B7", "SLC23A2", "ABCD1", "CLN6", "ALDH1A1", "ACSL1", "HSD17B4") |
| 32 | HALLMARK_G2M_CHECKPOINT | 0.409644 | 0.64007 | -0.339759 | -1.052402 | 3329 | 30 | c("CCND1", "SLC38A1", "TOP2A", "SLC12A2", "MT2A", "HNRNPU", "SMC4", "NUMA1", "SMAD3", "MYC") |
| 33 | HALLMARK_TNFA_SIGNALING_VIA_NFKB | 0.432958 | 0.655998 | -0.28993 | -1.035202 | 4003 | 77 | c("CCN1", "TNC", "CCND1", "FOSB", "DUSP4", "EFNA1", "PLK2", "NR4A1", "GEM", "EGR1", "INHBA", "PMEPA1", "RHOB", "DUSP1", "HES1", "FOS", "BTG2", "SOCS3", "EIF1", "KLF2") |
| 34 | HALLMARK_IL2_STAT5_SIGNALING | 0.472648 | 0.69507 | 0.196043 | 0.991083 | 431 | 65 | c("CAPG", "PLIN2", "CTSZ", "SPP1", "SMPDL3A", "HK2", "CD48", "IGF2R", "CAPN3", "TNFRSF21", "IRF8", "IL10RA", "CSF1", "TNFRSF1B") |
| 35 | HALLMARK_HEME_METABOLISM | 0.494454 | 0.706363 | -0.290222 | -0.995111 | 4412 | 56 | c("HBB", "HBD", "BCAM", "SLC4A1", "SELENBP1", "PRDX2", "ARHGEF12", "C3", "SLC25A37", "BTG2", "FBXO9") |
| 36 | HALLMARK_CHOLESTEROL_HOMEOSTASIS | 0.542028 | 0.73247 | 0.252126 | 0.93542 | 1250 | 21 | c("LGMN", "LGALS3", "PLAUR", "FABP5", "PPARG", "CXCL16", "ATF5", "ANXA5") |

| | | | | | | | | |
|---|---|---|---|---|---|---|---|---|
| 37 | HALLMARK_SPERMATOGENESIS | 0.535019 | 0.73247 | -0.485834 | -0.963569 | 3215 | 5 | c("GSTM3", "SLC12A2") |
| 38 | HALLMARK_WNT_BETA_CATENIN_SIGNALING | 0.635883 | 0.836689 | -0.421307 | -0.879493 | 3972 | 6 | c("CTNNB1", "CSNK1E", "MYC") |
| 39 | HALLMARK_PANCREAS_BETA_CELLS | 0.664643 | 0.852106 | -0.463206 | -0.859595 | 3830 | 4 | c("AKT3", "SRP9") |
| 40 | HALLMARK_MITOTIC_SPINDLE | 0.726044 | 0.88542 | -0.243167 | -0.829634 | 6447 | 54 | c("PALLD", "ARHGAP29", "FLNB", "TOP2A", "NET1", "SHROOM1", "ARHGEF12", "CTTN", "GSN", "SMC4", "FARP1", "NUMA1", "DST", "WASL", "CYTH2", "SPTAN1", "RANBP9", "ARFIP2", "EZR", "YWHAE", "MYH9") |
| 41 | HALLMARK_UV_RESPONSE_UP | 0.720456 | 0.88542 | -0.250411 | -0.825074 | 6187 | 43 | c("RET", "EPCAM", "FOSB", "IGFBP2", "NR4A1", "RHOB", "FOS", "BTG2") |
| 42 | HALLMARK_E2F_TARGETS | 0.748219 | 0.890737 | -0.268785 | -0.783994 | 5776 | 22 | c("TOP2A", "RBBP7", "MCM7", "POLD2", "SMC4", "HMGB2", "MYC", "NME1", "LUC7L3", "NUDT21") |
| 43 | HALLMARK_MYC_TARGETS_V1 | 0.795262 | 0.924723 | -0.231725 | -0.766102 | 6847 | 44 | c("CLNS1A", "IMPDH2", "CBX3", "GNL3", "HSP90AB1", "ILF2", "HNRNPU", "MCM7", "POLD2", "RACK1", "MYC", "RNPS1", "NHP2", "YWHAE", "LSM7", "CCT3", "BUB3", "NME1") |
| 44 | HALLMARK_PROTEIN_SECRETION | 0.815929 | 0.927192 | 0.184439 | 0.786018 | 1382 | 34 | c("ABCA1", "IGF2R", "M6PR", "CD63", "PPT1", "GLA", "CTSC", "STX7", "CLTA", "SNX2", "ATP6V1H", "LAMP2", "SEC22B", "STX12", "VAMP3", "SOD1", "AP3B1", "KIF1B", "VPS4B", "YKT6", "COPE") |
| 45 | HALLMARK_ADIPOGENESIS | 0.904054 | 0.972976 | -0.195691 | -0.689036 | 8272 | 68 | c("LAMA4", "SPARCL1", "COL4A1", "HSPB8", "CAVIN1", "FABP4", "PHLDB1", "C3", "TOB1", "CMBL", "GPAT4", "CCNG2", "PDCD4") |
| 46 | HALLMARK_UNFOLDED_PROTEIN_RESPONSE | 0.953517 | 0.972976 | -0.185371 | -0.577547 | 7794 | 31 | c("STC2", "XBP1", "EEF2", "DDIT4", "SHC1", "DNAJA4", "FUS", "NHP2", "VEGFA", "LSM1", "BANF1") |
| 47 | HALLMARK_MYC_TARGETS_V2 | 0.941575 | 0.972976 | 0.197587 | 0.589913 | 2916 | 11 | c("HK2", "CDK4", "HSPE1", "PA2G4", "IMP4", "FARSA", "SRM", "NOP16") |
| 48 | HALLMARK_FATTY_ACID_METABOLISM | 0.922944 | 0.972976 | 0.160431 | 0.67735 | 1604 | 33 | c("IL4I1", "ME1", "MGLL", "ALDH1A1", "ACSL1", "CD36", "HSD17B4") |
| 49 | HALLMARK_OXIDATIVE_PHOSPHORYLATION | 0.913183 | 0.972976 | 0.151631 | 0.761679 | 851 | 64 | c("FDX1", "POR", "ATP6V0C", "TCIRG1", "ATP6V0B", "TIMM8B", "ATP6V1C1", "ATP5F1E", "ATP6V1D", "ATP6V1H", "ATP6V1G1", "ETFB", "ATP6V1F", "GPX4", "DLD", "ATP6AP1", "SDHB", "NDUFS3", "COX7A2", "BAX", "ECHS1", "ATP5PD", "ATP6V1E1", "NDUFS7", "ACAT1", "TIMM13", "ACO2", "ATP5MG", "NDUFB6", "HSPA9", "PHB2", "ATP5PF", "IMMT", "NDUFS6", "NDUFB8") |
| 50 | HALLMARK_DNA_REPAIR | 0.999634 | 0.999634 | -0.105877 | -0.331658 | 8199 | 32 | c("BCAM", "IMPDH2", "NME3", "BRF2", "NME4", "POLR2H", "CANT1", "NELFCD", "NME1", "VPS28", "NUDT21", "POLR2I", "DDB1", "NFX1", "NCBP2", "AAAS", "EIF1B", "APRT", "POLR2K", "STX3", "POLR2E", "ITPA", "RALA", "GPX4", "HCLS1", "ADRM1", "ELOA", "AGO4", "SDCBP", "POLE4", "BCAP31") |

| Cluster 22 | pathway | pval | padj | ES | NES | nMoreExtreme | size | leadingEdge |
|---|---|---|---|---|---|---|---|---|
| 1 | HALLMARK_COMPLEMENT | 0.0001 | 0.001006 | -0.558291 | -1.933614 | 0 | 48 | c("C1S", "C1R", "COL4A2", "C1QA", "APOC1", "SERPINE1", "C3", "MMP14", "LTF", "C1QC", "PLAT", "LGMN", "SERPING1", "CTSL", "CTSD", "CD36", "LIPA", "LRP1") |
| 2 | HALLMARK_EPITHELIAL_MESENCHYMAL_TRANSITION | 1.00E-04 | 0.001006 | -0.707616 | -2.62161 | 0 | 102 | c("HTRA1", "COL5A2", "CCN2", "SFRP4", "CCN1", "FBN1", "COL4A1", "COL4A2", "FSTL1", "COL11A1", "COL12A1", "CDH11", "ELN", "THY1", "PDGFRB", "ADAM12", "PCOLCE", "INHBA", "NNMT", "DST", "SERPINE1", "LOXL1", "DPYSL3", "LAMC1", "LOX", "THBS2", "FAP", "COMP", "CTHRC1", "CALD1", "GEM", "MMP14", "TPM2", "COL16A1", "FBLN2", "CXCL12", "GJA1", "ACTA2", "IGFBP3", "PRRX1", "LOXL2", "JUN", "PMEPA1", "FBLN1", "DAB2", "TNC", "PMP22", "TAGLN", "FUCA1", "SERPINH1", "LRRC15", "ITGAV", "TGFBI", "MYL9", "TNFRSF12A", "MXRA5", "PFN2", "LRP1") |
| 3 | HALLMARK_UV_RESPONSE_DN | 0.0001 | 0.001006 | -0.565646 | -1.950357 | 0 | 46 | c("COL5A2", "CCN1", "COL11A1", "PDGFRB", "SERPINE1", "NRP1", "LAMC1", "GJA1", "FHL2", "EFEMP1", "PLPP3", "DAB2", "PMP22", "ANXA4", "SDC2", "CITED2", "IGF1R", "AKT3", "RUNX1", "RBPMS") |
| 4 | HALLMARK_COAGULATION | 0.0001 | 0.001006 | -0.672779 | -2.308431 | 0 | 44 | c("HTRA1", "MMP11", "CTSK", "C1S", "FBN1", "C1R", "C1QA", "CRIP2", "APOC1", "SERPINE1", "COMP", "S100A1", "PLAU", "C3", "A2M", "MMP14", "PLAT", "MMP9", "LGMN", "SERPING1", "GSN") |
| 5 | HALLMARK_KRAS_SIGNALING_UP | 0.000101 | 0.001006 | -0.586696 | -1.982913 | 0 | 39 | c("MMP11", "INHBA", "GPNMB", "NRP1", "APOD", "TMEM176B", "PLAU", "SPARCL1", "IGFBP3", "ENG", "PRRX1", "PLAT", "MMP9", "FUCA1", "SERPINA3", "ANKH", "EMP1", "SEMA3B", "CPE", "MAFB") |
| 6 | HALLMARK_ESTROGEN_RESPONSE_EARLY | 0.0007 | 0.005834 | -0.467903 | -1.684468 | 6 | 71 | c("SLC7A2", "ADCY1", "RHOD", "CXCL12", "GJA1", "PGR", "FHL2", "SLC2A1", "MAPT", "PAPSS2", "SEC14L2", "NBL1", "RHOBTB3", "IGF1R", "TTC39A", "CA12", "FARP1", "HSPB8", "GREB1", "SCNN1A", "FOS", "SEMA3B", "MYOF", "ABHD2", "RET", "CELSR1", "BHLHE40", "CANT1", "MAST4", "CHPT1", "SH3BP5", "UGCG", "MYC", "LRIG1", "NRIP1", "ELOVL5") |
| 7 | HALLMARK_MYOGENESIS | 0.002007 | 0.014339 | -0.490872 | -1.688003 | 19 | 45 | c("COL4A2", "ADAM12", "APOD", "GPX3", "TPM2", "PVALB", "SORBS3", "IGFBP3", "NQO1", "TAGLN", "GSN", "IGFBP7", "CD36", "TNNT1", "CNN3", "HSPB8", "AGRN", "PDLIM7", "BHLHE40", "PLXNB2", "COL1A1", "AKT2", "SCD") |
| 8 | HALLMARK_TNFA_SIGNALING_VIA_NFKB | 0.003202 | 0.020014 | -0.450886 | -1.598244 | 31 | 60 | c("CCN1", "NR4A1", "INHBA", "SERPINE1", "GEM", "PLAU", "EGR1", "FOSB", "JUN", "PMEPA1", "PLPP3", "PHLDA1", "TNC", "PLK2", "SGK1", "IER3", "MAP3K8", "MARCKS", "KLF9", "FOS", "EFNA1") |
| 9 | HALLMARK_HYPOXIA | 0.01691 | 0.084551 | -0.415181 | -1.477861 | 168 | 63 | c("CCN2", "CCN1", "SERPINE1", "PAM", "LOX", "SLC2A1", "CAVIN1", "IGFBP3", "HMOX1", "JUN", "SDC3", "TGFB3", "SDC2", "CITED2", "TGFBI", "CA12", "IER3", "DDIT4", "FOS", "EFNA1") |
| 10 | HALLMARK_ESTROGEN_RESPONSE_LATE | 0.016405 | 0.084551 | -0.409333 | -1.466389 | 163 | 67 | c("CXCL14", "LTF", "CXCL12", "PGR", "MAPT", "CDH1", "PAPSS2", "NBL1", "SERPINA3", "MDK", "SGK1", "CA12", "FARP1", "HSPB8", "SCNN1A", "FOS", "SEMA3B", "SCUBE2", "MYOF", "CPE", "ABHD2", "RET", "PRLR", "LSR", "IDH2", "CHPT1", "UGDH", "NRIP1", "ELOVL5", "ATP2B4", "PRSS23", "CD9", "BCL2", "ID2", "NCOR2", "DLG5", "SERPINA1") |
| 11 | HALLMARK_ANDROGEN_RESPONSE | 0.028416 | 0.126301 | -0.443139 | -1.490838 | 281 | 38 | c("BMPR1B", "MAF", "ELL2", "PMEPA1", "SELENOP", "ITGAV", "SGK1", "ANKH", "ARID5B", "ABHD2", "ACSL3", "ACTN1", "TPD52", "DBI", "SCD", "ELOVL5", "INPP4B", "B4GALT1", "ADRM1") |
| 12 | HALLMARK_IL2_STAT5_SIGNALING | 0.030312 | 0.126301 | -0.424665 | -1.460329 | 301 | 45 | c("NRP1", "S100A1", "PHLDA1", "ANXA4", "IGF1R", "ITGAV", "MAP3K8", "NFKBIZ", "EMP1", "CD81", "BHLHE40", "SPP1", "ENPP1", "MYC", "LRIG1", "BMPR2", "CKAP4", "CAPG", "ITGA6", "AHR", "CYFIP1", "BCL2", "CTSZ", "RNH1", "HIPK2") |
| 13 | HALLMARK_APICAL_JUNCTION | 0.033941 | 0.130541 | -0.407211 | -1.428603 | 338 | 54 | c("FBN1", "CDH11", "THY1", "CD276", "COL16A1", "SORBS3", "CDH1", "SDC3", "MMP9", "MDK", "TSPAN4", "AKT3", "TGFBI", "MYL9", "PARVA", "CERCAM", "CTNND1") |
| 14 | HALLMARK_XENOBIOTIC_METABOLISM | 0.041767 | 0.149168 | -0.417816 | -1.429079 | 415 | 43 | c("SERPINE1", "TMEM176B", "HMOX1", "PAPSS2", "FBLN1", "PDK4", "NQO1", "CD36", "CYB5A", "EPHX1", "ALDH2", "COMT", "PGRMC1", "BCAR1", "MT2A", "UGDH", "MAN1A1", "ELOVL5", "ARPP19", "CNDP2", "CAT", "MCCC2") |

| | | | | | | | | |
|---|---|---|---|---|---|---|---|---|
| 15 | HALLMARK_MYC_TARGETS_V1 | 0.047619 | 0.15873 | 0.244556 | 1.793388 | 0 | 49 | c("CNBP", "FAM120A", "SMARCC1", "HNRNPU", "EIF4A1", "PPIA", "HNRNPA2B1", "NOP56", "RNPS1", "HNRNPD", "HDGF", "PA2G4", "CBX3", "SRSF2", "CLNS1A", "APEX1", "EEF1B2", "DHX15", "CYC1", "HSP90AB1", "EIF3B", "BUB3", "TUFM", "POLD2", "PCBP1", "PSMA7", "SRM", "SNRPD2", "HSPE1", "SF3B3", "TXNL4A", "IMPDH2", "GNL3", "NCBP2", "NDUFAB1", "HNRNPC", "NME1", "HNRNPA3", "EIF4H", "LSM7", "MCM7", "YWHAE", "ILF2", "NHP2", "CANX") |
| 16 | HALLMARK_TGF_BETA_SIGNALING | 0.065738 | 0.205432 | -0.451622 | -1.43002 | 641 | 25 | c("SERPINE1", "ENG", "CDH1", "PMEPA1", "LTBP2", "ID3", "THBS1", "SKI", "BMPR2", "SKIL", "FNTA", "ID2", "NCOR2", "HIPK2", "CTNNB1", "JUNB", "TGIF1", "RAB31", "ARID4B") |
| 17 | HALLMARK_ANGIOGENESIS | 0.093175 | 0.274044 | -0.487624 | -1.400614 | 870 | 15 | c("COL5A2", "FSTL1", "NRP1", "ITGAV", "FGFR1", "SPP1", "COL3A1", "STC1", "VEGFA") |
| 18 | HALLMARK_GLYCOLYSIS | 0.103583 | 0.287731 | -0.381036 | -1.3103 | 1031 | 45 | c("PAM", "IGFBP3", "SDC3", "EGLN3", "SDC2", "CITED2", "TGFBI", "IER3", "CHPF", "DDIT4", "AGRN", "CYB5A", "SDC1", "GFPT1", "PLOD1", "PKM", "ARPP19", "B4GALT1", "COL5A1", "ELF3", "STC1", "VEGFA", "AKR1A1", "IDUA") |
| 19 | HALLMARK_APOPTOSIS | 0.157199 | 0.35727 | -0.357529 | -1.241271 | 1568 | 49 | c("PDGFRB", "GPX3", "HMOX1", "JUN", "PLAT", "GSN", "TNFRSF12A", "IER3", "ANKH", "ANXA1", "EMP1", "LGALS3") |
| 20 | HALLMARK_HEDGEHOG_SIGNALING | 0.155772 | 0.35727 | -0.518901 | -1.310976 | 1343 | 9 | c("THY1", "NRP1", "DPYSL2", "CELSR1") |
| 21 | HALLMARK_E2F_TARGETS | 0.152542 | 0.35727 | 0.224518 | 1.27285 | 35 | 25 | c("TOP2A", "SNRPB", "NOP56", "HNRNPD", "HMGB2", "PA2G4", "LUC7L3", "SRSF2", "NAA38", "RBBP7", "PNN", "POLD2", "SSRP1", "NUDT21", "PRKDC", "SMC4", "ILF3", "STAG1", "TUBB", "NME1", "MCM7", "CDKN1B") |
| 22 | HALLMARK_REACTIVE_OXYGEN_SPECIES_PATHWAY | 0.136533 | 0.35727 | -0.457442 | -1.334343 | 1285 | 16 | c("MPO", "GPX3", "NQO1") |
| 23 | HALLMARK_INFLAMMATORY_RESPONSE | 0.166021 | 0.360916 | -0.396167 | -1.271167 | 1628 | 27 | c("SLC7A2", "INHBA", "SERPINE1", "MMP14", "ITGA5", "STAB1", "MYC", "BTG2", "AHR", "SRI", "ADRM1", "CYBB", "BST2", "NFKBIA", "PLAUR") |
| 24 | HALLMARK_IL6_JAK_STAT3_SIGNALING | 0.213947 | 0.414343 | -0.397305 | -1.230963 | 2073 | 22 | c("A2M", "HMOX1", "JUN", "TNFRSF12A", "CD36", "MAP3K8") |
| 25 | HALLMARK_ADIPOGENESIS | 0.215459 | 0.414343 | -0.338297 | -1.184575 | 2151 | 53 | c("COL4A1", "FABP4", "LAMA4", "GPX3", "PHLDB1", "C3", "SPARCL1", "CAVIN1") |
| 26 | HALLMARK_UV_RESPONSE_UP | 0.206155 | 0.414343 | -0.363571 | -1.214619 | 2042 | 36 | c("NR4A1", "GPX3", "MMP14", "FOSB", "HMOX1", "UROD", "FOS", "EPHX1", "RET", "TFRC", "BTG2", "BSG", "CLTB", "YKT6", "CREG1") |
| 27 | HALLMARK_APICAL_SURFACE | 0.241534 | 0.416438 | -0.509045 | -1.204684 | 1989 | 7 | c("THY1", "B4GALT1", "SULF2", "PLAUR") |
| 28 | HALLMARK_P53_PATHWAY | 0.230754 | 0.416438 | -0.34016 | -1.178128 | 2301 | 48 | c("HMOX1", "JUN", "FUCA1", "S100A10", "CTSD", "PLK2", "IER3", "DDIT4", "FOS", "CD81", "EPHX1", "SDC1", "RAP2B", "SEC61A1", "PLXNB2", "IFI30", "BTG2", "IP6K2", "NUPR1") |
| 29 | HALLMARK_KRAS_SIGNALING_DN | 0.234687 | 0.416438 | -0.455732 | -1.218342 | 2114 | 11 | c("BMPR1B", "SELENOP", "SGK1", "BTG2", "SKIL", "IDUA", "IGFBP2") |
| 30 | HALLMARK_NOTCH_SIGNALING | 0.282908 | 0.471513 | 0.338532 | 1.128593 | 431 | 8 | c("NOTCH2", "APH1A", "ARRB1", "HES1", "RBX1", "LFNG") |
| 31 | HALLMARK_WNT_BETA_CATENIN_SIGNALING | 0.373791 | 0.584049 | -0.510942 | -1.092891 | 2860 | 5 | c("CSNK1E", "MYC", "NCOR2", "CTNNB1") |
| 32 | HALLMARK_MYC_TARGETS_V2 | 0.369257 | 0.584049 | 0.337427 | 1.054108 | 650 | 7 | c("NOP56", "PA2G4", "CBX3", "SRM", "HSPE1", "GNL3") |
| 33 | HALLMARK_FATTY_ACID_METABOLISM | 0.458622 | 0.674444 | -0.320664 | -1.028905 | 4499 | 27 | c("BMPR1B", "UROD", "S100A10", "CD36", "EPHX1", "PCBD1", "HSD17B4", "UGDH", "ELOVL5") |
| 34 | HALLMARK_ALLOGRAFT_REJECTION | 0.456482 | 0.674444 | -0.324098 | -1.026227 | 4457 | 25 | c("THY1", "INHBA", "MMP9") |
| 35 | HALLMARK_INTERFERON_GAMMA_RESPONSE | 0.503033 | 0.679774 | -0.299022 | -0.991398 | 4975 | 34 | c("C1S", "C1R", "SERPING1", "ARID5B", "IFI27", "STAT3", "IFI30", "MT2A", "SOCS3", "SRI", "STAT2", "BST2", "NFKBIA", "LGALS3BP", "CFB") |
| 36 | HALLMARK_MTORC1_SIGNALING | 0.49985 | 0.679774 | -0.287608 | -0.998519 | 4988 | 49 | c("SLC2A1", "EGLN3", "LGMN", "SERPINH1", "SYTL2", "DDIT4", "TFRC", "ACSL3", "BHLHE40", "IFI30", "BTG2", "GGA2", "SCD", "ELOVL5", "NUPR1", "PIK3R3", "YKT6", "M6PR", "CD9", "STC1", "CANX", "ITGB2", "SERP1", "ATP2A2", "USO1", "ENO1") |
| 37 | HALLMARK_HEME_METABOLISM | 0.498899 | 0.679774 | -0.283196 | -0.997951 | 4985 | 56 | c("C3", "ACP5", "SLC2A1", "ELL2", "SLC30A1", "UROD", "UBAC1", "SLC4A1", "TFRC", "OPTN", "CIR1", "CAST", "ARHGEF12", "BCAM", "BTG2", "BSG") |
| 38 | HALLMARK_CHOLESTEROL_HOMEOSTASIS | 0.537053 | 0.706648 | -0.320045 | -0.958362 | 5130 | 18 | c("LGMN", "TNFRSF12A", "SEMA3B", "LGALS3", "SCD", "CD9", "CTNNB1", "PLAUR", "ECH1", "ANXA5") |
| 39 | HALLMARK_INTERFERON_ALPHA_RESPONSE | 0.559785 | 0.717673 | -0.330219 | -0.935675 | 5205 | 14 | c("C1S", "IFI27", "IFI30", "STAT2", "BST2", "LGALS3BP") |
| 40 | HALLMARK_PANCREAS_BETA_CELLS | 0.617324 | 0.771655 | -0.418693 | -0.895574 | 4724 | 5 | c("AKT3", "MAFB") |
| 41 | HALLMARK_BILE_ACID_METABOLISM | 0.66989 | 0.816939 | -0.308906 | -0.842713 | 6095 | 12 | c("OPTN", "HSD17B4", "IDH2", "SCP2", "CAT") |

| | | | | | | | | |
|---|---|---|---|---|---|---|---|---|
| 42 | HALLMARK_G2M_CHECKPOINT | 0.738462 | 0.879121 | 0.136744 | 0.841236 | 95 | 31 | c("TOP2A", "SMARCC1", "HNRNPU", "PAFAH1B1", "YTHDC1", "SLC38A1", "CUL3", "HSPA8", "NOTCH2", "HNRNPD", "KIF5B", "TOP1", "BCL3", "SRSF2", "BUB3", "ATRX", "SMC4", "SMAD3", "ILF3", "STAG1", "NUMA1", "CDKN1B", "PRPF4B", "NCL", "CUL4A", "SS18", "MYC", "RBM14", "MT2A", "ABL1") |
| 43 | HALLMARK_MITOTIC_SPINDLE | 0.881229 | 0.974725 | -0.20748 | -0.733085 | 8806 | 58 | c("PALLD", "DST", "SHROOM1", "GSN", "FARP1", "MARCKS", "ARHGAP29", "TRIO", "ABL1", "RAB3GAP1", "BCAR1", "PCM1", "ABR", "ARHGEF12") |
| 44 | HALLMARK_DNA_REPAIR | 0.948658 | 0.974725 | -0.173217 | -0.569224 | 9367 | 32 | c("TMED2", "ERCC1", "SEC61A1", "CANT1", "BCAM", "POLR2I", "ADRM1", "NME3", "APRT", "POLR2E", "BRF2", "NME1", "POLR2A", "SUPT5H", "CSTF3", "NCBP2", "POLR2H", "NELFCD", "IMPDH2", "NME4", "NUDT21", "DCTN4", "SSRP1", "POLD4", "RBX1", "HCLS1") |
| 45 | HALLMARK_PROTEIN_SECRETION | 0.947225 | 0.974725 | -0.175603 | -0.586655 | 9386 | 36 | c("DST", "PAM", "TMED2", "TPD52", "SNX2", "ANP32E", "YKT6", "M6PR", "CD63", "MON2", "LAMP2", "COPE", "USO1", "STX7", "PPT1", "COPB2", "LMAN1", "ERGIC3", "RER1") |
| 46 | HALLMARK_UNFOLDED_PROTEIN_RESPONSE | 0.858329 | 0.974725 | -0.215953 | -0.709664 | 8475 | 32 | c("DNAJA4", "DDIT4", "HERPUD1", "BANF1", "SRPRA", "EIF4G1", "DCTN1", "VEGFA", "SHC1", "NHP2", "LSM1", "EIF4EBP1", "ATP6V0D1", "SERP1", "ATF4") |
| 47 | HALLMARK_PI3K_AKT_MTOR_SIGNALING | 0.954248 | 0.974725 | 0.104845 | 0.621095 | 145 | 29 | c("PFN1", "UBE2D3", "TNFRSF1A", "CXCR4", "ARF1", "GSK3B", "MYD88", "RAC1", "ACACA", "AKT1", "ACTR3", "CALR", "MAPK1", "SMAD2", "ITPR2", "GRB2", "ARPC3", "PLA2G12A", "DUSP3", "ARHGDIA", "PAK4", "CFL1", "MKNK2", "CDKN1B") |
| 48 | HALLMARK_PEROXISOME | 0.95523 | 0.974725 | -0.174996 | -0.542186 | 9259 | 22 | c("ERCC1", "HSD17B4", "IDH2", "ELOVL5", "SCP2", "CAT") |
| 49 | HALLMARK_SPERMATOGENESIS | 0.895353 | 0.974725 | 0.280176 | 0.664624 | 2446 | 4 | c("GSTM3", "PEBP1", "TALDO1") |
| 50 | HALLMARK_OXIDATIVE_PHOSPHORYLATION | 0.9987 | 0.9987 | -0.109581 | -0.3932 | 9984 | 68 | c("PDK4", "ATP1B1", "MAOB", "CYB5A", "NDUFB6", "NDUFS4", "UQCRB", "CYB5R3", "IDH2", "VDAC3", "SLC25A6", "COX4I1", "UQCR11", "NDUFB7", "UQCR10", "ECH1", "ATP6V0E1", "UQCRQ", "ATP5F1D", "NDUFC1", "RHOT2", "COX6C", "NDUFAB1", "ISCU", "ATP5F1B", "COX7A2", "NDUFA3", "TIMM13", "PDHA1", "ECI1", "NDUFB2", "COX7C", "IDH3B", "NDUFS7", "COX7A2L", "HADHA", "NDUFS6", "MGST3", "ATP6V0C", "NDUFC2", "NDUFA4", "ATP6AP1", "ACADVL", "CYC1", "ATP5MC2", "NDUFA2", "ATP5PD", "ATP5MF", "ECHS1", "COX6A1", "ATP6V1F", "NDUFV2", □ "UQCRC1", "NDUFA1", "LRPPRC", "COX8A", "NDUFB8", "ATP6V0B", "ETFB", "ACAA1") |

| Cluster 23 | pathway | pval | padj | ES | NES | nMoreExtreme | size | leadingEdge |
|---|---|---|---|---|---|---|---|---|
| 1 | HALLMARK_TNFA_SIGNALING_VIA_NFKB | 0.000297 | 0.001722 | -0.534423 | -2.826305 | 0 | 66 | c("FOSB", "PHLDA1", "SERPINE1", "EGR1", "CCN1", "DUSP1", "FOS", "NR4A1", "MAP3K8", "GEM", "EIF1", "CD44", "PLAUR", "KLF6", "MYC", "VEGFA", "SOCS3", "JUN", "JUNB", "NFE2L2", "NFKBIA", "EFNA1", "PNRC1", "FOSL2", "PLAU", "TNC", "BTG1", "CEBPD", "PLK2", "KLF2", "SGK1") |
| 2 | HALLMARK_HYPOXIA | 0.000296 | 0.001722 | -0.482497 | -2.624921 | 0 | 74 | c("IGFBP3", "SERPINE1", "CCN1", "DUSP1", "FOS", "SDC2", "STC2", "LOX", "BGN", "PLAUR", "KLF6", "VEGFA", "CXCR4", "COL5A1", "JUN", "TPD52", "CCN2", "WSB1", "MT2A", "CAVIN1", "SLC2A1", "EFNA1", "DCN", "TGFBI", "PNRC1", "CITED2", "SIAH2", "FOSL2", "CCNG2", "BTG1", "ANXA2") |
| 3 | HALLMARK_APOPTOSIS | 0.000294 | 0.001722 | -0.355597 | -1.876987 | 0 | 65 | c("ANXA1", "EMP1", "LGALS3", "CD44", "CTNNB1", "BGN", "TOP2A", "MMP2", "DPYD", "ANKH", "JUN", "GPX3", "TIMP3", "LUM", "TXNIP", "DCN", "IGF2R", "CLU", "RARA", "CREBBP", "CFLAR", "GSN", "PPT1", "SLC20A1", "IER3", "IFITM3", "ERBB2", "SAT1") |
| 4 | HALLMARK_COMPLEMENT | 0.000283 | 0.001722 | -0.483179 | -2.420858 | 0 | 52 | c("APOC1", "SERPINE1", "CD36", "LGALS3", "MMP14", "PLAUR", "CTSD", "CD55", "LTF", "HSPA1A", "GNAI2", "LIPA", "S100A9", "LRP1", "CTSB", "SERPINA1", "CTSS", "LGMN", "SERPING1", "ANXA5", "FCER1G", "CD46", "CLU", "FN1") |
| 5 | HALLMARK_EPITHELIAL_MESENCHYMAL_TRANSITIO N | 0.00031 | 0.001722 | -0.677461 | -3.940058 | 0 | 101 | c("IGFBP3", "SERPINE1", "ACTA2", "CCN1", "POSTN", "ADAM12", "MGP", "DPYSL3", "GEM", "LOX", "MXRA5", "ITGA5", "COL6A2", "THY1", "SPP1", "CD44", "MMP14", "COL16A1", "BGN", "PLAUR", "LAMC1", "ITGB1", "MMP2", "TPM4", "MYL9", "CTHRC1", "VEGFA", "COL6A3", "NNMT", "COL5A1", "JUN", "PCOLCE", "COPA", "CCN2", "LOXL2", "DAB2", "PRRX1", "VCAN", "COL5A2", "FAP", "GJA1", "LRP1", "THBS2", "TIMP3", "LUM", "SFRP4", "TAGLN", "LRRC15", "DCN", "TGFBI", "COL11A1", "CDH11", "COL1A1", "QSOX1", "LGALS1", "HTRA1", "FBLN1", □ "PFN2", "TNC", "PPIB", "FBLN2", "CALD1", "SDC1", "WIPF1", "FN1", "ELN", "ITGAV", "THBS1", "PMP22", "SPARC") |
| 6 | HALLMARK_ANGIOGENESIS | 0.000254 | 0.001722 | -0.699542 | -2.319206 | 0 | 13 | c("POSTN", "SPP1", "VEGFA", "NRP1", "VCAN", "COL5A2", "LUM") |
| 7 | HALLMARK_COAGULATION | 0.000272 | 0.001722 | -0.586481 | -2.745317 | 0 | 39 | c("APOC1", "SERPINE1", "S100A1", "ANXA1", "MMP9", "MMP14", "MMP2", "MMP11", "CTSK", "LRP1", "TIMP3", "CTSB", "SERPINA1", "LGMN", "SERPING1", "HTRA1", "PLAU", "CLU", "FN1", "THBS1", "SPARC", "S100A13", "GSN") |
| 8 | HALLMARK_IL2_STAT5_SIGNALING | 0.000281 | 0.001722 | -0.408159 | -2.028677 | 0 | 50 | c("PHLDA1", "S100A1", "MAP3K8", "EMP1", "SPP1", "MUC1", "CD44", "KLF6", "NFKBIZ", "MYC", "NRP1", "COL6A1", "ITGA6") |
| 9 | HALLMARK_KRAS_SIGNALING_UP | 0.000279 | 0.001722 | -0.558629 | -2.716287 | 0 | 46 | c("IGFBP3", "GPNMB", "SPARCL1", "MMP9", "TMEM176B", "EMP1", "SPP1", "CPE", "PLAUR", "MMP11", "CXCR4", "ITGB2", "ANKH", "NRP1", "LAPTM5", "SERPINA3", "PRRX1", "SEMA3B", "APOD", "CTSS", "PLAU", "FCER1G") |
| 10 | HALLMARK_SPERMATOGENESIS | 0.00066 | 0.0033 | 0.627316 | 1.979096 | 3 | 14 | c("HSPA2", "SPATA6", "PEBP1", "MLLT10", "MAST2", "MAP7", "TALDO1", "NF2", "SIRT1", "PHKG2", "STRBP") |
| 11 | HALLMARK_UV_RESPONSE_DN | 0.001144 | 0.005202 | -0.35552 | -1.853504 | 3 | 60 | c("SERPINE1", "CCN1", "DUSP1", "SDC2", "LAMC1", "MYC", "RUNX1", "NRP1", "DAB2", "COL5A2", "GJA1", "EFEMP1", "SRI", "ANXA4", "COL11A1", "COL1A1", "CITED2", "RBPMS", "ANXA2", "ADD3", "INPP4B", "PMP22", "AKT3", "TGFBR2", "BHLHE40") |
| 12 | HALLMARK_ESTROGEN_RESPONSE_EARLY | 0.002711 | 0.011295 | -0.312869 | -1.730499 | 8 | 79 | c("FOS", "PAPSS2", "KRT19", "RHOBTB3", "STC2", "MUC1", "CD44", "PGR", "AMFR", "HSPB8", "GREB1", "MED13L", "MYC", "TOB1", "GJA1", "SEMA3B", "SLC2A1") |
| 13 | HALLMARK_ESTROGEN_RESPONSE_LATE | 0.003012 | 0.011585 | -0.308064 | -1.703924 | 9 | 79 | c("FOS", "PAPSS2", "KRT19", "CPE", "CD44", "PGR", "AMFR", "HSPB8", "TOP2A", "LTF", "TOB1", "SERPINA3", "S100A9", "SEMA3B", "CDH1", "SERPINA1", "PRLR", "SIAH2", "IDH2", "ADD3", "COX6C", "SGK1", "CHPT1", "TSPAN13", "MAPT", "IGFBP4", "ZFP36", "AFF1", "RAB31", "DLG5", "PLAAT3", "CXCL12", "NBL1", "FARP1") |
| 14 | HALLMARK_MYOGENESIS | 0.003675 | 0.013127 | -0.361272 | -1.783308 | 12 | 49 | c("IGFBP3", "ADAM12", "CD36", "STC2", "COL6A2", "HSPB8", "ITGB1", "COL6A3", "AEBP1", "GPX3", "TAGLN", "APOD", "COL1A1", "NQO1", "PVALB", "IGFBP7", "CLU", "SCD", "SPARC", "PDLIM7", "GSN", "BHLHE40") |
| 15 | HALLMARK_CHOLESTEROL_HOMEOSTASIS | 0.004678 | 0.015593 | -0.502211 | -1.881649 | 17 | 19 | c("LGALS3", "CTNNB1", "PLAUR", "ACTG1", "SEMA3B", "PNRC1", "LGMN", "ANXA5", "S100A11", "CLU") |
| 16 | HALLMARK_INFLAMMATORY_RESPONSE | 0.006143 | 0.019197 | -0.397524 | -1.806529 | 22 | 35 | c("SERPINE1", "ITGA5", "MMP14", "PLAUR", "KLF6", "CD55", "MYC", "IL1R1", "NFKBIA", "SRI") |

| | | | | | | | | |
|---|---|---|---|---|---|---|---|---|
| 17 | HALLMARK_TGF_BETA_SIGNALING | 0.016729 | 0.046468 | -0.406592 | -1.710632 | 62 | 27 | c("SERPINE1", "CTNNB1", "SKIL", "JUNB", "CDH1", "RHOA", "PPP1CA", "THBS1", "ID3", "SLC20A1", "RAB31", "FNTA") |
| 18 | HALLMARK_NOTCH_SIGNALING | 0.016727 | 0.046468 | 0.774878 | 1.59911 | 96 | 4 | c("CCND1", "RBX1", "HES1", "ARRB1") |
| 19 | HALLMARK_REACTIVE_OXYGEN_SPECIES_PATHWAY | 0.031819 | 0.083735 | -0.455356 | -1.650319 | 123 | 17 | c("FTL", "EGLN2", "JUNB", "GPX3", "MPO", "NQO1") |
| 20 | HALLMARK_APICAL_JUNCTION | 0.036364 | 0.090909 | -0.271455 | -1.456072 | 123 | 69 | c("MMP9", "ACTB", "THY1", "COL16A1", "ACTG1", "ITGB1", "MMP2", "MYL9", "GNAI2", "CD99", "VCAN", "EXOC4", "CDH1", "CNN2", "TGFBI", "CDH11") |
| 21 | HALLMARK_IL6_JAK_STAT3_SIGNALING | 0.040806 | 0.097156 | -0.384962 | -1.557127 | 153 | 24 | c("CD36", "MAP3K8", "CD44", "SOCS3", "JUN", "IL1R1") |
| 22 | HALLMARK_XENOBIOTIC_METABOLISM | 0.053337 | 0.121221 | -0.298738 | -1.452586 | 190 | 46 | c("SERPINE1", "CD36", "PAPSS2", "TMEM176B", "APOE", "IL1R1", "MT2A", "NQO1", "PDK4", "FBLN1") |
| 23 | HALLMARK_GLYCOLYSIS | 0.057225 | 0.124401 | -0.271185 | -1.413824 | 199 | 60 | c("IGFBP3", "SDC2", "STC2", "CD44", "PPIA", "VEGFA", "CXCR4", "COL5A1", "VCAN", "DCN", "TGFBI", "EGLN3", "QSOX1", "CITED2") |
| 24 | HALLMARK_PANCREAS_BETA_CELLS | 0.073723 | 0.153589 | -0.596588 | -1.501759 | 302 | 6 | c("SPCS1", "SRP9", "SEC11A", "AKT3", "MAFB") |
| 25 | HALLMARK_UNFOLDED_PROTEIN_RESPONSE | 0.082086 | 0.164171 | -0.309894 | -1.397953 | 306 | 34 | c("STC2", "ATF4", "VEGFA", "FUS", "SPCS1", "SEC11A", "YWHAZ", "ATP6V0D1", "HERPUD1", "EIF4G1") |
| 26 | HALLMARK_PROTEIN_SECRETION | 0.114171 | 0.21956 | -0.299793 | -1.341233 | 426 | 33 | c("ARF1", "VPS45", "TPD52", "RAB2A", "IGF2R", "USO1", "CD63", "TMED2", "NAPA", "PPT1", "SNX2", "RER1", "ERGIC3") |
| 27 | HALLMARK_MYC_TARGETS_V1 | 0.120942 | 0.223967 | -0.25292 | -1.290512 | 420 | 56 | c("HSP90AB1", "PPIA", "MYC", "HNRNPC", "HNRNPU", "CBX3", "PSMD3", "SNRPD2", "YWHAE", "APEX1", "FAM120A", "NME1", "RACK1", "GNL3", "CANX", "CCT3", "HNRNPD", "PCBP1", "EIF4A1", "VDAC3", "RNPS1", "HNRNPA3") |
| 28 | HALLMARK_BILE_ACID_METABOLISM | 0.155025 | 0.27683 | 0.404151 | 1.304287 | 940 | 15 | c("FDXR", "HSD3B7", "PXMP2", "RXRA", "AMACR", "DHCR24", "GSTK1") |
| 29 | HALLMARK_WNT_BETA_CATENIN_SIGNALING | 0.172809 | 0.297947 | -0.484361 | -1.293594 | 707 | 7 | c("CTNNB1", "MYC") |
| 30 | HALLMARK_APICAL_SURFACE | 0.203067 | 0.327527 | -0.446344 | -1.249241 | 820 | 8 | c("THY1", "PLAUR") |
| 31 | HALLMARK_ALLOGRAFT_REJECTION | 0.196848 | 0.327527 | -0.267939 | -1.217636 | 736 | 35 | c("MMP9", "CD47", "THY1", "ITGB2", "TPD52", "B2M", "CTSS", "STAT1", "NME1", "ETS1", "SRGN") |
| 32 | HALLMARK_HEDGEHOG_SIGNALING | 0.224586 | 0.350915 | -0.434495 | -1.216079 | 907 | 8 | c("THY1", "VEGFA", "NRP1", "LDB1") |
| 33 | HALLMARK_ADIPOGENESIS | 0.246364 | 0.373279 | -0.212433 | -1.143596 | 829 | 71 | c("FABP4", "SPARCL1", "CD36", "HSPB8", "UBC", "MGST3", "TOB1", "APOE", "GPX3", "CAVIN1", "RTN3", "CMPK1", "LAMA4", "CCNG2", "CD151", "TKT", "COX6A1") |
| 34 | HALLMARK_ANDROGEN_RESPONSE | 0.295727 | 0.434893 | -0.22789 | -1.108095 | 1058 | 46 | c("STEAP4", "KRT19", "ANKH", "TPD52", "B2M", "SLC38A2", "ARID5B", "ACSL3", "INPP4B", "SGK1", "ITGAV", "SCD", "SMS") |
| 35 | HALLMARK_PI3K_AKT_MTOR_SIGNALING | 0.412074 | 0.588677 | 0.247075 | 1.035908 | 2586 | 36 | c("ITPR2", "SFN", "ARPC3", "PIK3R3", "TRAF2", "ECSIT", "MKNK2") |
| 36 | HALLMARK_HEME_METABOLISM | 0.435362 | 0.60467 | -0.193688 | -1.01482 | 1501 | 62 | c("HBB", "VEZF1", "HBD", "MGST3", "ACP5", "TNS1", "SLC2A1", "CTSB", "SLC4A1", "SNCA", "RBM5", "SLC25A37", "TFRC", "ARHGEF12", "LRP10", "PICALM", "OPTN", "GYPC") |
| 37 | HALLMARK_UV_RESPONSE_UP | 0.488021 | 0.659487 | -0.193834 | -0.97892 | 1710 | 54 | c("FOSB", "FOS", "NR4A1", "MMP14", "JUNB", "GPX3", "NFKBIA", "HNRNPU") |
| 38 | HALLMARK_G2M_CHECKPOINT | 0.589476 | 0.775627 | -0.201485 | -0.915639 | 2206 | 35 | c("TOP2A", "SLC38A1", "MYC", "MT2A", "HNRNPU", "PRPF4B") |
| 39 | HALLMARK_DNA_REPAIR | 0.60884 | 0.780564 | 0.210806 | 0.899209 | 3856 | 38 | c("NELFCD", "BCAM", "USP11", "ZNF707", "RBX1", "POLR2G", "POLB", "GTF2B", "CSTF3", "NCBP2", "NME3", "GPX4") |
| 40 | HALLMARK_MITOTIC_SPINDLE | 0.694396 | 0.846825 | -0.168044 | -0.863842 | 2403 | 58 | c("ARHGAP29", "SHROOM1", "TOP2A", "PCM1", "NET1", "ARHGAP4", "PALLD", "PREX1", "YWHAE") |
| 41 | HALLMARK_PEROXISOME | 0.691633 | 0.846825 | 0.212494 | 0.841139 | 4339 | 29 | c("DLG4", "HSD3B7", "ABCC5", "ACOX1", "NUDT19", "DHCR24", "GSTK1", "SEMA3C", "HSD11B2", "CLN6", "CRAT", "BCL10", "FIS1", "ECH1", "ECI2") |
| 42 | HALLMARK_P53_PATHWAY | 0.723033 | 0.860753 | -0.162625 | -0.847844 | 2526 | 60 | c("S100A10", "TM4SF1", "FOS", "CTSD", "TOB1", "JUN") |
| 43 | HALLMARK_INTERFERON_GAMMA_RESPONSE | 0.805398 | 0.926279 | -0.156087 | -0.793418 | 2834 | 55 | c("IFITM2", "SOCS3", "MT2A", "NFKBIA", "B2M", "SAMHD1", "TXNIP", "SRI", "STAT1", "SERPING1", "BTG1", "ARID5B") |
| 44 | HALLMARK_FATTY_ACID_METABOLISM | 0.833651 | 0.926279 | -0.160777 | -0.746599 | 3056 | 38 | c("S100A10", "CD36", "GLUL", "LGALS1", "APEX1", "FH") |
| 45 | HALLMARK_KRAS_SIGNALING_DN | 0.827816 | 0.926279 | -0.198585 | -0.719721 | 3225 | 17 | c("EFHD1", "SKIL") |
| 46 | HALLMARK_INTERFERON_ALPHA_RESPONSE | 0.947256 | 0.986725 | -0.138047 | -0.604217 | 3537 | 31 | c("CD47", "IFITM2", "B2M", "TXNIP") |
| 47 | HALLMARK_MTORC1_SIGNALING | 0.936391 | 0.986725 | 0.138262 | 0.656621 | 6123 | 58 | c("SLC7A11", "QDPR", "FDXR", "PIK3R3") |
| 48 | HALLMARK_MYC_TARGETS_V2 | 0.935585 | 0.986725 | 0.184181 | 0.594394 | 5678 | 15 | c("HSPE1", "RABEPK", "IMP4", "DCTPP1", "PA2G4", "WDR74", "PES1", "SORD", "NOP16", "CDK4", "FARSA", "SRM") |

| | | | | | | | | |
|---|---|---|---|---|---|---|---|---|
| 49 | HALLMARK_E2F_TARGETS | 0.983155 | 0.992793 | -0.11693 | -0.52313 | 3676 | 33 | c("TOP2A", "MYC", "NME1", "TUBB", "TFRC", "SNRPB", "HNRNPD", "PRKDC", "DDX39A", "SHMT1", "NUP153", "MCM7", "CDKN1B", "SRSF1", "TUBG1", "XRCC6", "RBBP7", "NBN", "CDK4", "STAG1", "POLD2", "CBX5", "PNN", "SSRP1", "DEK", "PA2G4", "PAN2", "NAA38", "DCTPP1", "RAD1", "CSE1L", "PPP1R8", "USP1") |
| 50 | HALLMARK_OXIDATIVE_PHOSPHORYLATION | 0.992793 | 0.992793 | 0.105597 | 0.534056 | 6611 | 78 | c("TIMM10", "COX8A", "IDH3A", "NDUFS6", "NDUFV2", "ETFB", "ALDH6A1", "NDUFB5", "CYB5A", "ACO2", "COX7A2", "ATP5F1B", "COX7A2L", "NDUFA7", "NDUFA4", "SDHA", "HTRA2", "NDUFC2", "GPX4", "GOT2", "ABCB7", "NDUFAB1", "SLC25A11", "GRPEL1") |

| Cluster 24 | pathway | pval | padj | ES | NES | nMoreExtreme | size | leadingEdge |
|---|---|---|---|---|---|---|---|---|
| 1 | HALLMARK_HYPOXIA | 0.00034 | 0.001889 | -0.439596 | -2.36245 | 0 | 73 | c("MIF", "STC2", "BGN", "DCN", "BHLHE40", "CCN2", "COL5A1", "CCN1", "CA12", "JUN", "SERPINE1", "ANXA2", "HMOX1", "TGFB3", "IGFBP3", "STC1", "TPD52", "MT2A", "TGFBI", "GAA", "GPC1", "PAM", "SDC4", "SDC2", "HDLBP", "DUSP1", "SIAH2", "MYH9", "SDC3", "CAVIN1", "EFNA1", "FBP1") |
| 2 | HALLMARK_CHOLESTEROL_HOMEOSTASIS | 0.000284 | 0.001889 | -0.603052 | -2.25691 | 0 | 19 | c("FASN", "ACTG1", "CD9", "SCD", "CLU", "LGALS3", "S100A11", "CTNNB1", "LGMN", "SEMA3B", "ANXA5") |
| 3 | HALLMARK_APOPTOSIS | 0.000332 | 0.001889 | -0.520519 | -2.712809 | 0 | 64 | c("TIMP3", "BGN", "LUM", "DCN", "MMP2", "CCND1", "HSPB1", "LMNA", "IFITM3", "RHOB", "PLAT", "CLU", "JUN", "TIMP2", "TIMP1", "GSN", "HMOX1", "GPX4", "PDGFRB", "ERBB2", "LGALS3", "SQSTM1", "SPTAN1", "RARA", "CTNNB1", "PEA15", "PPT1", "DAP", "CD44", "DAP3", "APP", "PDCD4", "EMP1") |
| 4 | HALLMARK_ESTROGEN_RESPONSE_EARLY | 0.000349 | 0.001889 | -0.684056 | -3.735158 | 0 | 78 | c("STC2", "FLNB", "KRT19", "MUC1", "NBL1", "SLC39A6", "MLPH", "HSPB8", "CCND1", "FASN", "SEC14L2", "KRT8", "TFF1", "BHLHE40", "MAPT", "PRSS23", "RET", "XBP1", "HES1", "CA12", "CELSR1", "SLC7A2", "LRIG1", "ADCY1", "GREB1", "PGR", "RHOBTB3", "GFRA1", "GAB2", "TOB1", "ELF3", "TFF3", "SCNN1A", "CLDN7", "FAM102A", "MYOF", "THSD4", "IGFBP4", "FARP1", "SLC9A3R1", "IGF1R", "FHL2", "RARA", "TTC39A", "GJA1", "SEMA3B", "NRIP1", "NCOR2", "CD44", "PAPSS2", "RHOD", "PLAAT3") |
| 5 | HALLMARK_ESTROGEN_RESPONSE_LATE | 0.00034 | 0.001889 | -0.588381 | -3.128858 | 0 | 70 | c("FLNB", "KRT19", "COX6C", "CXCL14", "NBL1", "SERPINA3", "HSPB8", "CCND1", "SCUBE2", "TFF1", "MAPT", "CD9", "PRSS23", "AGR2", "RET", "PRLR", "XBP1", "CA12", "DLG5", "UGDH", "PGR", "CPE", "TOB1", "CDH1", "TFF3", "SCNN1A", "FAM102A", "MYOF", "IGFBP4", "FARP1", "TSPAN13", "SLC9A3R1", "LSR", "MDK", "SEMA3B", "RABEP1", "NRIP1", "NCOR2", "CD44", "LLGL2", "PAPSS2", "PLAAT3", "SIAH2", "PDCD4", "DCXR") |
| 6 | HALLMARK_MYOGENESIS | 0.000322 | 0.001889 | -0.648005 | -3.240713 | 0 | 54 | c("STC2", "COL3A1", "COL6A2", "COL6A3", "AEBP1", "HSPB8", "APOD", "BHLHE40", "TAGLN", "PVALB", "CNN3", "IGFBP7", "SCD", "CLU", "SPARC", "GSN", "COL1A1", "SPDEF", "NQO1", "IGFBP3", "ITGB5", "COL4A2", "ADAM12", "TNNT1", "ITGB1", "GAA", "SPTAN1", "MYO1C", "ATP6AP1", "SYNGR2", "AGRN", "SORBS3", "PLXNB2", "APP") |
| 7 | HALLMARK_EPITHELIAL_MESENCHYMAL_TRANSITION | 0.000378 | 0.001889 | -0.738954 | -4.293055 | 0 | 105 | c("POSTN", "FN1", "MGP", "COL3A1", "COL6A2", "COL6A3", "TIMP3", "HTRA1", "BGN", "SPP1", "LUM", "ELN", "COMP", "DCN", "MMP2", "COL5A2", "FBLN1", "SFRP4", "THBS2", "ACTA2", "FBLN2", "CCN2", "LGALS1", "COL5A1", "CCN1", "COL1A2", "COL11A1", "RHOB", "MMP14", "TAGLN", "MXRA5", "SPARC", "JUN", "THY1", "CDH11", "TIMP1", "SERPINE1", "COL12A1", "COL4A1", "CTHRC1", "DPYSL3", "TPM4", "CALD1", "MYL9", "FSTL1", "COL1A1", "ECM1", "IGFBP3", "PDGFRB", "LOXL2", "ITGB5", "COL16A1", "COL4A2", "ADAM12", "FLNA", "FAP", □ "TGFBI", "LRP1", "IGFBP4", "LOXL1", "PRRX1", "ITGB1", "GPC1", "PMP22", "THBS1", "VIM", "GEM", "GJA1", "LAMC1", "SDC4", "PCOLCE", "CALU", "ITGAV", "COPA", "CAPG") |
| 8 | HALLMARK_OXIDATIVE_PHOSPHORYLATION | 0.000332 | 0.001889 | -0.440559 | -2.296081 | 0 | 64 | c("COX6C", "NDUFC2", "ATP1B1", "MAOB", "GPX4", "ATP5F1B", "COX7C", "ATP6AP1", "PDK4", "NDUFA4", "ATP6V0C", "COX6A1", "UQCR11", "FH", "UQCRQ", "UQCRB", "COX6B1", "CYB5A", "ATP6V1F", "NDUFAB1", "ATP5MC2", "NDUFB2", "ATP6V0E1", "NDUFA1", "ECHS1", "COX8A", "COX4I1", "MGST3", "UQCR10", "ATP5PD", "ATP5MF", "ECI1", "HADHA", "IDH2", "NDUFC1", "NDUFS4", "UQCRC1", "ATP5F1E", "CYB5R3") |
| 9 | HALLMARK_UV_RESPONSE_DN | 0.000323 | 0.001889 | -0.466098 | -2.297301 | 0 | 52 | c("COL3A1", "COL5A2", "BHLHE40", "CCN1", "COL1A2", "COL11A1", "EFEMP1", "SERPINE1", "ANXA2", "COL1A1", "PDGFRB", "ERBB2", "INPP4B", "NRP1", "IGF1R", "FHL2", "IGFBP5", "PMP22", "GJA1", "LAMC1", "RBPMS", "SDC2", "PIK3R3", "DUSP1") |
| 10 | HALLMARK_COAGULATION | 0.000313 | 0.001889 | -0.768435 | -3.63645 | 0 | 44 | c("FN1", "C3", "CFB", "TIMP3", "MMP11", "HTRA1", "COMP", "MMP2", "CTSB", "APOC1", "CD9", "PRSS23", "MMP14", "PLAT", "CLU", "SPARC", "S100A13", "TIMP1", "CTSK", "GSN", "SERPINE1", "A2M", "S100A1", "SERPING1", "C1R", "C1S", "LRP1", "CAPN2", "THBS1", "CRIP2", "C1QA", "LGMN", "CSRP1") |

| | | | | | | | | |
|---|---|---|---|---|---|---|---|---|
| 11 | HALLMARK_E2F_TARGETS | 0.000425 | 0.001931 | 0.405634 | 1.910143 | 2 | 69 | c("BRCA2", "CENPE", "LBR", "ESPL1", "CENPM", "HMMR", "AURKB", "CHEK2", "BIRC5", "CIT", "TCF19", "SPC24", "MXD3", "MKI67", "RRM2", "ANP32E", "CKS1B", "DCK", "TACC3", "HELLS", "POLE4", "EZH2", "PRPS1", "HMGA1", "HMGB2", "KIF22", "TFRC", "PLK1", "PCNA", "ASF1B", "MYBL2", "MLH1", "POLE", "TBRG4", "SMC6", "PNN", "SMC4", "MCM2", "HUS1", "MCM6", "CDKN1B", "STAG1", "TRA2B", "LIG1", "NUP107", "LMNB1", "MCM3", "DDX39A", "SRSF2", "LUC7L3", "PA2G4", "MYC") |
| 12 | HALLMARK_ANGIOGENESIS | 0.000562 | 0.002163 | -0.599989 | -2.213033 | 1 | 18 | c("POSTN", "COL3A1", "SPP1", "LUM", "COL5A2", "TIMP1", "FSTL1", "STC1", "NRP1", "FGFR1") |
| 13 | HALLMARK_HEME_METABOLISM | 0.000544 | 0.002163 | 0.353174 | 1.802052 | 3 | 104 | c("EPB42", "SPTA1", "NFE2", "HMBS", "ANK1", "KEL", "SLC4A1", "TRIM58", "KLF1", "ALAS2", "TMCC2", "ERMAP", "AHSP", "SPTB", "CA1", "CPOX", "CA2", "OSBP2", "SLC25A37", "SNCA", "HBD", "SYNJ1", "RANBP10", "DMTN", "USP15", "CCND3", "RCL1", "KAT2B", "ABCB6", "AGPAT4", "KDM7A", "HTRA2", "LMO2", "PPOX", "CAT", "BPGM", "TFRC", "SIDT2", "MAP2K3", "TENT5C", "PIGQ", "SLC2A1", "UROD", "FECH", "XPO7", "UBAC1", "RBM38", "MPP1", "ALAD", "SLC25A38", "LPIN2", "EPB41", "AQP3", "EZH1", "RBM5", "NARF") |
| 14 | HALLMARK_ADIPOGENESIS | 0.000662 | 0.002205 | -0.368948 | -1.887879 | 1 | 60 | c("C3", "HSPB8", "SPARCL1", "APOE", "COL4A1", "TOB1", "PHLDB1", "GPX4", "GPAT4", "CD151", "CMBL", "VEGFB", "COX6A1", "UQCR11", "UQCRQ", "UBC", "PDCD4", "REEP5", "NDUFAB1", "CAVIN1", "SOD1", "ECHS1", "COX8A", "CCNG2", "MGST3", "UQCR10", "APLP2") |
| 15 | HALLMARK_APICAL_JUNCTION | 0.00066 | 0.002205 | -0.408895 | -2.125247 | 1 | 63 | c("MMP2", "ACTG1", "PARVA", "JUP", "THY1", "CDH11", "CDH1", "MYL9", "CLDN7", "ACTB", "COL16A1", "EVL", "TGFBI", "CTNNA1", "ITGB1", "CD276", "CLDN4", "MDK", "NECTIN2", "CERCAM", "PIK3R3", "CTNND1", "SORBS3", "FBN1", "MYH9", "SDC3", "LIMA1", "WASL", "AKT3") |
| 16 | HALLMARK_ANDROGEN_RESPONSE | 0.000942 | 0.002943 | -0.423003 | -1.992047 | 2 | 43 | c("KRT19", "AZGP1", "CCND1", "KRT8", "BMPR1B", "SCD", "DHCR24", "SPDEF", "B2M", "DBI", "INPP4B", "TPD52", "SELENOP") |
| 17 | HALLMARK_COMPLEMENT | 0.001367 | 0.004011 | -0.338705 | -1.842874 | 3 | 76 | c("FN1", "C3", "GATA3", "CFB", "CTSB", "CTSD", "APOC1", "MMP14", "PLAT", "CLU", "S100A13", "TIMP2", "TIMP1", "SERPINE1", "SERPING1", "COL4A2", "C1R", "C1S", "LGALS3", "CTSL", "LRP1", "C1QC", "LIPA", "C1QA", "LGMN", "CSRP1") |
| 18 | HALLMARK_ALLOGRAFT_REJECTION | 0.001444 | 0.004011 | 0.41083 | 1.857932 | 9 | 57 | c("ELANE", "MAP4K1", "PRKCB", "TLR1", "CD96", "NCF4", "LTB", "ZAP70", "ACHE", "FGR", "IL4R", "WAS", "IL2RG", "IL16", "SRGN", "ITGAL", "LYN", "CCND3", "TLR2", "TRAF2", "ST8SIA4", "JAK2", "LCP2", "IRF8", "HCLS1", "TAP2", "ELF4", "FYB1", "PTPN6") |
| 19 | HALLMARK_XENOBIOTIC_METABOLISM | 0.00223 | 0.005868 | -0.389725 | -1.875907 | 6 | 47 | c("CFB", "FBLN1", "MCCC2", "JUP", "UGDH", "SERPINE1", "APOE", "SPINT2", "HMOX1", "NQO1", "MT2A", "ESR1", "IGFBP4", "PDK4", "EPHX1", "PAPSS2", "DDAH2", "DCXR", "CYB5A", "FBP1", "ATP2A2") |
| 20 | HALLMARK_GLYCOLYSIS | 0.002576 | 0.006394 | -0.378503 | -1.879792 | 7 | 53 | c("MIF", "STC2", "DCN", "COL5A1", "ELF3", "TFF3", "IGFBP3", "STC1", "TGFBI", "FUT8", "GPC1", "PAM", "IDUA", "SDC2", "PKM", "HDLBP", "CD44", "AGRN", "CYB5A", "SDC3", "IL13RA1", "SOD1", "SDC1", "DDIT4", "PPIA", "CHPF", "COPB2") |
| 21 | HALLMARK_IL2_STAT5_SIGNALING | 0.002685 | 0.006394 | -0.339746 | -1.783718 | 7 | 66 | c("MUC1", "COL6A1", "SPP1", "CD81", "BHLHE40", "IFITM3", "RHOB", "XBP1", "LRIG1", "S100A1", "GPX4", "PHLDA1", "ECM1", "NRP1", "AHNAK", "IGF1R", "CTSZ", "CYFIP1", "MYO1C", "ENPP1", "ITGAV", "CAPG", "CD44", "SYNGR2", "EMP1", "AHR", "BMPR2", "MAPKAPK2", "ALCAM") |
| 22 | HALLMARK_TGF_BETA_SIGNALING | 0.005262 | 0.011958 | -0.45276 | -1.84795 | 17 | 25 | c("SERPINE1", "CDH1", "THBS1", "CTNNB1", "RHOA", "NCOR2", "SKIL", "FNTA", "PMEPA1", "LTBP2", "BMPR2", "ENG", "RAB31", "TGIF1") |
| 23 | HALLMARK_FATTY_ACID_METABOLISM | 0.005857 | 0.012733 | -0.386097 | -1.749425 | 18 | 37 | c("MIF", "GLUL", "FASN", "BMPR1B", "LGALS1", "UGDH", "DHCR24", "S100A10", "PCBD1", "HSP90AA1", "EPHX1", "FH") |
| 24 | HALLMARK_INFLAMMATORY_RESPONSE | 0.006637 | 0.013827 | 0.385153 | 1.707183 | 45 | 51 | c("SELL", "CSF3R", "CLEC5A", "TLR1", "FPR1", "BEST1", "SEMA4D", "P2RY2", "IL4R", "IL7R", "AQP9", "PTPRE", "ITGB3", "MXD1", "LYN", "CD48", "TLR2", "CYBB", "LCP2", "IRF1", "NAMPT", "PIK3R5", "C5AR1") |
| 25 | HALLMARK_TNFA_SIGNALING_VIA_NFKB | 0.011933 | 0.023866 | -0.289851 | -1.547615 | 34 | 71 | c("CCND1", "NR4A1", "BHLHE40", "EIF1", "CCN1", "RHOB", "HES1", "EGR1", "JUN", "SERPINE1", "PHLDA1", "DUSP4", "SQSTM1", "FOSB", "GEM", "PLK2", "SDC4", "CD44", "DUSP1", "PLAU", "PMEPA1", "EFNA1", "TNC", "NFKBIA") |
| 26 | HALLMARK_NOTCH_SIGNALING | 0.014095 | 0.027105 | -0.699264 | -1.753191 | 57 | 6 | c("CCND1", "HES1", "NOTCH3") |

| | | | | | | | | |
|---|---|---|---|---|---|---|---|---|
| 27 | HALLMARK_G2M_CHECKPOINT | 0.017418 | 0.032256 | 0.319526 | 1.53774 | 123 | 77 | c("BRCA2", "CENPE", "PRC1", "LBR", "ESPL1", "HMMR", "AURKB", "BIRC5", "MKI67", "KIF20B", "NUP50", "E2F3", "CKS1B", "TFDP1", "E2F1", "TACC3", "MAPK14", "SLC7A5", "EZH2", "HIRA", "KMT5A", "HMGA1", "KIF22", "UBE2C", "WRN", "PLK1", "MYBL2", "SNRPD1", "TENT4A", "NUSAP1", "E2F4", "POLE", "ARID4A", "TPX2", "ODF2", "SMC4", "MTF2", "MCM2", "HUS1", "MCM6", "CDKN1B", "SFPQ", "STAG1", "TRA2B", "CUL4A", "LMNB1", "MCM3", "FOXN3", "DDX39A", "LIG3", "SRSF2") |
| 28 | HALLMARK_APICAL_SURFACE | 0.019968 | 0.035656 | -0.503076 | -1.715711 | 73 | 14 | c("GSTM3", "GATA3", "HSPB1", "SULF2", "THY1") |
| 29 | HALLMARK_IL6_JAK_STAT3_SIGNALING | 0.030471 | 0.052535 | 0.389236 | 1.551342 | 203 | 33 | c("CSF3R", "CD38", "ITGA4", "LTB", "IL17RA", "IL4R", "IL2RG", "CSF2RB", "ITGB3", "CSF2RA", "CBL", "BAK1", "TLR2", "IRF1", "PIK3R5", "MYD88", "PIM1") |
| 30 | HALLMARK_PROTEIN_SECRETION | 0.076757 | 0.127928 | -0.307166 | -1.391783 | 248 | 37 | c("TMED2", "TPD52", "TOM1L1", "PAM", "ARF1", "CD63", "PPT1", "SEC31A", "COPE", "RAB2A", "NAPA", "SOD1", "ARFGEF2", "ERGIC3", "COPB2", "LAMP2") |
| 31 | HALLMARK_MTORC1_SIGNALING | 0.093833 | 0.144371 | -0.251973 | -1.313218 | 282 | 64 | c("NUPR1", "BHLHE40", "CD9", "XBP1", "IFI30", "CANX", "SCD", "DHCR24", "CALR", "STC1", "SLC9A3R1", "SQSTM1", "IGFBP5", "SYTL2", "LGMN") |
| 32 | HALLMARK_P53_PATHWAY | 0.097935 | 0.144371 | -0.25073 | -1.311025 | 293 | 65 | c("NUPR1", "CD81", "CTSD", "TM4SF1", "IFI30", "JUN", "HMOX1", "TOB1", "S100A10", "ZFP36L1", "RACK1", "PLK2", "EPHX1", "EPS8L2", "TAX1BP3", "PLXNB2", "APP") |
| 33 | HALLMARK_KRAS_SIGNALING_UP | 0.094396 | 0.144371 | -0.253716 | -1.308237 | 282 | 62 | c("CFB", "MMP11", "SERPINA3", "SPP1", "APOD", "SPARCL1", "JUP", "PLAT", "GPNMB", "CPE", "IGFBP3", "NRP1", "TSPAN13", "PRRX1") |
| 34 | HALLMARK_KRAS_SIGNALING_DN | 0.098172 | 0.144371 | -0.45607 | -1.430408 | 375 | 11 | c("BMPR1B", "EFHD1", "SELENOP", "IDUA", "SKIL") |
| 35 | HALLMARK_MYC_TARGETS_V1 | 0.140097 | 0.200138 | -0.252497 | -1.253997 | 434 | 53 | c("CLNS1A", "CANX", "HSP90AB1", "PCBP1", "YWHAE", "SNRPD2", "RACK1", "CCT3", "IMPDH2", "NME1", "ILF2", "HNRNPU", "NDUFAB1", "CBX3", "NHP2", "PPIA", "HNRNPC", "GNL3", "RNPS1", "TUFM", "CNBP", "PSMD3", "POLD2", "TXNL4A") |
| 36 | HALLMARK_HEDGEHOG_SIGNALING | 0.166319 | 0.230998 | -0.410069 | -1.286132 | 636 | 11 | c("CELSR1", "THY1", "NRP1", "MYH9", "ADGRG1", "DPYSL2") |
| 37 | HALLMARK_UNFOLDED_PROTEIN_RESPONSE | 0.195461 | 0.261542 | -0.268322 | -1.214144 | 645 | 36 | c("STC2", "EEF2", "XBP1", "CALR", "YWHAZ", "SEC31A", "LSM1", "ATP6V0D1", "EIF4G1", "DDIT4", "NHP2", "SPCS1", "EIF4A2", "LSM4") |
| 38 | HALLMARK_UV_RESPONSE_UP | 0.198772 | 0.261542 | -0.24031 | -1.184436 | 614 | 52 | c("EPCAM", "NR4A1", "RET", "RHOB", "MMP14", "HMOX1", "SELENOW", "GRINA", "SQSTM1", "FOSB") |
| 39 | HALLMARK_REACTIVE_OXYGEN_SPECIES_PATHWAY | 0.277051 | 0.355194 | 0.331277 | 1.151104 | 1799 | 20 | c("MPO", "CDKN2D", "FES", "HHEX", "MSRA", "CAT", "TXNRD2", "MBP") |
| 40 | HALLMARK_PI3K_AKT_MTOR_SIGNALING | 0.391362 | 0.471793 | 0.252878 | 1.049696 | 2654 | 39 | c("PRKCB", "MAPK10", "DAPP1", "SLA", "IL2RG", "E2F1", "IRAK4", "TRAF2", "PDK1", "MAP2K3", "MYD88", "SLC2A1", "PIKFYVE", "RALB", "ACTR2", "GRK2") |
| 41 | HALLMARK_BILE_ACID_METABOLISM | 0.396306 | 0.471793 | 0.362701 | 1.052059 | 2445 | 11 | c("KLF1", "AQP9", "CAT", "ACSL1", "HSD17B11") |
| 42 | HALLMARK_SPERMATOGENESIS | 0.393404 | 0.471793 | -0.314488 | -1.046919 | 1490 | 13 | c("GSTM3", "SLC12A2", "PEBP1") |
| 43 | HALLMARK_MITOTIC_SPINDLE | 0.421823 | 0.490492 | 0.209125 | 1.02454 | 3026 | 84 | c("ANLN", "BRCA2", "CENPE", "TUBA4A", "PRC1", "CENPJ", "ESPL1", "DOCK2", "BIRC5", "SSH2", "TUBD1", "RHOF", "KNTC1", "KIF20B", "CNTRL", "CEP192", "TUBGCP3", "ARHGAP27", "KIF22", "ARHGAP4", "PLK1", "PPP4R2", "NUSAP1", "CDK5RAP2", "BCL2L11", "PCNT", "TPX2", "CLASP1", "NIN", "RASA2", "KIF1B", "ATG4B", "SMC4", "HDAC6", "EPB41", "MAP3K11") |
| 44 | HALLMARK_MYC_TARGETS_V2 | 0.467321 | 0.531047 | 0.326715 | 0.996892 | 2902 | 13 | c("SLC19A1", "RCL1", "PLK1", "NOP2", "TBRG4") |
| 45 | HALLMARK_PANCREAS_BETA_CELLS | 0.562262 | 0.624736 | 0.4259 | 0.932074 | 3250 | 5 | c("ELP4", "LMO2") |
| 46 | HALLMARK_DNA_REPAIR | 0.606513 | 0.645227 | 0.22432 | 0.905969 | 4059 | 35 | c("PNP", "AGO4", "ZNF707", "POLE4", "POLA1", "ZWINT", "ERCC5", "VPS37B", "STX3", "HCLS1", "PCNA", "POLL", "ELL") |
| 47 | HALLMARK_PEROXISOME | 0.594441 | 0.645227 | -0.238598 | -0.892948 | 2095 | 19 | c("CRABP2", "DHCR24", "SEMA3C", "SOD1", "FIS1", "ELOVL5", "IDH2", "CNBP") |
| 48 | HALLMARK_INTERFERON_ALPHA_RESPONSE | 0.630205 | 0.656464 | -0.195368 | -0.897459 | 2027 | 39 | c("IFITM3", "IFI30", "B2M", "IFI27", "C1S", "LGALS3BP", "CD74", "IFITM2", "BST2", "LY6E") |
| 49 | HALLMARK_INTERFERON_GAMMA_RESPONSE | 0.68927 | 0.703336 | 0.182841 | 0.873681 | 4868 | 75 | c("CD38", "FPR1", "IL4R", "VCAM1", "CSF2RB", "CASP8", "NLRC5", "ISG20", "PNP", "ST8SIA4", "JAK2", "LCP2", "BPGM", "IRF8", "PSMB9", "IRF1", "TRAFD1", "NAMPT", "FGL2", "MYD88", "PIM1", "RNF31", "NMI", "PTPN6", "PLSCR1", "SP110", "NCOA3", "SLC25A28", "CIITA", "IRF2", "CCL5", "EIF4E3", "TNFAIP3") |
| 50 | HALLMARK_WNT_BETA_CATENIN_SIGNALING | 0.942264 | 0.942264 | 0.206791 | 0.580224 | 5760 | 10 | c("FRAT1", "AXIN1", "CCND2", "NOTCH1", "MAML1", "MYC") |

| Cluster 25 | pathway | pval | padj | ES | NES | nMoreExtreme | size | leadingEdge |
|---|---|---|---|---|---|---|---|---|
| 1 | HALLMARK_HYPOXIA | 0.000112 | 0.000735 | -0.569476 | -2.098863 | 0 | 64 | c("DCN", "CCN1", "STC2", "CCN2", "BGN", "COL5A1", "JUN", "CA12", "LOX", "SERPINE1", "SDC2", "STC1", "IGFBP3", "ANXA2", "BHLHE40", "TGFBI", "HMOX1", "GPC1", "EFNA1", "FOS", "TPD52", "DTNA", "MIF", "SDC4", "CAVIN1", "TGFB3", "MT2A", "DUSP1", "SDC3", "DDIT4", "FBP1") |
| 2 | HALLMARK_ESTROGEN_RESPONSE_EARLY | 0.00011 | 0.000735 | -0.675537 | -2.544556 | 0 | 77 | c("SLC7A2", "STC2", "MAPT", "RET", "NBL1", "HSPB8", "TFF1", "MLPH", "CCND1", "KRT19", "IGFBP4", "PRSS23", "CA12", "SCNN1A", "HES1", "PGR", "ADCY1", "KRT8", "MUC1", "SLC39A6", "GJA1", "GREB1", "GFRA1", "CLDN7", "THSD4", "ELF3", "FLNB", "CELSR1", "BHLHE40", "RHOD", "SEC14L2", "MYOF", "FASN", "FARP1", "XBP1", "SEMA3B", "FOS", "PAPSS2", "RHOBTB3", "PLAAT3", "TTC39A", "TFF3", "FAM102A", "MAST4", "TOB1") |
| 3 | HALLMARK_ESTROGEN_RESPONSE_LATE | 0.00011 | 0.000735 | -0.599823 | -2.242906 | 0 | 72 | c("CXCL14", "CPE", "MAPT", "RET", "SERPINA3", "NBL1", "HSPB8", "TFF1", "AGR2", "SCUBE2", "CCND1", "PRLR", "KRT19", "IGFBP4", "PRSS23", "CA12", "SCNN1A", "PGR", "CD9", "MDK", "FLNB", "DLG5", "UGDH", "MYOF", "LSR", "CDH1", "FARP1", "XBP1", "COX6C", "SEMA3B", "FOS", "PAPSS2", "PLAAT3", "TFF3", "FAM102A", "TOB1", "SERPINA1", "LLGL2", "MAPK13", "SGK1", "SLC9A3R1") |
| 4 | HALLMARK_MYOGENESIS | 0.000115 | 0.000735 | -0.649124 | -2.320315 | 0 | 53 | c("COL6A2", "PVALB", "STC2", "HSPB8", "COL6A3", "APOD", "ADAM12", "ITGB5", "COL4A2", "COL3A1", "CLU", "COL1A1", "CNN3", "SPDEF", "IGFBP7", "IGFBP3", "TAGLN", "BHLHE40", "ERBB3", "SPARC", "NQO1", "SCD", "DTNA", "PLXNB2", "GPX3", "SORBS3", "TNNT1", "AEBP1", "APP", "SPTAN1", "AGRN", "TPM2", "MYO1C", "ITGB1") |
| 5 | HALLMARK_APICAL_JUNCTION | 0.000115 | 0.000735 | -0.578118 | -2.072112 | 0 | 54 | c("MMP2", "CDH11", "PARVA", "CD276", "THY1", "FBN1", "CLDN7", "MYL9", "MDK", "CLDN4", "COL16A1", "JUP", "TGFBI", "CDH1", "NECTIN2", "SORBS3", "CERCAM", "LIMA1", "SDC3", "PCDH1", "ACTG1", "EVL", "MAPK13", "CTNND1", "TSPAN4", "ITGB1", "PIK3R3", "AKT3", "VCAN", "CTNNA1", "SHC1") |
| 6 | HALLMARK_EPITHELIAL_MESENCHYMAL_TRANSITION | 0.000106 | 0.000735 | -0.783614 | -3.059631 | 0 | 104 | c("POSTN", "DCN", "ELN", "SPP1", "COL6A2", "CCN1", "MMP2", "CCN2", "FBLN1", "COL11A1", "HTRA1", "CDH11", "COL6A3", "LUM", "LRRC15", "SFRP4", "ACTA2", "BGN", "ADAM12", "MXRA5", "THBS2", "COL5A2", "ITGB5", "LOXL1", "TIMP3", "COL4A2", "FSTL1", "IGFBP4", "PDGFRB", "COL5A1", "JUN", "COL12A1", "FBLN2", "COL3A1", "MMP14", "CTHRC1", "THY1", "COMP", "COL1A1", "LOXL2", "TIMP1", "LOX", "FBN1", "FAP", "CALD1", "COL1A2", "MGP", "SERPINE1", "DPYSL3", "COL4A1", "GJA1", "MYL9", "RHOB", "NNMT", "FN1", "PMEPA1", "IGFBP3", □ "TAGLN", "PRRX1", "ECM1", "ITGAV", "COL16A1", "LRP1", "SPARC", "TPM4", "PMP22", "DST", "TGFBI", "PCOLCE", "TNC", "LGALS1", "GPC1", "LAMC1", "GEM", "PFN2", "SDC4", "SDC1", "SERPINH1", "FUCA1") |
| 7 | HALLMARK_UV_RESPONSE_DN | 0.000116 | 0.000735 | -0.623317 | -2.188201 | 0 | 47 | c("CCN1", "COL11A1", "COL5A2", "INPP4B", "PDGFRB", "IGFBP5", "COL3A1", "COL1A1", "EFEMP1", "COL1A2", "ERBB2", "SERPINE1", "GJA1", "SDC2", "ANXA2", "BHLHE40", "PMP22", "PLPP3", "LAMC1", "RBPMS", "DUSP1", "DAB2", "NRP1", "APBB2", "ANXA4", "PIK3R3", "AKT3") |
| 8 | HALLMARK_COAGULATION | 0.000118 | 0.000735 | -0.750737 | -2.602154 | 0 | 44 | c("MMP2", "S100A1", "A2M", "HTRA1", "C1R", "TIMP3", "C3", "PLAT", "PRSS23", "MMP11", "MMP14", "CLU", "COMP", "APOC1", "CD9", "TIMP1", "FBN1", "SERPINE1", "C1S", "S100A13", "CTSK", "FN1", "CRIP2", "C1QA", "LRP1", "SPARC", "SERPING1", "LGMN", "CFB", "CTSB", "THBS1", "SERPINA1", "CAPN2", "CSRP1") |
| 9 | HALLMARK_TNFA_SIGNALING_VIA_NFKB | 0.000226 | 0.00113 | -0.52478 | -1.909458 | 1 | 59 | c("CCN1", "NR4A1", "CCND1", "JUN", "EGR1", "HES1", "SERPINE1", "RHOB", "PMEPA1", "PLK2", "PHLDA1", "DUSP4", "BHLHE40", "TNC", "PLPP3", "EFNA1", "FOS", "FOSB", "GEM", "SDC4", "DUSP1", "SOCS3", "SGK1", "NFKBIA", "MARCKS", "EIF1", "KLF9", "NINJ1", "SQSTM1", "CD44", "PLAU", "INHBA", "SAT1", "FOSL2", "IER3", "BCL3", "VEGFA", "JUNB", "ZFP36", "IER2", "B4GALT1", "TGIF1", "KLF6", "MAP3K8", "MCL1") |
| 10 | HALLMARK_APOPTOSIS | 0.000223 | 0.00113 | -0.505273 | -1.870019 | 1 | 66 | c("DCN", "MMP2", "LUM", "BGN", "CCND1", "TIMP3", "PDGFRB", "PLAT", "JUN", "CLU", "TIMP1", "ERBB2", "IFITM3", "RHOB", "ERBB3", "EMP1", "HSPB1", "HMOX1", "TIMP2", "GPX3", "GPX4", "APP", "PEA15", "LMNA", "SPTAN1", "CTNNB1", "RARA", "TNFRSF12A", "PPT1", "GSN", "LGALS3", "SQSTM1", "CD44") |

| | | | | | | | | |
|---|---|---|---|---|---|---|---|---|
| 11 | HALLMARK_CHOLESTEROL_HOMEOSTASIS | 0.000684 | 0.003109 | -0.69878 | -1.932817 | 4 | 15 | c("CLU", "CD9", "LGMN", "FASN", "SCD", "SEMA3B", "ACTG1", "ALCAM", "ANXA5", "CTNNB1", "S100A11", "TNFRSF12A", "LGALS3", "PMVK") |
| 12 | HALLMARK_ANGIOGENESIS | 0.000808 | 0.003368 | -0.679319 | -1.934111 | 5 | 17 | c("POSTN", "SPP1", "LUM", "COL5A2", "FSTL1", "COL3A1", "TIMP1", "STC1", "ITGAV", "APP", "FGFR1", "NRP1") |
| 13 | HALLMARK_G2M_CHECKPOINT | 0.001083 | 0.003869 | 0.541719 | 2.913423 | 0 | 76 | c("E2F2", "KIF2C", "NDC80", "CDC20", "KIF11", "HMMR", "CCNB2", "PTTG1", "MKI67", "PLK1", "AURKB", "TACC3", "PRC1", "FBXO5", "TROAP", "MYBL2", "NUSAP1", "LBR", "TFDP1", "CCNA2", "SLC7A5", "KIF23", "CCNF", "EZH2", "E2F4", "E2F1", "BIRC5", "INCENP", "KIF20B", "TPX2", "SLC7A1", "TMPO", "KIF22", "HIRA", "TOP2A", "TOP1", "SMC4", "LMNB1", "MCM5", "POLE", "WRN", "CDC27", "NSD2", "KMT5A", "MCM2", "MCM6", "UBE2C") |
| 14 | HALLMARK_E2F_TARGETS | 0.001052 | 0.003869 | 0.606359 | 3.193507 | 0 | 71 | c("KIF18B", "KIF2C", "E2F8", "CDC20", "SPC24", "HMMR", "CCNB2", "RRM2", "PTTG1", "MKI67", "PLK1", "CIT", "AURKB", "MXD3", "TACC3", "SPC25", "MYBL2", "TFRC", "LBR", "HMGB2", "EZH2", "BIRC5", "CENPM", "PSIP1", "PCNA", "ANP32E", "ASF1A", "ASF1B", "DCK", "ATAD2", "TMPO", "NCAPD2", "KIF22", "TOP2A", "SMC4", "TUBG1", "LMNB1", "MCM5", "POLD1", "POLE", "LIG1", "MCM2", "MCM6", "DNMT1", "HELLS") |
| 15 | HALLMARK_IL2_STAT5_SIGNALING | 0.001248 | 0.00416 | -0.486257 | -1.760984 | 10 | 57 | c("SPP1", "S100A1", "COL6A1", "ENPP1", "MUC1", "IFITM3", "RHOB", "PHLDA1", "ECM1", "BHLHE40", "ITGAV", "EMP1", "XBP1", "AHR", "GPX4", "NRP1", "AHNAK", "ALCAM", "CD81", "CYFIP1", "LRIG1", "MYO1C", "ANXA4", "CTSZ", "BMPR2", "SYNGR2", "IGF1R", "SERPINB6", "NFKBIZ", "P4HA1", "CAPG", "CD44", "MAPKAPK2", "ITGA6") |
| 16 | HALLMARK_HEME_METABOLISM | 0.001996 | 0.006238 | 0.647689 | 3.748659 | 0 | 114 | c("RHAG", "EPB42", "CA1", "TMCC2", "AHSP", "HBD", "SLC4A1", "SPTA1", "TRIM58", "SNCA", "E2F2", "HBB", "ALAS2", "ANK1", "RBM38", "OSBP2", "DMTN", "SPTB", "HMBS", "NFE2", "FECH", "KEL", "TENT5C", "KLF1", "SLC25A37", "CA2", "TSPAN5", "GYPC", "BPGM", "KAT2B", "CPOX", "ERMAP", "XPO7", "FBXO7", "UROD", "TRAK2", "EPB41", "LMO2", "RANBP10", "CAT", "ALAD", "MAP2K3", "TFRC", "USP15", "GLRX5", "PRDX2", "ABCB6", "PPOX", "PIGQ", "UBAC1", "PPP2R5B", "GCLC", "HTRA2", "SLC6A8", "NARF", "TFDP2", "SLC2A1", "GCLM", □<br>"BNIP3L", "KLF3", "KDM7A", "HAGH", "RIOK3", "BLVRB", "RCL1", "MXI1", "FBXO9", "DCAF11", "MPP1", "CCND3", "PDZK1IP1", "TOP1", "SLC25A38", "MFHAS1", "MBOAT2", "CDC27", "UROS", "NCOA4", "UCP2", "SLC66A2", "SELENBP1") |
| 17 | HALLMARK_COMPLEMENT | 0.00374 | 0.011001 | -0.462783 | -1.679259 | 32 | 58 | c("C1R", "COL4A2", "C3", "PLAT", "MMP14", "GATA3", "CLU", "APOC1", "TIMP1", "SERPINE1", "C1S", "S100A13", "FN1", "C1QA", "LRP1", "C1QC", "SERPING1", "LGMN", "LIPA", "CTSD", "CFB", "TIMP2", "CTSB", "SERPINA1", "CSRP1", "ANXA5") |
| 18 | HALLMARK_KRAS_SIGNALING_UP | 0.004304 | 0.011956 | -0.484896 | -1.702264 | 36 | 47 | c("SPP1", "CPE", "SERPINA3", "APOD", "GPNMB", "PLAT", "SPARCL1", "MMP11", "IGFBP3", "PRRX1", "JUP", "EMP1", "TMEM176B", "SEMA3B", "CFB", "MAFB", "FUCA1") |
| 19 | HALLMARK_GLYCOLYSIS | 0.00576 | 0.015159 | -0.470997 | -1.676537 | 49 | 52 | c("DCN", "STC2", "COL5A1", "SDC2", "STC1", "ELF3", "IGFBP3", "TGFBI", "GPC1", "MIF", "EGLN3", "SDC1", "IL13RA1", "CHPF", "TFF3", "SDC3", "DDIT4", "AGRN", "PLOD1", "VCAN", "PKM", "FUT8", "PAM", "P4HA1", "CD44", "PGLS", "CYB5A", "IER3", "VEGFA", "GFPT1", "HDLBP", "ALDH9A1", "PYGB", "PPIA", "IDUA", "B4GALT1") |
| 20 | HALLMARK_ANDROGEN_RESPONSE | 0.008401 | 0.021003 | -0.488084 | -1.674347 | 70 | 41 | c("AZGP1", "CCND1", "INPP4B", "KRT19", "BMPR1B", "SPDEF", "KRT8", "SELENOP", "PMEPA1", "ITGAV", "DHCR24", "SCD", "TPD52") |
| 21 | HALLMARK_NOTCH_SIGNALING | 0.008859 | 0.021093 | -0.739242 | -1.667211 | 57 | 7 | c("NOTCH3", "CCND1", "HES1") |
| 22 | HALLMARK_P53_PATHWAY | 0.014317 | 0.032538 | -0.441346 | -1.581891 | 124 | 54 | c("NUPR1", "TM4SF1", "JUN", "PLK2", "IFI30", "HMOX1", "S100A10", "CTSD", "FOS", "EPS8L2", "EPHX1", "PLXNB2", "SDC1", "FUCA1", "APP", "TRAF4", "ZFP36L1", "TOB1", "DDIT4") |
| 23 | HALLMARK_APICAL_SURFACE | 0.022006 | 0.04784 | -0.634827 | -1.626892 | 153 | 11 | c("GATA3", "THY1", "GSTM3", "SULF2", "HSPB1", "APP") |
| 24 | HALLMARK_XENOBIOTIC_METABOLISM | 0.024081 | 0.050169 | -0.440566 | -1.546642 | 206 | 47 | c("FBLN1", "IGFBP4", "APOE", "SERPINE1", "UGDH", "JUP", "PDK4", "TMEM176B", "ESR1", "NQO1", "HMOX1", "CFB", "EPHX1", "BCAR1", "PAPSS2", "MT2A", "FBP1", "SPINT2", "PGRMC1") |
| 25 | HALLMARK_ALLOGRAFT_REJECTION | 0.025607 | 0.051215 | 0.321678 | 1.505363 | 38 | 43 | c("ELANE", "ACHE", "NCF4", "FGR", "WAS", "PRKCB", "ZAP70", "SRGN", "LTB", "IL2RG", "PTPN6", "CCND3", "LCP2", "FYB1", "HCLS1", "TLR2", "LYN", "JAK2", "TAP2", "ELF4") |

| | | | | | | | | |
|---|---|---|---|---|---|---|---|---|
| 26 | HALLMARK_TGF_BETA_SIGNALING | 0.034951 | 0.067214 | -0.504457 | -1.539538 | 269 | 23 | c("SERPINE1", "PMEPA1", "LTBP2", "CDH1", "THBS1", "CTNNB1", "ID3", "SKIL", "BMPR2", "NCOR2", "JUNB", "RHOA", "ENG", "TGIF1", "FNTA", "HIPK2") |
| 27 | HALLMARK_MYC_TARGETS_V2 | 0.038282 | 0.070893 | 0.518872 | 1.606471 | 114 | 11 | c("PLK1", "RCL1", "PRMT3", "SLC19A1", "MCM5", "MCM4", "MYC", "NOP16", "PA2G4", "NOP56") |
| 28 | HALLMARK_MITOTIC_SPINDLE | 0.049516 | 0.088421 | 0.252167 | 1.349258 | 45 | 75 | c("KIF2C", "NDC80", "KIF11", "CCNB2", "PLK1", "EPB41", "PRC1", "FBXO5", "NUSAP1", "TUBA4A", "KIF23", "BIRC5", "INCENP", "DOCK2", "KIF20B", "TPX2", "KIF22", "TOP2A", "SMC4", "ARHGAP4", "SSH2", "LMNB1", "BCL2L11", "CDC27") |
| 29 | HALLMARK_ADIPOGENESIS | 0.06492 | 0.111931 | -0.396134 | -1.401425 | 559 | 50 | c("HSPB8", "C3", "SPARCL1", "APOE", "PHLDB1", "COL4A1", "LAMA4", "FABP4", "CD151", "GPX3", "CAVIN1", "GPX4", "TOB1", "VEGFB", "GPAT4", "RAB34", "COX6A1", "CCNG2", "SCP2") |
| 30 | HALLMARK_KRAS_SIGNALING_DN | 0.123178 | 0.205297 | -0.531474 | -1.362026 | 861 | 11 | c("EFHD1", "BMPR1B", "SELENOP", "IGFBP2", "SGK1", "SKIL") |
| 31 | HALLMARK_PROTEIN_SECRETION | 0.254212 | 0.397206 | -0.364633 | -1.178866 | 2051 | 30 | c("DST", "TPD52", "TOM1L1", "TMED2", "SEC31A", "PPT1", "PAM", "CD63", "RAB2A", "COPE", "ARFGEF2", "USO1", "ERGIC3", "SNX2", "LMAN1", "LAMP2", "COPB2", "VPS45", "ARF1") |
| 32 | HALLMARK_FATTY_ACID_METABOLISM | 0.24708 | 0.397206 | -0.356611 | -1.187134 | 2030 | 35 | c("BMPR1B", "UGDH", "FASN", "DHCR24", "S100A10", "LGALS1", "PCBD1", "EPHX1", "MIF") |
| 33 | HALLMARK_REACTIVE_OXYGEN_SPECIES_PATHWAY | 0.274315 | 0.415629 | 0.309817 | 1.137499 | 690 | 18 | c("MPO", "CAT", "CDKN2D", "PRDX2", "GCLC", "GCLM", "MSRA") |
| 34 | HALLMARK_OXIDATIVE_PHOSPHORYLATION | 0.293974 | 0.432315 | -0.324232 | -1.138241 | 2526 | 47 | c("ATP1B1", "MAOB", "PDK4", "COX6C", "GPX4", "NDUFC2", "ATP6V1F", "NDUFA4", "NDUFB2", "COX7C", "ATP6AP1", "COX6A1", "CYB5A", "UQCRB", "ATP6V0E1", "ATP5F1B", "ECI1", "ATP5PD", "UQCRQ", "UQCR10", "NDUFAB1", "ATP5MC2", "NDUFA3", "GLUD1") |
| 35 | HALLMARK_PEROXISOME | 0.337966 | 0.482809 | -0.38618 | -1.114961 | 2528 | 18 | c("CRABP2", "SEMA3C", "DHCR24") |
| 36 | HALLMARK_UNFOLDED_PROTEIN_RESPONSE | 0.446007 | 0.602712 | -0.323702 | -1.02544 | 3551 | 27 | c("STC2", "XBP1", "DDIT4", "SEC31A", "YWHAZ", "SHC1", "SPCS1", "CALR", "VEGFA", "EEF2", "NHP2", "EIF4A2", "EIF4EBP1", "LSM1", "KIF5B", "SERP1", "EIF4G1") |
| 37 | HALLMARK_UV_RESPONSE_UP | 0.440298 | 0.602712 | -0.291777 | -1.028844 | 3786 | 49 | c("NR4A1", "RET", "MMP14", "RHOB", "HMOX1", "FOS", "FOSB", "EPHX1", "GPX3", "SELENOW", "IGFBP2", "EPCAM", "NFKBIA", "ATP6V1F", "PPT1", "SQSTM1") |
| 38 | HALLMARK_INTERFERON_ALPHA_RESPONSE | 0.487899 | 0.609873 | -0.321899 | -0.993581 | 3809 | 24 | c("C1S", "IFITM3", "LGALS3BP", "IFI30", "IFI27", "LY6E", "CD74", "IFITM2", "BST2", "B2M") |
| 39 | HALLMARK_MTORC1_SIGNALING | 0.466113 | 0.609873 | -0.279466 | -1.008601 | 4098 | 56 | c("NUPR1", "IGFBP5", "CD9", "STC1", "BHLHE40", "LGMN", "IFI30", "DHCR24", "XBP1", "SCD", "EGLN3", "SERPINH1", "SYTL2", "DDIT4", "SLC9A3R1", "PIK3R3", "CANX", "ACSL3") |
| 40 | HALLMARK_BILE_ACID_METABOLISM | 0.478883 | 0.609873 | 0.298275 | 0.976339 | 1371 | 13 | c("KLF1", "CAT", "GCLM") |
| 41 | HALLMARK_INTERFERON_GAMMA_RESPONSE | 0.549679 | 0.667041 | -0.271239 | -0.948572 | 4707 | 46 | c("C1R", "C1S", "IFITM3", "SERPING1", "LGALS3BP", "IFI30", "CFB", "IFI27", "MT2A", "SOCS3", "NFKBIA", "LY6E", "CD74", "IFITM2", "BST2", "B2M", "STAT3", "ARID5B") |
| 42 | HALLMARK_MYC_TARGETS_V1 | 0.560314 | 0.667041 | 0.196473 | 0.949631 | 784 | 49 | c("CDC20", "TFDP1", "CCNA2", "PCNA", "TYMS", "MCM5", "NCBP1", "MCM2", "MCM6", "SRPK1", "DEK", "MCM7", "RRM1", "DUT", "MCM4", "HDGF", "SRSF1", "SRSF2", "TRA2B", "USP1", "MYC", "NOP16", "VDAC3", "PA2G4", "EIF3B", "NOP56", "SF3B3", "HNRNPD", "HNRNPU", "HNRNPC", "FAM120A", "TUFM", "NME1", "PCBP1", "ILF2", "HSPE1", "NDUFAB1", "SNRPD2", "RNPS1", "PSMD3", "RACK1", "PPIA", "YWHAE", "NHP2", "IMPDH2", "CCT3", "CLNS1A", "CANX", "HSP90AB1") |
| 43 | HALLMARK_WNT_BETA_CATENIN_SIGNALING | 0.614767 | 0.694876 | -0.411931 | -0.888299 | 3904 | 6 | c("CTNNB1", "NCOR2", "CSNK1E", "NCSTN", "MYC", "NOTCH1") |
| 44 | HALLMARK_HEDGEHOG_SIGNALING | 0.602748 | 0.694876 | -0.361204 | -0.898162 | 4123 | 10 | c("THY1", "CELSR1", "NRP1", "DPYSL2", "ADGRG1", "VEGFA") |
| 45 | HALLMARK_PI3K_AKT_MTOR_SIGNALING | 0.625389 | 0.694876 | 0.210657 | 0.901888 | 1206 | 30 | c("MAP2K3", "PRKCB", "SLA", "E2F1", "SLC2A1", "IL2RG", "GRK2", "PIKFYVE", "CXCR4", "MKNK2", "PPP2R1B", "ACTR3", "CLTC") |
| 46 | HALLMARK_SPERMATOGENESIS | 0.65311 | 0.709902 | 0.266943 | 0.854528 | 1910 | 12 | c("KIF2C", "CCNB2", "EZH2") |
| 47 | HALLMARK_PANCREAS_BETA_CELLS | 0.684415 | 0.728101 | -0.409991 | -0.839286 | 4228 | 5 | c("MAFB", "AKT3", "SPCS1") |
| 48 | HALLMARK_IL6_JAK_STAT3_SIGNALING | 0.707057 | 0.736517 | -0.260663 | -0.825741 | 5630 | 27 | c("A2M", "JUN", "CD9", "HMOX1", "IL13RA1", "SOCS3", "LTBR", "TNFRSF12A", "CD44", "IL1R1") |
| 49 | HALLMARK_INFLAMMATORY_RESPONSE | 0.798722 | 0.815022 | -0.222107 | -0.761926 | 6749 | 41 | c("SLC7A2", "MMP14", "TIMP1", "SERPINE1") |

| | | | | | | | | |
|---|---|---|---|---|---|---|---|---|
| 50 | HALLMARK_DNA_REPAIR | 0.897436 | 0.897436 | 0.158367 | 0.715782 | 1539 | 37 | c("PNP", "PCNA", "TYMS", "POLD1", "HCLS1", "ZWINT", "LIG1", "AGO4", "VPS37B", "DUT", "STX3", "AK1", "GUK1", "EIF1B") |

| Cluster 26 | pathway | pval | padj | ES | NES | nMoreExtreme | size | leadingEdge |
|---|---|---|---|---|---|---|---|---|
| 1 | HALLMARK_APICAL_JUNCTION | 0.000558 | 0.011181 | 0.455671 | 1.907404 | 4 | 75 | c("DSC3", "COL17A1", "CX3CL1", "ACTG2", "WNK4", "ITGB4", "CDH3", "ITGA2", "LAMB3", "CLDN11", "EGFR", "ITGA3", "LAMC2", "IRS1", "NECTIN1", "LAMA3", "AMIGO2") |
| 2 | HALLMARK_COMPLEMENT | 0.000739 | 0.011181 | -0.382545 | -1.967638 | 0 | 51 | c("SERPINA1", "S100A9", "CTSL", "PLAUR", "FN1", "CTSD", "PIM1", "LIPA", "CTSB", "CTSS", "LGMN", "HSPA1A", "PFN1", "APOC1", "PRKCD") |
| 3 | HALLMARK_OXIDATIVE_PHOSPHORYLATION | 0.000894 | 0.011181 | -0.372159 | -2.033522 | 0 | 66 | c("COX6C", "NDUFC2", "COX6B1", "NDUFB2", "ATP6V0C", "TCIRG1", "MGST3", "COX8A", "ATP5MF", "UQCRC1", "ATP6AP1", "NDUFS6", "FH", "COX7C", "ATP6V1F", "NDUFS7", "NDUFA1", "NDUFA2", "ATP6V0B", "UQCR11", "NDUFC1", "ATP5PD", "COX6A1", "ATP5F1E", "NDUFA4") |
| 4 | HALLMARK_HEME_METABOLISM | 0.000685 | 0.011181 | -0.597028 | -3.008757 | 0 | 46 | c("HBB", "HBD", "SLC4A1", "BLVRB", "ACP5", "MPP1", "SNCA", "HAGH", "CTSB", "TFRC", "PRDX2", "MGST3", "SLC2A1", "BSG", "FBXO7", "GYPC", "RAD23A", "FBXO9", "SELENBP1", "GDE1") |
| 5 | HALLMARK_TNFA_SIGNALING_VIA_NFKB | 0.001656 | 0.015733 | 0.417018 | 1.781857 | 14 | 85 | c("CXCL2", "AREG", "LAMB3", "ATF3", "TANK", "DUSP2", "F3", "PNRC1", "PDE4B", "PER1", "KLF9", "MYC", "MAFF", "DUSP5", "HES1", "EGR1", "JAG1", "STAT5A", "NFIL3", "JUN", "EGR3", "IRS2", "TNFAIP3", "SGK1", "DUSP4", "TRIP10", "PLK2", "TNC", "TLR2", "SDC4", "FOSB", "NFE2L2", "NFAT5", "PMEPA1", "IER5", "KLF6", "ZFP36", "KLF4", "NFKBIA", "TSC22D1") |
| 6 | HALLMARK_ESTROGEN_RESPONSE_EARLY | 0.002203 | 0.015733 | -0.29998 | -1.765867 | 1 | 90 | c("RHOBTB3", "RET", "HSPB8", "BLVRB", "ABHD2", "PRSS23", "GAB2", "ADCY1", "PAPSS2", "GJA1", "ITPK1", "SEC14L2", "MAST4", "SEMA3B", "MUC1", "SLC2A1", "STC2", "GREB1", "RHOD", "CCND1", "PGR", "SLC7A2", "FAM102A", "FLNB", "CANT1") |
| 7 | HALLMARK_ESTROGEN_RESPONSE_LATE | 0.00206 | 0.015733 | -0.320262 | -1.835001 | 1 | 82 | c("COX6C", "RET", "HSPB8", "BLVRB", "SERPINA1", "TOP2A", "S100A9", "ABHD2", "PRSS23", "PAPSS2", "AGR2", "ITPK1", "SEMA3B", "DCXR", "CDH1", "CCND1", "PGR", "TSPAN13", "FAM102A", "FLNB") |
| 8 | HALLMARK_KRAS_SIGNALING_DN | 0.002696 | 0.016851 | 0.616559 | 1.881765 | 20 | 17 | c("SOX10", "KRT5", "KRT15", "TFCP2L1", "ZBTB16", "TFAP2B", "TCF7L1", "DTNB", "SGK1", "IGFBP2") |
| 9 | HALLMARK_INFLAMMATORY_RESPONSE | 0.00772 | 0.042887 | 0.456262 | 1.693721 | 64 | 39 | c("CX3CL1", "EREG", "MET", "RGS1", "ITGB8", "TNFSF10", "F3", "PDE4B", "MYC", "CALCRL", "P2RY2", "ACVR2A", "AHR") |
| 10 | HALLMARK_MTORC1_SIGNALING | 0.010959 | 0.054795 | -0.336594 | -1.696283 | 15 | 46 | c("SCD", "IFI30", "CORO1A", "SQSTM1", "ITGB2", "LGMN", "P4HA1", "TFRC", "TXNRD1", "SLC2A1", "GSR", "STC1", "GGA2", "CALR") |
| 11 | HALLMARK_KRAS_SIGNALING_UP | 0.022704 | 0.1032 | 0.379198 | 1.532292 | 199 | 61 | c("PIGR", "EREG", "SLPI", "APOD", "ITGA2", "ANO1", "GUCY1A1", "ALDH1A3", "SOX9", "DUSP6", "KCNN4", "SPRY2", "PRDM1", "PEG3", "TRIB2", "ANGPTL4", "IGFBP3", "TNFAIP3") |
| 12 | HALLMARK_NOTCH_SIGNALING | 0.029179 | 0.113315 | 0.589516 | 1.603429 | 217 | 11 | c("MAML2", "FZD7", "TCF7L2", "HES1", "JAG1") |
| 13 | HALLMARK_REACTIVE_OXYGEN_SPECIES_PATHWAY | 0.029462 | 0.113315 | -0.476763 | -1.736993 | 68 | 15 | c("MPO", "FTL", "PRDX2", "TXNRD1", "GSR") |
| 14 | HALLMARK_P53_PATHWAY | 0.038044 | 0.135872 | 0.348053 | 1.447105 | 339 | 72 | c("KRT17", "TP63", "ITGB4", "ZBTB16", "PERP", "LDHB", "ATF3", "PDGFA", "TPD52L1", "PTPN14", "SFN", "JAG2", "TXNIP", "DDIT4", "JUN", "F2R", "ST14", "STEAP3", "SESN1", "FBXW7", "FOXO3", "EPS8L2", "PLK2", "CDKN2B", "APP", "IER5", "KLF4", "KIF13B", "TSC22D1", "CCNG1", "TRAF4", "BLCAP", "FOS", "DNTTIP2", "RALGDS", "TSPYL2", "VWA5A", "SDC1", "IER3", "RACK1", "BTG2", "NOTCH1") |
| 15 | HALLMARK_COAGULATION | 0.054489 | 0.181631 | -0.304923 | -1.45555 | 87 | 38 | c("SERPINA1", "MMP9", "PRSS23", "FN1", "CTSB", "LGMN", "APOC1", "SPARC", "TIMP3", "COMP", "PLAU") |
| 16 | HALLMARK_MYOGENESIS | 0.090888 | 0.284026 | 0.342425 | 1.35452 | 791 | 54 | c("MYH11", "KLF5", "MYLK", "ITGB4", "APOD", "NAV2", "DMD", "CRYAB", "LAMA2", "TPD52L1", "EFS", "EPHB3", "SVIL", "ABLIM1", "WWTR1", "IGFBP3", "TAGLN") |
| 17 | HALLMARK_XENOBIOTIC_METABOLISM | 0.119505 | 0.351484 | -0.27815 | -1.32775 | 192 | 38 | c("HMOX1", "BLVRB", "APOE", "PAPSS2", "DCXR", "GSR", "MCCC2", "DHRS7", "BCAR1", "EPHX1") |
| 18 | HALLMARK_WNT_BETA_CATENIN_SIGNALING | 0.137804 | 0.35882 | 0.471734 | 1.342035 | 1043 | 13 | c("MYC", "FZD8", "JAG2", "JAG1", "ADAM17", "CSNK1E", "MAML1", "NOTCH1", "HDAC2", "CCND2", "NUMB") |
| 19 | HALLMARK_ANDROGEN_RESPONSE | 0.143528 | 0.35882 | -0.255681 | -1.270225 | 213 | 44 | c("BMPR1B", "SCD", "STEAP4", "ABHD2", "GSR", "CCND1") |
| 20 | HALLMARK_FATTY_ACID_METABOLISM | 0.142324 | 0.35882 | -0.302374 | -1.311547 | 266 | 27 | c("BMPR1B", "FH", "PSME1", "GLUL", "EPHX1", "HSP90AA1", "LGALS1", "IDH3B", "ALDH9A1", "HSD17B4", "PDHA1") |
| 21 | HALLMARK_UV_RESPONSE_UP | 0.171026 | 0.407205 | -0.246122 | -1.222737 | 254 | 44 | c("HMOX1", "RET", "SQSTM1", "GRINA", "TFRC", "BSG", "PRKCD", "ARRB2", "HNRNPU", "ATP6V1F", "PPT1", "POLR2H", "EPHX1") |

| | | | | | | | | |
|---|---|---|---|---|---|---|---|---|
| 22 | HALLMARK_DNA_REPAIR | 0.24046 | 0.546501 | -0.250087 | -1.17291 | 396 | 36 | c("BRF2", "NME4", "CANT1", "NME1", "HCLS1", "POLD4", "NUDT21", "POLR2H", "GUK1", "NCBP2", "GPX4", "TMED2", "IMPDH2", "ADRM1", "SEC61A1", "POLR2I", "VPS28", "CSTF3", "DUT", "STX3", "ITPA", "DCTN4", "SURF1", "TAF10", "ERCC1", "TP53", "DGUOK", "SSRP1", "POLR1D", "XPC", "NFX1") |
| 23 | HALLMARK_HYPOXIA | 0.268655 | 0.559698 | 0.271062 | 1.146996 | 2422 | 80 | c("ERRFI1", "EGFR", "MT2A", "ATF3", "F3", "PNRC1", "AK4", "KLF7", "CCNG2", "MAFF", "NFIL3", "DDIT4", "JUN", "TES", "IRS2", "ANGPTL4", "IGFBP3", "TNFAIP3", "FOXO3", "MXI1", "KLHL24", "SDC4", "EXT1", "KLF6", "ZFP36", "CITED2", "NDST1", "ETS1", "PAM", "CCN2", "PDK1", "DUSP1", "FOS", "MAP3K1") |
| 24 | HALLMARK_BILE_ACID_METABOLISM | 0.26087 | 0.559698 | 0.376473 | 1.181527 | 2051 | 19 | c("TFCP2L1", "ABCA8", "BCAR3", "DIO2", "DHCR24") |
| 25 | HALLMARK_APICAL_SURFACE | 0.292689 | 0.561732 | 0.391023 | 1.154459 | 2241 | 15 | c("CX3CL1", "EFNA5", "CRYBG1") |
| 26 | HALLMARK_E2F_TARGETS | 0.30031 | 0.561732 | -0.232696 | -1.110777 | 484 | 38 | c("TOP2A", "ANP32E", "TFRC", "HMGB2", "NME1", "SNRPB", "NUDT21", "ILF3", "HNRNPD", "POLD2", "HELLS", "LUC7L3", "PA2G4", "NOP56", "PNN", "PRKDC", "DUT", "NUP153", "NUP107", "TUBB", "TBRG4", "TP53", "SRSF2", "SSRP1", "CDK4", "CNOT9", "MTHFD2", "SHMT1", "SMC3", "CDKN1B", "MCM7", "STAG1", "USP1") |
| 27 | HALLMARK_EPITHELIAL_MESENCHYMAL_TRANSITION | 0.303335 | 0.561732 | 0.25689 | 1.116461 | 2773 | 95 | c("OXTR", "SFRP1", "AREG", "MYLK", "DST", "ITGA2", "LAMC2", "PDLIM4", "LAMA3", "LAMA2", "FZD8", "MGP", "COL7A1", "ACTA2", "JUN", "ANPEP", "IGFBP3", "TAGLN", "TNFAIP3", "SNAI2", "IGFBP2", "TPM1", "TNC", "SDC4", "TPM2", "TPM4", "PMEPA1", "FBLN5", "MYL9", "SLIT2", "CALD1") |
| 28 | HALLMARK_CHOLESTEROL_HOMEOSTASIS | 0.484174 | 0.733596 | 0.303429 | 0.989771 | 3869 | 22 | c("ERRFI1", "ATF3", "PNRC1", "JAG1", "NFIL3", "TP53INP1") |
| 29 | HALLMARK_INTERFERON_ALPHA_RESPONSE | 0.473845 | 0.733596 | -0.233376 | -0.981765 | 932 | 24 | c("IFI30", "IFITM2", "TRIM25", "PSME1", "LGALS3BP", "B2M", "BST2") |
| 30 | HALLMARK_MYC_TARGETS_V2 | 0.417755 | 0.733596 | 0.354301 | 1.046039 | 3199 | 15 | c("DUSP2", "MYC", "WDR43", "MYBBP1A", "DDX18", "PPAN", "PES1", "SRM") |
| 31 | HALLMARK_GLYCOLYSIS | 0.469308 | 0.733596 | 0.255316 | 1.006086 | 4074 | 53 | c("MET", "DSC2", "EGFR", "SOX9", "ELF3", "AK4", "DDIT4", "IRS2", "ANGPTL4", "IGFBP3") |
| 32 | HALLMARK_UV_RESPONSE_DN | 0.451044 | 0.733596 | 0.248397 | 1.021903 | 4016 | 67 | c("KIT", "NFIB", "MET", "RND3", "IRS1", "F3", "MYC", "RBPMS", "ADGRL2", "ACVR2A", "SNAI2", "AKT3", "DDAH1", "PTEN", "ERBB2", "RUNX1", "DLC1", "FBLN5", "PHF3", "CITED2", "MGMT", "DUSP1", "SCAF8", "PLPP3", "ATXN1", "CAV1", "MIOS", "BHLHE40", "MAP1B", "ICA1", "TGFBR2", "CELF2", "CDC42BPA", "NIPBL", "ATRN", "CCN1", "ZMIZ1", "ADD3", "CAP2", "CDKN1B", "ATP2B4") |
| 33 | HALLMARK_ALLOGRAFT_REJECTION | 0.430452 | 0.733596 | -0.226195 | -1.020775 | 751 | 31 | c("MMP9", "ITGB2", "CTSS", "SPI1", "CAPG", "NME1", "TAPBP", "HCLS1", "B2M", "AKT1") |
| 34 | HALLMARK_MITOTIC_SPINDLE | 0.605363 | 0.850026 | 0.220268 | 0.919814 | 5417 | 74 | c("SORBS2", "DST", "NCK2", "CDC42EP1", "ARHGAP29", "ALS2", "SPTBN1", "ARHGAP5", "CDK5RAP2", "BCL2L11", "LLGL1", "AKAP13", "ALMS1", "ABL1", "RASAL2", "EPB41L2", "SUN2", "RALBP1", "TLK1", "MYO1E", "SOS1", "FARP1", "RABGAP1", "TRIO", "SHROOM1", "ARHGEF7", "WASF2", "SYNPO", "ATG4B", "ARHGEF12", "CD2AP", "PCM1", "PALLD", "HOOK3", "UXT", "KIF1B", "CDC42BPA", "VCL", "GSN", "PKD2", "SMC3") |
| 35 | HALLMARK_IL6_JAK_STAT3_SIGNALING | 0.616105 | 0.850026 | -0.220463 | -0.86534 | 1315 | 19 | c("HMOX1", "PIM1") |
| 36 | HALLMARK_PROTEIN_SECRETION | 0.581974 | 0.850026 | 0.260285 | 0.92607 | 4816 | 32 | c("GLA", "DST", "EGFR", "CAV2", "SCRN1", "ATP1A1", "VPS4B", "PAM", "ADAM10", "ERGIC3", "STX16", "ICA1", "TOM1L1", "KIF1B") |
| 37 | HALLMARK_ANGIOGENESIS | 0.62902 | 0.850026 | 0.301733 | 0.875525 | 4811 | 14 | c("PDGFA", "JAG2", "JAG1", "PTK2", "TNFRSF21", "APP") |
| 38 | HALLMARK_INTERFERON_GAMMA_RESPONSE | 0.648411 | 0.853172 | 0.231112 | 0.884182 | 5528 | 45 | c("MT2A", "TNFSF10", "ARID5B", "PDE4B", "PELI1", "AUTS2", "TXNIP", "TNFAIP3", "CASP4", "NCOA3", "FGL2", "SLC25A28") |
| 39 | HALLMARK_IL2_STAT5_SIGNALING | 0.689324 | 0.861655 | 0.216551 | 0.862232 | 6023 | 56 | c("TNFSF10", "MYC", "MAFF", "NFIL3", "AHR", "SPRY4", "HIPK2", "WLS", "TNFRSF21", "DHRS3", "PRNP", "NCOA3", "SPRED2", "LRIG1", "FGL2", "PLPP1", "KLF6", "SNX9", "IFNGR1", "MYO1E", "SWAP70", "BMPR2", "PLAGL1", "PLSCR1", "IRF6", "PLEC") |
| 40 | HALLMARK_SPERMATOGENESIS | 0.674505 | 0.861655 | 0.300654 | 0.838812 | 5076 | 12 | c("SLC12A2", "NPHP1", "IFT88", "STAM2") |
| 41 | HALLMARK_TGF_BETA_SIGNALING | 0.797045 | 0.905733 | 0.21519 | 0.749976 | 6526 | 29 | c("LTBP2", "BCAR3", "WWTR1", "HIPK2", "SPTBN1", "ACVR1", "PMEPA1", "BMPR2", "CDK9", "SKI", "TRIM33", "ENG", "SMURF2", "TGIF1", "KLF10") |
| 42 | HALLMARK_G2M_CHECKPOINT | 0.796087 | 0.905733 | 0.196161 | 0.781045 | 6956 | 56 | c("EFNA5", "DMD", "MT2A", "SLC12A2", "MYC", "MEIS1", "HOXC10", "SLC38A1", "SS18", "ABL1", "RASAL2", "SFPQ", "HUS1", "TFDP1") |
| 43 | HALLMARK_HEDGEHOG_SIGNALING | 0.793844 | 0.905733 | 0.286527 | 0.734051 | 5802 | 9 | c("TLE1", "ETS2", "DPYSL2", "ADGRG1", "PML", "RASA1", "MYH9") |
| 44 | HALLMARK_PI3K_AKT_MTOR_SIGNALING | 0.74858 | 0.905733 | 0.223421 | 0.794911 | 6195 | 32 | c("EGFR", "VAV3", "MAPK8", "SFN", "ITPR2") |
| 45 | HALLMARK_ADIPOGENESIS | 0.822177 | 0.91353 | 0.18749 | 0.762479 | 7258 | 63 | c("MYLK", "TANK", "SLC5A6", "CAVIN2", "SNCG", "CCNG2", "STAT5A", "ATL2", "PHLDB1", "ANGPTL4", "LIFR", "PDCD4") |

| | | | | | | | | |
|---|---|---|---|---|---|---|---|---|
| 46 | HALLMARK_APOPTOSIS | 0.92939 | 0.98255 | 0.159002 | 0.642507 | 8186 | 61 | c("EREG", "ATF3", "TNFSF10", "TXNIP", "JUN", "F2R", "EGR3", "PTK2", "ANXA1", "PDCD4", "BCL2L11", "CASP4", "ERBB2", "APP") |
| 47 | HALLMARK_MYC_TARGETS_V1 | 0.962899 | 0.98255 | -0.116308 | -0.630748 | 1115 | 64 | c("CLNS1A", "BUB3", "HNRNPU", "PCBP1", "NME1", "PSMD3", "GNL3", "HNRNPD", "POLD2", "NCBP2", "EIF3B", "IMPDH2", "PA2G4", "UBA2", "CUL1", "DHX15", "SRPK1", "NOP56", "RAD23B", "DUT", "APEX1", "VDAC1", "SF3B3", "HNRNPA3", "NHP2", "NOP16", "YWHAE", "EIF4H", "SMARCC1", "PSMD1", "RNPS1", "EIF4G2", "KPNB1", "HDGF", "FAM120A", "LSM2", "GLO1", "SRSF2", "SNRPD2", "CDK4", "SF3A1", "EIF3J", "MCM7", "CCT3", "SNRPA", "HNRNPR", "PHB2", "USP1", "SET", "SRM", "SRSF3", "HDAC2", "POLE3", "RACK1", "DDX18", "ORC2", "SERBP1", □ "EEF1B2", "TFDP1") |
| 48 | HALLMARK_PEROXISOME | 0.956296 | 0.98255 | -0.133061 | -0.566243 | 1837 | 25 | c("TOP2A", "TSPO", "FIS1") |
| 49 | HALLMARK_PANCREAS_BETA_CELLS | 0.929736 | 0.98255 | 0.309258 | 0.617147 | 6099 | 4 | c("AKT3", "SRP9") |
| 50 | HALLMARK_UNFOLDED_PROTEIN_RESPONSE | 0.995973 | 0.995973 | 0.112101 | 0.419027 | 8407 | 40 | c("ATF3", "ERN1", "DDIT4", "SLC1A4", "HERPUD1", "EXOC2") |

| Cluster 27 | pathway | pval | padj | ES | NES | nMoreExtreme | size | leadingEdge |
|---|---|---|---|---|---|---|---|---|
| 1 | HALLMARK_TNFA_SIGNALING_VIA_NFKB | 0.000167 | 0.004136 | 0.532634 | 2.564474 | 0 | 86 | c("FOSB", "NR4A3", "KLF4", "NR4A1", "SOCS3", "FOS", "ZFP36", "DUSP1", "ATF3", "JUNB", "G0S2", "MAFF", "EGR1", "KLF2", "EGR2", "IL7R", "NR4A2", "KLF9", "IER2", "ACKR3", "JUN", "PER1", "CCN1", "TRIP10", "NFIL3", "ETS2", "PPP1R15A", "CEBPD", "NAMPT", "CCL5", "DNAJB4", "CDKN1A", "TNFAIP3", "JAG1", "PLPP3", "GEM", "DRAM1", "KLF6", "PNRC1", "ICAM1", "PTPRE", "GADD45B", "FOSL2", "TIPARP", "CSF1", "TNFAIP2", "MCL1", "SOD2", "EGR3") |
| 2 | HALLMARK_ESTROGEN_RESPONSE_EARLY | 0.000246 | 0.004136 | -0.573437 | -2.76914 | 0 | 76 | c("RET", "ADCY1", "MAPT", "KRT19", "SLC39A6", "PGR", "KRT8", "TTC39A", "SLC9A3R1", "SLC2A1", "STC2", "TFF3", "SCNN1A", "SEC14L2", "XBP1", "GREB1", "CLDN7", "MUC1", "MLPH", "SIAH2", "MYB", "CA12", "ABHD2", "THSD4", "KDM4B", "SLC7A2", "FASN", "ELF3", "RHOD", "CCND1", "RHOBTB3", "TFF1", "CELSR1", "IGF1R", "GAB2", "MAST4", "CANT1", "RAB31", "TOB1", "GFRA1", "ELOVL5", "FLNB", "BAG1", "PRSS23") |
| 3 | HALLMARK_ESTROGEN_RESPONSE_LATE | 0.000248 | 0.004136 | -0.558036 | -2.631978 | 0 | 69 | c("SERPINA1", "TOP2A", "RET", "AGR2", "MAPT", "KRT19", "CDH1", "PGR", "PRLR", "SLC9A3R1", "S100A9", "TFF3", "SERPINA3", "SCNN1A", "LTF", "XBP1", "COX6C", "SIAH2", "MYB", "CA12", "ABHD2", "LLGL2", "CCND1", "IDH2", "TFF1", "TSPAN13", "DCXR", "MAPK13", "SCUBE2", "CD9", "LSR", "RAB31", "TOB1", "ELOVL5", "FLNB", "BAG1", "PRSS23") |
| 4 | HALLMARK_KRAS_SIGNALING_UP | 0.000686 | 0.008573 | 0.449966 | 1.955061 | 3 | 54 | c("KLF4", "F13A1", "APOD", "G0S2", "AKAP12", "IL7R", "GNG11", "CFH", "SPARCL1", "ADGRL4", "ANGPTL4", "TFPI", "ADGRA2", "EMP1", "EPB41L3", "TNFRSF1B", "PPP1R15A", "PECAM1", "IL10RA", "PLVAP", "GUCY1A1", "ENG", "SPRY2", "ETS1", "TNFAIP3", "CCND2", "DUSP6", "SCN1B", "GYPC", "NRP1", "C3AR1", "TMEM176B") |
| 5 | HALLMARK_ADIPOGENESIS | 0.003208 | 0.032078 | 0.384928 | 1.733053 | 18 | 62 | c("GPX3", "ITIH5", "FABP4", "ITGA7", "C3", "CAVIN2", "ADIPOQ", "SPARCL1", "ME1", "FZD4", "LIPE", "SNCG", "ANGPTL4", "LIFR", "CAVIN1", "CD36", "COL15A1", "LPL", "SCARB1") |
| 6 | HALLMARK_HYPOXIA | 0.003858 | 0.032153 | 0.362312 | 1.676388 | 22 | 71 | c("GPC3", "FOS", "ZFP36", "DUSP1", "CCN5", "ATF3", "CAV1", "MAFF", "DCN", "AKAP12", "EGFR", "CDKN1C", "SRPX", "ACKR3", "JUN", "ANGPTL4", "CCN1", "CAVIN1", "CAVIN3", "NFIL3", "PPP1R15A", "SCARB1", "ETS1", "CDKN1A", "TNFAIP3", "PAM", "RBPJ", "KLF6", "PNRC1", "NDRG1", "FOSL2", "TIPARP", "PDGFB", "NR3C1") |
| 7 | HALLMARK_ANDROGEN_RESPONSE | 0.005573 | 0.039808 | -0.443879 | -1.84011 | 22 | 38 | c("BMPR1B", "SPDEF", "KRT19", "TPD52", "KRT8", "INPP4B", "AZGP1", "ABHD2", "CCND1", "DHCR24", "SCD") |
| 8 | HALLMARK_MYOGENESIS | 0.019922 | 0.099612 | 0.340894 | 1.534802 | 117 | 62 | c("CFD", "GPX3", "FXYD1", "PTGIS", "FHL1", "APOD", "MYH11", "ITGA7", "SOD3", "GSN", "CRYAB", "ABLIM1", "IGF1", "LAMA2", "COX7A1", "BIN1", "CD36", "COL15A1", "MEF2C", "CDKN1A", "PRNP", "PTP4A3", "IGFBP7", "COL4A2", "GADD45B", "AK1") |
| 9 | HALLMARK_INTERFERON_GAMMA_RESPONSE | 0.016257 | 0.099612 | 0.373002 | 1.589676 | 95 | 49 | c("SOCS3", "C1R", "SERPING1", "C1S", "CFH", "ST3GAL5", "MX2", "ARID5B", "IL10RA", "NAMPT", "VAMP5", "CCL5", "CDKN1A", "TNFAIP3", "SAMHD1", "ICAM1", "FGL2", "TXNIP", "TNFAIP2", "SOD2", "IRF8", "CASP4", "JAK2", "APOL6", "PML", "IFITM3") |
| 10 | HALLMARK_INFLAMMATORY_RESPONSE | 0.018021 | 0.099612 | 0.384925 | 1.585792 | 105 | 42 | c("GPC3", "STAB1", "IL7R", "CD14", "TNFRSF1B", "IL10RA", "NAMPT", "CCL5", "CDKN1A", "SCN1B", "KLF6", "C5AR1", "C3AR1", "LPAR1", "OSMR", "ICAM1", "PTPRE", "CSF1") |
| 11 | HALLMARK_IL6_JAK_STAT3_SIGNALING | 0.033779 | 0.140746 | 0.449727 | 1.551272 | 196 | 21 | c("SOCS3", "LEPR", "ACVRL1", "CD14", "A2M", "JUN", "TNFRSF1B", "CD36", "OSMR", "CSF1", "TNFRSF1A", "IL1R1", "TNFRSF21", "IFNGR1") |
| 12 | HALLMARK_UV_RESPONSE_UP | 0.033625 | 0.140746 | 0.392661 | 1.52945 | 195 | 33 | c("FOSB", "GPX3", "NR4A1", "FOS", "ATF3", "JUNB", "CDKN1C") |
| 13 | HALLMARK_HEME_METABOLISM | 0.039059 | 0.150226 | -0.360655 | -1.524301 | 160 | 41 | c("SLC4A1", "HBD", "HBB", "SLC2A1", "SLC25A37", "ACP5") |
| 14 | HALLMARK_OXIDATIVE_PHOSPHORYLATION | 0.060092 | 0.19848 | -0.3511 | -1.455492 | 247 | 38 | c("COX6C", "NDUFC2", "IDH2", "MAOB", "ATP1B1", "COX6A1", "UQCRQ", "NDUFS2", "TCIRG1", "UQCRC1", "ECI1", "FH", "UQCR10", "ATP6V0B", "NDUFAB1", "ATP5PD", "VDAC3", "TIMM13", "NDUFA4", "MGST3", "NDUFA2", "ATP5MC2") |
| 15 | HALLMARK_UV_RESPONSE_DN | 0.063514 | 0.19848 | 0.306178 | 1.386053 | 375 | 64 | c("DUSP1", "TGFBR3", "GRK5", "CAV1", "FBLN5", "TGFBR2", "ID1", "DAB2", "FYN", "TFPI", "EFEMP1", "CCN1", "DLC1", "ADD3", "PMP22", "PLPP3", "PTPRM", "LAMC1", "NRP1", "LPAR1", "TJP1", "NFIB", "PDGFRB", "NR3C1", "MAP1B", "SMAD7", "PPARG", "CELF2", "MGLL", "NR1D2") |

| | | | | | | | | |
|---|---|---|---|---|---|---|---|---|
| 16 | HALLMARK_COAGULATION | 0.055821 | 0.19848 | 0.331435 | 1.423868 | 326 | 51 | c("CFD", "GSN", "MAFF", "C3", "CFI", "C1R", "SERPING1", "C1S", "VWF", "RAPGEF3", "CFH", "C1QA", "C2", "FYN", "A2M", "MMP2", "THBD", "PECAM1", "LGMN", "DUSP6", "LRP1", "FBN1", "ANXA1", "PDGFB", "CTSK", "ISCU", "GNG12") |
| 17 | HALLMARK_MTORC1_SIGNALING | 0.082384 | 0.242307 | -0.336317 | -1.394211 | 339 | 38 | c("SLC9A3R1", "SLC2A1", "EGLN3", "XBP1", "IFI30", "DHCR24", "SCD", "CD9", "STC1", "ELOVL5", "NUPR1", "SYTL2", "ACACA", "LTA4H", "ACSL3") |
| 18 | HALLMARK_UNFOLDED_PROTEIN_RESPONSE | 0.110081 | 0.275202 | -0.369145 | -1.36941 | 462 | 24 | c("STC2", "XBP1", "LSM1", "EIF4EBP1", "DNAJA4", "NHP2", "LSM4", "VEGFA", "SPCS1", "EIF4A3", "EIF4G1") |
| 19 | HALLMARK_E2F_TARGETS | 0.104137 | 0.275202 | -0.420775 | -1.408479 | 437 | 17 | c("TOP2A", "SMC4", "ANP32E", "NME1", "HMGB2", "MCM7", "SNRPB") |
| 20 | HALLMARK_EPITHELIAL_MESENCHYMAL_TRANSITION | 0.10759 | 0.275202 | 0.253736 | 1.268601 | 644 | 107 | c("SFRP1", "SLIT3", "TGFBR3", "ABI3BP", "DCN", "FBLN5", "FBLN2", "FBLN1", "GREM1", "LAMA2", "IGFBP4", "FSTL1", "DAB2", "JUN", "MMP2", "GAS1", "CXCL12", "CCN1", "SFRP4", "ELN", "TNFAIP3", "SLIT2", "PMP22", "PCOLCE", "FERMT2", "GLIPR1", "GEM", "NNMT", "LRP1", "ECM2", "LAMC1", "THBS2", "MYL9", "CALD1", "MFAP5", "FBN1", "COL4A2", "GADD45B") |
| 21 | HALLMARK_MYC_TARGETS_V1 | 0.137716 | 0.318938 | -0.340852 | -1.324049 | 573 | 29 | c("CLNS1A", "NME1", "HSP90AB1", "HNRNPC", "HDGF", "NHP2", "IMPDH2", "MCM7", "LSM7", "CBX3", "NDUFAB1", "CANX", "VDAC3", "PPIA") |
| 22 | HALLMARK_BILE_ACID_METABOLISM | 0.140333 | 0.318938 | 0.453952 | 1.323527 | 809 | 12 | c("ABCA6", "ABCA8", "LIPE", "ALDH1A1") |
| 23 | HALLMARK_TGF_BETA_SIGNALING | 0.154629 | 0.336149 | 0.354401 | 1.281992 | 896 | 25 | c("JUNB", "ID1", "CDKN1C", "ID3", "PPP1R15A", "ENG", "SMAD1", "TJP1", "SKI", "SPTBN1", "LTBP2", "KLF10", "SMAD7", "SMURF2") |
| 24 | HALLMARK_WNT_BETA_CATENIN_SIGNALING | 0.173998 | 0.347995 | 0.589754 | 1.272195 | 984 | 5 | c("CCND2", "JAG1", "RBPJ", "NOTCH1") |
| 25 | HALLMARK_APOPTOSIS | 0.16828 | 0.347995 | 0.278277 | 1.226647 | 990 | 57 | c("GPX3", "TGFBR3", "ATF3", "IGFBP6", "CAV1", "GSN", "DCN", "CD14", "JUN", "MMP2", "EMP1", "LMNA", "CDKN1A", "CCND2", "ANXA1", "GADD45B", "TXNIP", "PDGFRB", "MCL1", "SOD2", "EGR3", "BTG2", "CASP4", "SMAD7") |
| 26 | HALLMARK_PEROXISOME | 0.197425 | 0.379663 | -0.3803 | -1.24795 | 827 | 16 | c("TOP2A", "CRABP2", "IDH2", "DHCR24", "ELOVL5") |
| 27 | HALLMARK_COMPLEMENT | 0.25253 | 0.467649 | 0.251601 | 1.137797 | 1496 | 63 | c("MAFF", "C3", "C1R", "SERPING1", "C1S", "CTSC", "CFH", "ME1", "C1QA", "C2", "C1QC", "FYN", "GNG2", "CD36", "CCL5", "PRCP", "TNFAIP3", "LGMN", "DUSP6", "LRP1", "DOCK9", "SH2B3", "COL4A2", "PDGFB", "CBLB") |
| 28 | HALLMARK_P53_PATHWAY | 0.290698 | 0.519103 | 0.247516 | 1.1111 | 1724 | 61 | c("KLF4", "FOS", "ATF3", "ZBTB16", "JUN", "TSPYL2", "PPP1R15A", "TCN2", "SESN1", "COQ8A", "CDKN1A", "CCND2", "PLK3", "PITPNC1", "DRAM1", "NDRG1", "PTPRE", "AK1", "TXNIP", "ISCU", "S100A4", "BTG2", "NOTCH1", "ITGB4", "TAX1BP3", "XPC", "CD81", "CTSF", "FBXW7", "MXD4", "ZFP36L1", "STOM", "EPHX1") |
| 29 | HALLMARK_GLYCOLYSIS | 0.313692 | 0.540849 | -0.287723 | -1.106898 | 1298 | 28 | c("STC2", "TFF3", "EGLN3", "SDC1", "ELF3", "MIF", "STC1") |
| 30 | HALLMARK_DNA_REPAIR | 0.37708 | 0.628467 | -0.317813 | -1.06383 | 1585 | 17 | c("BRF2", "NME4", "CANT1", "NME1", "POLR2H", "IMPDH2", "POLD4", "BCAM", "TAF10") |
| 31 | HALLMARK_CHOLESTEROL_HOMEOSTASIS | 0.648591 | 0.746307 | -0.244893 | -0.860896 | 2738 | 20 | c("FASN", "ALCAM", "SCD", "CD9", "PMVK", "ACTG1") |
| 32 | HALLMARK_G2M_CHECKPOINT | 0.468936 | 0.746307 | -0.263509 | -0.986907 | 1969 | 25 | c("TOP2A", "SMC4", "CCND1") |
| 33 | HALLMARK_NOTCH_SIGNALING | 0.613151 | 0.746307 | 0.349853 | 0.886309 | 3505 | 8 | c("JAG1", "FZD7", "NOTCH3", "TCF7L2", "NOTCH1", "ARRB1") |
| 34 | HALLMARK_PROTEIN_SECRETION | 0.580336 | 0.746307 | -0.255508 | -0.910673 | 2419 | 21 | c("TOM1L1", "TPD52") |
| 35 | HALLMARK_APICAL_JUNCTION | 0.643347 | 0.746307 | 0.206119 | 0.895568 | 3751 | 54 | c("CD209", "CLDN5", "CD34", "EGFR", "VWF", "MMP2", "LIMA1", "PECAM1", "EPB41L2", "TSPAN4", "SLIT2", "MYL9", "CD99", "FBN1", "ICAM1", "TJP1", "RRAS", "MAP3K20", "GNAI2", "ARHGEF6", "ITGB4", "PARVA", "SIRPA", "ITGB1") |
| 36 | HALLMARK_HEDGEHOG_SIGNALING | 0.656751 | 0.746307 | 0.353135 | 0.857529 | 3730 | 7 | c("DPYSL2", "ETS2", "NRP1", "PML") |
| 37 | HALLMARK_MYC_TARGETS_V2 | 0.478281 | 0.746307 | -0.473917 | -0.982203 | 2091 | 4 | c("CBX3", "PA2G4", "MYC", "SRM") |
| 38 | HALLMARK_XENOBIOTIC_METABOLISM | 0.511611 | 0.746307 | -0.225193 | -0.964009 | 2114 | 43 | c("ESR1", "SPINT2", "APOE", "HMOX1", "CFB", "DCXR", "NQO1", "FBP1", "ELOVL5", "JUP", "PGD", "MCCC2") |
| 39 | HALLMARK_FATTY_ACID_METABOLISM | 0.606424 | 0.746307 | -0.228802 | -0.895317 | 2529 | 30 | c("BMPR1B", "FASN", "DHCR24", "MIF") |
| 40 | HALLMARK_REACTIVE_OXYGEN_SPECIES_PATHWAY | 0.591244 | 0.746307 | -0.278776 | -0.896821 | 2484 | 15 | c("MPO", "NQO1", "PRDX2", "NDUFS2", "ATOX1", "EGLN2") |
| 41 | HALLMARK_ANGIOGENESIS | 0.582263 | 0.746307 | -0.270594 | -0.905771 | 2448 | 17 | c("SPP1", "POSTN", "STC1") |
| 42 | HALLMARK_IL2_STAT5_SIGNALING | 0.649415 | 0.746307 | 0.203052 | 0.891701 | 3826 | 56 | c("ITIH5", "SHE", "MAFF", "CDKN1C", "DHRS3", "EMP1", "PLPP1", "TNFRSF1B", "NFIL3", "PLAGL1", "IL10RA", "CCND2", "PRNP", "NRP1", "KLF6", "NDRG1", "GADD45B", "FGL2", "CSF1") |
| 43 | HALLMARK_SPERMATOGENESIS | 0.532415 | 0.746307 | 0.444619 | 0.959115 | 3013 | 5 | c("SHE", "MAST2", "AGFG1") |
| 44 | HALLMARK_KRAS_SIGNALING_DN | 0.549088 | 0.746307 | -0.325303 | -0.924931 | 2348 | 10 | c("BMPR1B", "EFHD1") |
| 45 | HALLMARK_ALLOGRAFT_REJECTION | 0.69017 | 0.766855 | 0.217109 | 0.84566 | 4022 | 33 | c("EGFR", "STAB1", "C2", "CCL5", "ETS1", "CCND2", "NCK1", "ICAM1", "CSF1", "ELF4", "CD4", "SOCS5", "HCLS1", "IRF8", "JAK2", "GBP2", "IFNGR1", "TAP2") |
| 46 | HALLMARK_APICAL_SURFACE | 0.718553 | 0.781036 | -0.299976 | -0.792565 | 3078 | 8 | GATA3 |

| | | | | | | | | |
|---|---|---|---|---|---|---|---|---|
| 47 | HALLMARK_PANCREAS_BETA_CELLS | 0.74659 | 0.794244 | 0.311796 | 0.789896 | 4268 | 8 | c("STXBP1", "FOXO1", "LMO2") |
| 48 | HALLMARK_INTERFERON_ALPHA_RESPONSE | 0.883313 | 0.920118 | 0.187997 | 0.689013 | 5139 | 26 | c("C1S", "NCOA7", "CSF1", "TXNIP", "LPAR6", "TENT5A", "GBP2", "UBA7", "IFITM3", "TRIM25", "LAP3", "SP110", "PARP14", "CMTR1", "PSMB8", "LY6E", "IRF1") |
| 49 | HALLMARK_PI3K_AKT_MTOR_SIGNALING | 0.920827 | 0.93962 | -0.183416 | -0.613958 | 3872 | 17 | c("SLC2A1", "VAV3") |
| 50 | HALLMARK_MITOTIC_SPINDLE | 0.999319 | 0.999319 | 0.093243 | 0.398627 | 5870 | 50 | c("GSN", "BIN1", "ABL1", "EPB41L2", "SYNPO", "NCK1", "PKD2", "ARHGAP29", "SPTBN1", "BCR", "NCK2", "DST", "PXN", "SPTAN1", "PDLIM5", "RASAL2", "ARHGEF7", "ARHGDIA", "KIF1B", "SUN2", "CAPZB", "TRIO", "DOCK2", "KLC1", "AKAP13", "FLNA", "MAP3K11", "DOCK4") |

| Cluster 28 | pathway | pval | padj | ES | NES | nMoreExtreme | size | leadingEdge |
|---|---|---|---|---|---|---|---|---|
| 1 | HALLMARK_TNFA_SIGNALING_VIA_NFKB | 0.000474 | 0.009193 | -0.480376 | -2.404007 | 0 | 52 | c("GEM", "NR4A1", "FOSB", "DUSP1", "FOS", "MAP3K8", "JUN", "EGR1", "CCN1", "ZFP36", "BHLHE40", "BTG2", "PHLDA1", "EIF1", "CEBPD", "KLF6", "RHOB", "JUNB", "IER2", "PLK2", "MCL1", "SOD2", "CEBPB", "PFKFB3", "SERPINE1") |
| 2 | HALLMARK_ESTROGEN_RESPONSE_EARLY | 0.000552 | 0.009193 | -0.65716 | -3.570009 | 0 | 74 | c("GREB1", "HSPB8", "STC2", "PGR", "SLC39A6", "MAPT", "SLC7A2", "GAB2", "TOB1", "FOS", "KRT8", "ADCY1", "SEC14L2", "SLC9A3R1", "ELF3", "FASN", "RHOBTB3", "LRIG1", "SCNN1A", "CELSR1", "SIAH2", "RET", "RARA", "SLC2A1", "MYB", "BHLHE40", "NRIP1", "KDM4B", "PRSS23", "GFRA1", "MED13L", "ELOVL5", "CA12", "IGF1R", "XBP1", "MUC1", "AKAP1", "UGCG", "FLNB", "BLVRB", "MLPH", "CLDN7", "ITPK1", "TTC39A", "FHL2") |
| 3 | HALLMARK_ESTROGEN_RESPONSE_LATE | 0.000521 | 0.009193 | -0.551437 | -2.913374 | 0 | 66 | c("HSPB8", "SERPINA1", "CDH1", "SERPINA3", "PGR", "MAPT", "TOB1", "CXCL14", "FOS", "PRLR", "S100A9", "SLC9A3R1", "COX6C", "LTF", "SCNN1A", "SIAH2", "RET", "ZFP36", "SCUBE2", "MYB", "NRIP1", "PRSS23", "LLGL2", "DLG5", "ELOVL5", "CA12", "TOP2A", "XBP1", "FLNB", "DCXR", "BLVRB", "ITPK1", "UGDH", "LSR", "KRT19", "BCL2", "NCOR2", "RABEP1", "SEMA3B", "CD44", "ADD3", "FAM102A", "CHPT1", "BAG1", "CCND1") |
| 4 | HALLMARK_EPITHELIAL_MESENCHYMAL_TRANSITION | 0.000943 | 0.011788 | 0.400775 | 1.813849 | 7 | 96 | c("CDH2", "ITGB3", "COL1A1", "PCOLCE", "COL1A2", "CADM1", "SPARC", "SPP1", "TNC", "FZD8", "SERPINH1", "ANPEP", "SLIT2", "PLOD2", "ITGA2", "GJA1", "COL5A1", "TGFB1", "VCAM1", "SERPINE2", "PFN2", "SGCB", "LRRC15", "BMP1", "COL4A2", "SNAI2", "EMP3", "COL4A1", "P3H1", "TGFBI", "PMEPA1", "EDIL3", "TNFRSF12A", "COL16A1", "COL12A1", "PLOD1", "LOX", "MMP14", "ITGAV", "CDH11", "GPC1", "PPIB", "COL5A2", "PDLIM4", "FBN1", "TPM1", "NNMT", "EFEMP2", "NOTCH2", "ITGB5", "ID2", "BGN", "SLIT3", "SDC1") |
| 5 | HALLMARK_HYPOXIA | 0.003717 | 0.037175 | -0.323791 | -1.721704 | 6 | 68 | c("STC1", "STC2", "DUSP1", "FOS", "TPD52", "JUN", "CCN1", "SIAH2", "SLC2A1", "ZFP36", "CCNG2", "BHLHE40", "MAP3K1", "MIF", "CCN2", "KLF6", "CA12", "DTNA") |
| 6 | HALLMARK_REACTIVE_OXYGEN_SPECIES_PATHWAY | 0.015959 | 0.119966 | -0.489024 | -1.752368 | 49 | 15 | c("MPO", "NQO1", "JUNB", "FTL", "SOD2", "EGLN2", "PRDX2") |
| 7 | HALLMARK_HEME_METABOLISM | 0.016795 | 0.119966 | -0.326321 | -1.607485 | 36 | 48 | c("HBB", "SLC4A1", "HBD", "C3", "FBXO7", "SLC2A1", "SLC25A37", "SNCA", "BTG2", "BLVRB", "BCAM", "CIR1", "FBXO9", "VEZF1", "HAGH", "PRDX2", "TNS1", "GDE1") |
| 8 | HALLMARK_MTORC1_SIGNALING | 0.051246 | 0.311693 | -0.286106 | -1.426157 | 108 | 51 | c("STC1", "SCD", "SLC9A3R1", "NUPR1", "SLC2A1", "BHLHE40", "EGLN3", "BTG2", "ELOVL5", "XBP1", "CXCR4", "GGA2", "UBE2D3", "GSK3B", "LTA4H") |
| 9 | HALLMARK_UV_RESPONSE_DN | 0.056105 | 0.311693 | 0.367859 | 1.457321 | 436 | 47 | c("ITGB3", "COL1A1", "ID1", "COL1A2", "MMP16", "KIT", "LTBP1", "SDC2", "MAP1B", "GJA1", "SMAD7", "LPAR1", "SNAI2", "KCNMA1", "PTPRM", "SMAD3") |
| 10 | HALLMARK_COMPLEMENT | 0.087154 | 0.435769 | -0.273232 | -1.345965 | 191 | 48 | c("SERPINA1", "S100A9", "C3", "LTF", "CFB", "GATA3", "APOC1", "CD55") |
| 11 | HALLMARK_WNT_BETA_CATENIN_SIGNALING | 0.130243 | 0.517466 | 0.51596 | 1.359012 | 856 | 10 | c("HEY2", "FZD8", "JAG1", "HDAC5", "HDAC2", "ADAM17", "NOTCH1") |
| 12 | HALLMARK_ANDROGEN_RESPONSE | 0.157161 | 0.517466 | -0.265031 | -1.250136 | 371 | 40 | c("BMPR1B", "INPP4B", "KRT8", "SCD", "TPD52", "STEAP4", "ELOVL5", "AZGP1", "SPDEF") |
| 13 | HALLMARK_MYOGENESIS | 0.152778 | 0.517466 | -0.243706 | -1.235995 | 318 | 55 | c("APOD", "PVALB", "HSPB8", "STC2", "SCD", "TNNT1", "NQO1", "BHLHE40", "ERBB3") |
| 14 | HALLMARK_APICAL_SURFACE | 0.165589 | 0.517466 | -0.46171 | -1.305708 | 588 | 8 | c("SULF2", "GATA3", "HSPB1") |
| 15 | HALLMARK_FATTY_ACID_METABOLISM | 0.128 | 0.517466 | -0.293122 | -1.30338 | 319 | 32 | c("BMPR1B", "GLUL", "FASN", "MIF", "ELOVL5", "FH") |
| 16 | HALLMARK_UV_RESPONSE_UP | 0.163522 | 0.517466 | -0.252239 | -1.227859 | 363 | 46 | c("NR4A1", "FOSB", "FOS", "RET", "BTG2", "RHOB", "EPCAM", "JUNB", "ARRB2", "SOD2", "GRINA", "CREG1", "HNRNPU") |
| 17 | HALLMARK_CHOLESTEROL_HOMEOSTASIS | 0.310399 | 0.756492 | -0.318563 | -1.117858 | 996 | 14 | c("SCD", "FASN", "ALCAM", "ACTG1") |
| 18 | HALLMARK_TGF_BETA_SIGNALING | 0.329394 | 0.756492 | 0.318452 | 1.116898 | 2424 | 28 | c("SMAD6", "ID1", "ID3", "SMAD7", "TGFB1", "TGFBR1", "PMEPA1", "ACVR1", "SMAD3", "ENG", "ID2", "FURIN", "TJP1", "LTBP2") |
| 19 | HALLMARK_APICAL_JUNCTION | 0.33806 | 0.756492 | 0.266148 | 1.100883 | 2693 | 58 | c("MMP9", "ITGA10", "CERCAM", "SLIT2", "ITGA2", "LDLRAP1", "FSCN1", "RAC2", "VCAM1", "BMP1", "CD276", "TGFBI", "COL16A1", "CAP1", "CNN2", "CDH11", "EPB41L2", "VCL", "SRC", "FBN1", "ICAM1", "TJP1", "ARHGEF6", "HRAS", "CTNND1", "CD34", "ARPC2", "PTK2", "SHC1", "INPPL1", "VCAN", "GNAI2", "PKD1", "CD99", "NF2", "TAOK2", "MYH10") |
| 20 | HALLMARK_INFLAMMATORY_RESPONSE | 0.322743 | 0.756492 | 0.302776 | 1.11733 | 2437 | 34 | c("ITGB3", "SGMS2", "SCARF1", "IL4R", "LPAR1", "SPHK1", "EMP3", "CDKN1A", "PCDH7") |
| 21 | HALLMARK_ALLOGRAFT_REJECTION | 0.330597 | 0.756492 | 0.326913 | 1.116984 | 2415 | 25 | c("MMP9", "SOCS1", "IL4R", "TGFB1", "TAP2", "CD4", "DEGS1", "ICAM1", "C2", "ETS1", "CSF1") |

| | | | | | | | | |
|---|---|---|---|---|---|---|---|---|
| 22 | HALLMARK_KRAS_SIGNALING_UP | 0.347986 | 0.756492 | 0.291512 | 1.094315 | 2643 | 37 | c("MMP9", "ANKH", "CLEC4A", "SPP1", "ITGA2", "ALDH1A3", "ANGPTL4", "VWA5A", "CA2") |
| 23 | HALLMARK_PANCREAS_BETA_CELLS | 0.318468 | 0.756492 | 0.531941 | 1.132174 | 1970 | 5 | c("VDR", "SRPRB", "SEC11A", "LMO2") |
| 24 | HALLMARK_APOPTOSIS | 0.389327 | 0.778654 | -0.211923 | -1.043206 | 838 | 49 | c("JUN", "RARA", "HMGB2", "BTG2", "ERBB3", "HSPB1", "RHOB", "TOP2A", "TIMP3", "MCL1", "SOD2", "ERBB2", "DAP3", "CREBBP", "CD44") |
| 25 | HALLMARK_INTERFERON_GAMMA_RESPONSE | 0.375033 | 0.778654 | 0.290494 | 1.072006 | 2832 | 34 | c("SOCS1", "IL4R", "VCAM1", "ST3GAL5", "PSME2", "CFH", "PTPN2", "CDKN1A", "VAMP5", "PNP", "ZNFX1", "IFI44", "ICAM1") |
| 26 | HALLMARK_IL6_JAK_STAT3_SIGNALING | 0.436147 | 0.838743 | 0.309941 | 1.025797 | 3141 | 22 | c("ITGB3", "SOCS1", "IL4R", "TGFB1", "PTPN2", "TNFRSF12A", "BAK1") |
| 27 | HALLMARK_COAGULATION | 0.470957 | 0.840994 | 0.259984 | 0.997829 | 3607 | 41 | c("MMP9", "ITGB3", "CTSK", "SPARC", "ITGA2", "BMP1", "CFH", "THBD", "S100A1", "ARF4", "MMP14") |
| 28 | HALLMARK_IL2_STAT5_SIGNALING | 0.460651 | 0.840994 | 0.249273 | 1.008475 | 3634 | 52 | c("PTH1R", "SPP1", "NCS1", "SOCS1", "IL4R", "CA2", "TIAM1", "ENPP1", "CKAP4", "SPRY4", "P4HA1", "S100A1", "PLPP1", "AGER", "PRAF2", "ITGAV", "PNP", "TWSG1", "MYO1E") |
| 29 | HALLMARK_DNA_REPAIR | 0.5 | 0.862069 | -0.225465 | -0.974781 | 1283 | 29 | c("BCAM", "TMED2", "IMPDH2", "CANT1", "GPX4", "TAF10", "POLR2A", "APRT", "DDB1", "POLR2K", "ADRM1", "STX3", "TP53", "ERCC1", "POLR2E", "NME3", "SSRP1", "RBX1", "NCBP2", "AAAS", "AK1", "SEC61A1", "CSTF3", "BCAP31", "ITPA", "POLE4", "ELOA", "PNP", "TYMS") |
| 30 | HALLMARK_ADIPOGENESIS | 0.578455 | 0.903329 | -0.183479 | -0.942149 | 1175 | 58 | c("HSPB8", "TOB1", "SPARCL1", "C3", "CCNG2", "COX6A1", "GPAT4") |
| 31 | HALLMARK_INTERFERON_ALPHA_RESPONSE | 0.631813 | 0.903329 | 0.28069 | 0.868912 | 4404 | 17 | c("IL4R", "TENT5A", "PSME2", "IFI44", "MVB12A", "LPAR6", "CSF1", "LAP3", "LGALS3BP", "CNP") |
| 32 | HALLMARK_HEDGEHOG_SIGNALING | 0.580241 | 0.903329 | 0.387136 | 0.914135 | 3705 | 7 | c("HEY2", "NRP2") |
| 33 | HALLMARK_GLYCOLYSIS | 0.63233 | 0.903329 | 0.224635 | 0.885938 | 4916 | 46 | c("KDELR3", "PLOD2", "SDC2", "EXT1", "ANGPTL4", "COL5A1", "TPST1", "B3GALT6", "TGFBI", "P4HA2", "SLC25A13", "P4HA1", "PLOD1", "EXT2", "GPC1", "SLC16A3", "B4GALT2", "AGRN") |
| 34 | HALLMARK_ANGIOGENESIS | 0.571429 | 0.903329 | 0.295795 | 0.915672 | 3983 | 17 | c("PDGFA", "SPP1", "JAG1", "THBD", "LPL", "FGFR1", "ITGAV", "COL5A2", "S100A4") |
| 35 | HALLMARK_PEROXISOME | 0.614013 | 0.903329 | 0.270801 | 0.886791 | 4407 | 21 | c("CADM1", "CTPS1", "ACSL4", "CLN8") |
| 36 | HALLMARK_MYC_TARGETS_V2 | 0.671277 | 0.91972 | 0.319476 | 0.841482 | 4416 | 10 | c("SRM", "PPRC1", "CDK4", "PA2G4", "MCM4", "FARSA", "NOP56", "NOP16") |
| 37 | HALLMARK_XENOBIOTIC_METABOLISM | 0.680593 | 0.91972 | -0.179912 | -0.875781 | 1514 | 46 | c("CFB", "ESR1", "SPINT2", "NQO1", "MCCC2", "ELOVL5", "DCXR", "BLVRB", "UGDH", "MAN1A1", "JUP", "FBP1", "SERPINE1", "CYB5A") |
| 38 | HALLMARK_P53_PATHWAY | 0.715872 | 0.941937 | 0.203749 | 0.835698 | 5673 | 56 | c("EPHA2", "PDGFA", "VDR", "SOCS1", "PTPN14", "VWA5A", "TGFB1", "SPHK1", "ZMAT3", "CDKN1A", "BAK1", "BLCAP", "STEAP3", "S100A4", "MXD4", "SEC61A1", "AK1", "NDRG1", "SDC1", "CYFIP2", "CD82", "NOTCH1", "HRAS") |
| 39 | HALLMARK_KRAS_SIGNALING_DN | 0.759694 | 0.973967 | 0.267402 | 0.766646 | 5171 | 13 | c("COPZ2", "FGFR3", "ENTPD7") |
| 40 | HALLMARK_MITOTIC_SPINDLE | 0.848251 | 0.997206 | 0.18392 | 0.725363 | 6595 | 46 | c("CDC42EP1", "FSCN1", "TIAM1", "ARHGAP10", "CDK5RAP2", "MARCKS", "BCAR1", "DOCK4", "EPB41L2", "VCL", "NCK2", "MYO1E", "CCDC88A", "NOTCH2", "PDLIM5") |
| 41 | HALLMARK_G2M_CHECKPOINT | 0.799672 | 0.997206 | 0.219156 | 0.748803 | 5843 | 25 | c("TGFB1", "SNRPD1", "SFPQ", "MARCKS", "SMAD3", "CDK4", "TFDP1", "NOTCH2", "ODF2", "EWSR1", "FOXN3") |
| 42 | HALLMARK_NOTCH_SIGNALING | 0.825988 | 0.997206 | 0.268013 | 0.705933 | 5434 | 10 | c("NOTCH3", "JAG1", "NOTCH2", "NOTCH1", "LFNG", "RBX1", "TCF7L2", "CUL1") |
| 43 | HALLMARK_PROTEIN_SECRETION | 0.967806 | 0.997206 | -0.128281 | -0.56537 | 2434 | 31 | c("TPD52", "TOM1L1", "TMED2", "VPS45", "LAMP2", "RAB2A", "GOSR2", "SEC31A", "ARCN1", "STX12", "DST", "ERGIC3", "ABCA1", "SEC22B", "M6PR", "ADAM10", "VAMP3", "YKT6", "DNM1L", "COPB1", "RER1", "IGF2R", "LMAN1", "ARFGAP3", "TMED10", "CLTA", "CD63", "SEC24D", "ATP6V1H", "CTSC", "ATP1A1") |
| 44 | HALLMARK_UNFOLDED_PROTEIN_RESPONSE | 0.98634 | 0.997206 | 0.128542 | 0.476747 | 7436 | 35 | c("KDELR3", "SSR1", "PDIA5", "ALDH18A1", "SRPRB", "SPCS3", "YIF1A", "HYOU1", "SEC11A", "TATDN2", "HERPUD1") |
| 45 | HALLMARK_PI3K_AKT_MTOR_SIGNALING | 0.883513 | 0.997206 | -0.162164 | -0.682992 | 2373 | 26 | c("SLC2A1", "CXCR4", "GRB2", "UBE2D3", "VAV3", "GSK3B") |
| 46 | HALLMARK_E2F_TARGETS | 0.982371 | 0.997206 | 0.143202 | 0.47395 | 7076 | 22 | c("CTPS1", "PRDX4", "CDKN1A", "CDK4", "POLE4", "CNOT9", "RPA1", "PA2G4", "SSRP1", "POLD2", "MCM4", "NOP56", "TRA2B") |
| 47 | HALLMARK_MYC_TARGETS_V1 | 0.995556 | 0.997206 | 0.109447 | 0.441087 | 7839 | 51 | c("CTPS1", "SRM", "PRDX4", "SNRPD1", "EIF1AX", "PHB2", "SNRPA1", "CDK4", "TYMS", "HDAC2", "PSMD7") |

| | | | | | | | | |
|---|---|---|---|---|---|---|---|---|
| 48 | HALLMARK_OXIDATIVE_PHOSPHORYLATION | 0.997206 | 0.997206 | 0.10402 | 0.454177 | 8208 | 78 | c("TCIRG1", "SLC25A6", "ATP5PF", "OAT", "ATP6V0B", "PHB2", "ACADVL", "ATP6V1D", "ATP5MC1", "CPT1A", "ETFB", "ACO2", "ATP6V1H", "NDUFS3", "TOMM22", "NDUFA9", "ATP6V1E1", "ATP6V0C", "ETFA", "TIMM50", "ATP5F1D", "COX11", "NDUFB6", "CYB5R3", "ATP6V0E1", "NDUFB5", "ATP5MG", "ATP6V1C1", "COX7A2", "SDHA", "BAX", "ATP5MF", "ISCU", "IDH3B", "NDUFS7", "ATP5F1B", "NDUFS6", "DECR1", "NDUFB7", "ATP5F1E", "PDHB", "UQCR10", "COX8A", "NDUFB2", "ATP5F1A", "SLC25A11", "UQCRB", "SUCLG1", "ATP6V1F", "ATP6AP1", "PDHA1", <br>"ECH1", "TIMM13", "TIMM8B") |
| 49 | HALLMARK_BILE_ACID_METABOLISM | 0.935807 | 0.997206 | 0.242889 | 0.573527 | 5976 | 7 | c("SLC29A1", "SLC35B2", "HSD3B7") |
| 50 | HALLMARK_SPERMATOGENESIS | 0.970798 | 0.997206 | 0.22772 | 0.513151 | 6116 | 6 | c("SEPTIN4", "IP6K1", "NF2") |

| Cluster 29 | pathway | pval | padj | ES | NES | nMoreExtreme | size | leadingEdge |
|---|---|---|---|---|---|---|---|---|
| 1 | HALLMARK_IL6_JAK_STAT3_SIGNALING | 0.065067 | 0.542228 | -0.396826 | -1.46306 | 612 | 21 | c("HMOX1", "STAT3", "PTPN1", "TNFRSF12A", "IL13RA1", "STAT1", "SOCS3") |
| 2 | HALLMARK_ESTROGEN_RESPONSE_EARLY | 0.025968 | 0.542228 | -0.323855 | -1.478875 | 258 | 68 | c("SLC9A3R1", "SLC2A1", "UGCG", "GREB1", "PGR", "SLC7A2", "FAM102A", "SCNN1A", "TTC39A", "THSD4", "ADCY1", "CANT1", "RET", "CLDN7", "BCL2", "AKAP1", "SH3BP5", "BLVRB", "ABHD2", "RHOD", "SEC14L2", "HSPB8", "MAPT", "SIAH2", "KDM4B", "GJA1", "ELF3", "MYC") |
| 3 | HALLMARK_ESTROGEN_RESPONSE_LATE | 0.027574 | 0.542228 | -0.324958 | -1.479014 | 274 | 66 | c("SERPINA1", "SLC9A3R1", "AGR2", "PGR", "FAM102A", "SCNN1A", "RET", "DCXR", "LSR", "SERPINA3", "BCL2", "CDH1", "BLVRB", "S100A9", "TOP2A", "DLG5", "ABHD2", "COX6C", "LLGL2", "HSPB8", "LTF", "MAPT", "TSPAN13", "SGK1", "SCUBE2", "SIAH2") |
| 4 | HALLMARK_PROTEIN_SECRETION | 0.062398 | 0.542228 | -0.354073 | -1.448836 | 610 | 34 | c("ERGIC3", "ARF1", "COPB1", "LMAN1", "ANP32E", "IGF2R", "TMED2", "MAPK1", "SNX2", "MON2", "VPS45", "TOM1L1", "ARFGEF2", "TPD52") |
| 5 | HALLMARK_KRAS_SIGNALING_UP | 0.049609 | 0.542228 | -0.345525 | -1.460328 | 488 | 40 | c("MMP9", "INHBA", "ITGB2", "PLAUR", "MAFB", "SERPINA3", "NRP1", "CTSS", "TMEM176B", "PLAU", "LAPTM5", "IKZF1", "TSPAN13", "SPARCL1", "CXCR4", "JUP", "MMP11", "MAP3K1", "IGFBP3", "LCP1", "PRRX1", "FCER1G", "AKT2", "GYPC") |
| 6 | HALLMARK_PANCREAS_BETA_CELLS | 0.036244 | 0.542228 | -0.664061 | -1.575285 | 275 | 5 | c("MAFB", "AKT3", "SRP9") |
| 7 | HALLMARK_ALLOGRAFT_REJECTION | 0.100052 | 0.714656 | -0.350488 | -1.368952 | 964 | 27 | c("MMP9", "SRGN", "INHBA", "ITGB2", "STAT1", "PSMB10", "CTSS", "SPI1", "CAPG", "TPD52", "BCL3") |
| 8 | HALLMARK_OXIDATIVE_PHOSPHORYLATION | 0.117647 | 0.735294 | 0.153311 | 1.41607 | 1 | 70 | c("GPX4", "COX7A2", "NDUFV1", "NDUFB5", "GLUD1", "NDUFA1", "CYB5R3", "SLC25A6", "NDUFAB1", "ATP6V0E1", "NDUFS6", "NDUFS7", "OXA1L", "NDUFC1", "PDHA1", "COX7C", "IDH2", "ISCU", "CYC1", "NDUFB7", "CYB5A", "PDK4", "NDUFS4", "NDUFA4", "NDUFS2", "ECHS1", "NDUFB2", "COX6A1", "NDUFC2", "ATP5MG", "MGST3", "VDAC3", "NDUFV2", "HADHA", "ATP5F1D", "UQCR10", "ATP5PD", "UQCRC1", "NDUFA3", "ACADVL", "ATP5F1E", "UQCRQ", "ECH1", "ACAA1", "ATP5MC2", "IDH3B", "NDUFB8", "COX4I1", "ECI1", "UQCRB", "NDUFA2", "NDUFB6", □ "TIMM13", "ATP5F1B", "UQCR11", "LRPPRC", "ATP6V1F", "ATP6V0C", "RHOT2", "ATP5MF", "COX7A2L", "COX8A", "ATP1B1", "ATP6AP1", "ATP6V0B", "TCIRG1", "MAOB") |
| 9 | HALLMARK_DNA_REPAIR | 0.186956 | 0.791586 | -0.307668 | -1.250919 | 1825 | 33 | c("NME4", "TAF10", "IMPDH2", "CANT1", "NUDT21", "POLD4", "POLR2I", "TMED2", "SEC61A1", "BRF2") |
| 10 | HALLMARK_NOTCH_SIGNALING | 0.219865 | 0.791586 | 0.384083 | 1.21652 | 424 | 7 | c("APH1A", "NOTCH3", "NOTCH2", "RBX1", "HES1", "PSENEN", "ARRB1") |
| 11 | HALLMARK_MTORC1_SIGNALING | 0.173015 | 0.791586 | -0.286426 | -1.247636 | 1716 | 48 | c("SLC9A3R1", "SLC2A1", "CORO1A", "ITGB2", "ACTR3", "TXNRD1", "IFI30", "EDEM1", "PSMB5", "HSPE1", "P4HA1", "SYTL2", "ACSL3", "ACACA", "LTA4H", "TFRC", "CXCR4") |
| 12 | HALLMARK_P53_PATHWAY | 0.2105 | 0.791586 | -0.277572 | -1.209072 | 2088 | 48 | c("HMOX1", "TAX1BP3", "DCXR", "IFI30", "EPS8L2", "IP6K2", "SEC61A1", "ABCC5", "HEXIM1", "SP1", "TM4SF1", "RXRA", "PLXNB2", "SLC3A2", "CTSD", "PRMT2") |
| 13 | HALLMARK_HEME_METABOLISM | 0.194999 | 0.791586 | -0.271968 | -1.214666 | 1941 | 57 | c("HBB", "HBD", "ACP5", "SLC2A1", "SLC25A37", "UROD", "MPP1", "RBM5", "UCP2", "GLRX5", "FBXO9", "BLVRB", "RNF19A", "SLC4A1", "SNCA", "TFRC", "BCAM", "ADD1", "HAGH") |
| 14 | HALLMARK_KRAS_SIGNALING_DN | 0.221644 | 0.791586 | -0.425922 | -1.230256 | 1873 | 9 | c("SKIL", "BMPR1B", "SGK1", "EFHD1", "IGFBP2") |
| 15 | HALLMARK_E2F_TARGETS | 0.255741 | 0.85247 | -0.308116 | -1.183323 | 2449 | 25 | c("SNRPB", "NUDT21", "ANP32E", "CDKN1B", "RBBP7", "TOP2A", "SMC4", "TFRC", "MYC", "STAG1", "LUC7L3", "MCM7", "PRKDC", "HMGB2", "HNRNPD", "SSRP1", "NOP56") |
| 16 | HALLMARK_ANDROGEN_RESPONSE | 0.314251 | 0.982034 | -0.272133 | -1.126389 | 3084 | 36 | c("ACTN1", "SMS", "LMAN1", "GNAI3", "BMPR1B", "ABHD2", "ACSL3", "TPD52", "SGK1", "TMEM50A", "ELL2", "SPDEF", "DBI", "PMEPA1", "B4GALT1", "SRP19", "ITGAV", "MAF", "ADRM1", "UBE2I", "STEAP4", "ELOVL5", "SCD", "PA2G4", "INPP4B", "GSR") |
| 17 | HALLMARK_TNFA_SIGNALING_VIA_NFKB | 0.963558 | 0.994071 | -0.126453 | -0.568203 | 9597 | 60 | c("TNC", "INHBA", "PLAUR", "FOSL2", "SOCS3", "GEM", "PLAU", "SOD2", "NFKBIA", "SGK1", "KLF2", "BCL3", "CEBPB", "MYC", "PHLDA1", "EIF1", "MAP3K8", "LITAF", "SDC4", "PMEPA1", "TNIP1", "B4GALT1") |
| 18 | HALLMARK_HYPOXIA | 0.62342 | 0.994071 | -0.203192 | -0.919123 | 6212 | 63 | c("HMOX1", "TGFBI", "SLC2A1", "GAA", "PLAUR", "MT2A", "CDKN1B", "CITED2", "BCL2", "FOSL2", "P4HA1", "MIF", "TPD52", "CXCR4", "DTNA", "SIAH2", "MAP3K1", "IGFBP3", "HEXA", "IDS", "VHL", "PAM", "LOX", "TGFB3", "SDC4", "HK1") |

| | | | | | | | | |
|---|---|---|---|---|---|---|---|---|
| 19 | HALLMARK_CHOLESTEROL_HOMEOSTASIS | 0.940474 | 0.994071 | -0.167613 | -0.574129 | 8578 | 16 | c("PLAUR", "TNFRSF12A", "ATXN2") |
| 20 | HALLMARK_MITOTIC_SPINDLE | 0.923193 | 0.994071 | -0.144834 | -0.639027 | 9182 | 52 | c("ACTN4", "RANBP9", "ARHGEF2", "CNTRL", "TOP2A", "ARHGAP4", "SMC4", "SEPTIN9", "NUMA1", "CYTH2", "EZR", "SPTAN1", "NET1", "RHOT2", "ARFIP2", "YWHAE", "CAPZB", "LRPPRC", "PCM1") |
| 21 | HALLMARK_WNT_BETA_CATENIN_SIGNALING | 0.956927 | 0.994071 | -0.241931 | -0.57391 | 7286 | 5 | c("CSNK1E", "MYC", "CTNNB1") |
| 22 | HALLMARK_TGF_BETA_SIGNALING | 0.921801 | 0.994071 | 0.111222 | 0.681148 | 388 | 25 | c("TGIF1", "SMAD3", "IFNGR2", "ARID4B", "HIPK2", "FNTA", "BMPR2", "PPM1A", "UBE2D3", "SKI", "ID3", "ENG", "NCOR2", "ID2", "LTBP2") |
| 23 | HALLMARK_G2M_CHECKPOINT | 0.587383 | 0.994071 | -0.230101 | -0.923729 | 5716 | 31 | c("MT2A", "CDKN1B", "RBM14", "BUB3", "TOP2A", "SMC4", "NUMA1", "HNRNPU", "BCL3", "MYC", "ATRX", "STAG1") |
| 24 | HALLMARK_APOPTOSIS | 0.966623 | 0.994071 | -0.122735 | -0.541527 | 9614 | 52 | c("HMOX1", "ERBB2", "CDKN1B", "IGF2R", "TNFRSF12A") |
| 25 | HALLMARK_ADIPOGENESIS | 0.994071 | 0.994071 | -0.096898 | -0.428505 | 9891 | 53 | c("CD151", "RAB34", "PIM3", "UCP2", "PHLDB1", "CMBL", "HSPB8", "APLP2", "SPARCL1", "GPAT4", "G3BP2", "COX8A", "RTN3", "ESYT1", "UQCR11", "CCNG2", "BCL2L13", "CMPK1", "CD36", "DDT", "SOD1", "ECH1", "UQCRQ", "UBC", "TALDO1", "UQCRC1", "UQCR10", "TOB1", "MGST3", "YWHAG", "DHRS7", "STOM", "PFKFB3", "COX6A1", "COL4A1", "ECHS1", "SLC1A5", "LAMA4", "REEP5", "ALDH2", "NDUFB7", "CYC1", "TKT", "C3", "APOE", "CAVIN1") |
| 26 | HALLMARK_MYOGENESIS | 0.74202 | 0.994071 | -0.192682 | -0.817908 | 7322 | 41 | c("GAA", "HSPB8", "PVALB", "PLXNB2", "ADAM12", "DTNA", "PDLIM7", "ATP6AP1", "SPTAN1", "NQO1", "LSP1", "IGFBP3", "FLII", "SPDEF", "TNNT1", "AKT2") |
| 27 | HALLMARK_INTERFERON_ALPHA_RESPONSE | 0.988928 | 0.994071 | -0.133496 | -0.457267 | 9020 | 16 | c("IFI30", "TRIM25", "IFI27", "LY6E", "LAP3", "IFITM2", "STAT2", "PSME1", "BST2", "CD47", "C1S", "TXNIP") |
| 28 | HALLMARK_INTERFERON_GAMMA_RESPONSE | 0.488467 | 0.994071 | -0.238022 | -0.996682 | 4806 | 38 | c("STAT3", "MT2A", "PTPN1", "IFI30", "STAT1", "SOCS3", "PSMB10", "TRIM25", "RBCK1", "SOD2", "NFKBIA") |
| 29 | HALLMARK_APICAL_JUNCTION | 0.512495 | 0.994071 | -0.225903 | -0.984007 | 5085 | 48 | c("MMP9", "ACTN4", "ACTN1", "TGFBI", "CTNNA1", "AKT3", "CLDN7", "CDH1", "PARVA", "SYMPK", "VASP", "CLDN4", "PBX2", "JUP", "TSPAN4", "STX4", "CD276", "TIAL1", "CERCAM", "GNAI2", "AKT2", "B4GALT1", "MAPK13", "ITGB1", "EXOC4", "PIK3R3", "WASL", "INPPL1", "MSN", "THY1", "COL16A1", "CD99", "PECAM1", "CDH11", "ADAM15", "SDC3", "SORBS3", "MYH9") |
| 30 | HALLMARK_APICAL_SURFACE | 0.470508 | 0.994071 | -0.375301 | -0.997907 | 3796 | 7 | c("PLAUR", "GSTM3", "SULF2") |
| 31 | HALLMARK_HEDGEHOG_SIGNALING | 0.943563 | 0.994071 | -0.210161 | -0.582818 | 7790 | 8 | c("NRP1", "ADGRG1") |
| 32 | HALLMARK_COMPLEMENT | 0.789903 | 0.994071 | -0.179634 | -0.782465 | 7838 | 48 | c("SERPINA1", "PLAUR", "GNAI3", "S100A9", "CTSS", "CTSL", "APOC1", "LIPA", "LTF", "LTA4H", "CD46", "CTSD", "CEBPB", "STX4", "HSPA1A", "FCER1G", "GNAI2") |
| 33 | HALLMARK_UNFOLDED_PROTEIN_RESPONSE | 0.643378 | 0.994071 | -0.220241 | -0.884143 | 6261 | 31 | c("DCTN1", "NHP2", "BANF1", "EDEM1", "HERPUD1", "EIF4EBP1") |
| 34 | HALLMARK_PI3K_AKT_MTOR_SIGNALING | 0.545501 | 0.994071 | -0.23941 | -0.955198 | 5298 | 30 | c("SLC2A1", "ARF1", "CDKN1B", "ACTR3", "SMAD2", "MAPK1", "PAK4", "ACACA", "CXCR4", "PPP1CA", "ARPC3", "ITPR2", "CAB39", "MYD88", "GRB2", "TNFRSF1A", "PIK3R3", "GSK3B", "GRK2", "MKNK2", "STAT2") |
| 35 | HALLMARK_MYC_TARGETS_V1 | 0.447477 | 0.994071 | -0.235164 | -1.02779 | 4442 | 49 | c("EIF3B", "NHP2", "IMPDH2", "PSMD3", "PCBP1", "PSMA7", "HSPE1", "BUB3", "ILF2", "SNRPD2", "HNRNPU", "FAM120A", "TXNL4A", "MYC", "CLNS1A", "PPIA", "YWHAE", "MCM7", "NCBP2", "EIF4A1", "APEX1", "HNRNPC", "HNRNPD", "NOP56", "TUFM", "EEF1B2", "CBX3", "CCT3", "DHX15", "PA2G4", "HSP90AB1", "GNL3", "HDGF") |
| 36 | HALLMARK_MYC_TARGETS_V2 | 0.606939 | 0.994071 | -0.336085 | -0.893634 | 4897 | 7 | c("HSPE1", "MYC", "NOP56", "CBX3", "PA2G4", "GNL3") |
| 37 | HALLMARK_EPITHELIAL_MESENCHYMAL_TRANSITIO N | 0.977189 | 0.994071 | -0.12265 | -0.580476 | 9766 | 90 | c("TGFBI", "TNC", "INHBA", "MXRA5", "COLGALT1", "PLAUR", "DPYSL3", "LOXL2", "IL32", "TNFRSF12A") |
| 38 | HALLMARK_INFLAMMATORY_RESPONSE | 0.79098 | 0.994071 | -0.193436 | -0.755532 | 7628 | 27 | c("SLC7A2", "INHBA", "PLAUR", "GNAI3", "ITGA5", "CYBB", "NFKBIA", "MYC") |
| 39 | HALLMARK_XENOBIOTIC_METABOLISM | 0.915826 | 0.994071 | -0.149309 | -0.643167 | 9073 | 45 | c("HMOX1", "PGD", "MT2A", "DCXR", "PGRMC1", "PSMB10", "BLVRB", "TMEM176B", "NQO1", "JUP", "CNDP2", "MCCC2", "PAPSS2", "CD36", "DDT", "TNFRSF1A", "BCAR1", "DDAH2", "ELOVL5", "ECH1", "ESR1", "SERPINE1", "GSR", "NINJ1", "COMT", "CSAD") |
| 40 | HALLMARK_FATTY_ACID_METABOLISM | 0.512944 | 0.994071 | -0.254743 | -0.978343 | 4913 | 25 | c("SMS", "UROD", "FH", "BMPR1B", "MIF", "HSD17B4", "APEX1", "HSP90AA1", "CD36", "PCBD1", "ECI1", "IDH3B", "ELOVL5", "ACAA1", "ECH1", "ACADVL", "PSME1", "S100A10") |

| | | | | | | | | |
|---|---|---|---|---|---|---|---|---|
| 41 | HALLMARK_GLYCOLYSIS | 0.666566 | 0.994071 | -0.204364 | -0.877985 | 6602 | 44 | c("TGFBI", "CITED2", "IL13RA1", "FUT8", "P4HA1", "QSOX1", "RBCK1", "MIF", "CXCR4", "ELF3", "PYGL", "IGFBP3", "PPIA", "SDC1", "PAM", "PGLS", "B4GALT1", "EGLN3", "CHPF", "PLOD1", "SOD1", "TFF3", "GFPT1", "AKR1A1", "COPB2", "TALDO1", "DDIT4", "PKM", "PYGB") |
| 42 | HALLMARK_REACTIVE_OXYGEN_SPECIES_PATHWAY | 0.764745 | 0.994071 | -0.224381 | -0.768578 | 6975 | 16 | c("MPO", "TXNRD1", "SOD2", "NQO1", "LSP1") |
| 43 | HALLMARK_UV_RESPONSE_UP | 0.35408 | 0.994071 | -0.264564 | -1.09506 | 3475 | 36 | c("HMOX1", "MGAT1", "UROD", "RET", "NXF1", "WIZ", "SOD2", "TFRC", "NFKBIA", "ARRB2", "HNRNPU", "EIF5", "CREG1", "PRPF3", "IGFBP2", "ATP6V1F") |
| 44 | HALLMARK_UV_RESPONSE_DN | 0.938195 | 0.994071 | -0.140985 | -0.60092 | 9274 | 42 | c("ERBB2", "CDKN1B", "AKT3", "CITED2", "NRP1") |
| 45 | HALLMARK_ANGIOGENESIS | 0.467234 | 0.994071 | -0.327539 | -1.004216 | 4063 | 11 | c("LRPAP1", "NRP1") |
| 46 | HALLMARK_COAGULATION | 0.926943 | 0.994071 | -0.147251 | -0.613114 | 9109 | 37 | c("SERPINA1", "MMP9", "S100A1", "PLAU", "APOC1", "LTA4H", "CTSK", "MMP11", "CRIP2") |
| 47 | HALLMARK_IL2_STAT5_SIGNALING | 0.966182 | 0.994071 | -0.124913 | -0.536652 | 9570 | 44 | c("SERPINB6", "IGF2R", "NRP1", "BCL2", "P4HA1", "S100A1", "ITGA6", "CAPG", "MYC", "ECM1", "PHLDA1", "LRIG1", "MAP3K8", "ITGAV", "MYO1C", "ENPP1", "MAPKAPK2", "PIM1", "PLEC", "RNH1", "EMP1", "RABGAP1L", "AHR", "NFKBIZ", "CCND3") |
| 48 | HALLMARK_BILE_ACID_METABOLISM | 0.398998 | 0.994071 | 0.242815 | 1.01791 | 477 | 12 | c("ALDH9A1", "SCP2", "CAT", "DHCR24", "IDH2", "LONP2", "GSTK1", "SOD1", "SLC29A1", "OPTN", "HSD17B4", "RXRA") |
| 49 | HALLMARK_PEROXISOME | 0.777351 | 0.994071 | 0.140438 | 0.793576 | 404 | 22 | c("ALDH9A1", "SCP2", "CAT", "FIS1", "DHCR24", "SEMA3C", "IDH2", "ERCC1", "LONP2", "CNBP", "SMARCC1", "GSTK1", "ECH1", "ACAA1", "ELOVL5", "SOD1", "TSPO", "HSD17B4", "CRABP2", "SOD2", "TOP2A", "ABCC5") |
| 50 | HALLMARK_SPERMATOGENESIS | 0.954301 | 0.994071 | -0.244424 | -0.579823 | 7266 | 5 | GSTM3 |
